# Labels of aberrant Clusters of Differentiation gene expression in a compendium of systemic lupus erythematosus patients

**DOI:** 10.1101/277145

**Authors:** Trang T. Le, Nigel O. Blackwood, Matthew K. Breitenstein

**Author notes:** correspondence to:* mkbreit at pennmedicine dot upenn dot edu.

## Abstract

This author manuscript serves as an extended annotation of gene expression for all known clusters of differentiation (**CD**) within a compendium of systemic lupus erythematosus (**SLE**) patients. The overarching goal for this line of research is to enrich the perspective of the CD transcriptome with upstream gene expression features.

## Introduction

CDs are cell surface biomarkers that denote key biological differences between cell types and disease state. For each of the >400 known CDs^1^, distinct monoclonal antibodies (**mABs**) enable robust immunophenotyping and serve as scalable biomarkers for translational research. Annotation of CD molecules have been organized through a series of international meetings known as the Human Leucocyte Differentiation Antigens (**HLDA**) Workshops, affiliated with the Human Cell Differentiation Molecules (**HCDM**) organization.

### CD nomenclature

*<http://www.hcdm.org/>*

## Methods

A compendium containing human SLE gene expression data was previously collected, aggregated, and normalized by collaborators in the Greene Lab at the University of Pennsylvania. *<https://github.com/greenelab/rheum-plier-data/tree/master/sle-wb>* This compendium was slightly modified to include basic demographic information and exclude patients not belonging to classifications of healthy control, treatment naïve SLE, or SLE with exposure to various treatments – the modified dataset represents our ‘**SLE Compendium**’. Entrez gene ID to CD mapping was provided by HCDM^1^. The SLE Compendium dataset and R code corresponding to data pre-processing can be found on the Breitenstein Lab Github page.

### SLE Compendium

*<https://breitensteinlab.github.io/SLE-Compendium-2018/>*

Within our SLE Compendium, all known CDs gene expression of, were categorized as ‘aberrant’ or ‘non-aberrant’ based on the following criteria: *i)* two-tailed normalization at 20^th^ and 80^th^ percentile of relative gene expression. Specifically, the two tails encompassed ‘aberrant’ CD expression, whereas the middle distribution served as ‘non-aberrant’. Following visual inspection of all histograms, specific CDs were deemed to require manual adjustments. *ii)* Manually adjustment of two-tail normalization was applied when CDs followed apparent normal distribution but would require slight modification of thresholds to characterization feature variation. *iii)* Binarization CD features was applied when an obvious non-normal distribution was observed. Thresholds separated the expression values into low/ high groups (instead of non-aberrant/aberrant) to capture apparent patterns in the expression distributions.

In future research, CDs will be enriched with perspective of interdependent gene expression features using the integrated machine learning pipeline for aberrant biomarker enrichment (**i-mAB pipeline**).

### i-mAB pipeline

*<https://breitensteinlab.github.io/i-mAB/>*

## Results

Within the original study cohorts, multiple observations were generated for most patients. Observation characteristics of the SLE Compendium, including PMID for the 6 original studies^2,3,4,5,6,7^, can be found in **Table 1**. Additional sample/cohort can be ascertained from the Gene Expression Omnibus via GSE or GEO accession ID.

**Table 1.** SLE Compendium characteristics as ascertained from study of origin

|  | <u>Cohort 1</u><sup>2</sup> | <u>Cohort 2</u><sup>3</sup> | <u>Cohort 3</u><sup>4</sup> | <u>Cohort 4</u><sup>5</sup> | <u>Cohort 5</u><sup>6</sup> | <u>Cohort 6</u><sup>7</sup> | <u>Overall</u> |
| --- | --- | --- | --- | --- | --- | --- | --- |
| **Study PMID** | 18631455 | 23203821 | 24644022 | 25736140 | 27040498 | 26138472 | --- |
| **Study GEO identifier** | GSE11907 | GSE39088 | GSE49454 | GSE61635 | GSE65391 | GSE78193 | --- |
| **Healthy control\*** | **0** | **46** | **0** | **30** | **72** | **12** | **160** |
| median age (range) | --- | 34.5<br>(19-50) | --- | --- | 12<br>(6-21) | --- | 16<br>(6-50) |
| gender - female/male | --- | 34 | --- | --- | 57 | --- | 91 |
| **SLE-treatment naïve\*** | **37** | **21** | **177** | **99** | **924** | **32** | **1290** |
| median age (range) | 14<br>(8-17) | 43<br>(20-50) | 40<br>(18-71) | --- | 15<br>(6-19) | --- | 16<br>(6-71) |
| gender - female | 35 | 21 | 148 | --- | 817 | --- | 1021 |
| **SLE-various treatments\*** | **0** | **57** | **0** | **0** | **0** | **69** | **126** |
| median age (range) | --- | 36<br>(19-50) | --- | --- | --- | --- | 36<br>(19-50) |
| gender - female/male | --- | 57 | --- | --- | --- | --- | 57 |

\*observation characteristics include multiple observations per patient

### Gene Expression Omnibus

*<https://www.ncbi.nlm.nih.gov/geo/query/acc.cgi>*

By default, CD gene expression thresholds were labeled as ‘aberrant’ or ‘non-aberrant’, with ‘aberrant’ being further delineated as ‘low’ or ‘high’. Gene expression was stratified at 20 and 80 percentiles, with low expression being between 0 and 20, average between 20 and 80, and high between 80 and 100. Descriptive statistics of CD gene expression with corresponding Entrez ID can be found in **Supplement 1, Table 2**.

Overall the default two-tailed thresholds provided satisfactory characterization of ‘aberrant’ (including ‘low’ and ‘high’) vs. ‘non-aberrant’ normal gene expression distributions. However, some CDs required manual adjustment, including shifting of thresholds (n=3) and binary transformation for non-normal distributions (n=85) (**Supplement 2, Table 3**). Amongst CD genes requiring binary transformation, no clear data-driven hypothesis of ‘aberrant’ vs ‘average’ was practical so gene expression was labeled simply as ‘low’ or ‘high’ (i.e. no clear baseline comparison is available). Consensus review by the research team determined manual revisions of thresholds. Detailed labelling of all CD features (n=351) identified within the SLE Compendium can be found in **Supplement 3, Figures 1-290**. Included are histograms of CD gene expression distribution and descriptive statistics of expression and variation.

---

**Supplement 1. Table 2.** CD expression characteristics with the SLE Compendium

| Cluster of Differentiation | Entrez ID | 1<sup>st</sup> quartile | Median | Mean | Standard error | 3<sup>rd</sup> quartile |
| --- | --- | --- | --- | --- | --- | --- |
| CD1A | 909 | 0.2877 | 0.4131 | 0.4306 | 0.0049 | 0.5418 |
| CD1B | 910 | 0.0429 | 0.2810 | 0.2859 | 0.0067 | 0.4699 |
| CD1C | 911 | 0.4298 | 0.4847 | 0.5165 | 0.0046 | 0.6191 |
| CD1D | 912 | 0.4725 | 0.5790 | 0.5669 | 0.0047 | 0.6692 |
| CD1E | 913 | 0.0808 | 0.3933 | 0.3550 | 0.0072 | 0.5690 |
| CD2 | 914 | 0.5507 | 0.6660 | 0.6479 | 0.0045 | 0.7567 |
| CD3D | 915 | 0.5816 | 0.6579 | 0.6517 | 0.0037 | 0.7393 |
| CD3E | 916 | 0.3478 | 0.4621 | 0.4756 | 0.0050 | 0.5963 |
| CD3G | 917 | 0.4454 | 0.5510 | 0.5513 | 0.0045 | 0.6593 |
| CD4 | 920 | 0.4148 | 0.5369 | 0.5325 | 0.0055 | 0.6743 |
| CD5 | 921 | 0.4914 | 0.6062 | 0.5890 | 0.0047 | 0.7053 |
| CD6 | 923 | 0.5397 | 0.6628 | 0.6438 | 0.0048 | 0.7728 |
| CD7 | 924 | 0.4072 | 0.5400 | 0.5347 | 0.0052 | 0.6661 |
| CD8A | 925 | 0.4365 | 0.5553 | 0.5594 | 0.0049 | 0.6907 |
| CD8B | 926 | 0.3612 | 0.4399 | 0.4622 | 0.0044 | 0.5634 |
| CD9 | 928 | 0.2634 | 0.4078 | 0.4108 | 0.0056 | 0.5510 |
| CD10 | 4311 | 0.3475 | 0.4486 | 0.4754 | 0.0053 | 0.5955 |
| CD11A | 3683 | 0.3848 | 0.4876 | 0.4934 | 0.0047 | 0.5992 |
| CD11B | 3684 | 0.4461 | 0.5705 | 0.5660 | 0.0049 | 0.6893 |
| CD11C | 3687 | 0.3237 | 0.4352 | 0.4567 | 0.0051 | 0.5685 |
| CD13 | 290 | 0.4166 | 0.5281 | 0.5234 | 0.0048 | 0.6361 |
| CD14 | 929 | 0.4937 | 0.6117 | 0.5974 | 0.0047 | 0.7128 |
| CD15 | 2526 | 0.3056 | 0.3946 | 0.4032 | 0.0041 | 0.4802 |
| CD16B | 2215 | 0.6375 | 0.7064 | 0.6900 | 0.0039 | 0.7744 |
| CD18 | 3689 | 0.2797 | 0.3487 | 0.3843 | 0.0047 | 0.4451 |
| CD19 | 930 | 0.3390 | 0.4880 | 0.4796 | 0.0057 | 0.6270 |
| CD20 | 931 | 0.4250 | 0.5417 | 0.5368 | 0.0045 | 0.6543 |
| CD21 | 1380 | 0.4267 | 0.5343 | 0.5282 | 0.0051 | 0.6581 |
| CD22 | 933 | 0.2415 | 0.3850 | 0.3967 | 0.0051 | 0.5316 |
| CD23 | 2208 | 0.3318 | 0.3686 | 0.4186 | 0.0052 | 0.5378 |
| CD25 | 3559 | 0.3142 | 0.3952 | 0.4048 | 0.0042 | 0.4792 |
| CD26 | 1803 | 0.3800 | 0.4833 | 0.4955 | 0.0048 | 0.6179 |
| CD27 | 939 | 0.4434 | 0.5544 | 0.5574 | 0.0049 | 0.6732 |
| CD28 | 940 | 0.5163 | 0.6076 | 0.5988 | 0.0042 | 0.6912 |
| CD30 | 943 | 0.4002 | 0.5314 | 0.5164 | 0.0049 | 0.6416 |
| CD31 | 5175 | 0.5092 | 0.6208 | 0.6047 | 0.0047 | 0.7196 |
| CD33 | 945 | 0.4531 | 0.5563 | 0.5536 | 0.0045 | 0.6564 |
| CD34 | 947 | 0.0888 | 0.2385 | 0.3193 | 0.0072 | 0.5643 |
| CD35 | 1378 | 0.2969 | 0.3960 | 0.4141 | 0.0048 | 0.5101 |
| CD36 | 948 | 0.5503 | 0.6644 | 0.6425 | 0.0046 | 0.7539 |
| CD37 | 951 | 0.4015 | 0.5194 | 0.5212 | 0.0048 | 0.6362 |
| CD38 | 952 | 0.2643 | 0.3496 | 0.3615 | 0.0042 | 0.4382 |
| CD39 | 953 | 0.4219 | 0.5449 | 0.5388 | 0.0048 | 0.6571 |
| CD40 | 958 | 0.4295 | 0.5178 | 0.5264 | 0.0049 | 0.6269 |
| CD41 | 3674 | 0.2975 | 0.4246 | 0.4302 | 0.0051 | 0.5517 |
| CD42A | 2815 | 0.4343 | 0.5542 | 0.5424 | 0.0049 | 0.6606 |
| CD42B | 2811 | 0.2818 | 0.4168 | 0.4206 | 0.0055 | 0.5492 |
| CD43 | 6693 | 0.4208 | 0.5204 | 0.5248 | 0.0046 | 0.6187 |
| CD44 | 960 | 0.4447 | 0.5468 | 0.5461 | 0.0044 | 0.6526 |
| CD45 | 5788 | 0.4831 | 0.5932 | 0.5897 | 0.0049 | 0.7097 |
| CD46 | 4179 | 0.3236 | 0.4766 | 0.4776 | 0.0056 | 0.6164 |
| CD47 | 961 | 0.3421 | 0.4422 | 0.4758 | 0.0051 | 0.6049 |
| CD48 | 962 | 0.5302 | 0.6346 | 0.6206 | 0.0045 | 0.7340 |
| CD49B | 3673 | 0.0505 | 0.0692 | 0.2157 | 0.0057 | 0.3656 |
| CD49C | 3675 | 0.0874 | 0.1118 | 0.2535 | 0.0065 | 0.4513 |
| CD49D | 3676 | 0.4888 | 0.5827 | 0.5825 | 0.0044 | 0.6847 |
| CD49E | 3678 | 0.4647 | 0.5786 | 0.5710 | 0.0044 | 0.6751 |
| CD49F | 3655 | 0.4701 | 0.5373 | 0.5076 | 0.0043 | 0.5757 |
| CD50 | 3385 | 0.4232 | 0.5380 | 0.5340 | 0.0050 | 0.6507 |
| CD51 | 3685 | 0.4443 | 0.5668 | 0.5565 | 0.0046 | 0.6670 |
| CD52 | 1043 | 0.5706 | 0.6869 | 0.6614 | 0.0046 | 0.7794 |
| CD53 | 963 | 0.4696 | 0.5772 | 0.5712 | 0.0045 | 0.6843 |
| CD54 | 3383 | 0.4393 | 0.4668 | 0.4858 | 0.0041 | 0.5453 |
| CD55 | 1604 | 0.3661 | 0.4807 | 0.4874 | 0.0048 | 0.5927 |
| CD56 | 4684 | 0.4322 | 0.5026 | 0.4613 | 0.0043 | 0.5338 |
| CD57 | 27087 | 0.0730 | 0.1057 | 0.2654 | 0.0066 | 0.4719 |
| CD58 | 965 | 0.4668 | 0.5758 | 0.5660 | 0.0045 | 0.6744 |
| CD59 | 966 | 0.3650 | 0.4031 | 0.4285 | 0.0038 | 0.4862 |
| CD61 | 3690 | 0.3290 | 0.4558 | 0.4583 | 0.0051 | 0.5751 |
| CD62E | 6401 | 0.0770 | 0.0991 | 0.2399 | 0.0067 | 0.4248 |
| CD62L | 6402 | 0.7253 | 0.8035 | 0.7461 | 0.0049 | 0.8507 |
| CD62P | 6403 | 0.3499 | 0.3783 | 0.4157 | 0.0043 | 0.4860 |
| CD63 | 967 | 0.3594 | 0.4536 | 0.4675 | 0.0049 | 0.5648 |
| CD66A | 634 | 0.2915 | 0.3878 | 0.4013 | 0.0045 | 0.5012 |
| CD66B | 1088 | 0.1388 | 0.2813 | 0.3267 | 0.0057 | 0.4510 |
| CD66C | 4680 | 0.1392 | 0.2528 | 0.2982 | 0.0058 | 0.4030 |
| CD66D | 1084 | 0.4487 | 0.5544 | 0.5490 | 0.0047 | 0.6591 |
| CD66E | 1048 | 0.0563 | 0.0711 | 0.1684 | 0.0054 | 0.2331 |
| CD66F | 5669 | 0.0613 | 0.0806 | 0.2017 | 0.0063 | 0.3510 |
| CD68 | 968 | 0.3514 | 0.4872 | 0.4973 | 0.0055 | 0.6275 |
| CD69 | 969 | 0.3740 | 0.4842 | 0.4911 | 0.0048 | 0.6100 |
| CD70 | 970 | 0.2230 | 0.2760 | 0.3463 | 0.0050 | 0.4596 |
| CD71 | 7037 | 0.3510 | 0.4346 | 0.4396 | 0.0038 | 0.5131 |
| CD72 | 971 | 0.3064 | 0.4160 | 0.4328 | 0.0049 | 0.5519 |
| CD73 | 4907 | 0.4912 | 0.5631 | 0.5156 | 0.0051 | 0.5975 |
| CD74 | 972 | 0.4180 | 0.5258 | 0.5255 | 0.0048 | 0.6352 |
| CD79A | 973 | 0.3382 | 0.4927 | 0.4903 | 0.0055 | 0.6265 |
| CD79B | 974 | 0.3125 | 0.4823 | 0.4761 | 0.0058 | 0.6293 |
| CD80 | 941 | 0.0514 | 0.1602 | 0.2415 | 0.0063 | 0.4027 |
| CD81 | 975 | 0.5014 | 0.6132 | 0.6063 | 0.0045 | 0.7234 |
| CD82 | 3732 | 0.3397 | 0.4747 | 0.4727 | 0.0051 | 0.5940 |
| CD83 | 9308 | 0.3623 | 0.4244 | 0.4407 | 0.0040 | 0.5118 |
| CD84 | 8832 | 0.3429 | 0.4340 | 0.4523 | 0.0045 | 0.5422 |
| CD85A | 11025 | 0.6931 | 0.7578 | 0.7279 | 0.0038 | 0.8070 |
| CD85C | 10990 | 0.0850 | 0.4020 | 0.3630 | 0.0067 | 0.5531 |
| CD85D | 10288 | 0.3225 | 0.4431 | 0.4598 | 0.0053 | 0.5779 |
| CD85F | 353514 | 0.3677 | 0.5059 | 0.5027 | 0.0053 | 0.6397 |
| CD85G | 23547 | 0.4090 | 0.4471 | 0.4459 | 0.0041 | 0.4722 |
| CD85H | 11027 | 0.5010 | 0.6070 | 0.5915 | 0.0044 | 0.7014 |
| CD85J | 10859 | 0.3159 | 0.4139 | 0.4257 | 0.0046 | 0.5250 |
| CD85K | 11006 | 0.4451 | 0.5721 | 0.5530 | 0.0049 | 0.6757 |
| CD85M | 79166 | 0.0732 | 0.0925 | 0.1941 | 0.0058 | 0.2385 |
| CD86 | 942 | 0.4575 | 0.5648 | 0.5551 | 0.0046 | 0.6661 |
| CD87 | 5329 | 0.3865 | 0.5033 | 0.5014 | 0.0048 | 0.6157 |
| CD88 | 728 | 0.4475 | 0.5733 | 0.5605 | 0.0048 | 0.6804 |
| CD89 | 2204 | 0.2910 | 0.3992 | 0.4156 | 0.0049 | 0.5191 |
| CD90 | 7070 | 0.4564 | 0.5648 | 0.5543 | 0.0046 | 0.6762 |
| CD91 | 4035 | 0.3538 | 0.3758 | 0.4351 | 0.0045 | 0.5273 |
| CD92 | 23446 | 0.2598 | 0.3575 | 0.3815 | 0.0048 | 0.4715 |
| CD93 | 22918 | 0.5621 | 0.6614 | 0.6459 | 0.0043 | 0.7482 |
| CD94 | 3824 | 0.3531 | 0.4708 | 0.4715 | 0.0051 | 0.5947 |
| CD95 | 355 | 0.2904 | 0.3998 | 0.4154 | 0.0050 | 0.5267 |
| CD96 | 10225 | 0.4582 | 0.5713 | 0.5583 | 0.0046 | 0.6695 |
| CD97 | 976 | 0.6236 | 0.6971 | 0.6633 | 0.0046 | 0.7619 |
| CD98 | 6520 | 0.3606 | 0.4524 | 0.4941 | 0.0051 | 0.5967 |
| CD99 | 4267 | 0.4504 | 0.5680 | 0.5712 | 0.0050 | 0.6912 |
| CD100 | 10507 | 0.3385 | 0.4490 | 0.4607 | 0.0050 | 0.5779 |
| CD101 | 9398 | 0.2724 | 0.3138 | 0.3420 | 0.0047 | 0.3551 |
| CD102 | 3384 | 0.4566 | 0.5730 | 0.5620 | 0.0049 | 0.6810 |
| CD103 | 3682 | 0.4387 | 0.5648 | 0.5500 | 0.0050 | 0.6722 |
| CD104 | 3691 | 0.0816 | 0.1027 | 0.2067 | 0.0057 | 0.2799 |
| CD105 | 2022 | 0.3528 | 0.3820 | 0.4359 | 0.0046 | 0.5444 |
| CD106 | 7412 | 0.0866 | 0.1180 | 0.2716 | 0.0065 | 0.4658 |
| CD107A | 3916 | 0.4480 | 0.5595 | 0.5639 | 0.0050 | 0.6827 |
| CD107B | 3920 | 0.4848 | 0.6087 | 0.5944 | 0.0050 | 0.7172 |
| CD108 | 8482 | 0.0407 | 0.0579 | 0.2029 | 0.0058 | 0.3577 |
| CD109 | 135228 | 0.0000 | 0.0000 | 0.1030 | 0.0052 | 0.1858 |
| CD110 | 4352 | 0.3177 | 0.4492 | 0.4495 | 0.0054 | 0.5842 |
| CD111 | 5818 | 0.0928 | 0.1236 | 0.2683 | 0.0066 | 0.4653 |
| CD112 | 5819 | 0.3224 | 0.4273 | 0.4002 | 0.0044 | 0.4934 |
| CD113 | 25945 | 0.0681 | 0.0913 | 0.2426 | 0.0064 | 0.4338 |
| CD114 | 1441 | 0.5753 | 0.6686 | 0.6467 | 0.0043 | 0.7422 |
| CD115 | 1436 | 0.5198 | 0.6335 | 0.6169 | 0.0047 | 0.7290 |
| CD116 | 1438 | 0.4898 | 0.6032 | 0.5869 | 0.0048 | 0.7041 |
| CD117 | 3815 | 0.3513 | 0.4393 | 0.4171 | 0.0048 | 0.5119 |
| CD118 | 3977 | 0.0659 | 0.0853 | 0.1629 | 0.0050 | 0.1796 |
| CD119 | 3459 | 0.4095 | 0.5120 | 0.5233 | 0.0048 | 0.6231 |
| CD120A | 7132 | 0.5005 | 0.6198 | 0.6110 | 0.0047 | 0.7361 |
| CD120B | 7133 | 0.4559 | 0.5577 | 0.5536 | 0.0043 | 0.6504 |
| CD121A | 3554 | 0.0190 | 0.0244 | 0.1232 | 0.0049 | 0.1807 |
| CD121B | 7850 | 0.2424 | 0.3407 | 0.3743 | 0.0052 | 0.4773 |
| CD122 | 3560 | 0.3886 | 0.5172 | 0.5126 | 0.0052 | 0.6399 |
| CD123 | 3563 | 0.3700 | 0.4103 | 0.3952 | 0.0043 | 0.4318 |
| CD124 | 3566 | 0.2042 | 0.2736 | 0.3085 | 0.0043 | 0.3769 |
| CD125 | 3568 | 0.3568 | 0.4262 | 0.4325 | 0.0041 | 0.5189 |
| CD126 | 3570 | 0.3755 | 0.4928 | 0.4965 | 0.0051 | 0.6100 |
| CD127 | 3575 | 0.5461 | 0.6411 | 0.6247 | 0.0040 | 0.7205 |
| CD130 | 3572 | 0.3375 | 0.4164 | 0.4160 | 0.0041 | 0.5015 |
| CD131 | 1439 | 0.4445 | 0.5347 | 0.5372 | 0.0041 | 0.6284 |
| CD132 | 3561 | 0.3418 | 0.4780 | 0.4704 | 0.0053 | 0.5944 |
| CD133 | 8842 | 0.0488 | 0.0621 | 0.1466 | 0.0050 | 0.1944 |
| CD134 | 7293 | 0.3926 | 0.4894 | 0.4966 | 0.0045 | 0.5970 |
| CD135 | 2322 | 0.0291 | 0.0373 | 0.1480 | 0.0057 | 0.2172 |
| CD136 | 4486 | 0.0000 | 0.0000 | 0.1381 | 0.0070 | 0.1647 |
| CD137 | 3604 | 0.3478 | 0.4435 | 0.4429 | 0.0045 | 0.5422 |
| CD138 | 6382 | 0.3120 | 0.3694 | 0.3835 | 0.0044 | 0.4721 |
| CD140A | 5156 | 0.0972 | 0.1263 | 0.2032 | 0.0053 | 0.2115 |
| CD140B | 5159 | 0.3238 | 0.3561 | 0.4020 | 0.0049 | 0.4690 |
| CD141 | 7056 | 0.2608 | 0.2839 | 0.3207 | 0.0043 | 0.3411 |
| CD142 | 2152 | 0.0865 | 0.1460 | 0.2883 | 0.0067 | 0.4879 |
| CD143 | 1636 | 0.0946 | 0.1203 | 0.2143 | 0.0056 | 0.2871 |
| CD144 | 1003 | 0.1160 | 0.1465 | 0.2313 | 0.0054 | 0.2808 |
| CD146 | 4162 | 0.0703 | 0.0887 | 0.1900 | 0.0058 | 0.2591 |
| CD147 | 682 | 0.3429 | 0.4740 | 0.4832 | 0.0055 | 0.6138 |
| CD148 | 5795 | 0.4195 | 0.5279 | 0.5285 | 0.0045 | 0.6324 |
| CD150 | 6504 | 0.4758 | 0.6168 | 0.5923 | 0.0048 | 0.7180 |
| CD151 | 977 | 0.3240 | 0.4176 | 0.4358 | 0.0045 | 0.5333 |
| CD152 | 1493 | 0.5168 | 0.5735 | 0.5624 | 0.0046 | 0.6629 |
| CD153 | 944 | 0.2956 | 0.3934 | 0.3992 | 0.0045 | 0.4975 |
| CD154 | 959 | 0.3712 | 0.4881 | 0.4943 | 0.0053 | 0.6246 |
| CD155 | 5817 | 0.4646 | 0.5331 | 0.5162 | 0.0043 | 0.5983 |
| CD156A | 101 | 0.4237 | 0.5389 | 0.5380 | 0.0050 | 0.6521 |
| CD156C | 102 | 0.4647 | 0.5321 | 0.5452 | 0.0039 | 0.6233 |
| CD157 | 683 | 0.3466 | 0.4645 | 0.4743 | 0.0051 | 0.5949 |
| CD158E | 3811 | 0.3143 | 0.4080 | 0.4189 | 0.0043 | 0.5066 |
| CD158I | 3809 | 0.0000 | 0.0000 | 0.1134 | 0.0060 | 0.1442 |
| CD158K | 3812 | 0.3227 | 0.4613 | 0.4677 | 0.0053 | 0.6060 |
| CD160 | 11126 | 0.3611 | 0.4916 | 0.4939 | 0.0048 | 0.6118 |
| CD161 | 3820 | 0.4143 | 0.5328 | 0.5259 | 0.0049 | 0.6424 |
| CD162 | 6404 | 0.3945 | 0.5122 | 0.5080 | 0.0047 | 0.6193 |
| CD163 | 9332 | 0.2490 | 0.3195 | 0.3529 | 0.0043 | 0.4156 |
| CD164 | 8763 | 0.3328 | 0.4739 | 0.4945 | 0.0050 | 0.6267 |
| CD166 | 214 | 0.3513 | 0.4532 | 0.4597 | 0.0048 | 0.5668 |
| CD167A | 780 | 0.3657 | 0.4349 | 0.4419 | 0.0043 | 0.5196 |
| CD167B | 4921 | 0.0842 | 0.1105 | 0.2371 | 0.0063 | 0.3681 |
| CD168 | 3161 | 0.0473 | 0.1676 | 0.2173 | 0.0056 | 0.3415 |
| CD169 | 6614 | 0.0290 | 0.1402 | 0.2449 | 0.0072 | 0.4275 |
| CD170 | 8778 | 0.4018 | 0.5207 | 0.5186 | 0.0049 | 0.6382 |
| CD171 | 3897 | 0.0649 | 0.0819 | 0.2025 | 0.0061 | 0.3430 |
| CD172A | 140885 | 0.4620 | 0.5746 | 0.5634 | 0.0045 | 0.6771 |
| CD172G | 55423 | 0.3642 | 0.4814 | 0.4895 | 0.0049 | 0.6088 |
| CD174 | 2525 | 0.0000 | 0.0000 | 0.1298 | 0.0063 | 0.2475 |
| CD177 | 57126 | 0.0226 | 0.0311 | 0.1232 | 0.0047 | 0.1642 |
| CD178 | 356 | 0.4686 | 0.5054 | 0.5078 | 0.0044 | 0.5625 |
| CD179A | 7441 | 0.0409 | 0.0523 | 0.1288 | 0.0045 | 0.1056 |
| CD179B | 3543 | 0.5017 | 0.5881 | 0.5847 | 0.0044 | 0.6935 |
| CD180 | 4064 | 0.0896 | 0.3274 | 0.3306 | 0.0066 | 0.5309 |
| CD181 | 3577 | 0.5136 | 0.6410 | 0.6177 | 0.0049 | 0.7477 |
| CD182 | 3579 | 0.5253 | 0.6211 | 0.6064 | 0.0041 | 0.7002 |
| CD183 | 2833 | 0.4208 | 0.5164 | 0.5323 | 0.0046 | 0.6433 |
| CD184 | 7852 | 0.4024 | 0.4948 | 0.5076 | 0.0045 | 0.6019 |
| CD185 | 643 | 0.3929 | 0.5298 | 0.5033 | 0.0047 | 0.6156 |
| CD186 | 10663 | 0.4005 | 0.4329 | 0.4296 | 0.0042 | 0.4769 |
| CD191 | 1230 | 0.4444 | 0.5803 | 0.5620 | 0.0051 | 0.6919 |
| CD193 | 1232 | 0.4625 | 0.5722 | 0.5543 | 0.0049 | 0.6712 |
| CD194 | 1233 | 0.0459 | 0.0605 | 0.1963 | 0.0064 | 0.3592 |
| CD195 | 1234 | 0.3732 | 0.4295 | 0.4471 | 0.0042 | 0.5207 |
| CD196 | 1235 | 0.3381 | 0.4530 | 0.4578 | 0.0050 | 0.5753 |
| CD197 | 1236 | 0.4787 | 0.6108 | 0.5969 | 0.0050 | 0.7204 |
| CD199 | 10803 | 0.3392 | 0.4020 | 0.3840 | 0.0038 | 0.4403 |
| CD200 | 4345 | 0.0567 | 0.0819 | 0.2404 | 0.0062 | 0.4174 |
| CD201 | 10544 | 0.2714 | 0.3164 | 0.3688 | 0.0048 | 0.4618 |
| CD202B | 7010 | 0.0674 | 0.2322 | 0.2739 | 0.0064 | 0.4369 |
| CD203C | 5169 | 0.0818 | 0.1037 | 0.1722 | 0.0046 | 0.1822 |
| CD204 | 4481 | 0.0663 | 0.0921 | 0.2563 | 0.0064 | 0.4345 |
| CD205 | 4065 | 0.4357 | 0.5313 | 0.5450 | 0.0040 | 0.6345 |
| CD206 | 4360 | 0.0625 | 0.0808 | 0.1978 | 0.0054 | 0.3369 |
| CD207 | 50489 | 0.0724 | 0.0917 | 0.2102 | 0.0060 | 0.3352 |
| CD208 | 27074 | 0.2642 | 0.3410 | 0.3676 | 0.0043 | 0.4326 |
| CD209 | 30835 | 0.0000 | 0.0000 | 0.1070 | 0.0056 | 0.1425 |
| CD210 | 3587 | 0.4240 | 0.5205 | 0.5219 | 0.0044 | 0.6222 |
| CDW210B | 3588 | 0.5186 | 0.6042 | 0.5920 | 0.0041 | 0.6832 |
| CD212 | 3594 | 0.4045 | 0.5213 | 0.5310 | 0.0049 | 0.6489 |
| CD213A1 | 3597 | 0.4451 | 0.5561 | 0.5489 | 0.0048 | 0.6578 |
| CD213A2 | 3598 | 0.0658 | 0.0828 | 0.1504 | 0.0045 | 0.1479 |
| CD215 | 3601 | 0.0744 | 0.0945 | 0.1943 | 0.0057 | 0.2438 |
| CD217A | 23765 | 0.3543 | 0.4847 | 0.4848 | 0.0053 | 0.6106 |
| CD218A | 8809 | 0.2310 | 0.3237 | 0.3634 | 0.0053 | 0.4489 |
| CD218B | 8807 | 0.3551 | 0.4772 | 0.4696 | 0.0049 | 0.5795 |
| CD220 | 3643 | 0.0558 | 0.0707 | 0.1690 | 0.0054 | 0.2508 |
| CD221 | 3480 | 0.3040 | 0.3717 | 0.3908 | 0.0041 | 0.4642 |
| CD222 | 3482 | 0.3865 | 0.4883 | 0.4928 | 0.0047 | 0.5964 |
| CD223 | 3902 | 0.3208 | 0.4433 | 0.4631 | 0.0048 | 0.5783 |
| CD225 | 8519 | 0.6660 | 0.7433 | 0.7008 | 0.0047 | 0.7958 |
| CD226 | 10666 | 0.4493 | 0.5699 | 0.5684 | 0.0046 | 0.6877 |
| CD227 | 4582 | 0.2539 | 0.2935 | 0.3209 | 0.0038 | 0.3633 |
| CD228 | 4241 | 0.0977 | 0.1279 | 0.2904 | 0.0067 | 0.5091 |
| CD229 | 4063 | 0.4910 | 0.6255 | 0.6065 | 0.0049 | 0.7315 |
| CD230 | 5621 | 0.4669 | 0.5790 | 0.5668 | 0.0047 | 0.6783 |
| CD231 | 7102 | 0.0694 | 0.1267 | 0.1527 | 0.0038 | 0.2059 |
| CD232 | 10154 | 0.3927 | 0.5165 | 0.5161 | 0.0050 | 0.6311 |
| CD233 | 6521 | 0.3742 | 0.4941 | 0.4807 | 0.0052 | 0.5946 |
| CD234 | 2532 | 0.2298 | 0.2971 | 0.3469 | 0.0051 | 0.4538 |
| CD235A | 2993 | 0.0264 | 0.0336 | 0.0949 | 0.0040 | 0.1100 |
| CD235B | 2994 | 0.2290 | 0.3563 | 0.3744 | 0.0056 | 0.5058 |
| CD236 | 2995 | 0.4735 | 0.5625 | 0.5534 | 0.0043 | 0.6455 |
| CD238 | 3792 | 0.2072 | 0.2995 | 0.3301 | 0.0045 | 0.4292 |
| CD239 | 4059 | 0.1152 | 0.1459 | 0.2366 | 0.0059 | 0.2638 |
| CD240CE | 6006 | 0.1861 | 0.2264 | 0.2542 | 0.0041 | 0.2675 |
| CD240D | 6007 | 0.0252 | 0.0322 | 0.1400 | 0.0050 | 0.2184 |
| CD241 | 6005 | 0.0282 | 0.0361 | 0.1449 | 0.0051 | 0.2316 |
| CD242 | 3386 | 0.3286 | 0.3667 | 0.3986 | 0.0041 | 0.4481 |
| CD243 | 5243 | 0.3582 | 0.4697 | 0.4731 | 0.0050 | 0.5995 |
| CD244 | 51744 | 0.5086 | 0.5471 | 0.5625 | 0.0044 | 0.6383 |
| CD246 | 238 | 0.0540 | 0.0687 | 0.1963 | 0.0064 | 0.3157 |
| CD247 | 919 | 0.5804 | 0.6672 | 0.6598 | 0.0039 | 0.7547 |
| CD248 | 57124 | 0.3734 | 0.4714 | 0.4798 | 0.0047 | 0.5894 |
| CD249 | 2028 | 0.0632 | 0.0860 | 0.2344 | 0.0064 | 0.4235 |
| CD252 | 7292 | 0.2576 | 0.3412 | 0.3685 | 0.0047 | 0.4500 |
| CD253 | 8743 | 0.3552 | 0.4537 | 0.4613 | 0.0046 | 0.5625 |
| CD254 | 8600 | 0.0778 | 0.1061 | 0.2277 | 0.0060 | 0.3639 |
| CD257 | 10673 | 0.4844 | 0.6047 | 0.5812 | 0.0048 | 0.6959 |
| CD258 | 8740 | 0.3544 | 0.5111 | 0.5044 | 0.0057 | 0.6435 |
| CD262 | 8795 | 0.3919 | 0.5091 | 0.5013 | 0.0049 | 0.6212 |
| CD263 | 8794 | 0.3718 | 0.4614 | 0.4833 | 0.0048 | 0.5685 |
| CD264 | 8793 | 0.0537 | 0.0768 | 0.2575 | 0.0069 | 0.4421 |
| CD265 | 8792 | 0.0607 | 0.0770 | 0.1617 | 0.0049 | 0.2216 |
| CD266 | 51330 | 0.2305 | 0.2770 | 0.3496 | 0.0051 | 0.4592 |
| CD267 | 23495 | 0.3001 | 0.3167 | 0.3391 | 0.0042 | 0.3625 |
| CD269 | 608 | 0.2179 | 0.3244 | 0.3582 | 0.0049 | 0.4692 |
| CD270 | 8764 | 0.3937 | 0.5311 | 0.5315 | 0.0053 | 0.6555 |
| CD271 | 4804 | 0.0484 | 0.0626 | 0.1994 | 0.0059 | 0.3569 |
| CD272 | 151888 | 0.3200 | 0.4516 | 0.4583 | 0.0049 | 0.5837 |
| CD273 | 80380 | 0.0000 | 0.0000 | 0.0919 | 0.0048 | 0.1376 |
| CD274 | 29126 | 0.0346 | 0.0441 | 0.1299 | 0.0049 | 0.1768 |
| CD276 | 80381 | 0.0618 | 0.0801 | 0.2286 | 0.0069 | 0.3964 |
| CD277 | 11119 | 0.3237 | 0.4063 | 0.4270 | 0.0044 | 0.5089 |
| CD278 | 29851 | 0.4065 | 0.5303 | 0.5204 | 0.0051 | 0.6423 |
| CD279 | 5133 | 0.0390 | 0.0499 | 0.1709 | 0.0061 | 0.2742 |
| CD280 | 9902 | 0.0455 | 0.2031 | 0.2347 | 0.0058 | 0.3882 |
| CD281 | 7096 | 0.3368 | 0.4997 | 0.4942 | 0.0060 | 0.6577 |
| CD282 | 7097 | 0.2973 | 0.4055 | 0.4288 | 0.0050 | 0.5405 |
| CD283 | 7098 | 0.0974 | 0.1225 | 0.2435 | 0.0059 | 0.3783 |
| CD284 | 7099 | 0.3382 | 0.4540 | 0.4687 | 0.0051 | 0.5907 |
| CD286 | 10333 | 0.3518 | 0.4914 | 0.4930 | 0.0054 | 0.6335 |
| CD288 | 51311 | 0.4240 | 0.5298 | 0.5305 | 0.0048 | 0.6437 |
| CD289 | 54106 | 0.3317 | 0.3715 | 0.3772 | 0.0042 | 0.3984 |
| CD290 | 81793 | 0.4269 | 0.5439 | 0.5341 | 0.0046 | 0.6416 |
| CD292 | 657 | 0.0928 | 0.1299 | 0.2889 | 0.0065 | 0.4903 |
| CDW293 | 658 | 0.0771 | 0.1042 | 0.2290 | 0.0054 | 0.3880 |
| CD294 | 11251 | 0.2721 | 0.2885 | 0.3274 | 0.0045 | 0.3827 |
| CD295 | 3953 | 0.4433 | 0.5012 | 0.5038 | 0.0041 | 0.5759 |
| CD296 | 417 | 0.3463 | 0.3758 | 0.4412 | 0.0044 | 0.5359 |
| CD297 | 420 | 0.0485 | 0.0637 | 0.2164 | 0.0062 | 0.3780 |
| CD298 | 483 | 0.3326 | 0.4407 | 0.4449 | 0.0047 | 0.5547 |
| CD299 | 10332 | 0.0339 | 0.0427 | 0.1953 | 0.0067 | 0.3262 |
| CD300A | 11314 | 0.4993 | 0.6021 | 0.5909 | 0.0045 | 0.6974 |
| CD300C | 10871 | 0.4161 | 0.5457 | 0.5341 | 0.0051 | 0.6624 |
| CD301 | 10462 | 0.4865 | 0.5319 | 0.5278 | 0.0044 | 0.6119 |
| CD304 | 8829 | 0.0821 | 0.1085 | 0.2568 | 0.0065 | 0.4547 |
| CD305 | 3903 | 0.3634 | 0.4618 | 0.4744 | 0.0044 | 0.5776 |
| CD306 | 3904 | 0.2039 | 0.2878 | 0.3070 | 0.0044 | 0.3783 |
| CD307A | 115350 | 0.0862 | 0.1091 | 0.1929 | 0.0053 | 0.2121 |
| CD307B | 79368 | 0.3870 | 0.5031 | 0.4953 | 0.0047 | 0.6056 |
| CD307C | 115352 | 0.4221 | 0.5118 | 0.5191 | 0.0044 | 0.6154 |
| CD307D | 83417 | 0.0523 | 0.0663 | 0.1390 | 0.0046 | 0.1500 |
| CD307E | 83416 | 0.2651 | 0.3560 | 0.3732 | 0.0044 | 0.4562 |
| CD308 | 2321 | 0.1455 | 0.1845 | 0.2555 | 0.0054 | 0.2298 |
| CD309 | 3791 | 0.0792 | 0.1003 | 0.1694 | 0.0045 | 0.1944 |
| CD312 | 30817 | 0.4273 | 0.5262 | 0.5254 | 0.0046 | 0.6323 |
| CD315 | 5738 | 0.0863 | 0.1089 | 0.1899 | 0.0050 | 0.2524 |
| CD316 | 93185 | 0.1475 | 0.5214 | 0.4480 | 0.0070 | 0.6486 |
| CD317 | 684 | 0.4043 | 0.5385 | 0.5355 | 0.0054 | 0.6746 |
| CD318 | 64866 | 0.0909 | 0.2574 | 0.3190 | 0.0070 | 0.5427 |
| CD319 | 57823 | 0.3282 | 0.4635 | 0.4620 | 0.0048 | 0.5813 |
| CD320 | 51293 | 0.4975 | 0.5343 | 0.5618 | 0.0042 | 0.6432 |
| CD321 | 50848 | 0.3659 | 0.4622 | 0.4825 | 0.0047 | 0.5838 |
| CD322 | 58494 | 0.0578 | 0.0736 | 0.2095 | 0.0065 | 0.3546 |
| CD324 | 999 | 0.0830 | 0.1046 | 0.1786 | 0.0048 | 0.1851 |
| CD325 | 1000 | 0.1798 | 0.2115 | 0.2514 | 0.0044 | 0.2504 |
| CD326 | 4072 | 0.0374 | 0.0473 | 0.1227 | 0.0043 | 0.1631 |
| CD327 | 946 | 0.1152 | 0.4171 | 0.3868 | 0.0067 | 0.5691 |
| CD328 | 27036 | 0.3945 | 0.4399 | 0.4761 | 0.0041 | 0.5506 |
| CD329 | 27180 | 0.3735 | 0.4845 | 0.4874 | 0.0050 | 0.5985 |
| CD331 | 2260 | 0.0623 | 0.0815 | 0.1848 | 0.0057 | 0.2825 |
| CD332 | 2263 | 0.1078 | 0.1379 | 0.1910 | 0.0046 | 0.1685 |
| CD333 | 2261 | 0.0938 | 0.1198 | 0.2726 | 0.0064 | 0.4759 |
| CD334 | 2264 | 0.1030 | 0.1317 | 0.2100 | 0.0055 | 0.1767 |
| CD335 | 9437 | 0.0930 | 0.1176 | 0.2136 | 0.0056 | 0.3205 |
| CD336 | 9436 | 0.0406 | 0.0514 | 0.1626 | 0.0057 | 0.2566 |
| CD337 | 259197 | 0.3529 | 0.4894 | 0.4933 | 0.0046 | 0.6105 |
| CD338 | 9429 | 0.0000 | 0.0000 | 0.0981 | 0.0052 | 0.0917 |
| CD339 | 182 | 0.0478 | 0.0622 | 0.1937 | 0.0061 | 0.3276 |
| CD340 | 2064 | 0.4351 | 0.4842 | 0.4856 | 0.0044 | 0.5603 |
| CD344 | 8322 | 0.0000 | 0.0000 | 0.1506 | 0.0071 | 0.3500 |
| CD349 | 8326 | 0.0453 | 0.0572 | 0.1384 | 0.0049 | 0.1556 |
| CD350 | 11211 | 0.0368 | 0.0508 | 0.2128 | 0.0066 | 0.3945 |
| CD351 | 8395 | 0.2458 | 0.3268 | 0.3550 | 0.0044 | 0.4425 |
| CD353 | 56833 | 0.0388 | 0.0496 | 0.1923 | 0.0060 | 0.3306 |
| CD354 | 54210 | 0.4307 | 0.5610 | 0.5460 | 0.0052 | 0.6796 |
| CD355 | 56253 | 0.0568 | 0.0724 | 0.1858 | 0.0061 | 0.2610 |
| CD357 | 8784 | 0.0684 | 0.0893 | 0.2214 | 0.0061 | 0.3834 |
| CD358 | 27242 | 0.3184 | 0.3443 | 0.3893 | 0.0049 | 0.4793 |
| CD360 | 50615 | 0.5752 | 0.6418 | 0.6174 | 0.0045 | 0.7088 |
| CD361 | 2124 | 0.2721 | 0.3649 | 0.3831 | 0.0047 | 0.4680 |
| CD362 | 6383 | 0.2068 | 0.2533 | 0.3137 | 0.0050 | 0.3991 |
| CD363 | 1901 | 0.4069 | 0.5292 | 0.5368 | 0.0051 | 0.6612 |
| CD364 | 221476 | 0.4220 | 0.4661 | 0.4906 | 0.0047 | 0.5900 |
| CD366 | 84868 | 0.4126 | 0.5178 | 0.5167 | 0.0046 | 0.6249 |
| CD367 | 50856 | 0.5184 | 0.6120 | 0.6118 | 0.0042 | 0.7098 |
| CD369 | 64581 | 0.4805 | 0.5836 | 0.5811 | 0.0049 | 0.6980 |

By default, CD gene expression was stratified at 20 and 80 percentiles, with low expression being between 0 and 20, average between 20 and 80, and high between 80 and 100. Two-tailed thresholds were labeled as 'aberrant' or 'non-aberrant', with 'aberrant' being further delineated as 'low' or 'high'.

**Supplement 2. Table 3.** Clusters of differentiation gene expression requiring. Feature transformation included threshold shifting (n=3) or binary dichotomization (n=85).

| Cluster of Differentiation | Entrez ID | Feature transformation | Threshold | Figure # |
| --- | --- | --- | --- | --- |
| CD34 | 947 | Dichotomization | 0.18 | 38 |
| CD49B | 3673 | Dichotomization | 0.1 | 54 |
| CD49C | 3675 | Dichotomization | 0.18 | 55 |
| CD57 | 27087 | Dichotomization | 0.2 | 66 |
| CD62E | 6401 | Dichotomization | 0.2 | 70 |
| CD62L | 6402 | Dichotomization | 0.7 | 71 |
| CD66E | 1048 | Dichotomization | 0.12 | 78 |
| CD66F | 5669 | Dichotomization | 0.15 | 79 |
| CD72 | 971 | First threshold shifted | 0.35 | 84 |
| CD80 | 941 | Dichotomization | 0.1 | 89 |
| CD85C | 10990 | Dichotomization | 0.2 | 95 |
| CD85M | 79166 | Dichotomization | 0.2 | 102 |
| CD91 | 4035 | Dichotomization | 0.45 | 108 |
| CD104 | 3691 | Dichotomization | 0.2 | 121 |
| CD105 | 2022 | First threshold shifted | 0.3 | 122 |
| CD106 | 7412 | Dichotomization | 0.2 | 123 |
| CD108 | 8482 | Dichotomization | 0.15 | 126 |
| CD111 | 5818 | Dichotomization | 0.2 | 129 |
| CD113 | 25945 | Dichotomization | 0.15 | 131 |
| CD118 | 3977 | Dichotomization | 0.18 | 136 |
| CD121A | 3554 | Dichotomization | 0.1 | 140 |
| CD133 | 8842 | Dichotomization | 0.1 | 151 |
| CD135 | 2322 | Dichotomization | 0.1 | 153 |
| CD140A | 5156 | Dichotomization | 0.2 | 157 |
| CD142 | 2152 | Dichotomization | 0.2 | 160 |
| CD143 | 1636 | Dichotomization | 0.2 | 161 |
| CD144 | 1003 | Dichotomization | 0.25 | 162 |
| CD146 | 4162 | Dichotomization | 0.2 | 163 |
| CD154 | 959 | Dichotomization | 0.35 | 170 |
| CD167B | 4921 | Dichotomization | 0.2 | 185 |
| CD168 | 3161 | Dichotomization | 0.15 | 186 |
| CD169 | 6614 | Dichotomization | 0.1 | 187 |
| CD171 | 3897 | Dichotomization | 0.2 | 189 |
| CD179A | 7441 | Dichotomization | 0.15 | 195 |
| CD180 | 4064 | Dichotomization | 0.2 | 197 |
| CD194 | 1233 | Dichotomization | 0.1 | 206 |
| CD200 | 4345 | Dichotomization | 0.2 | 211 |
| CD202B | 7010 | Dichotomization | 0.15 | 213 |
| CD203C | 5169 | Dichotomization | 0.18 | 214 |
| CD204 | 4481 | Dichotomization | 0.2 | 215 |
| CD206 | 4360 | Dichotomization | 0.15 | 217 |
| CD207 | 50489 | Dichotomization | 0.15 | 218 |
| CD213A2 | 3598 | Dichotomization | 0.15 | 225 |
| CD215 | 3601 | Dichotomization | 0.2 | 226 |
| CD220 | 3643 | Dichotomization | 0.15 | 230 |
| CD225 | 8519 | Dichotomization | 0.63 | 234 |
| CD228 | 4241 | Dichotomization | 0.2 | 237 |
| CD231 | 7102 | First threshold shifted | 0.05 | 240 |
| CD235A | 2993 | Dichotomization | 0.07 | 244 |
| CD239 | 4059 | Dichotomization | 0.2 | 248 |
| CD240D | 6007 | Dichotomization | 0.1 | 250 |
| CD241 | 6005 | Dichotomization | 0.1 | 251 |
| CD246 | 238 | Dichotomization | 0.2 | 255 |
| CD249 | 2028 | Dichotomization | 0.2 | 258 |
| CD254 | 8600 | Dichotomization | 0.2 | 261 |
| CD264 | 8793 | Dichotomization | 0.15 | 266 |
| CD265 | 8792 | Dichotomization | 0.15 | 267 |
| CD271 | 4804 | Dichotomization | 0.15 | 272 |
| CD274 | 29126 | Dichotomization | 0.1 | 275 |
| CD276 | 80381 | Dichotomization | 0.2 | 276 |
| CD279 | 5133 | Dichotomization | 0.1 | 279 |
| CD280 | 9902 | Dichotomization | 0.15 | 280 |
| CD283 | 7098 | Dichotomization | 0.2 | 283 |
| CD292 | 657 | Dichotomization | 0.2 | 289 |
| CD297 | 420 | Dichotomization | 0.2 | 290 |
| CD299 | 10332 | Dichotomization | 0.1 | 294 |
| CD304 | 8829 | Dichotomization | 0.2 | 296 |
| CD307A | 115350 | Dichotomization | 0.2 | 300 |
| CD307D | 83417 | Dichotomization | 0.1 | 303 |
| CD309 | 3791 | Dichotomization | 0.2 | 306 |
| CD315 | 5738 | Dichotomization | 0.2 | 309 |
| CD316 | 93185 | Dichotomization | 0.2 | 311 |
| CD318 | 64866 | Dichotomization | 0.2 | 312 |
| CD322 | 58494 | Dichotomization | 0.15 | 314 |
| CD324 | 999 | Dichotomization | 0.2 | 318 |
| CD326 | 4072 | Dichotomization | 0.1 | 319 |
| CD331 | 2260 | Dichotomization | 0.2 | 321 |
| CD333 | 2261 | Dichotomization | 0.2 | 325 |
| CD334 | 2264 | Dichotomization | 0.2 | 327 |
| CD335 | 9437 | Dichotomization | 0.2 | 328 |
| CD336 | 9436 | Dichotomization | 0.1 | 329 |
| CD339 | 182 | Dichotomization | 0.1 | 330 |
| CD349 | 8326 | Dichotomization | 0.1 | 333 |
| CD350 | 11211 | Dichotomization | 0.1 | 336 |
| CD353 | 56833 | Dichotomization | 0.1 | 337 |
| CD355 | 56253 | Dichotomization | 0.15 | 339 |
| CD357 | 8784 | Dichotomization | 0.15 | 341 |
| CDW293 | 658 | Dichotomization | 0.2 | 342 |

**Supplement 3. Figures 1 – 351.**
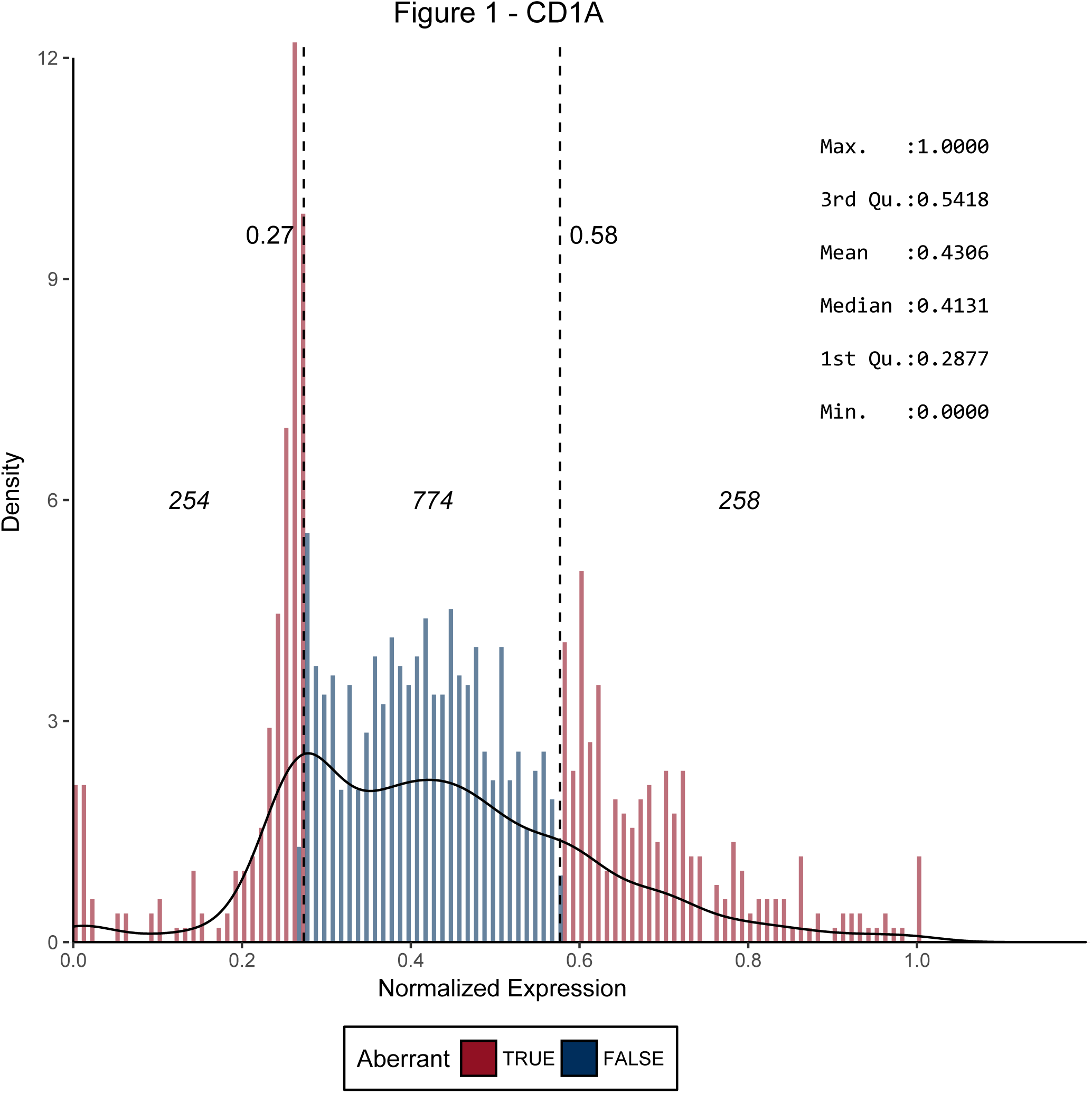
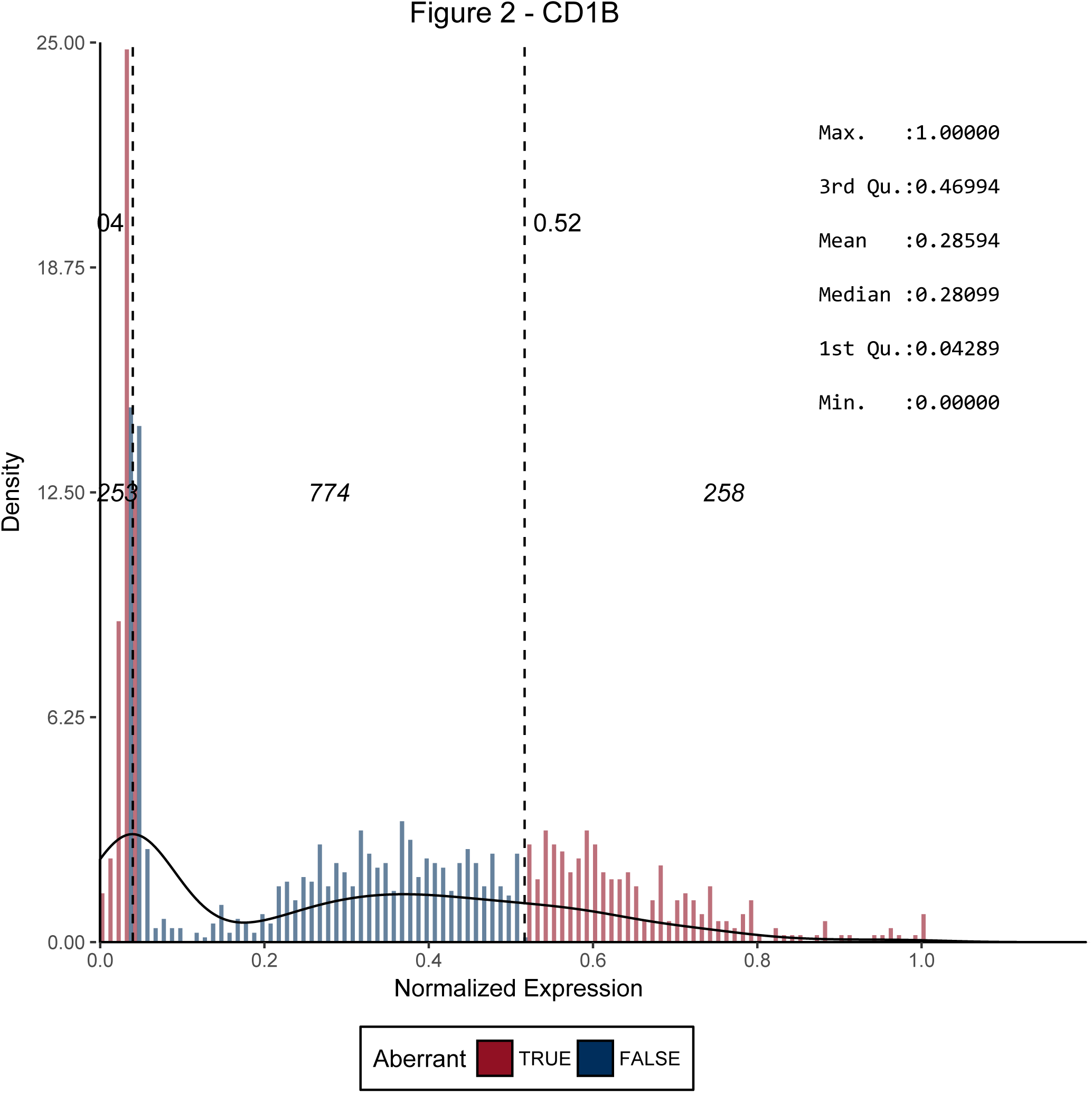
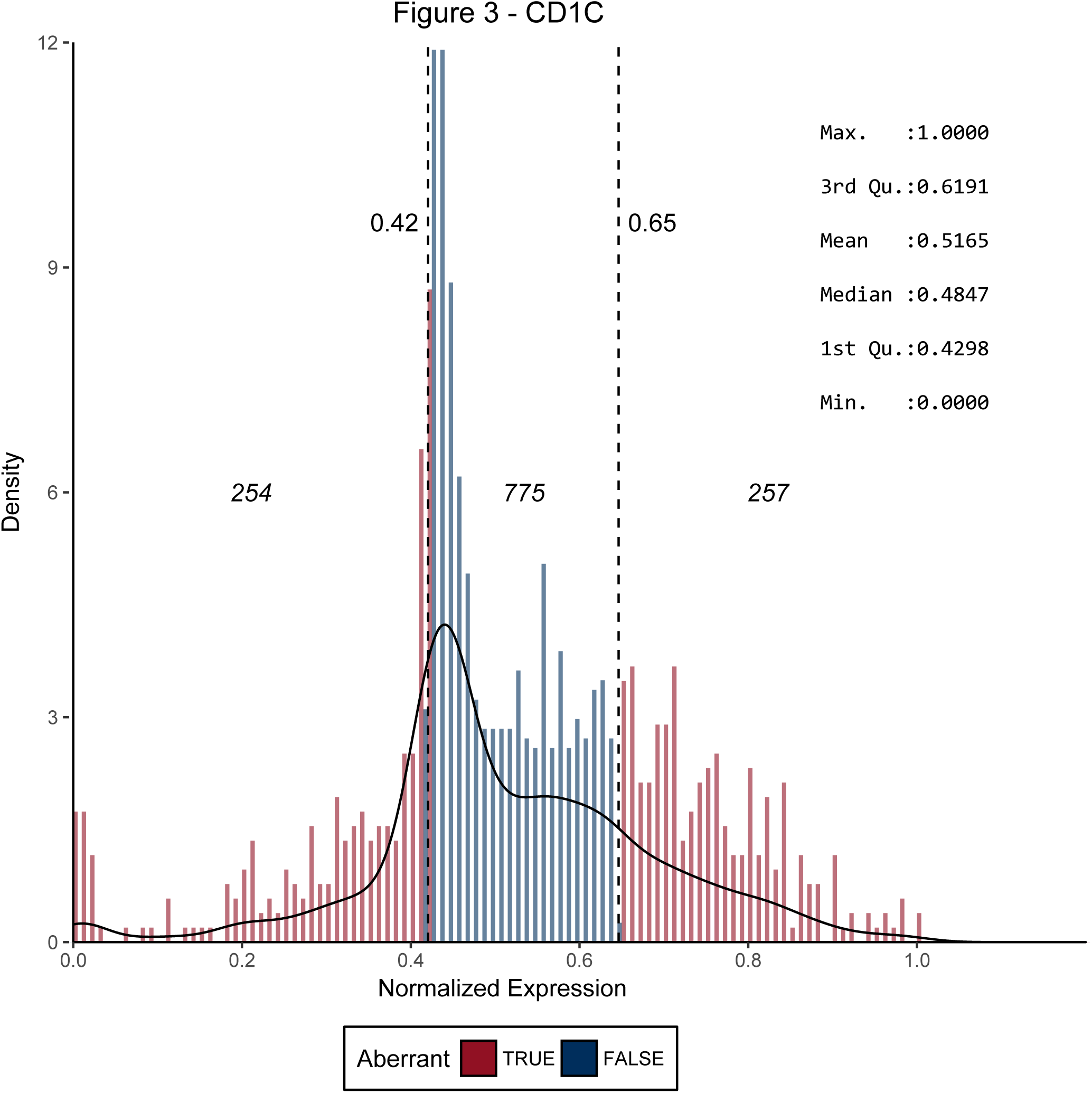
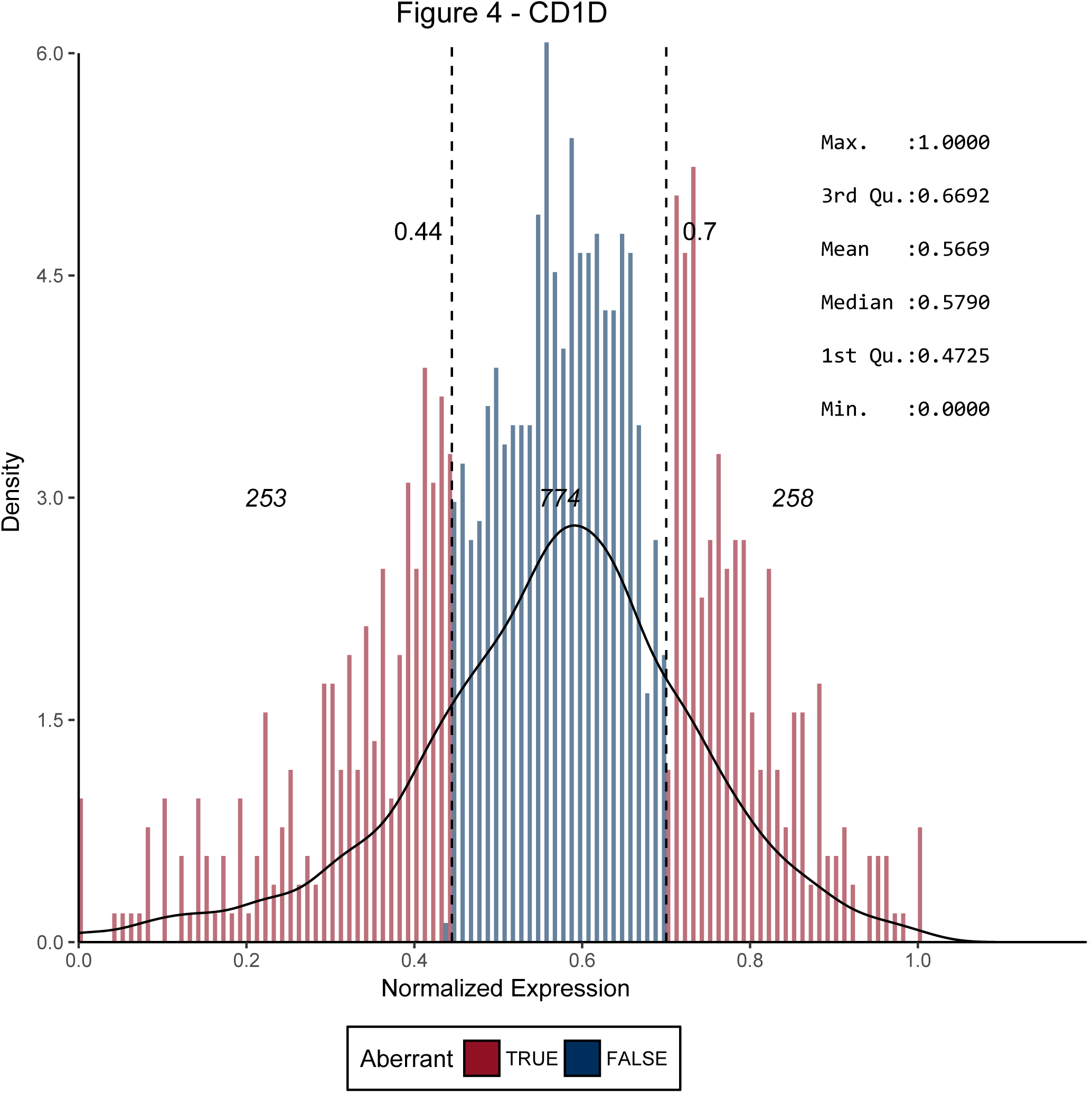
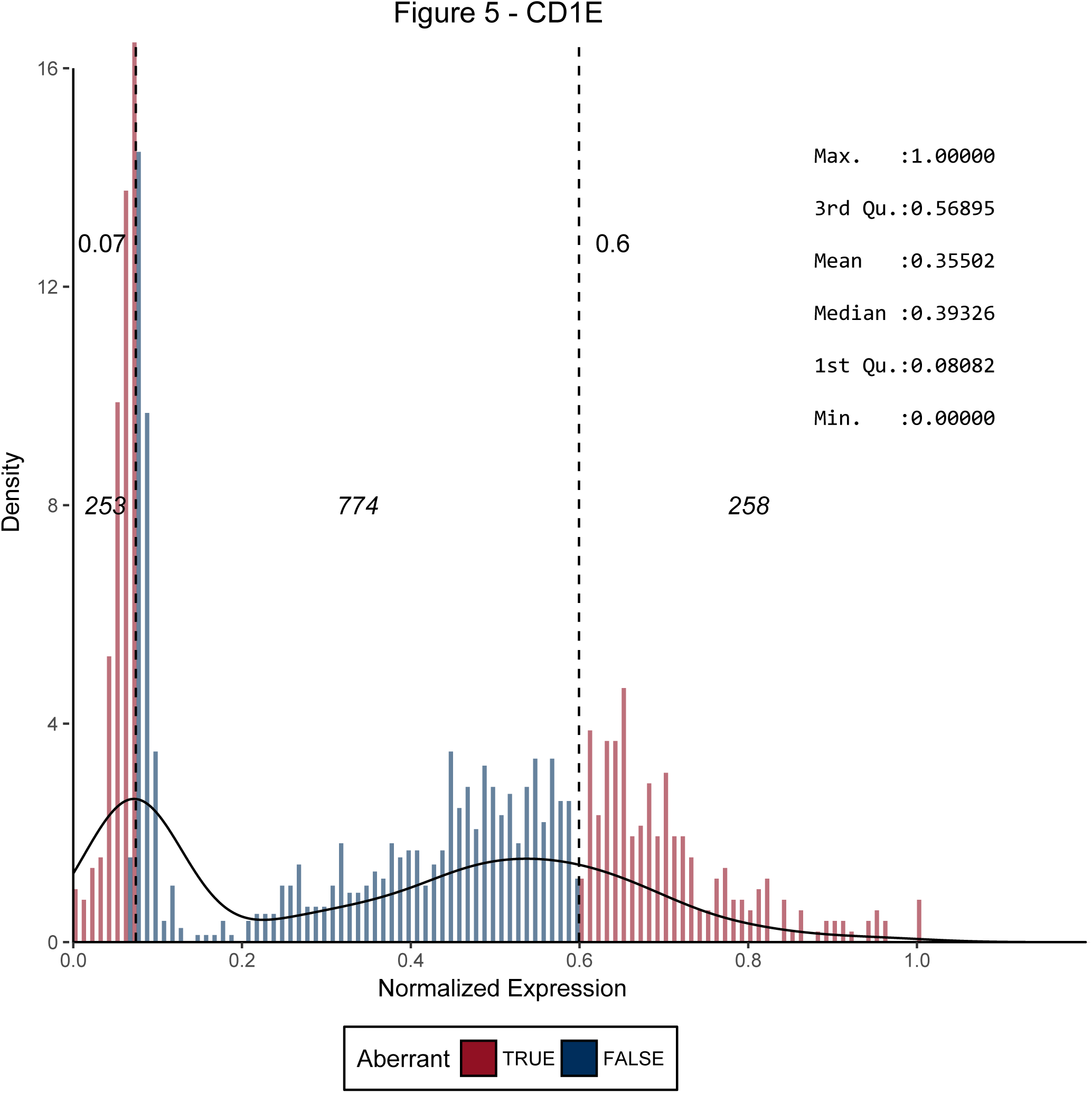
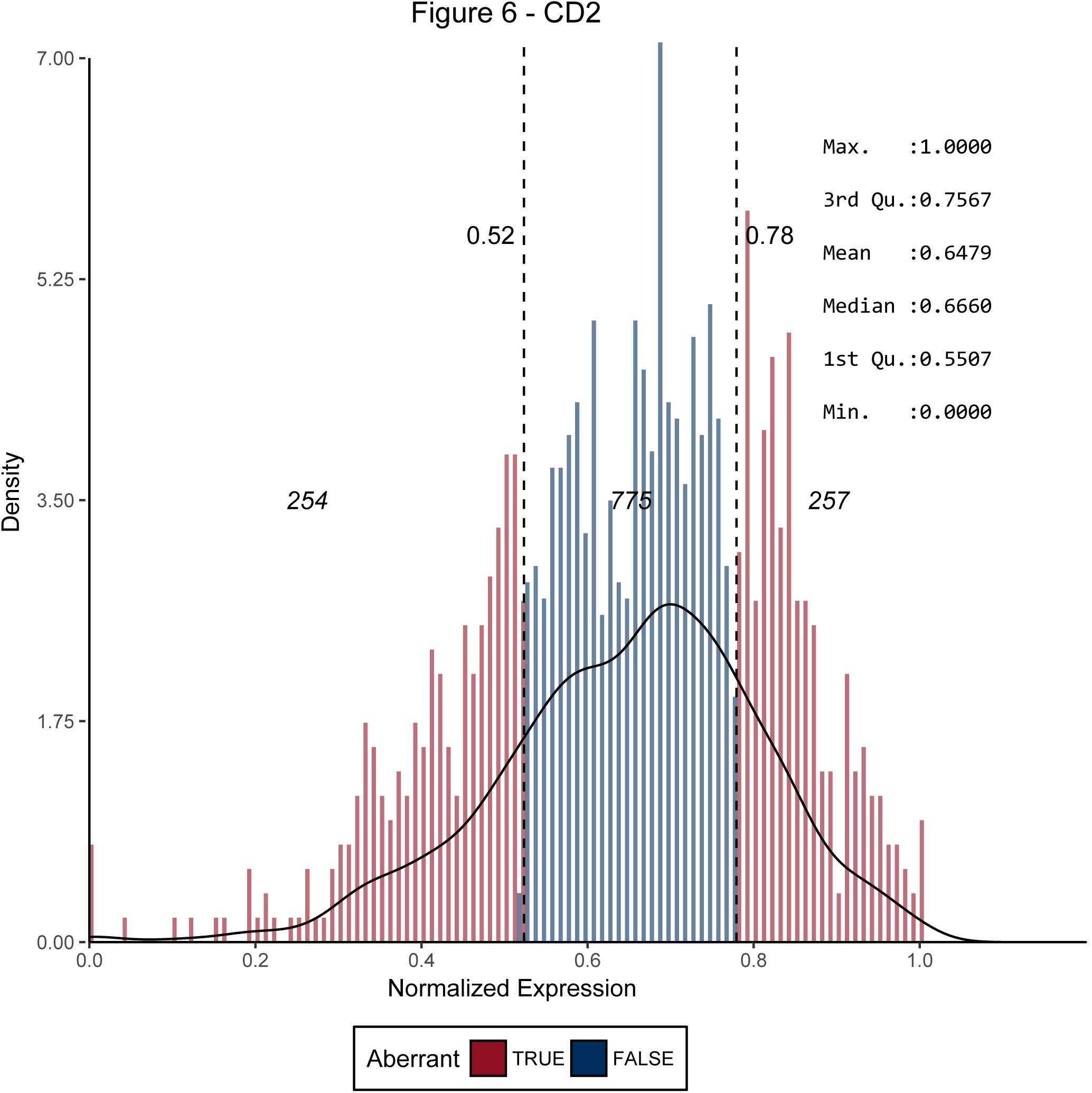
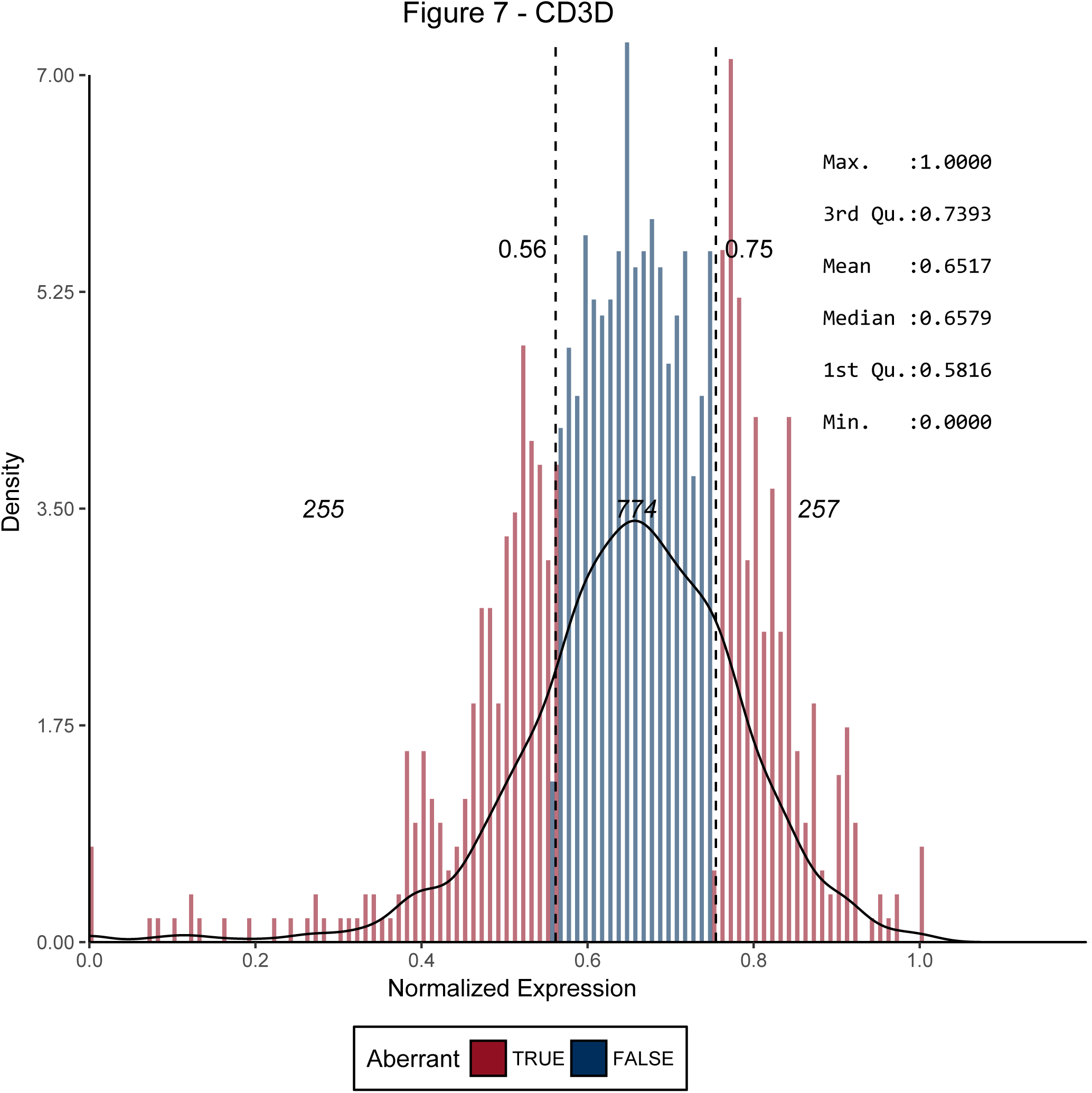
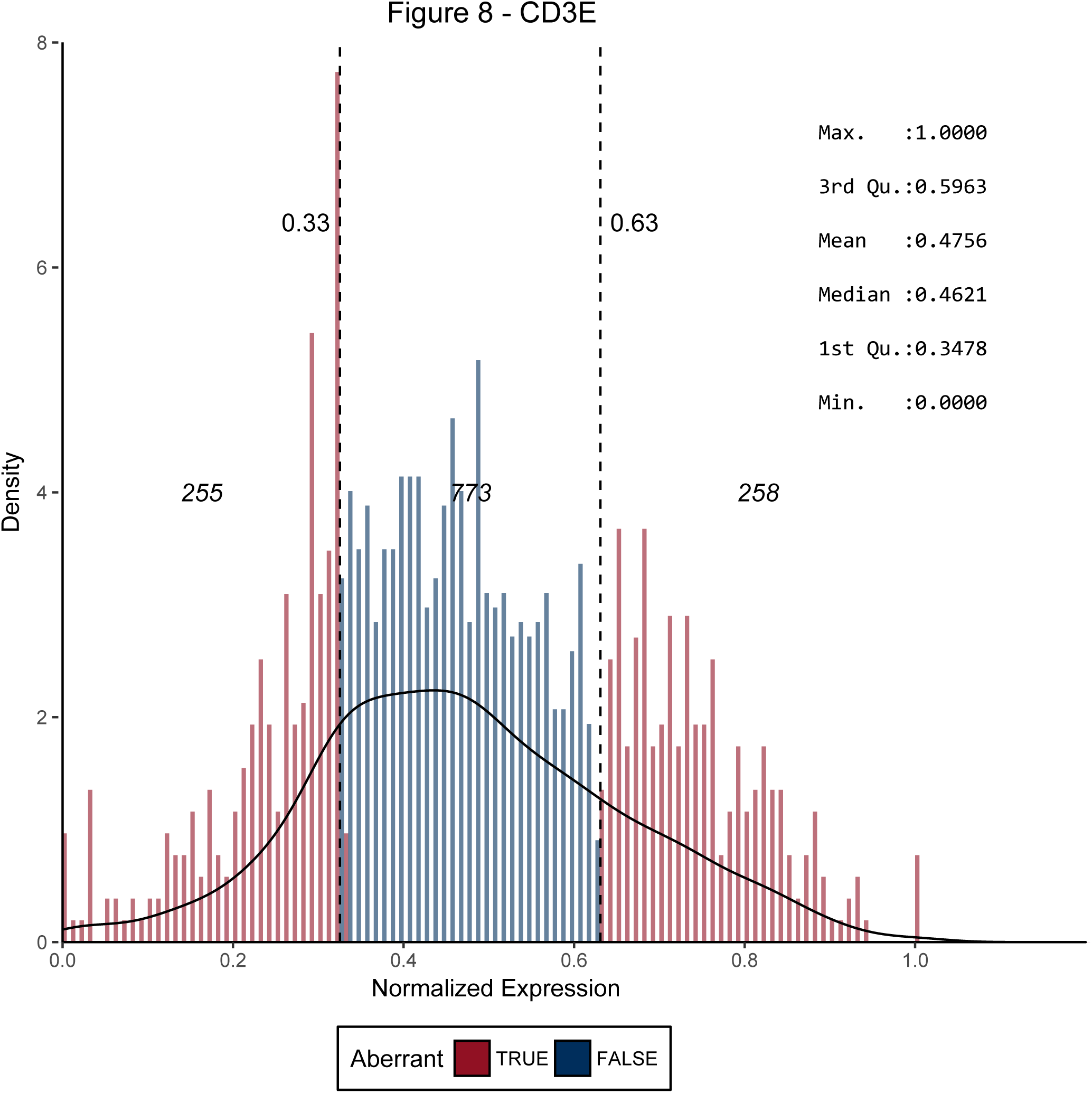
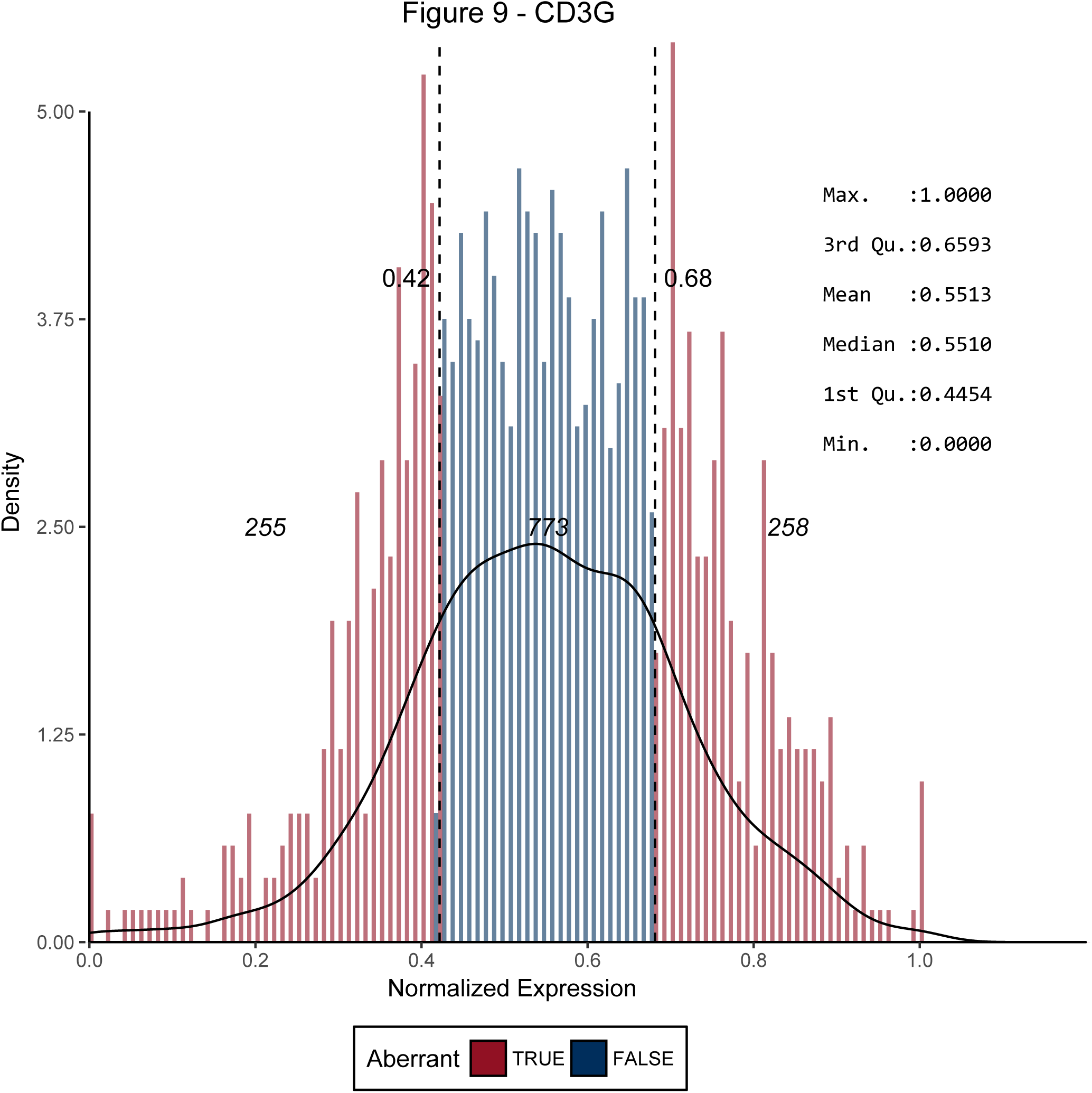
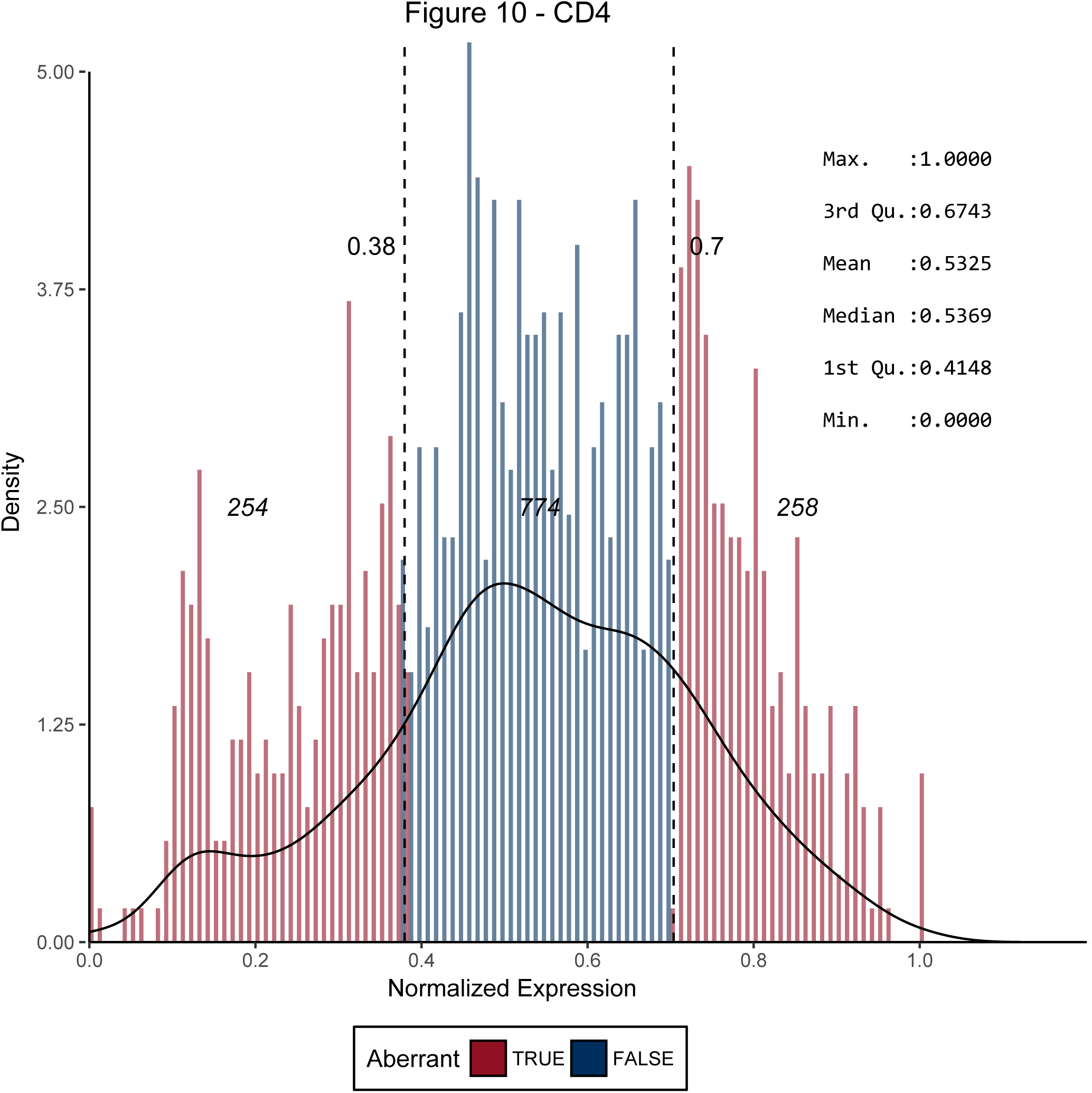
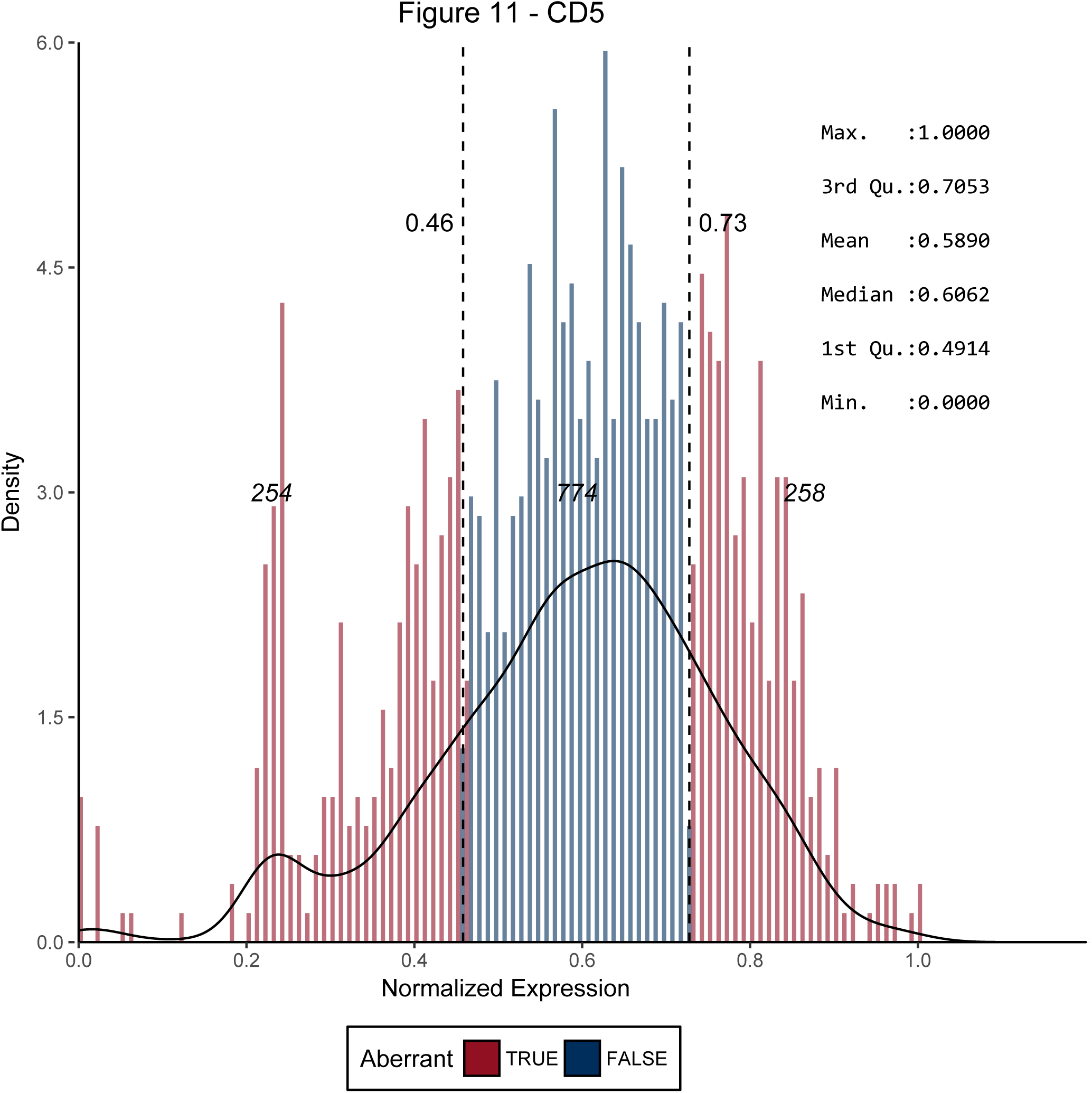
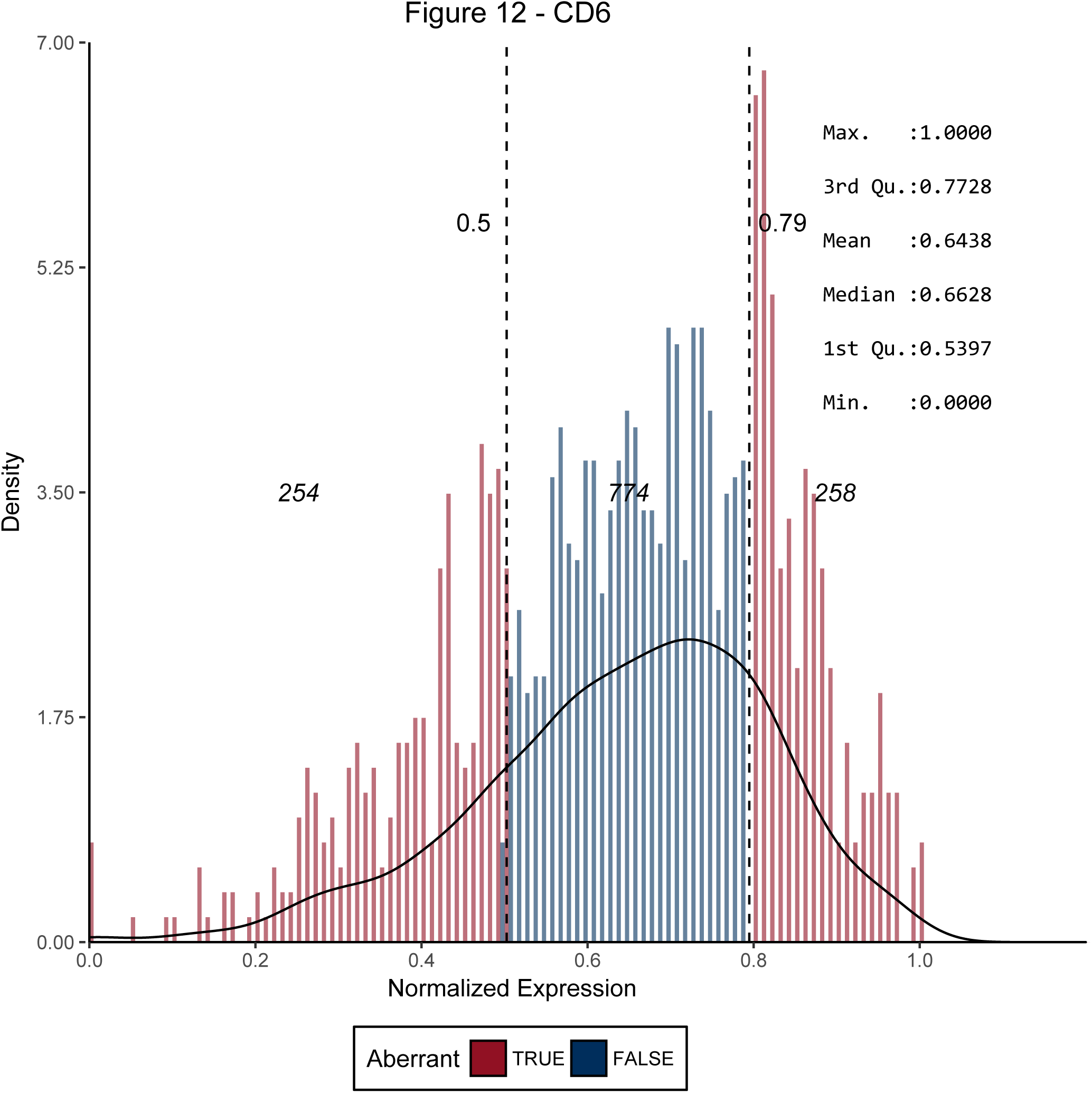
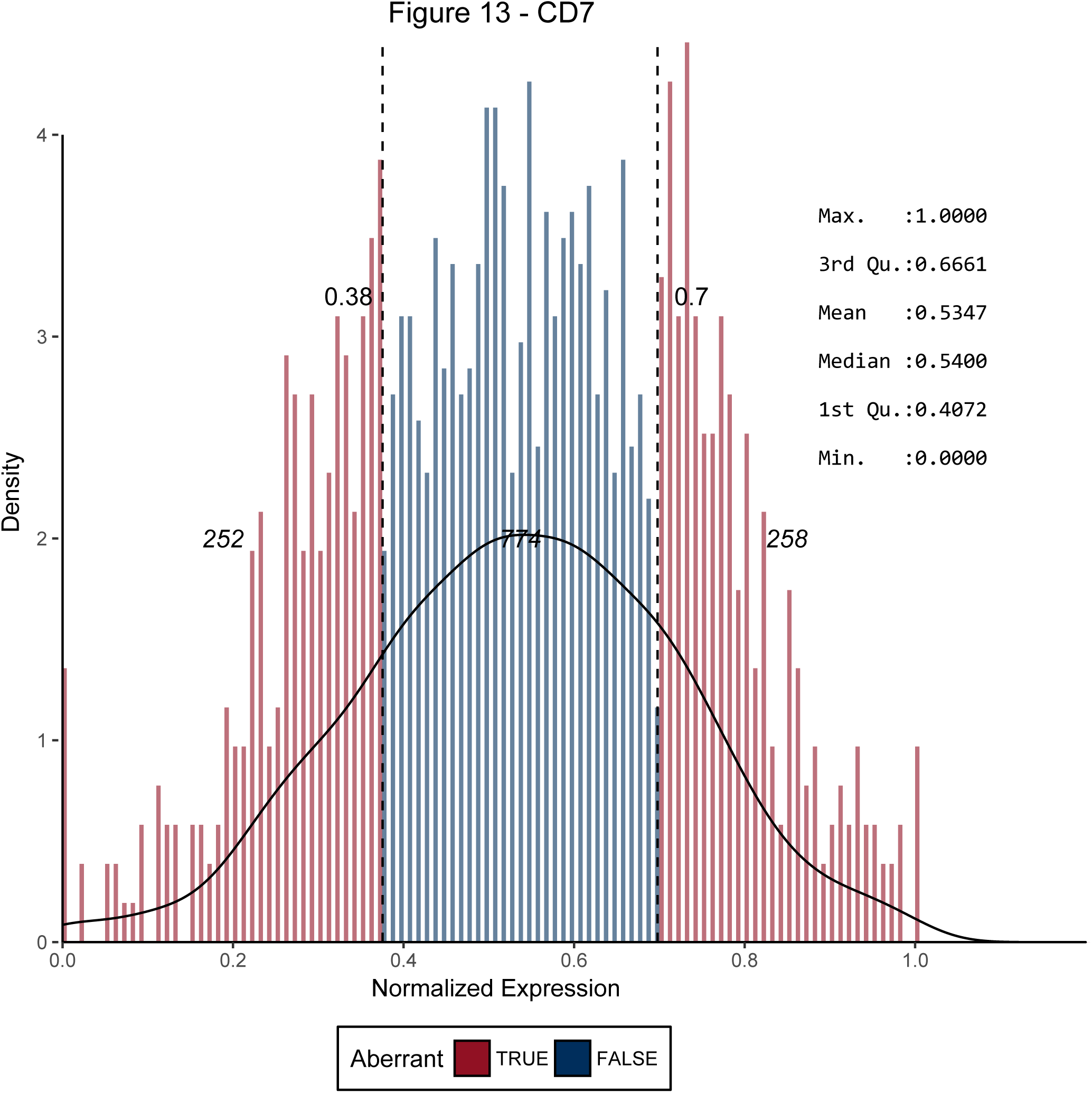
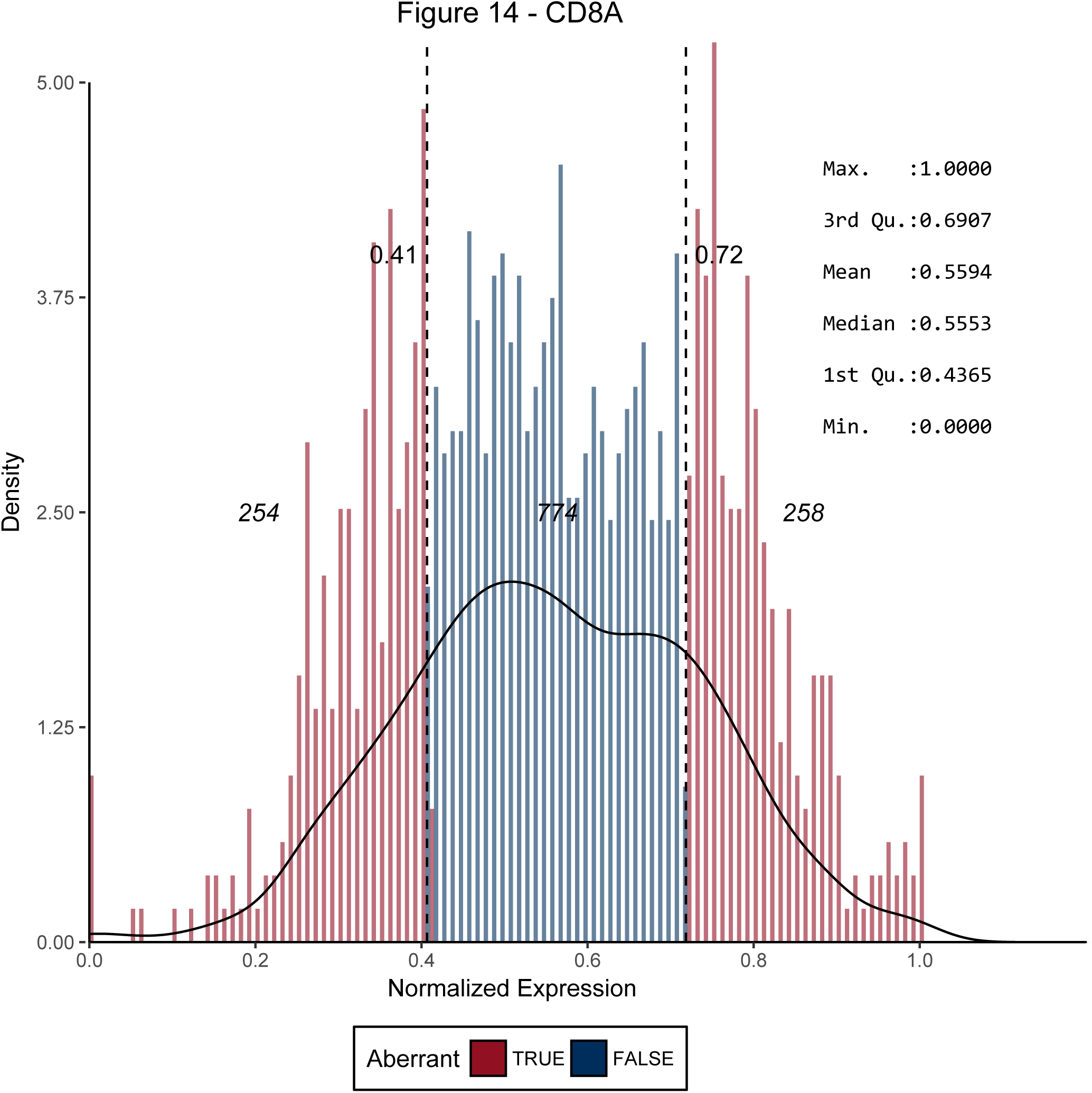
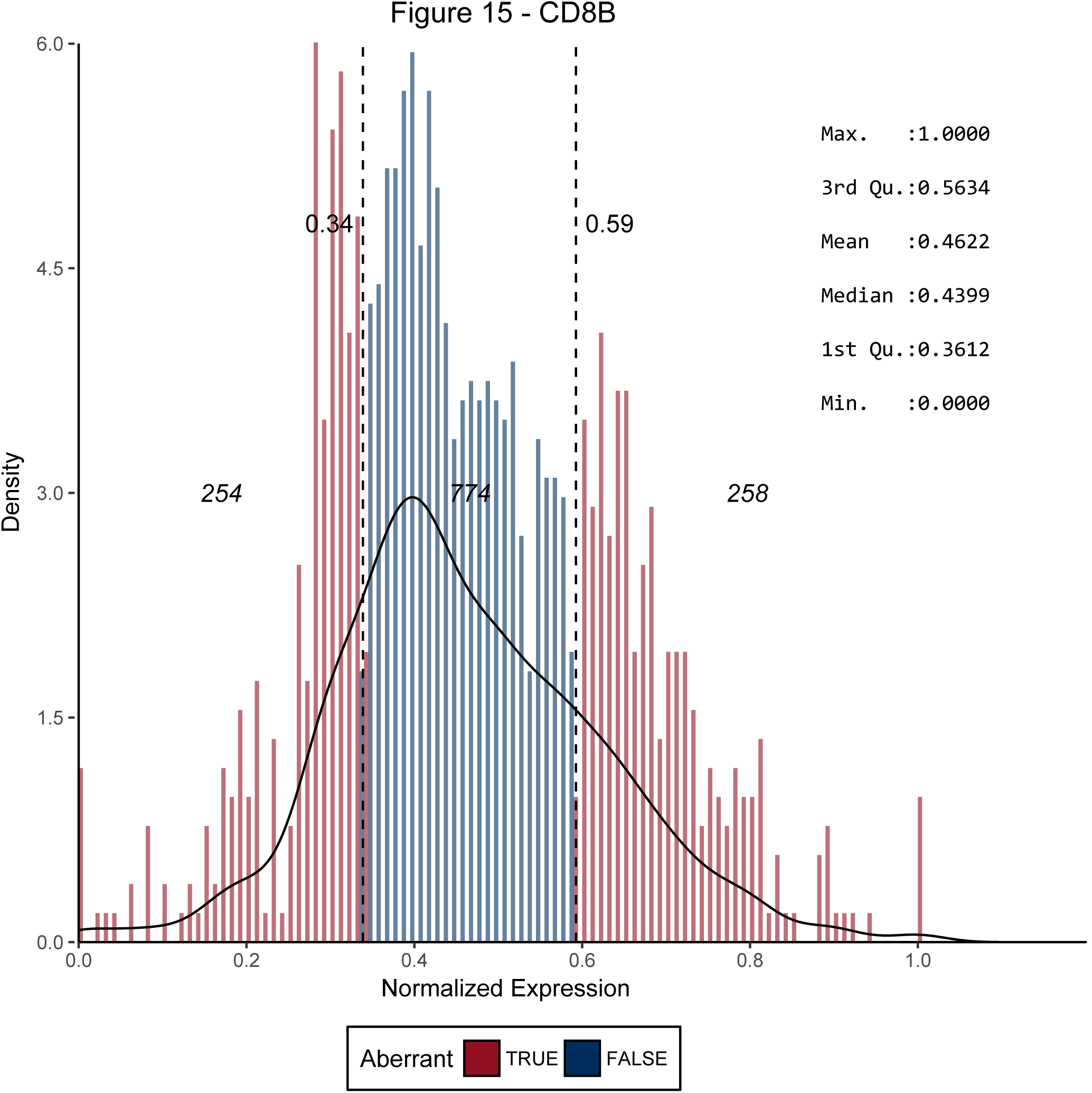
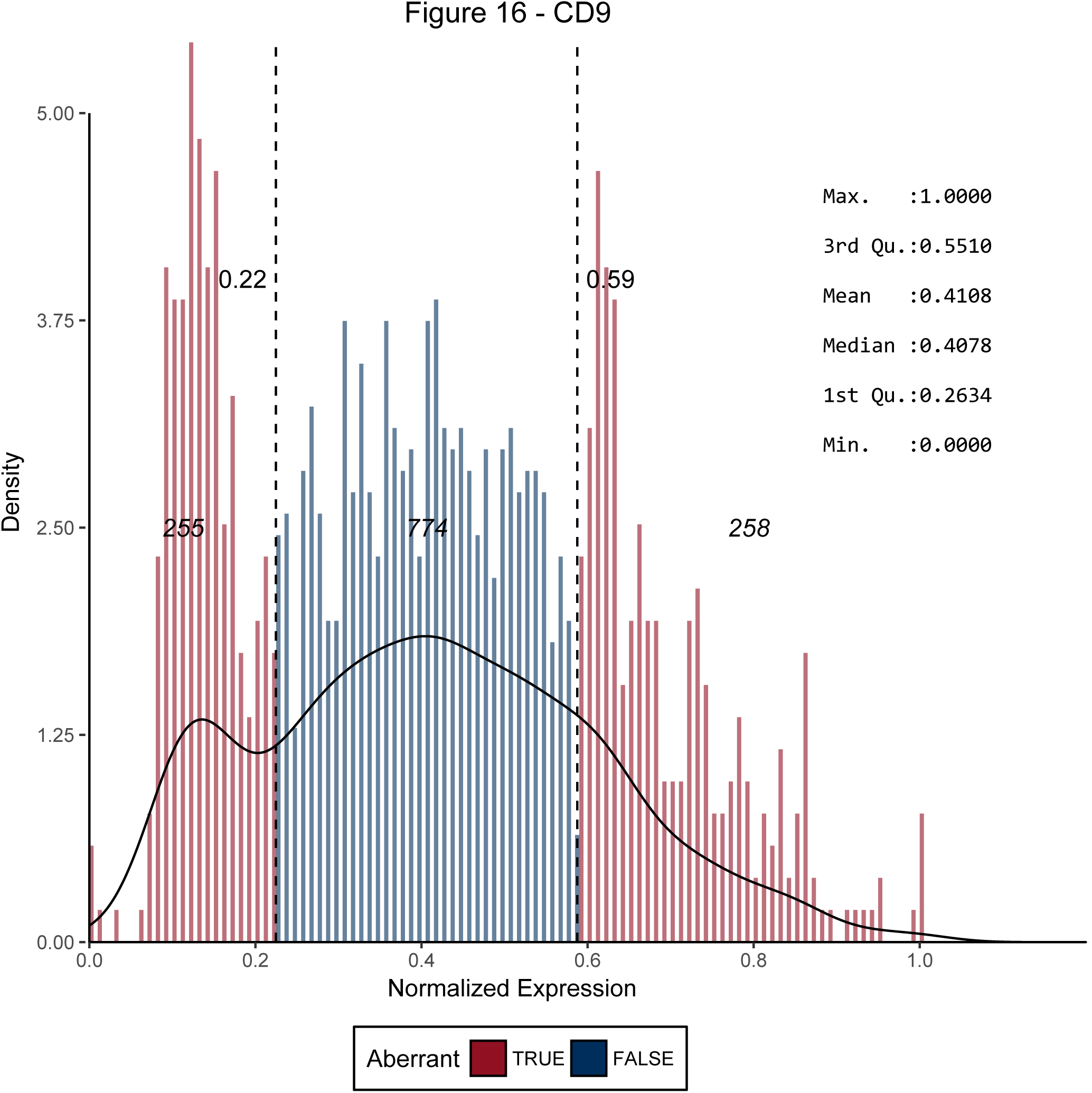
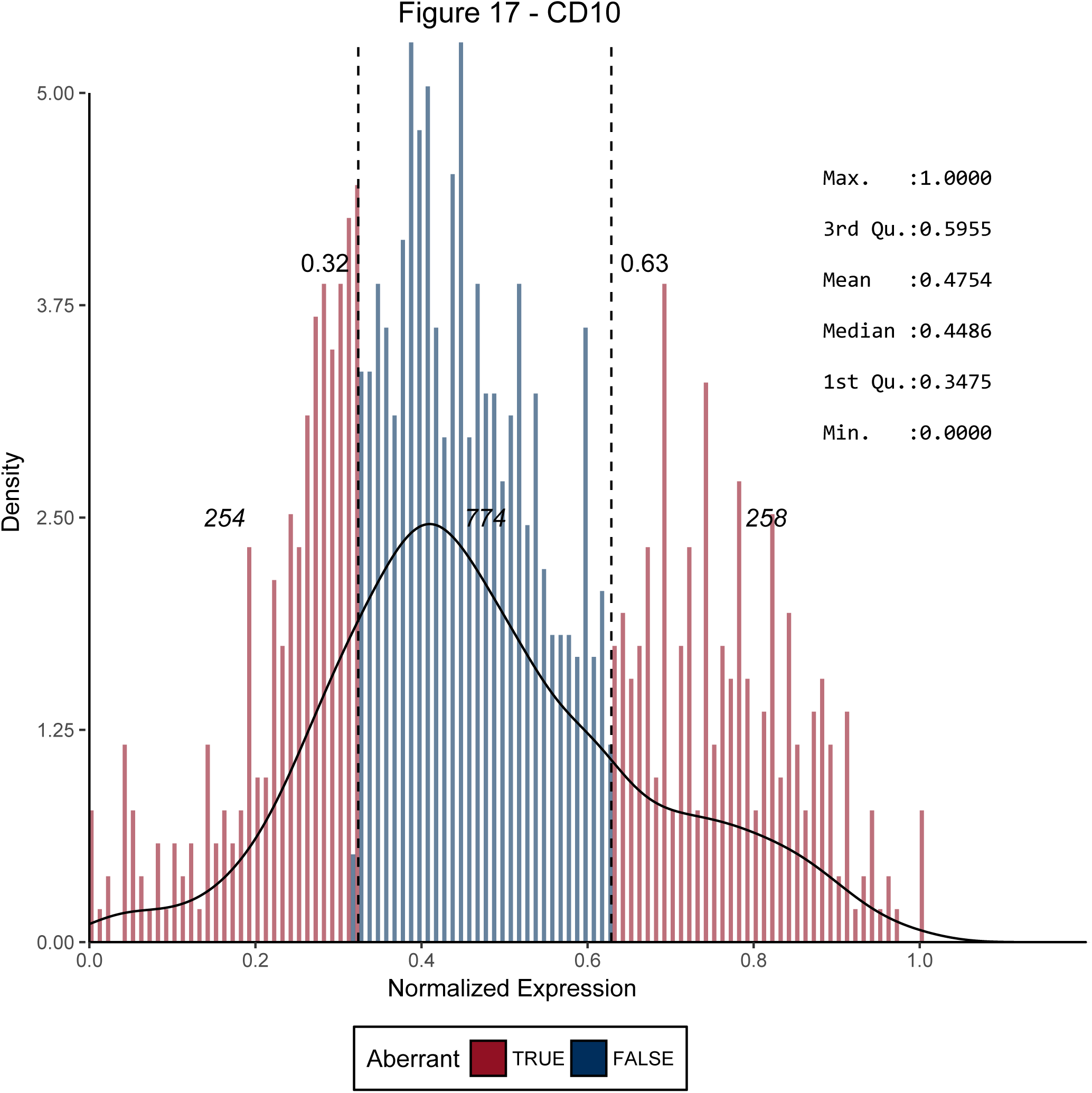
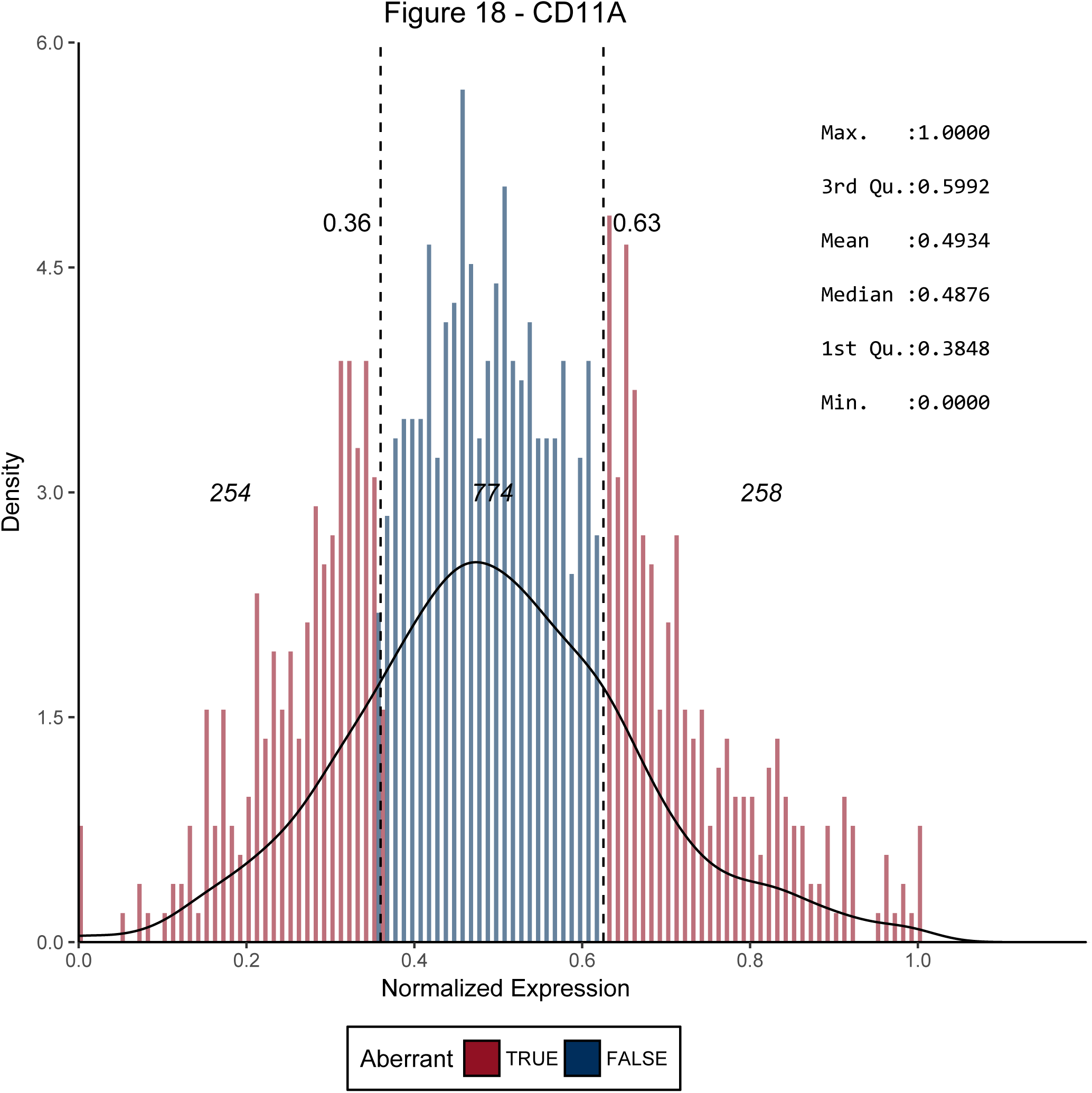
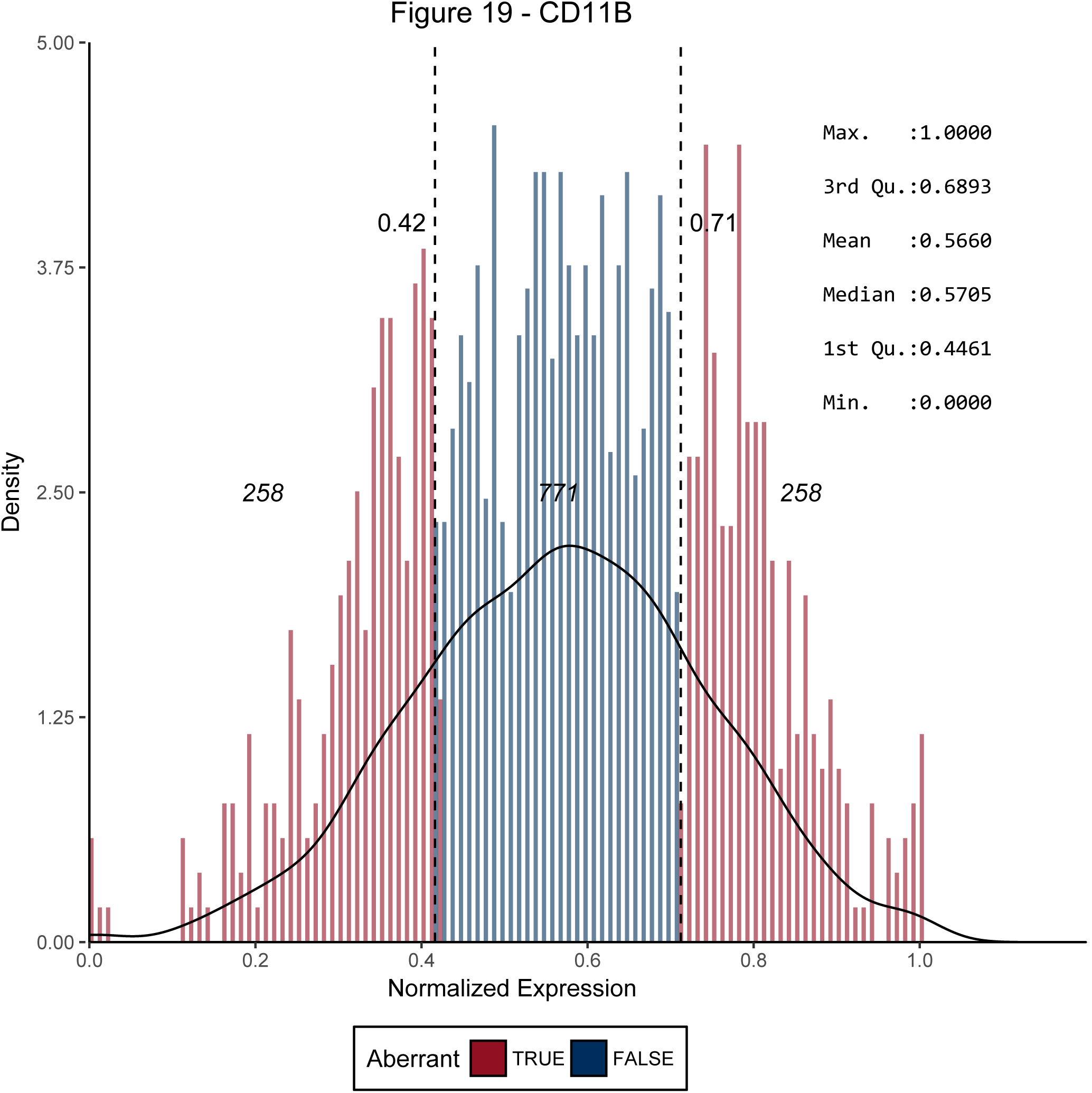
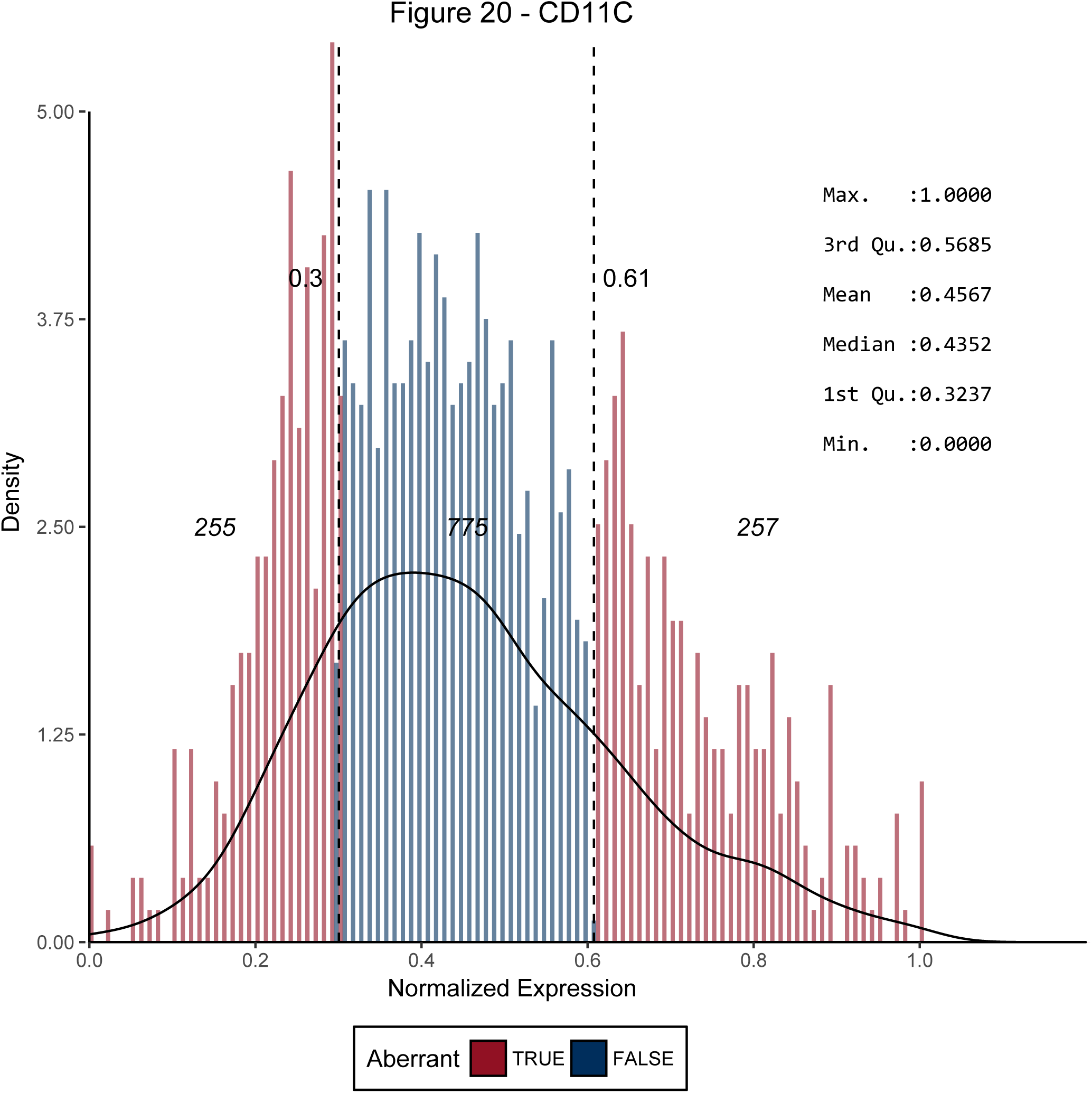
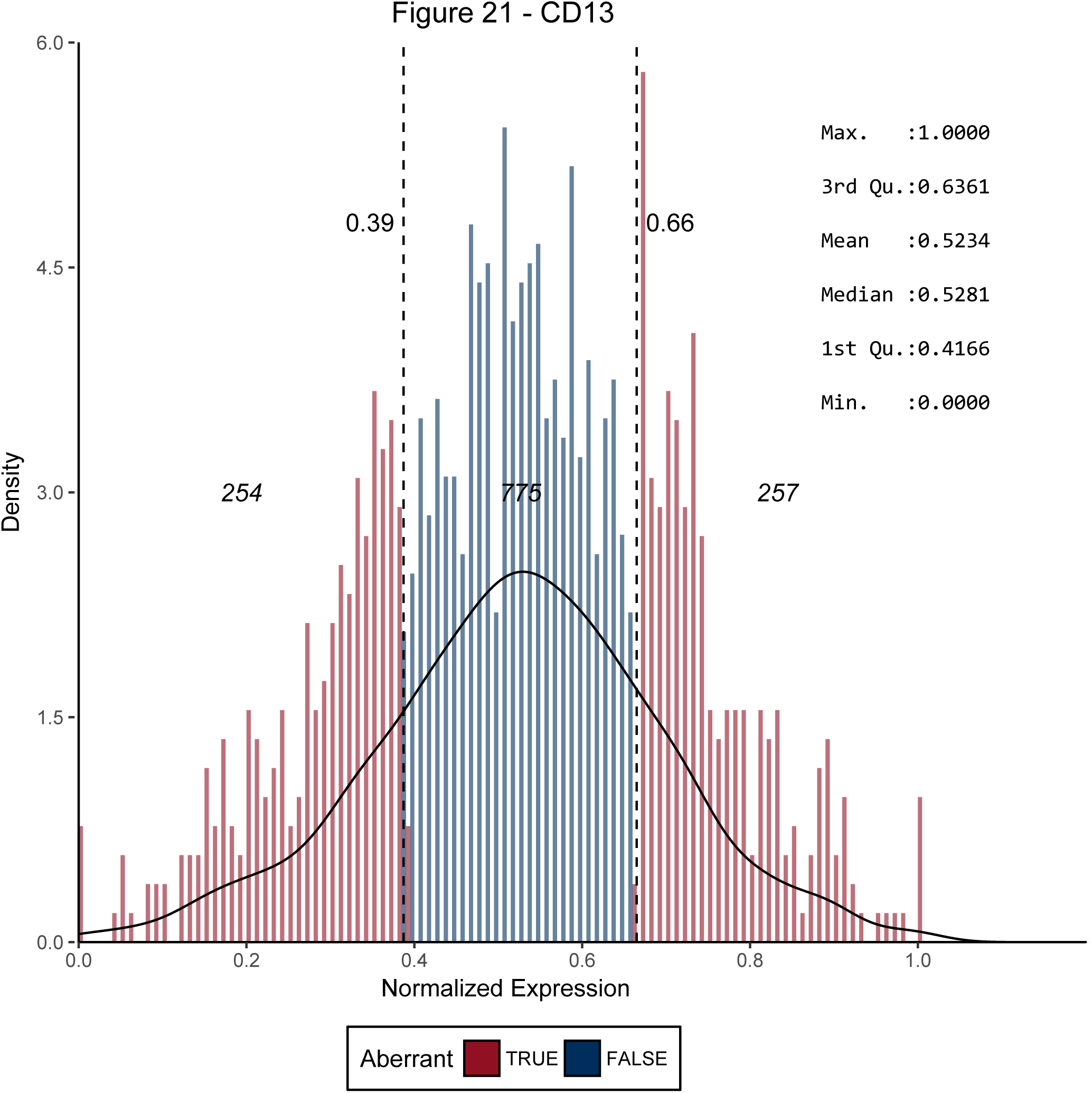
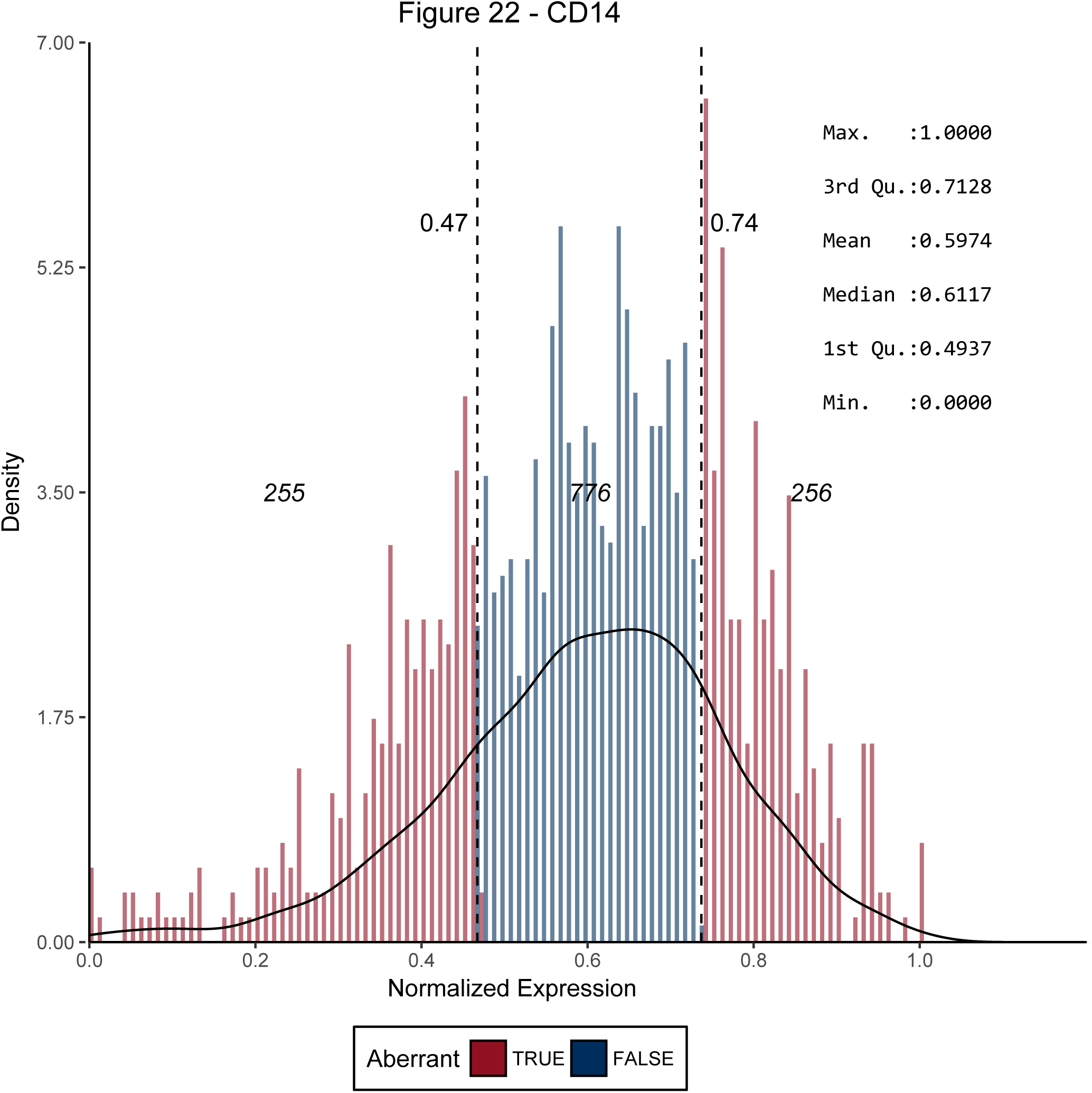
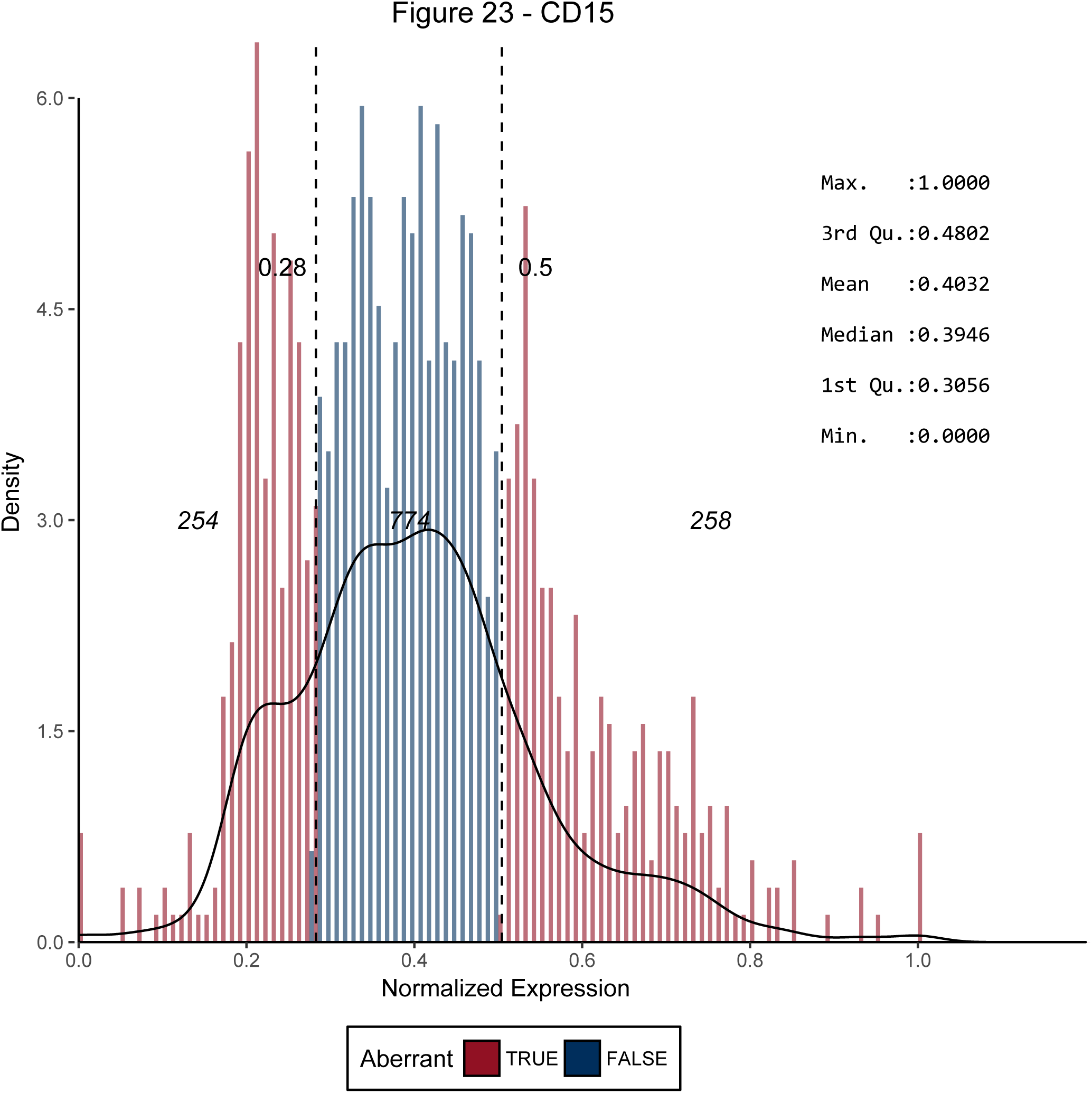
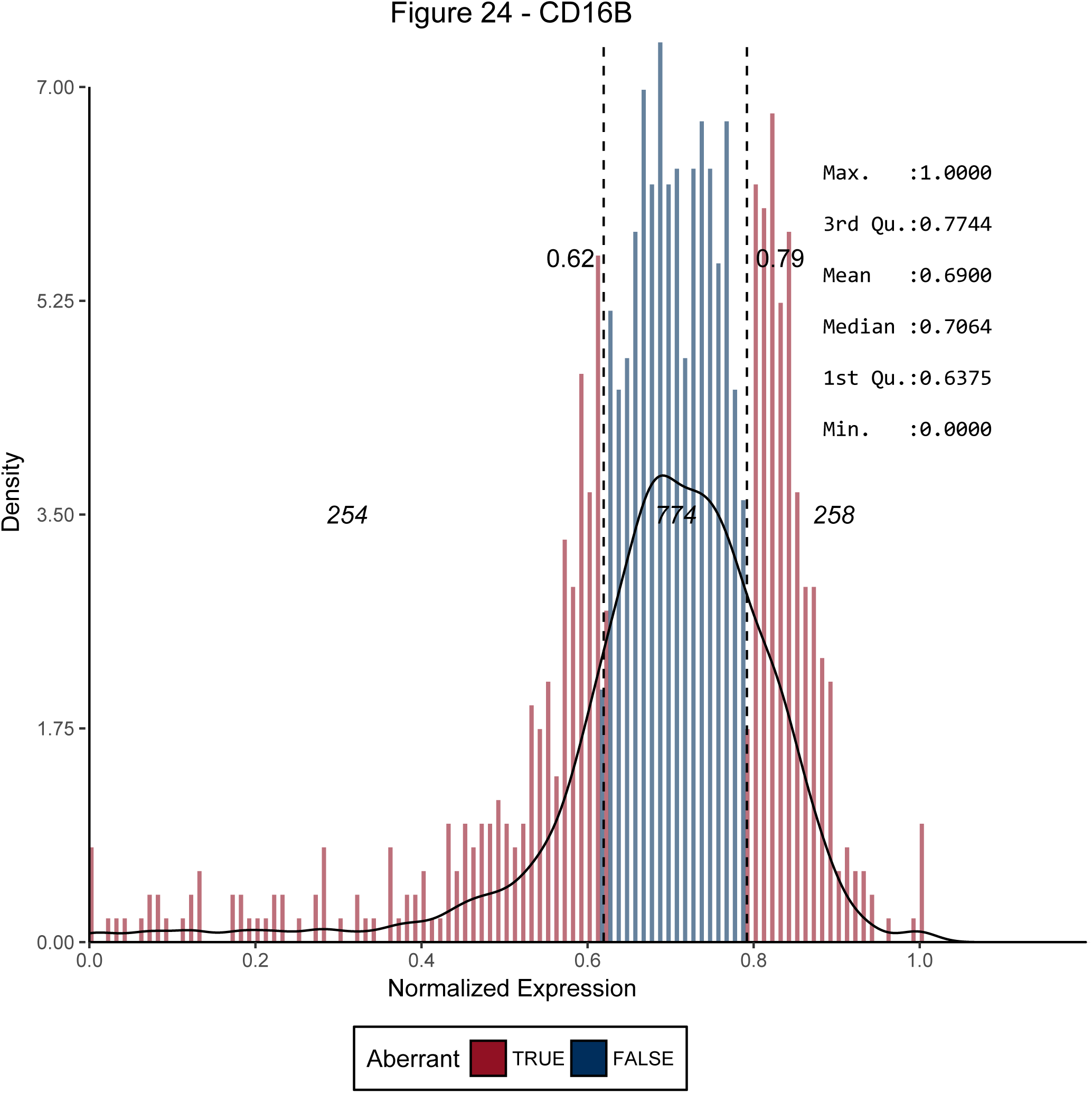
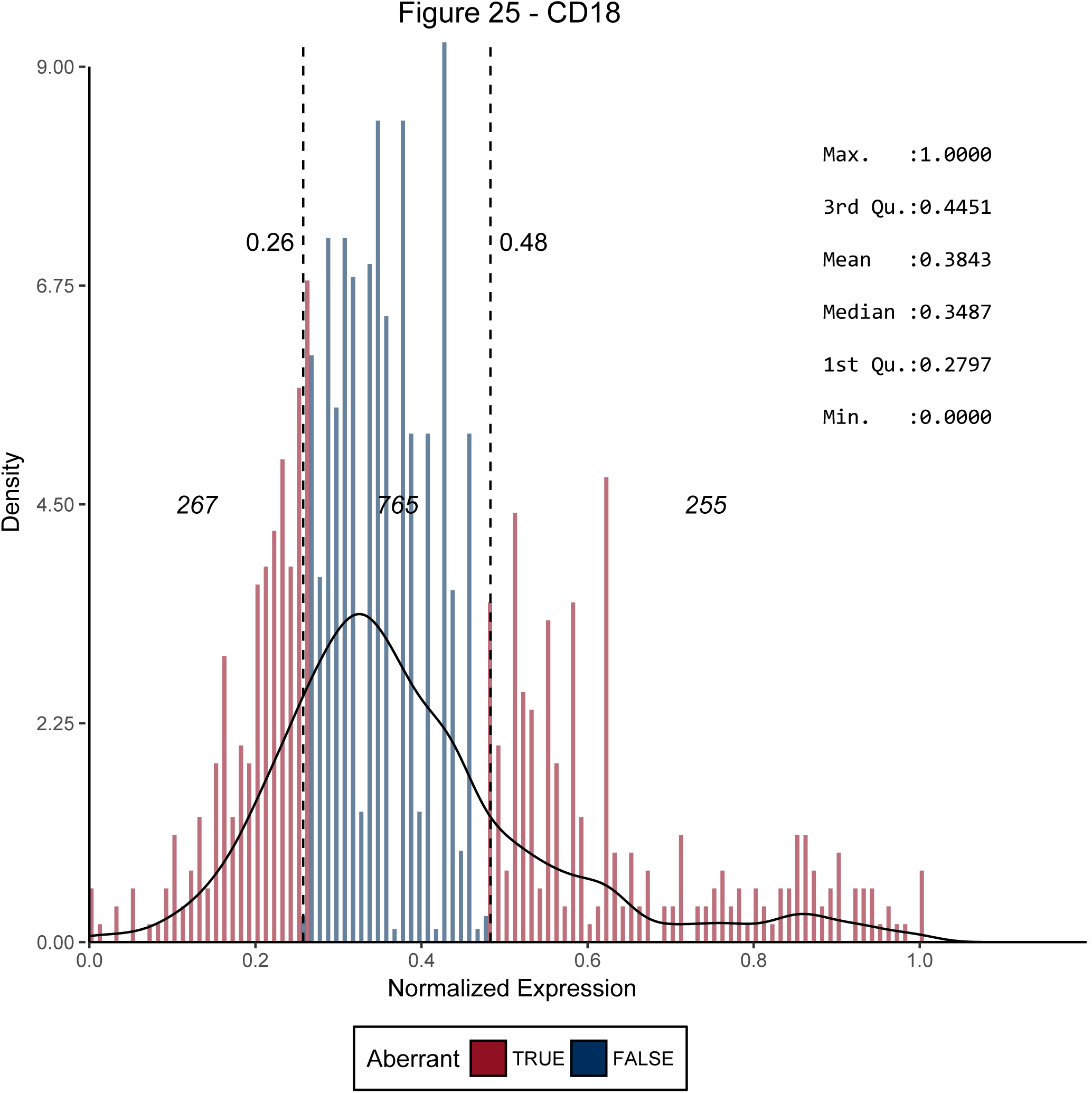
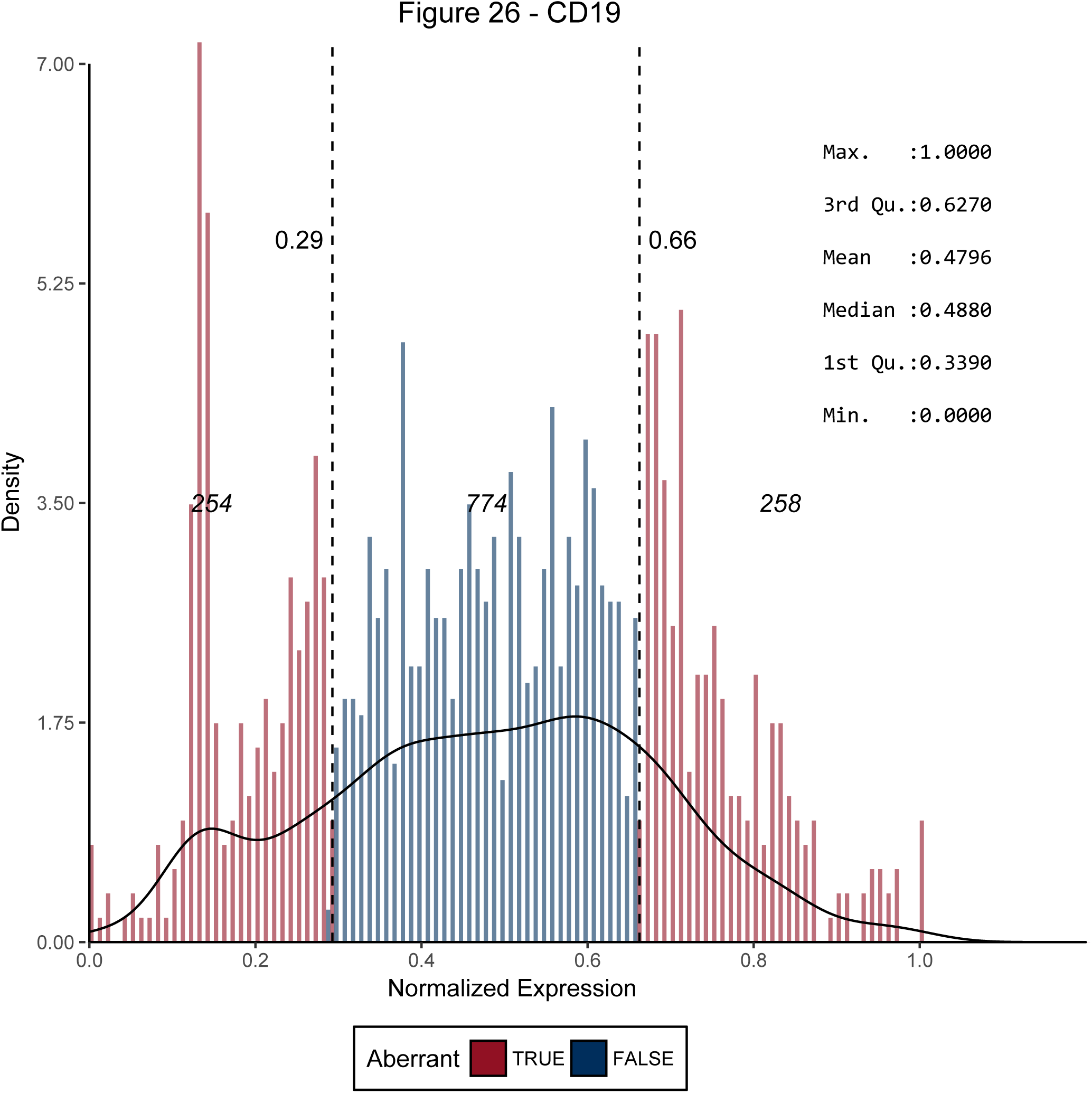
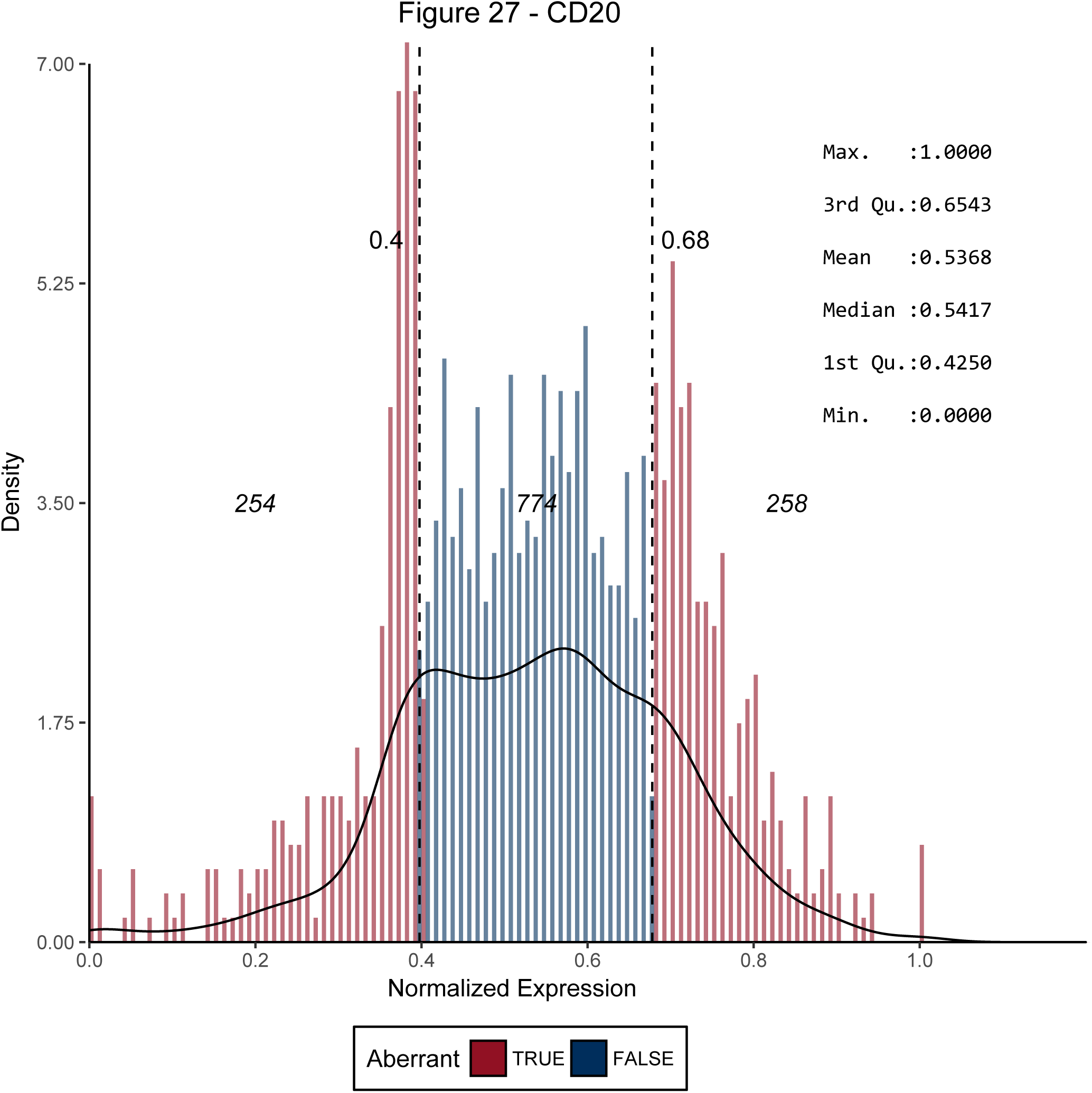
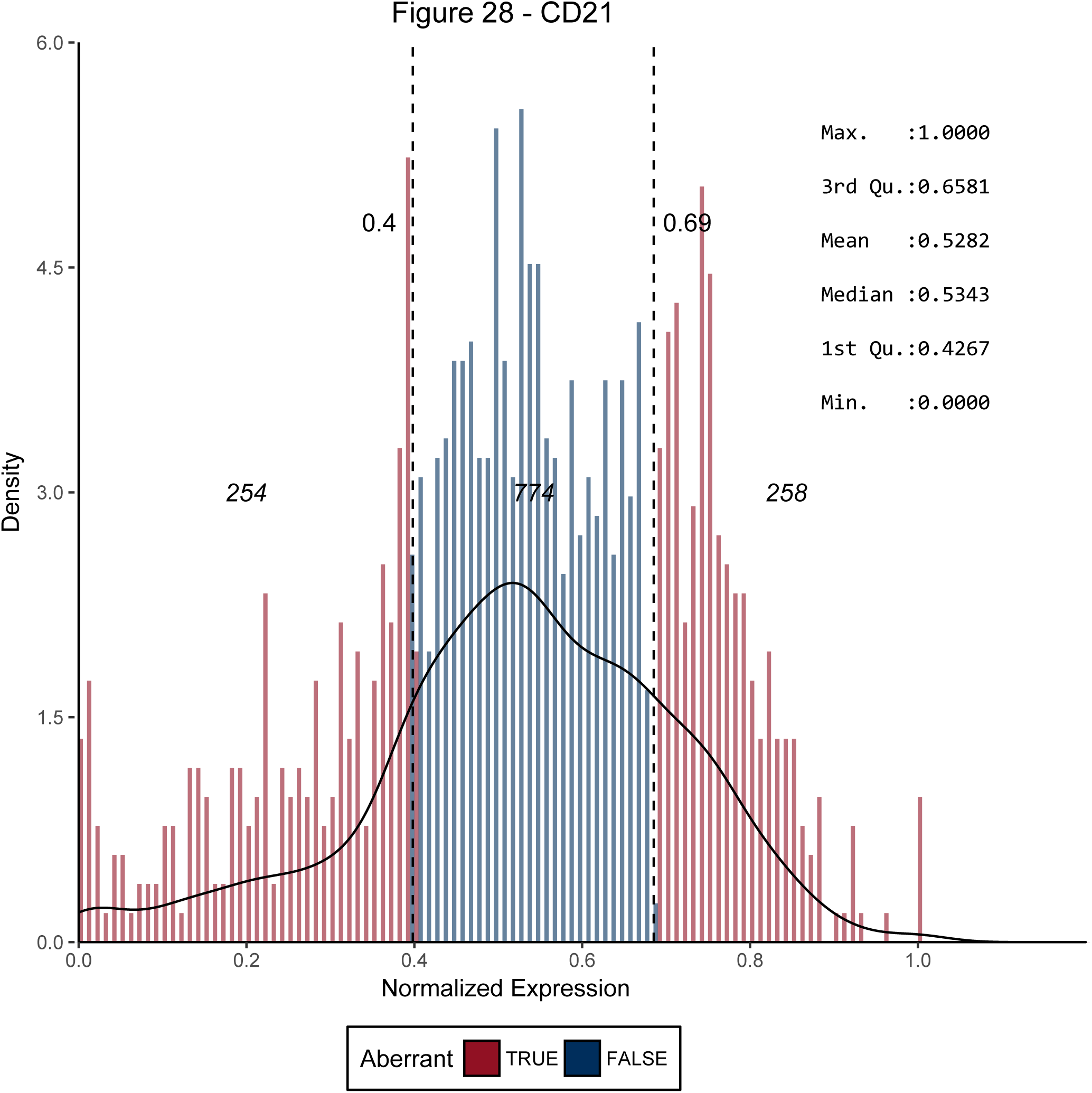
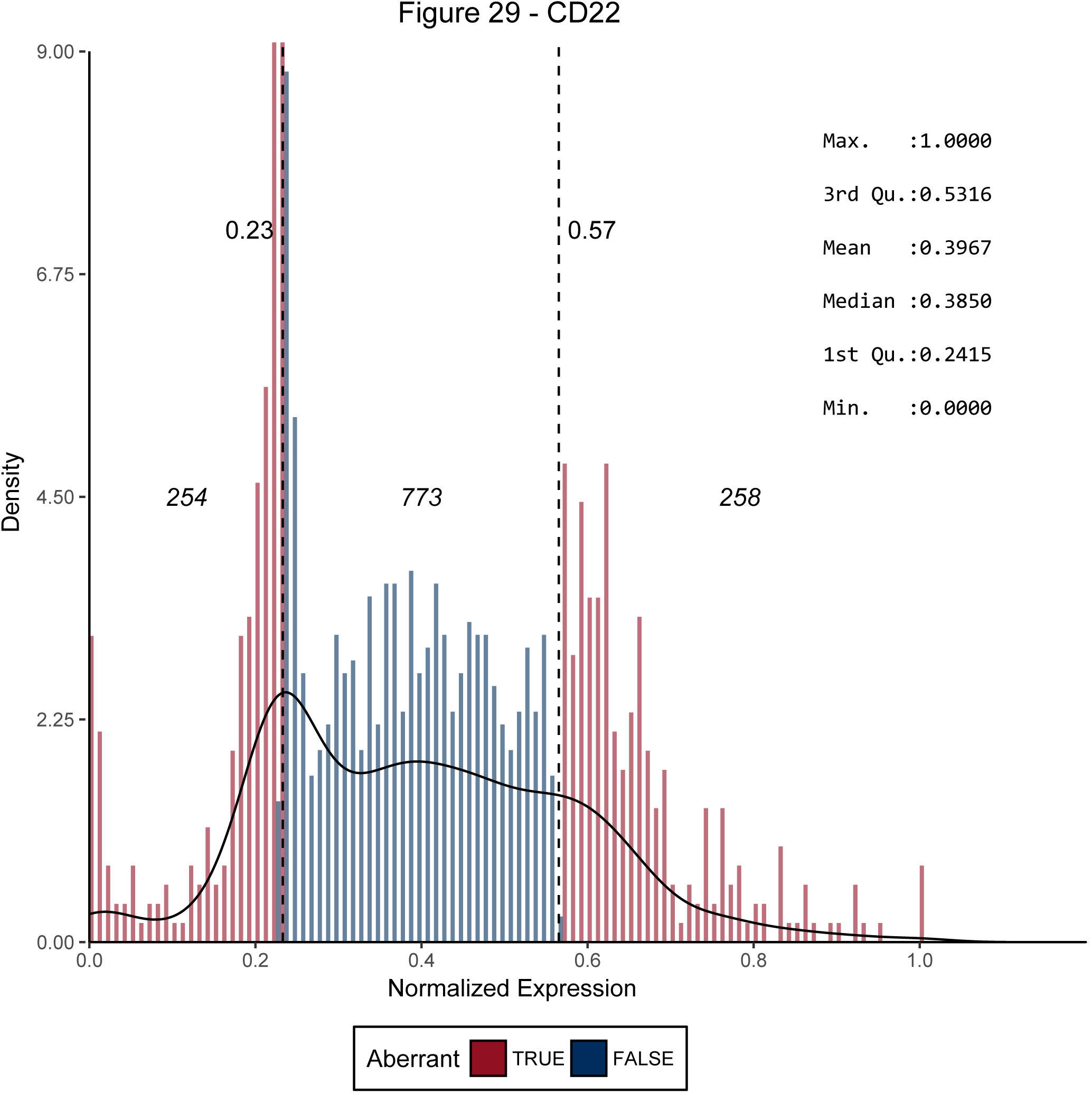
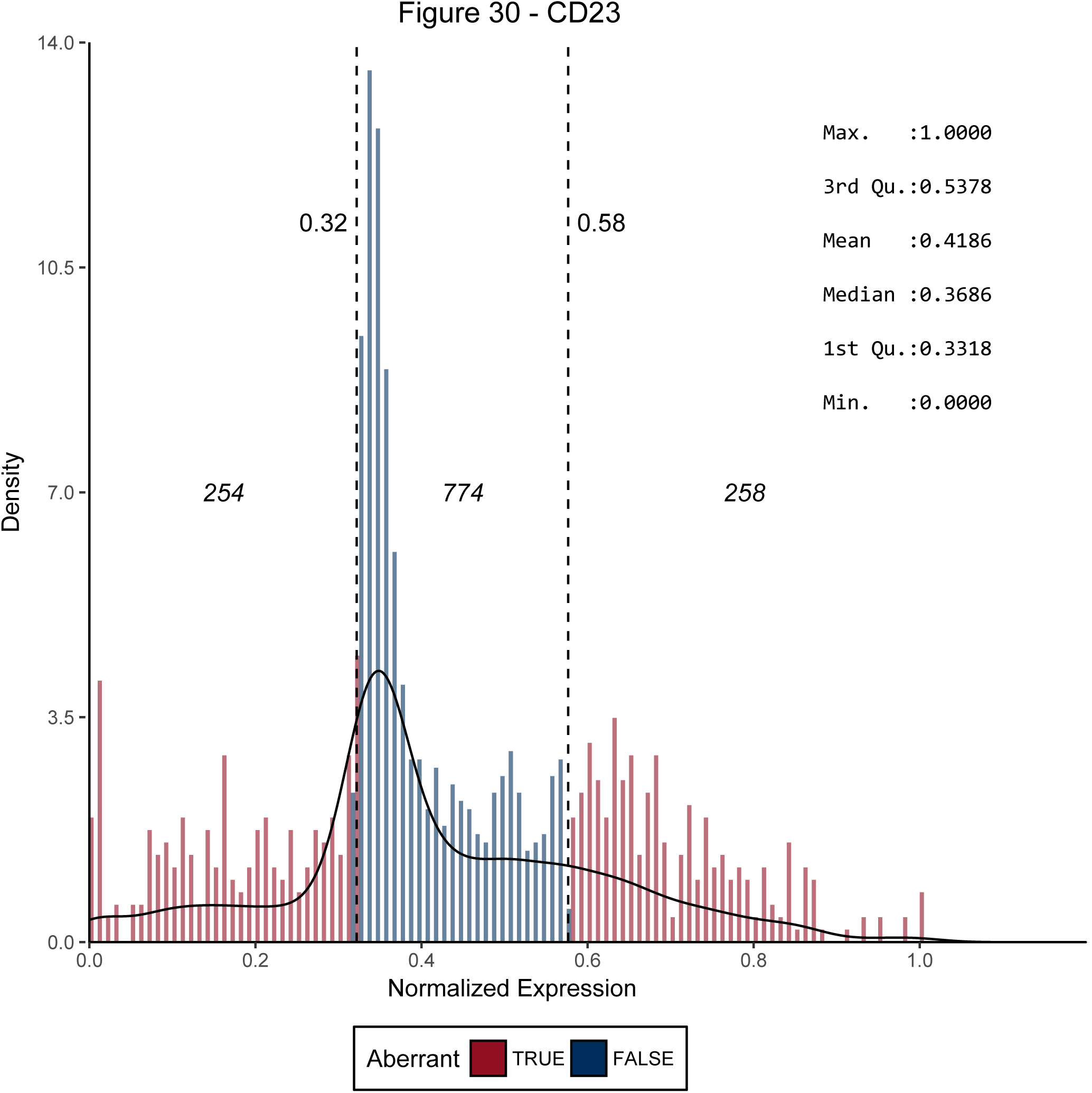
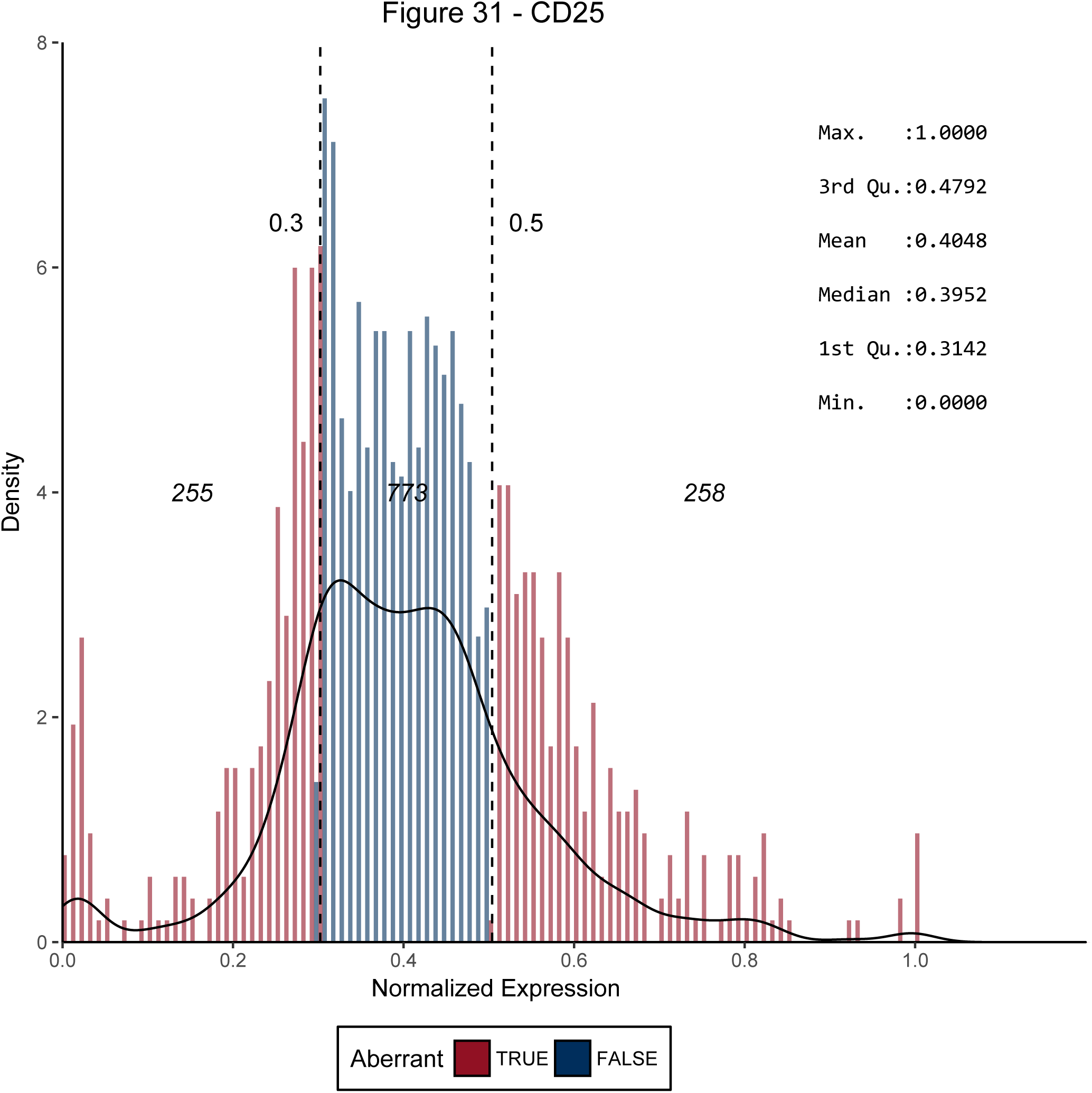
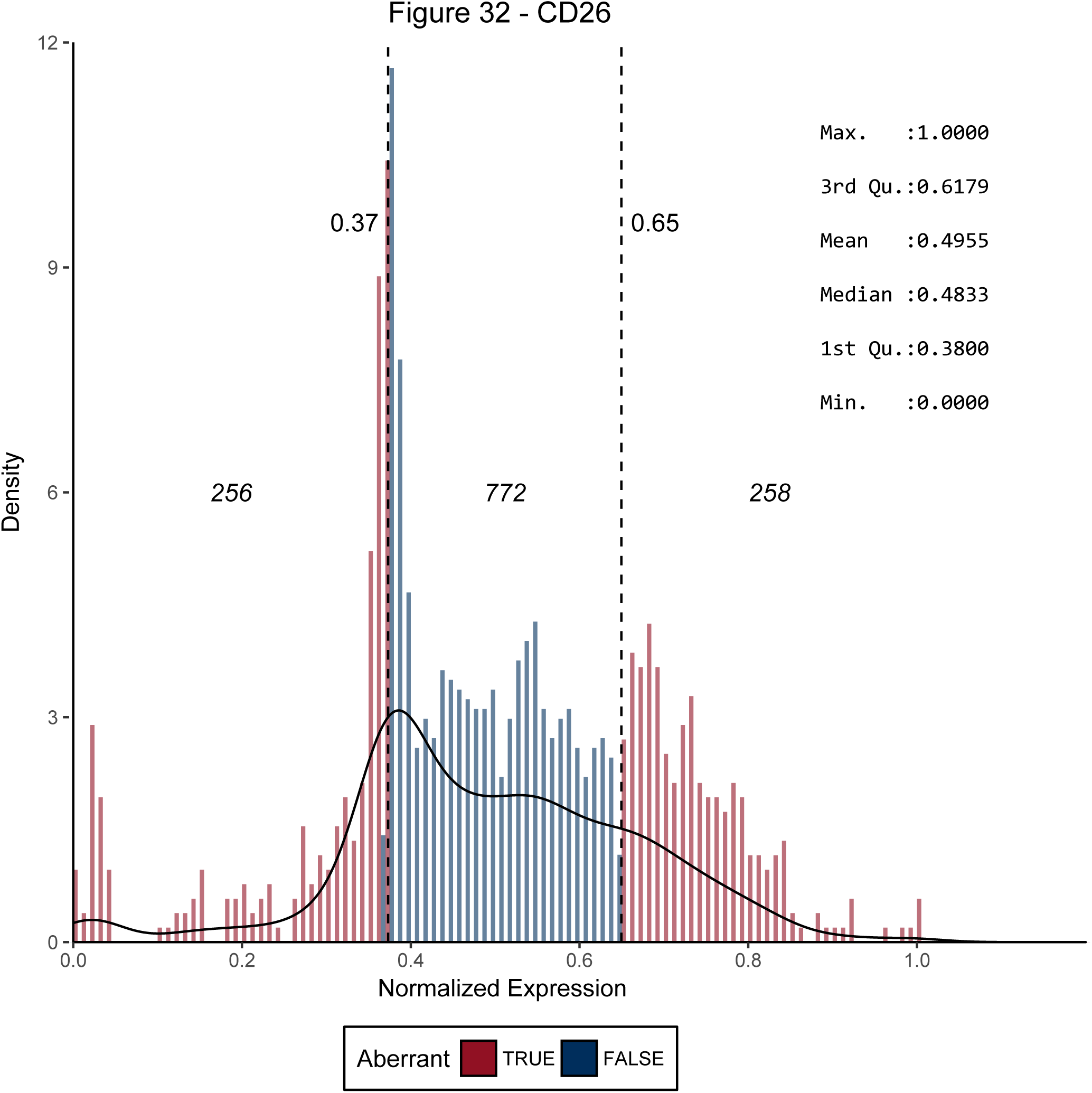
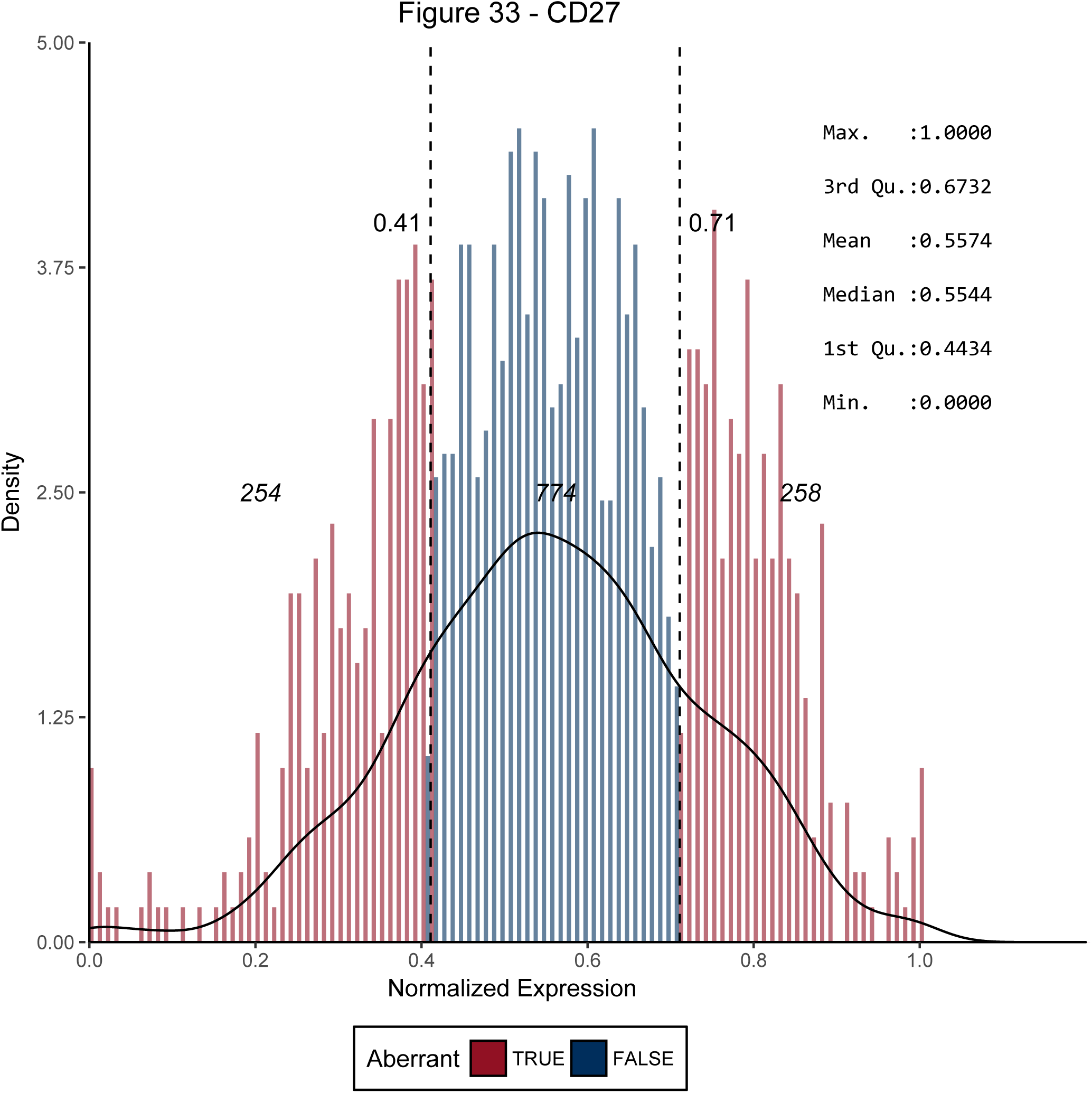
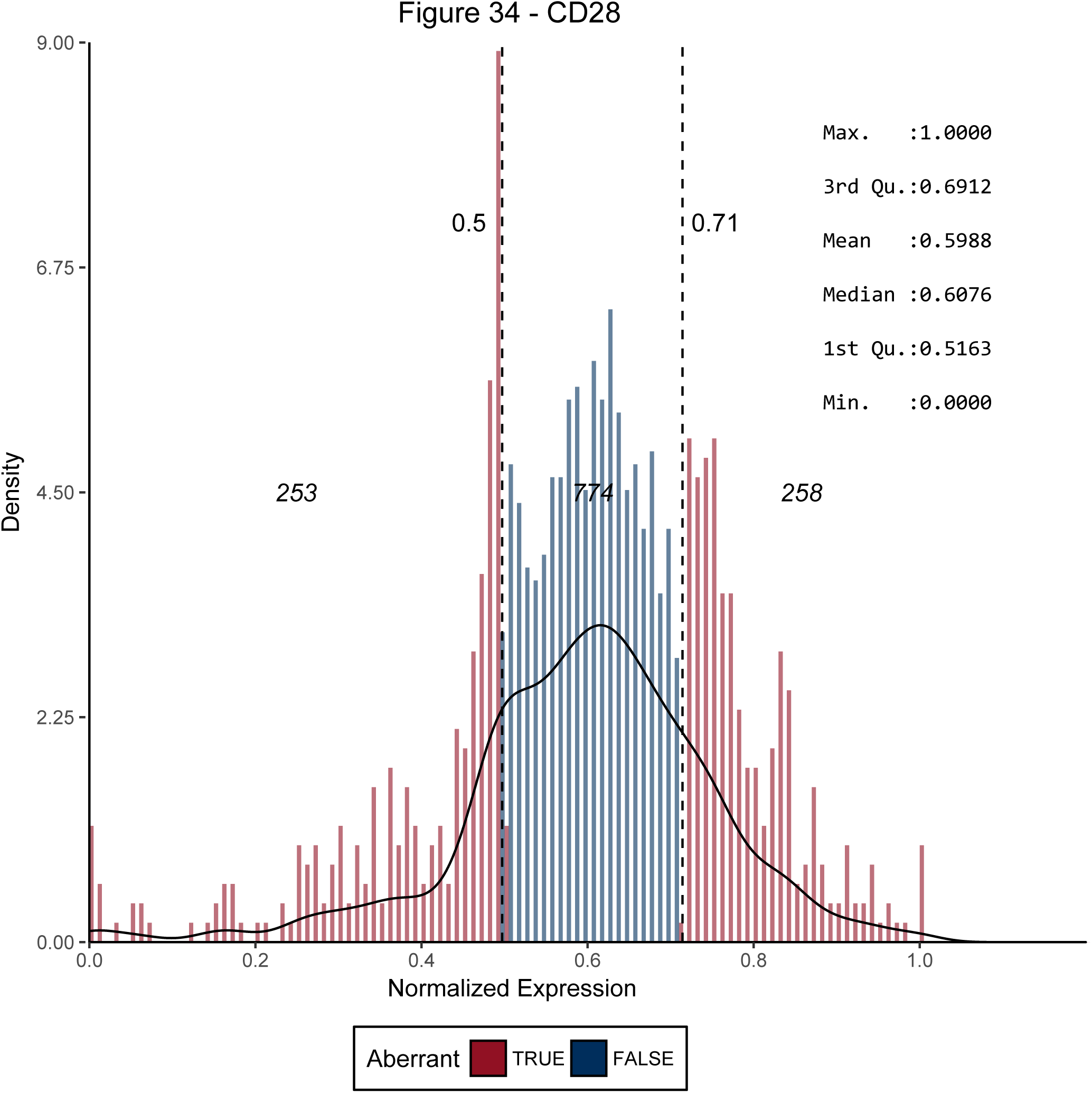
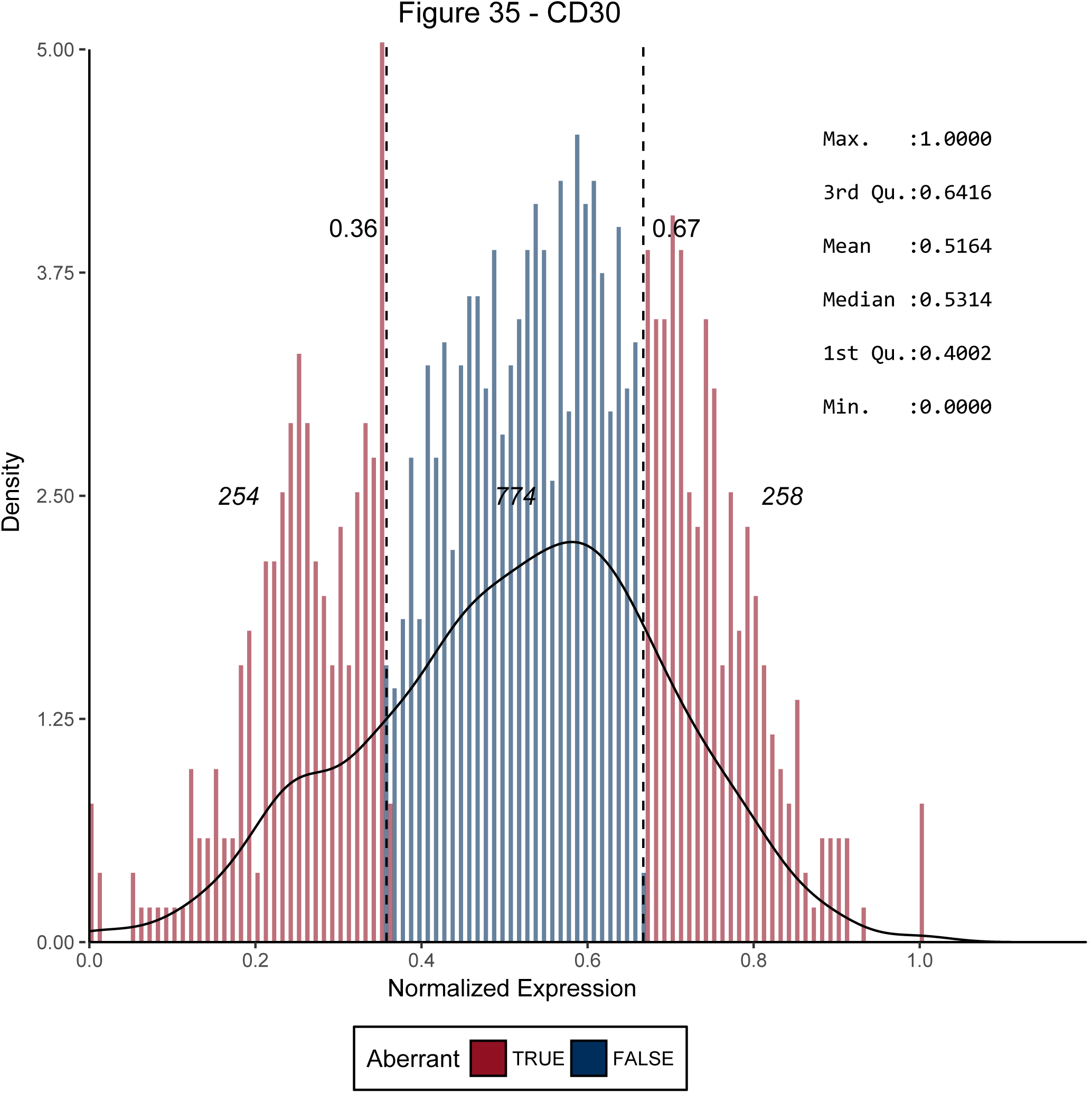
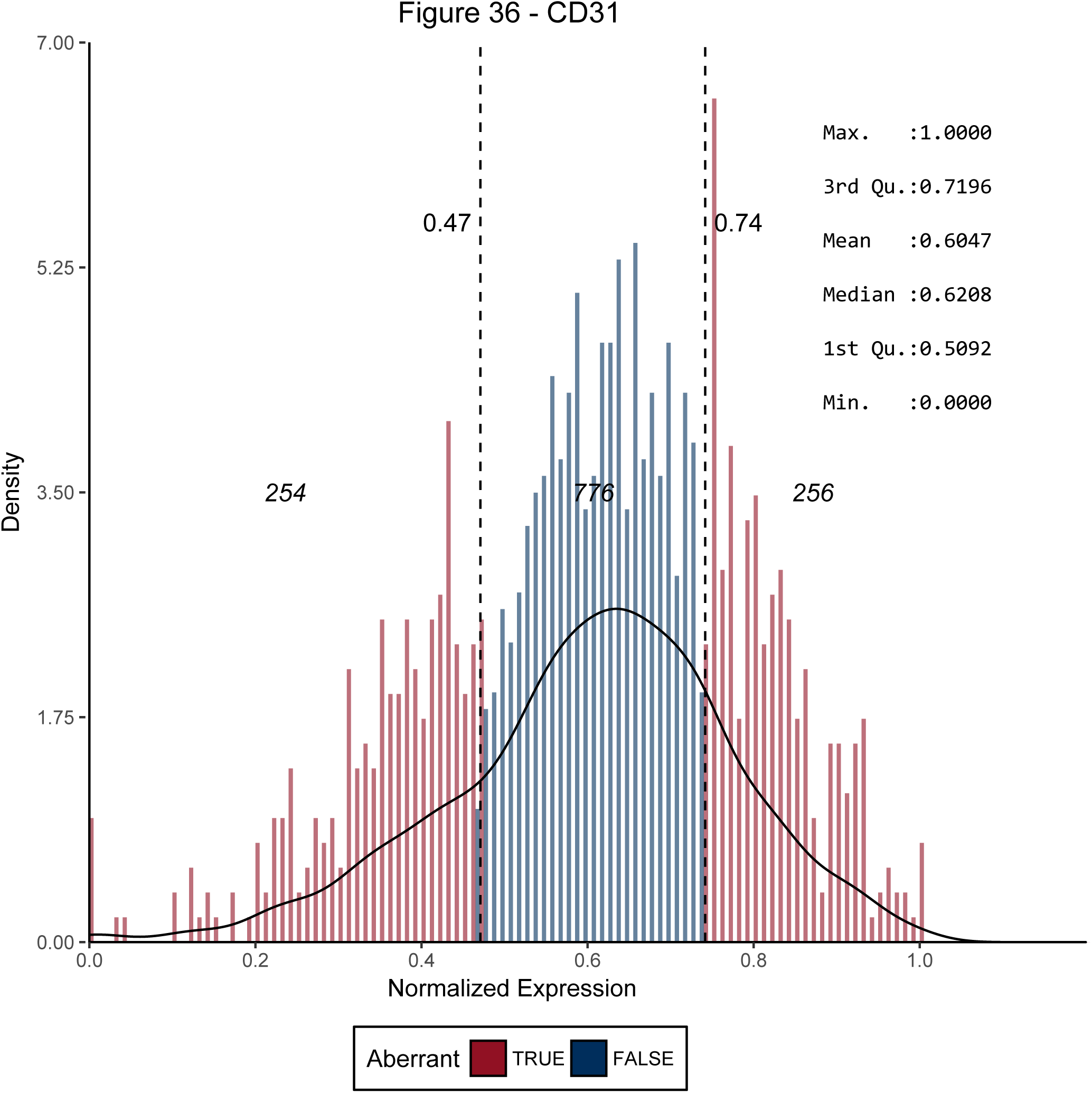
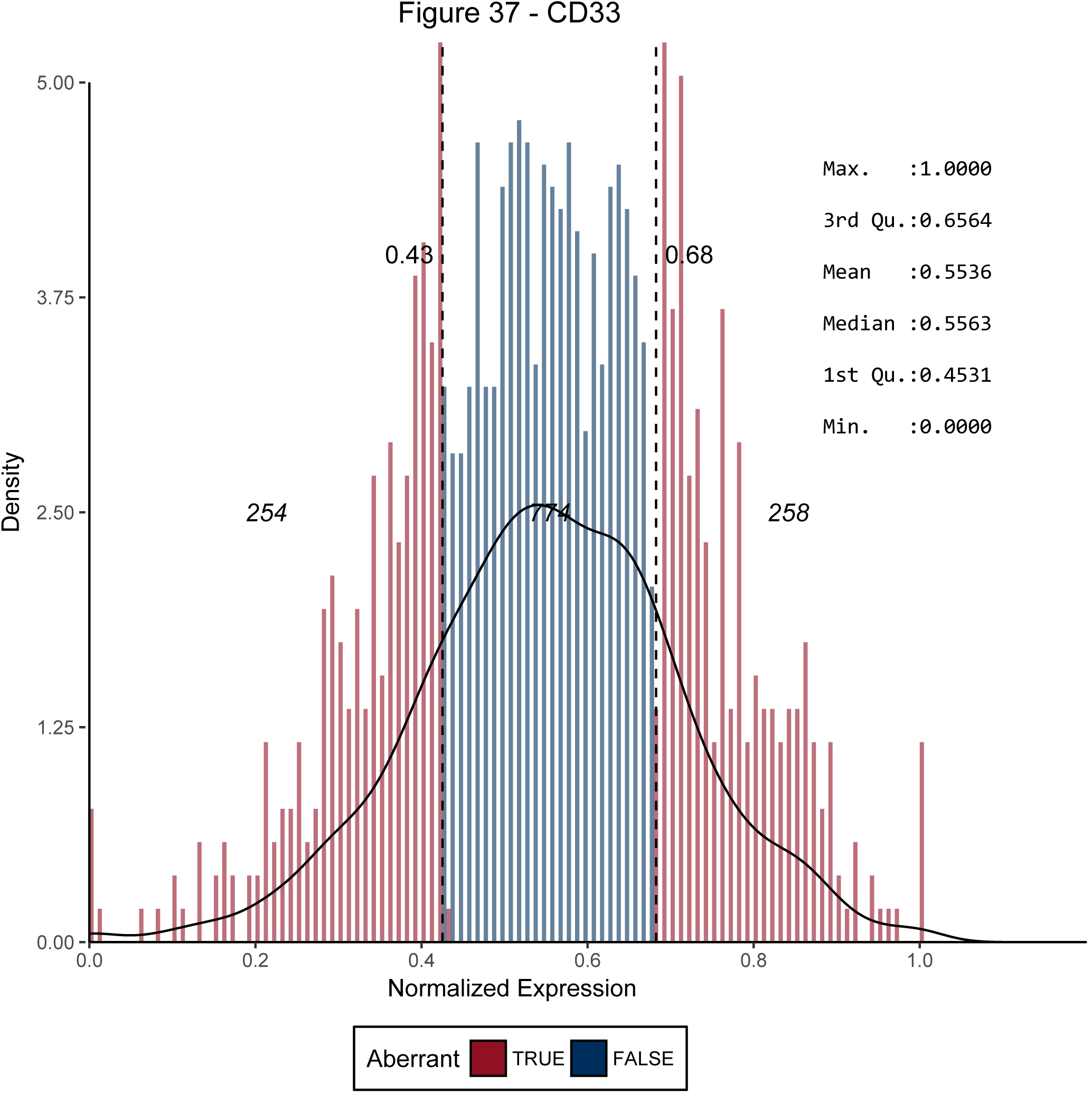
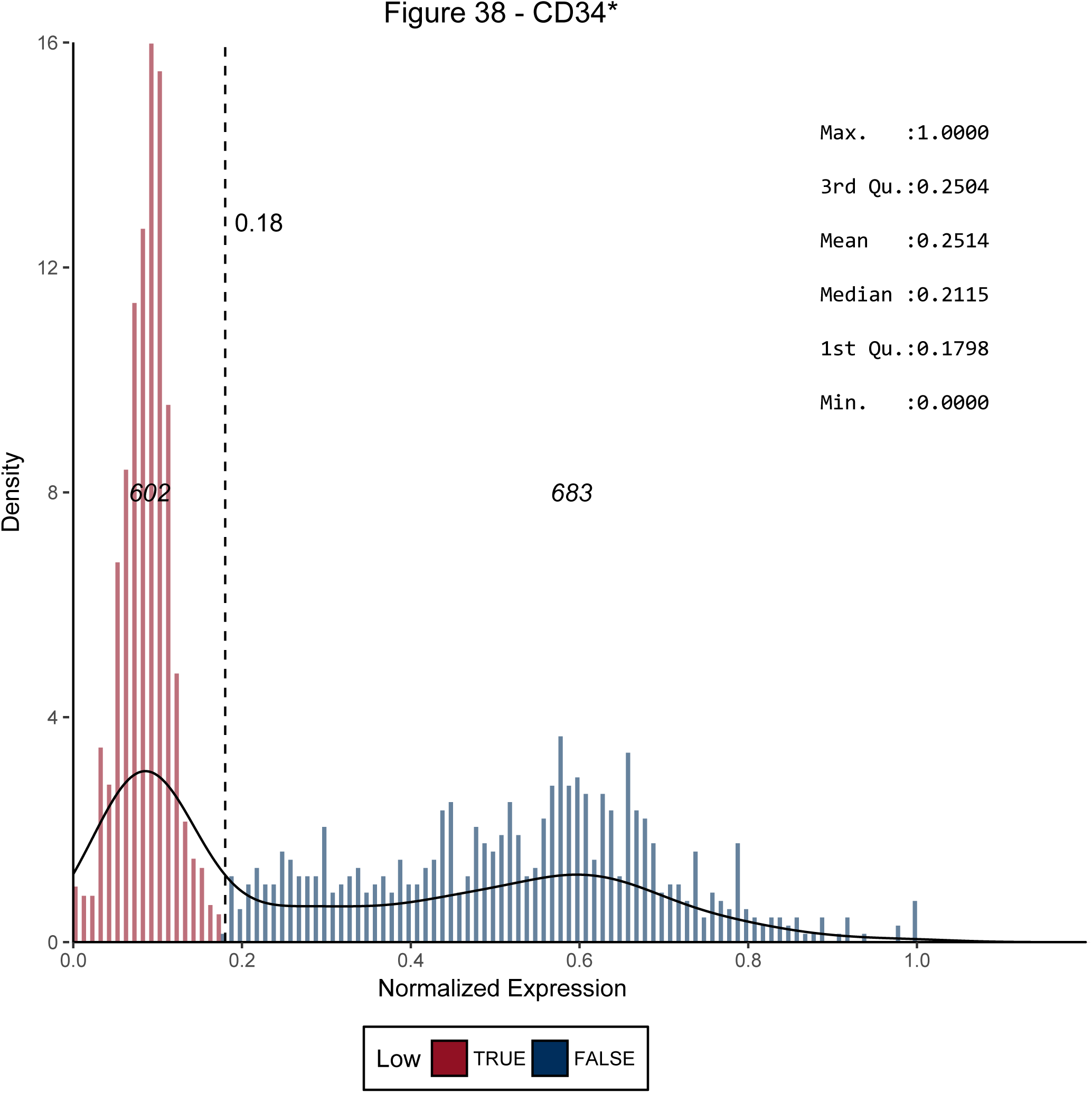
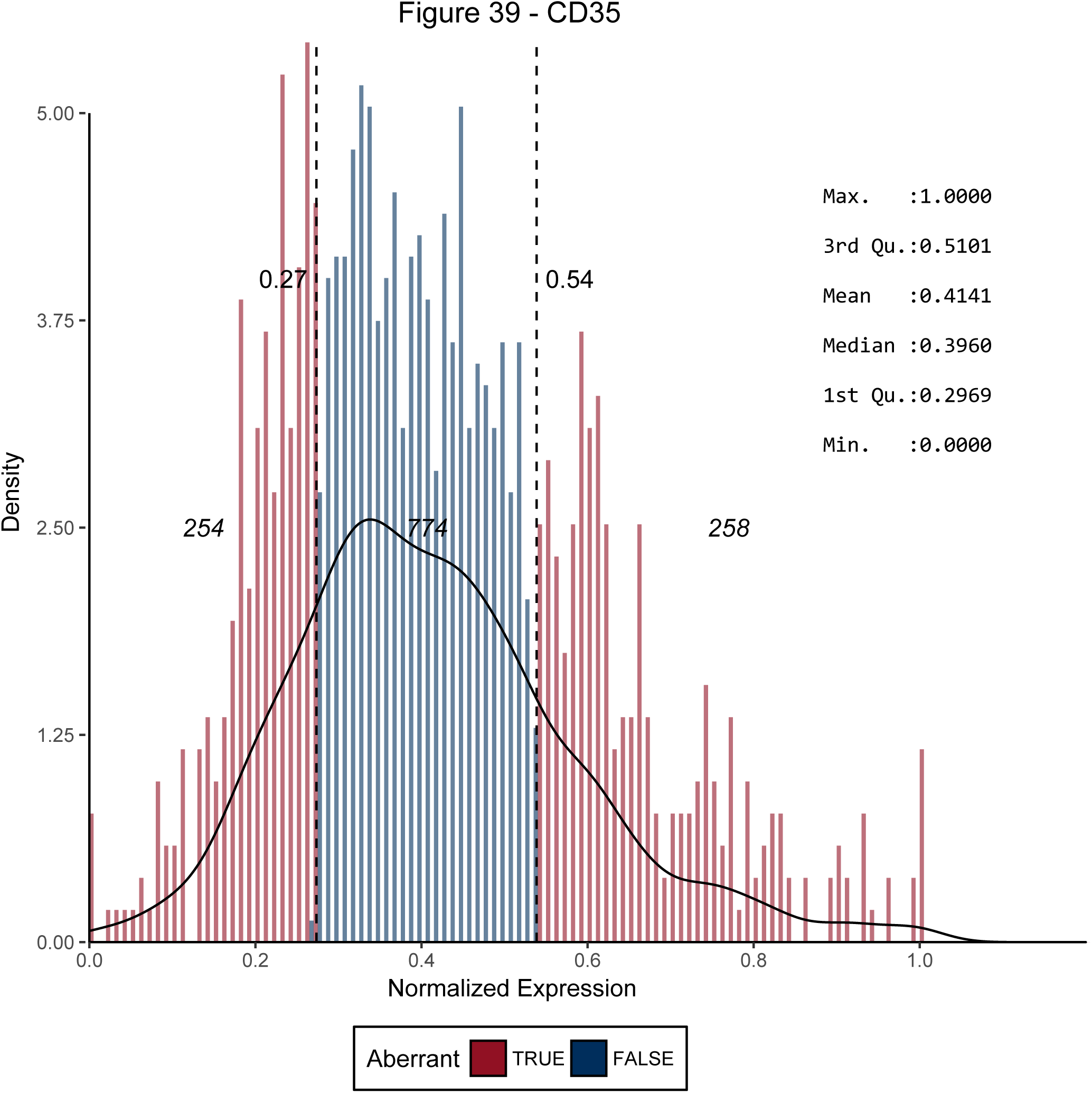
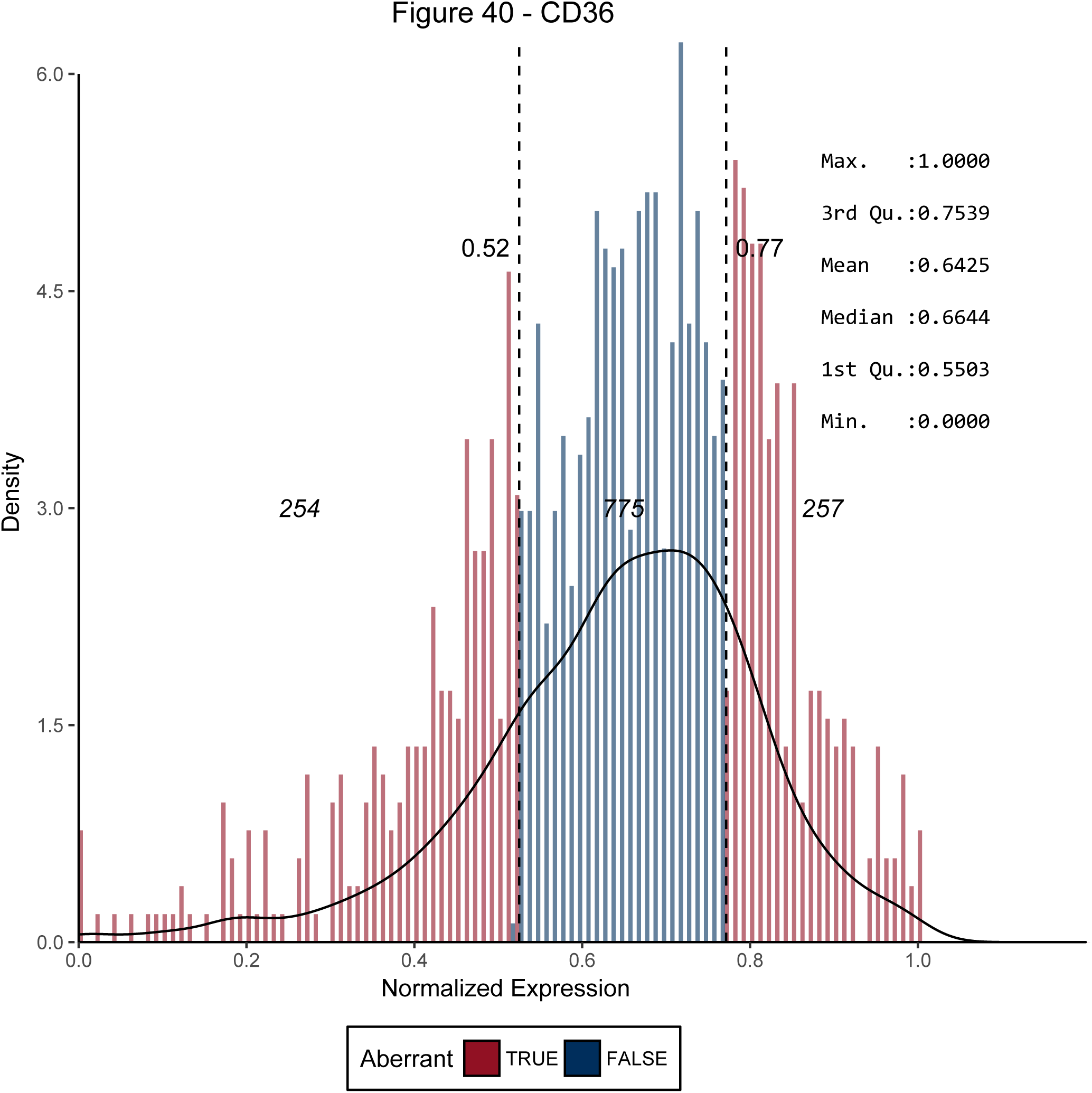
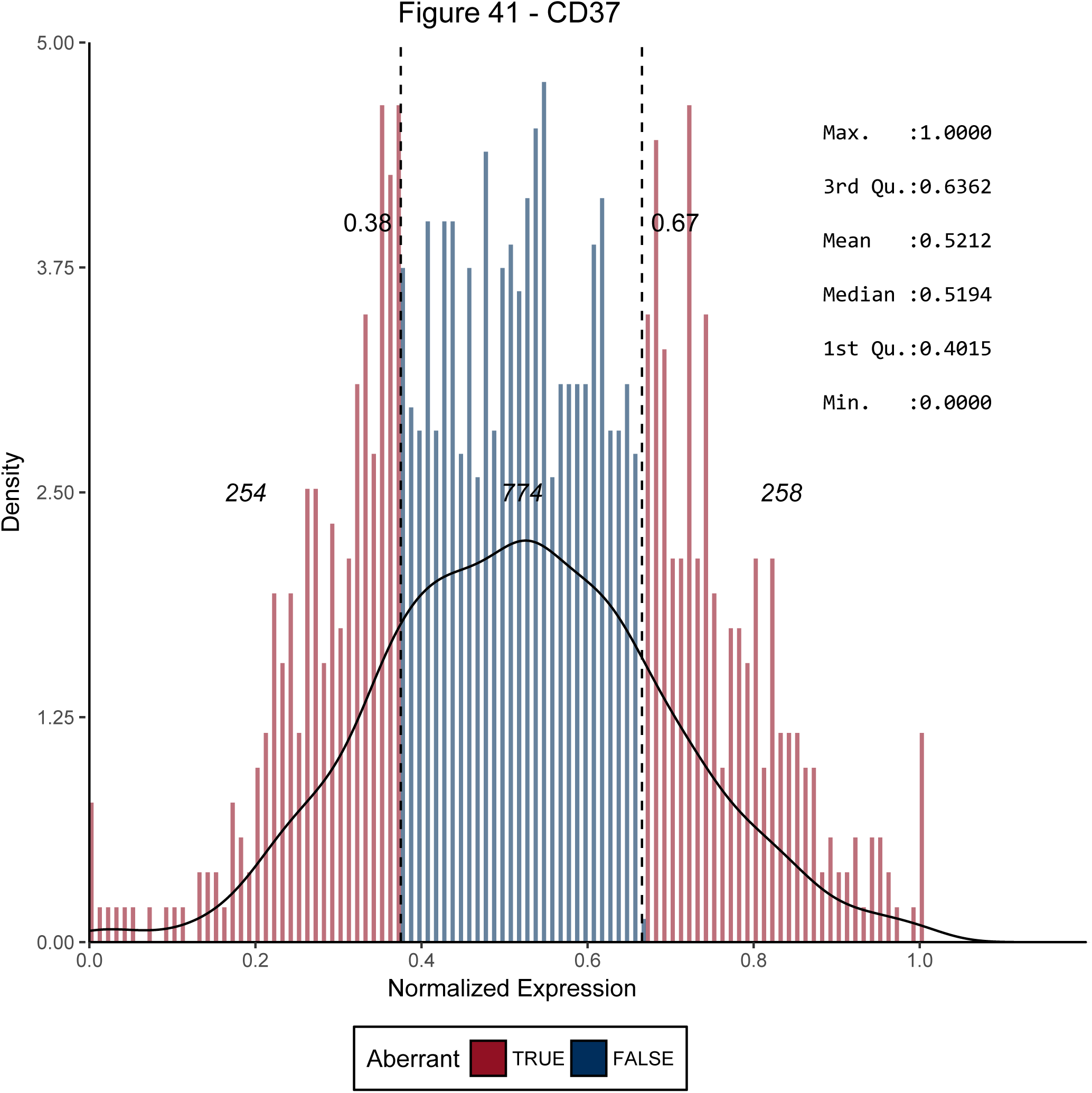
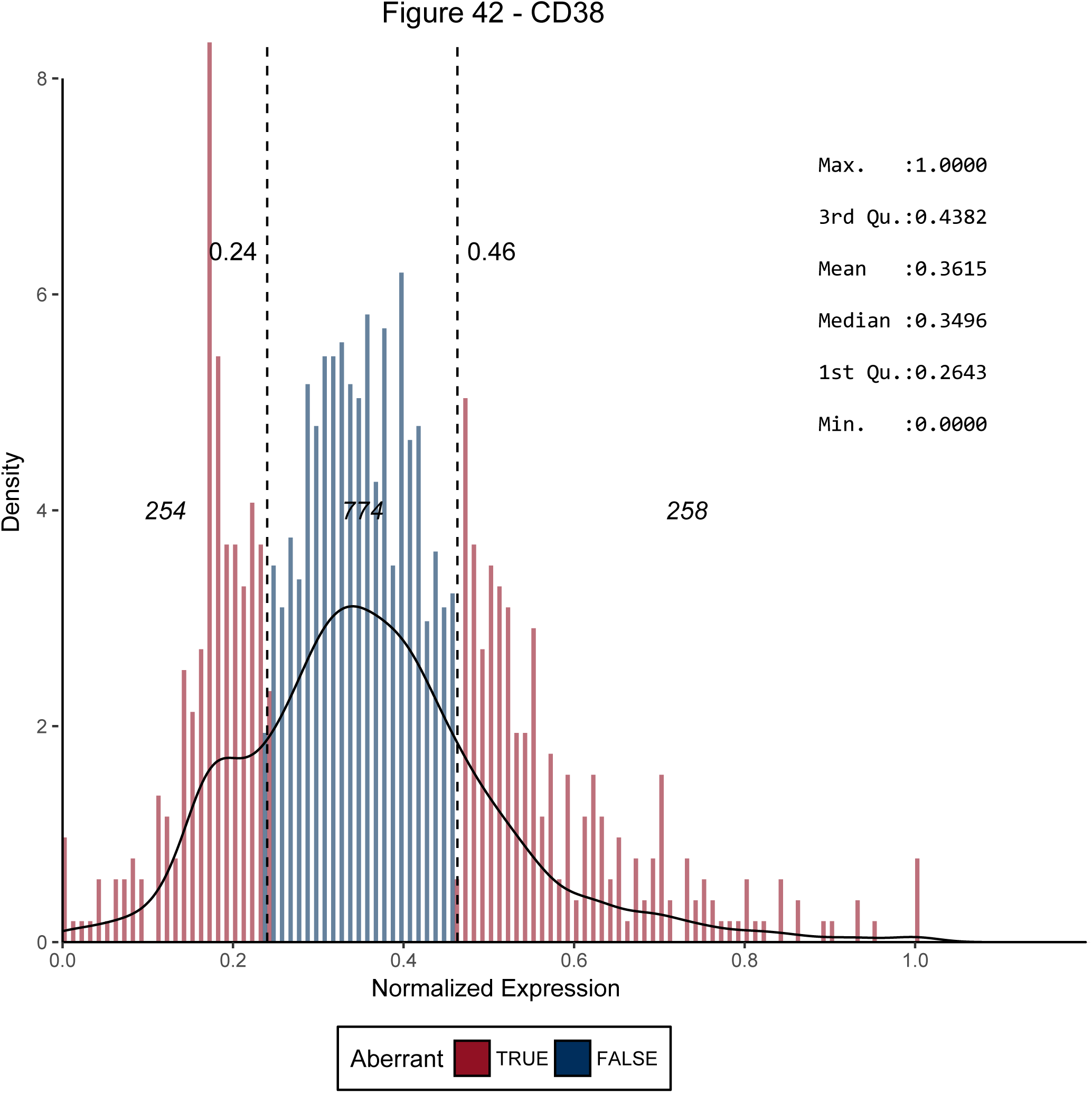
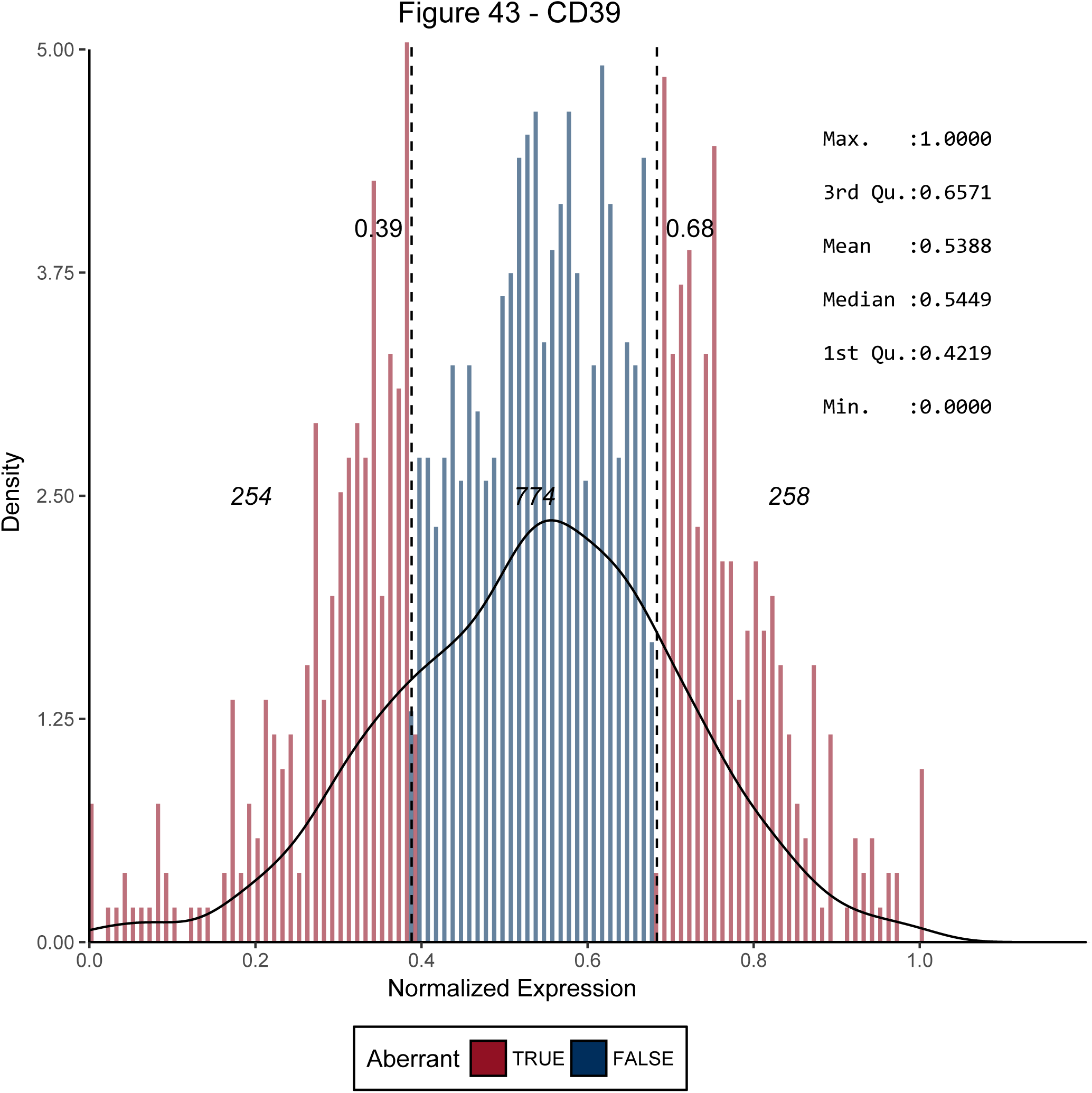
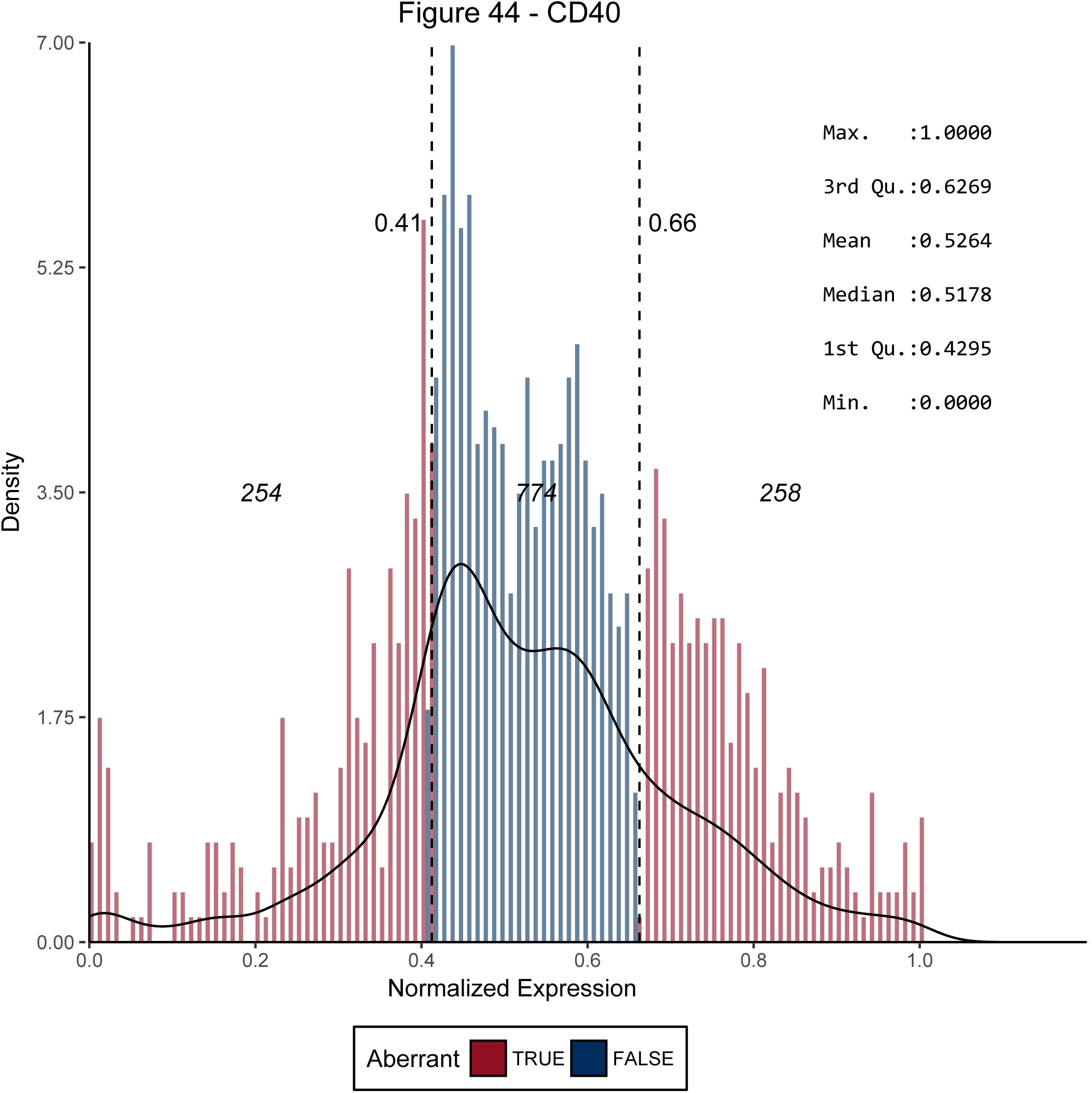
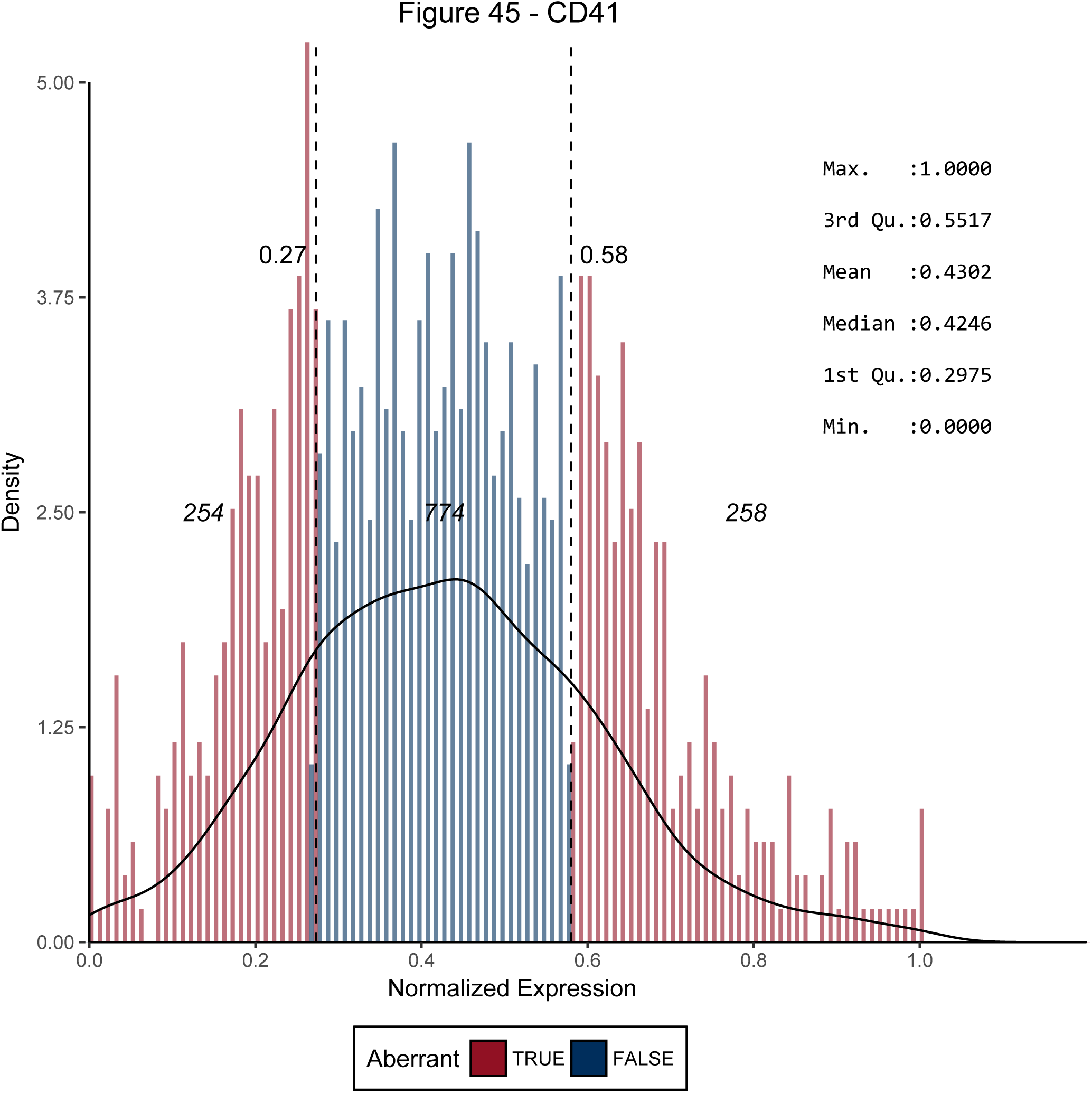
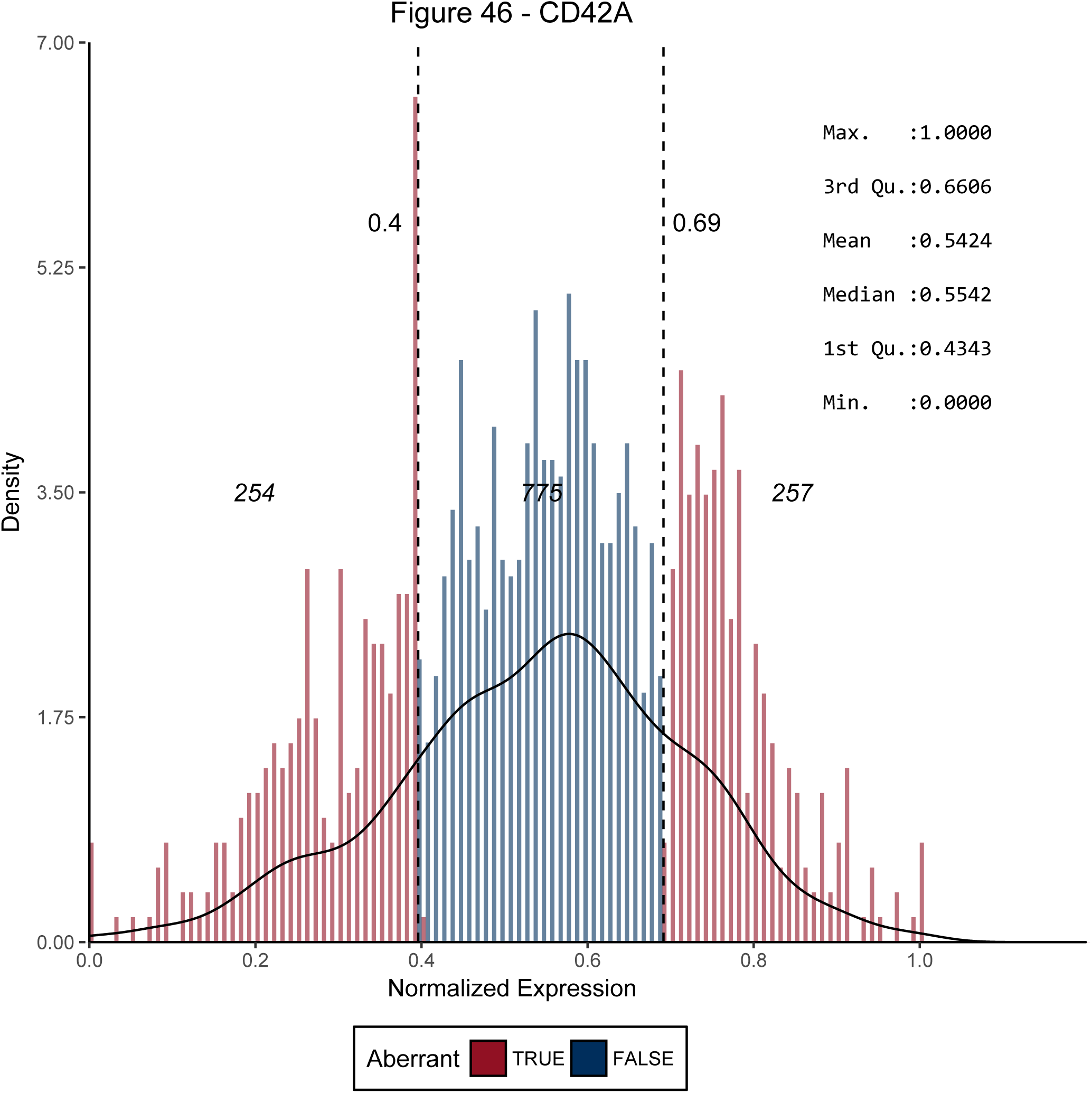
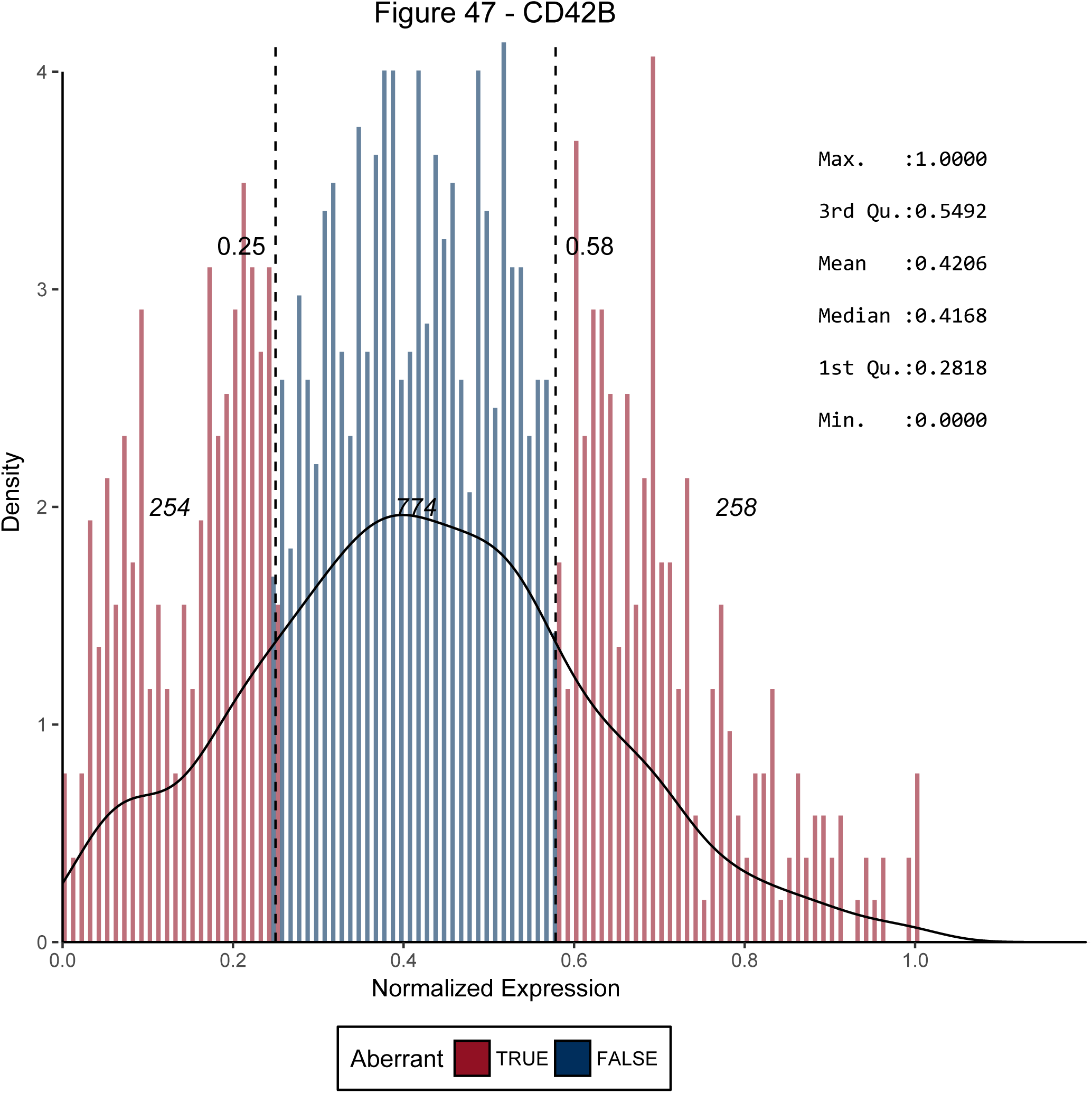
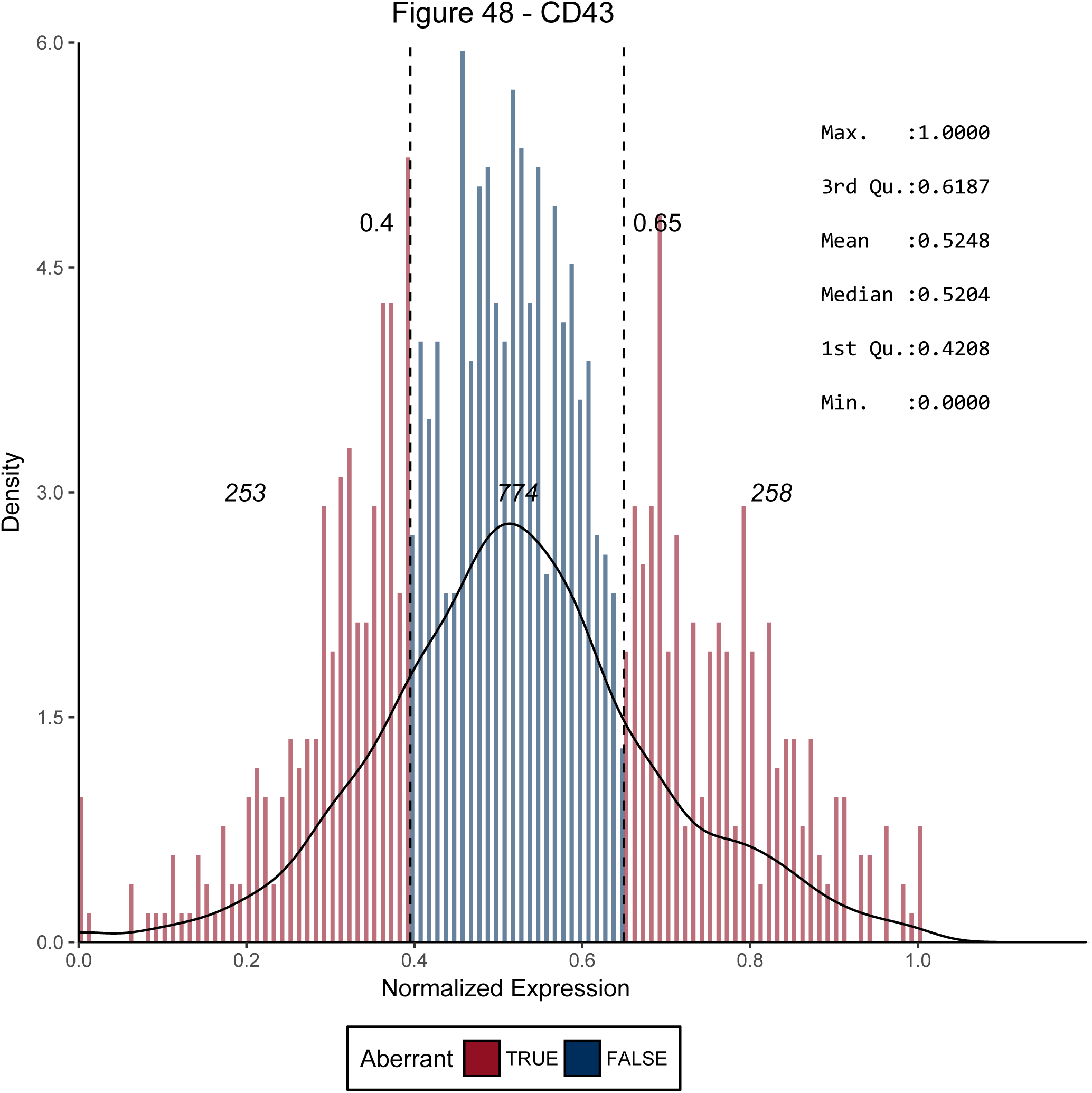
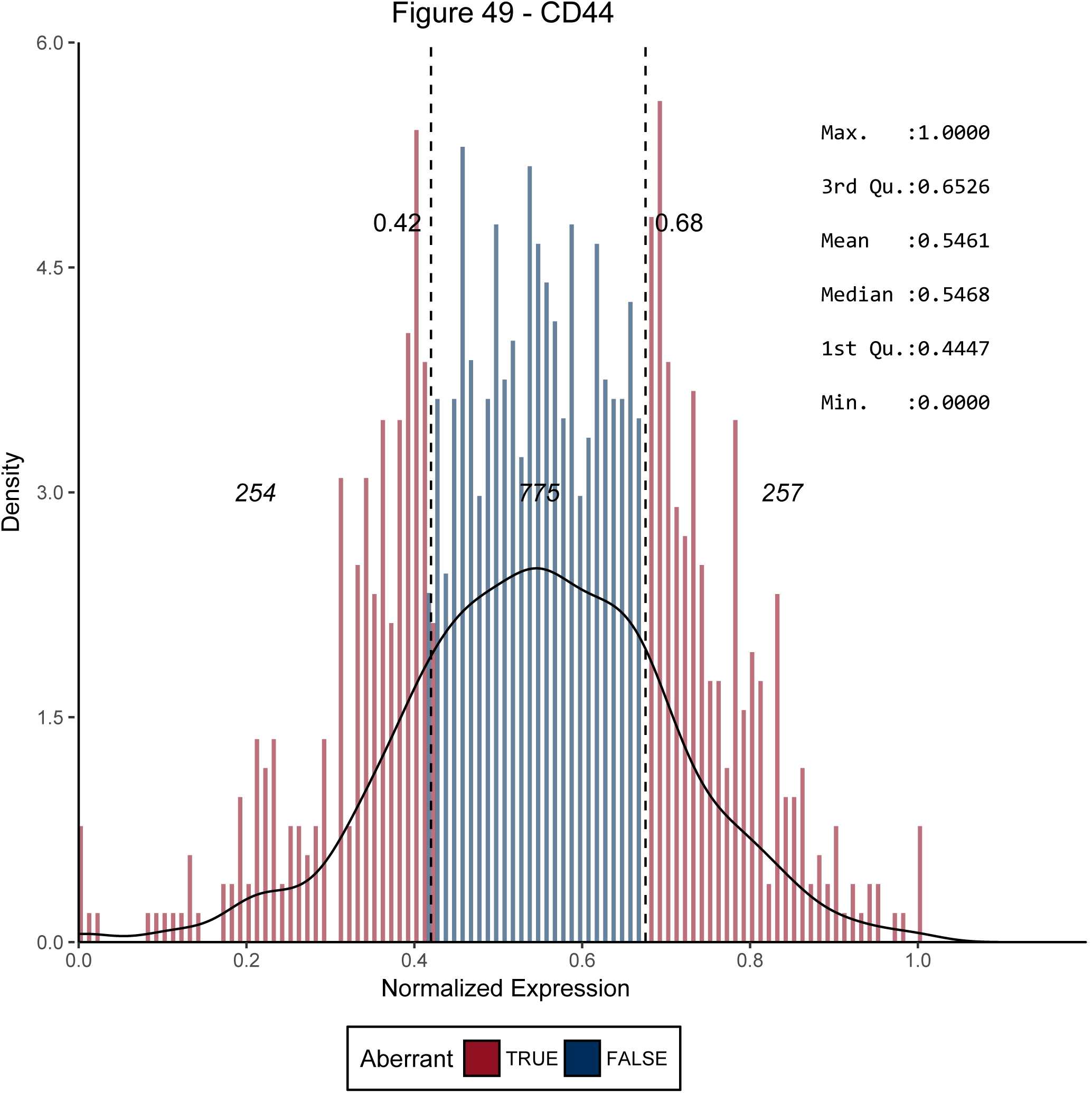
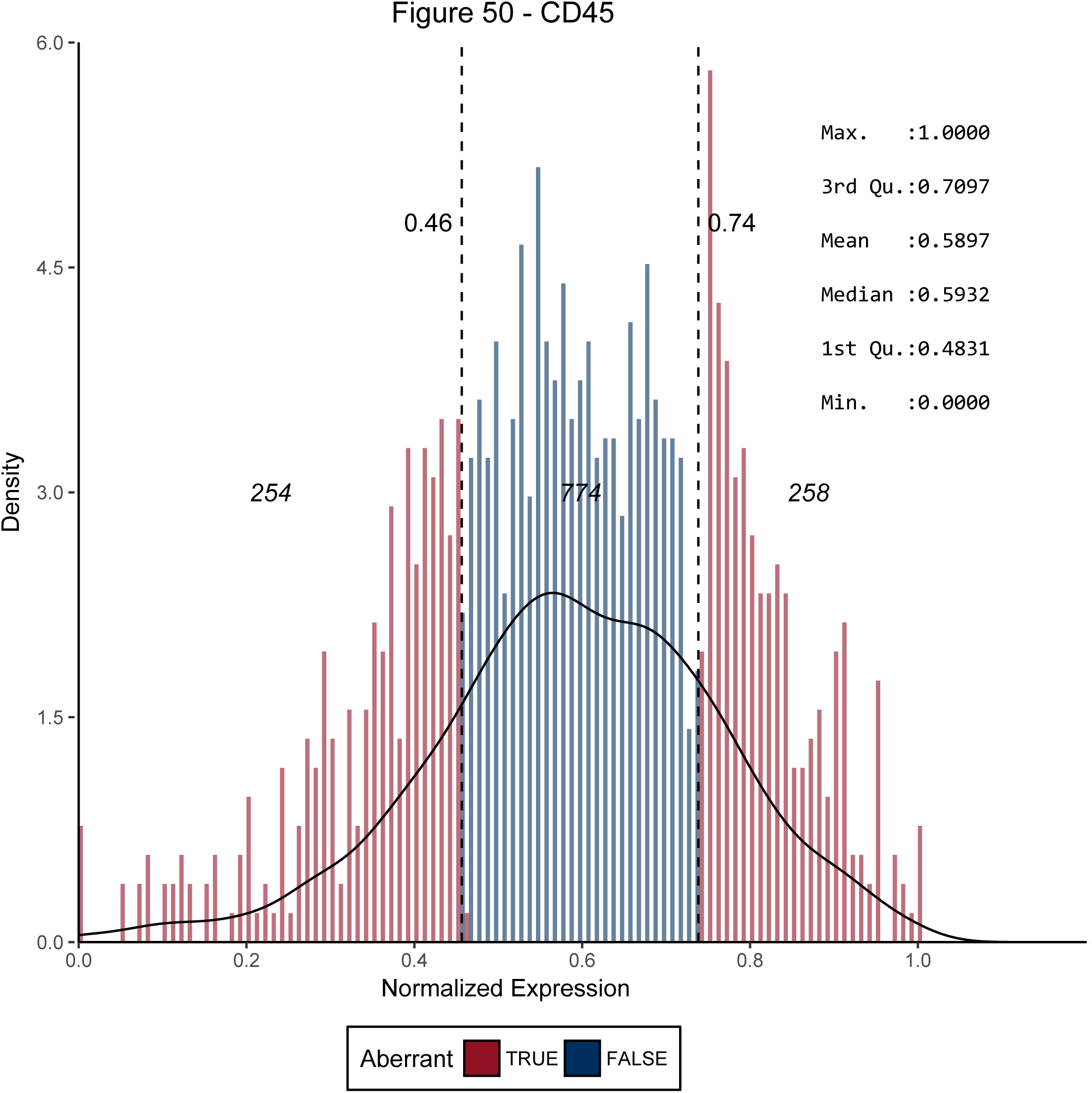
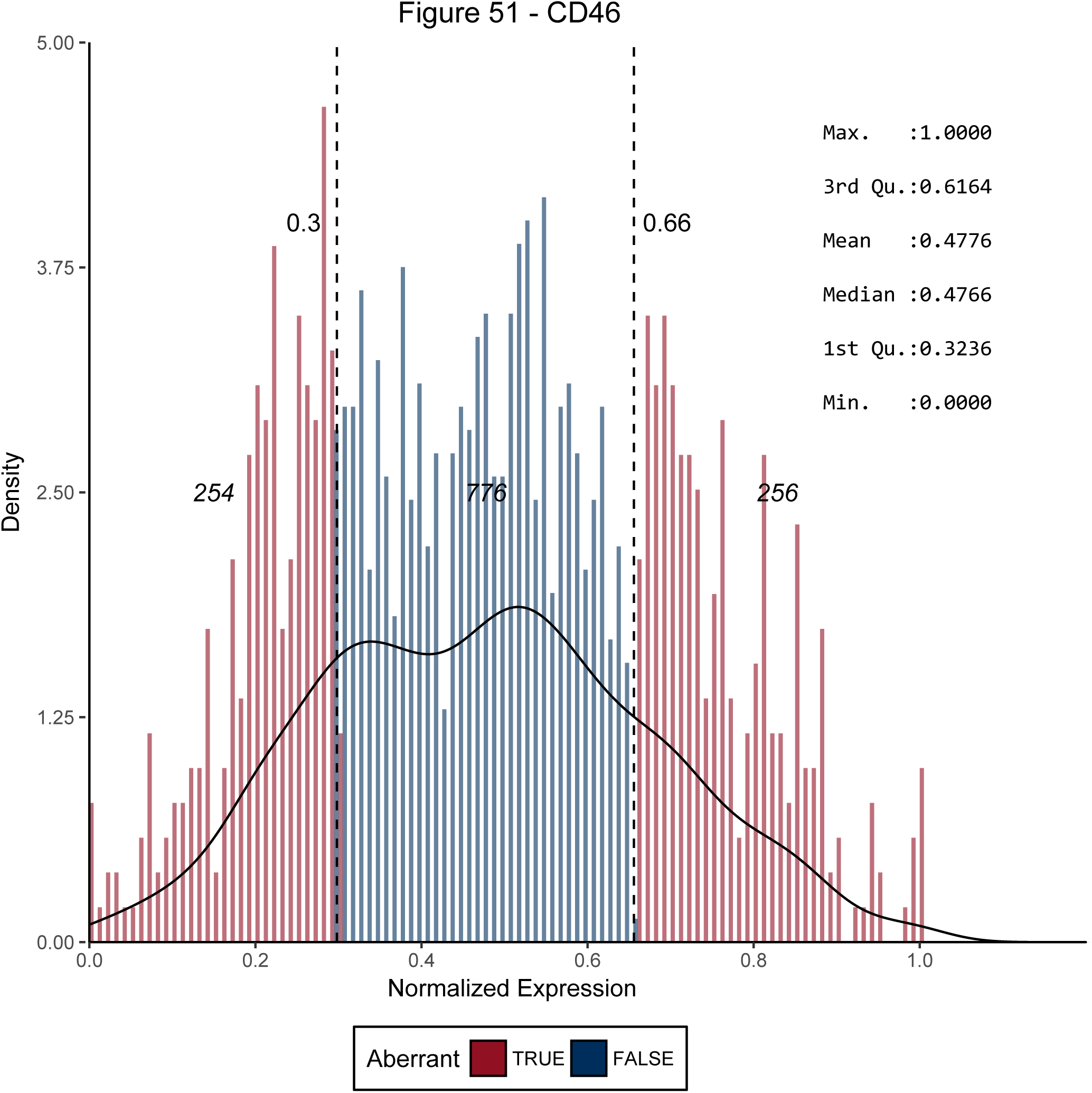
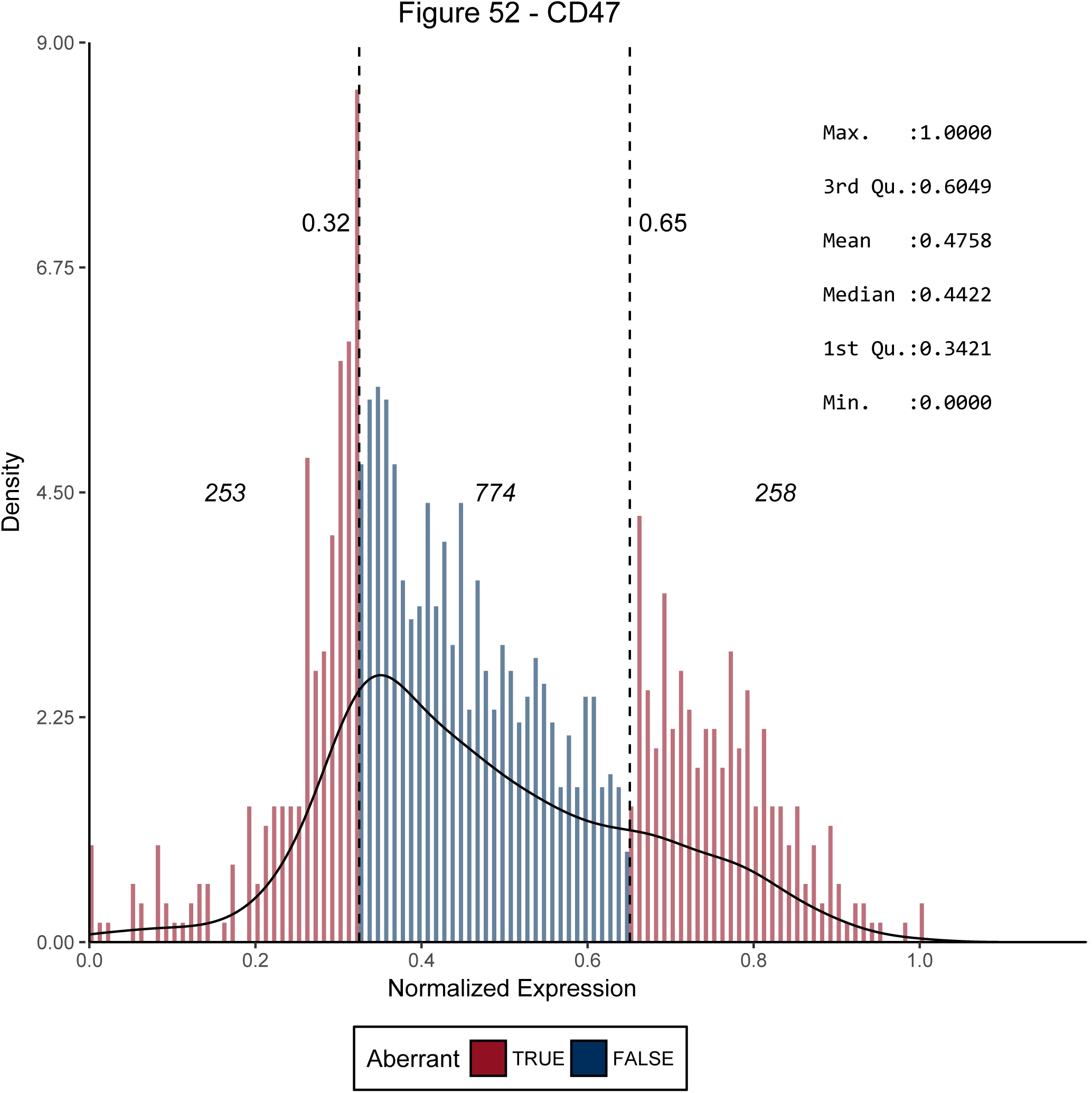
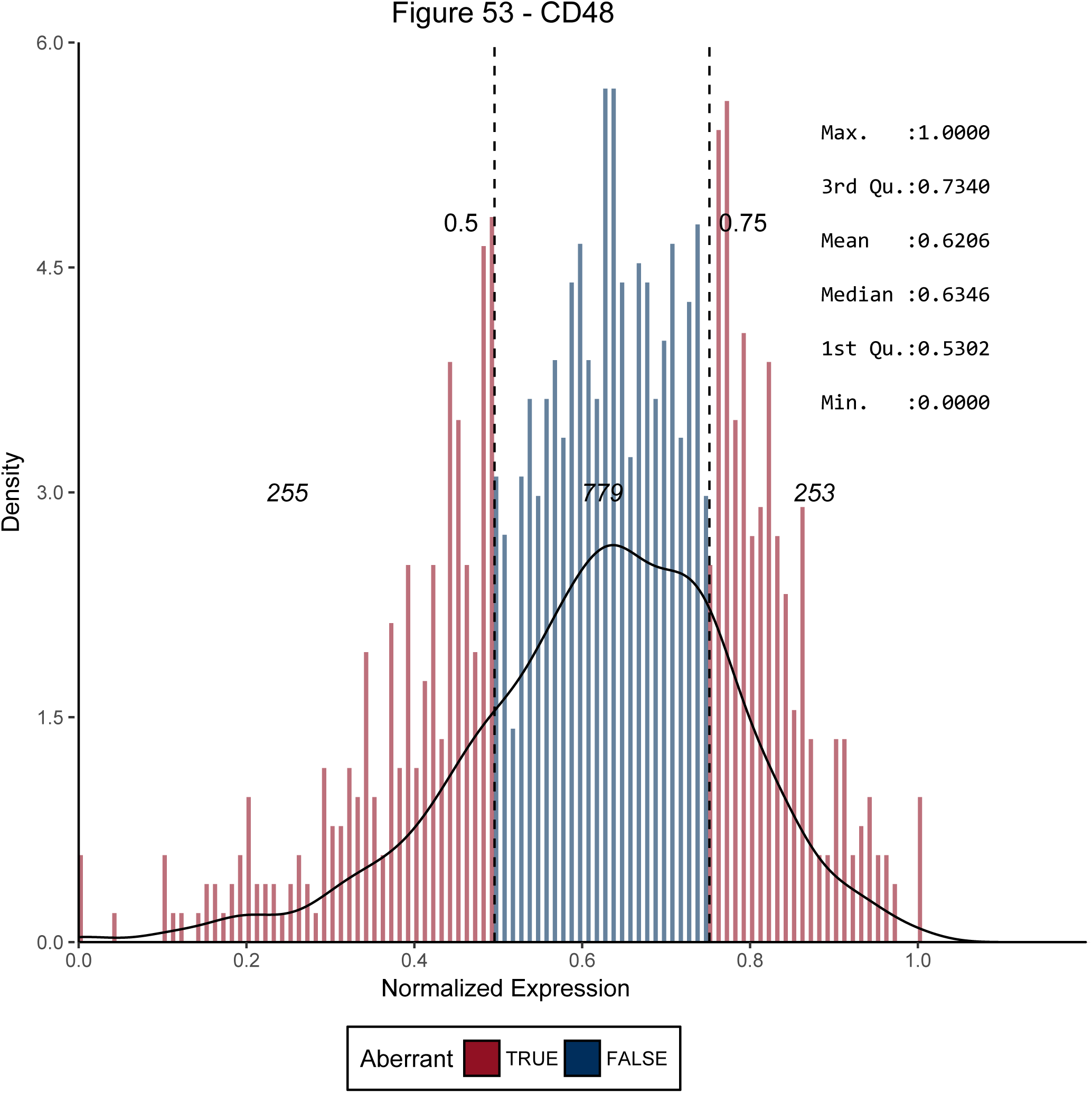
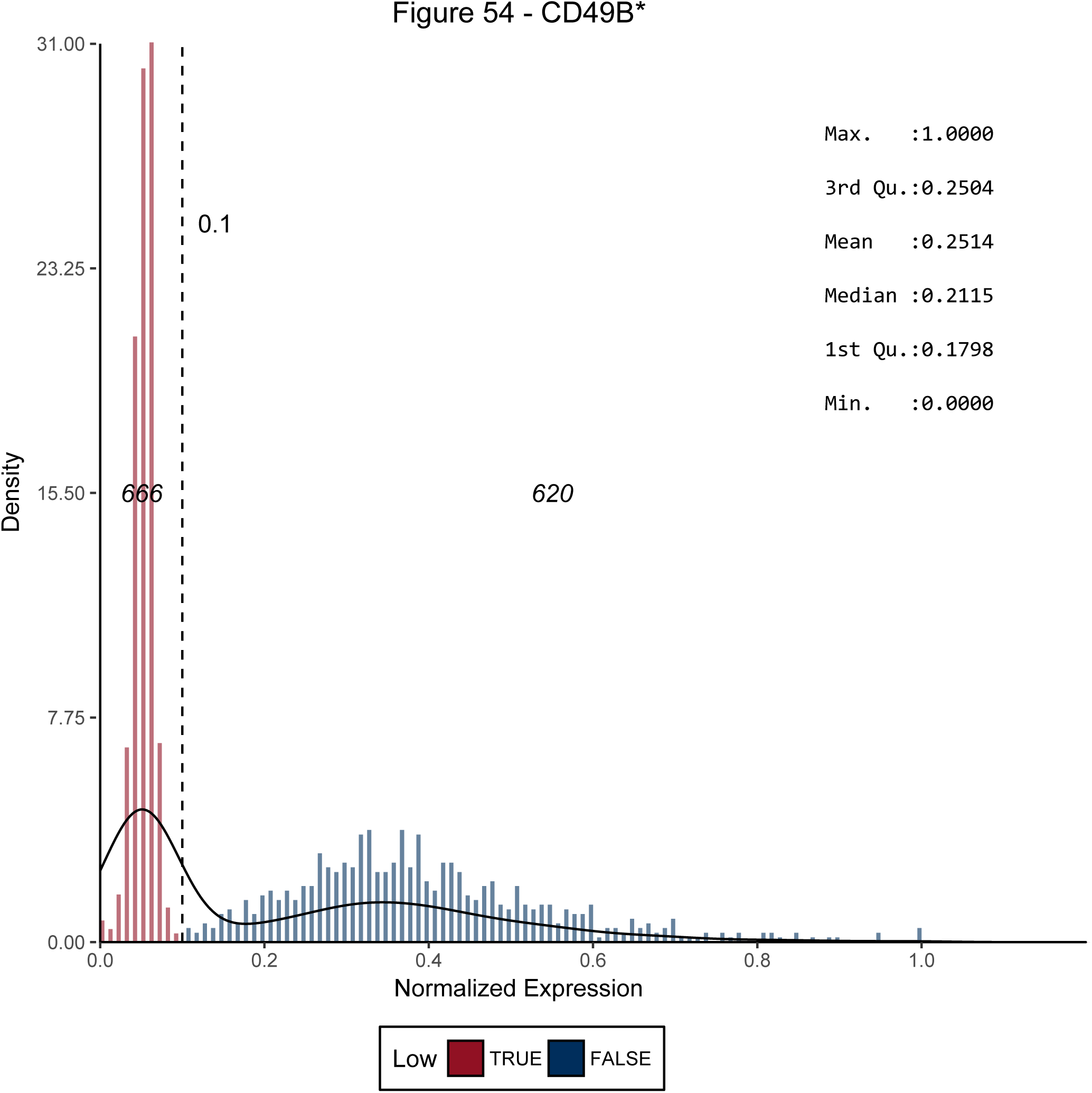
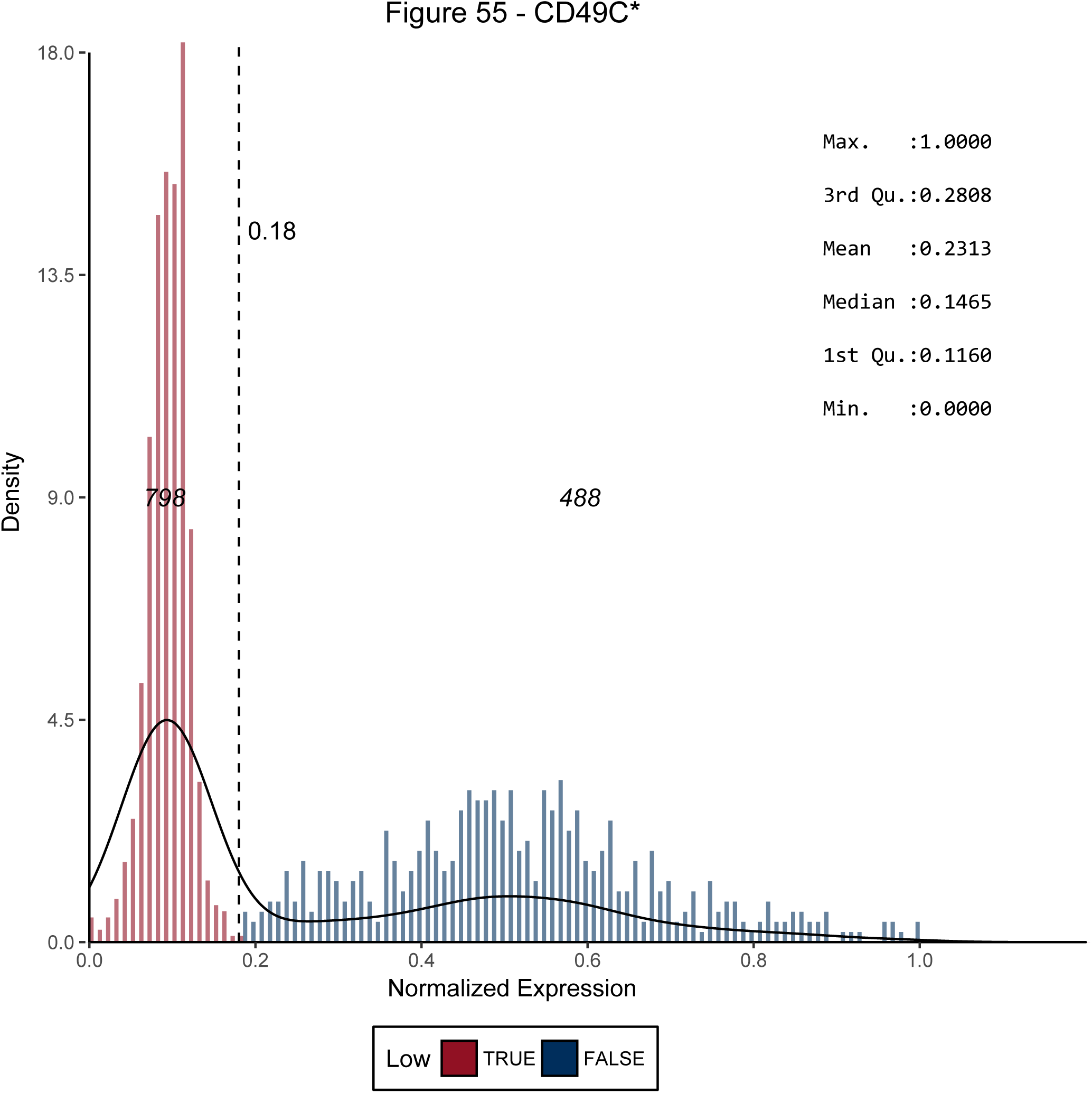
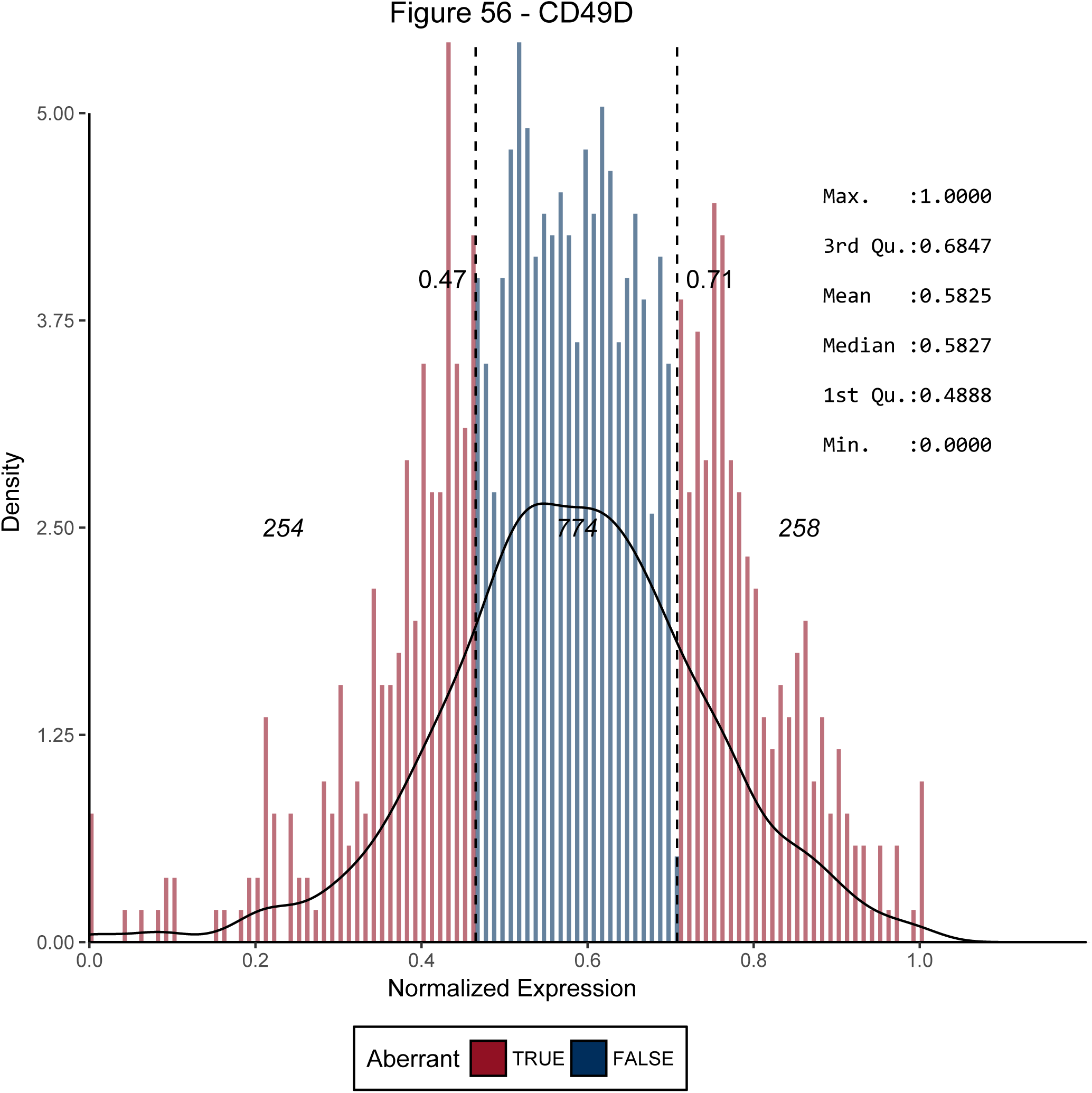
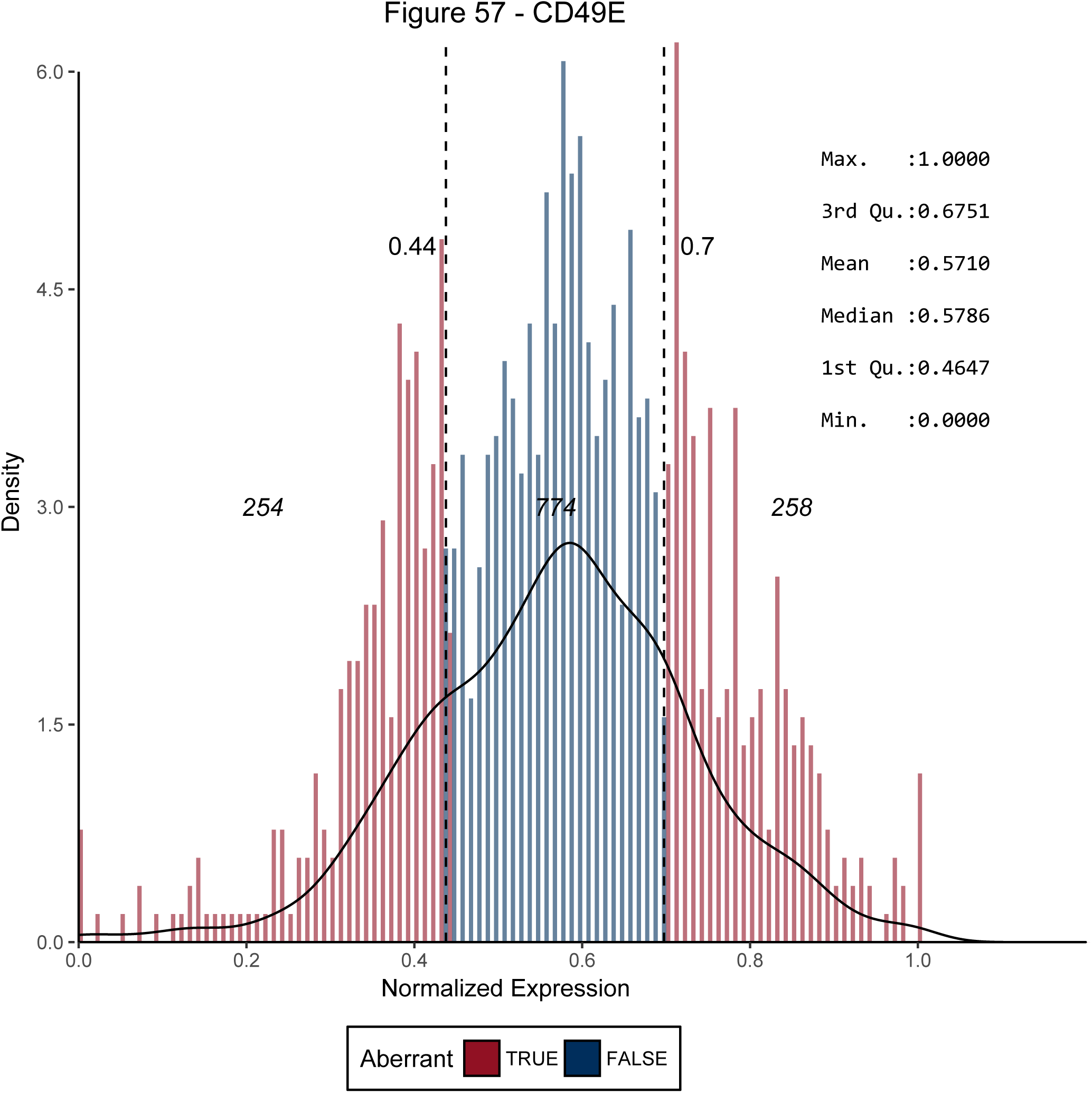
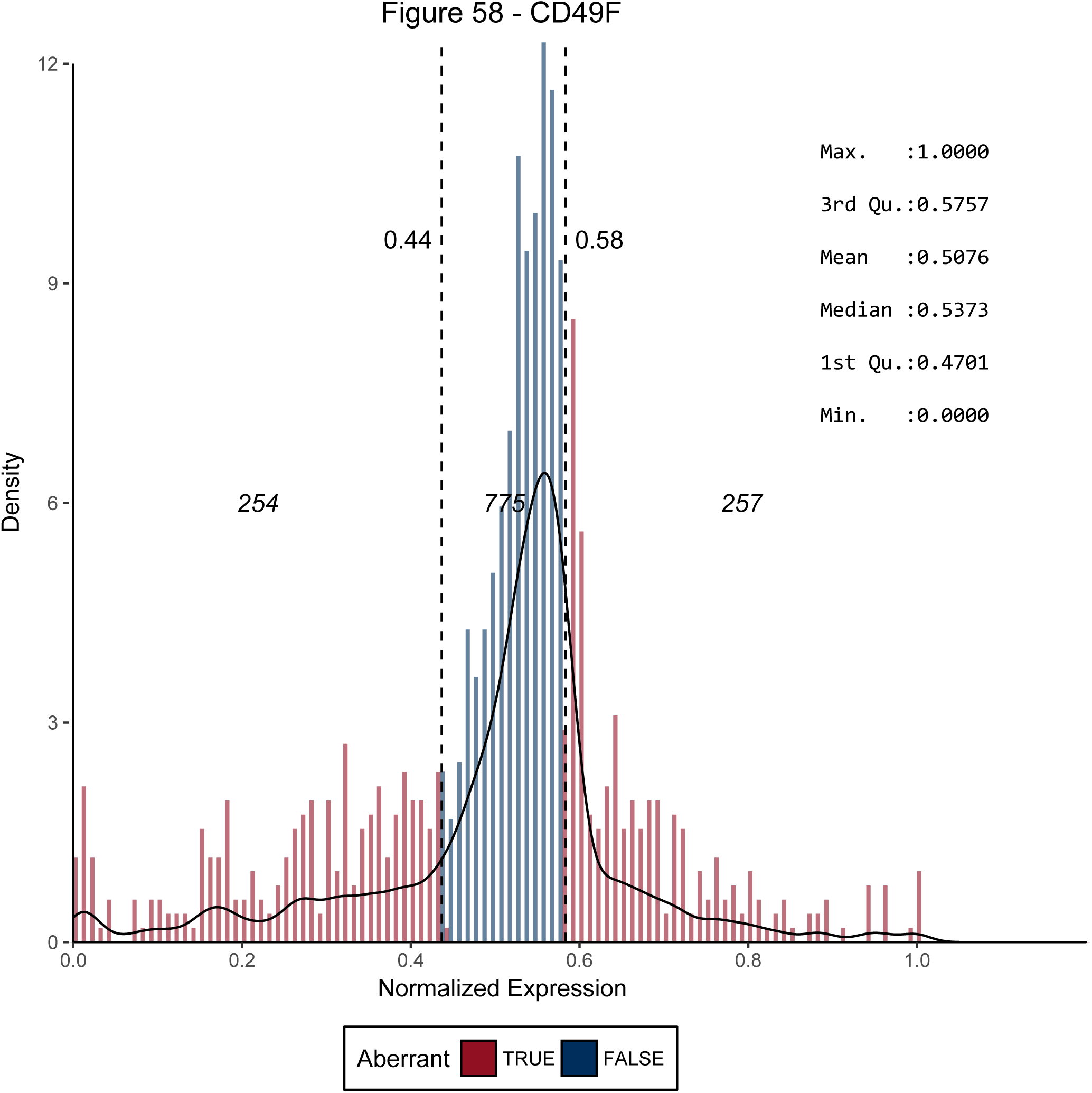
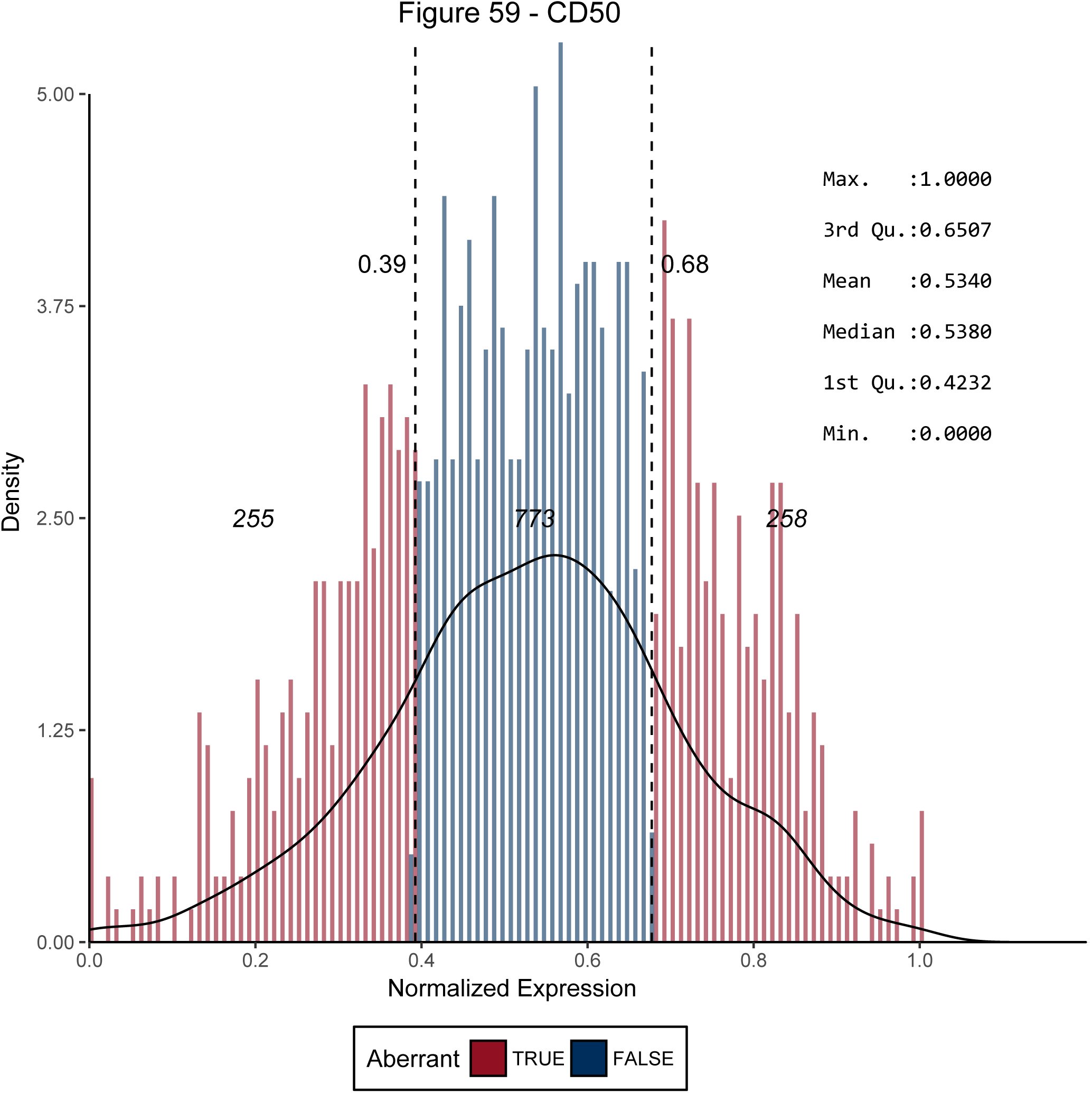
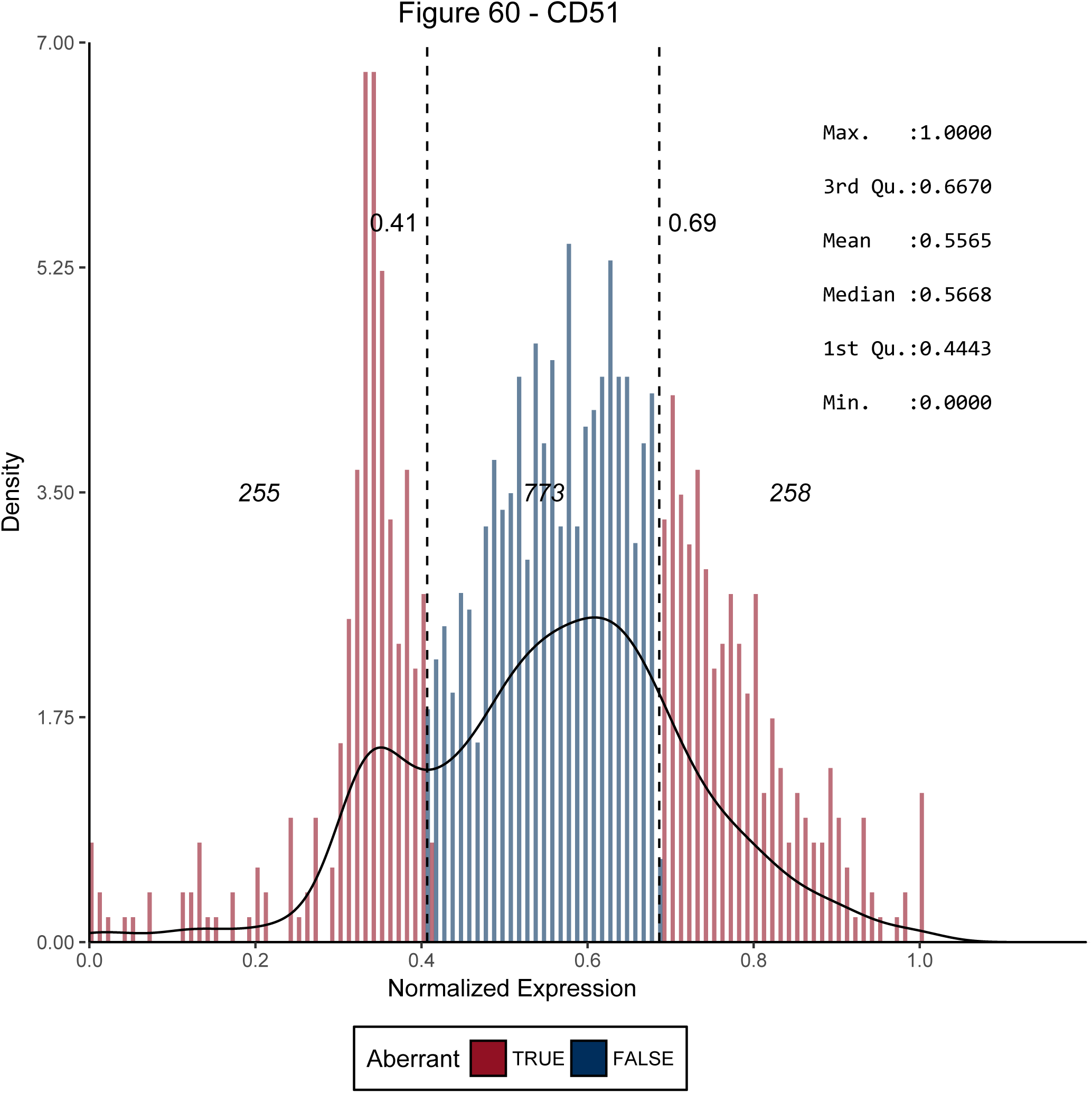
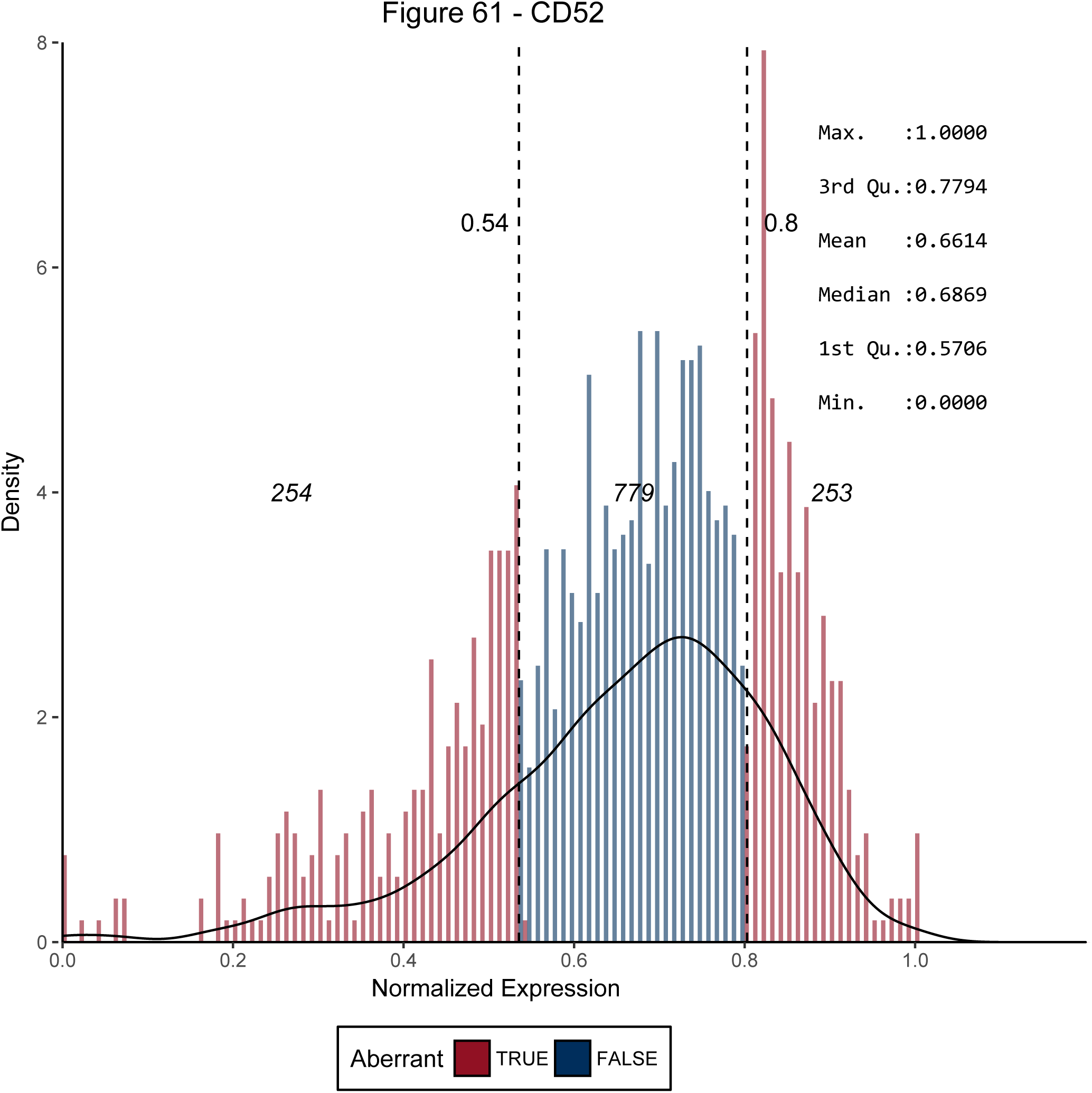
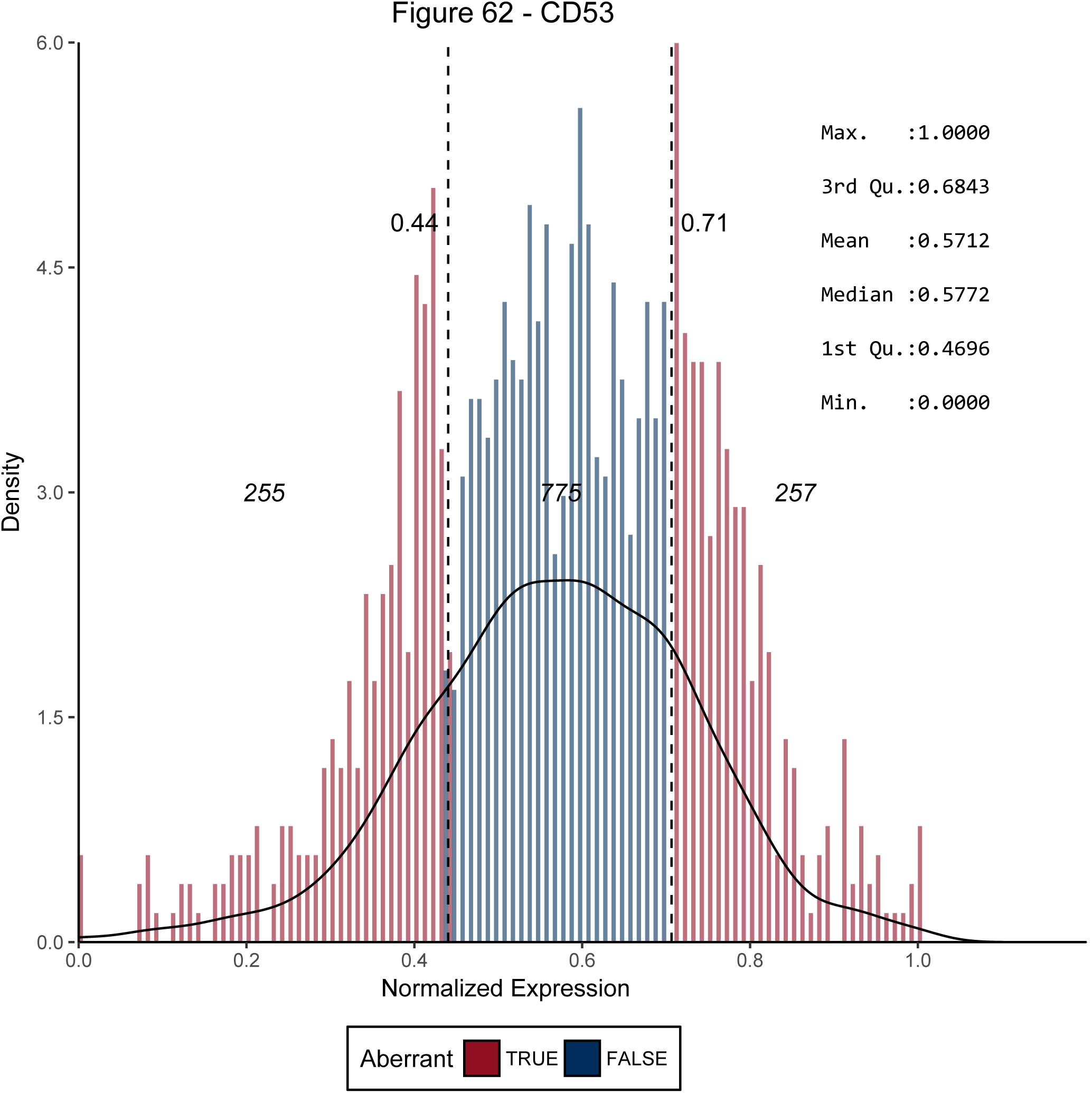
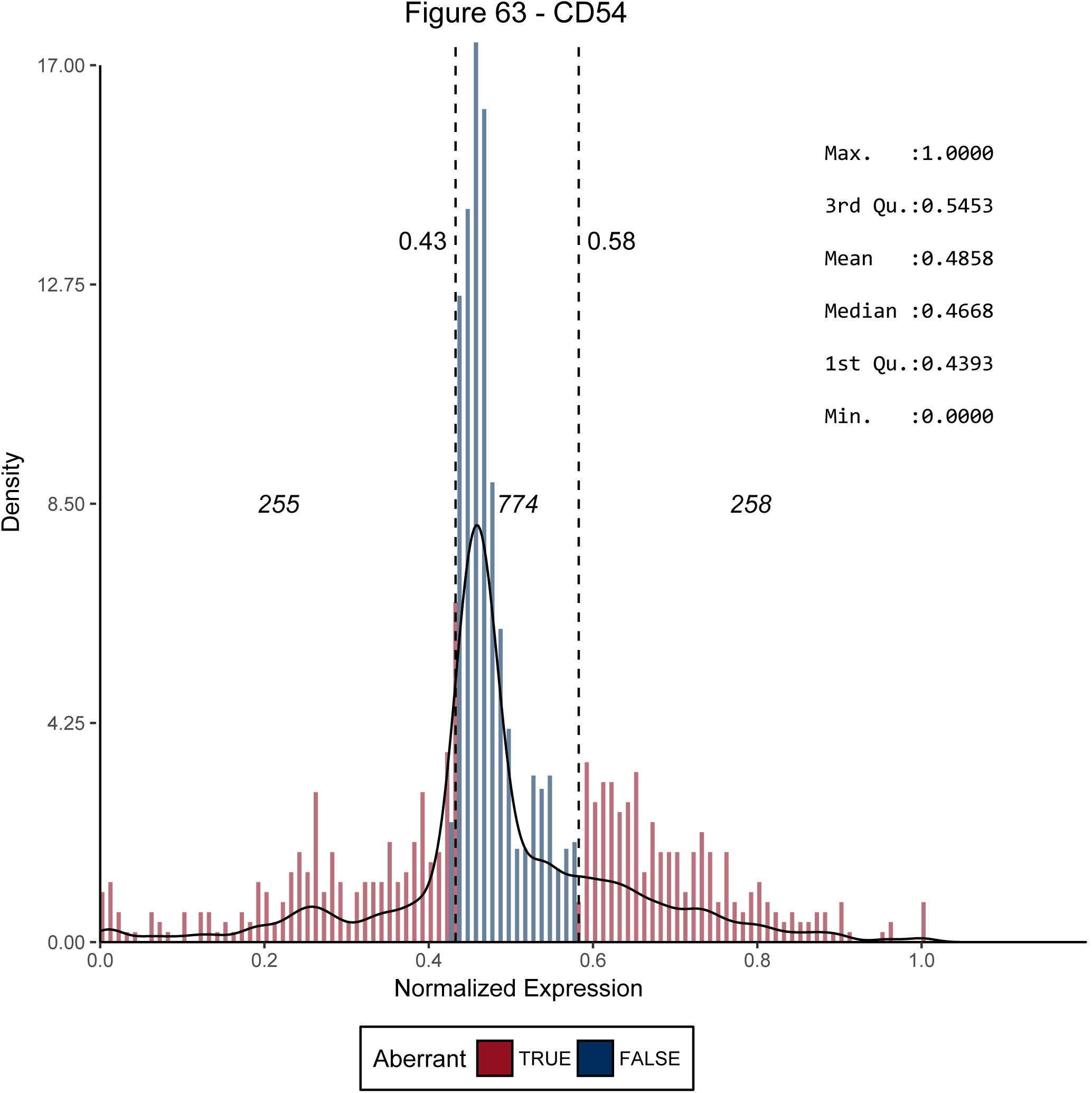
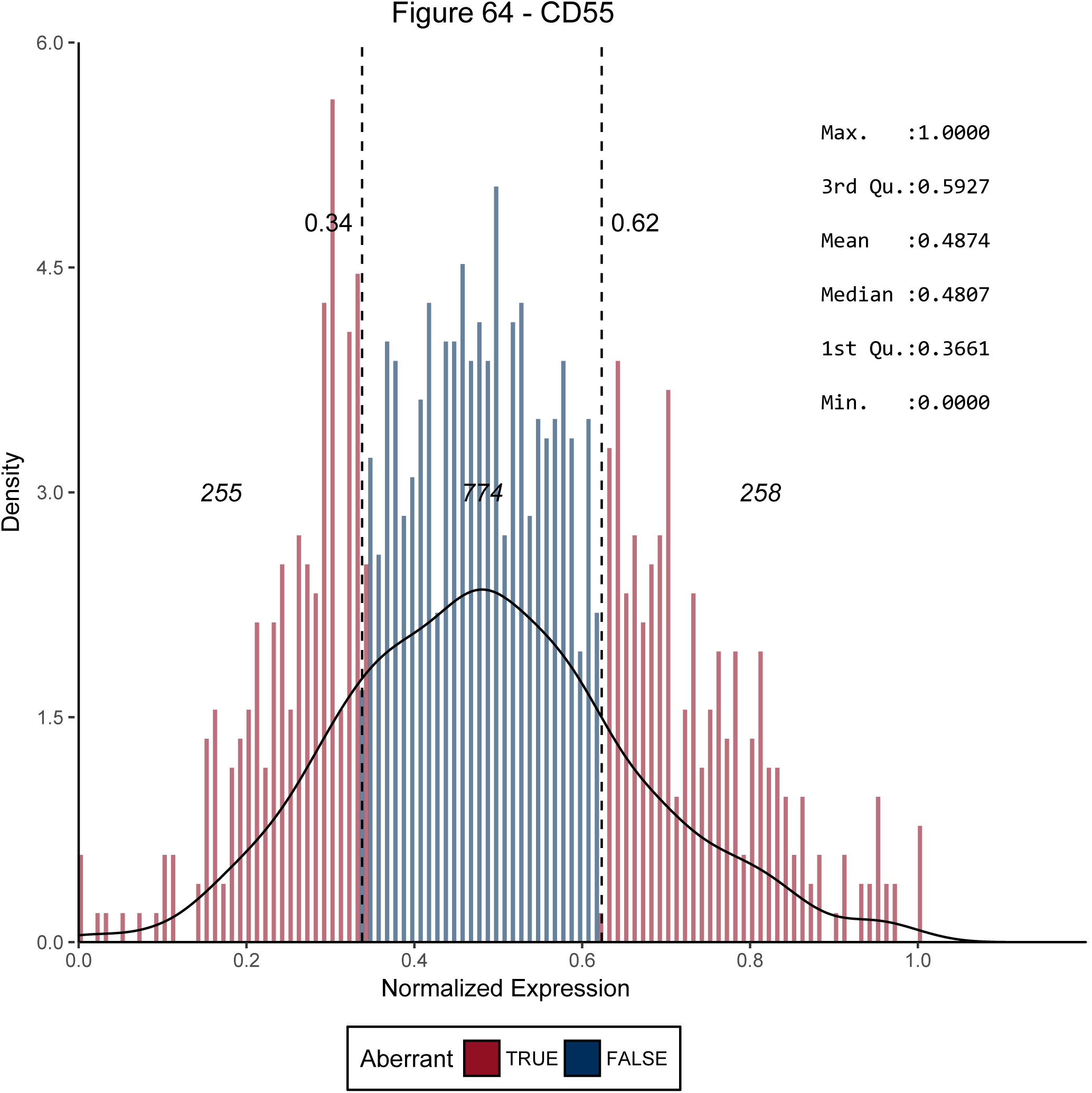
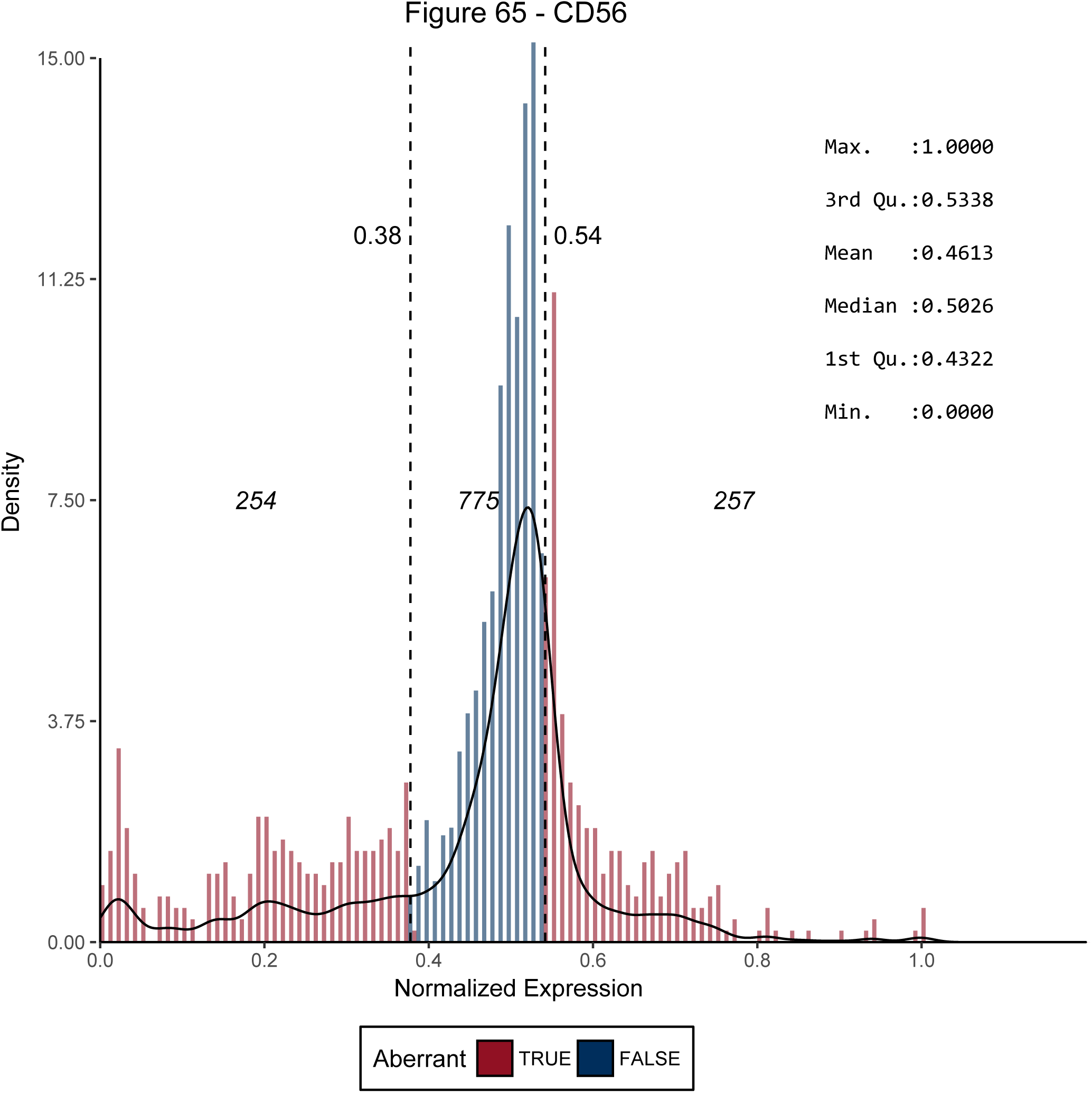
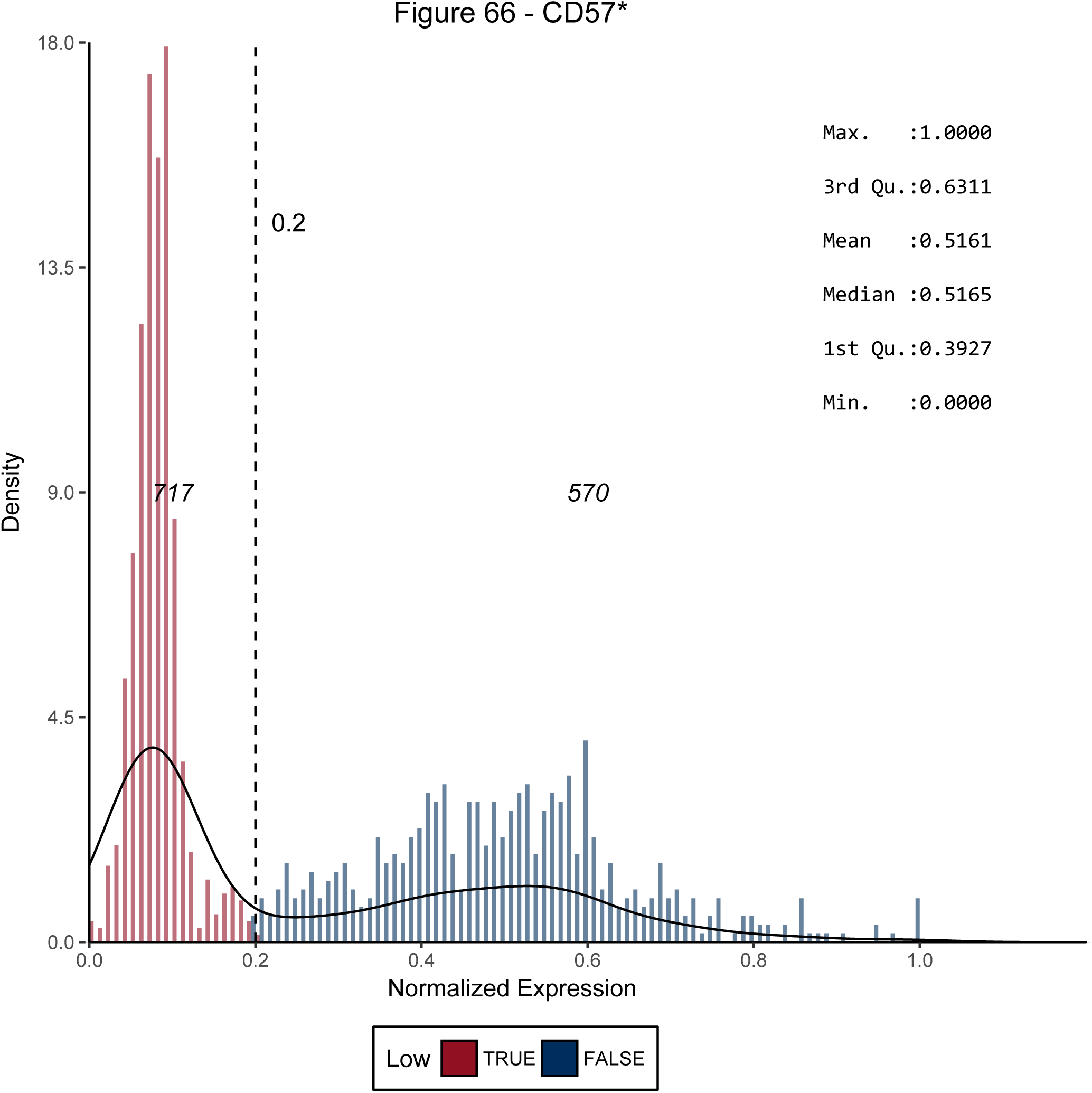
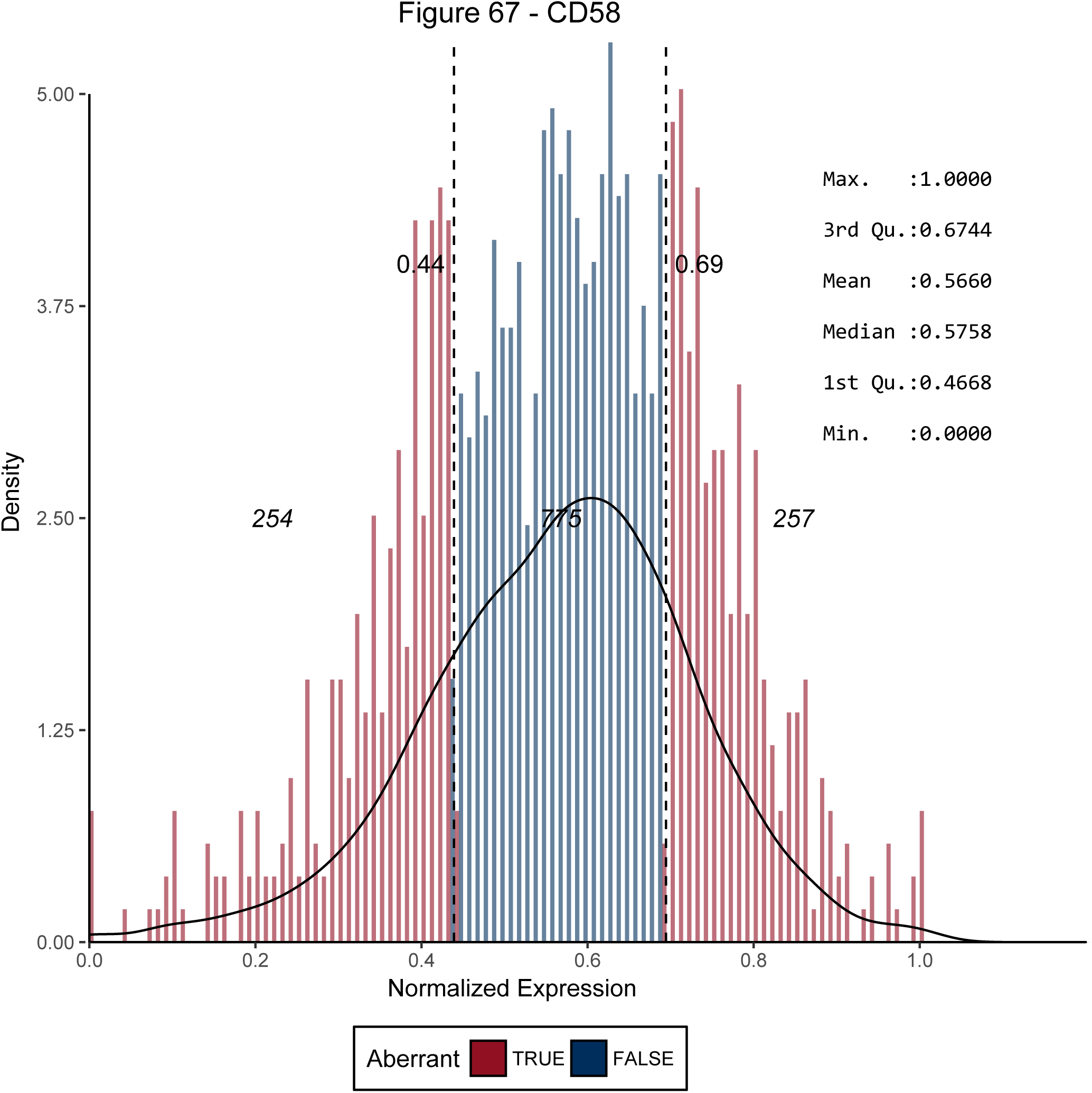
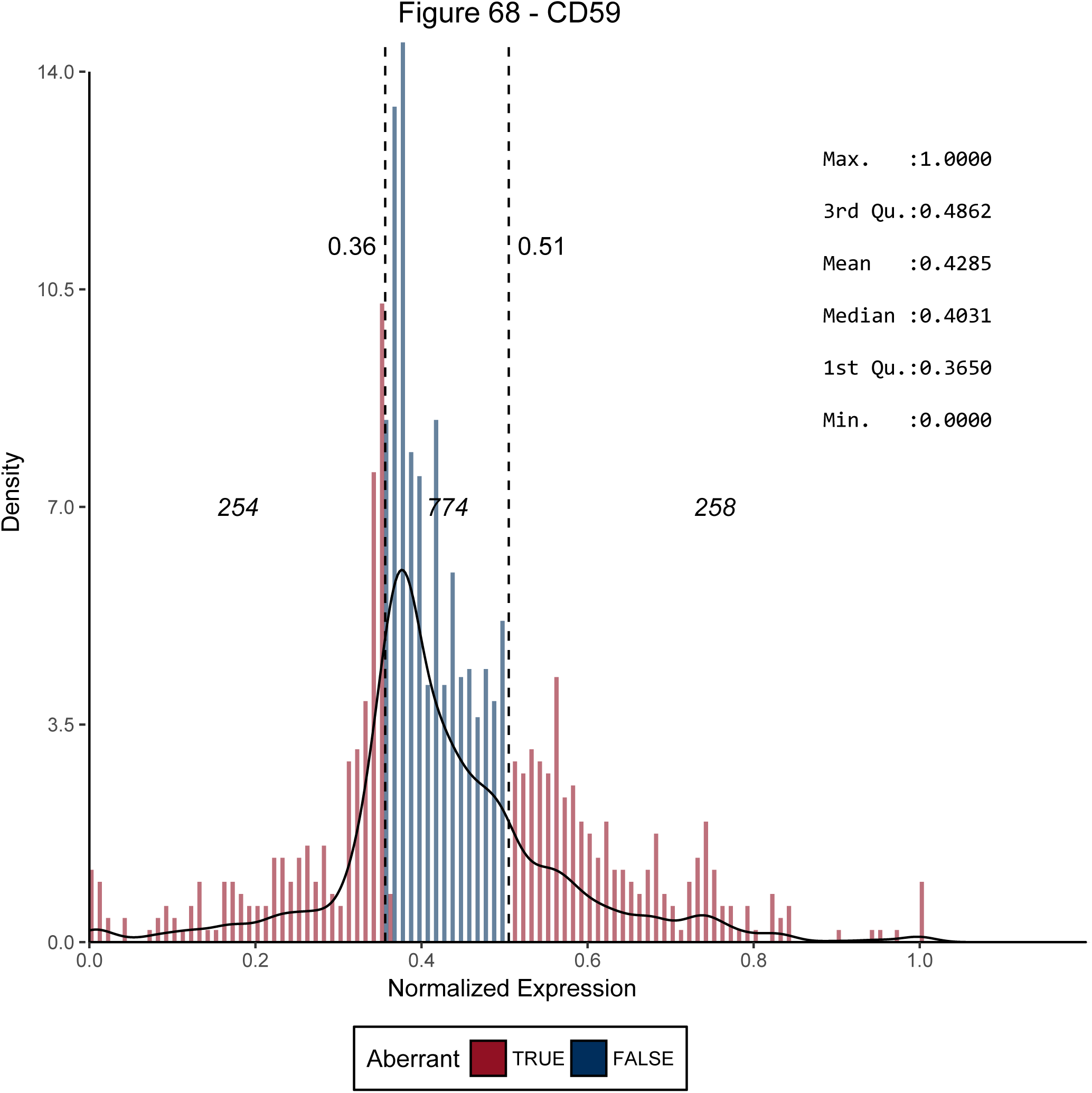
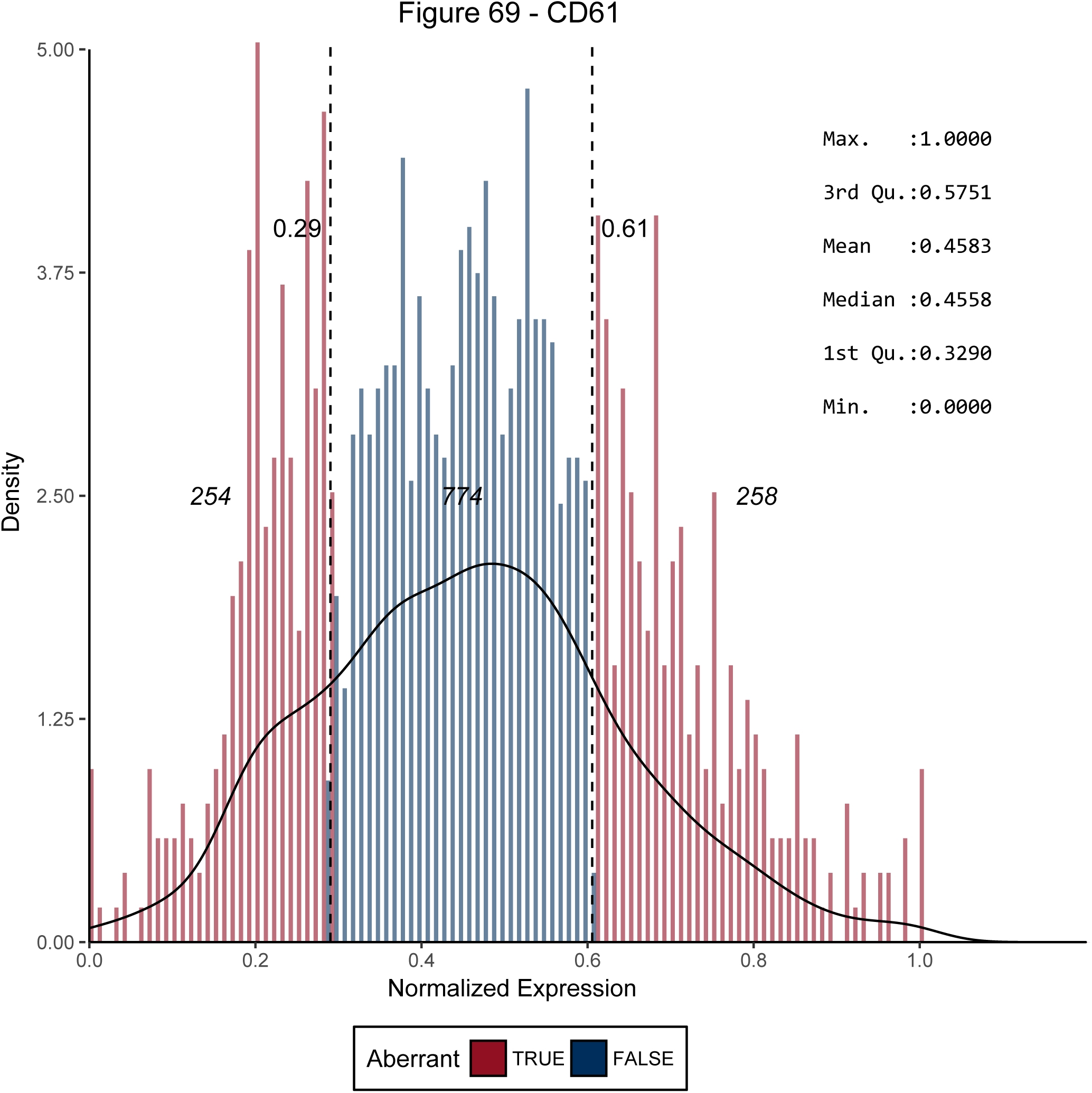
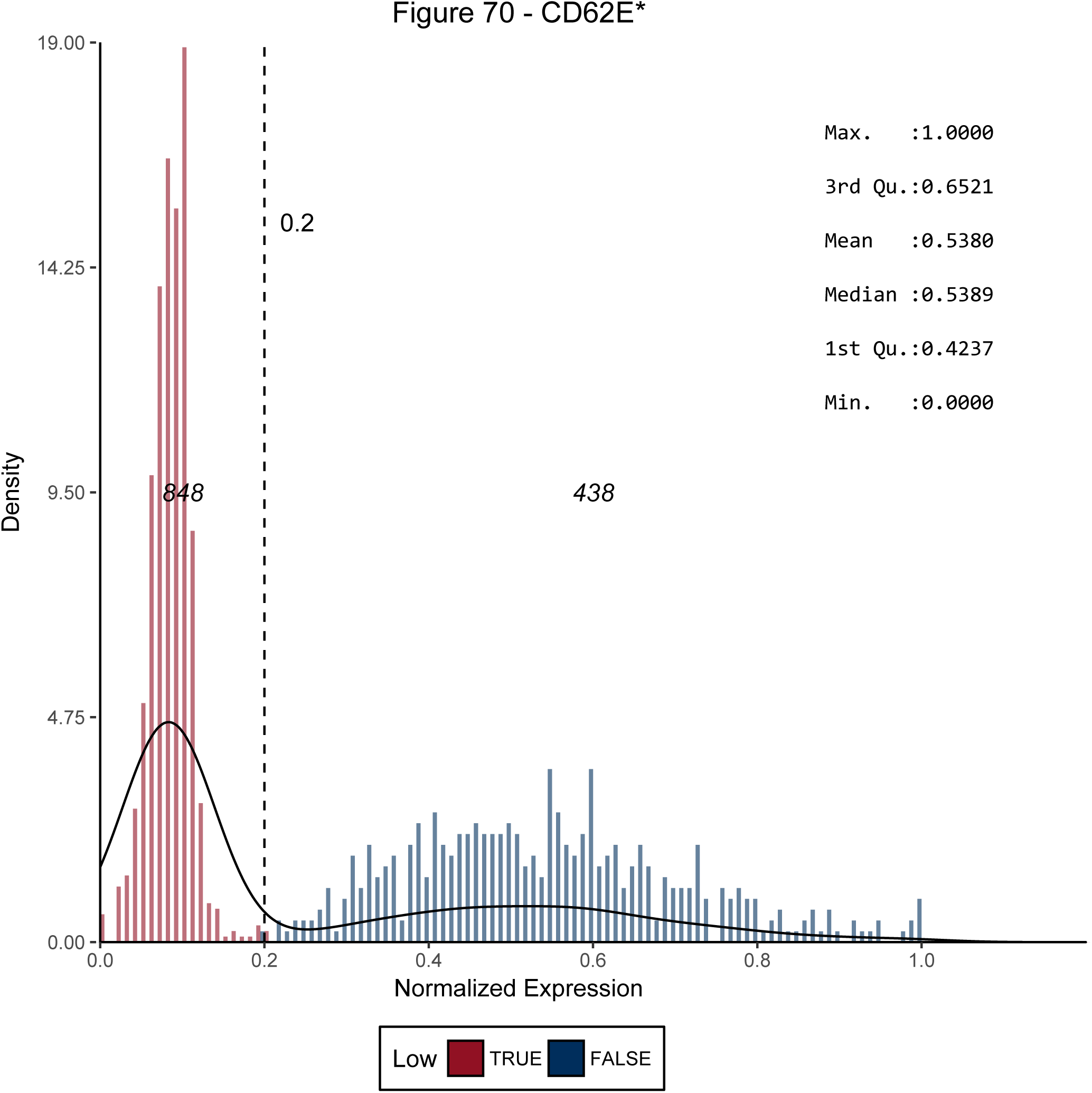
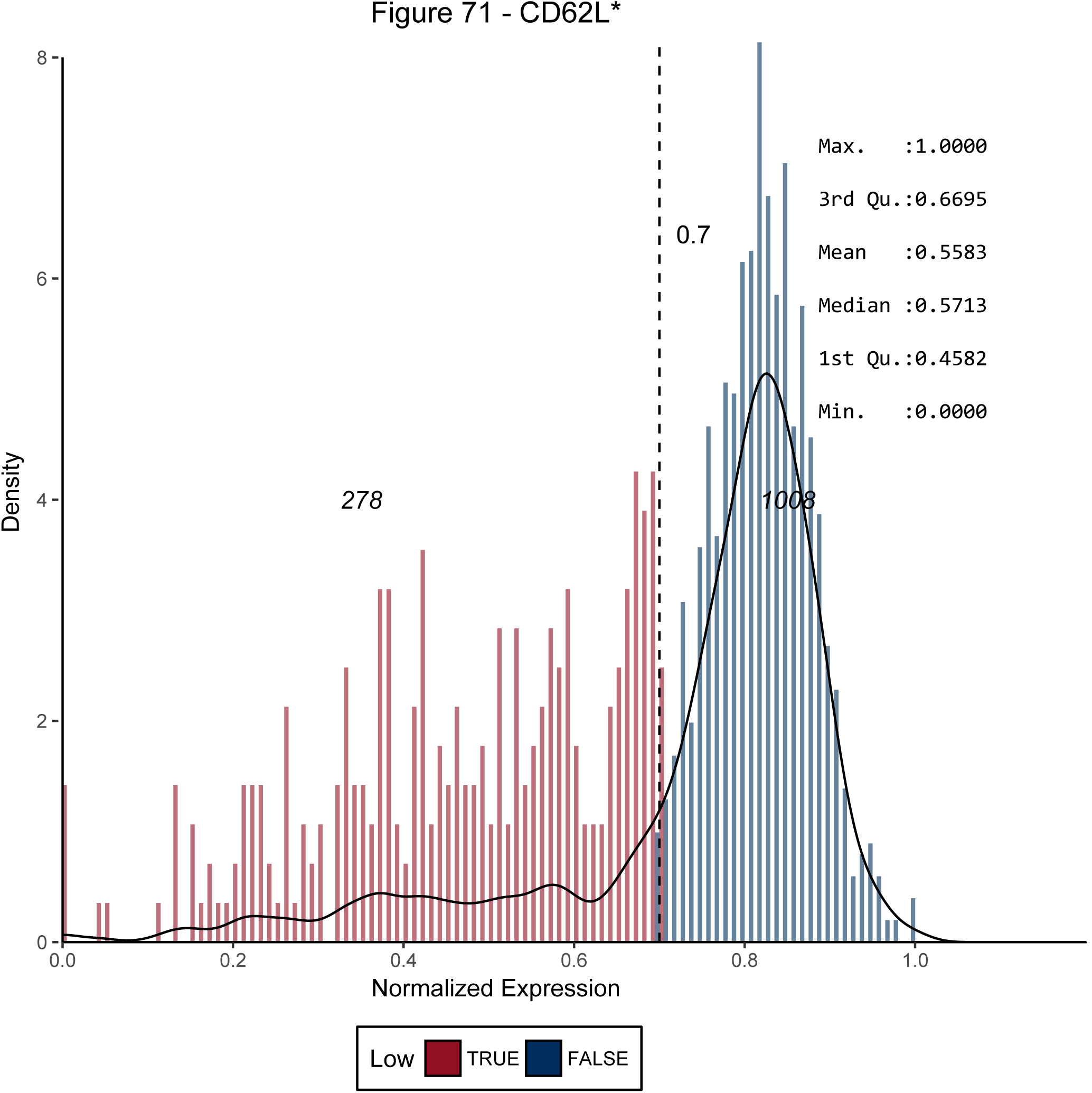
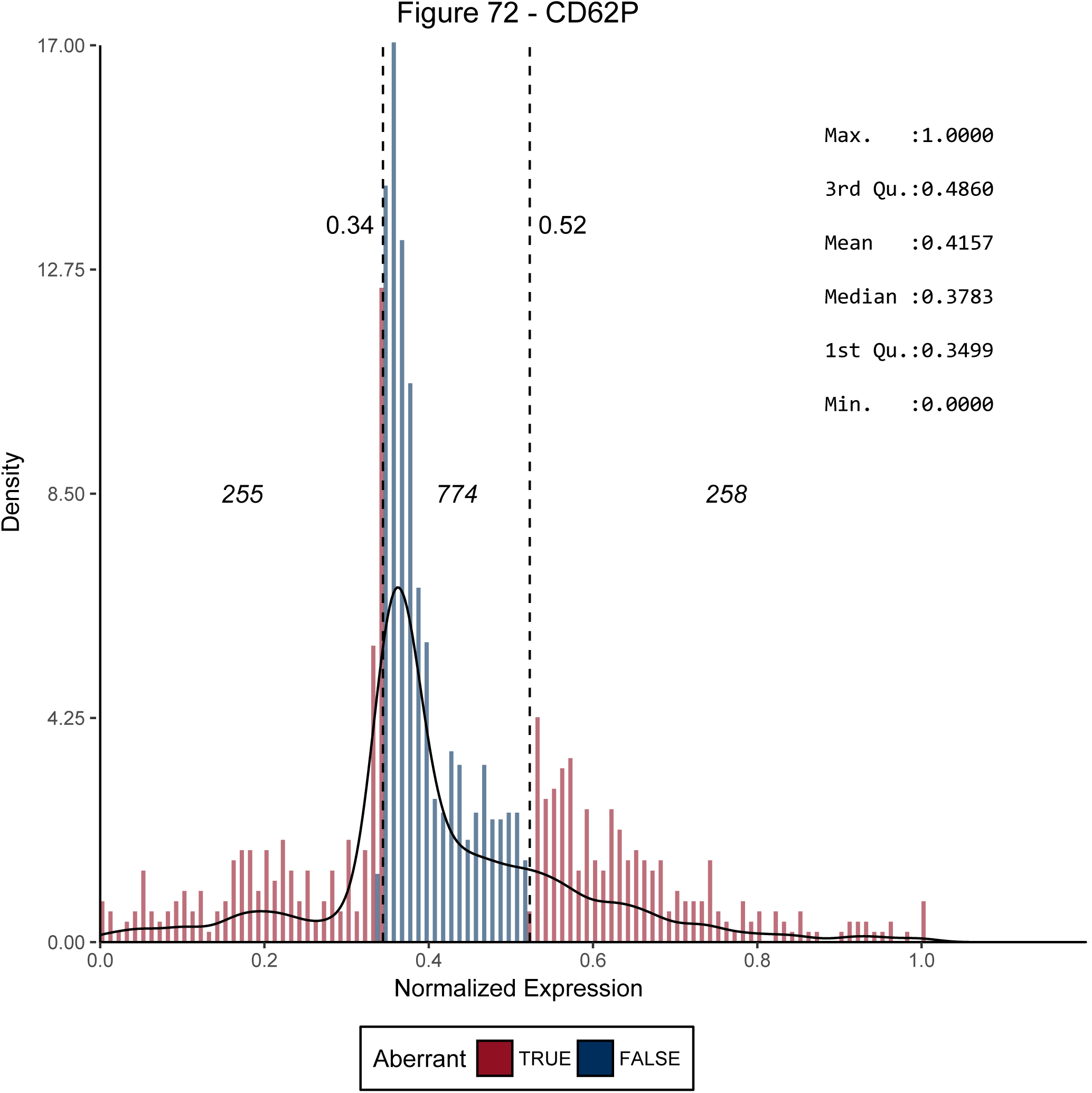
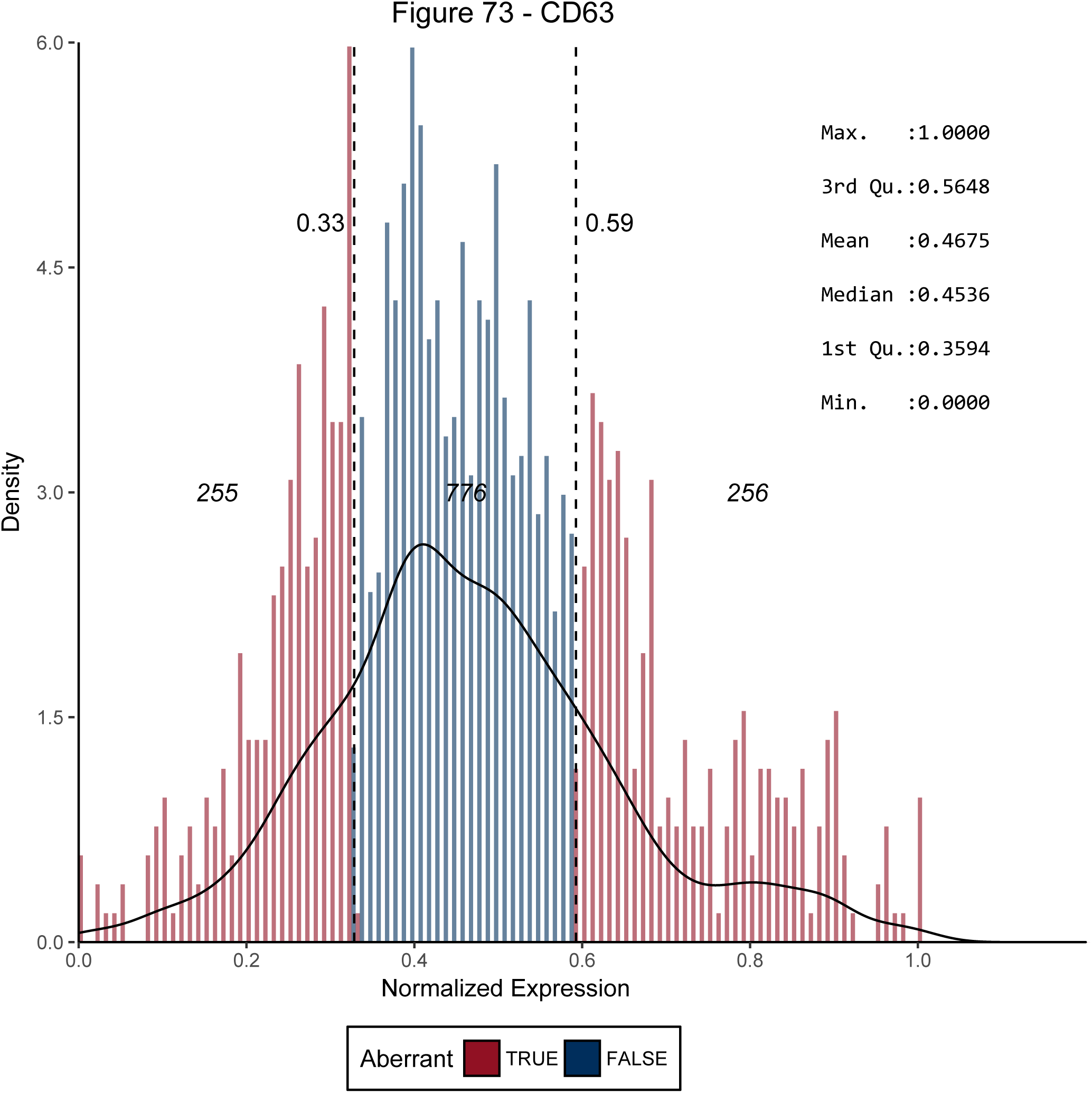
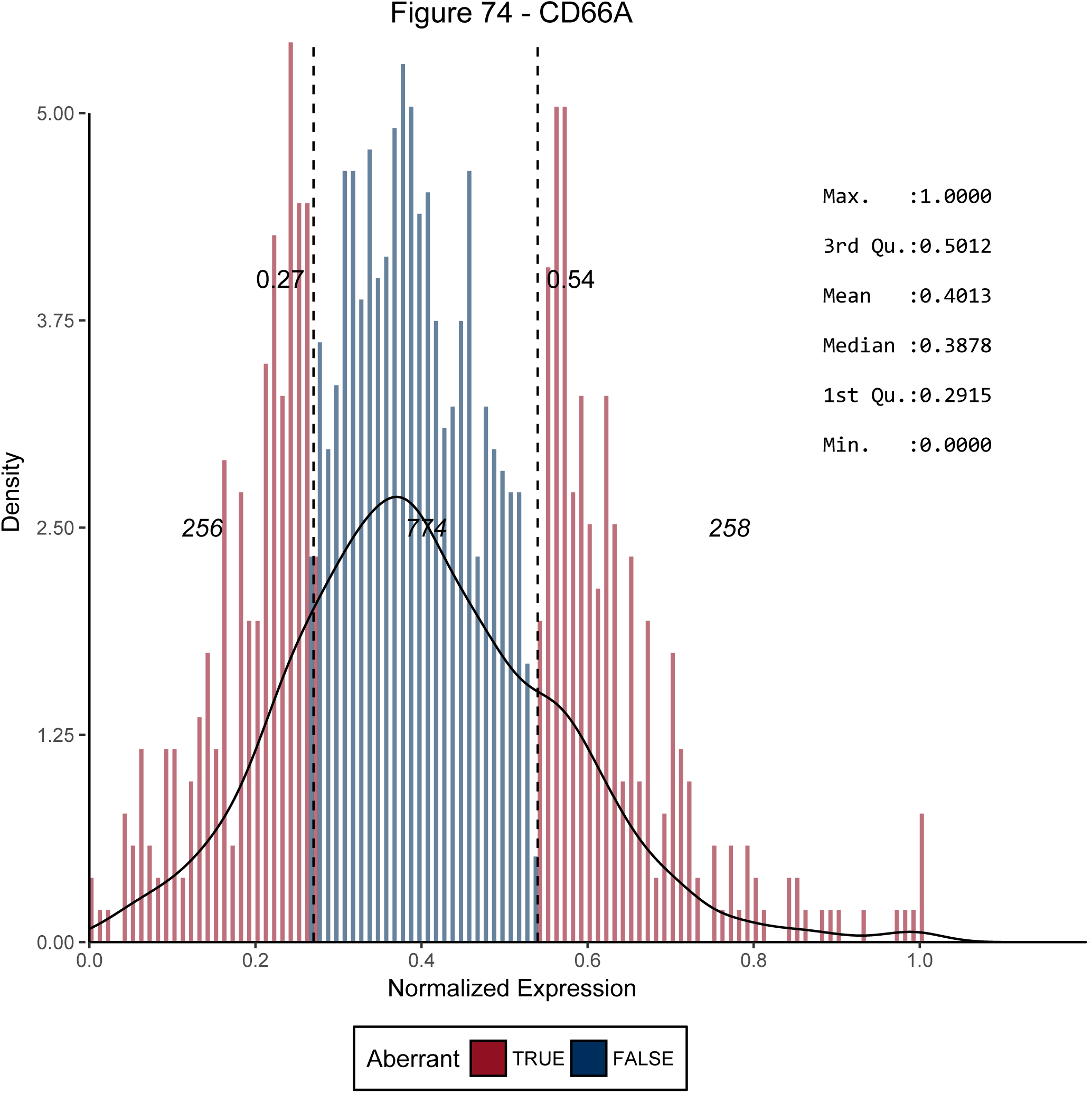
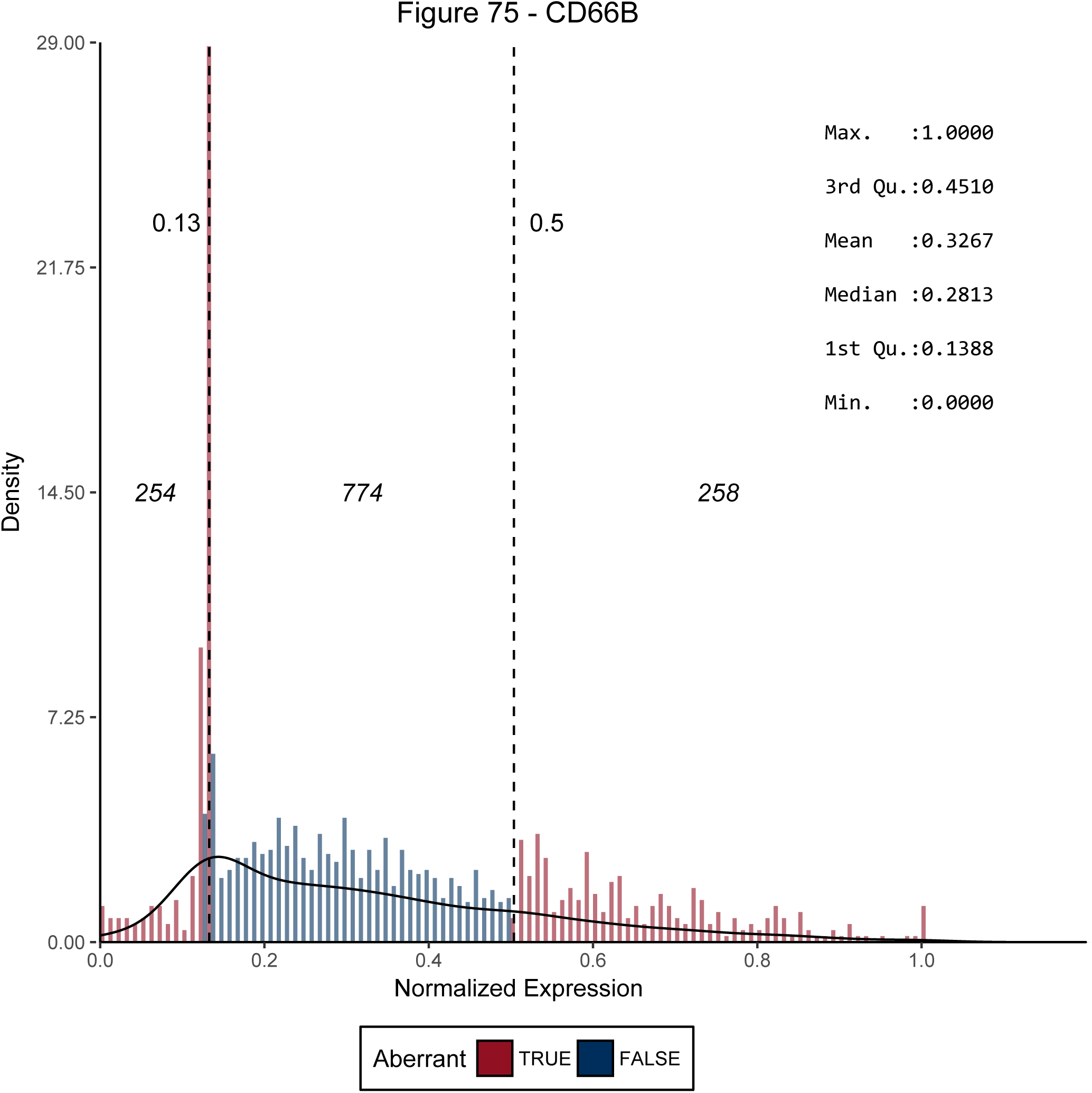
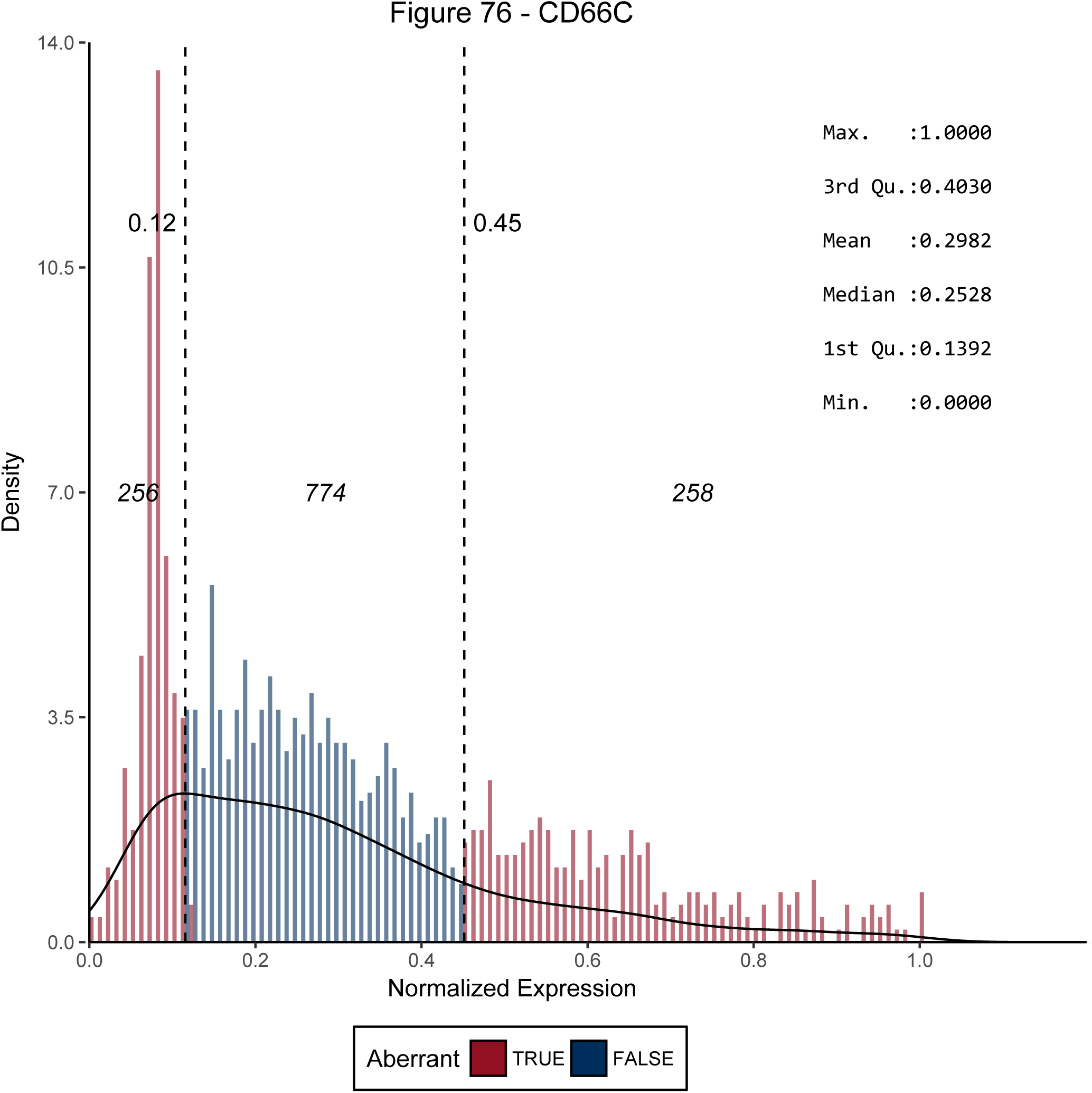
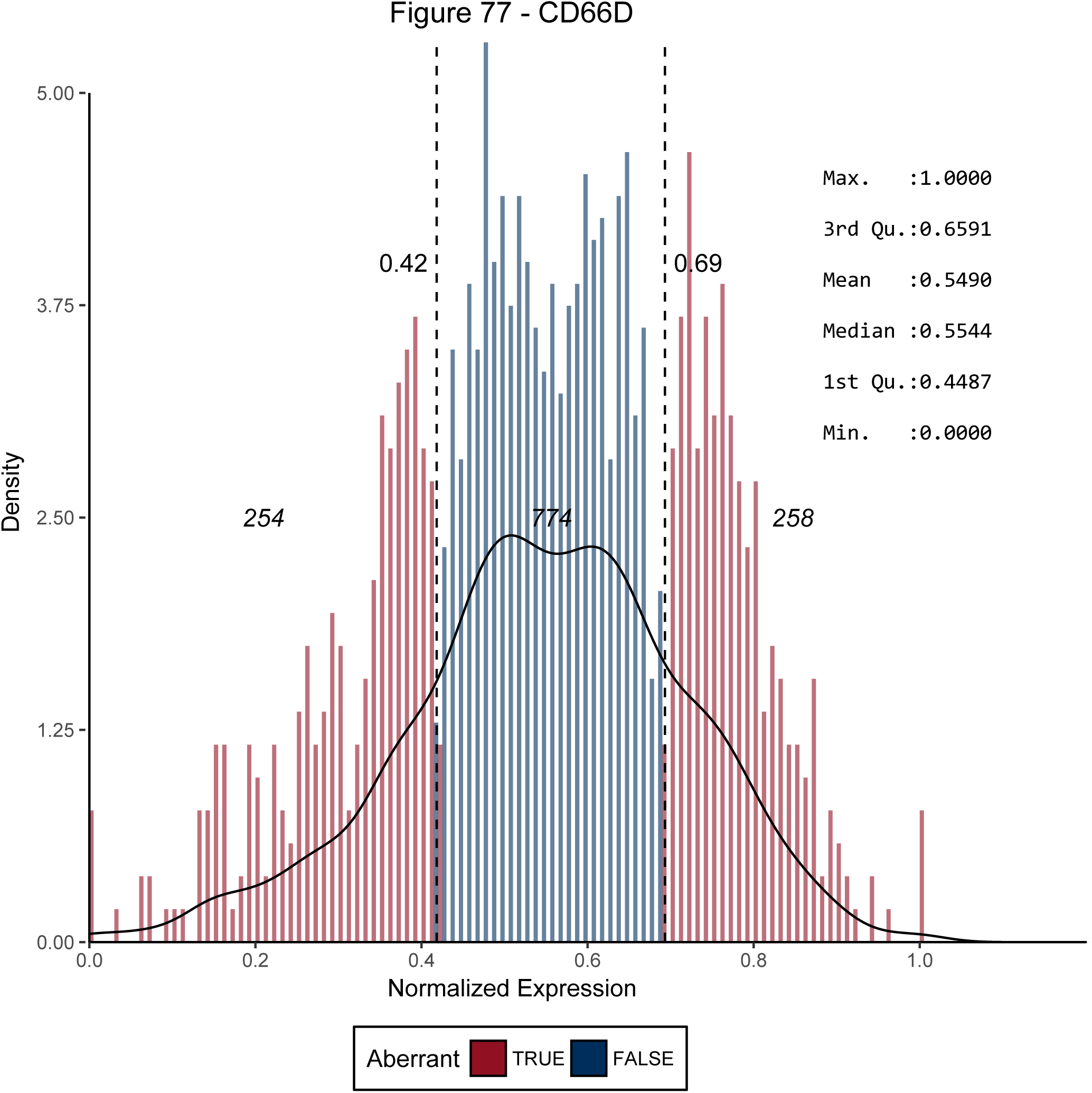
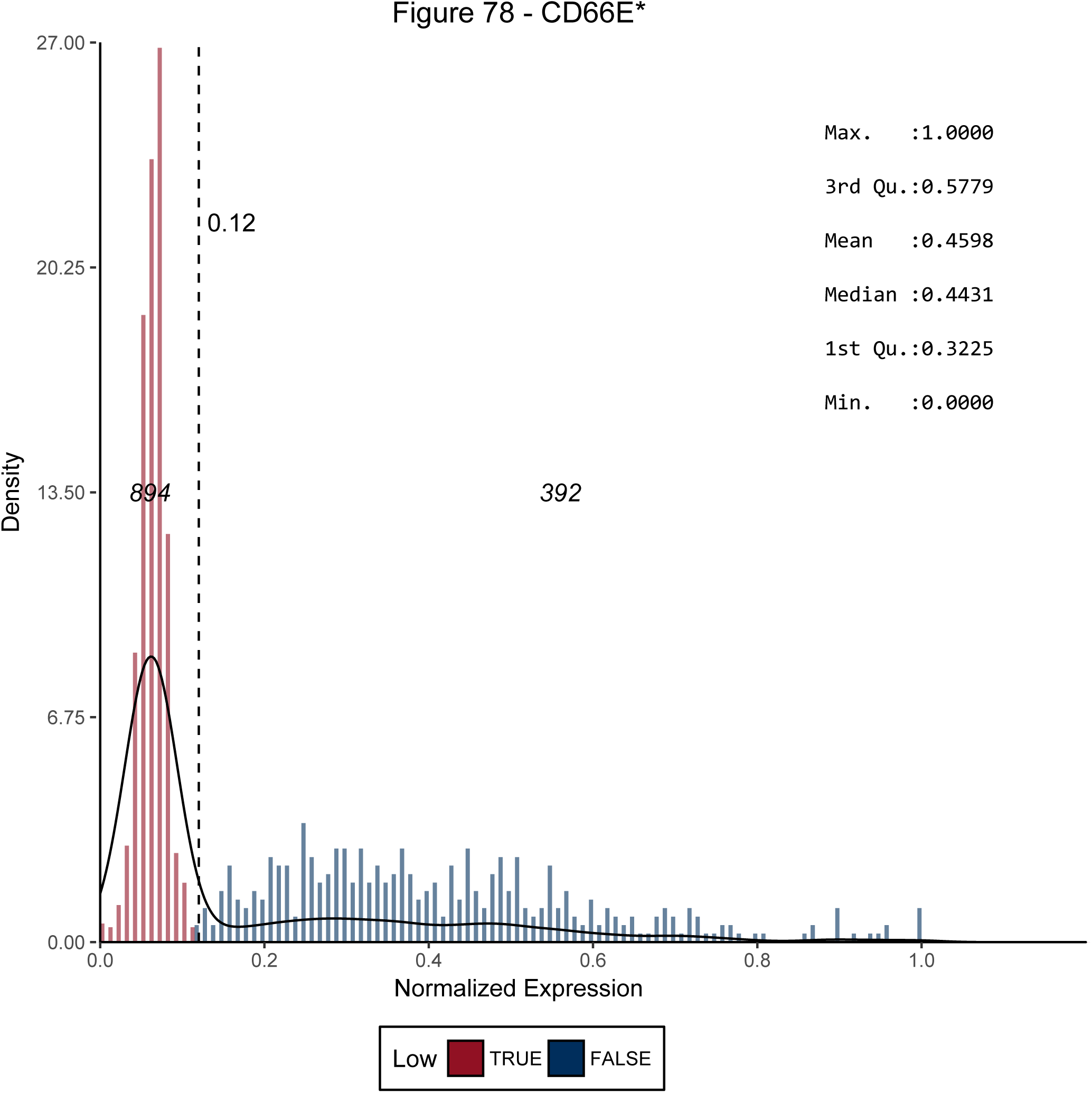
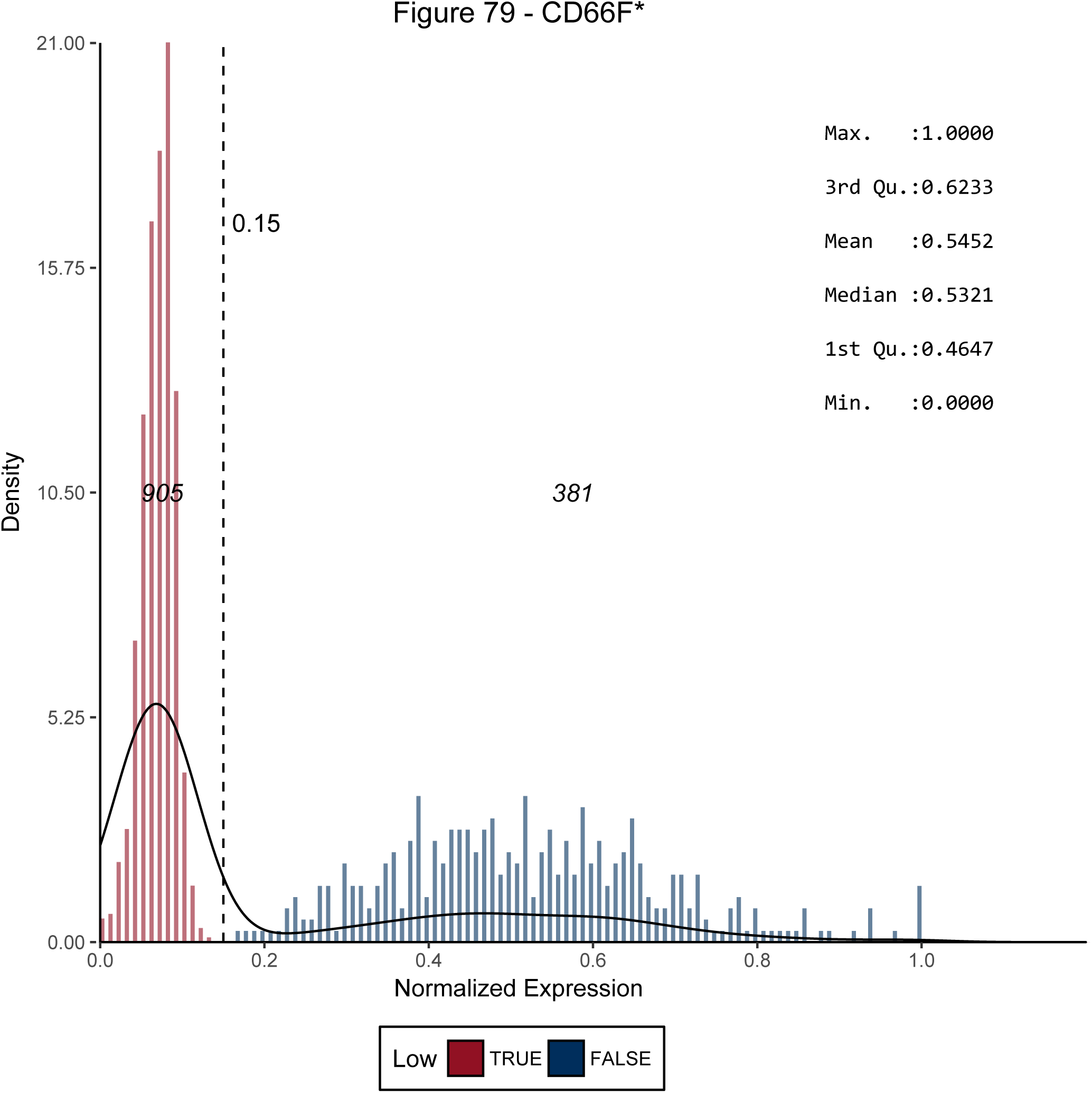
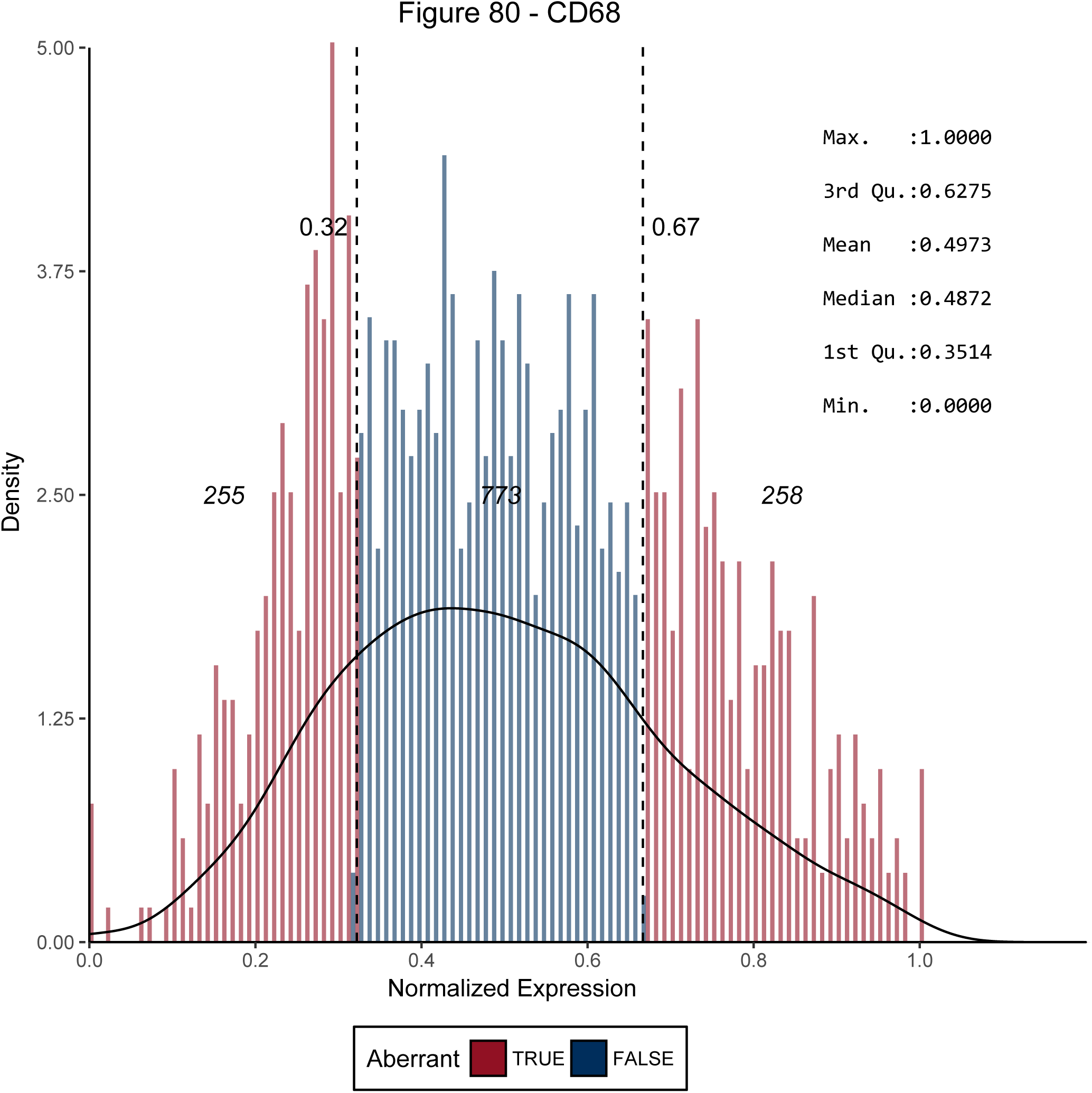
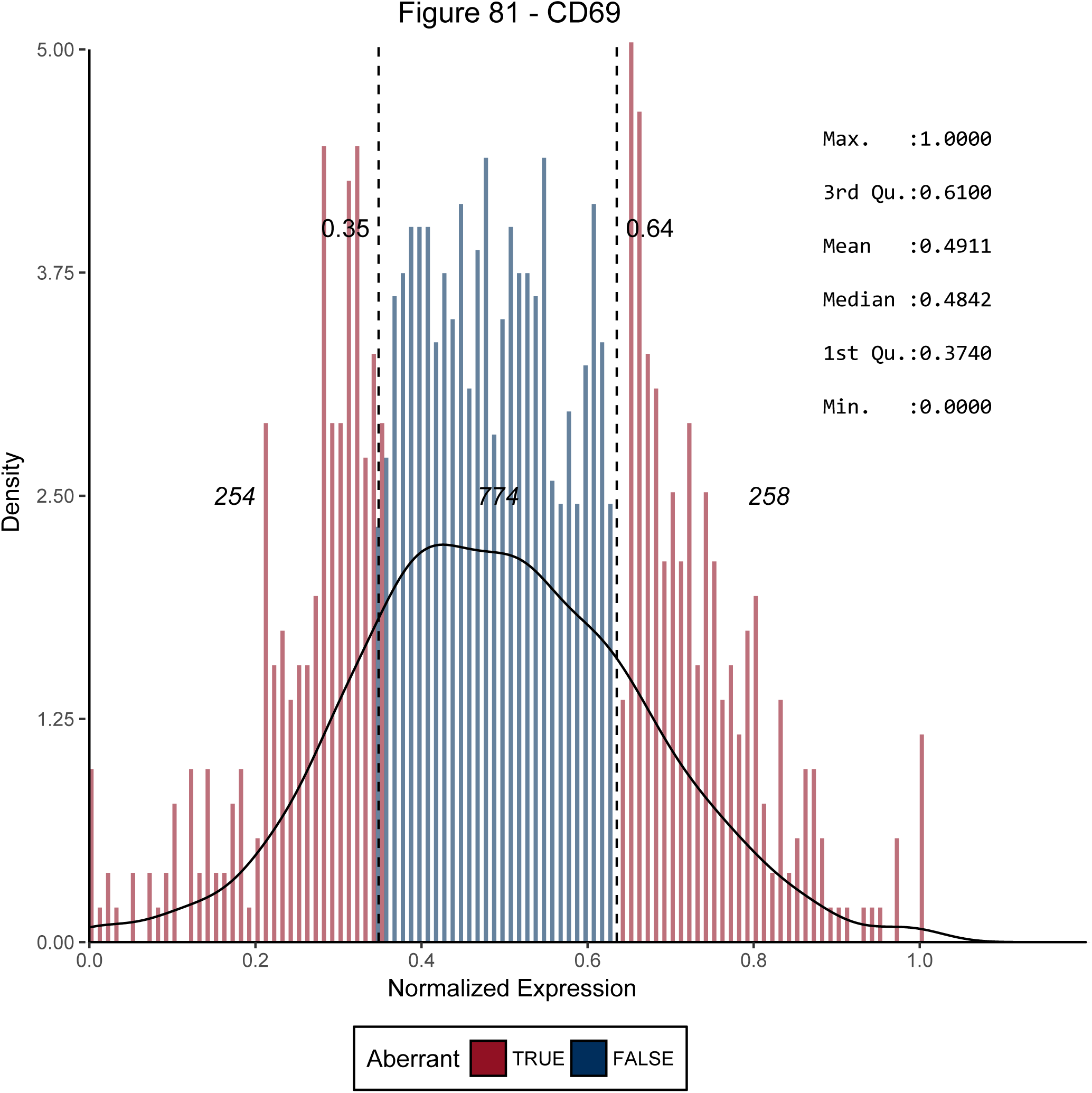
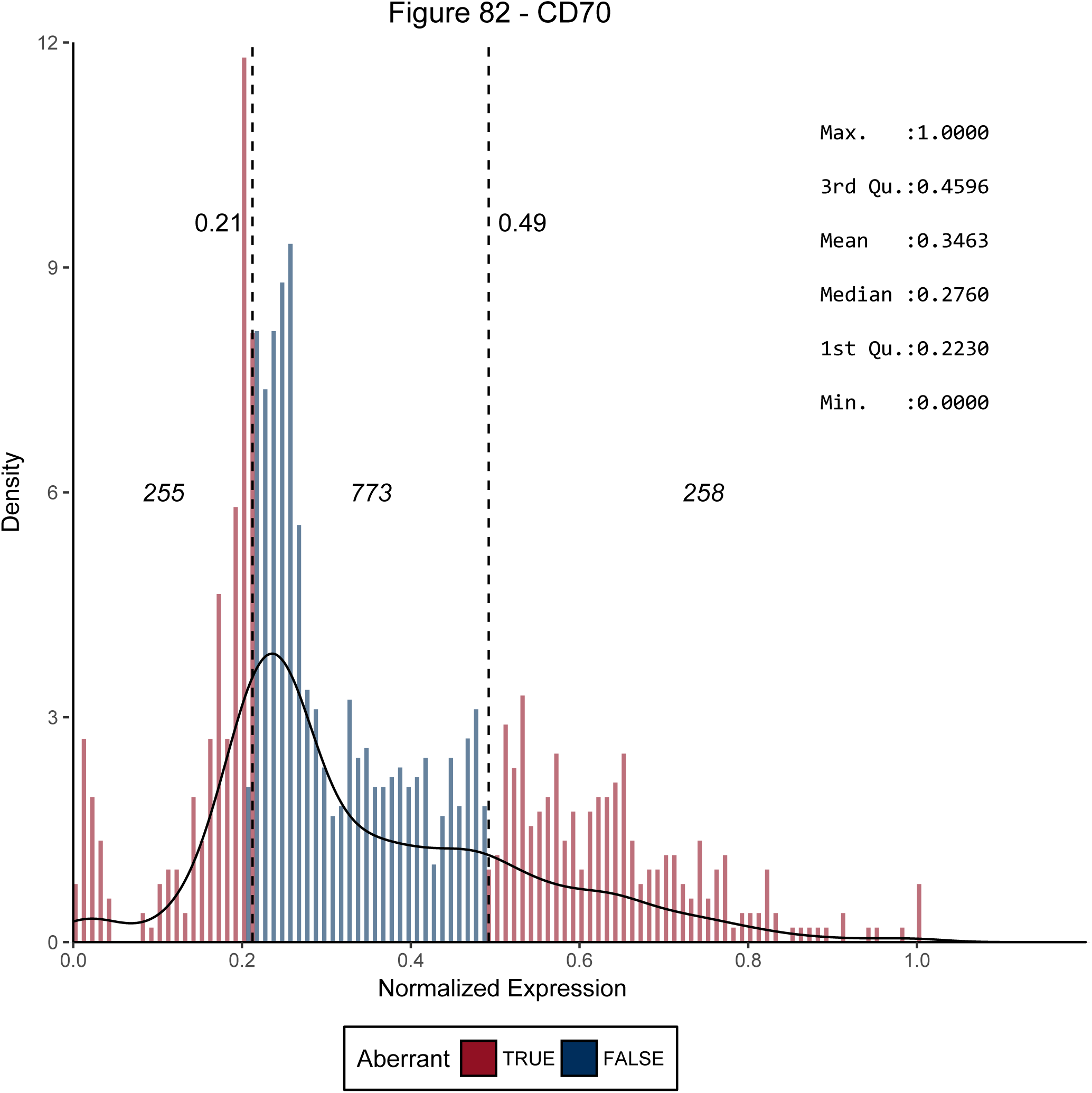
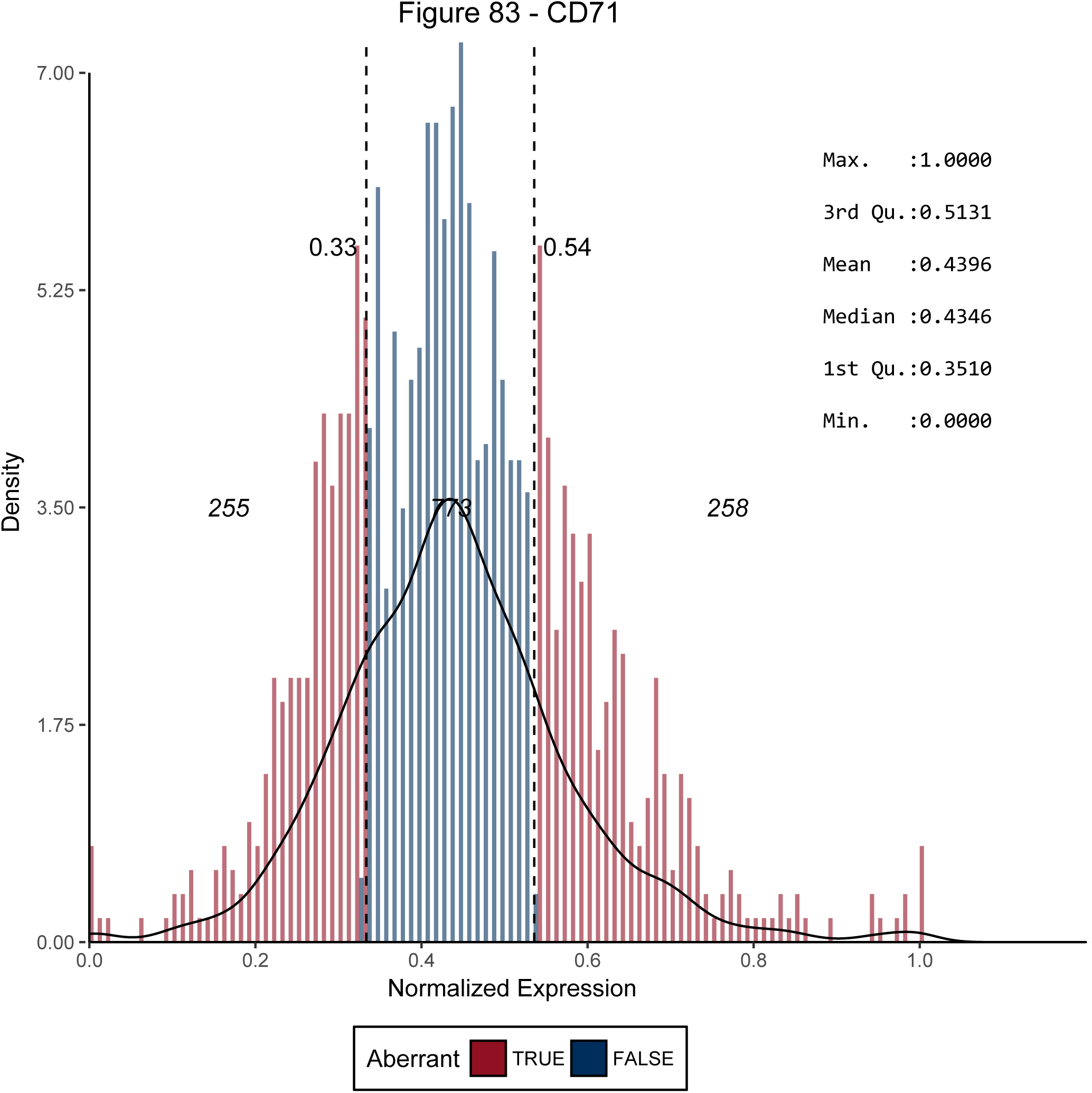
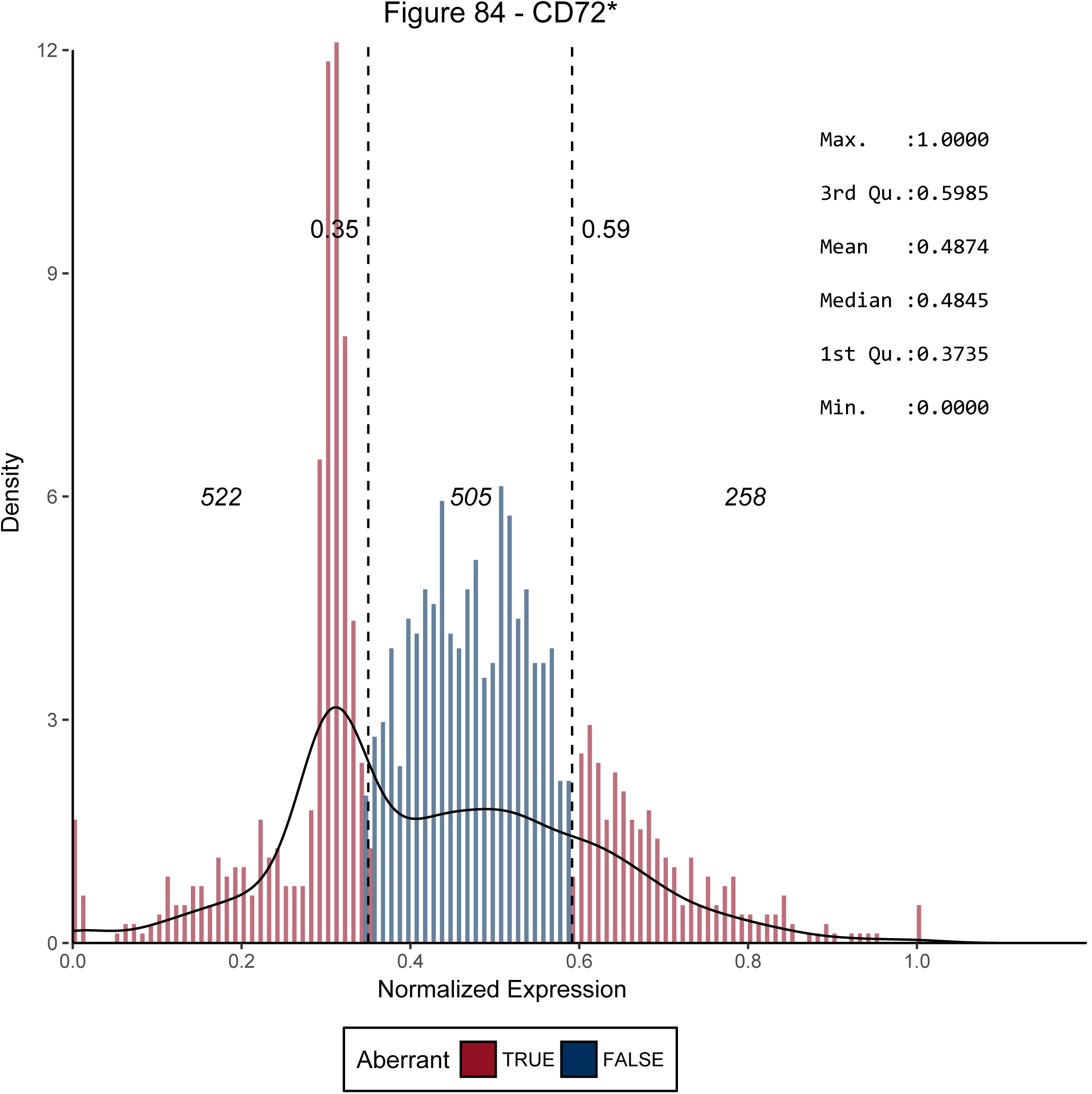
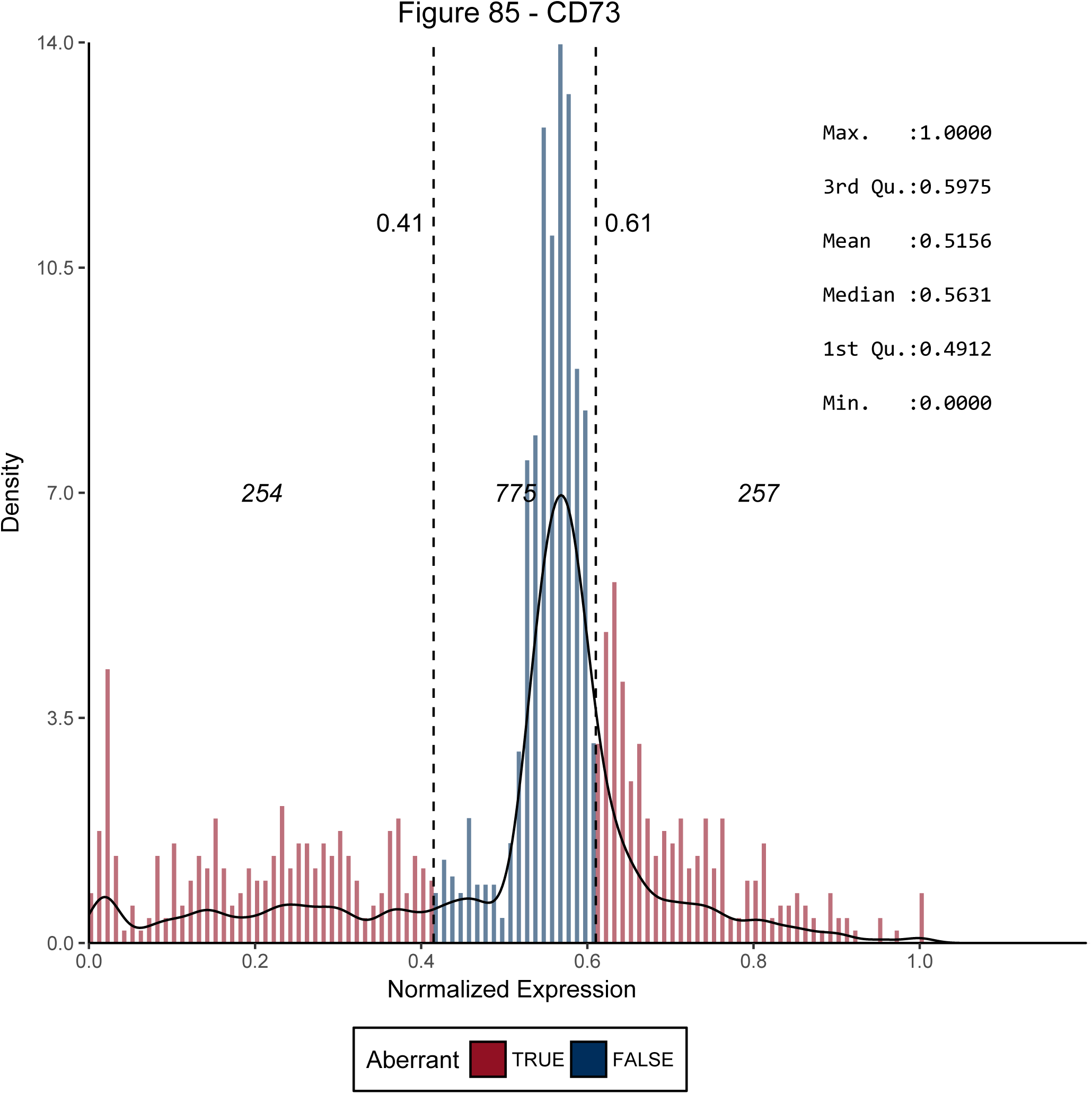
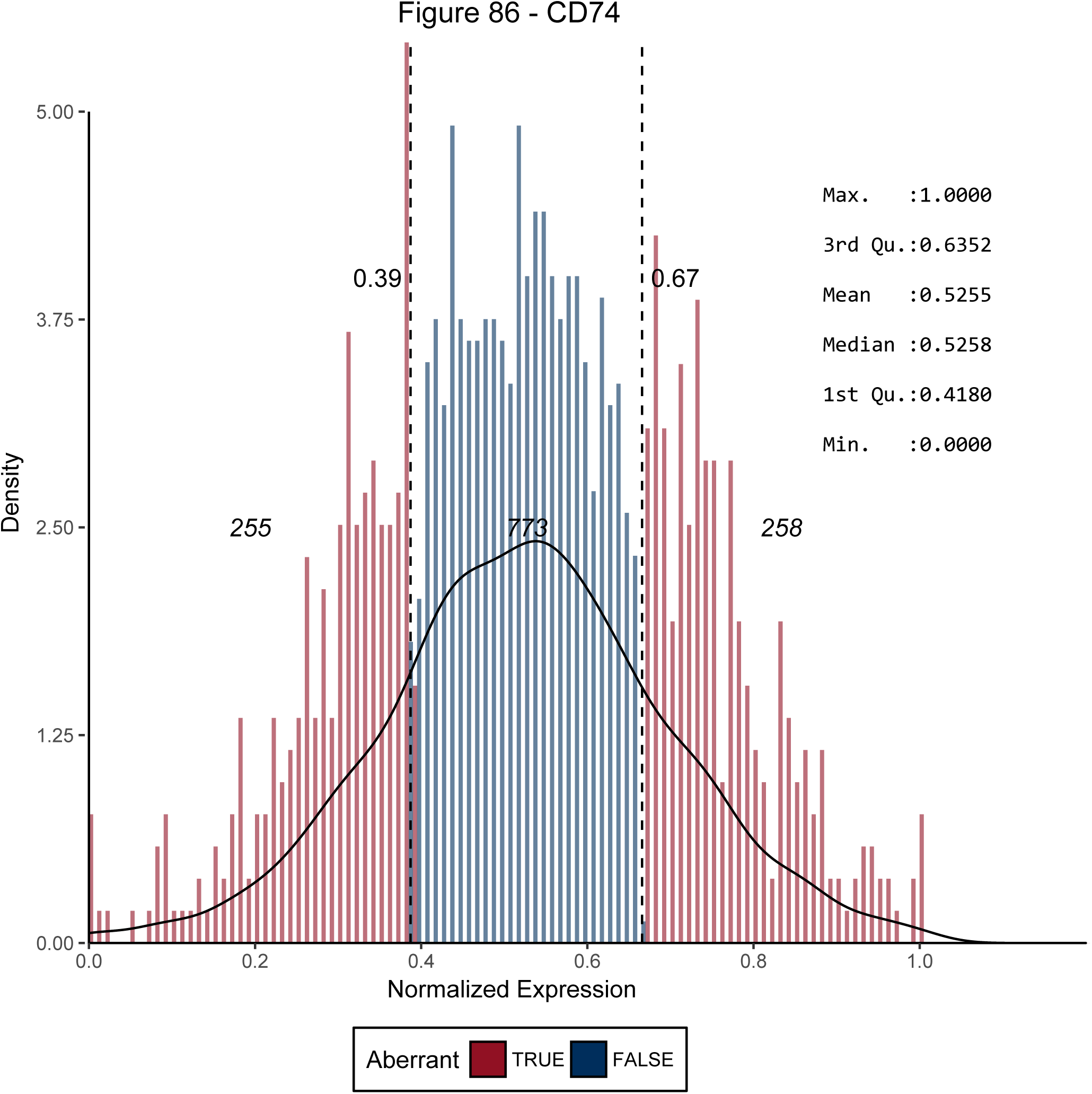
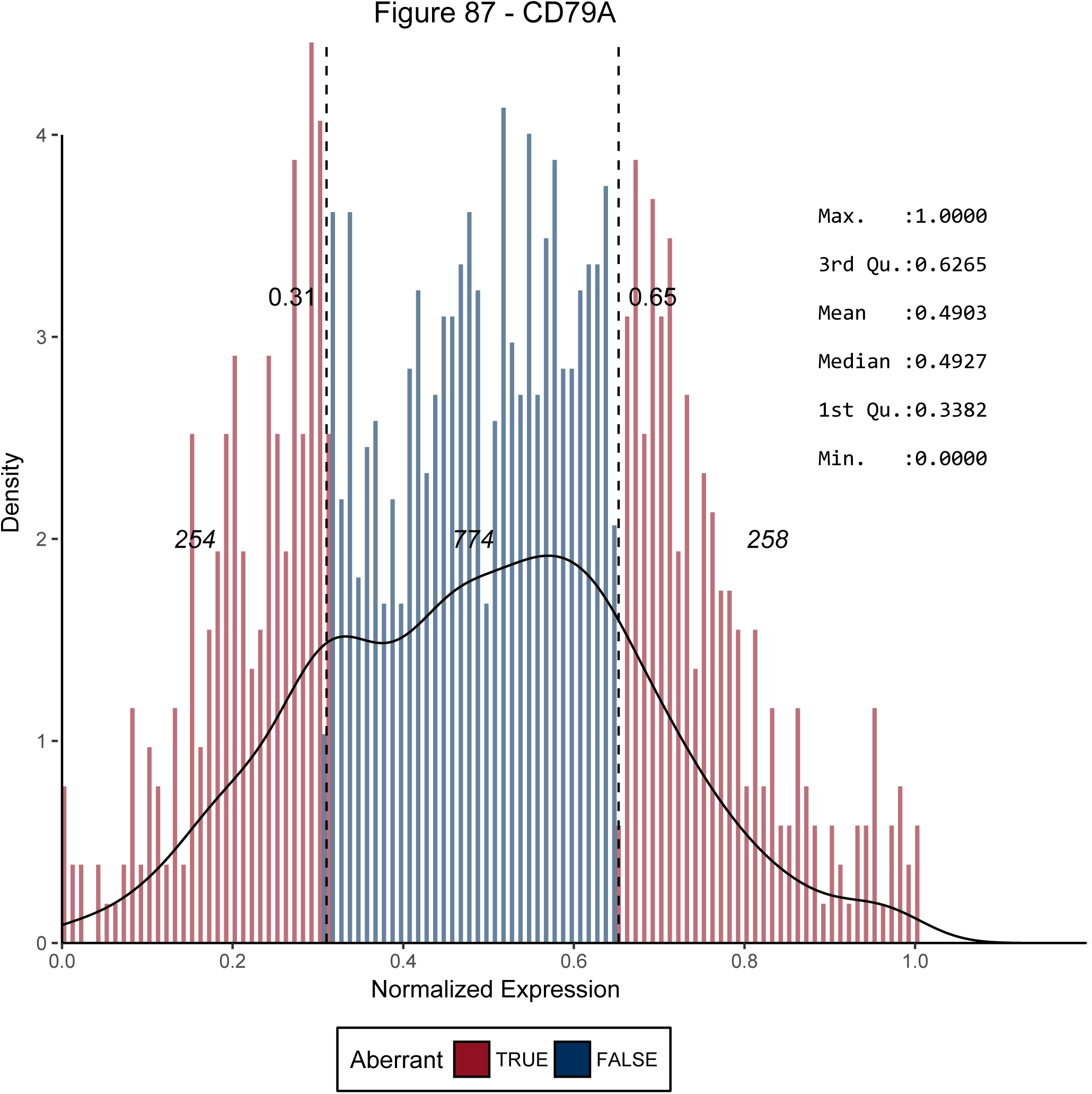
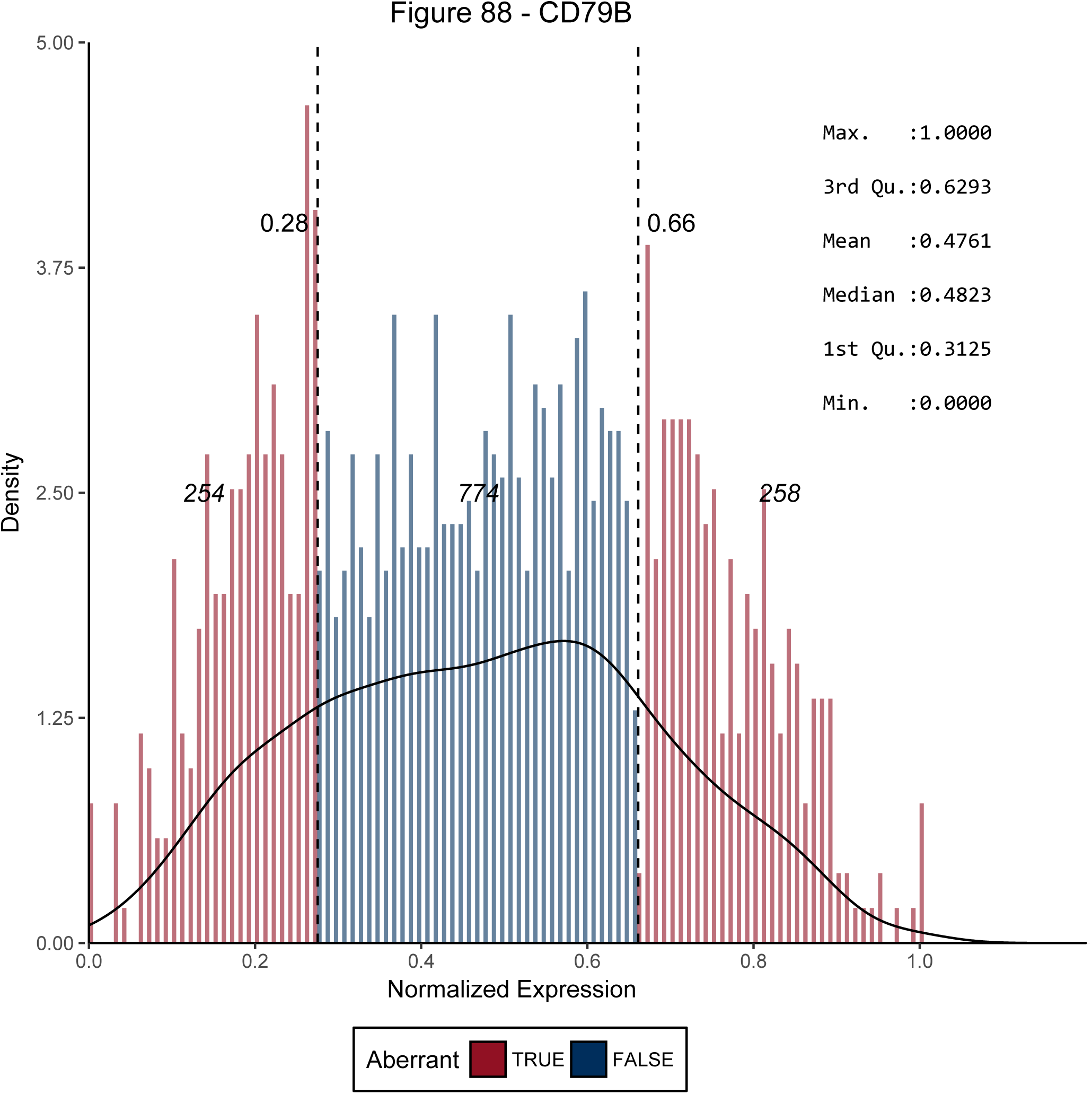
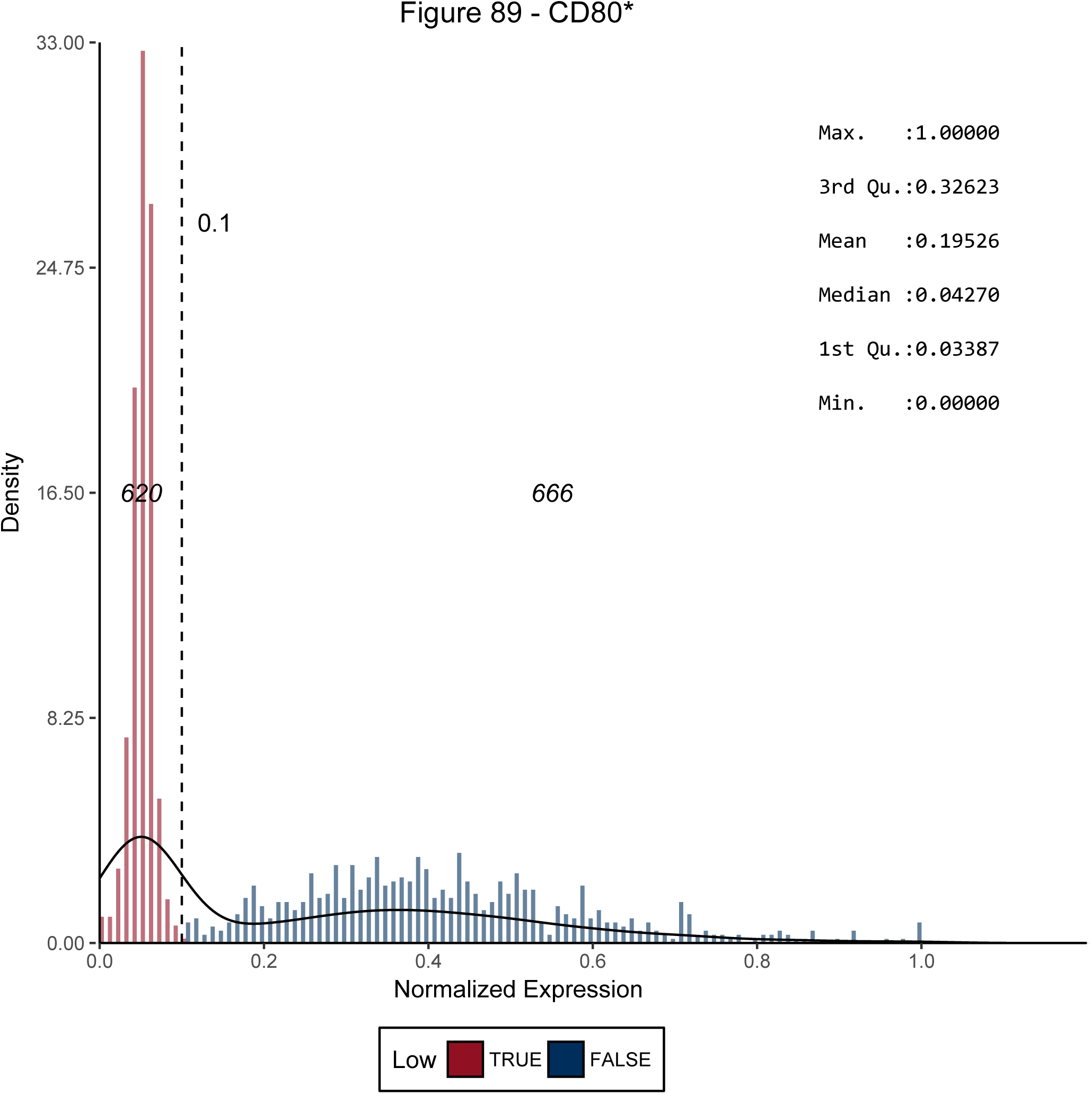
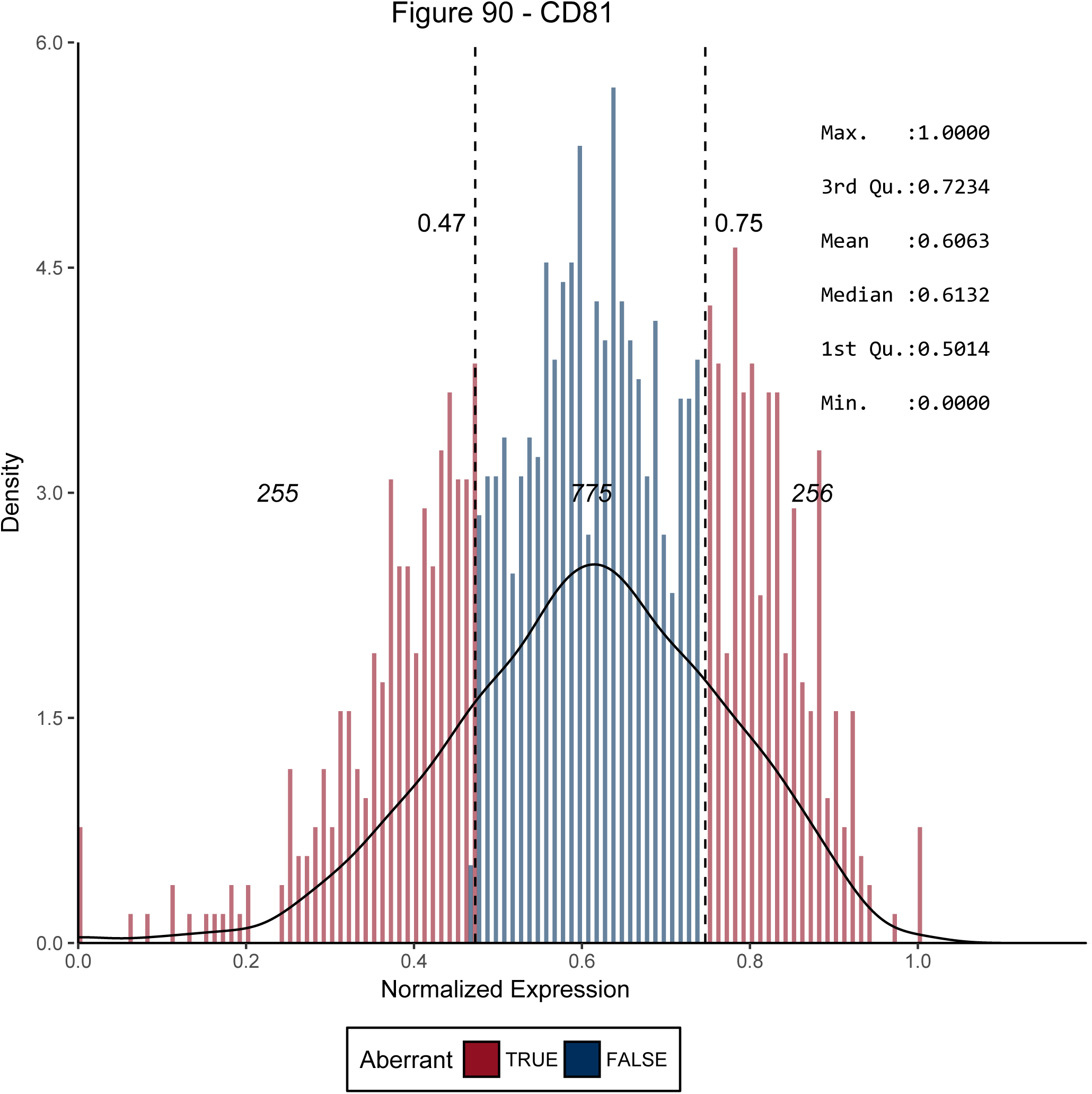
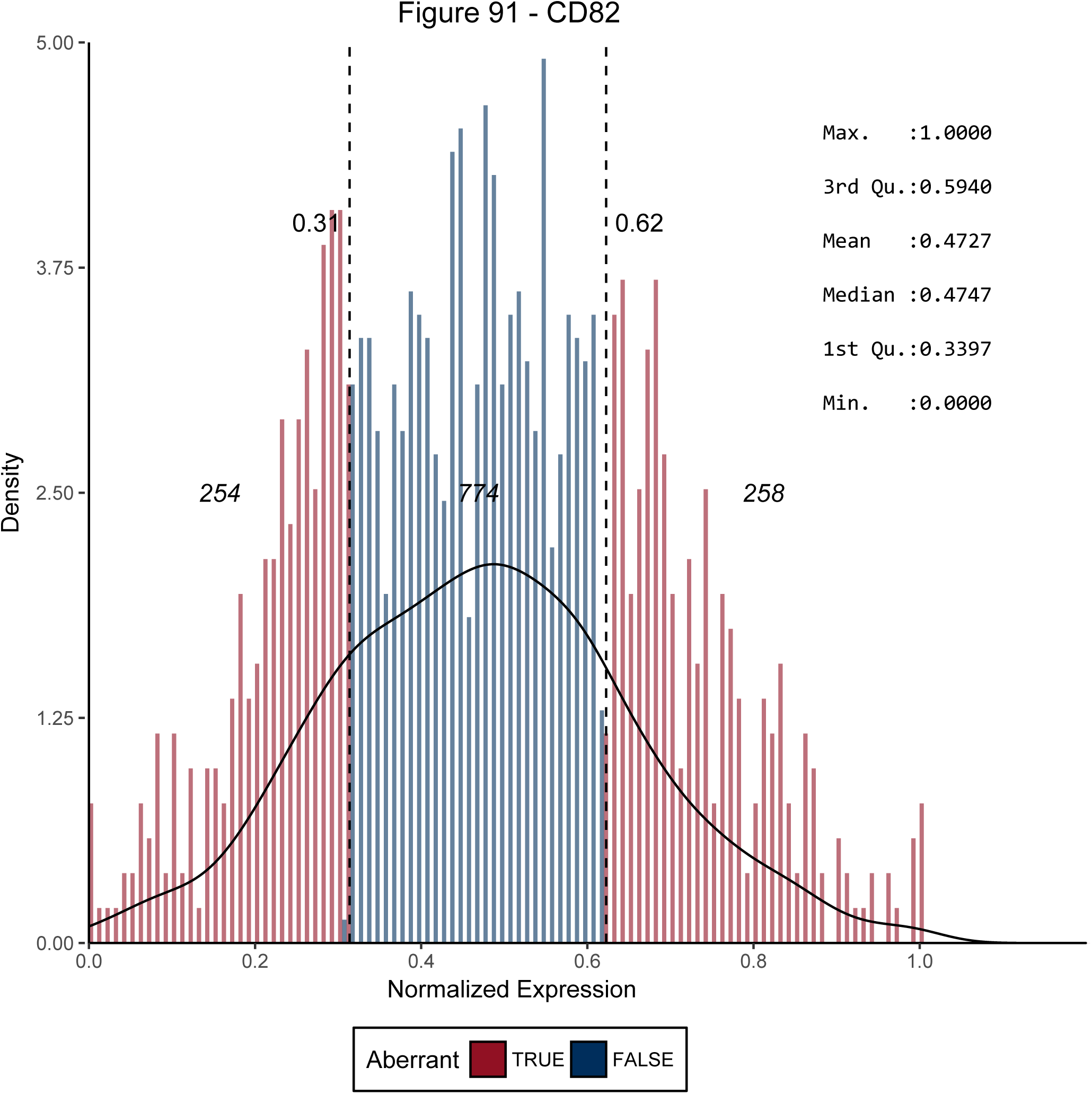
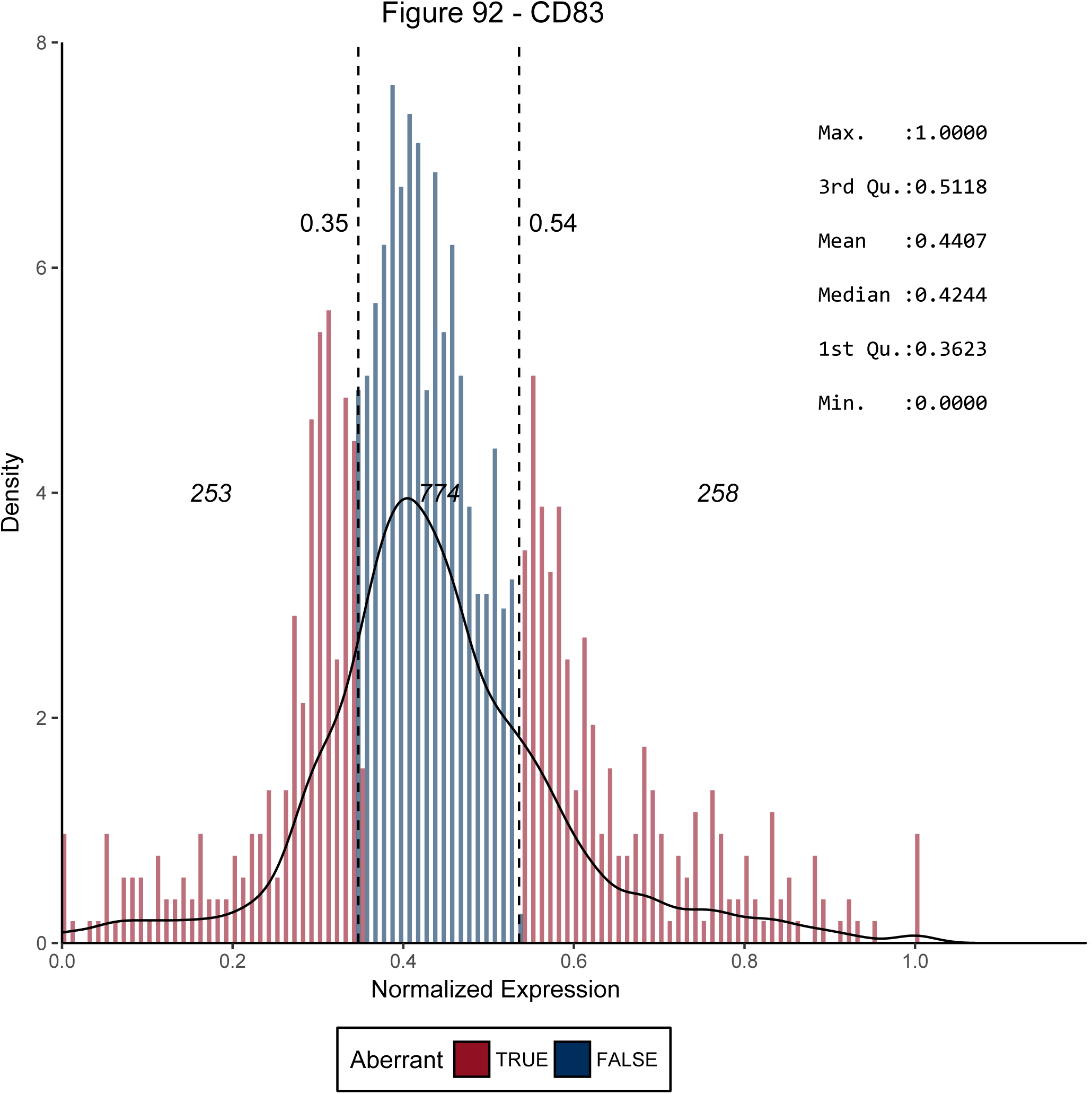
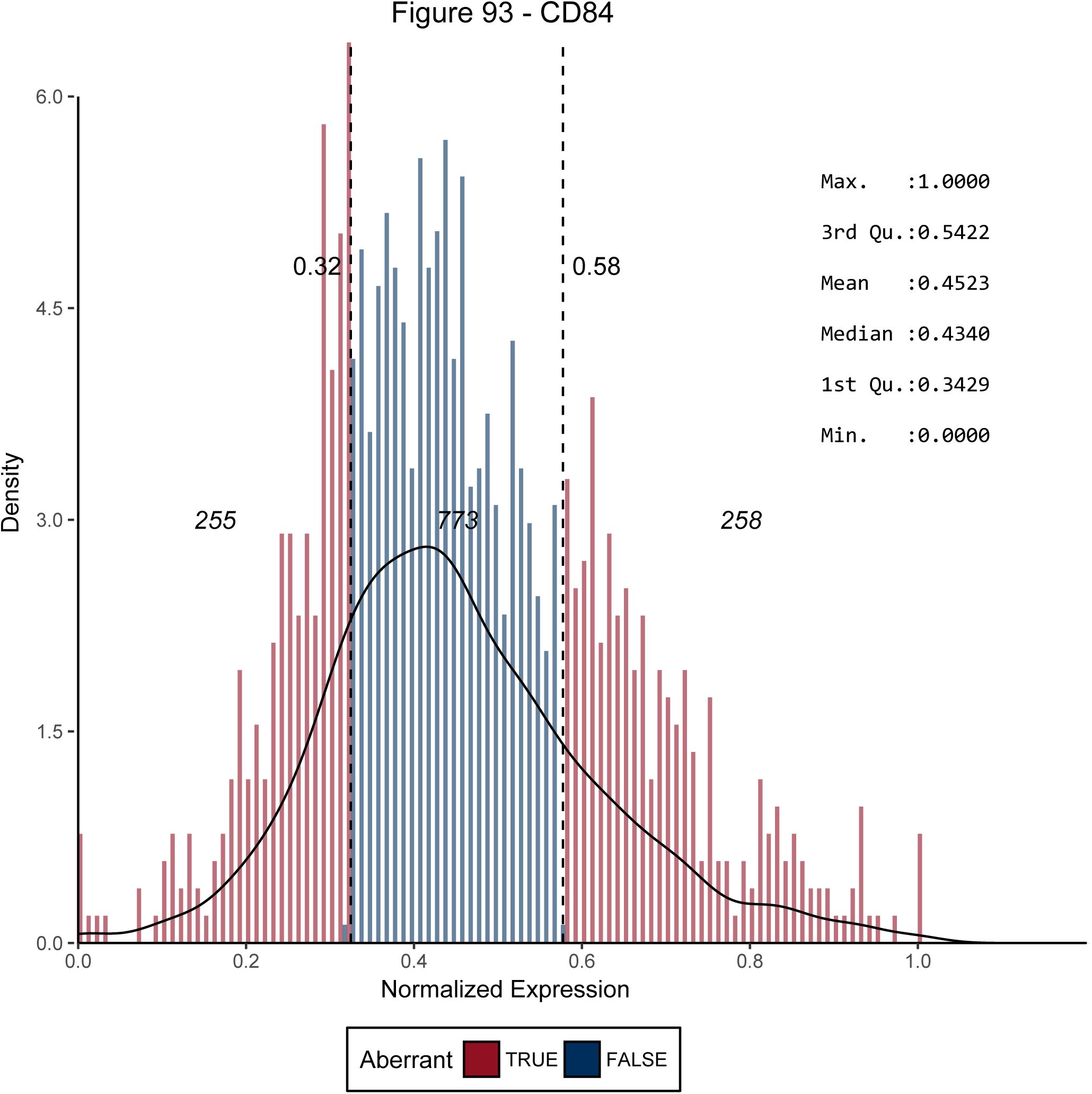
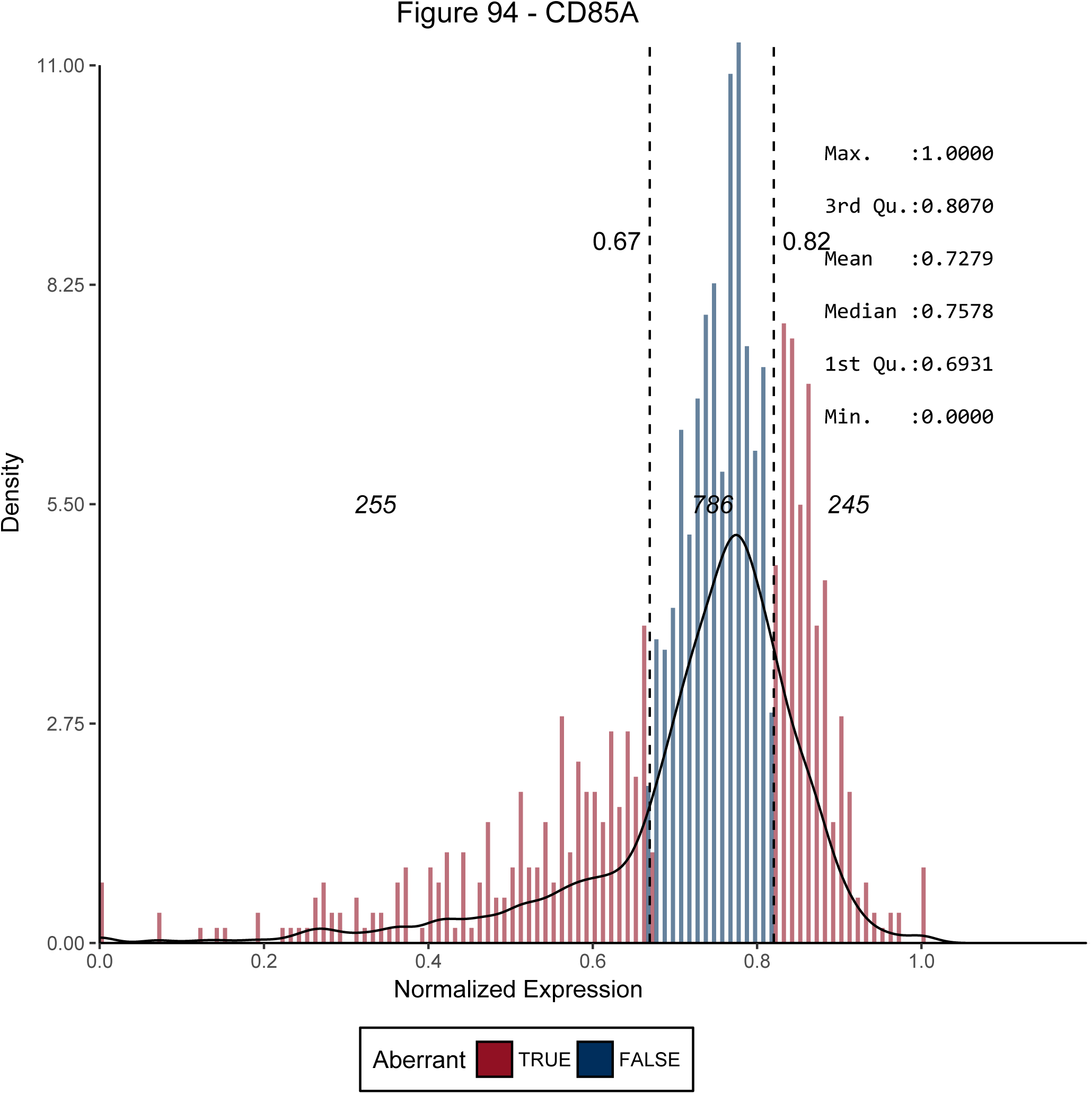
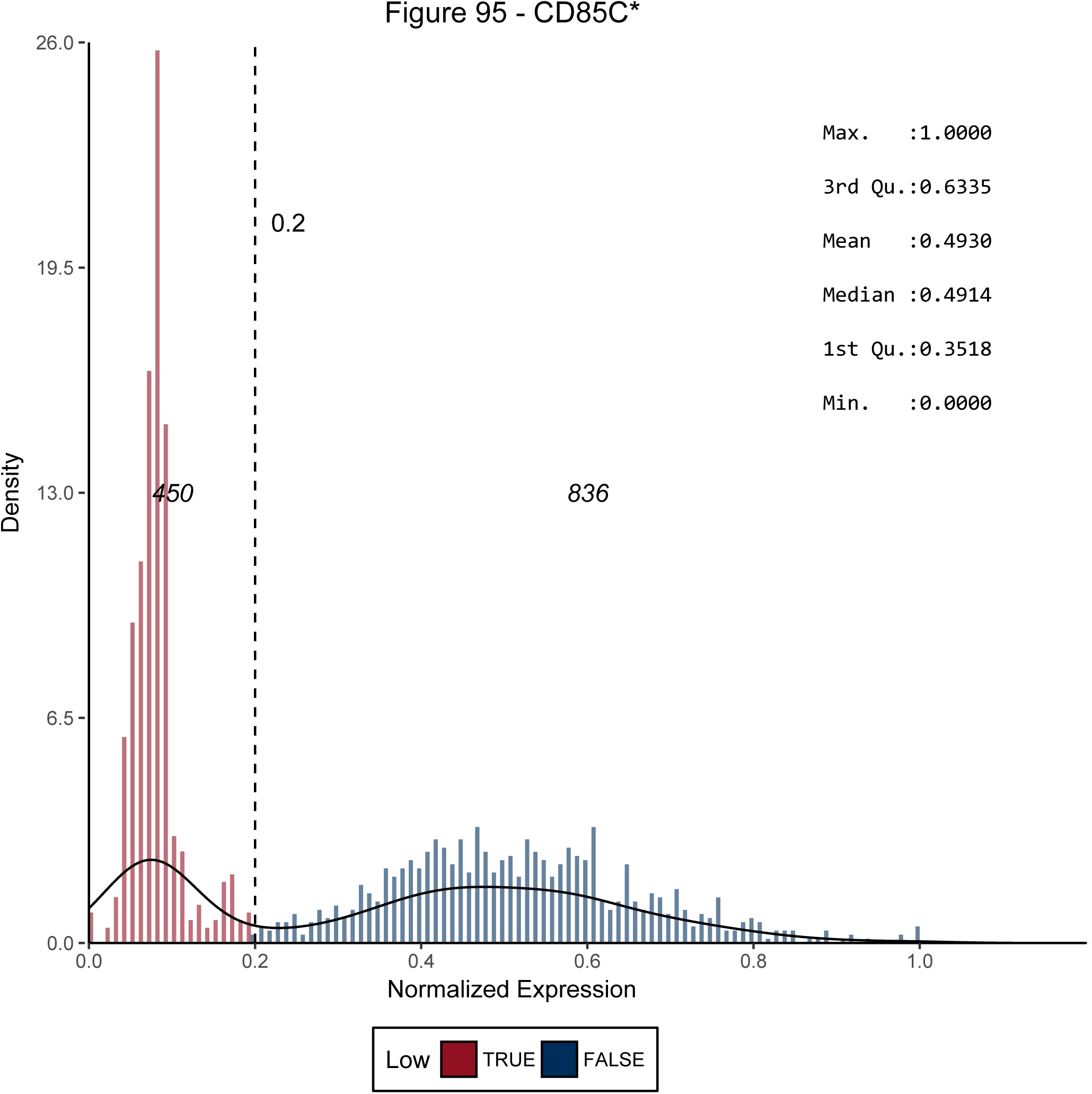
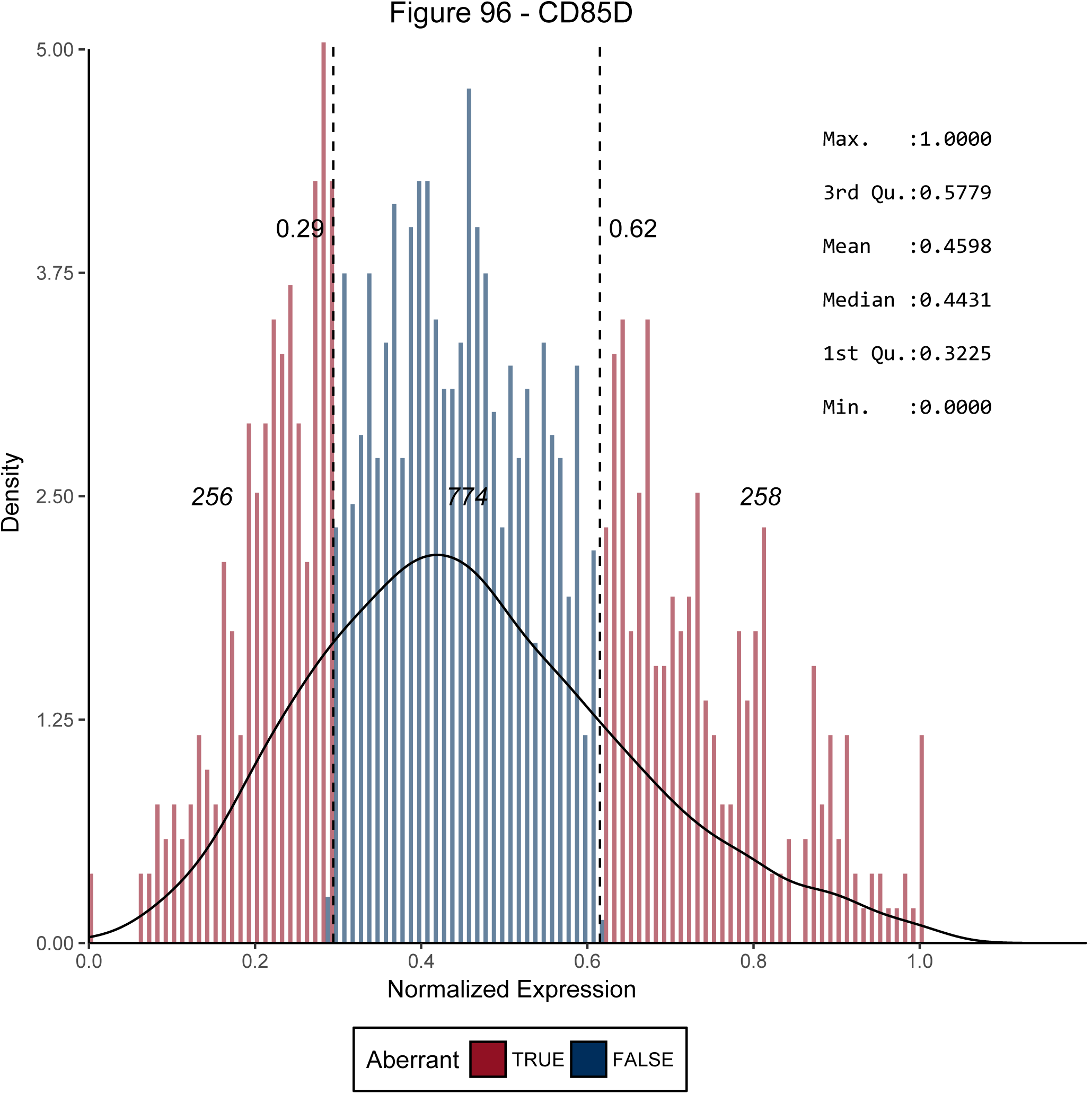
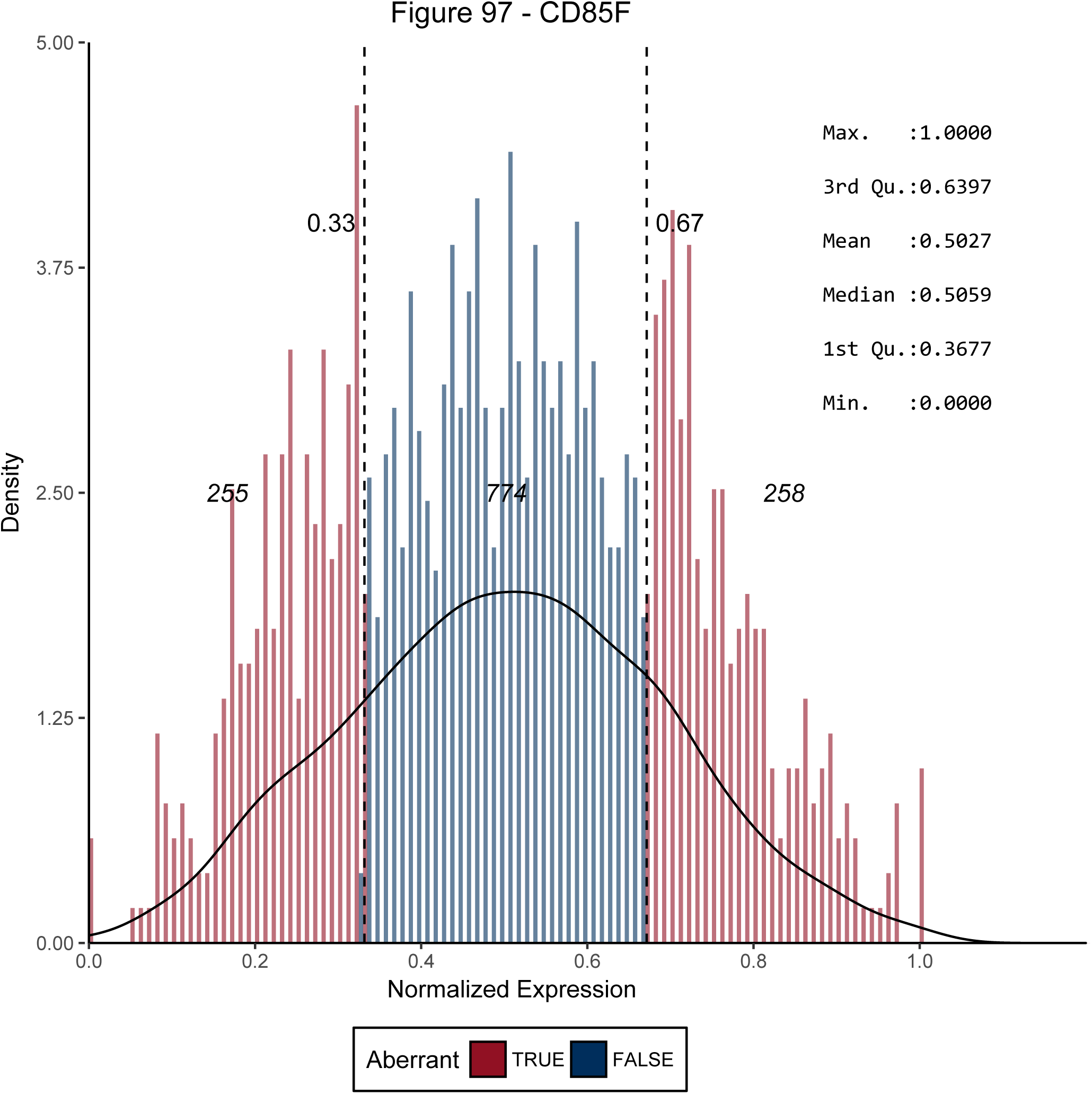
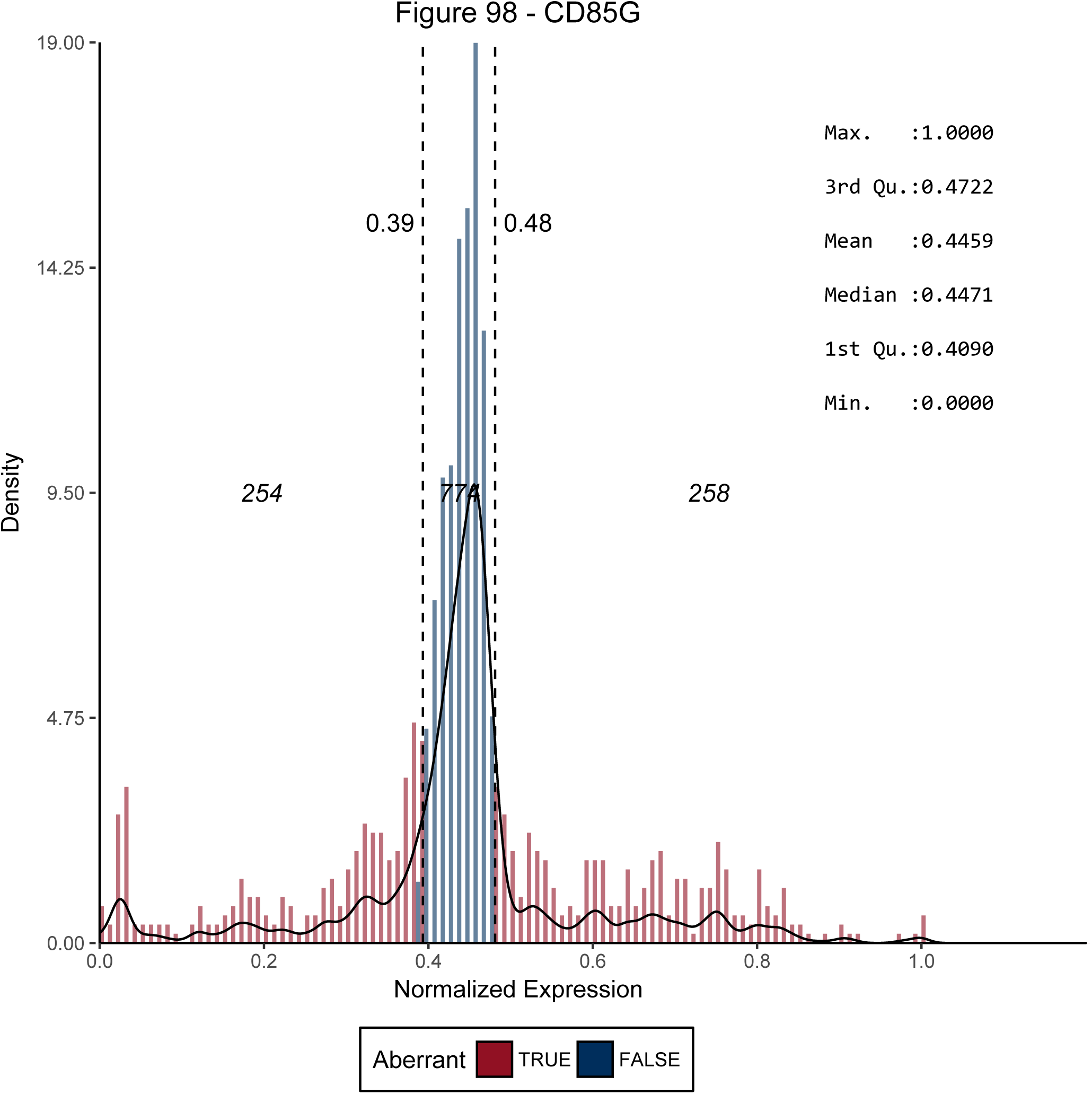
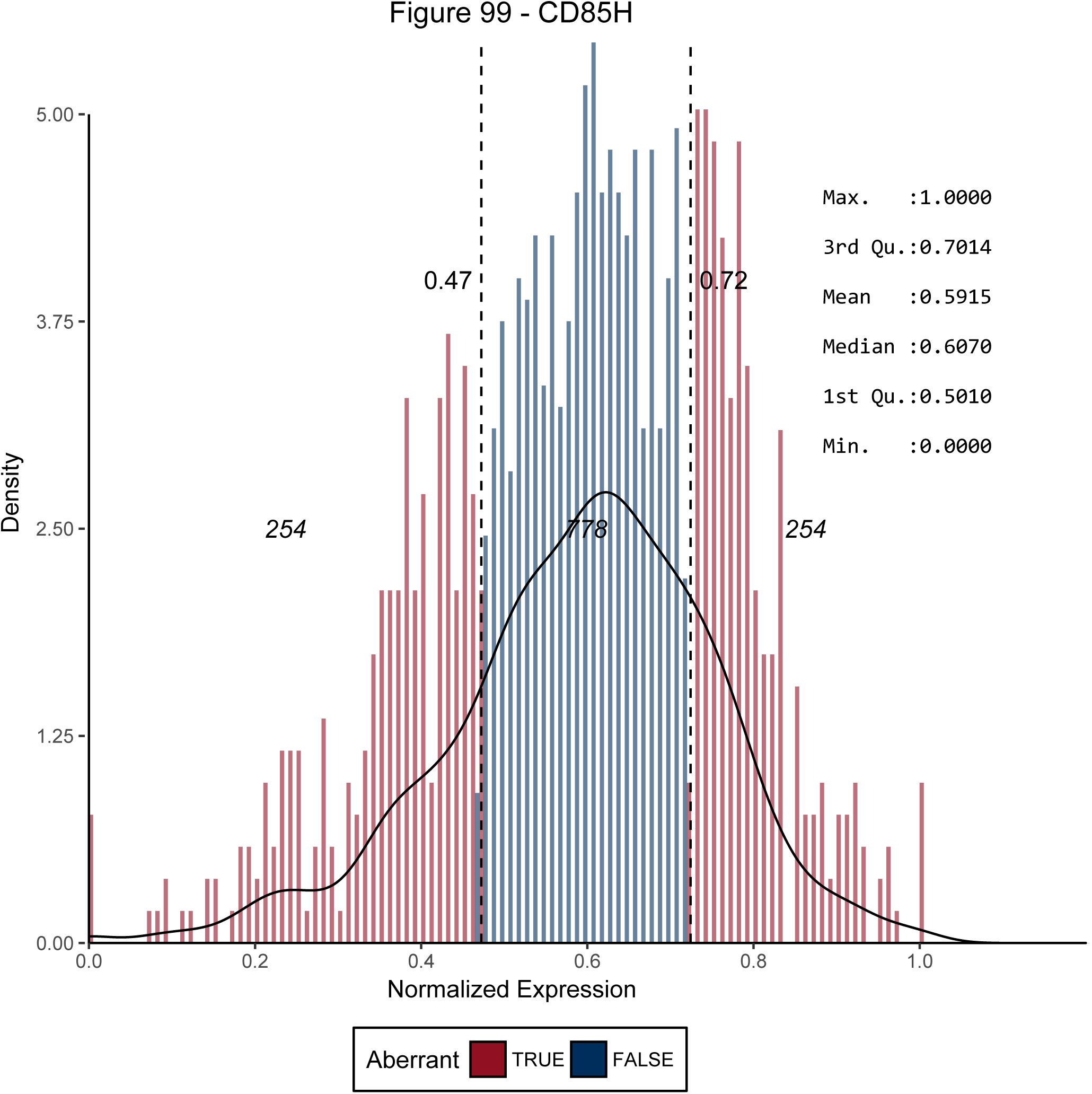
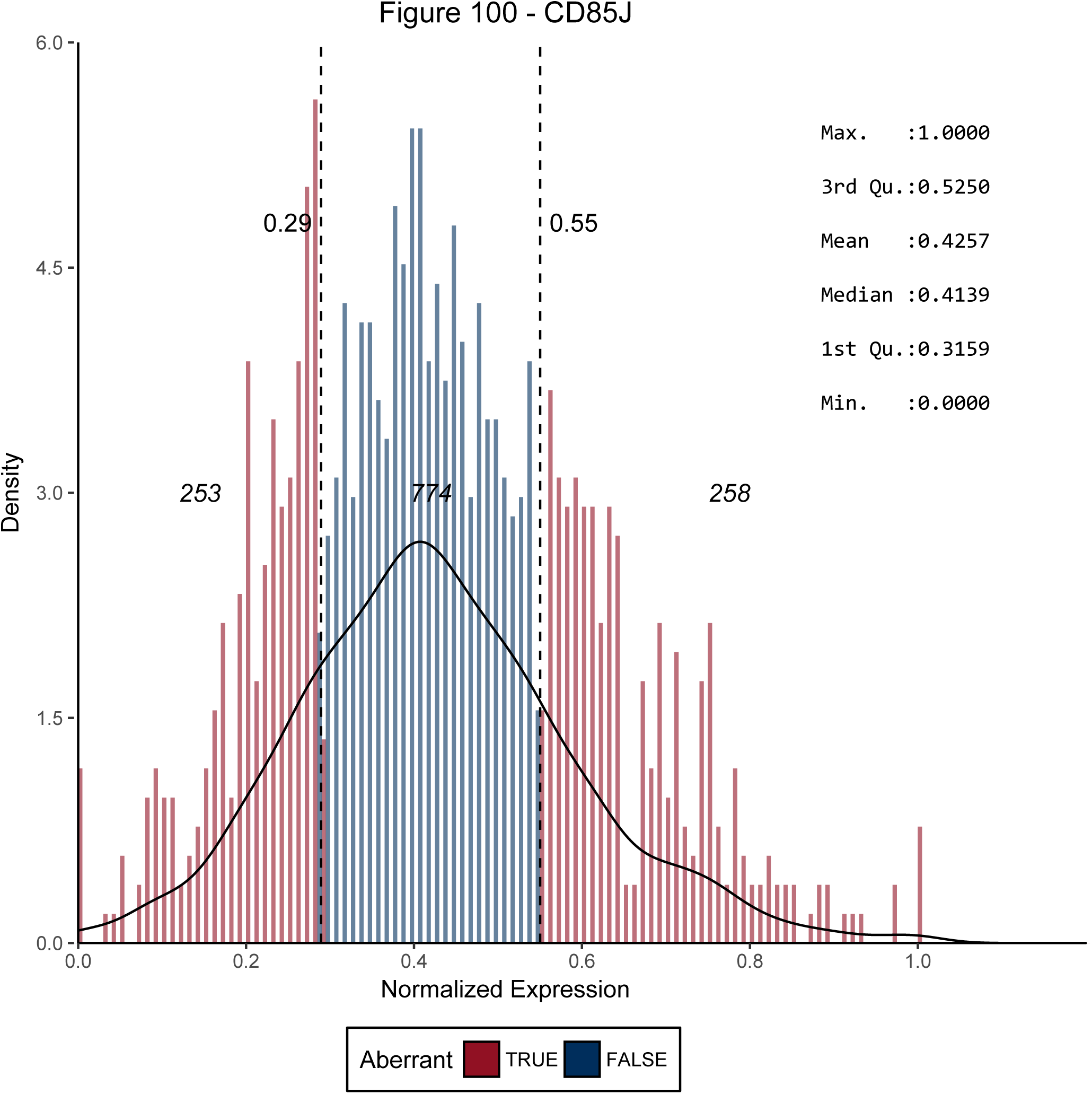
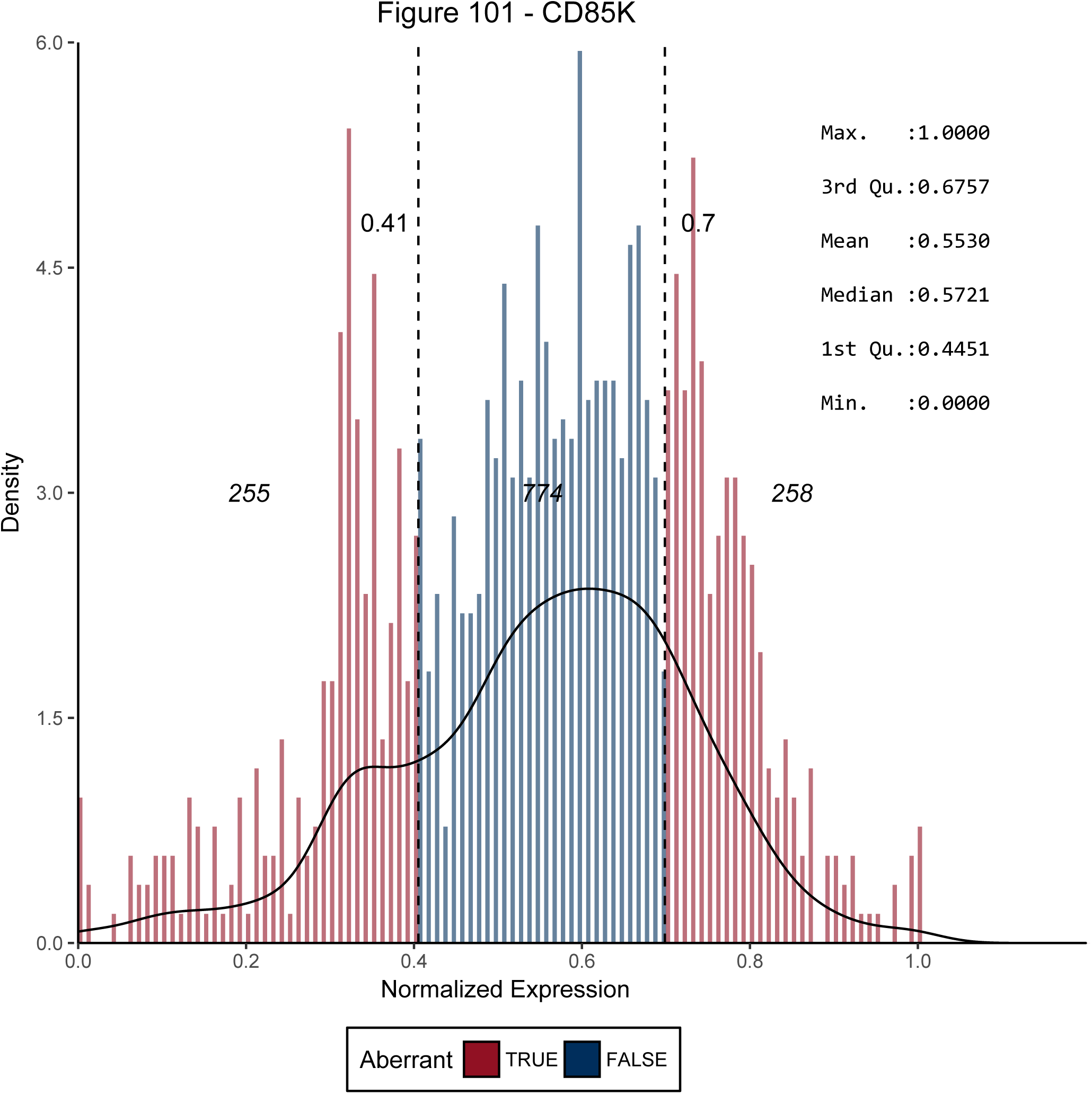
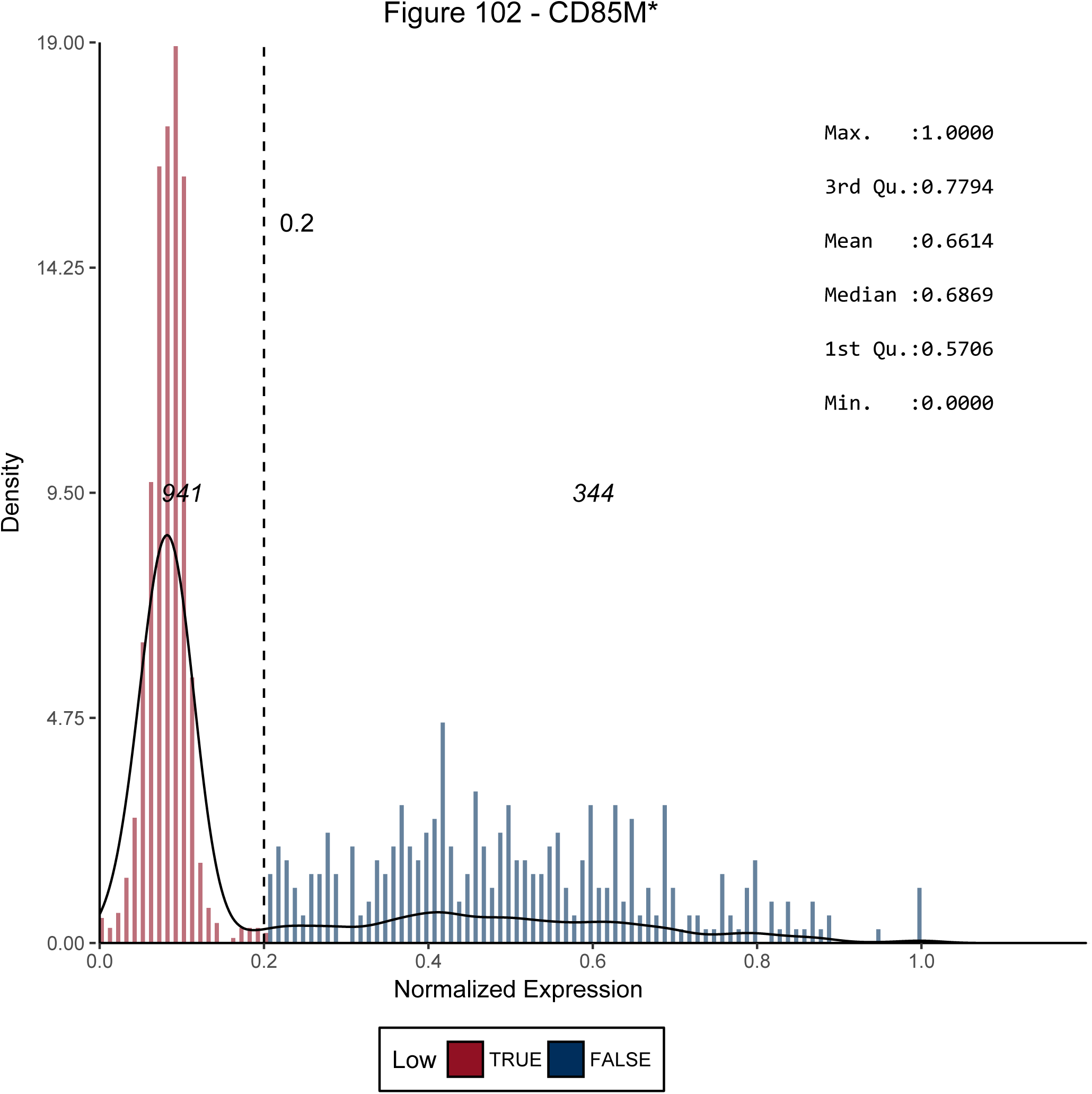
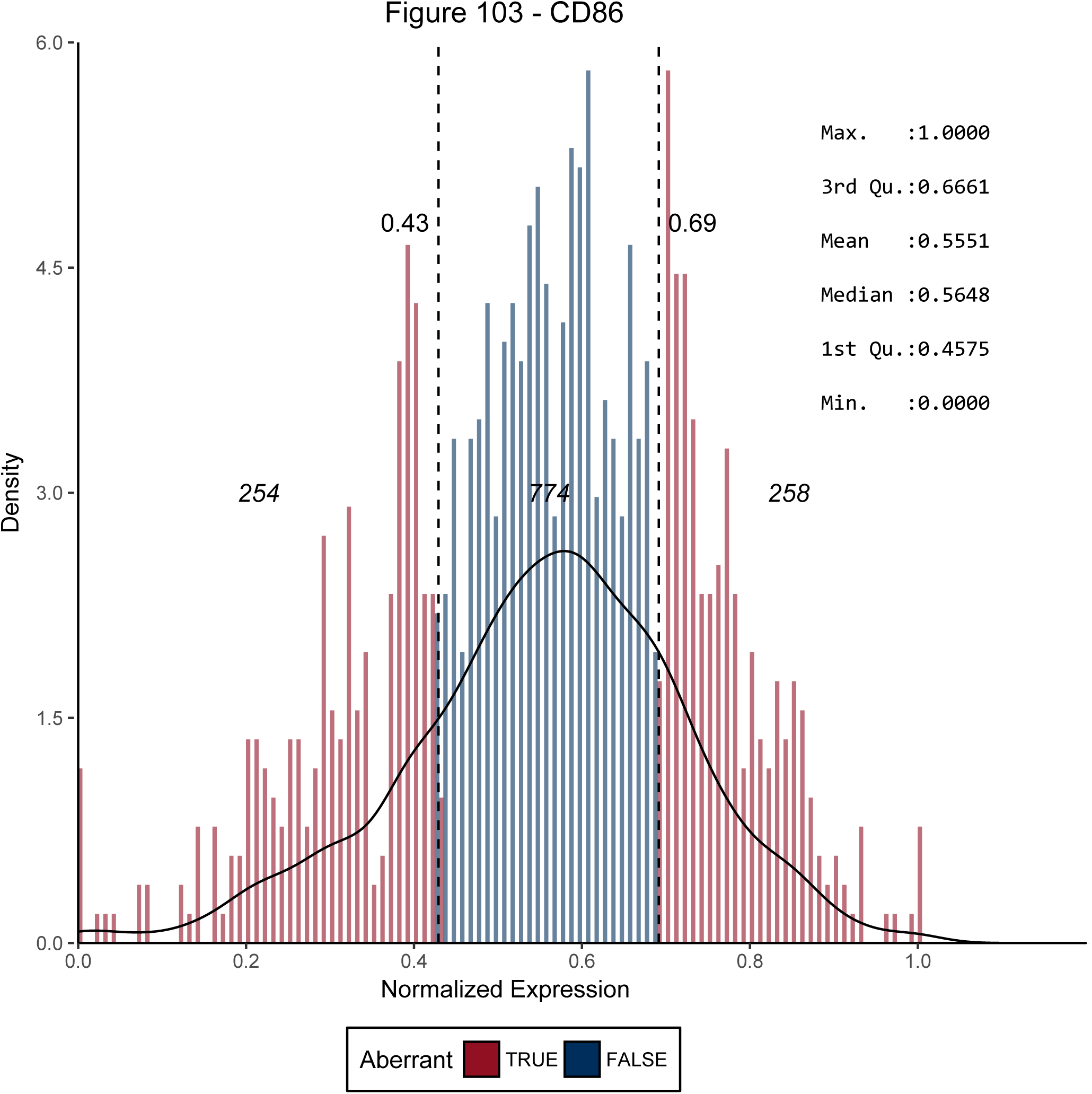
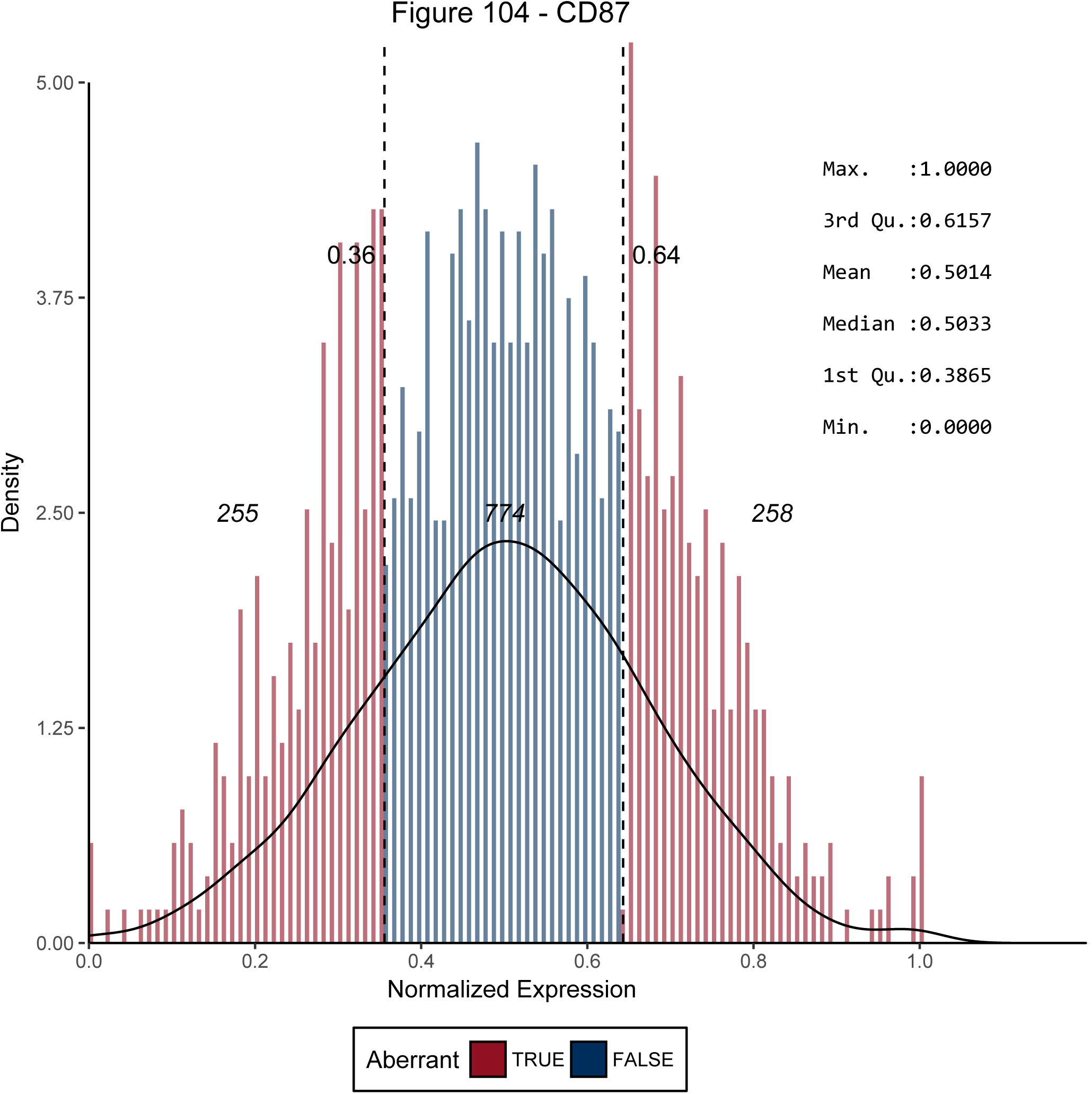
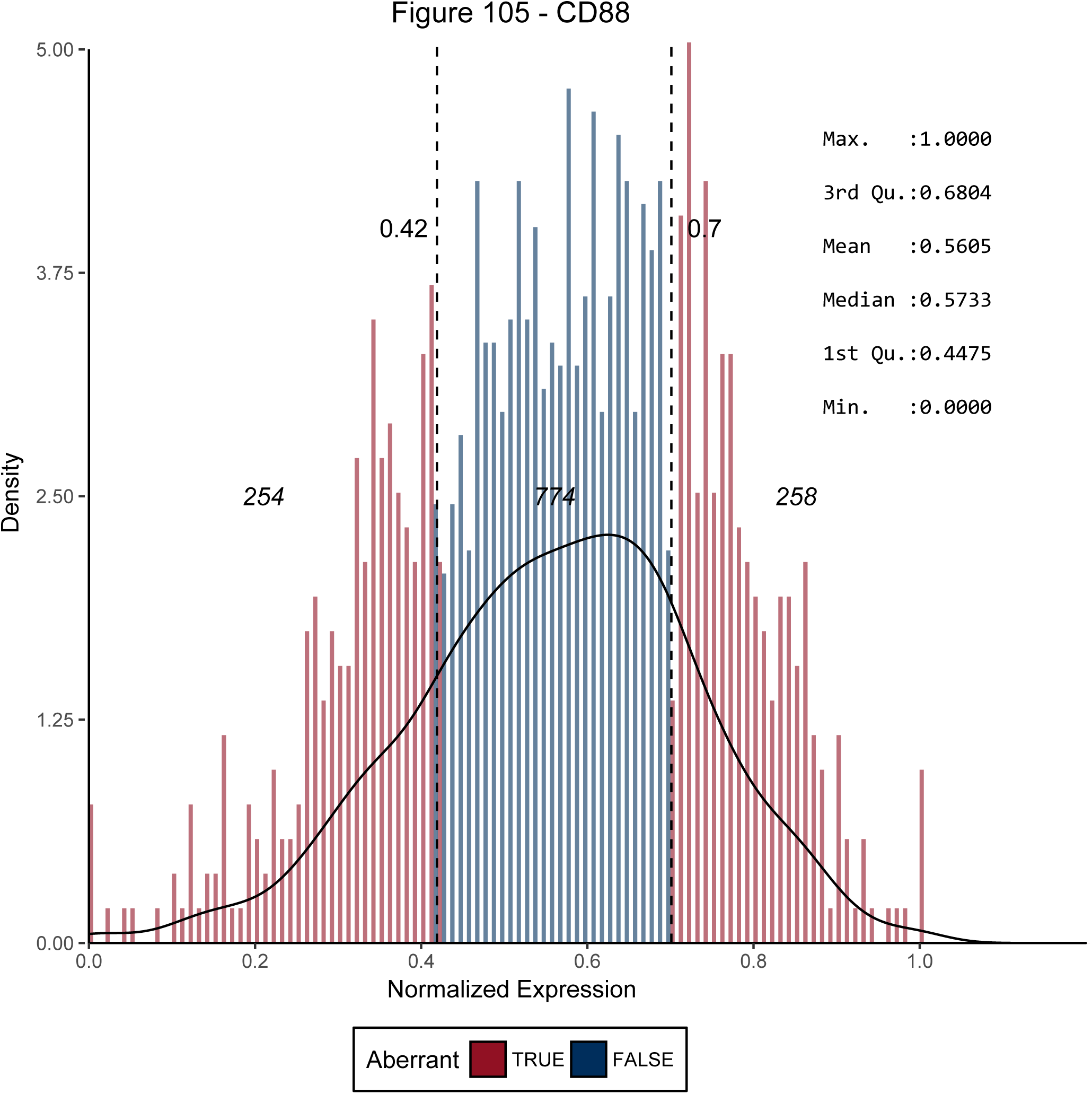
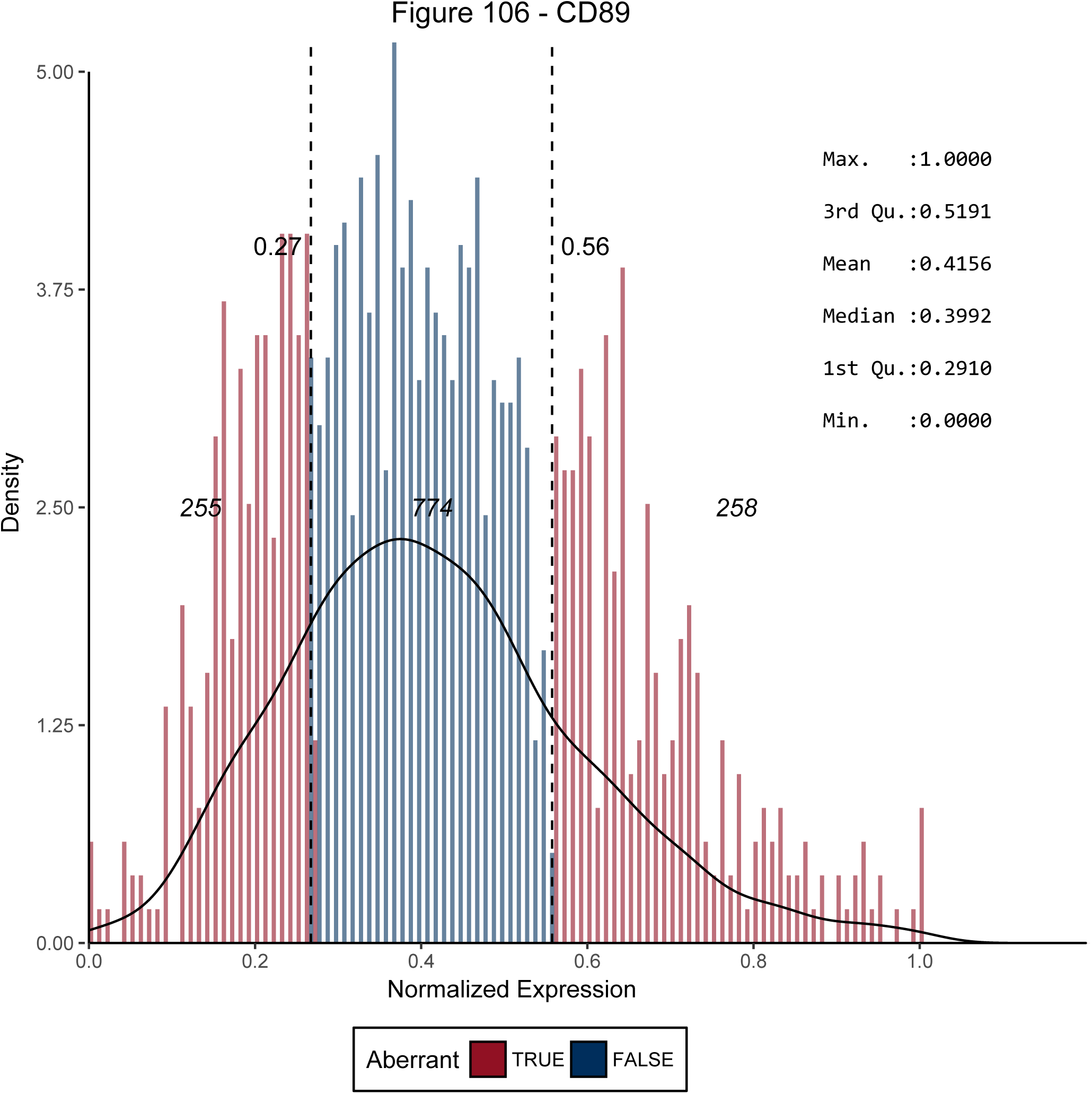
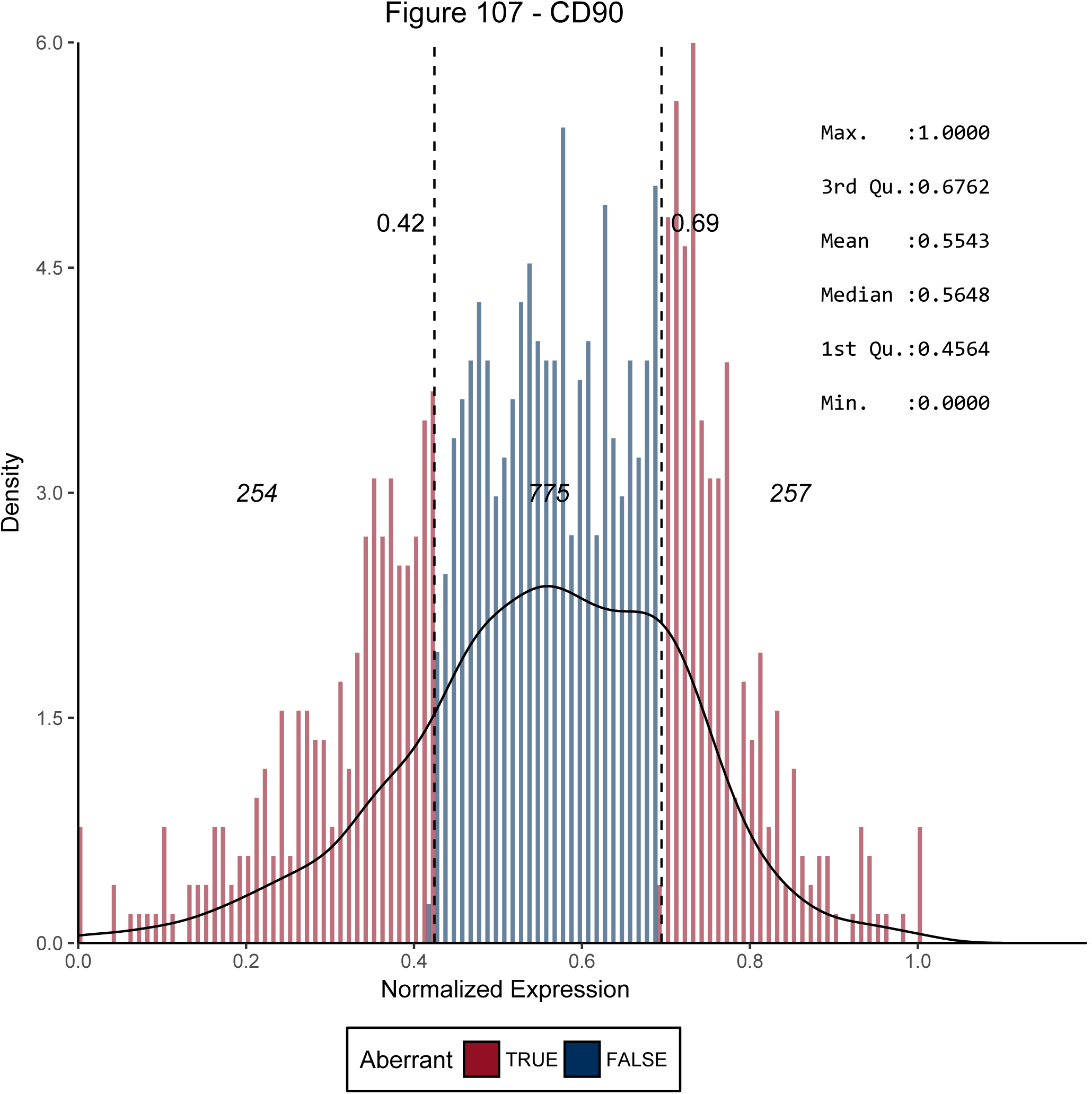
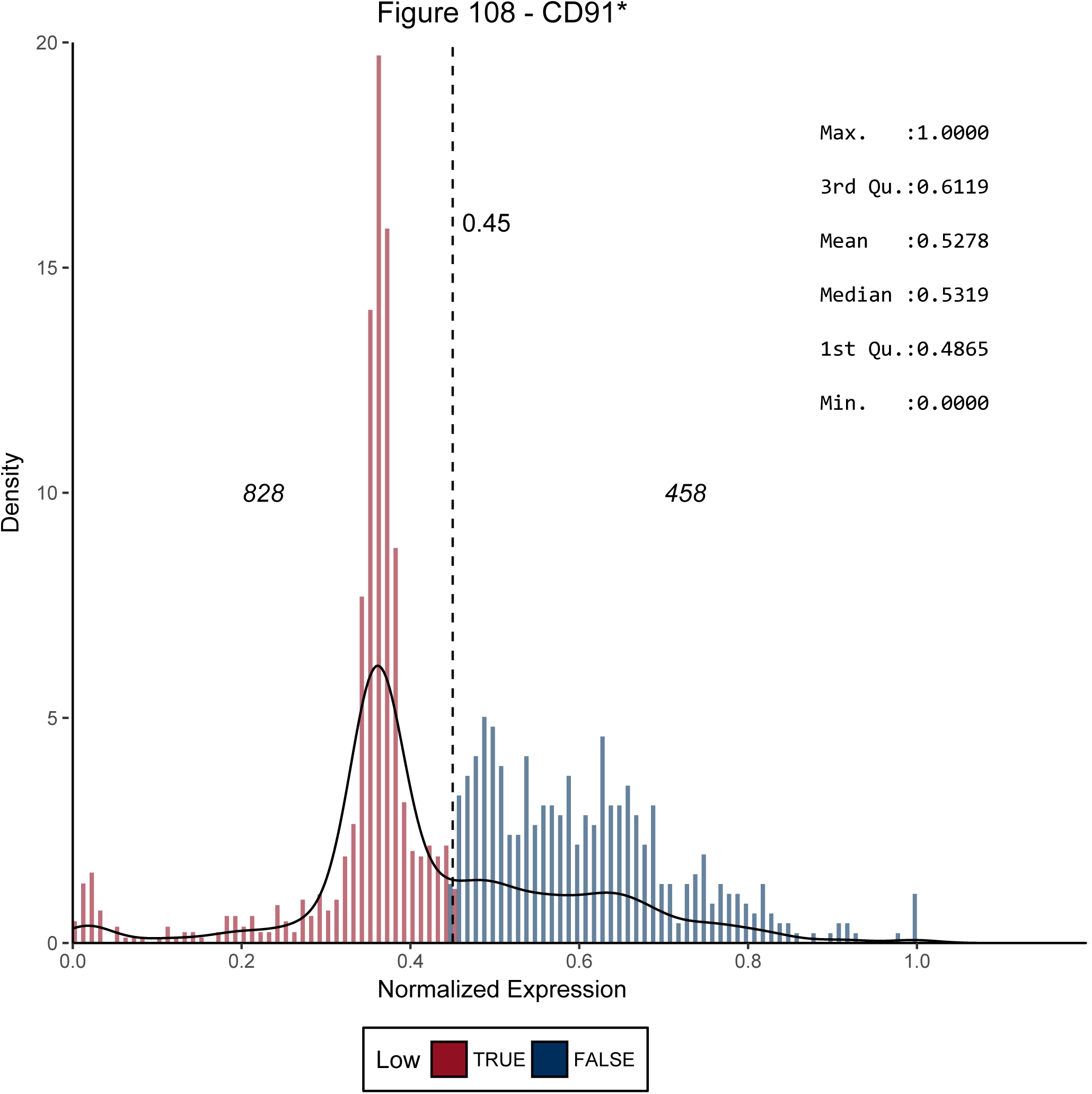
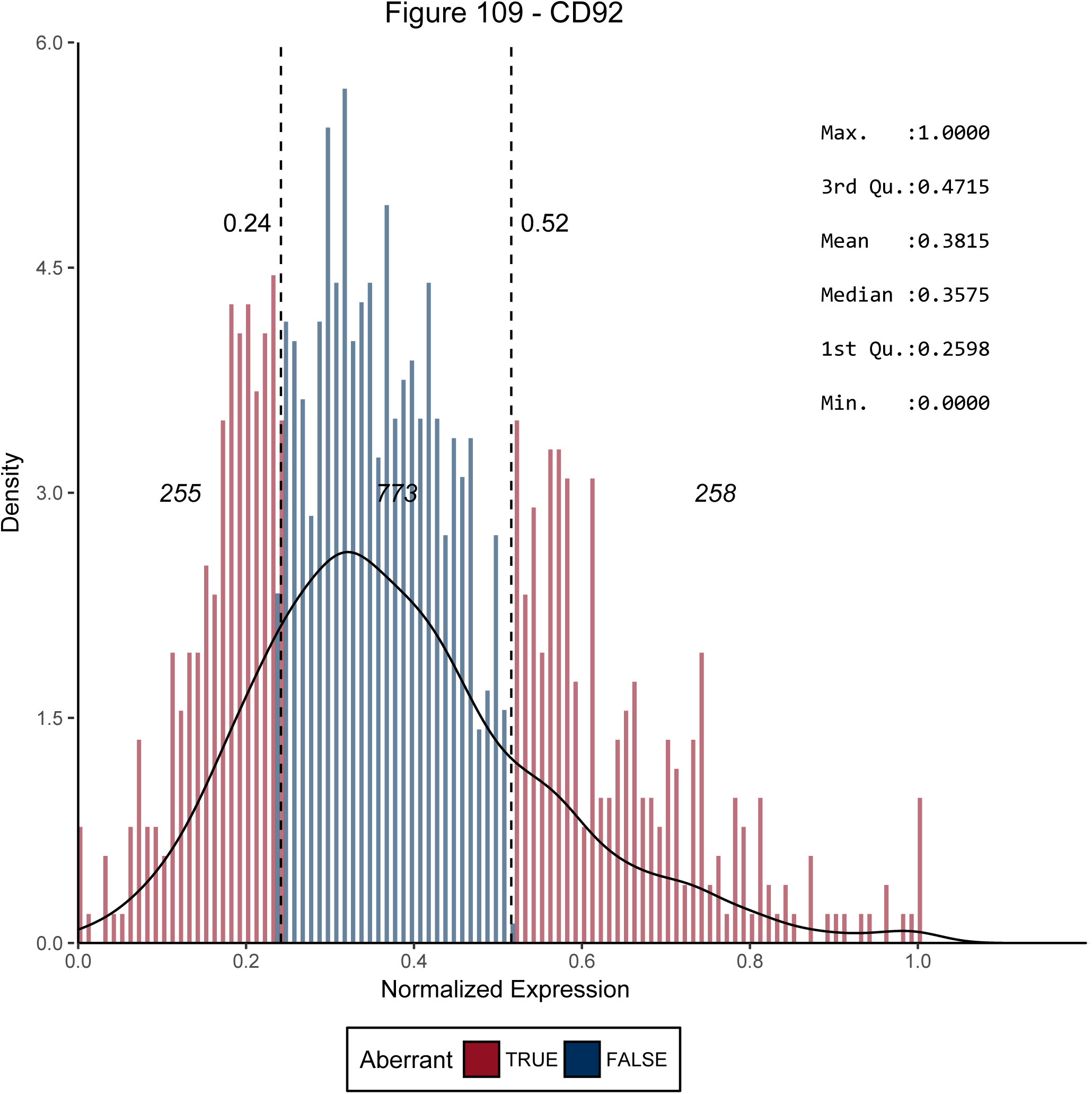
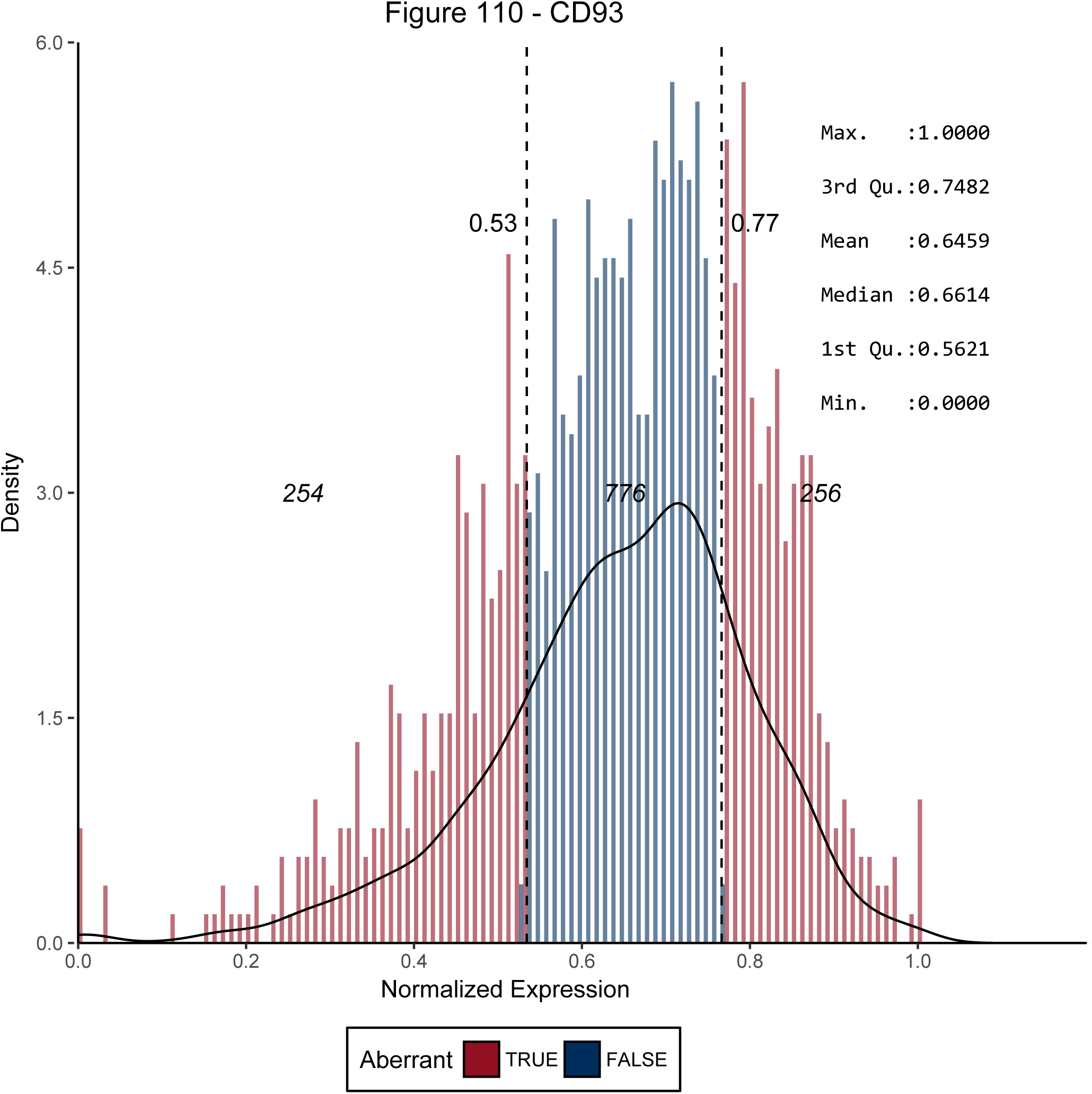
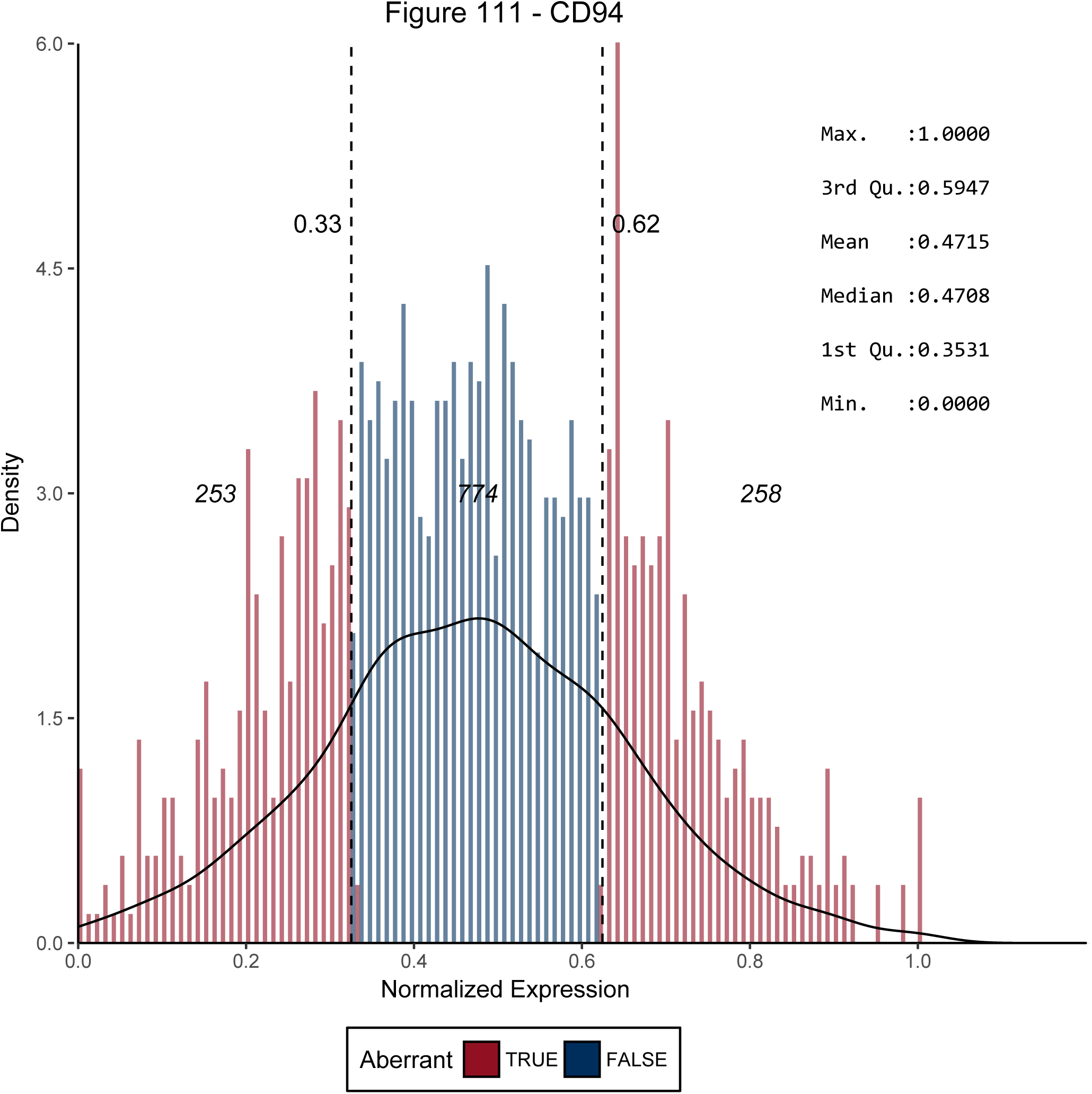
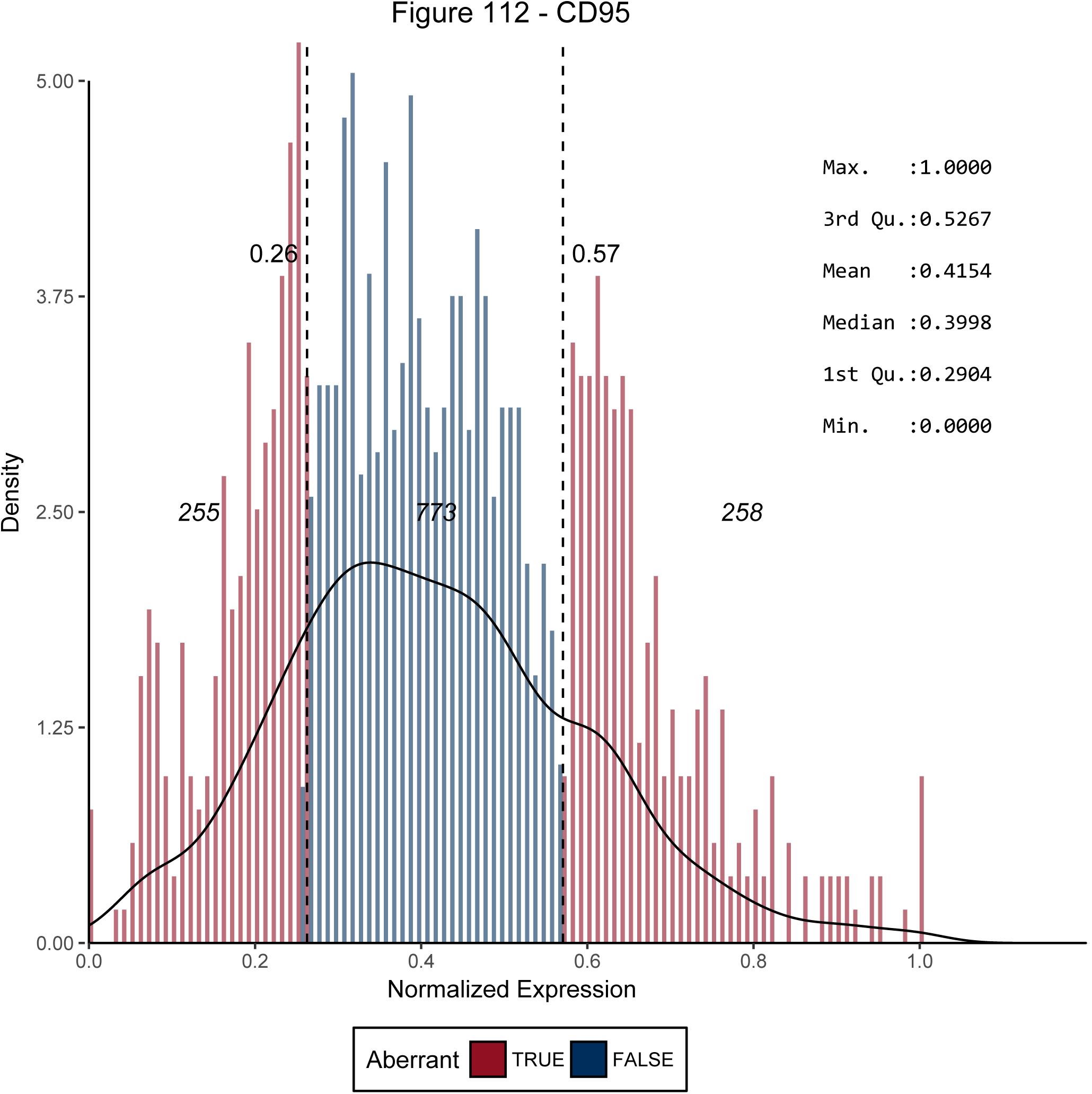
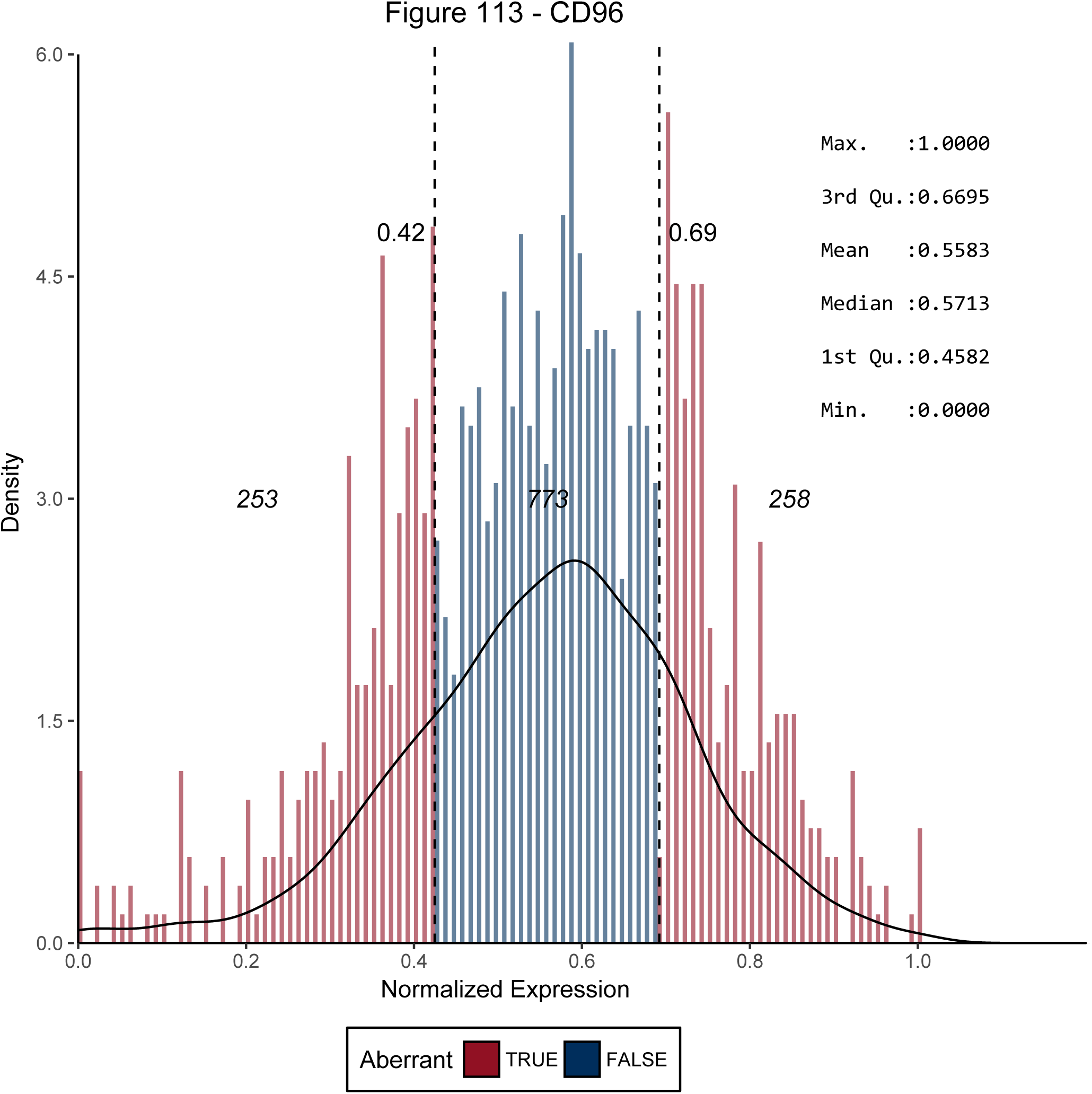
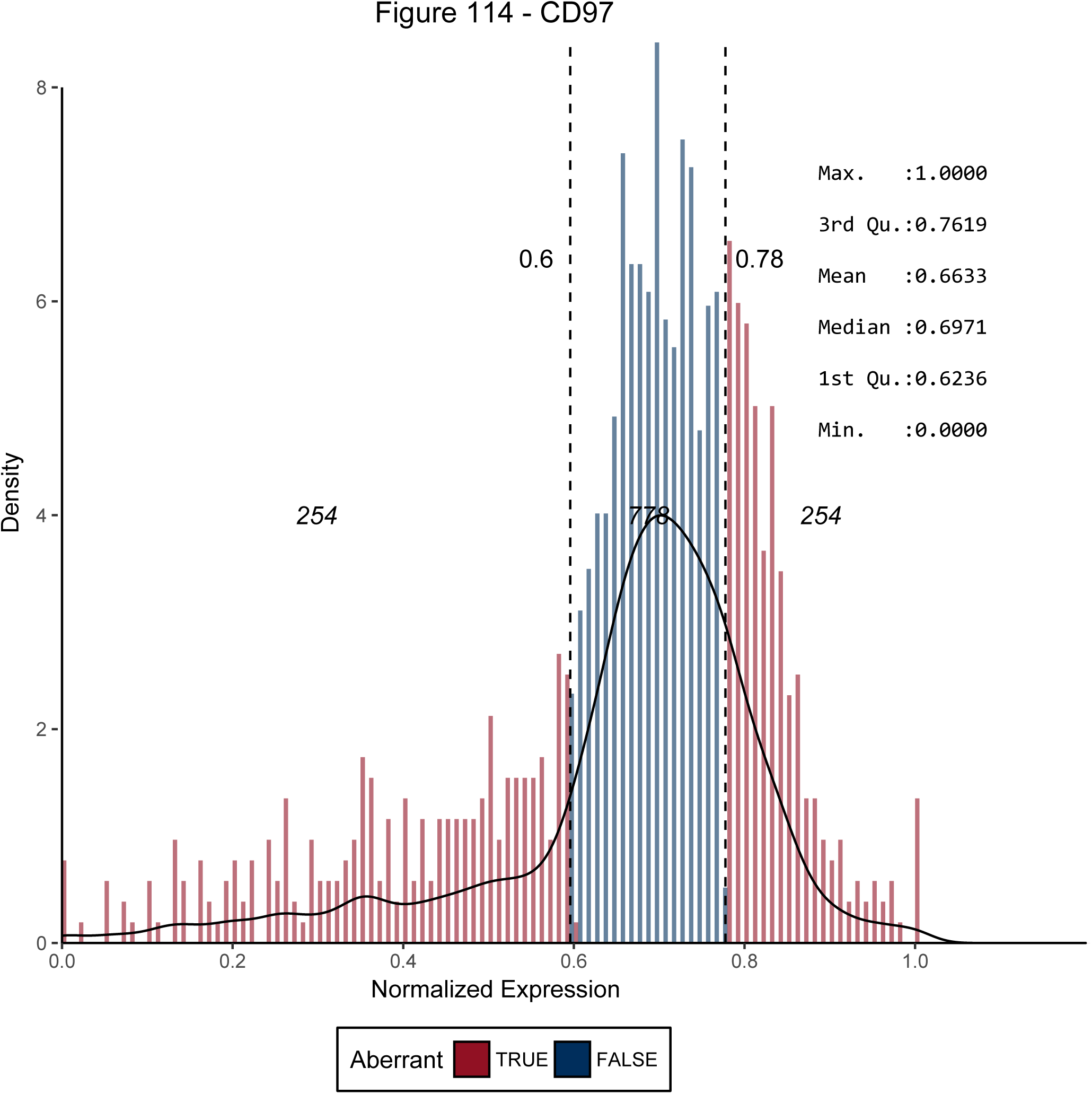
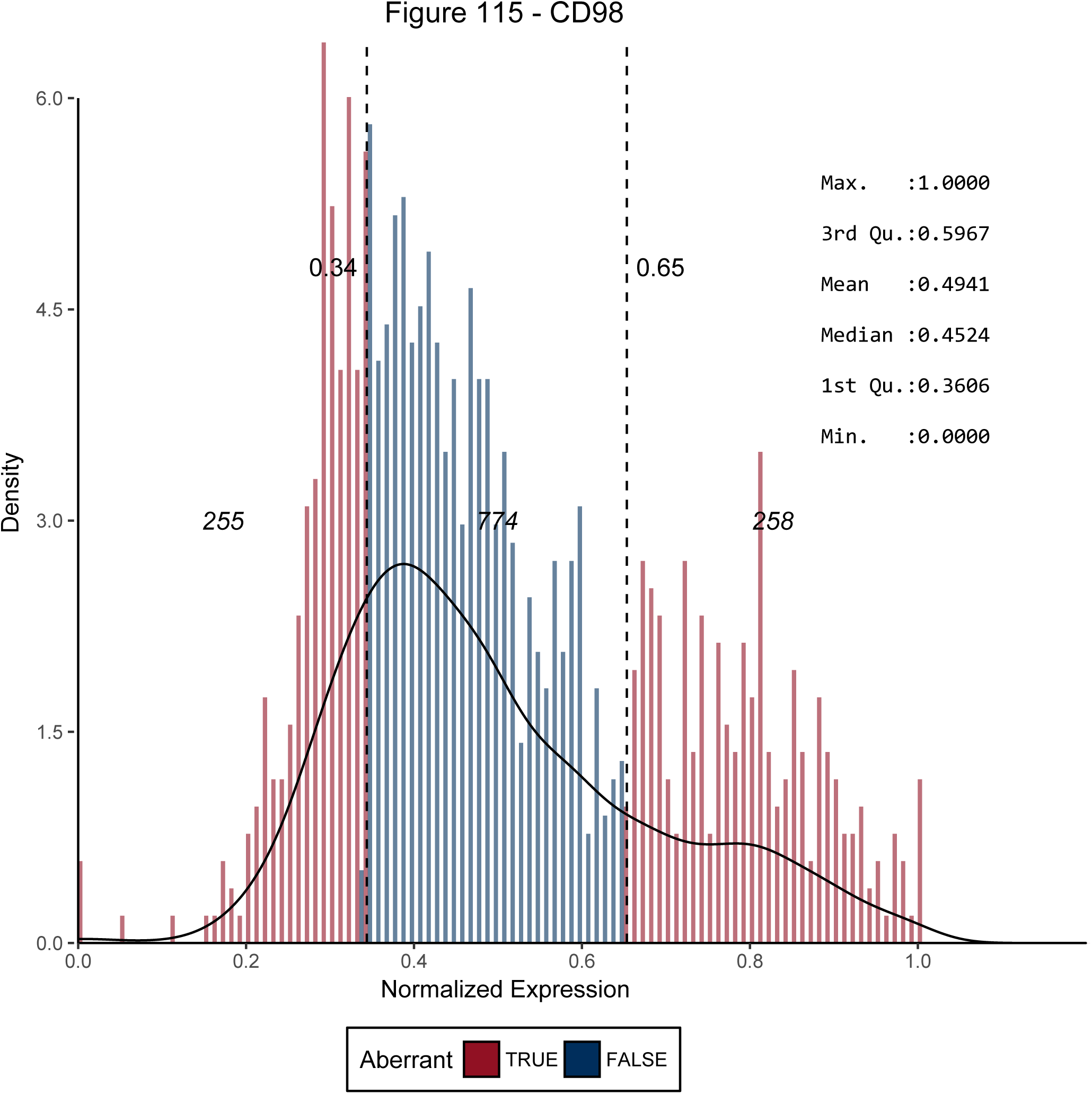
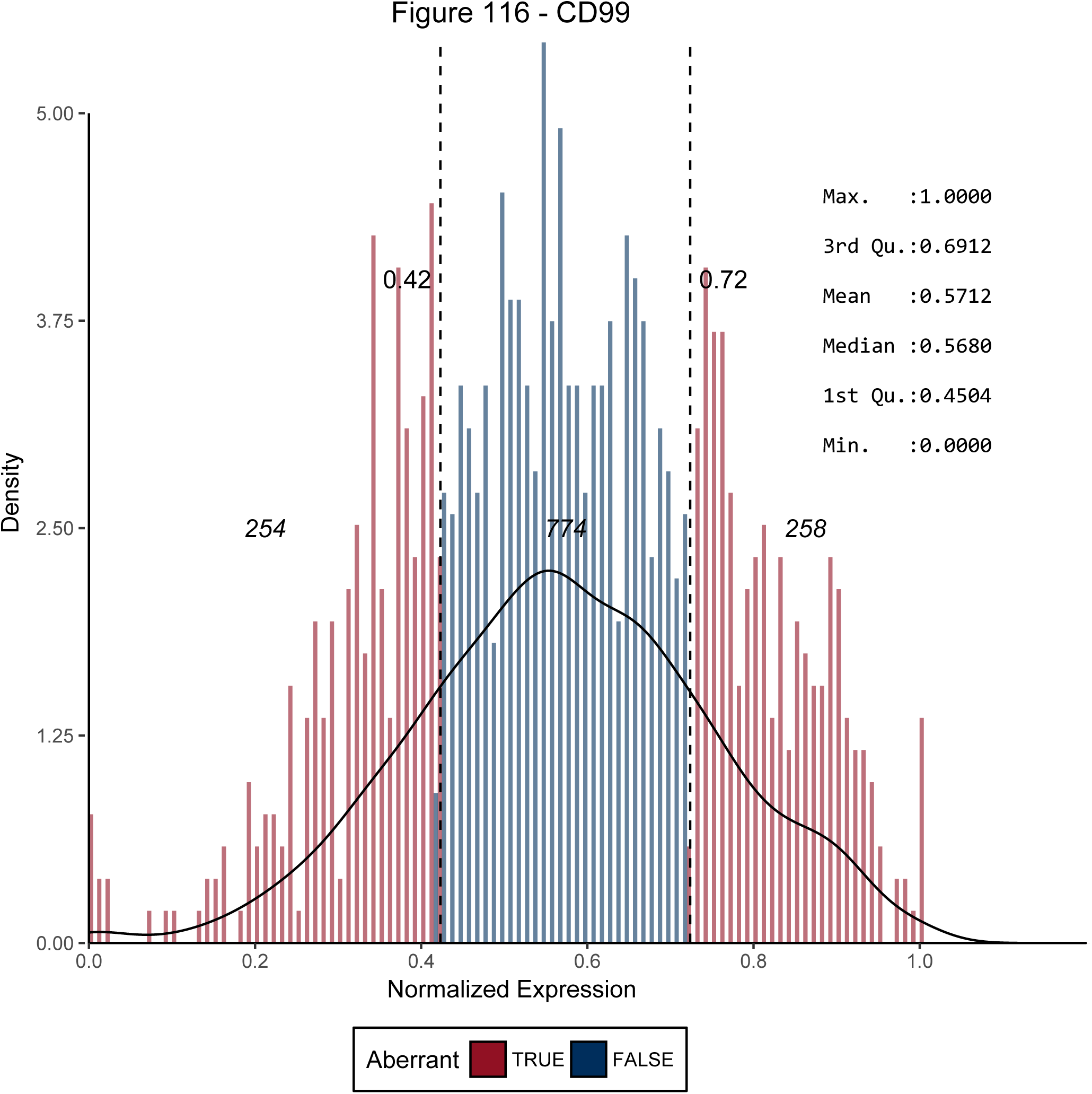
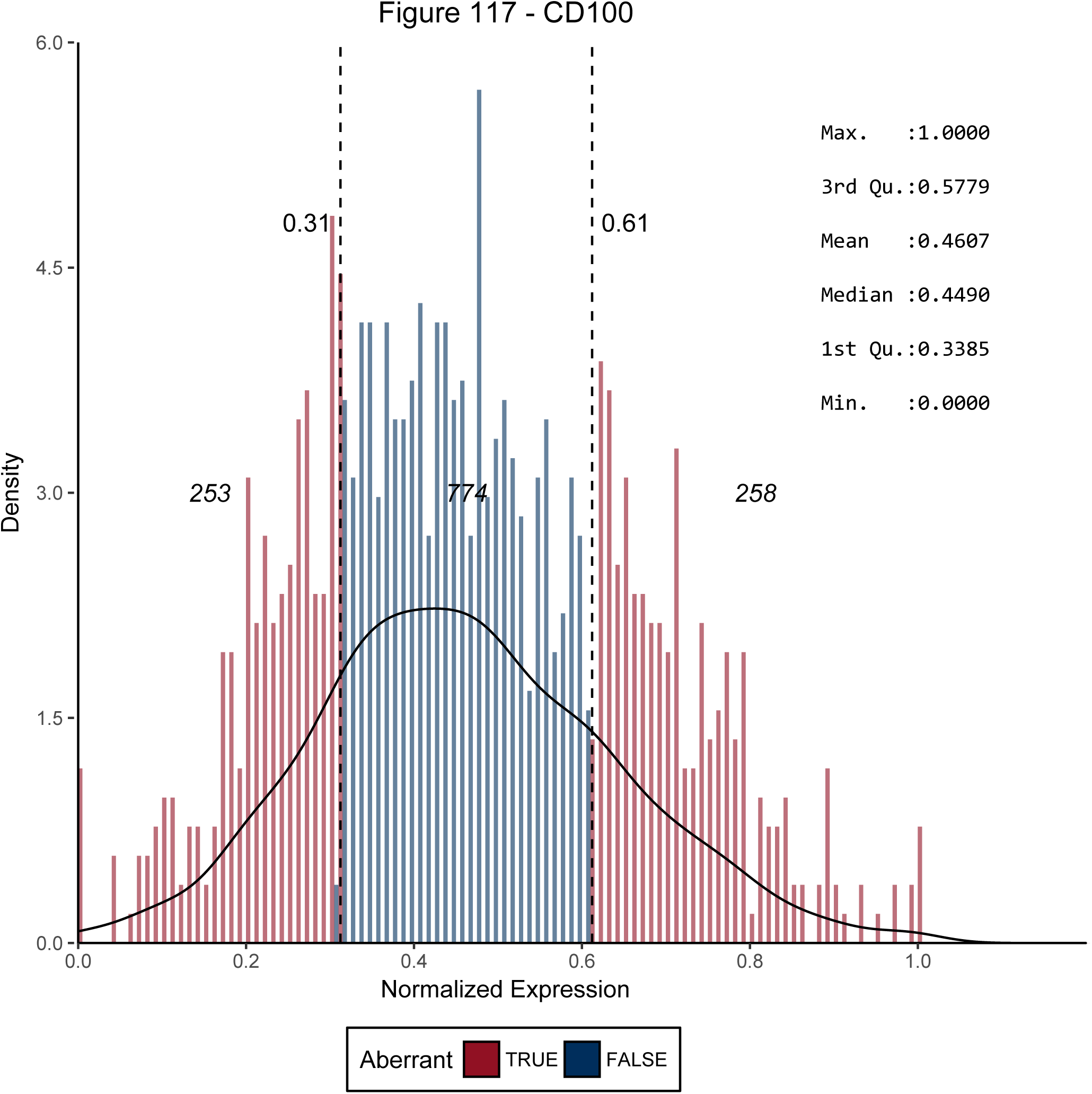
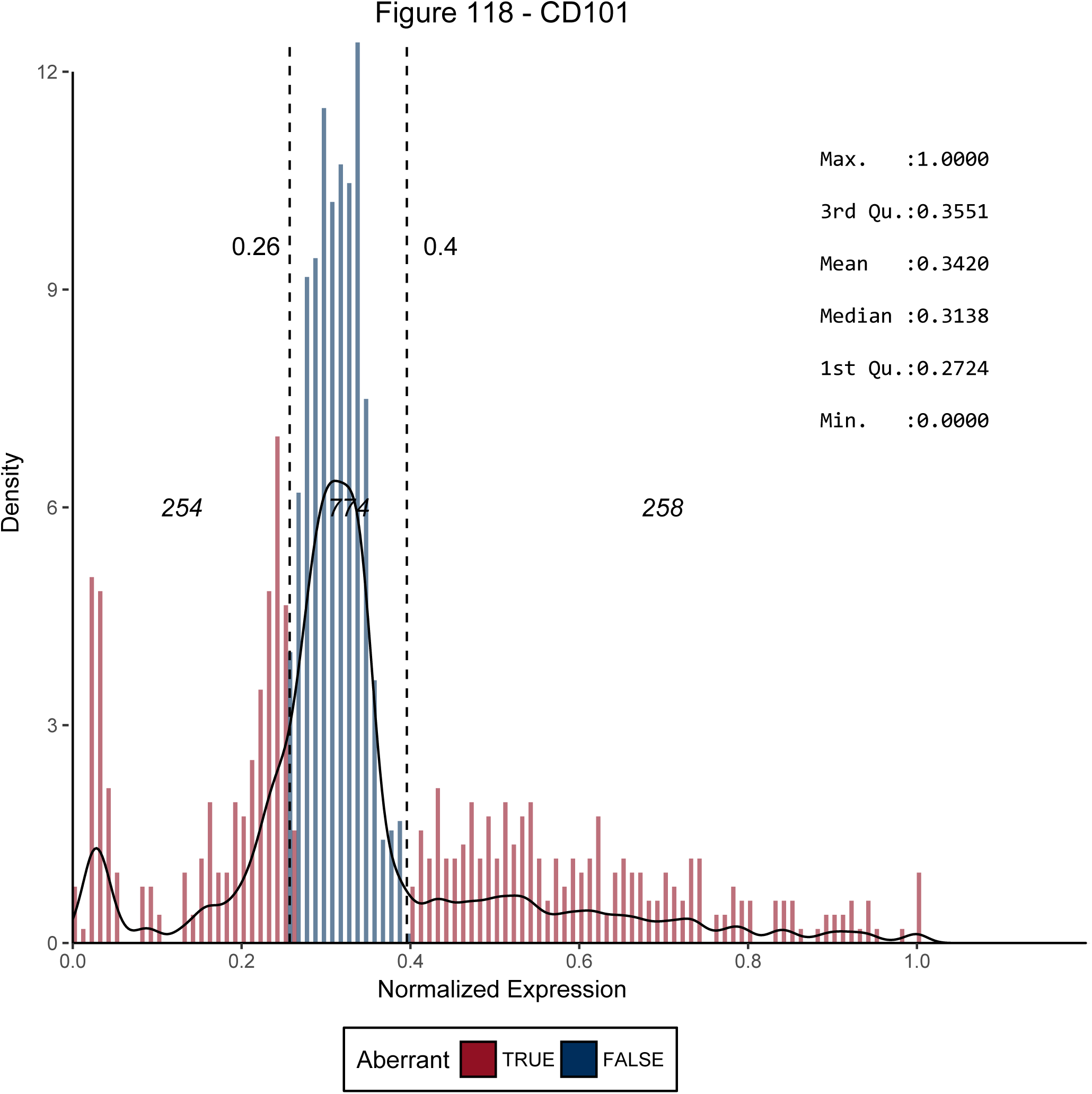
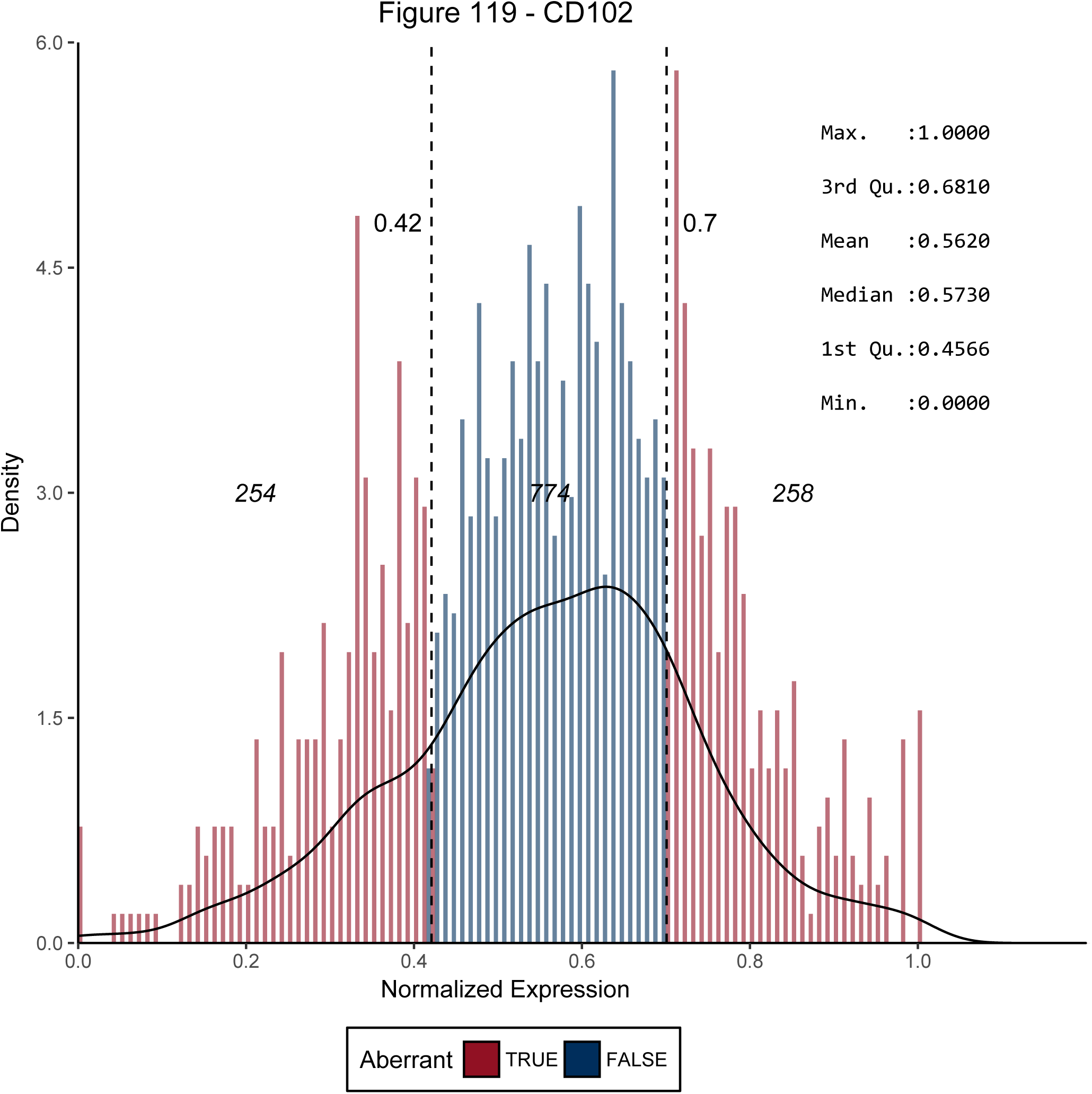
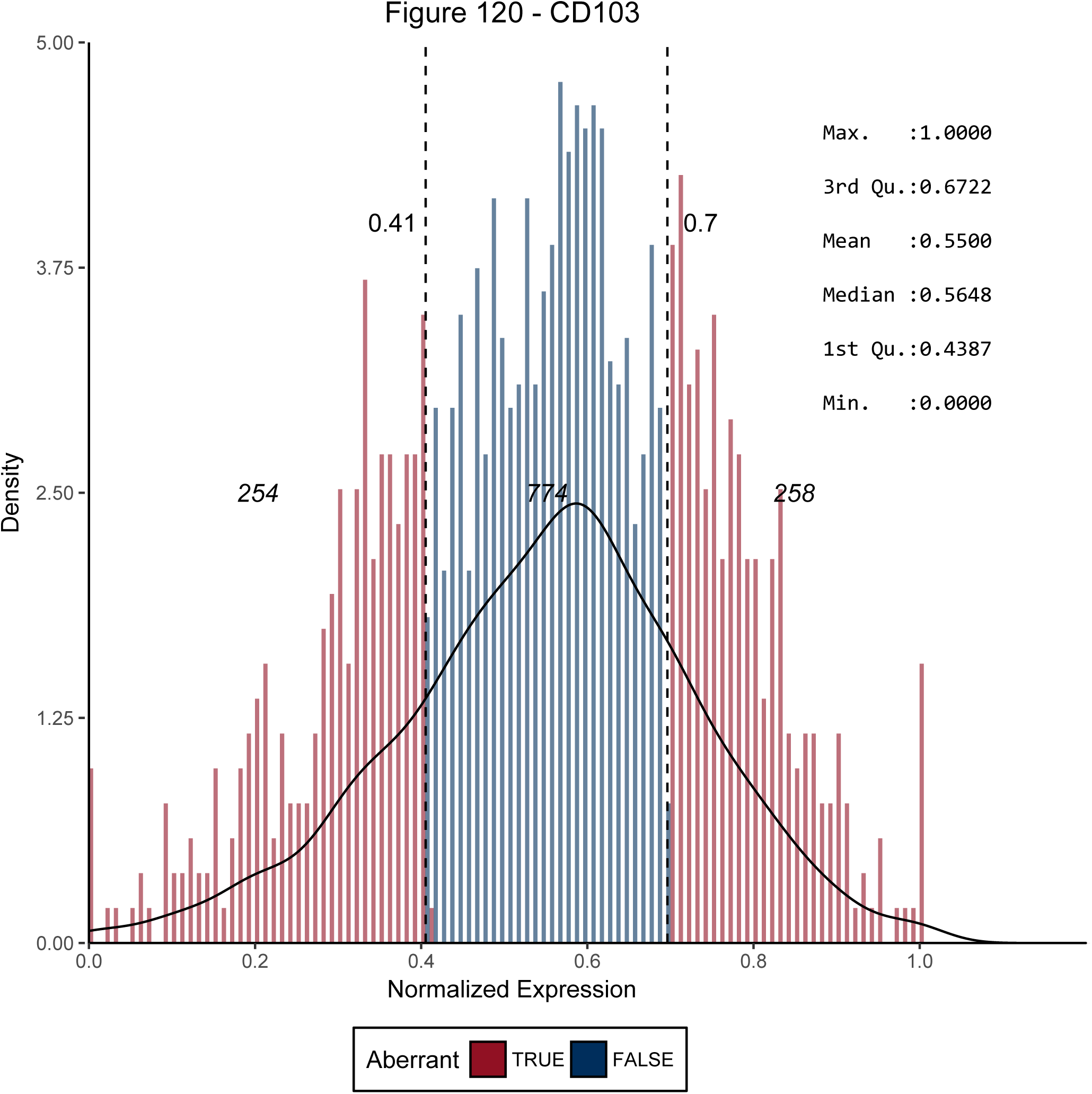
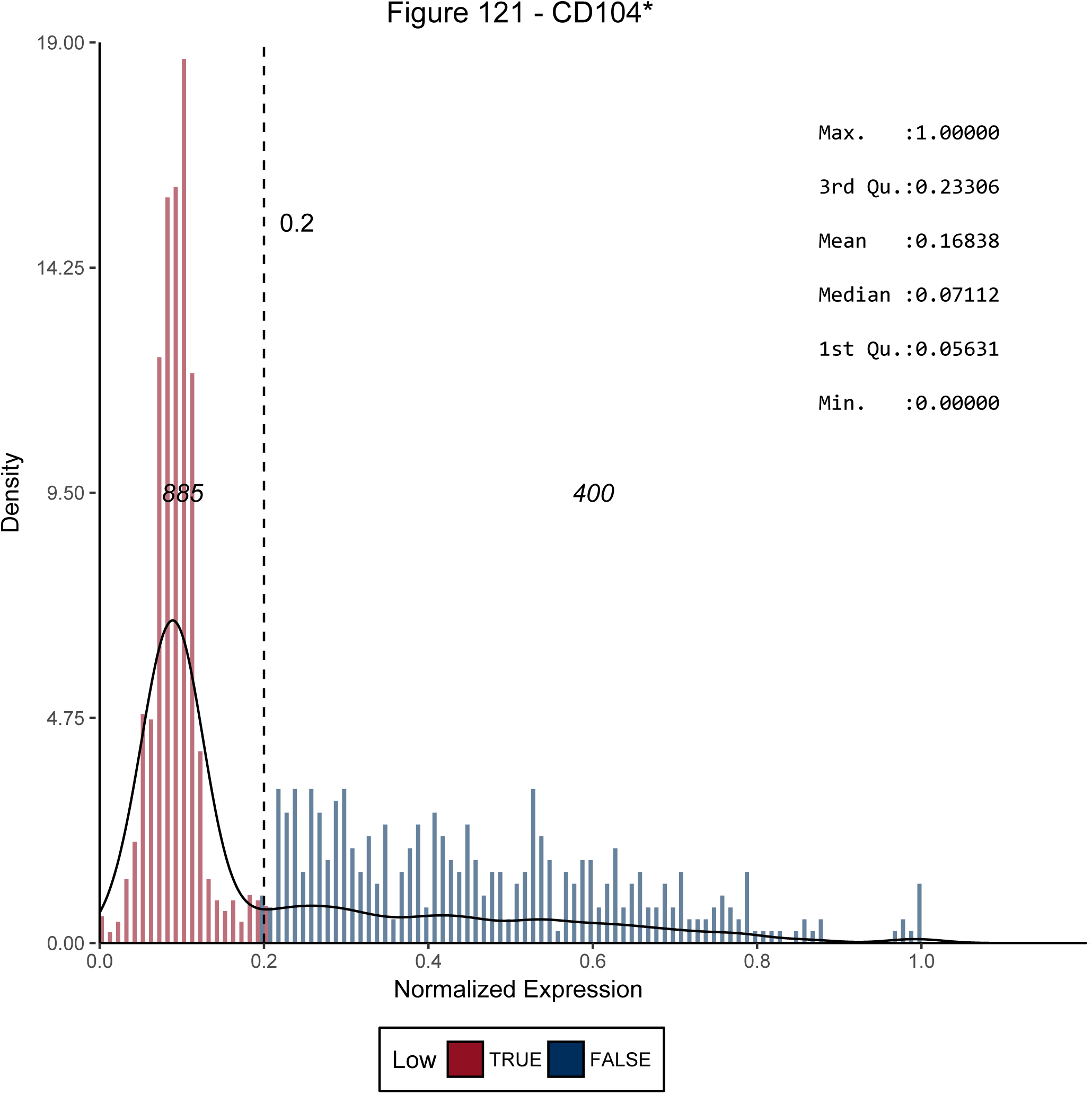
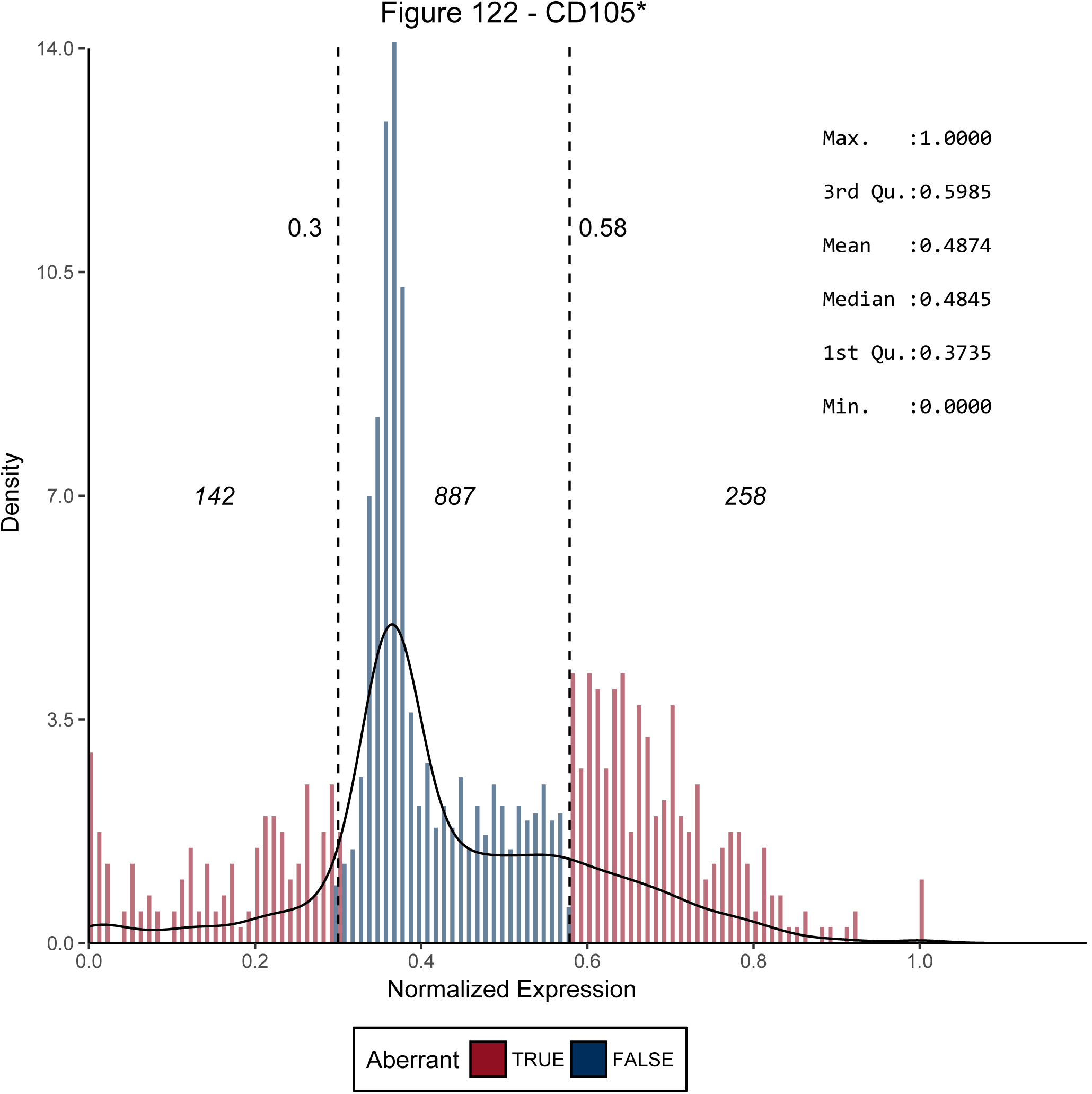
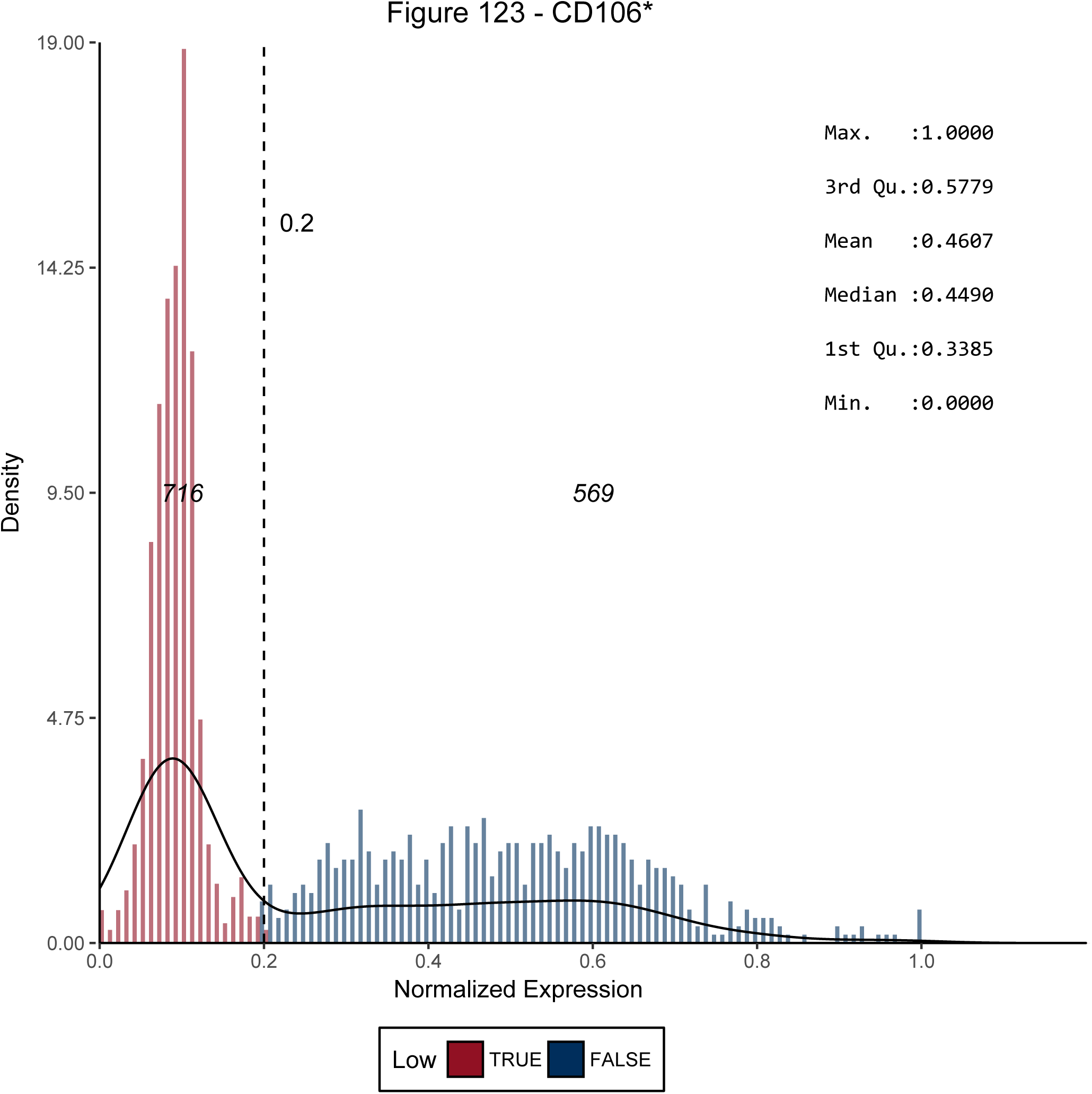
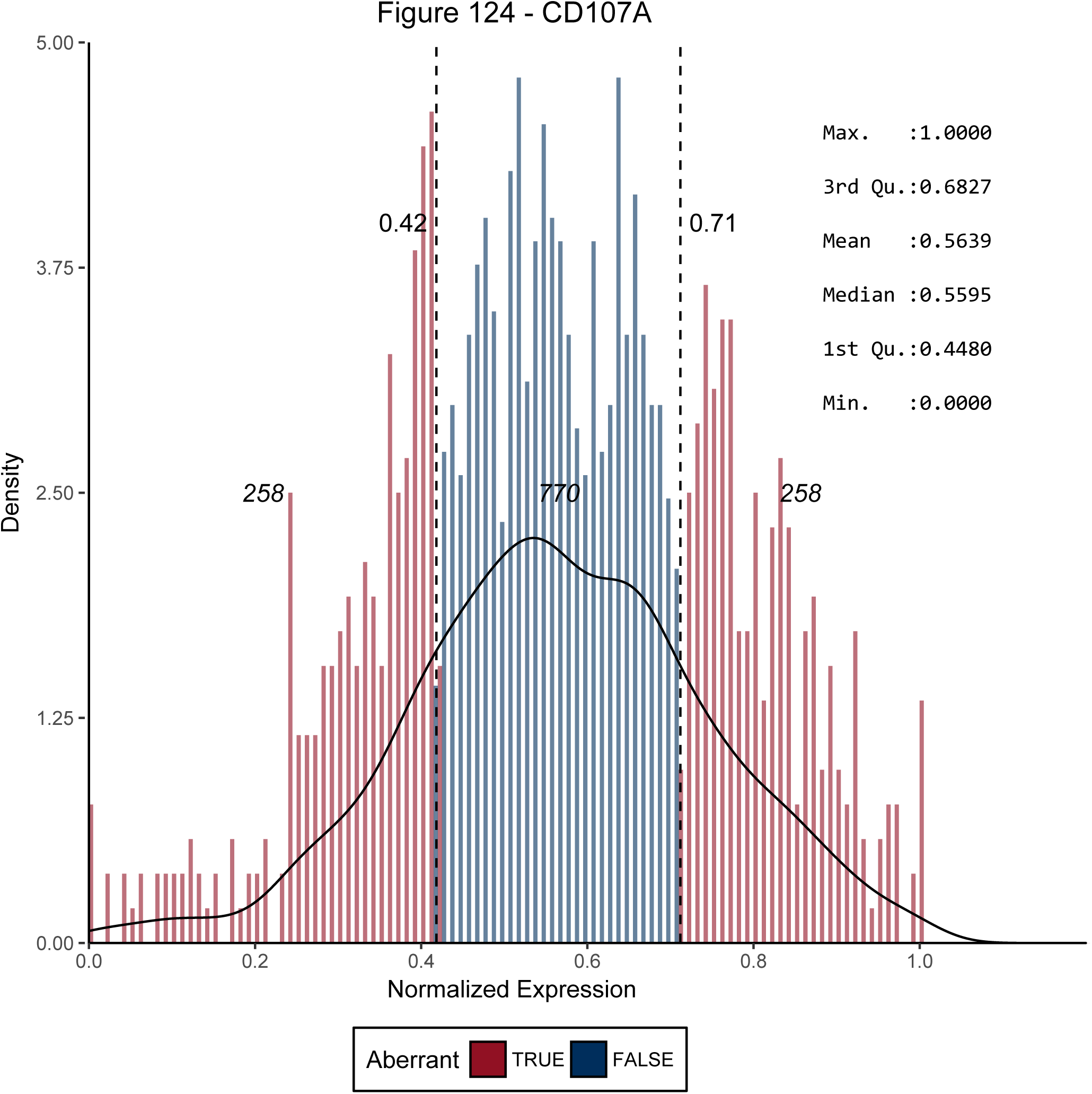
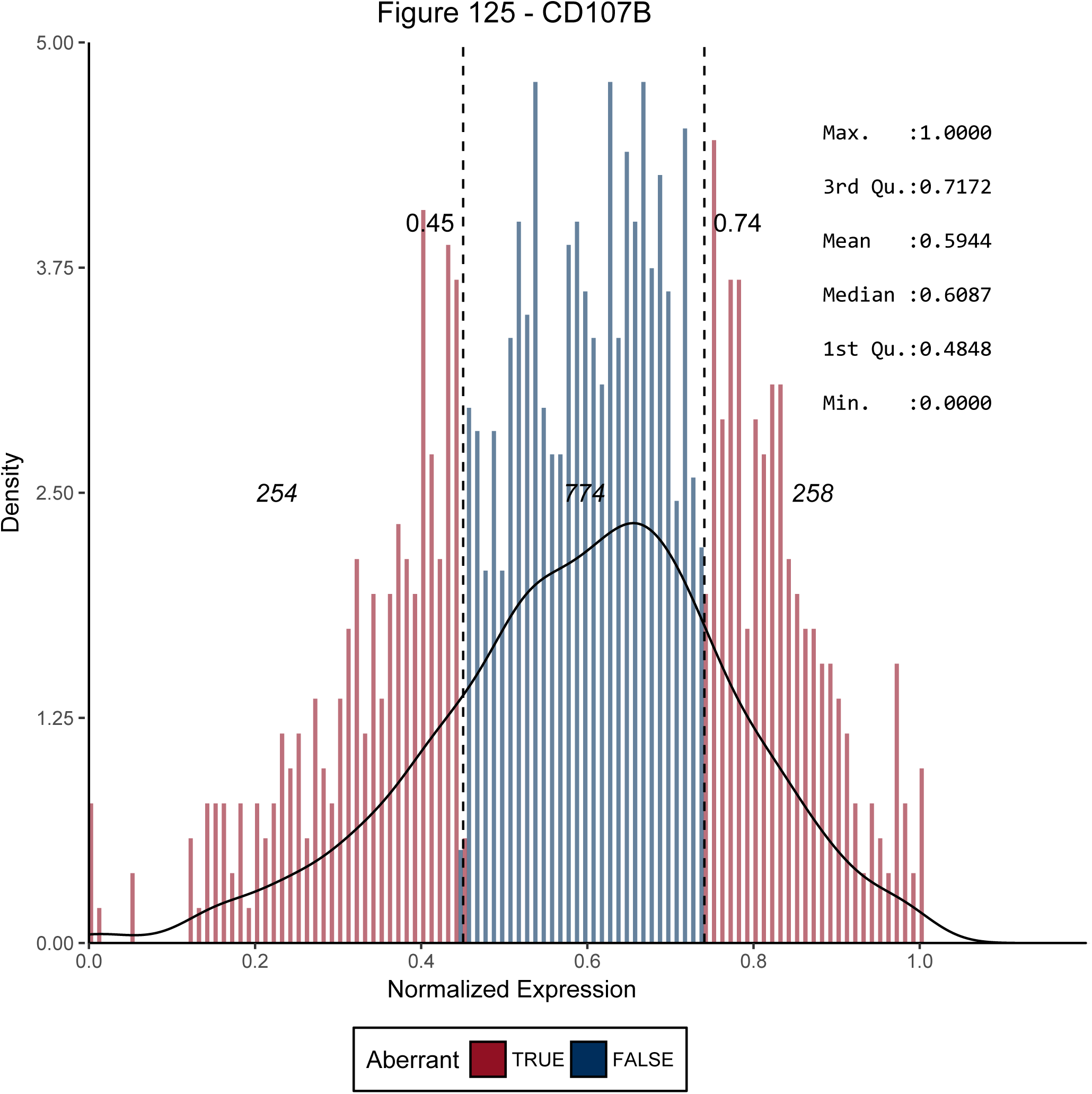
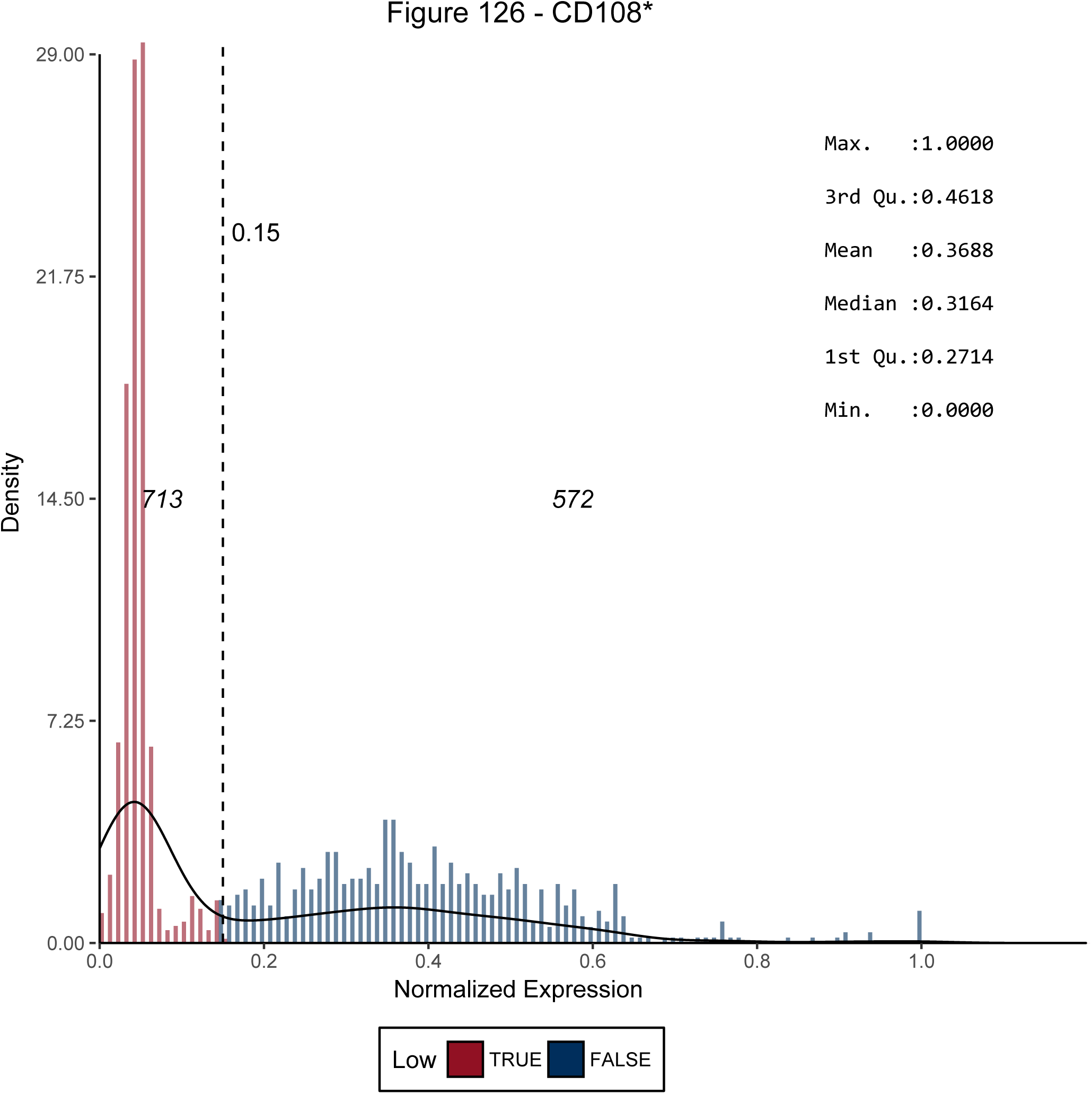
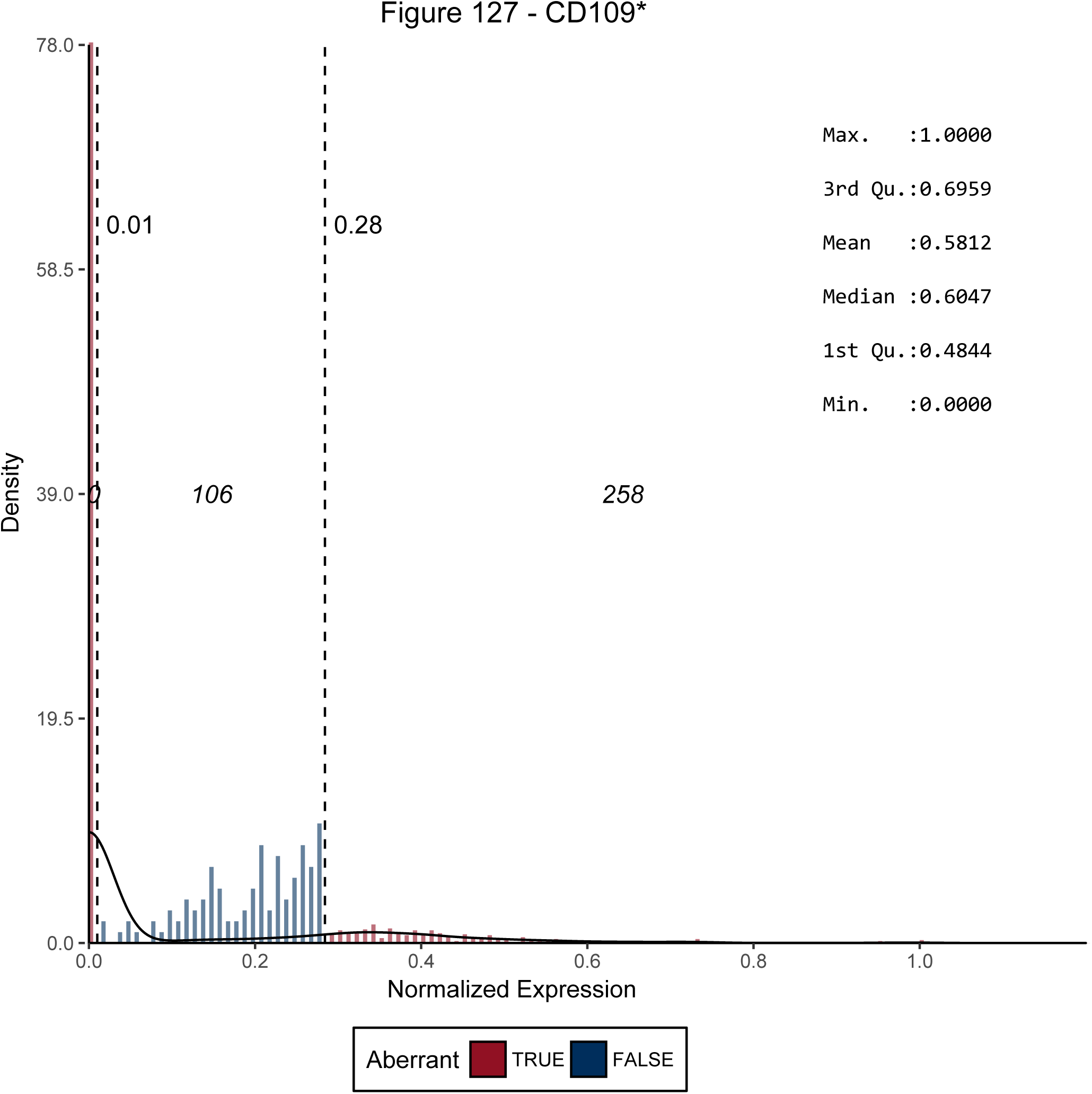
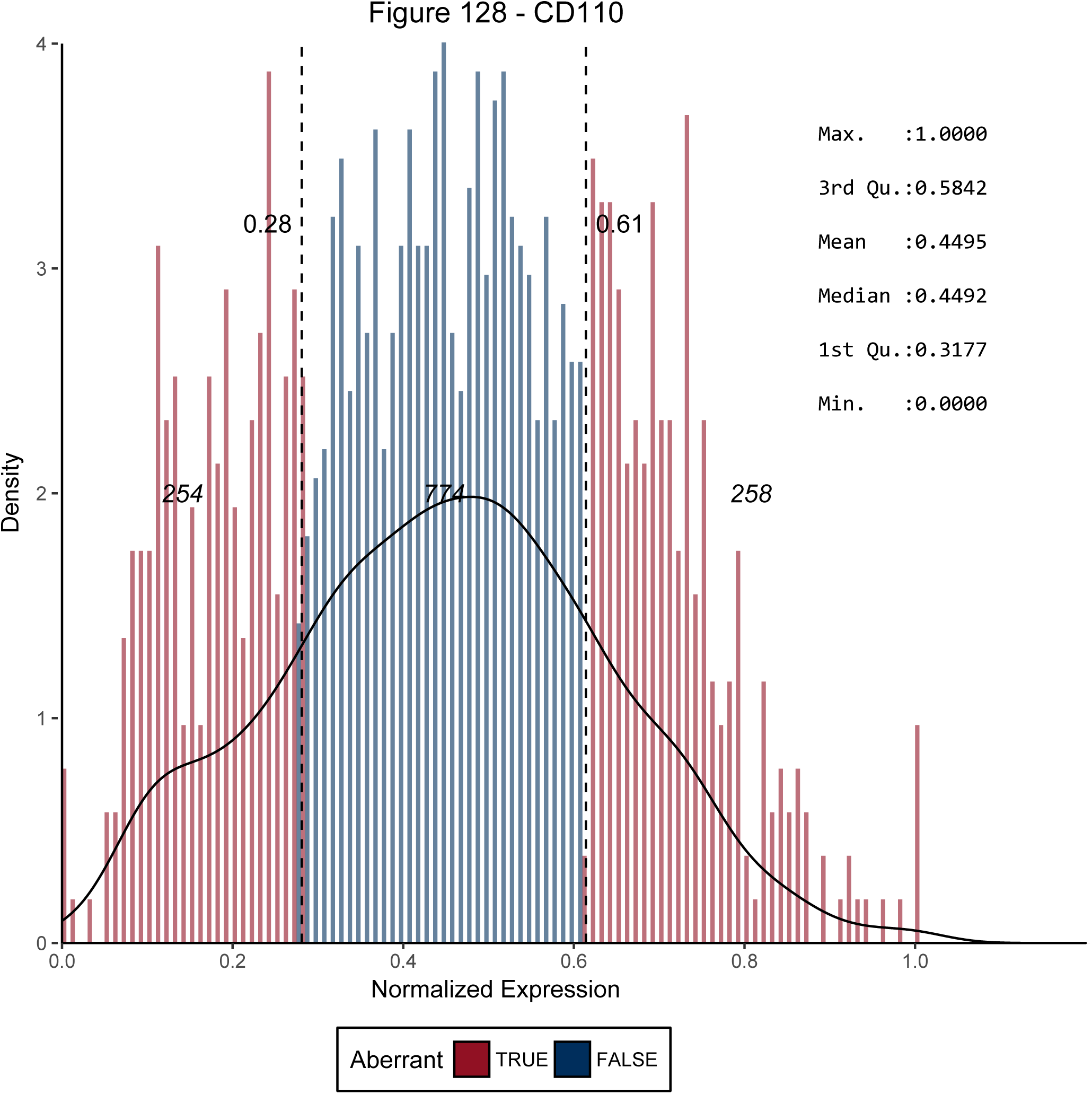
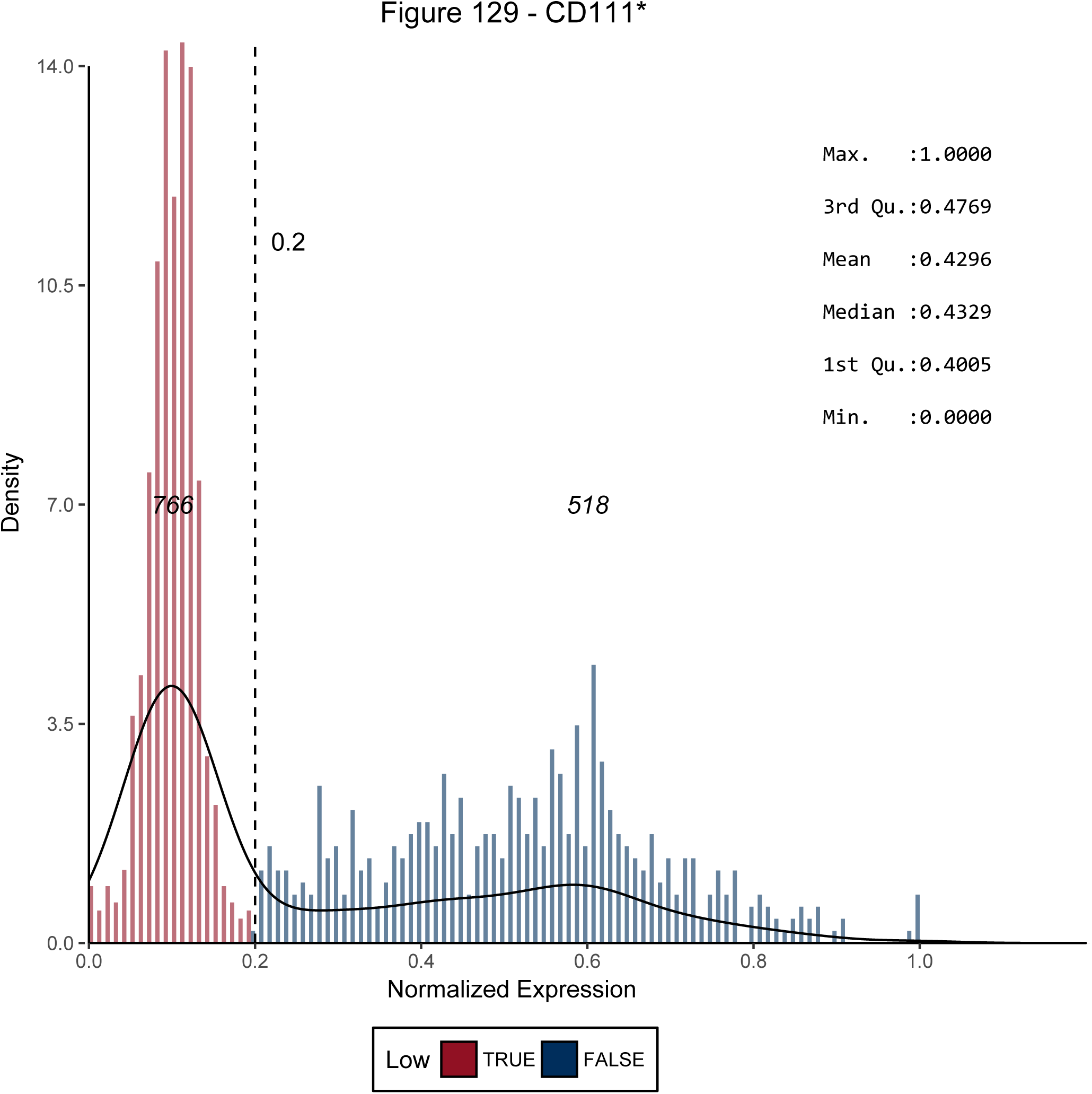
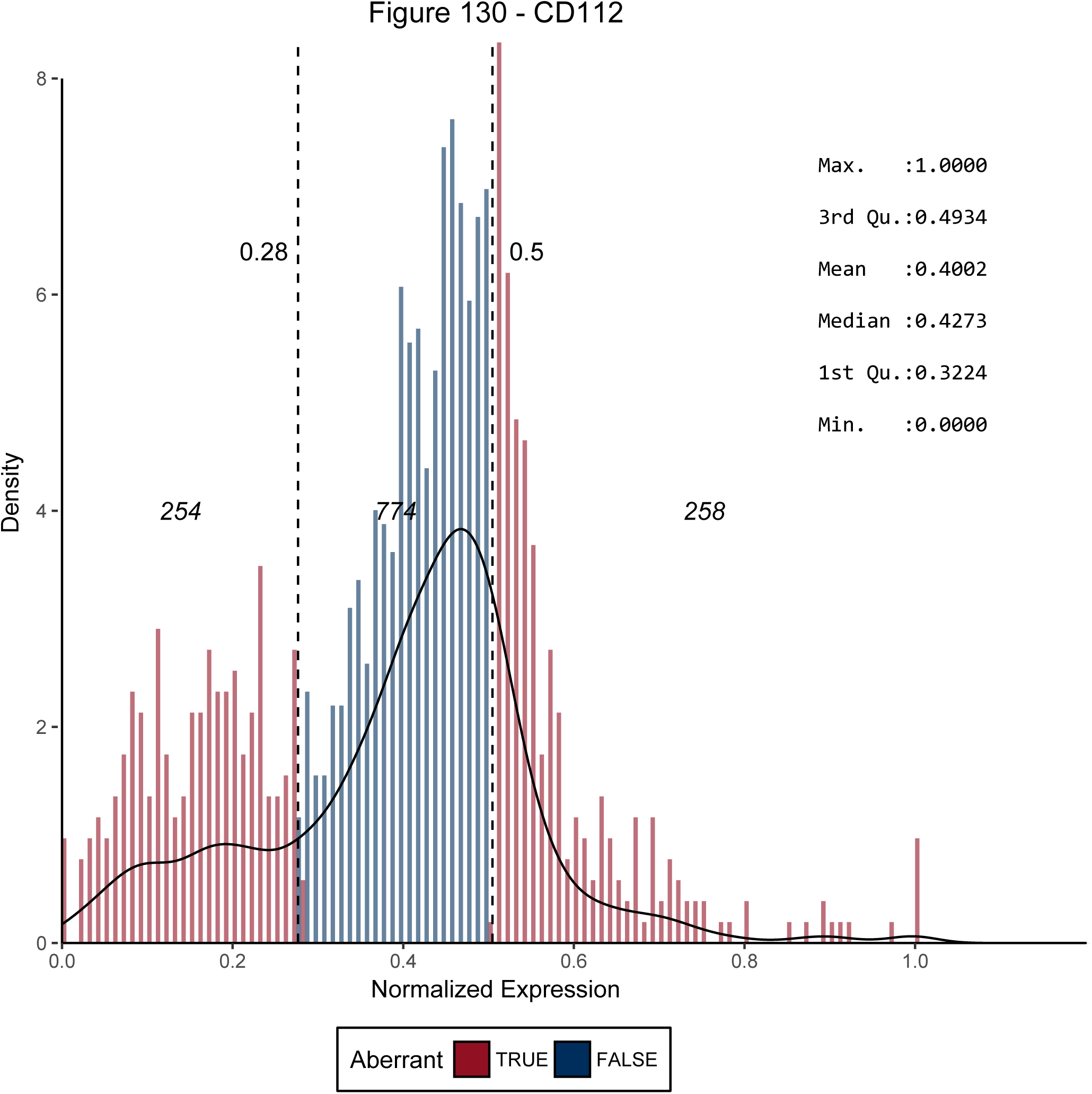
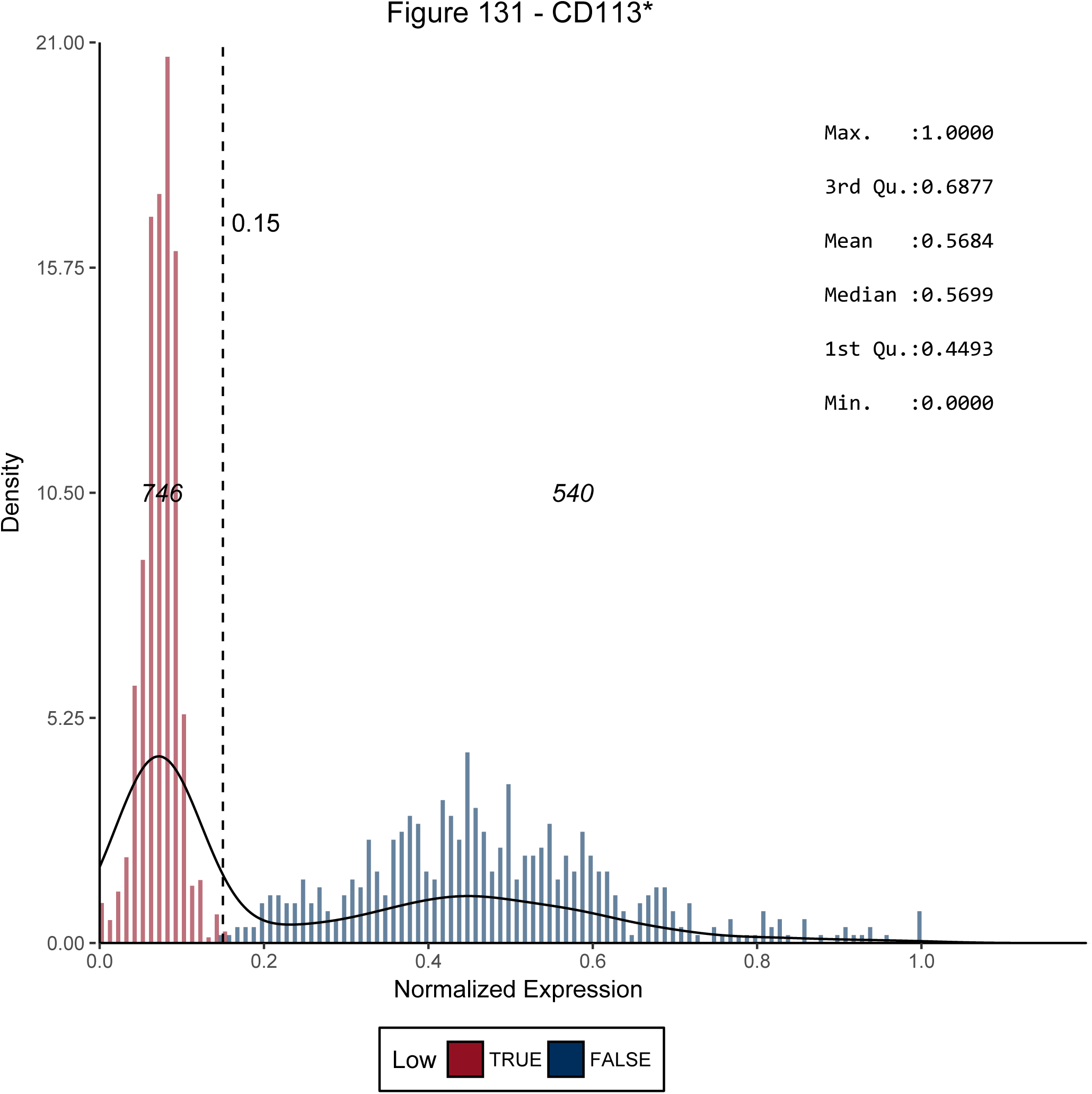
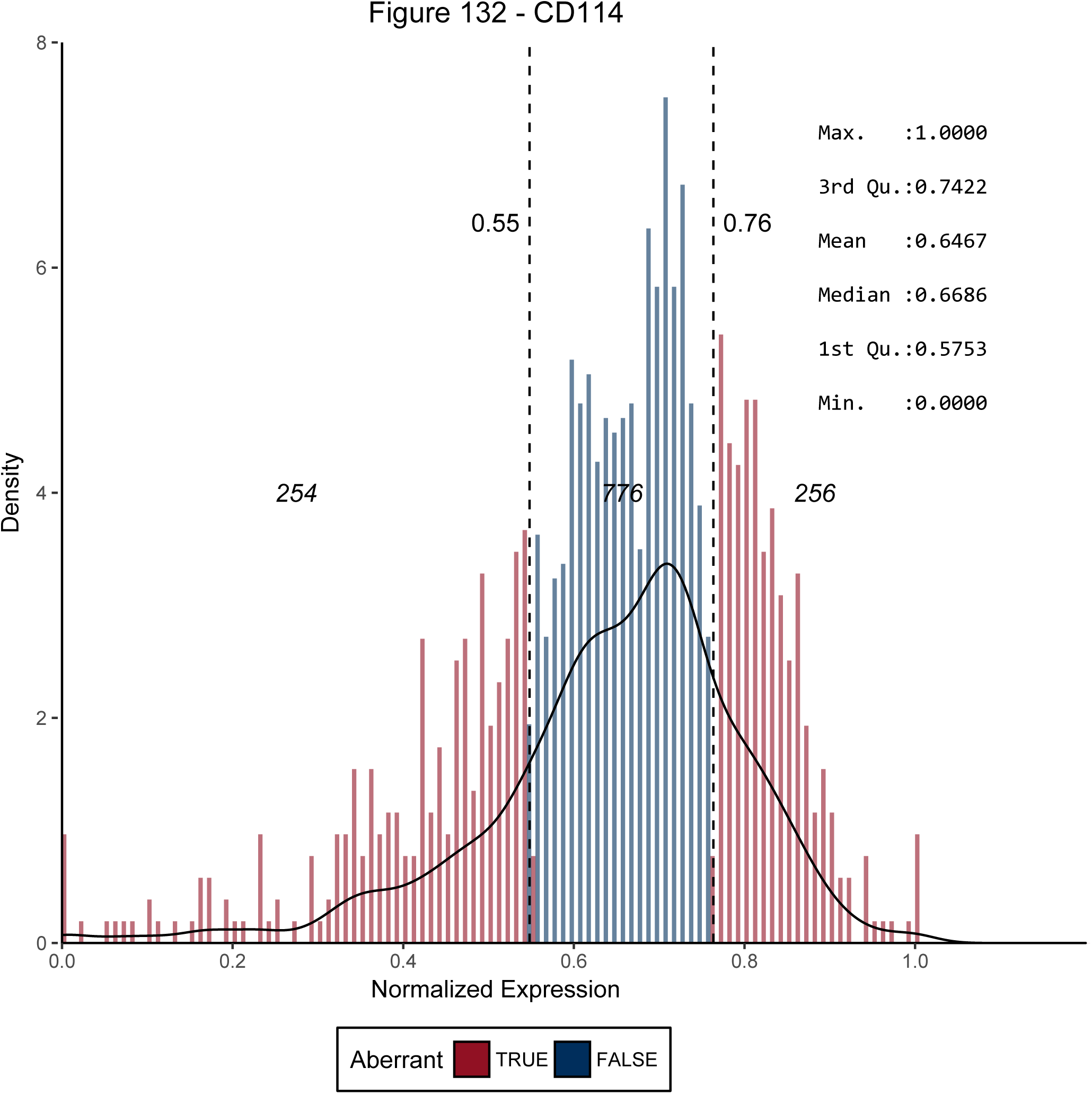
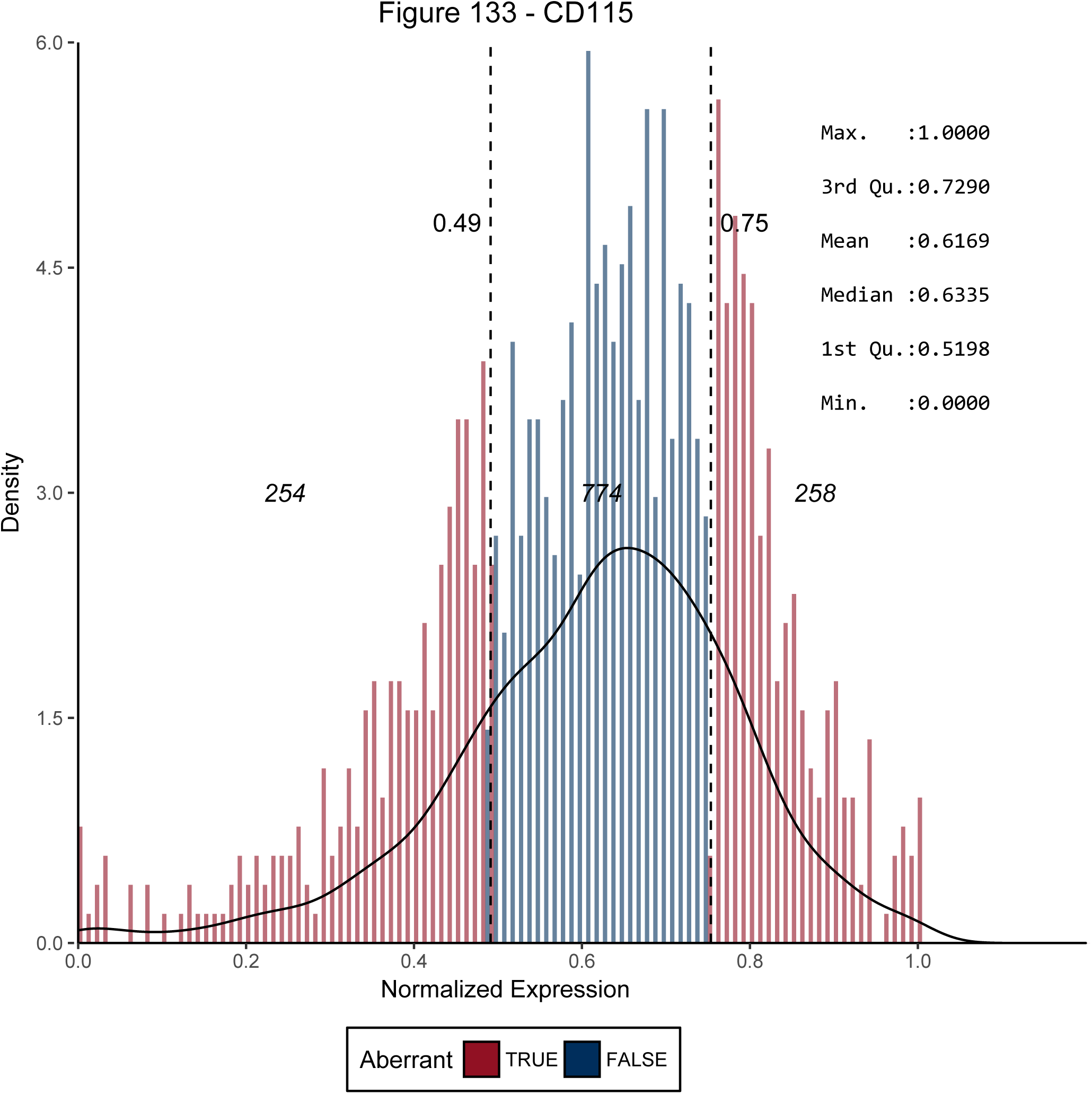
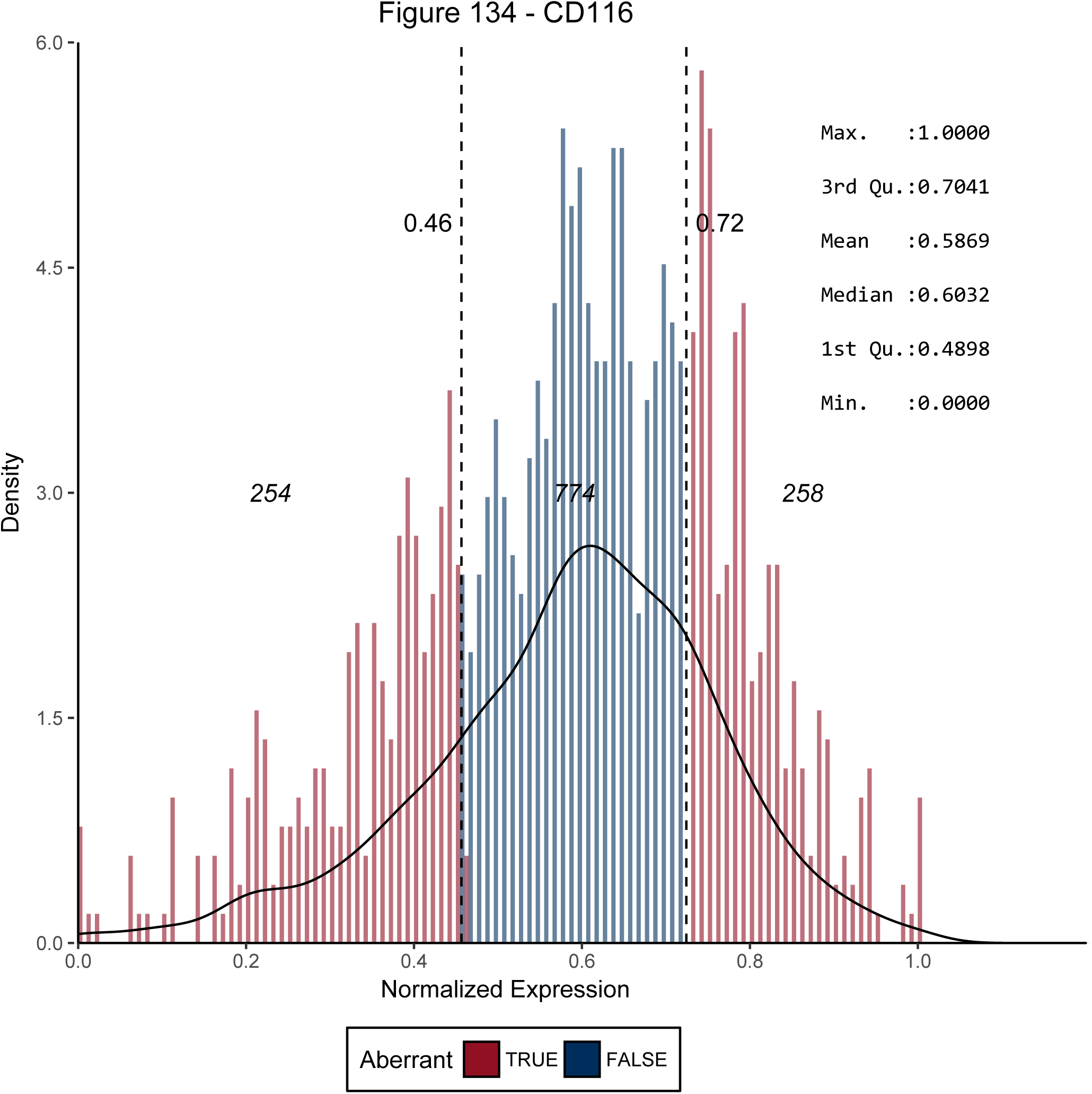
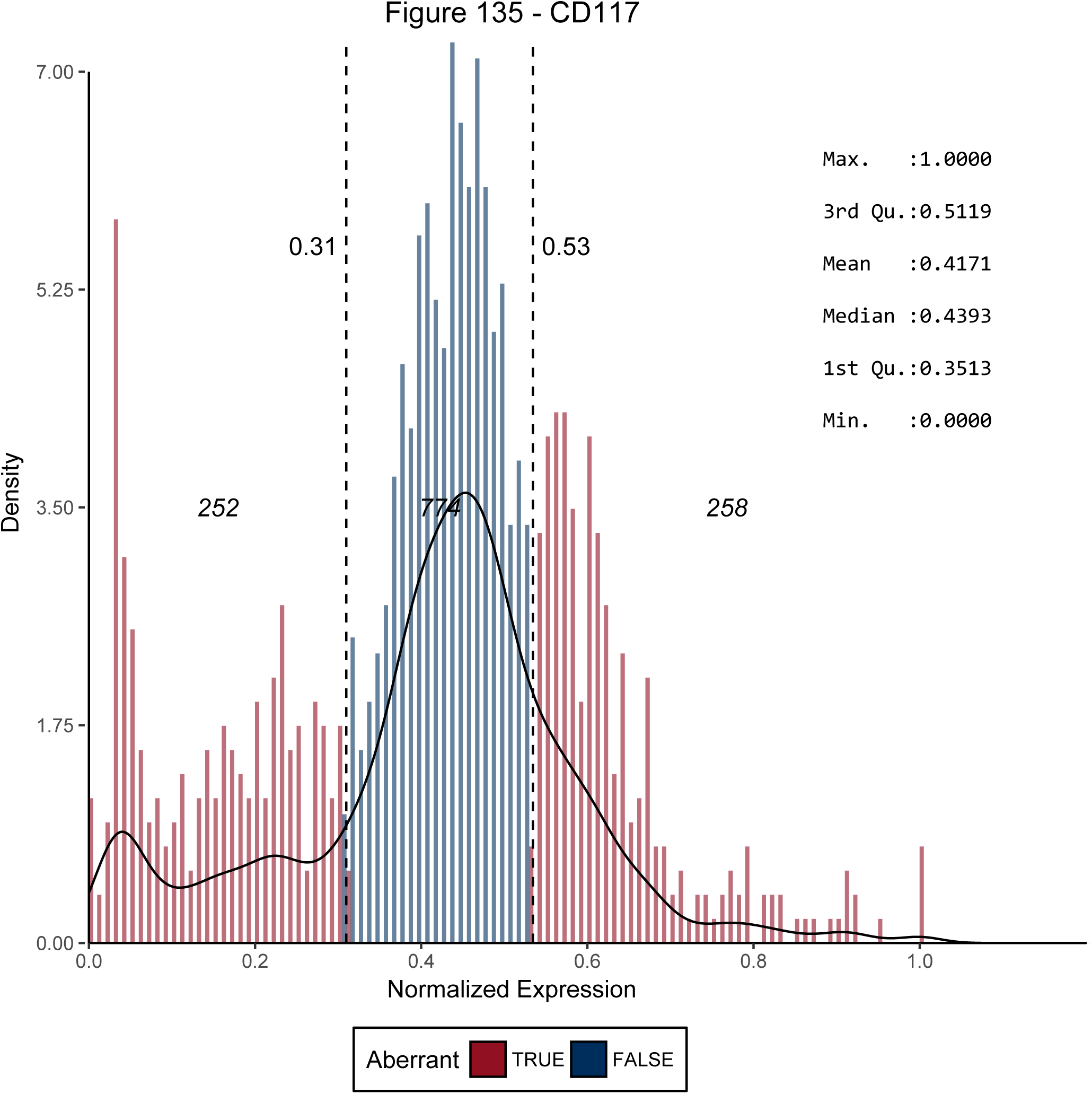
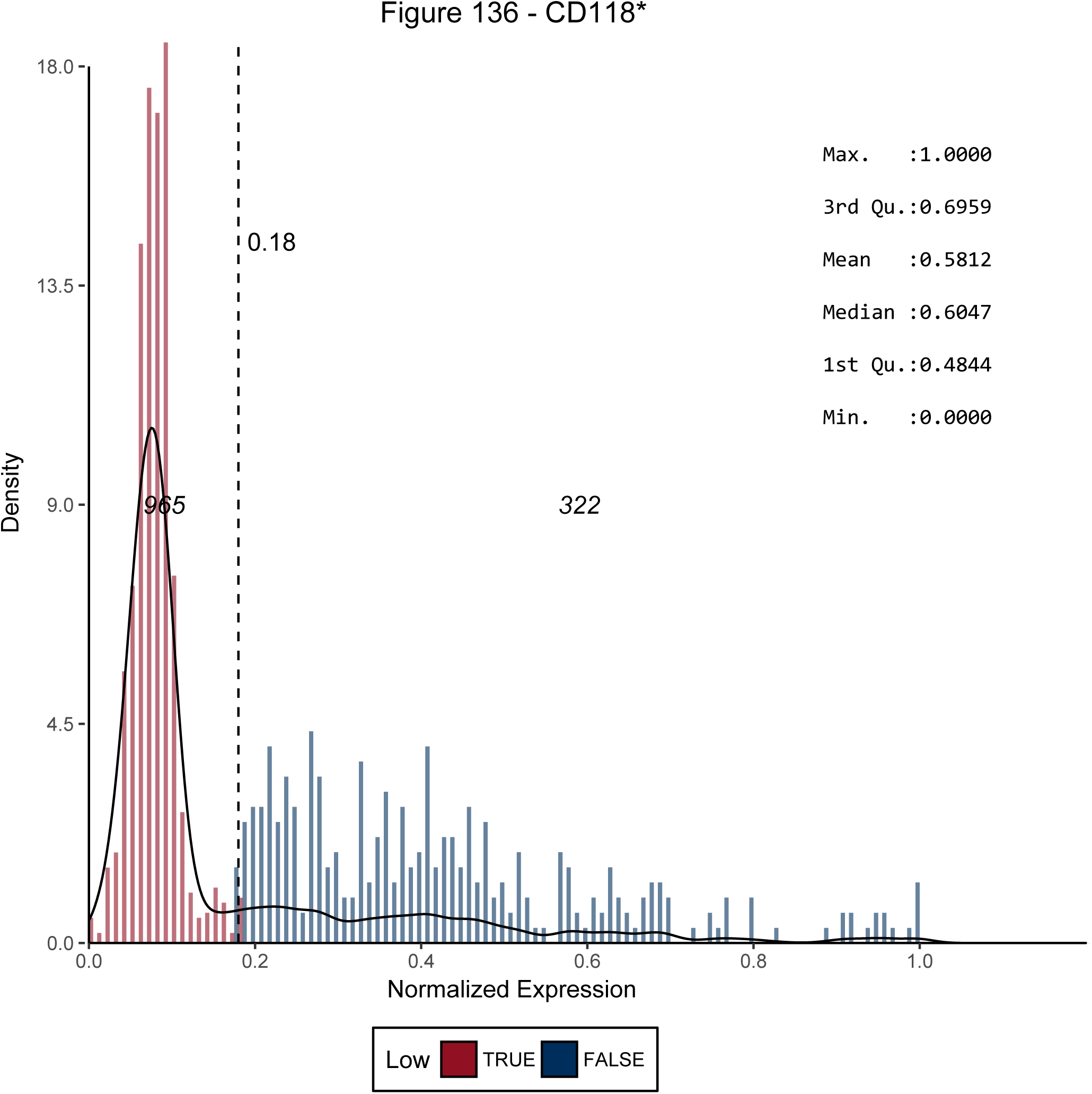
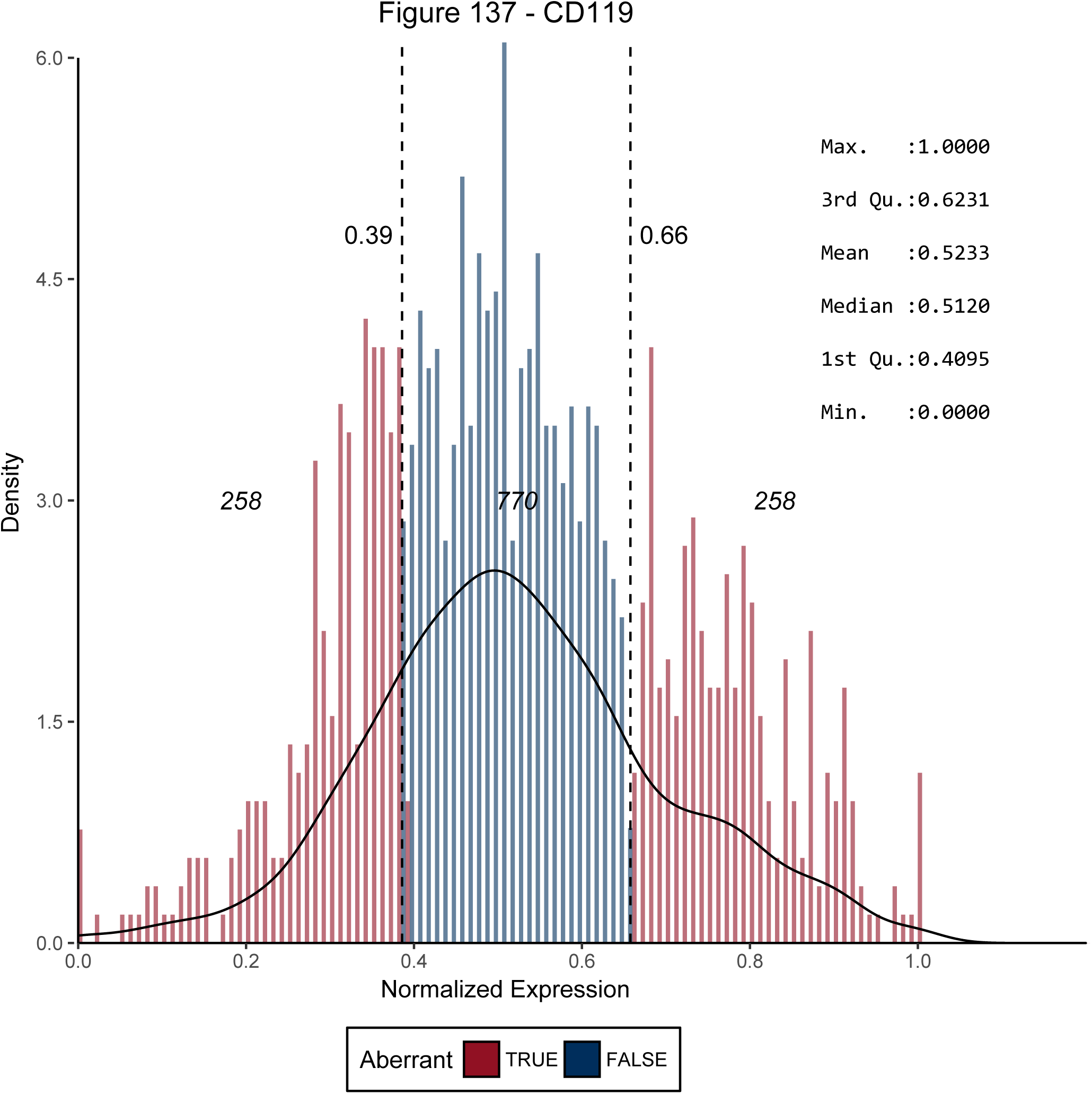
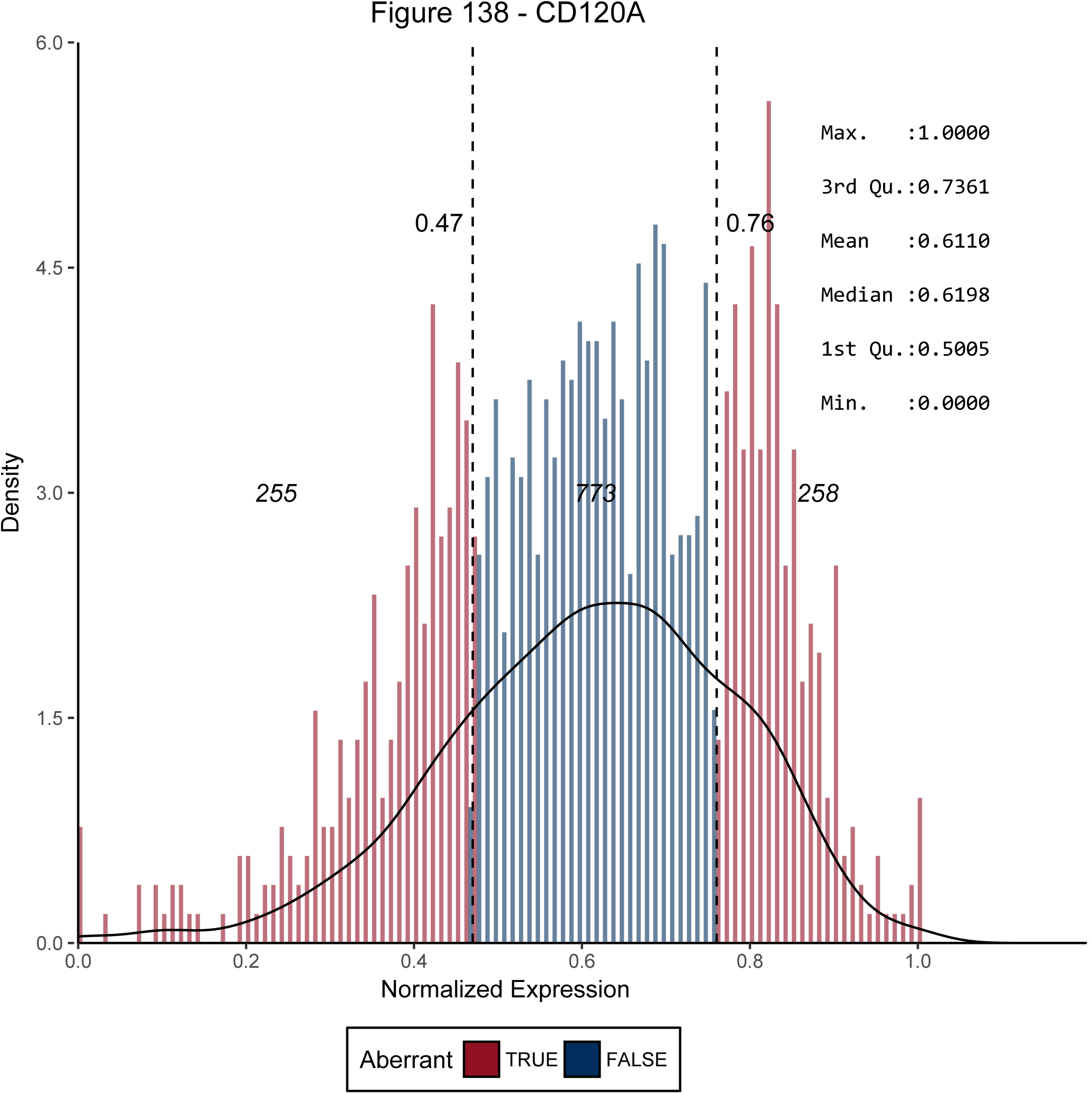
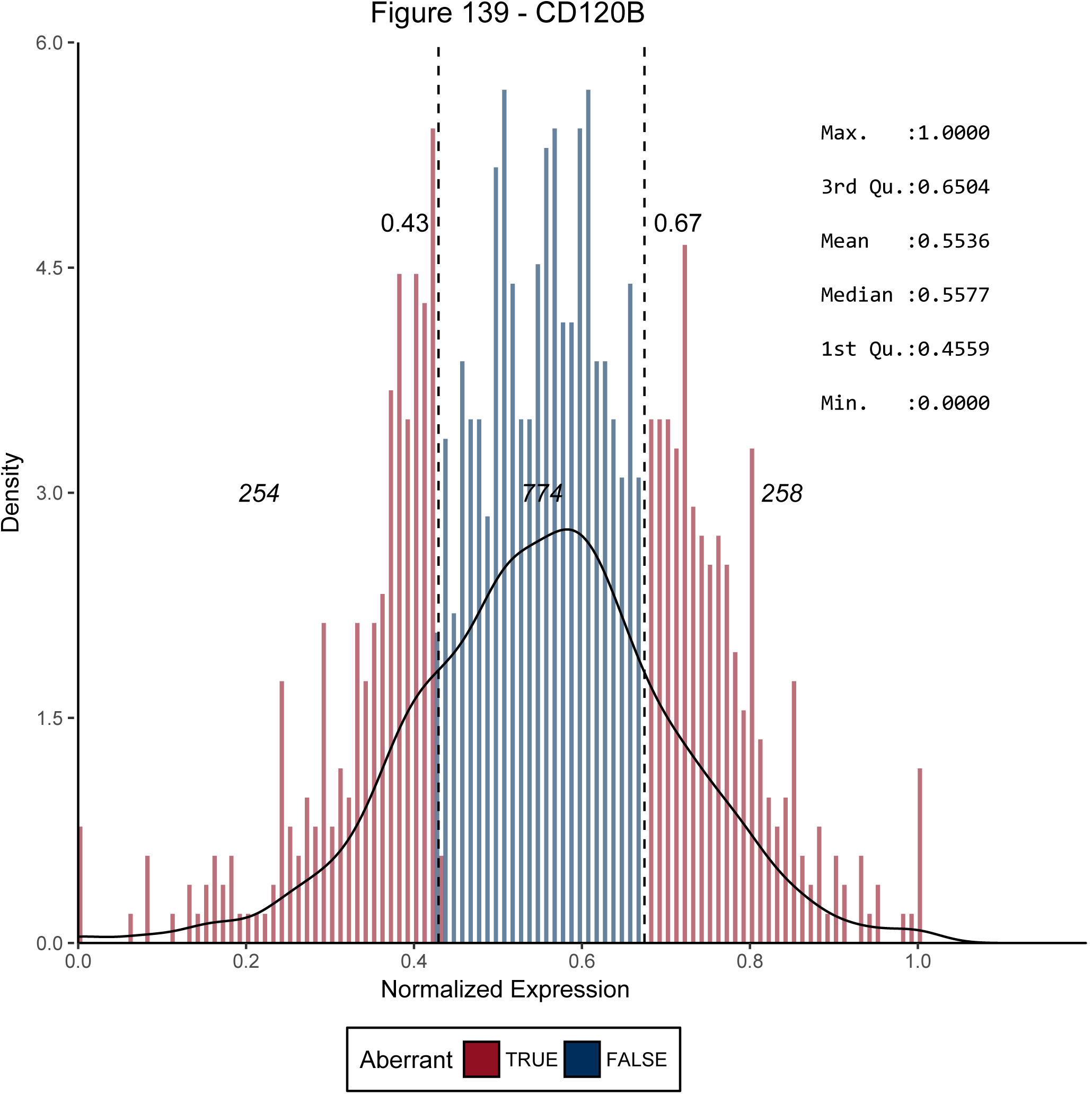
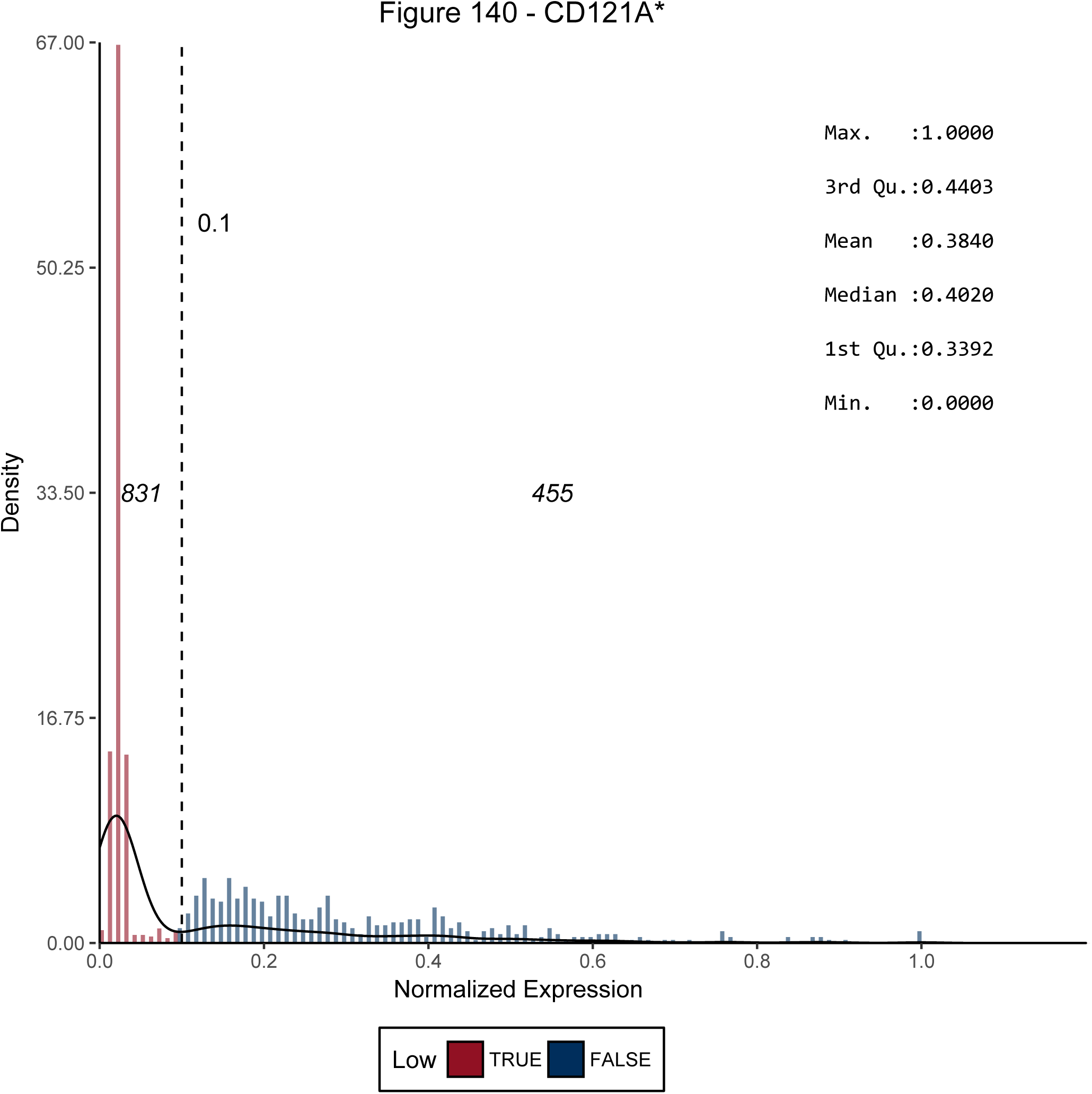
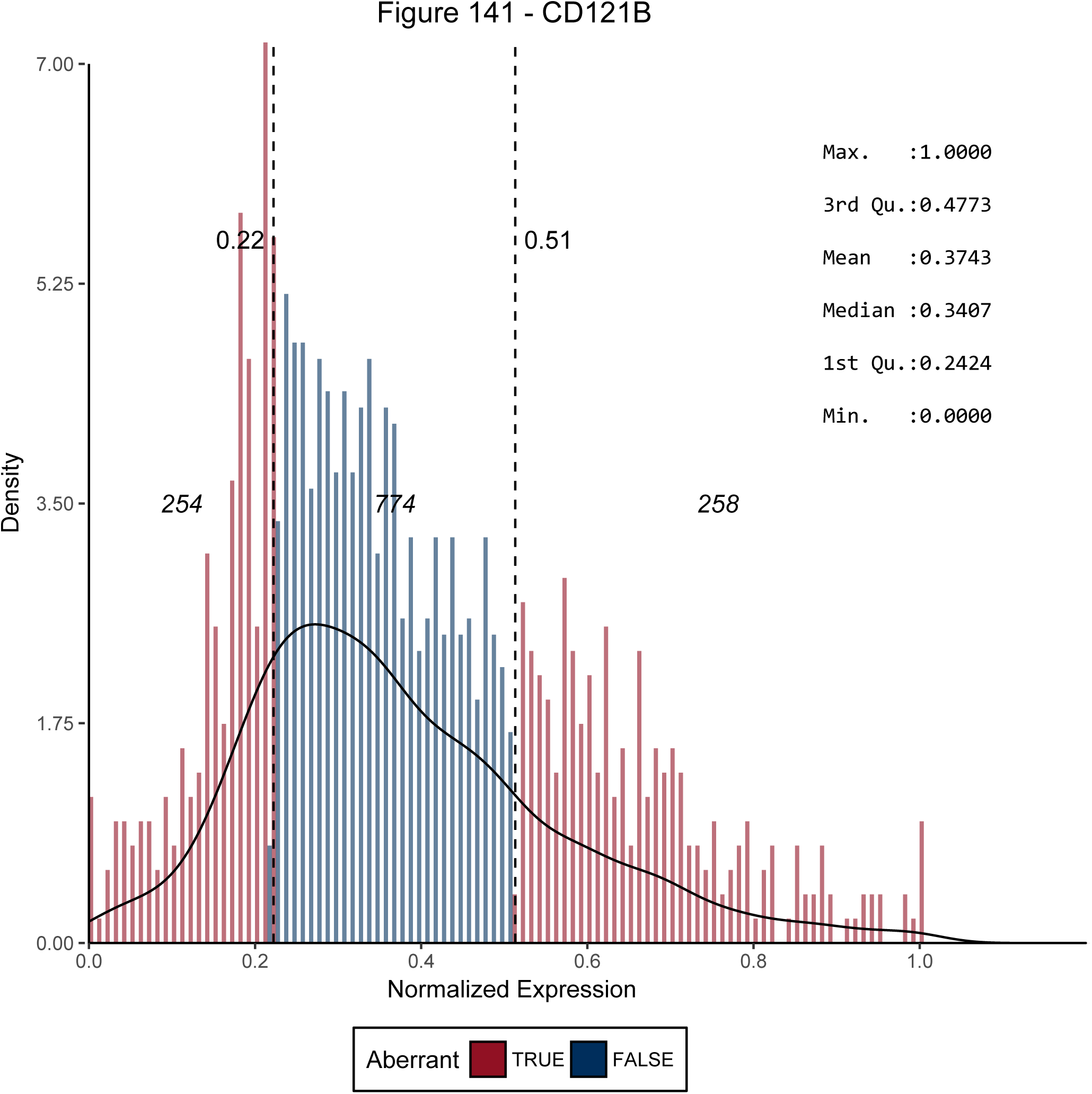
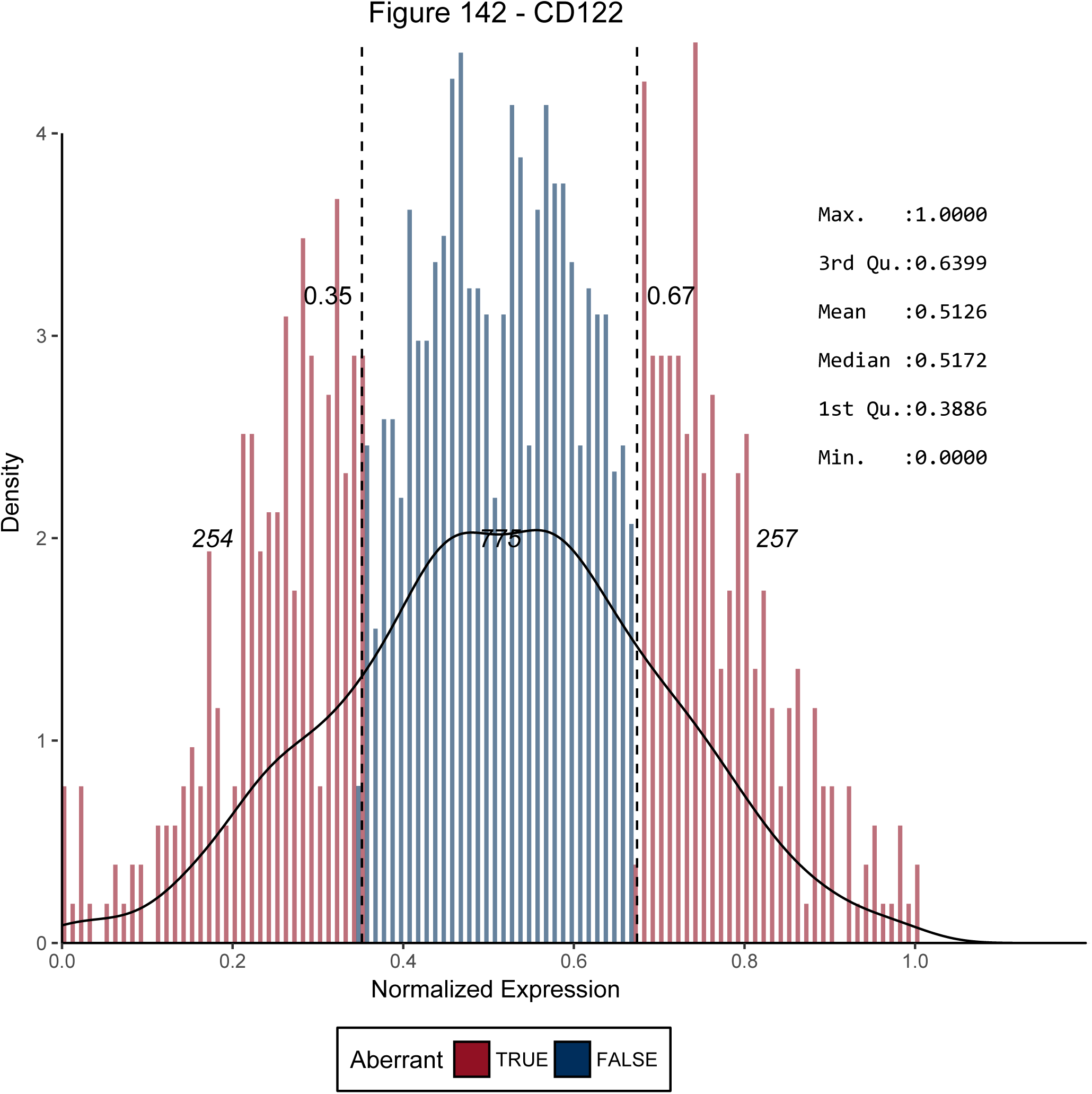
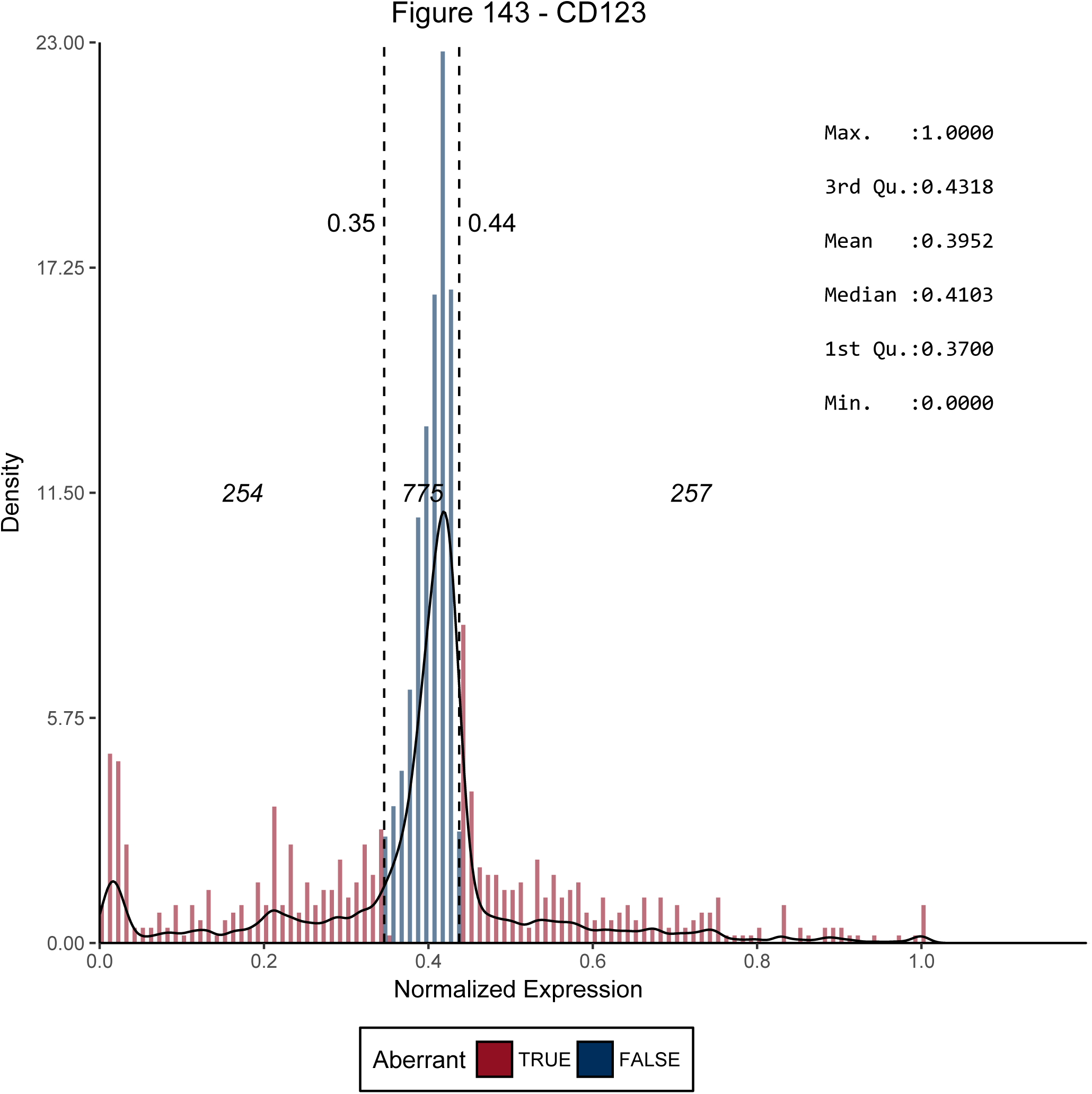
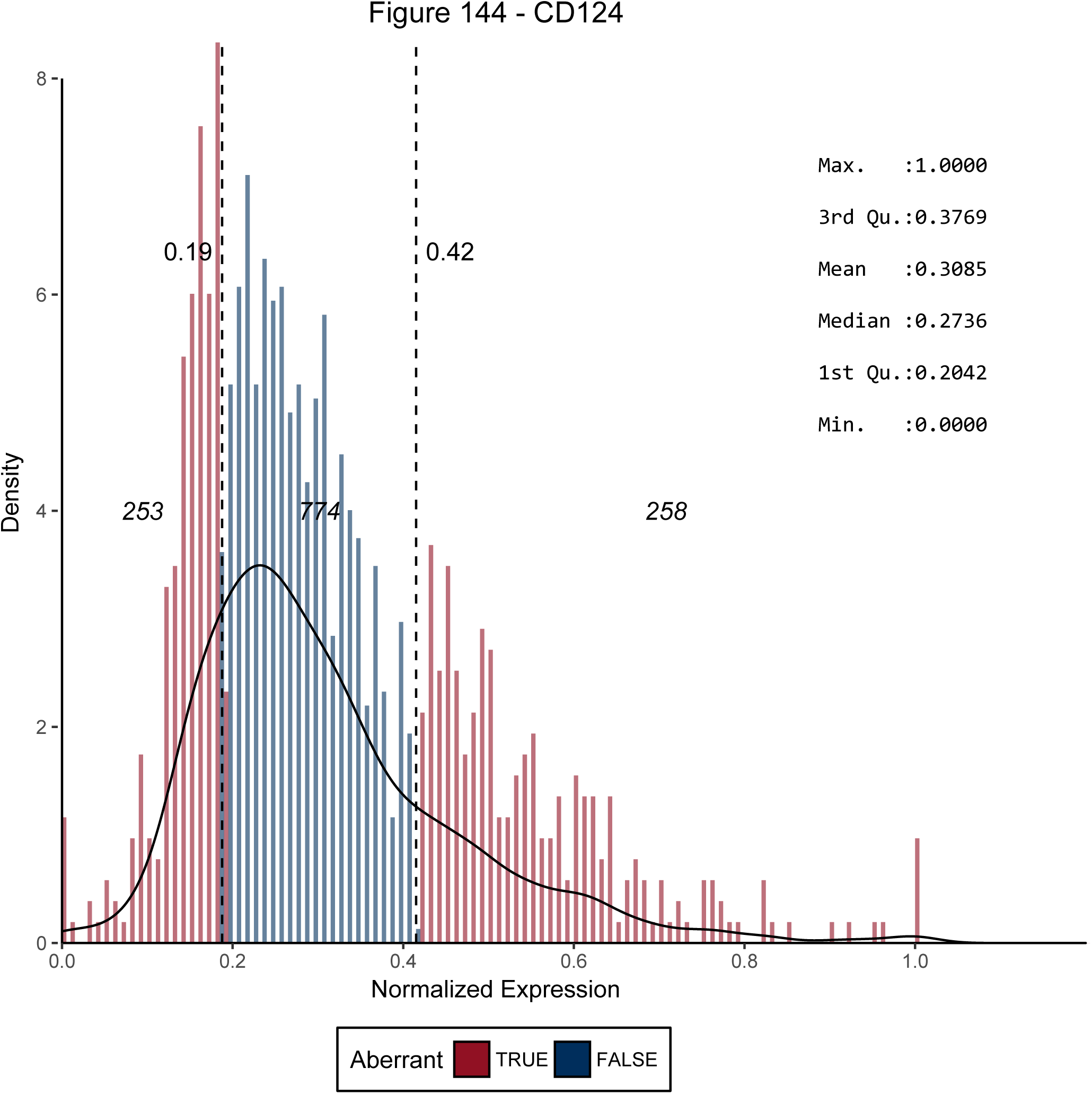
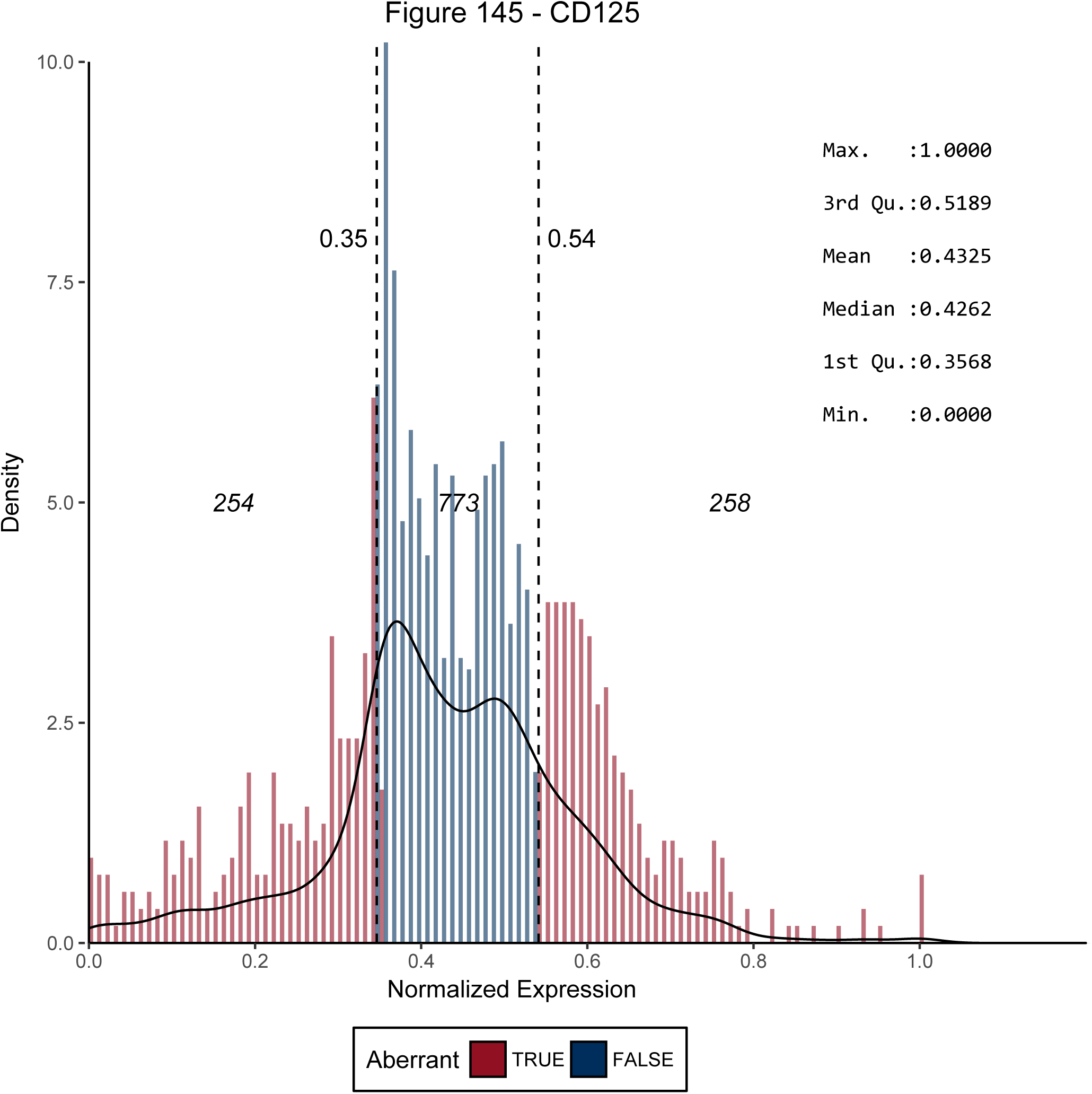
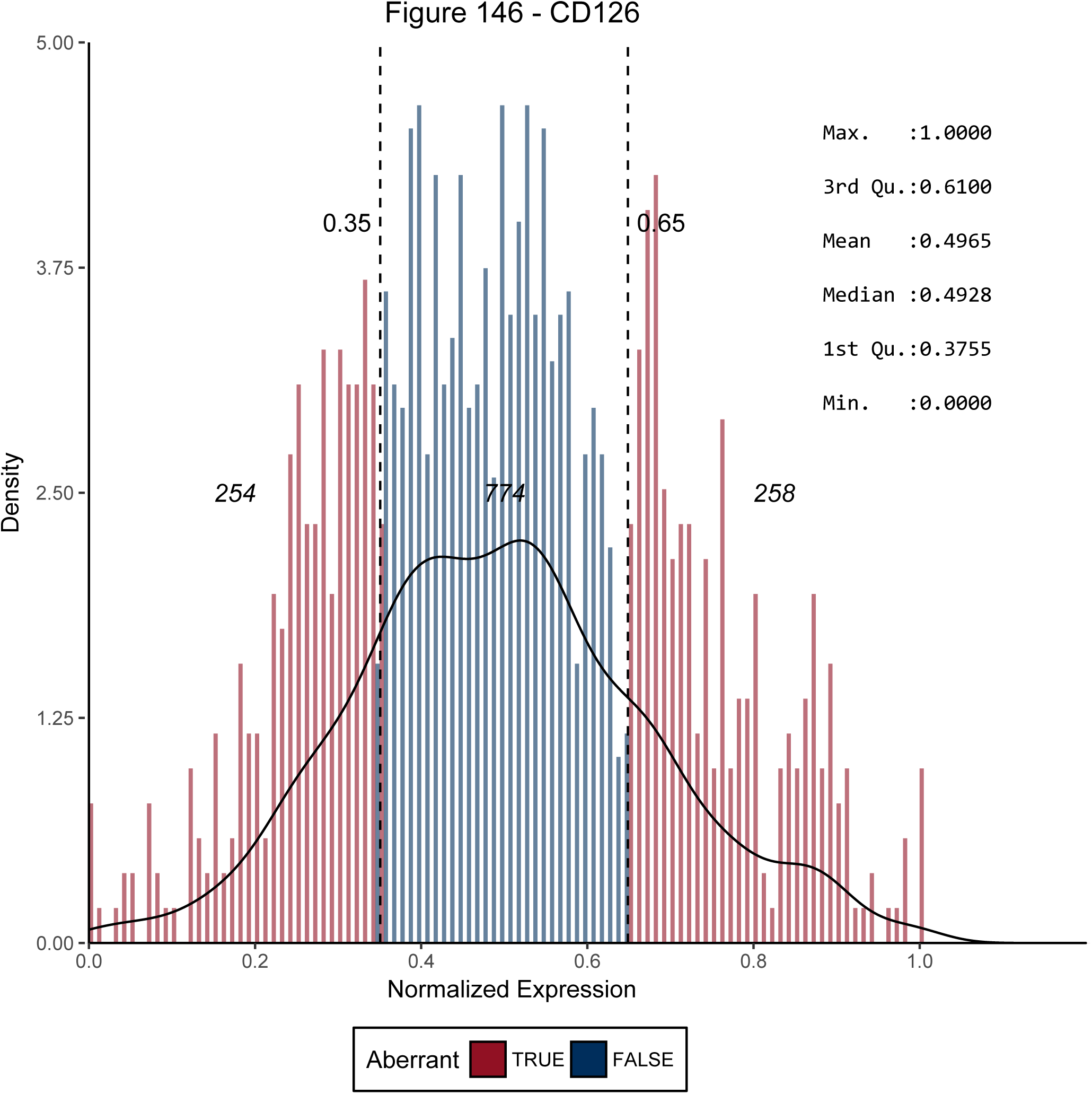
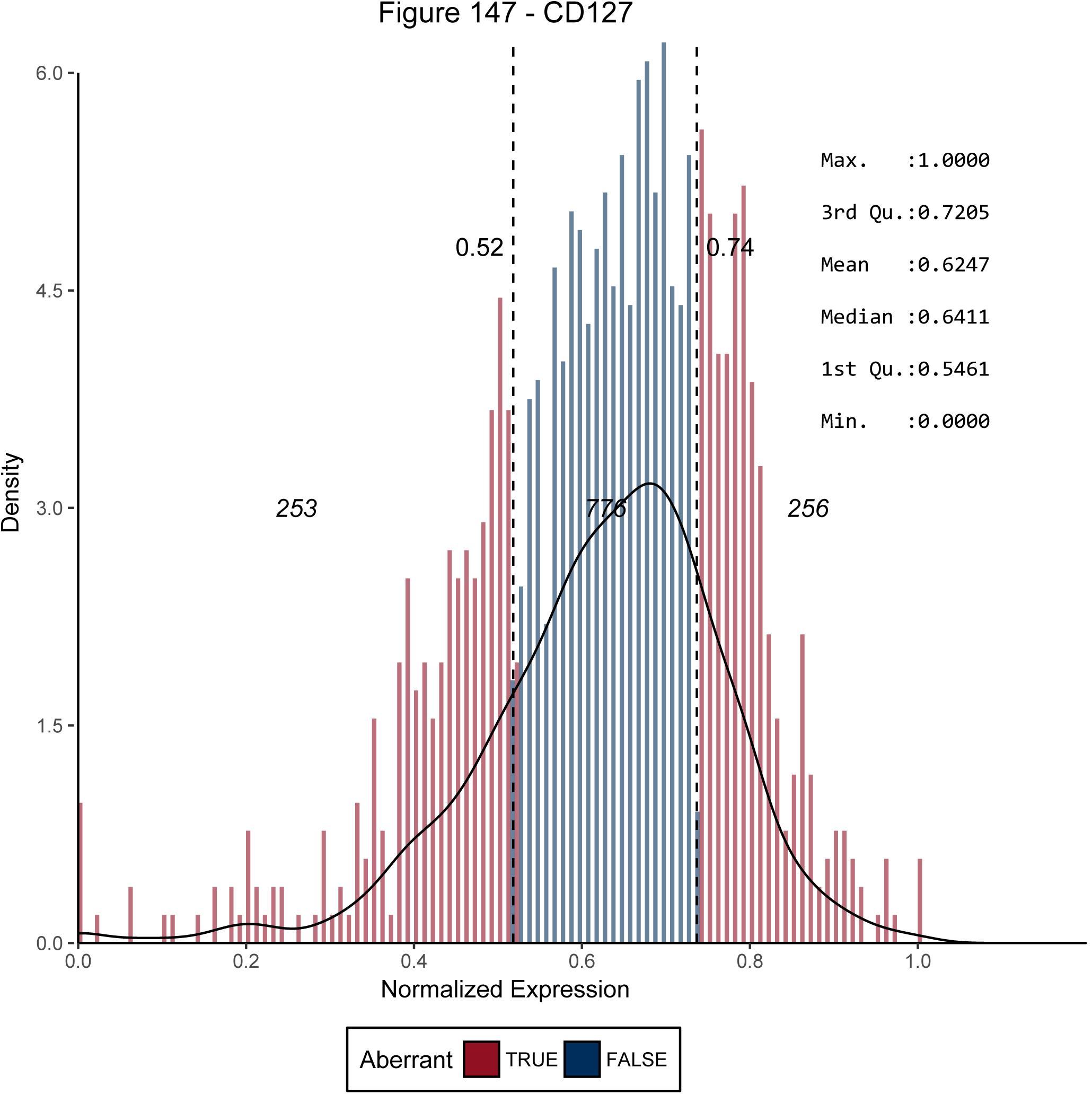
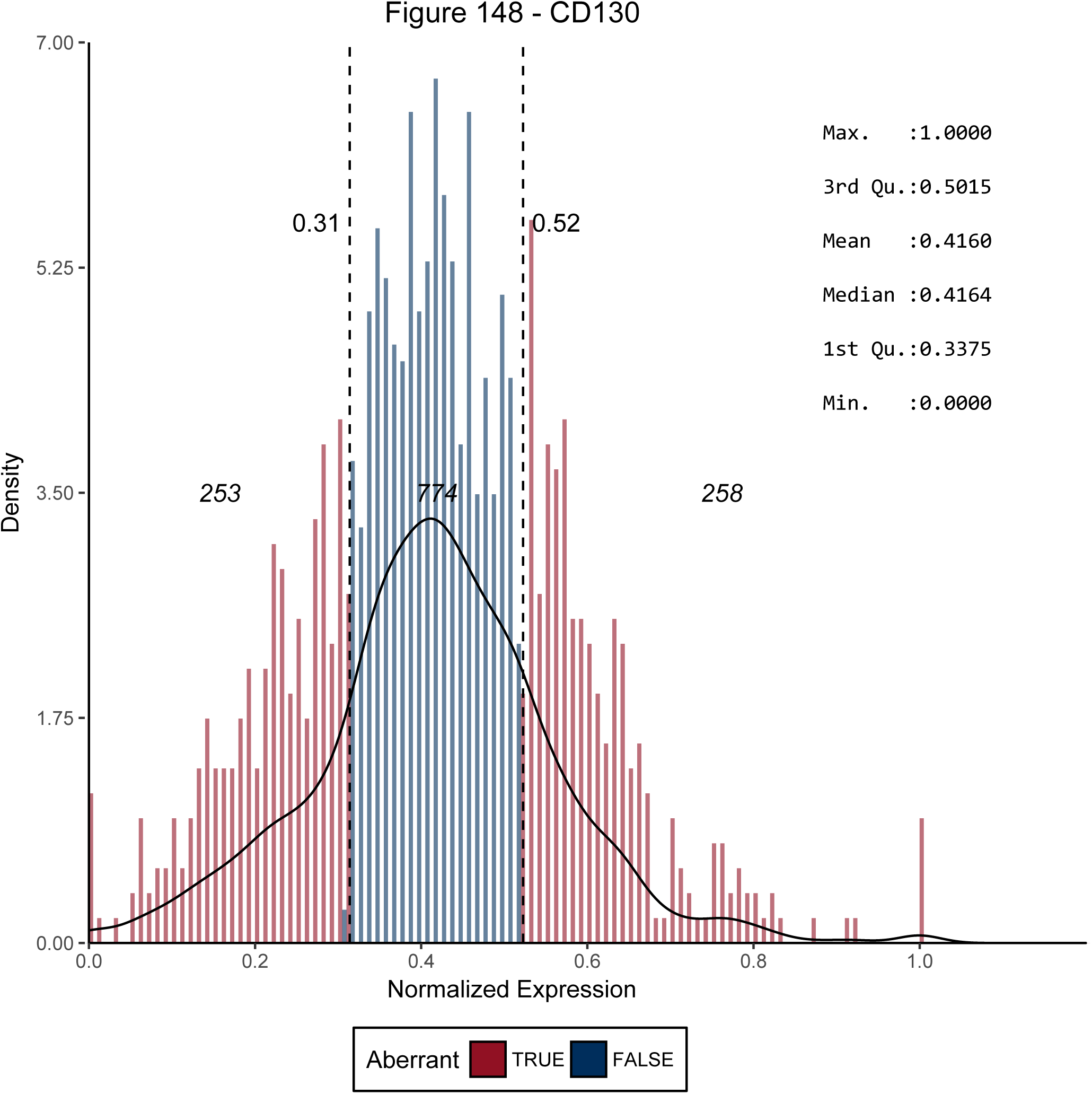
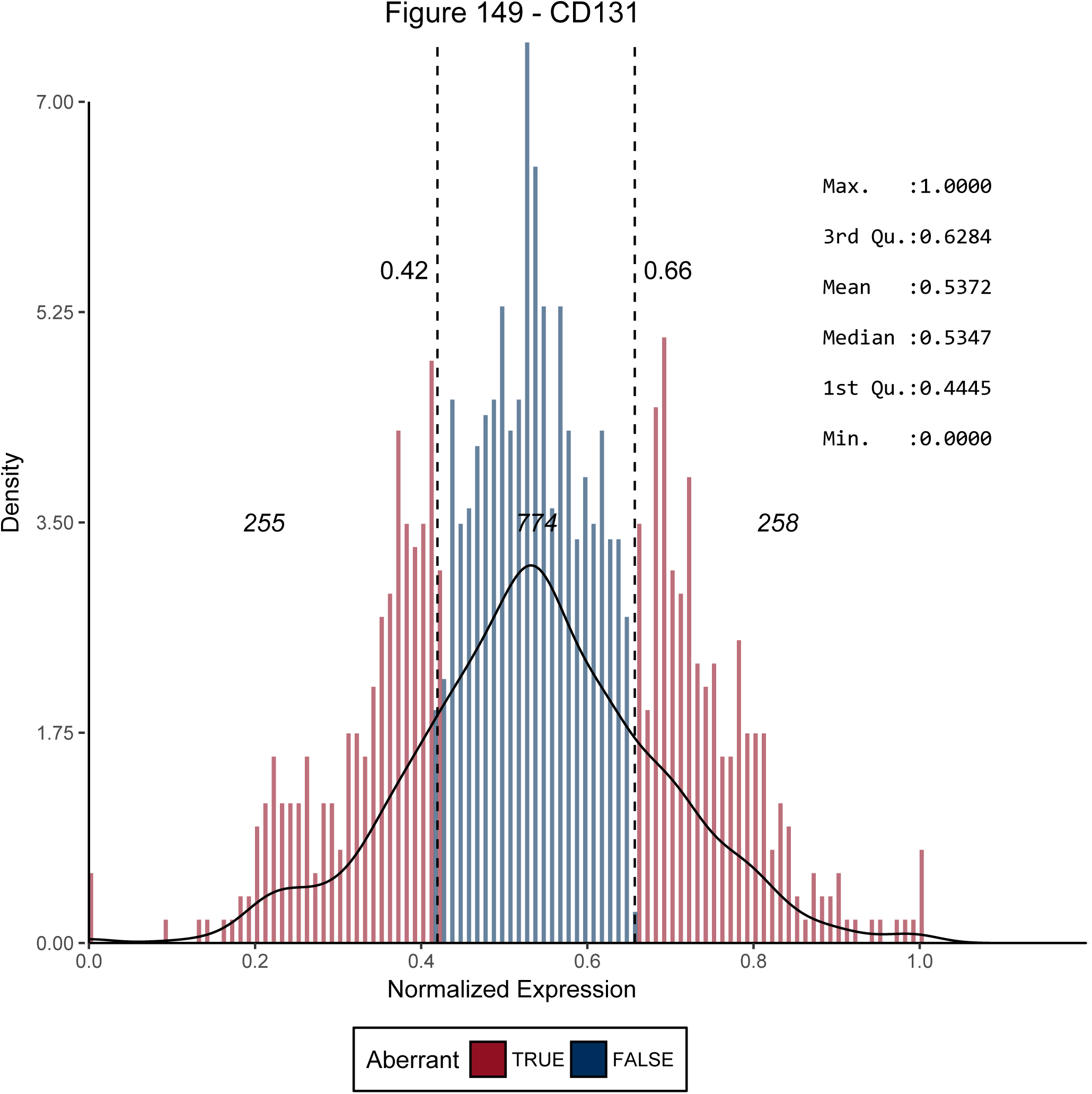
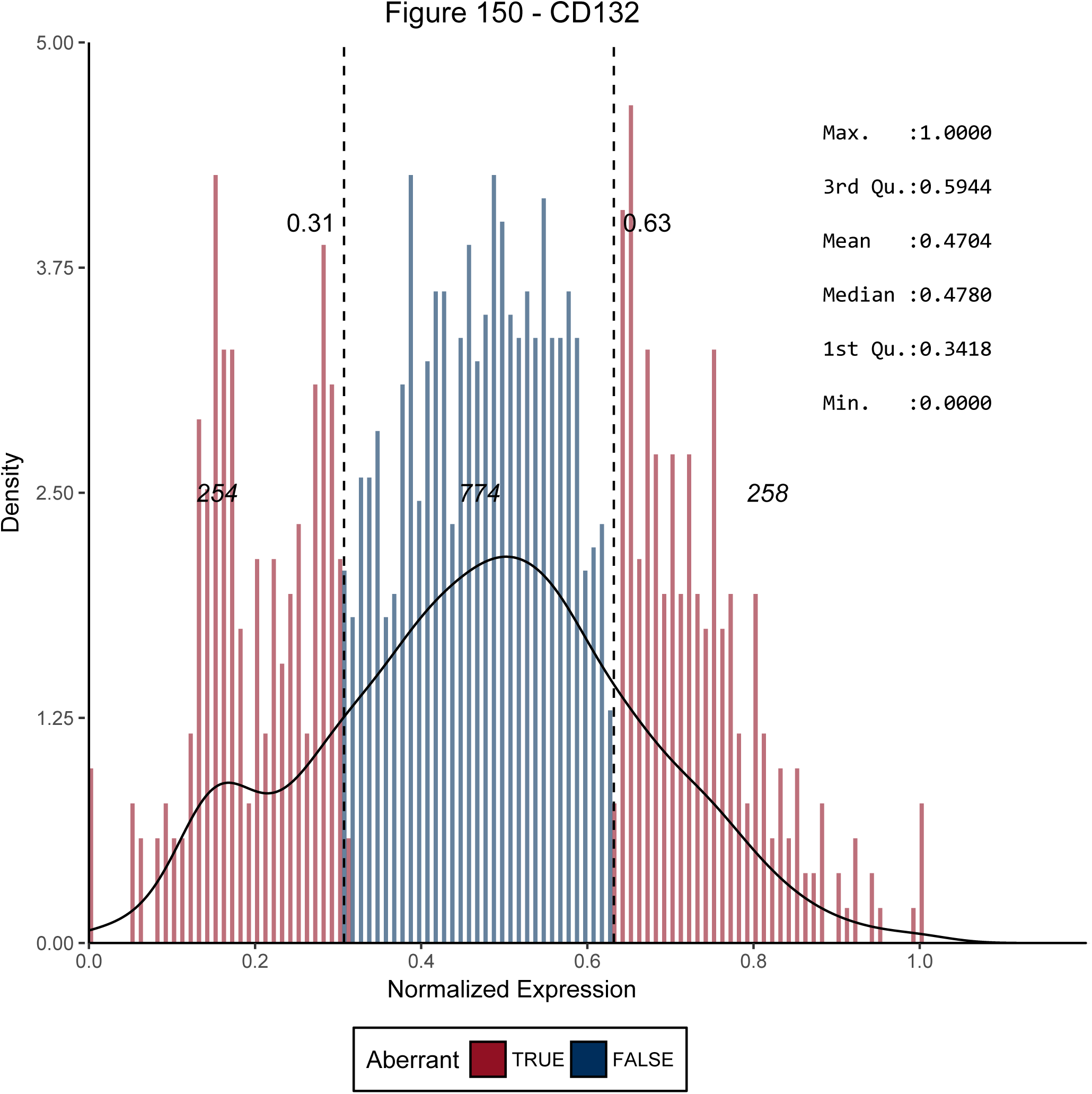
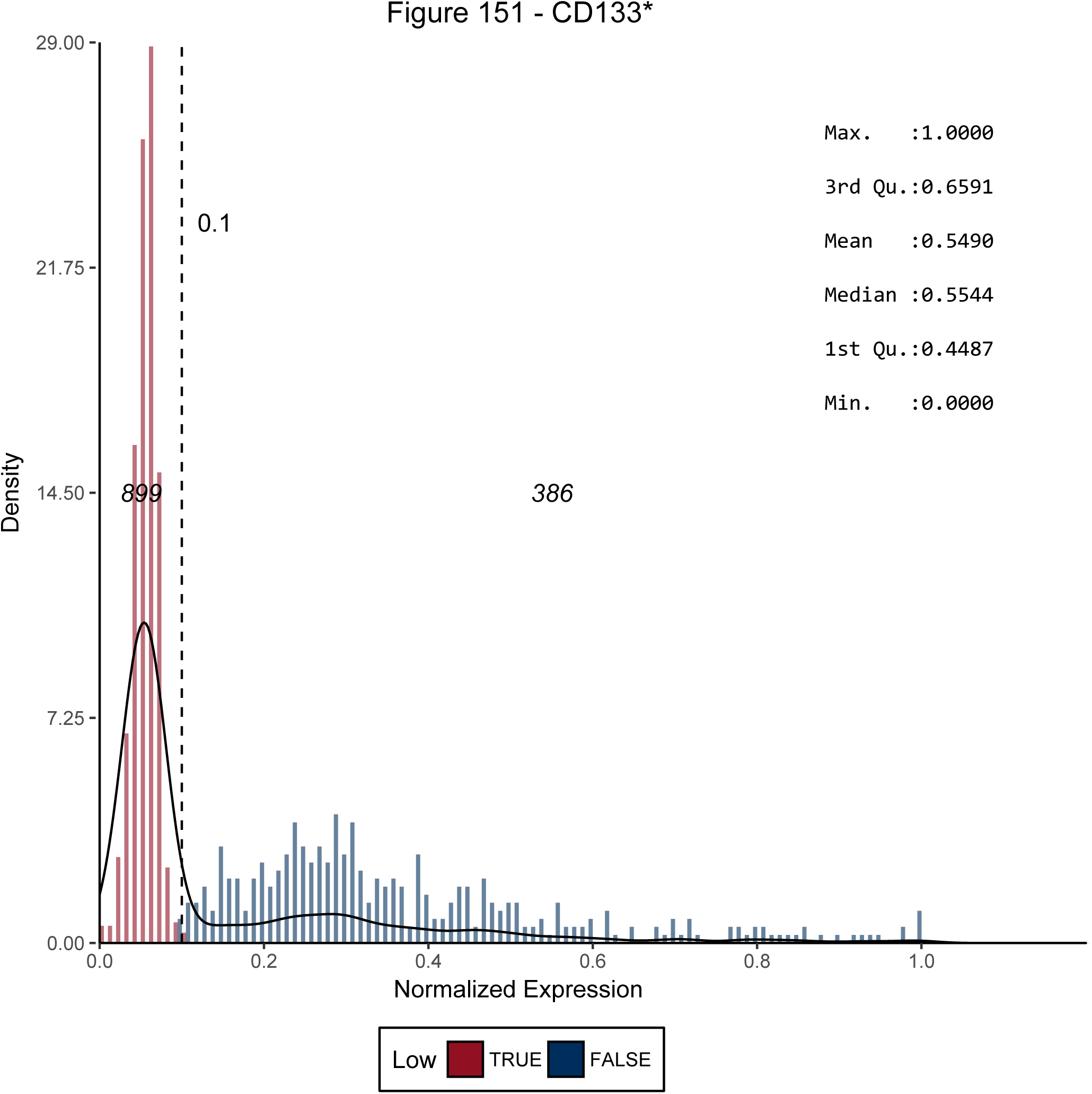
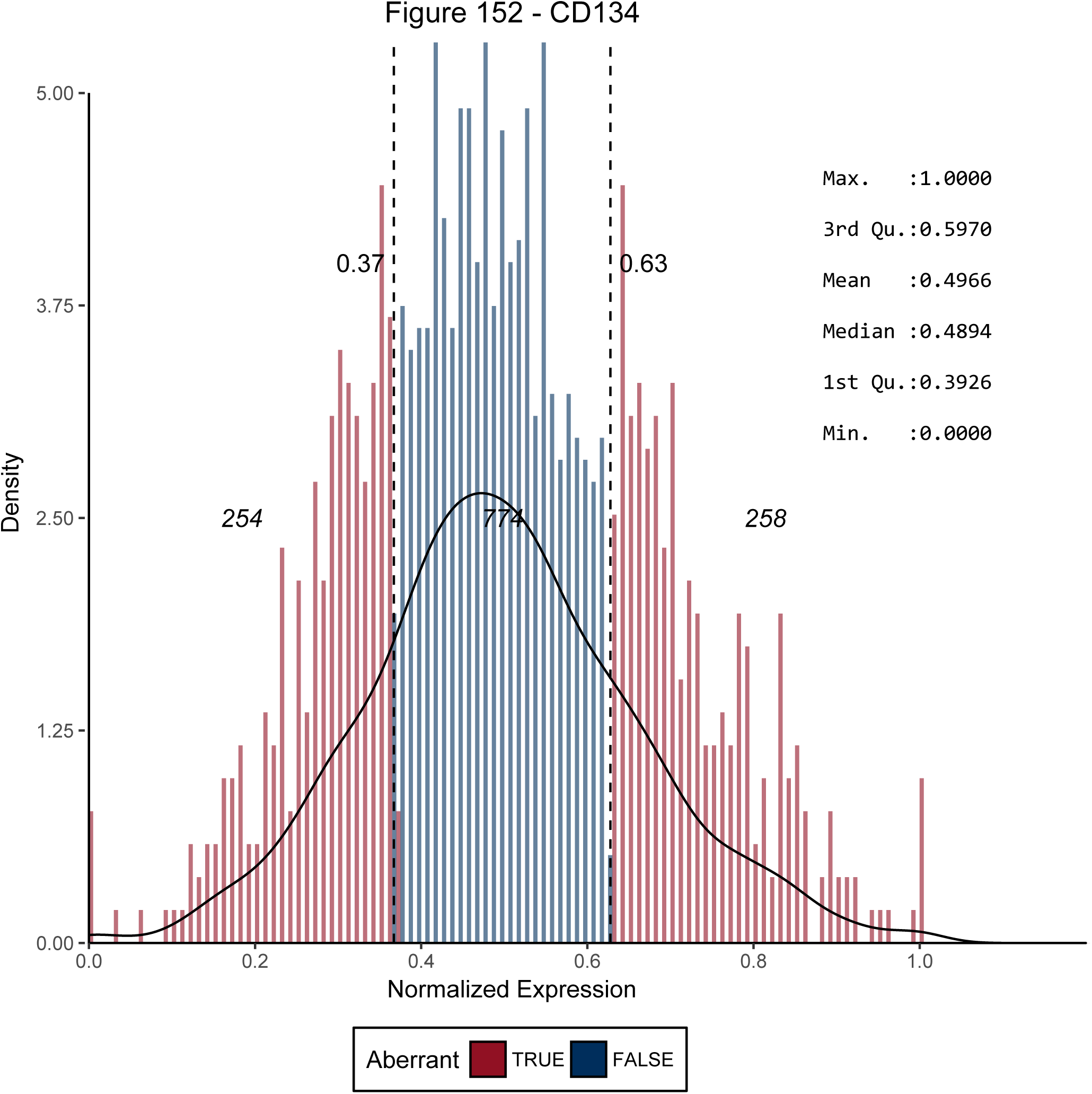
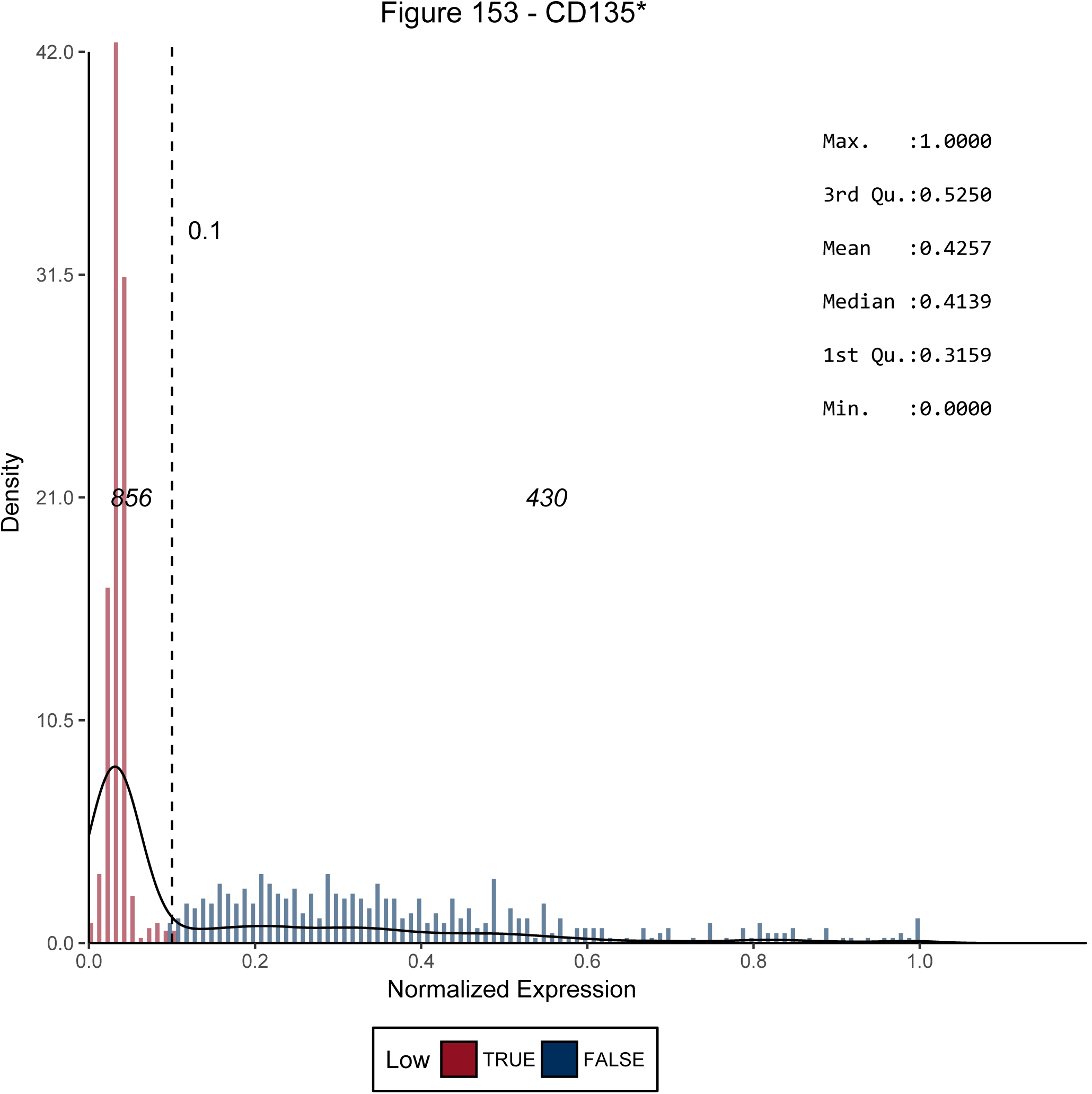
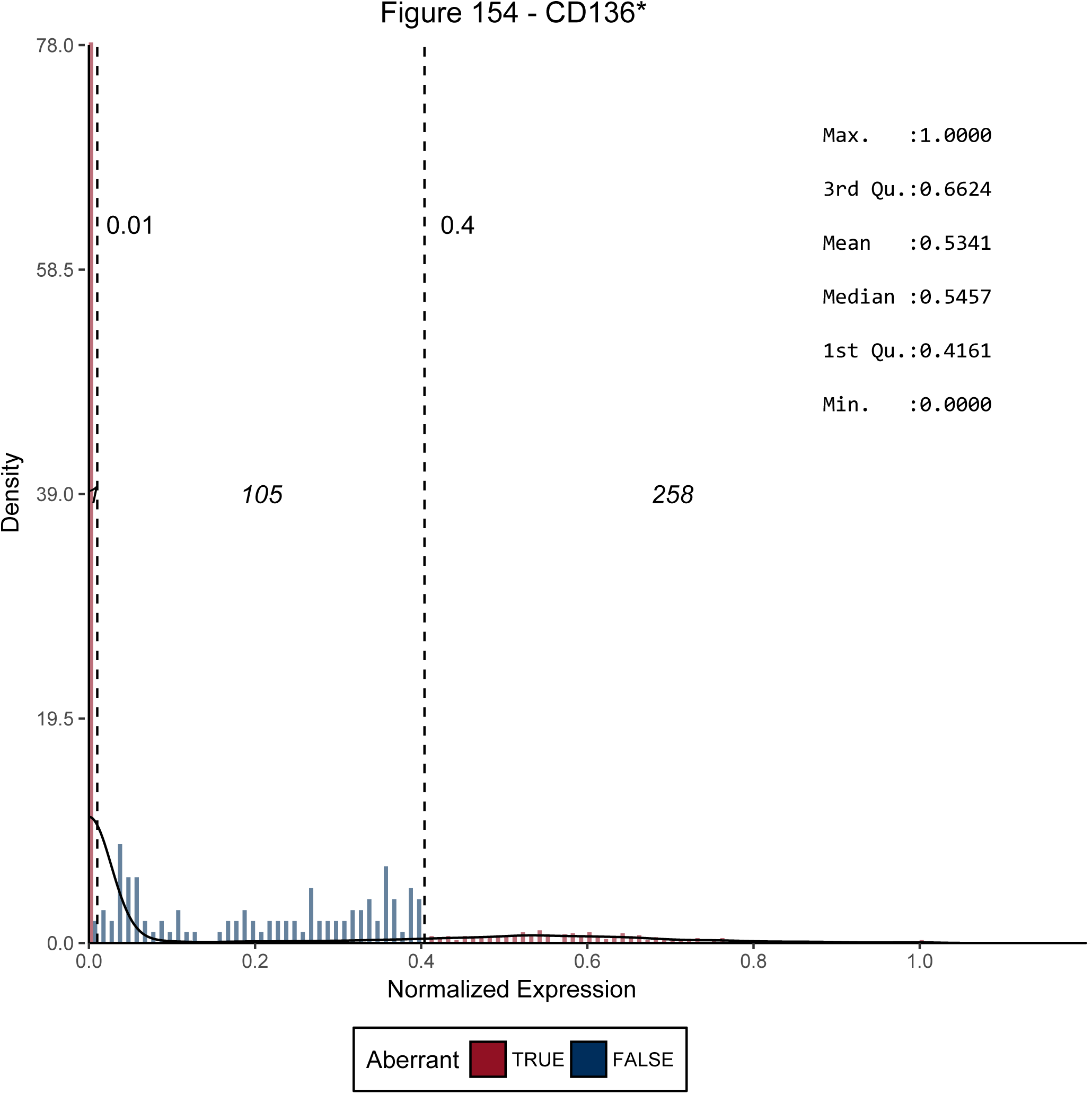
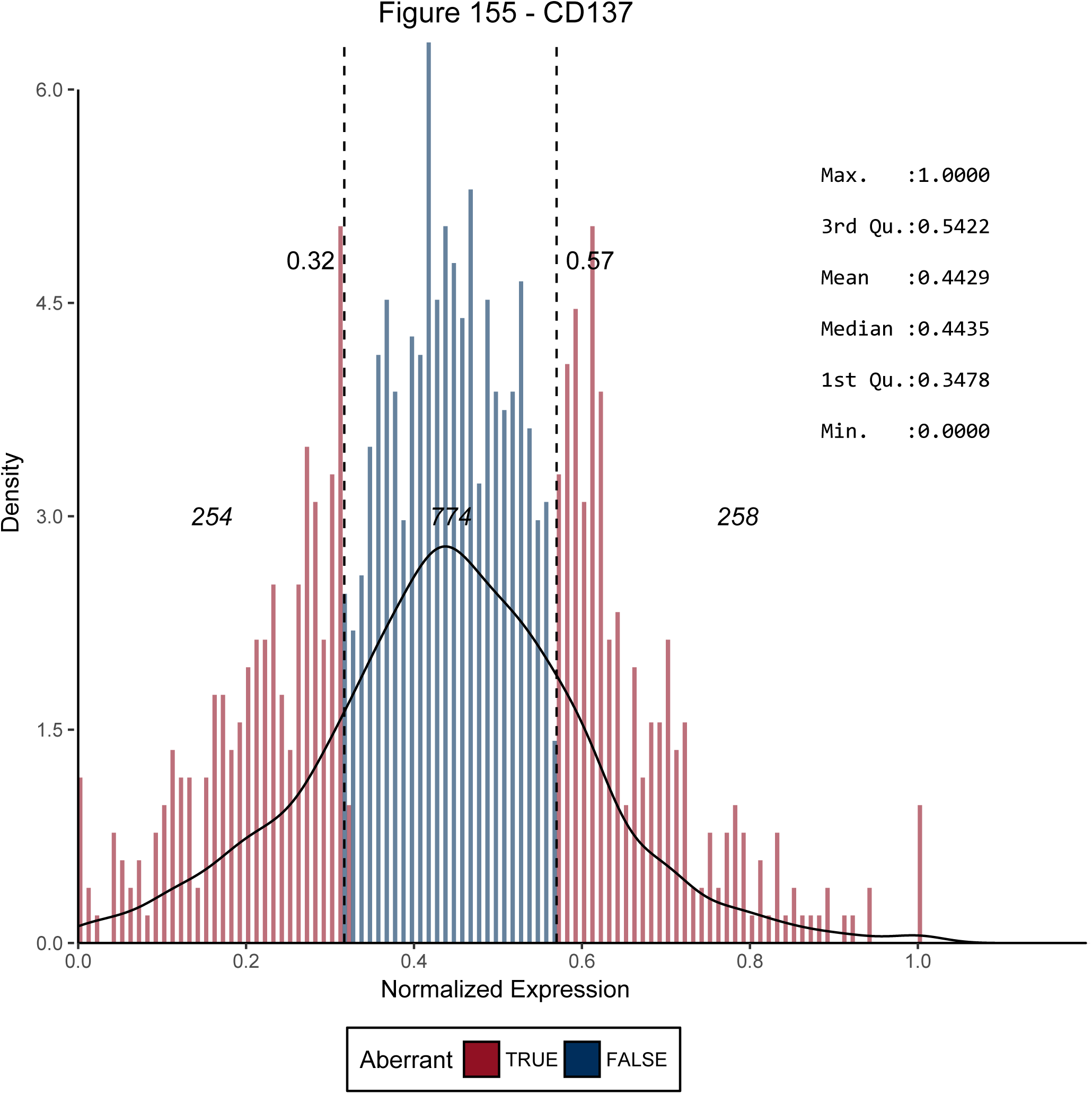
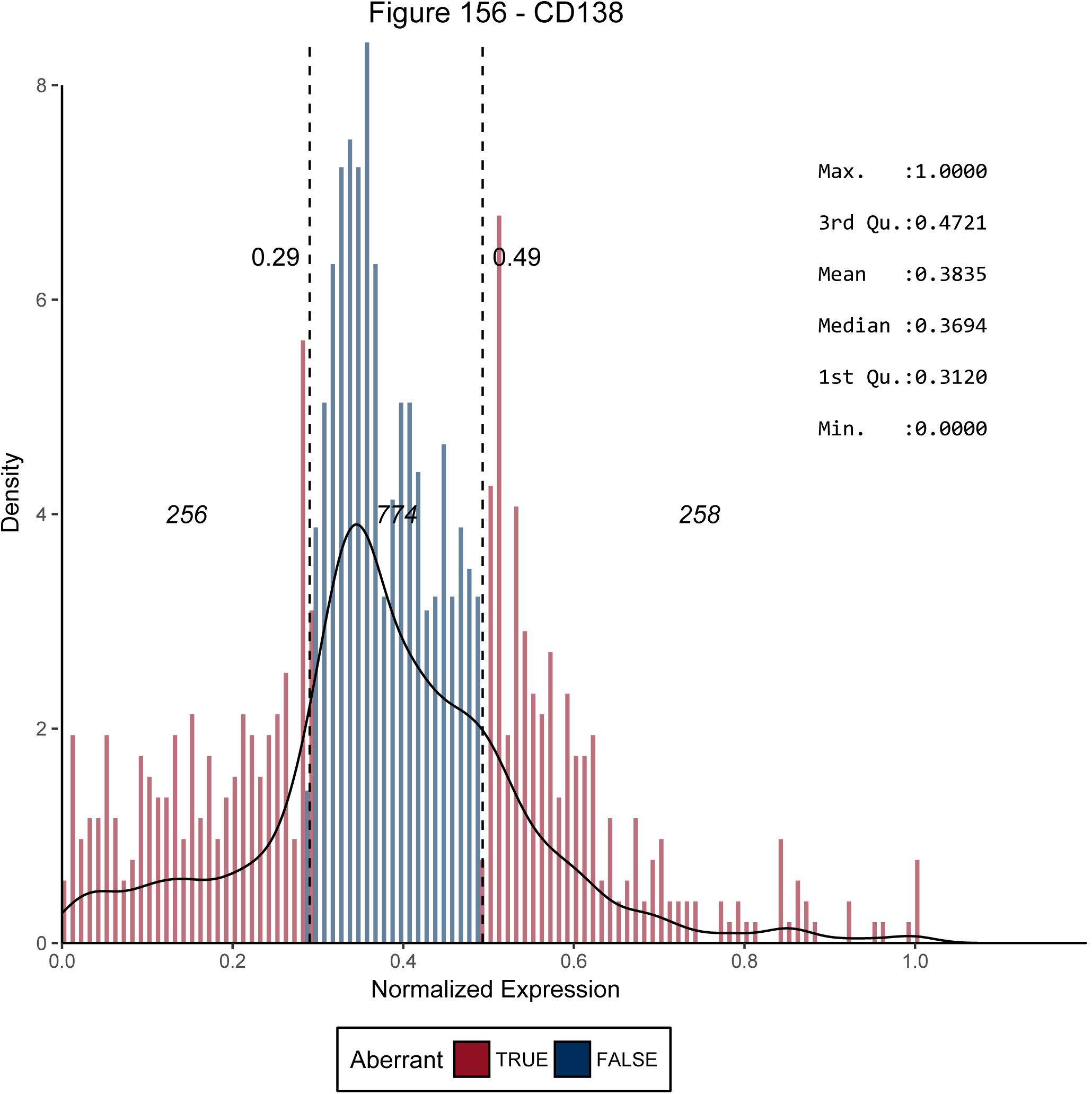
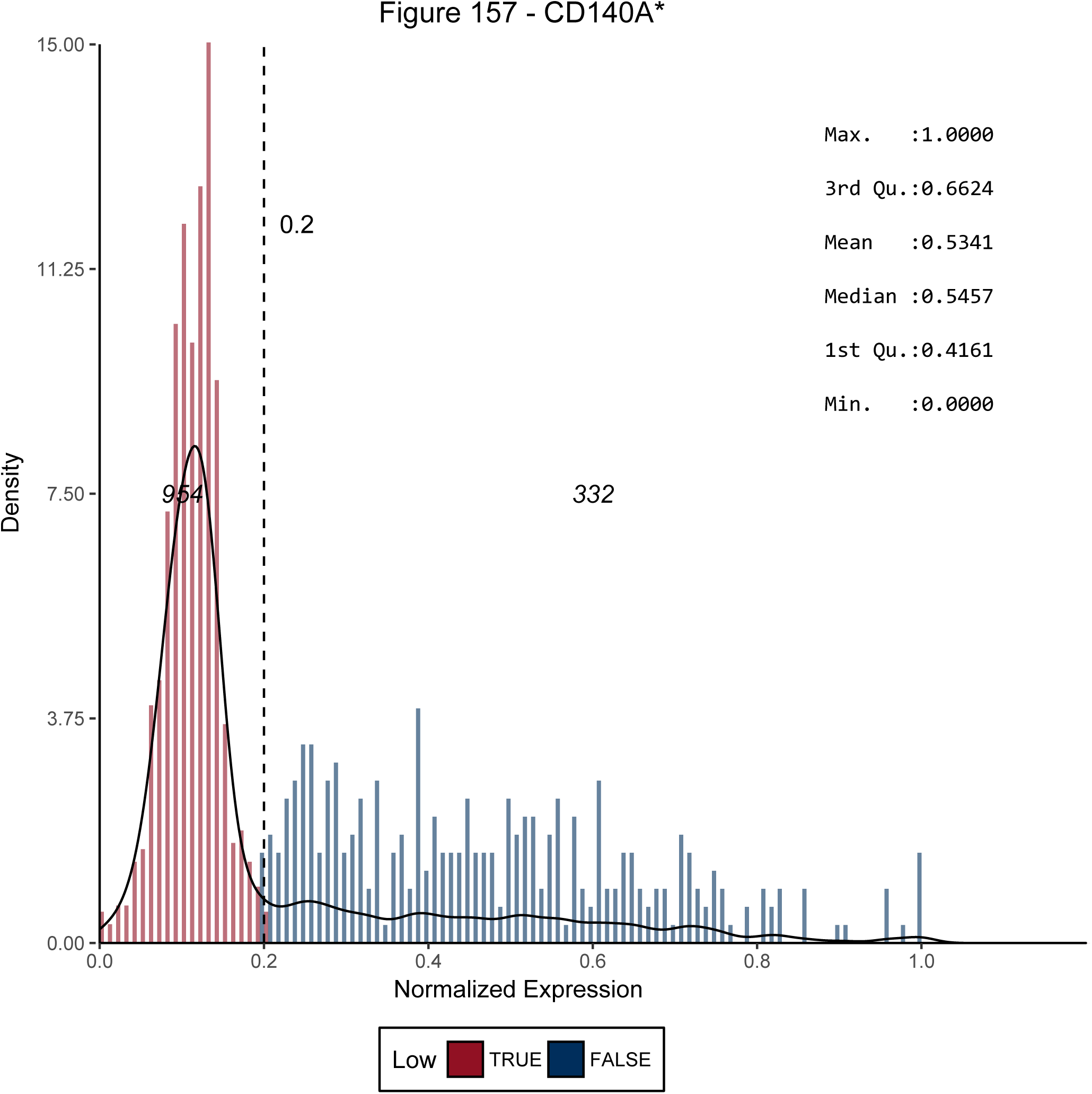
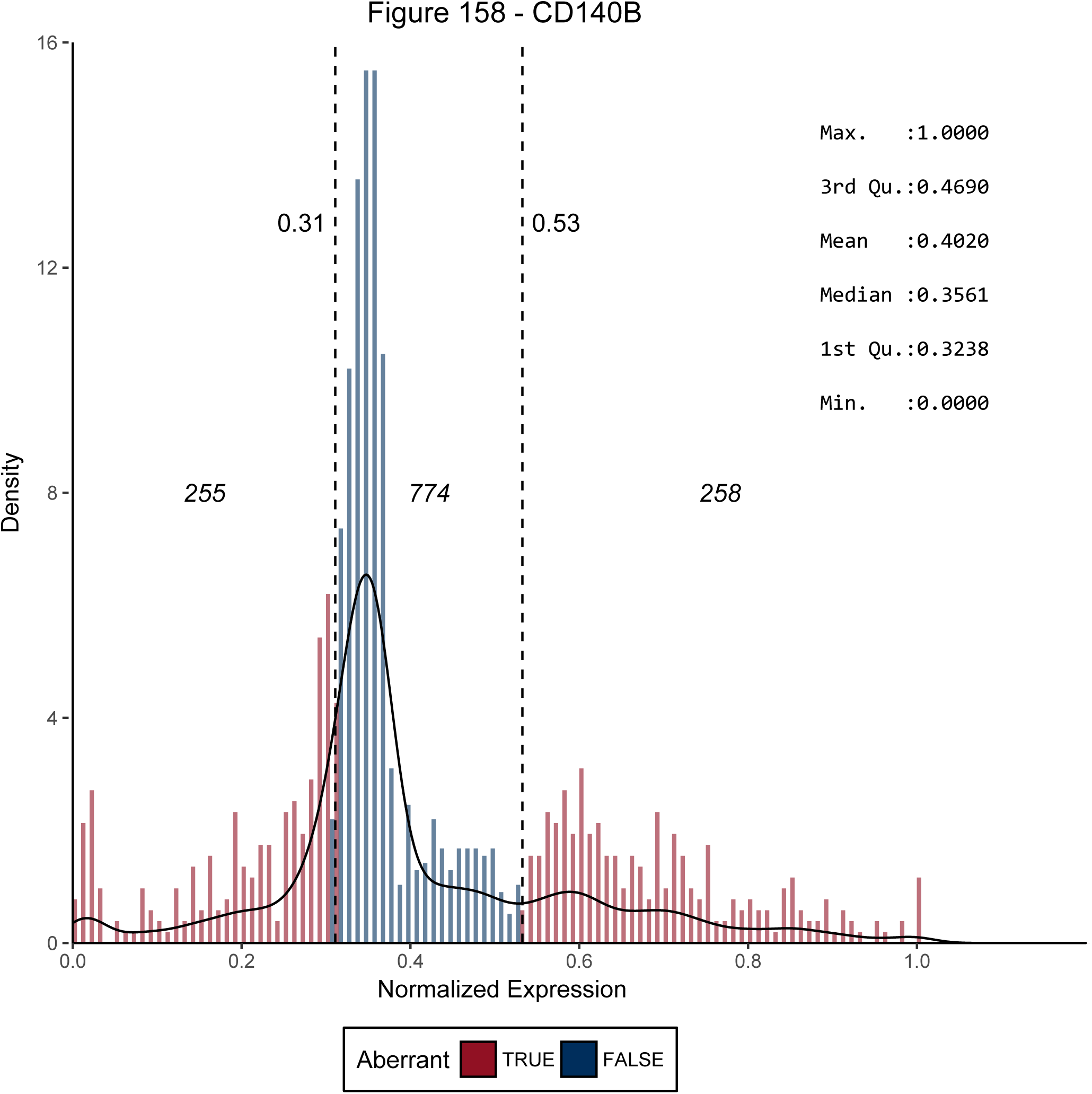
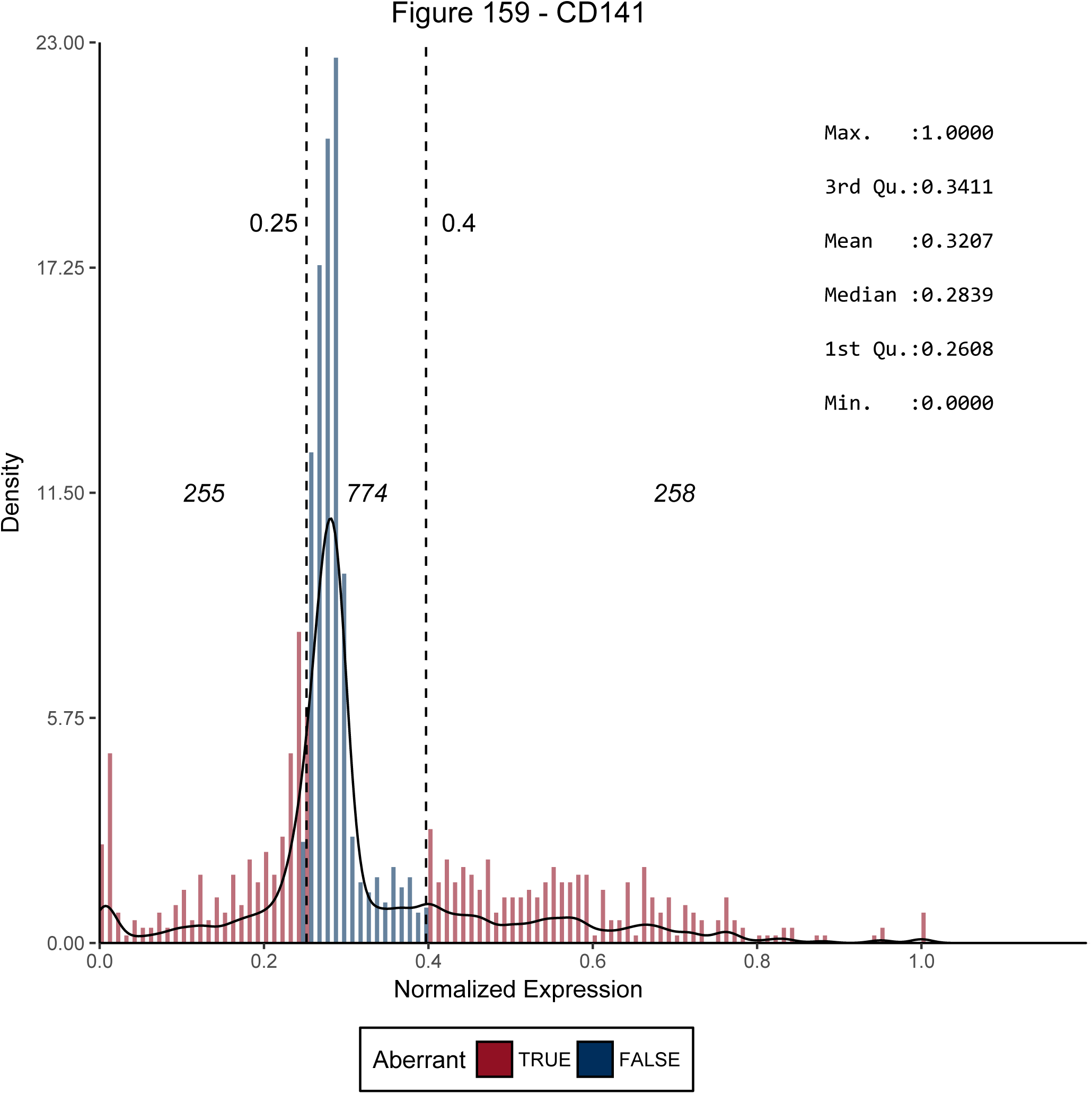
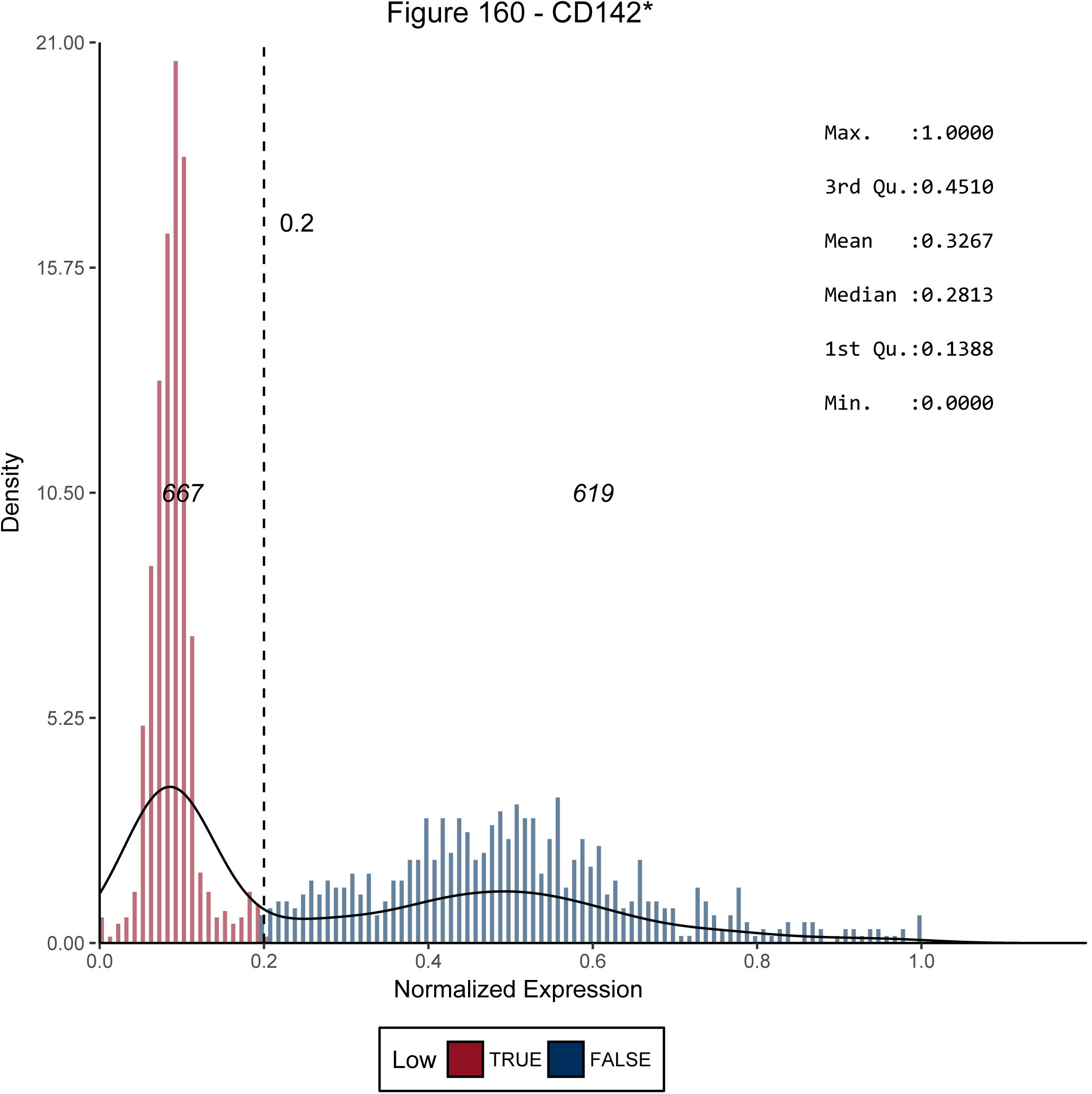
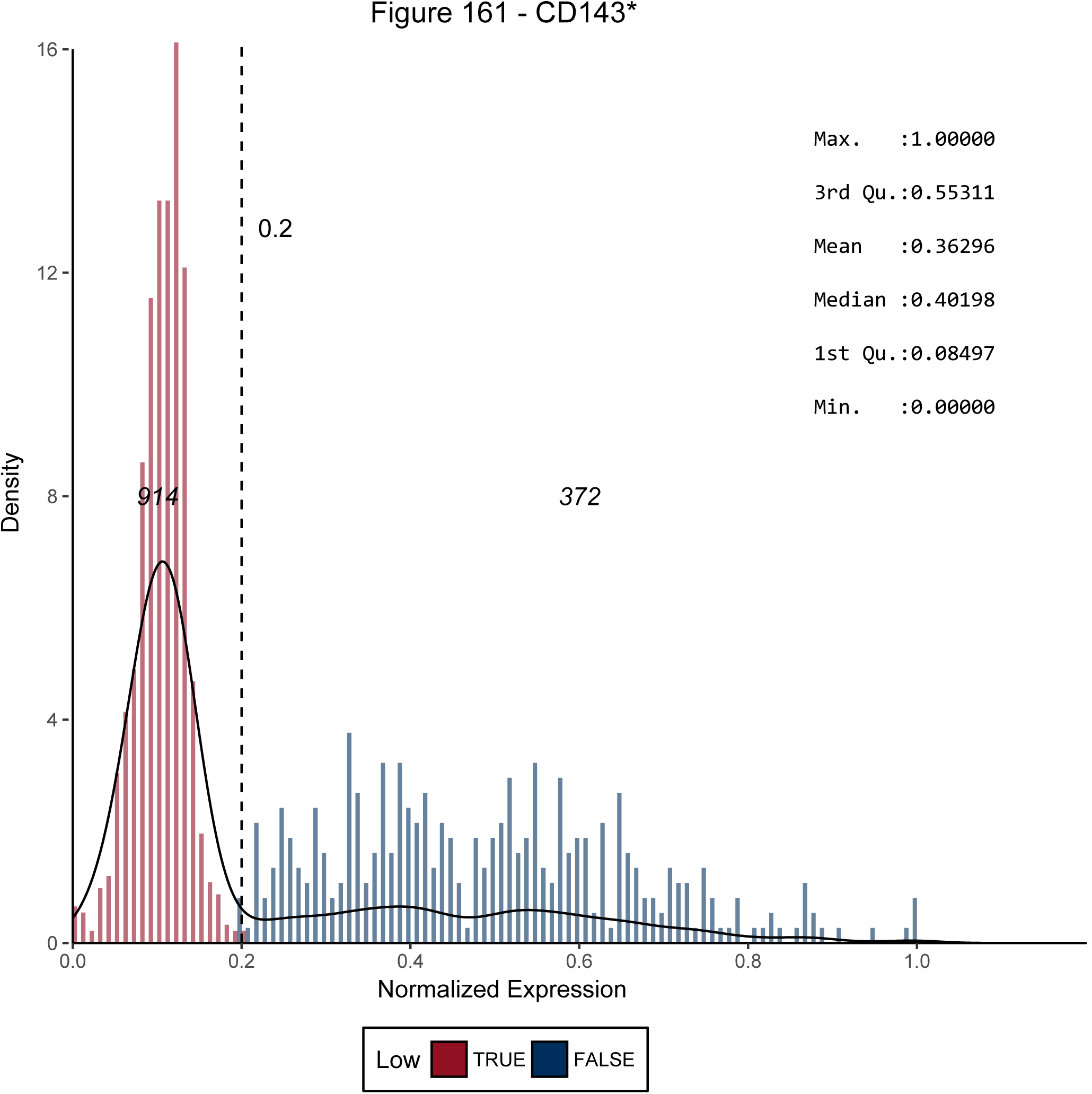
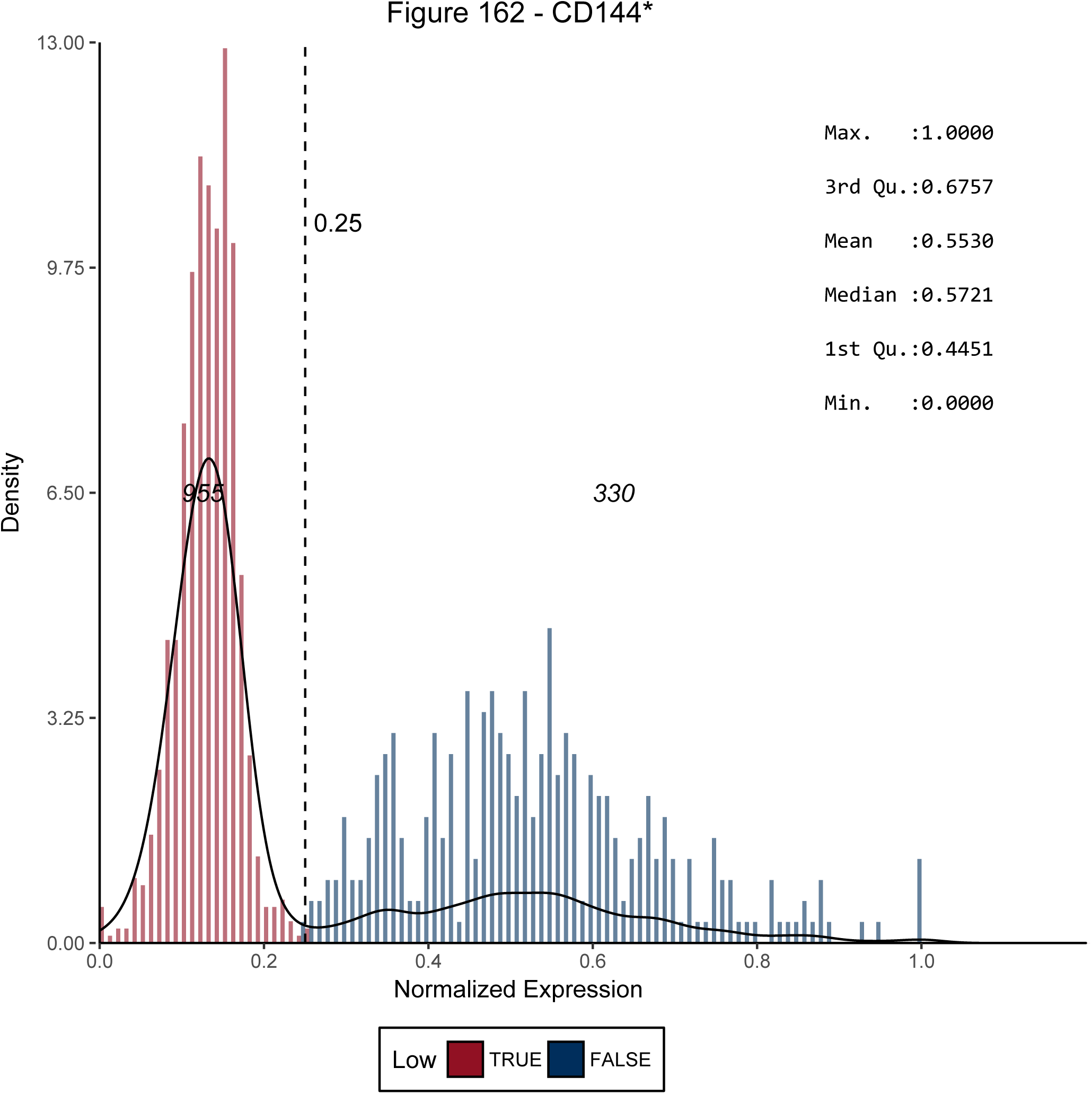
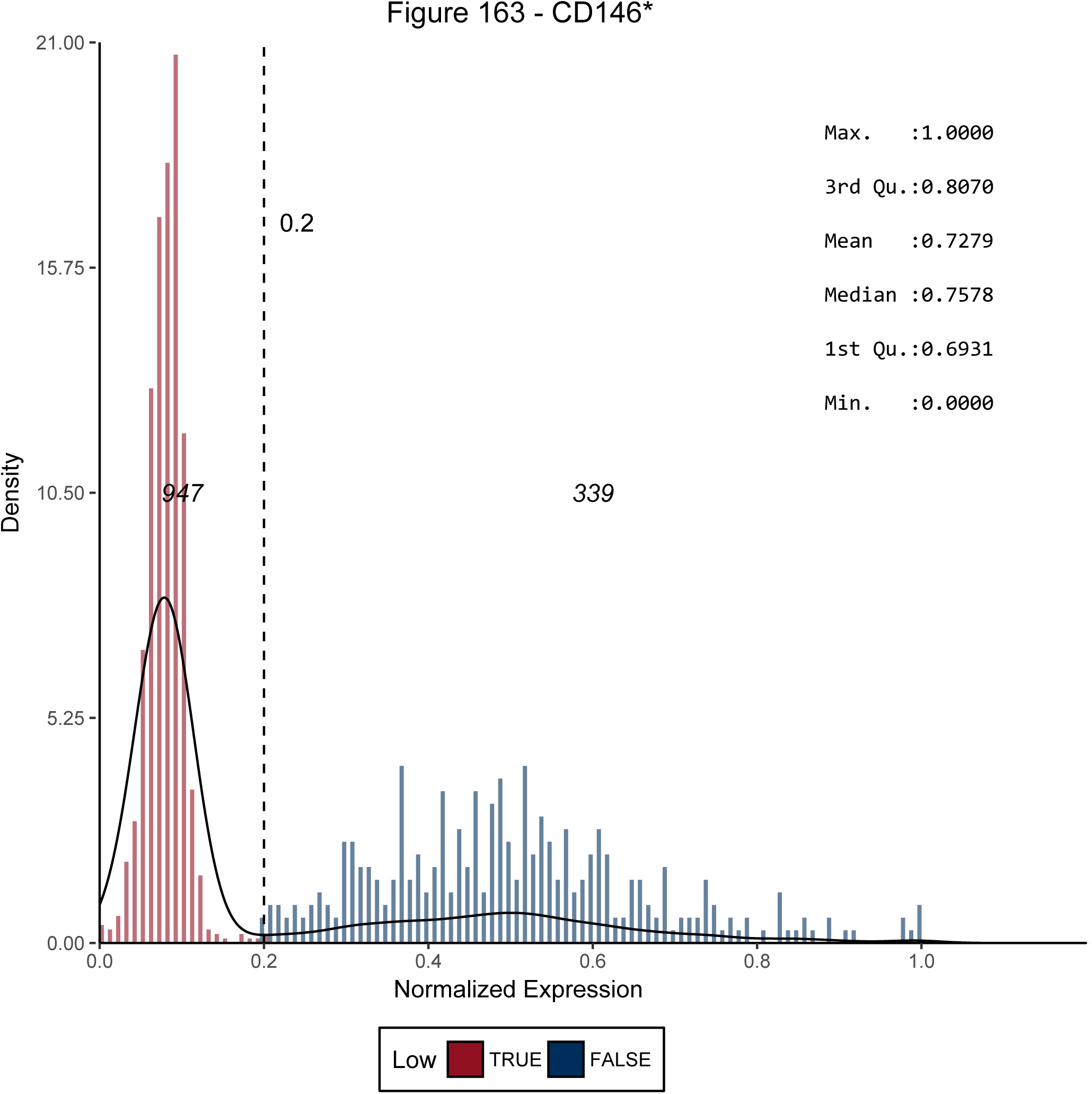
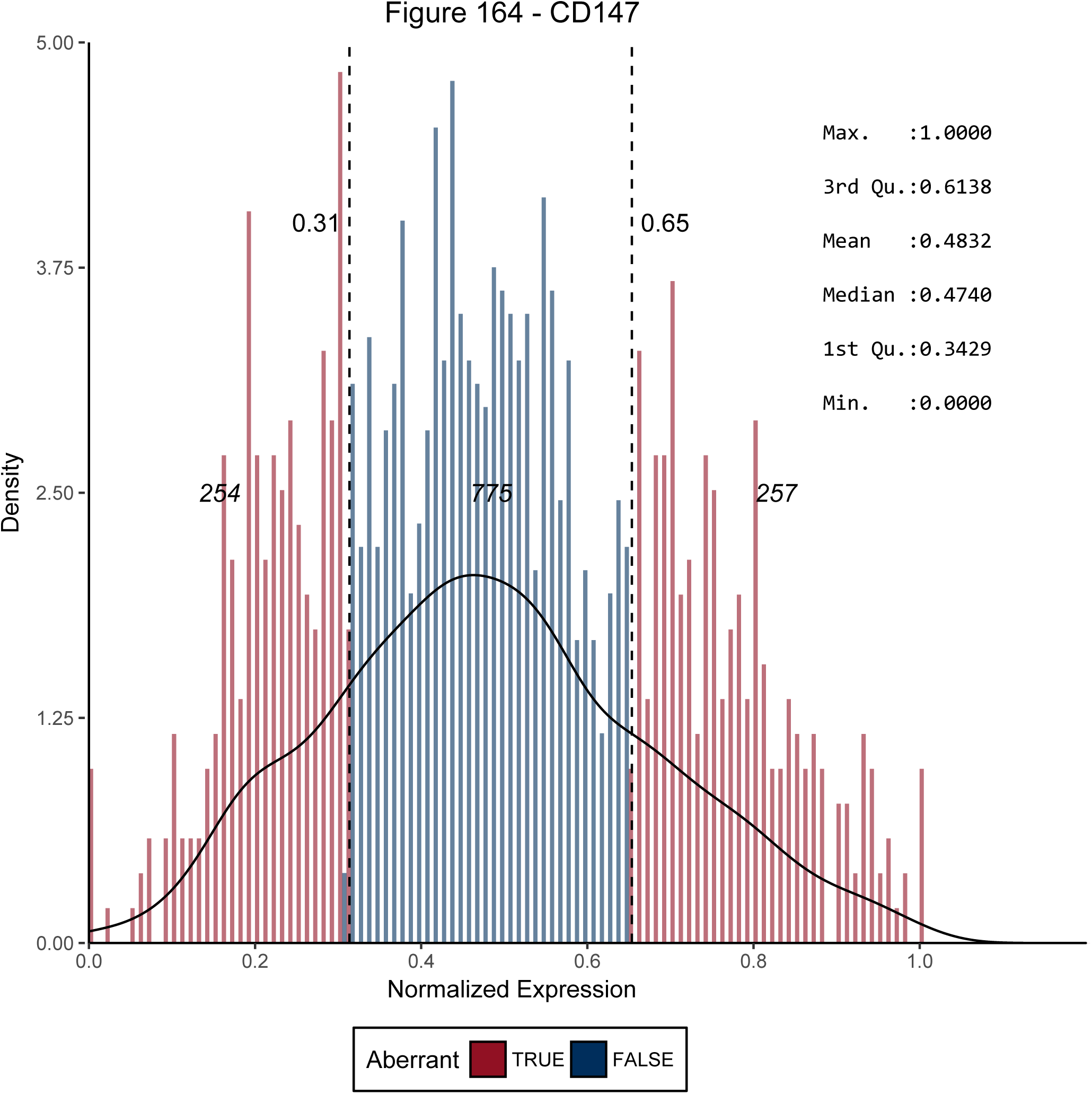
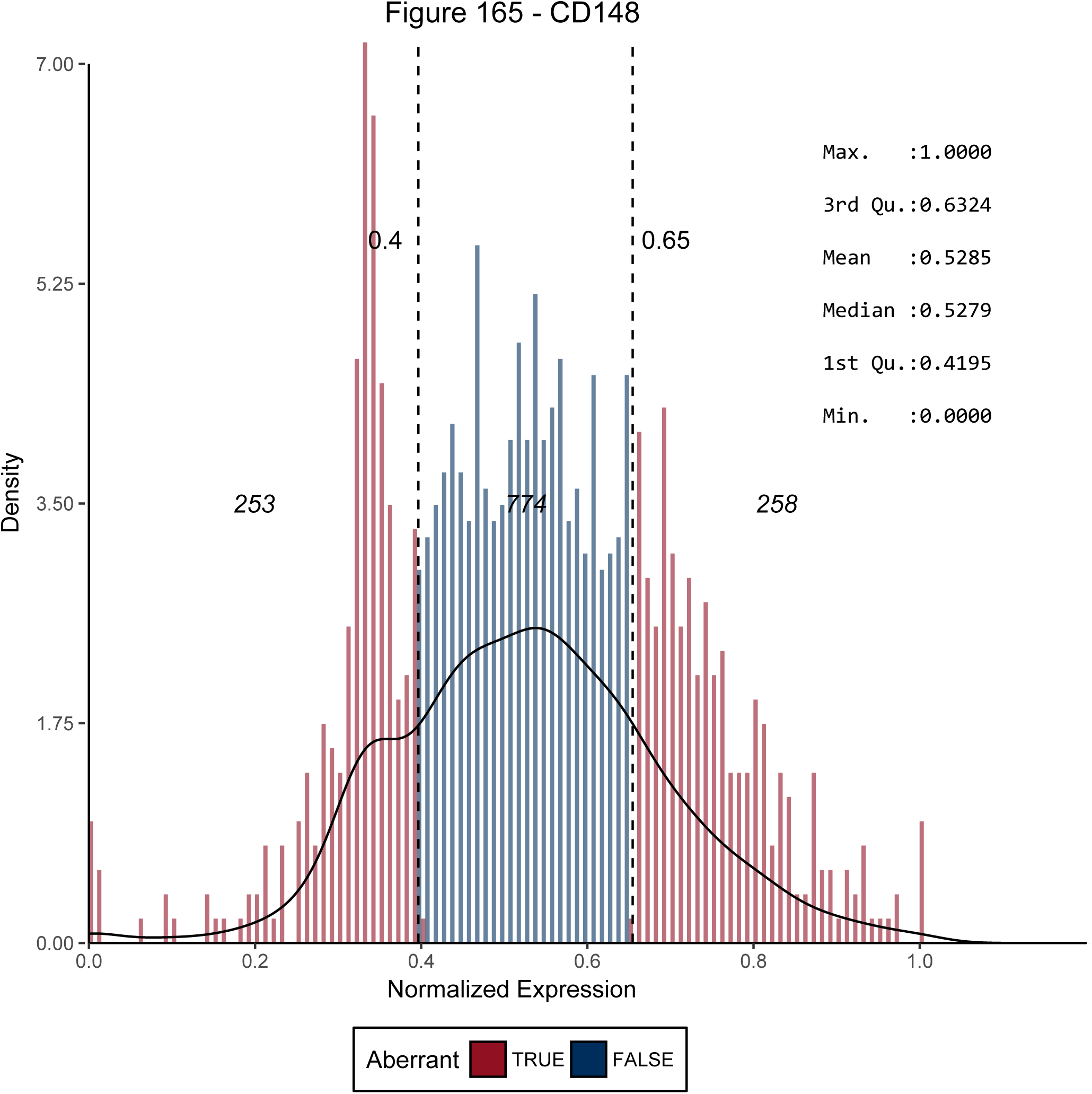
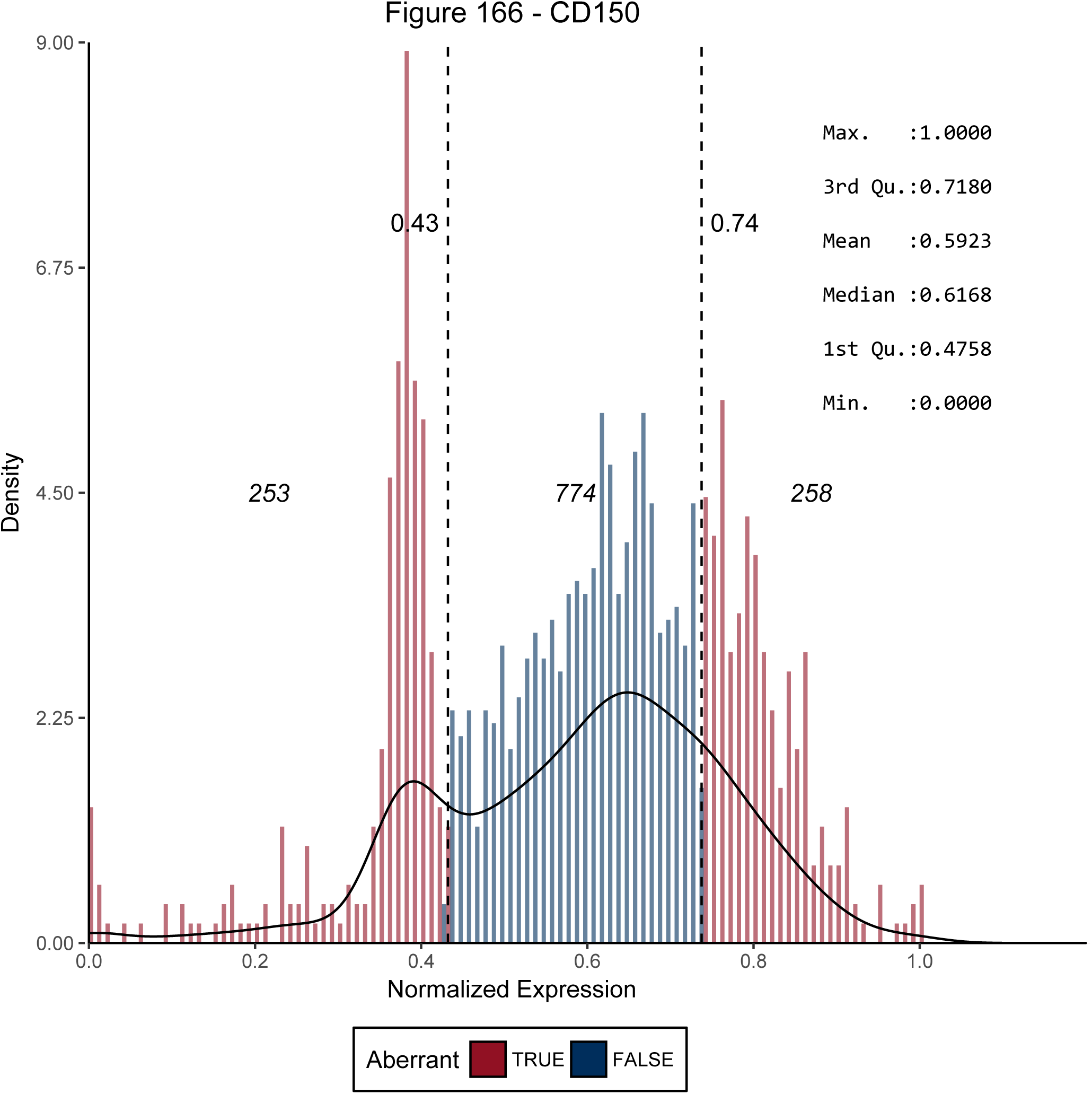
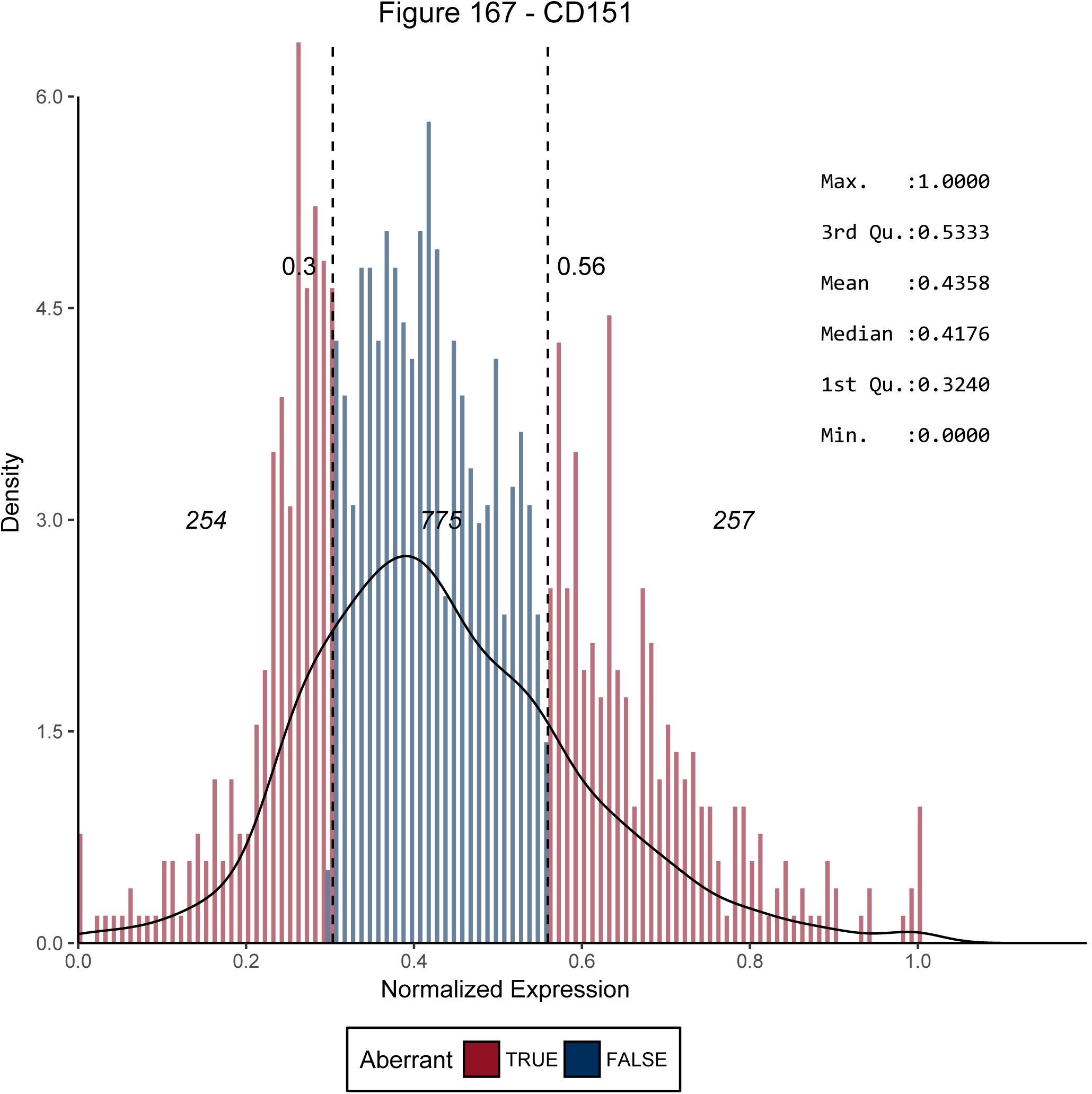
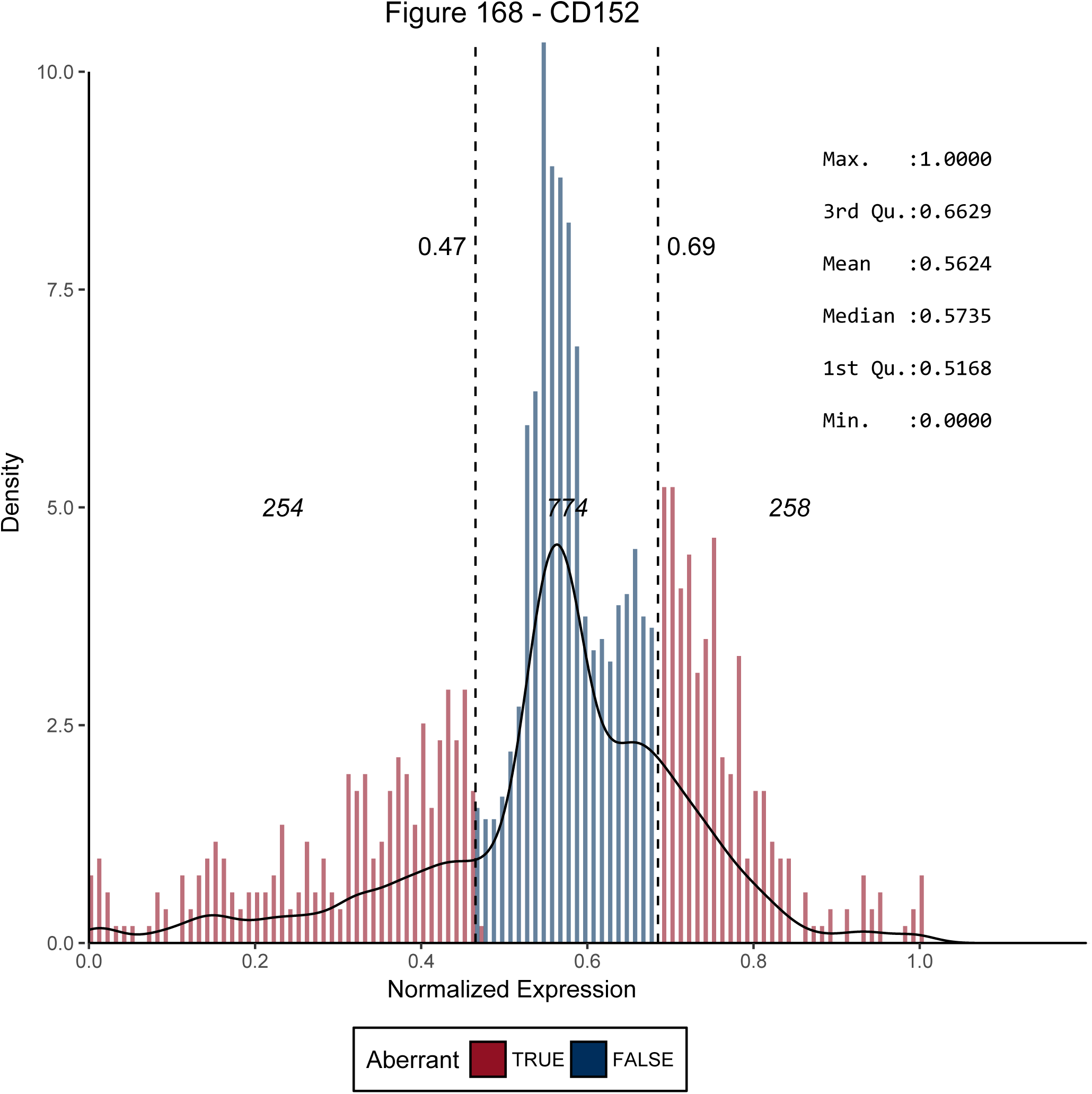
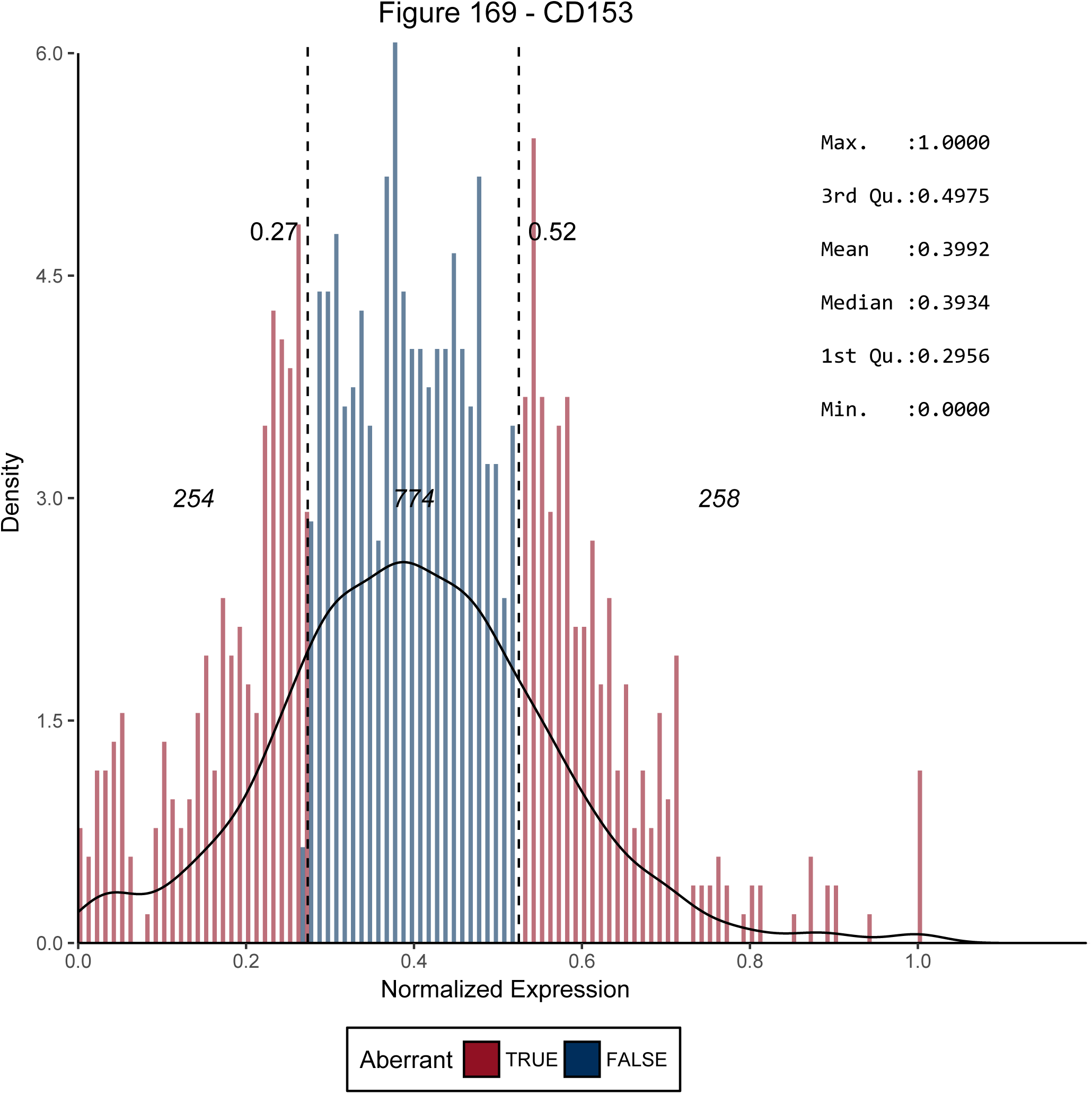
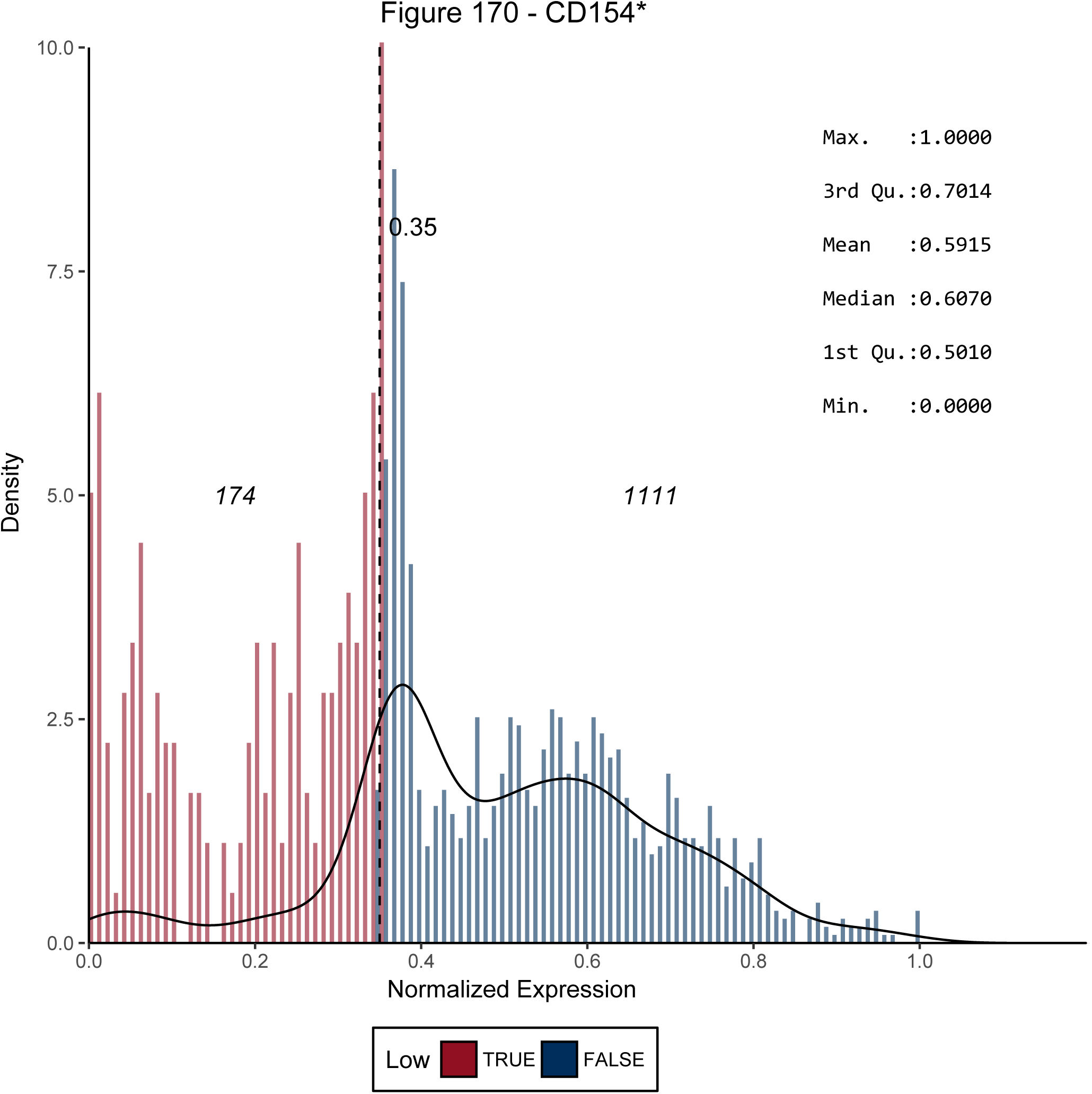
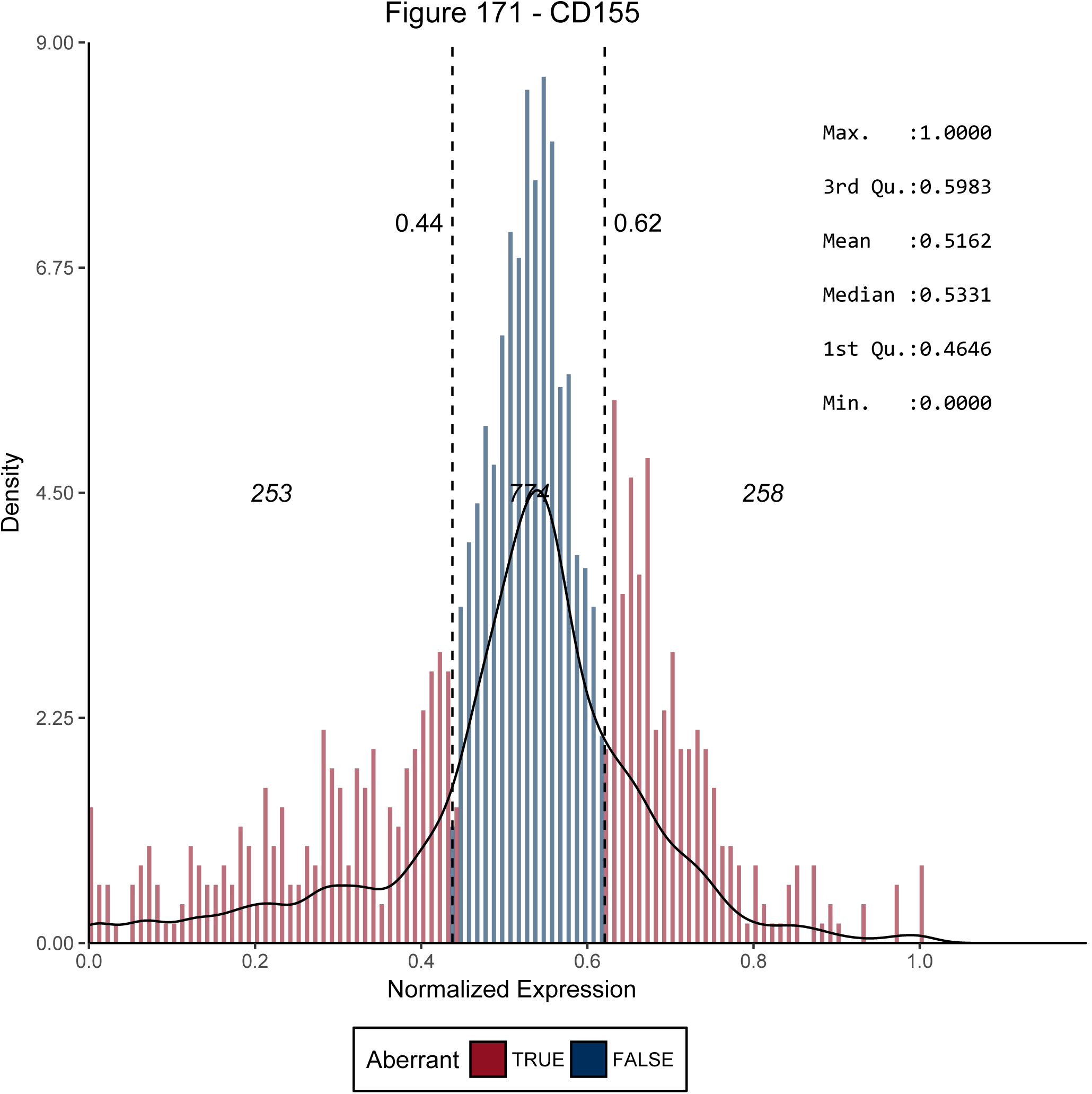
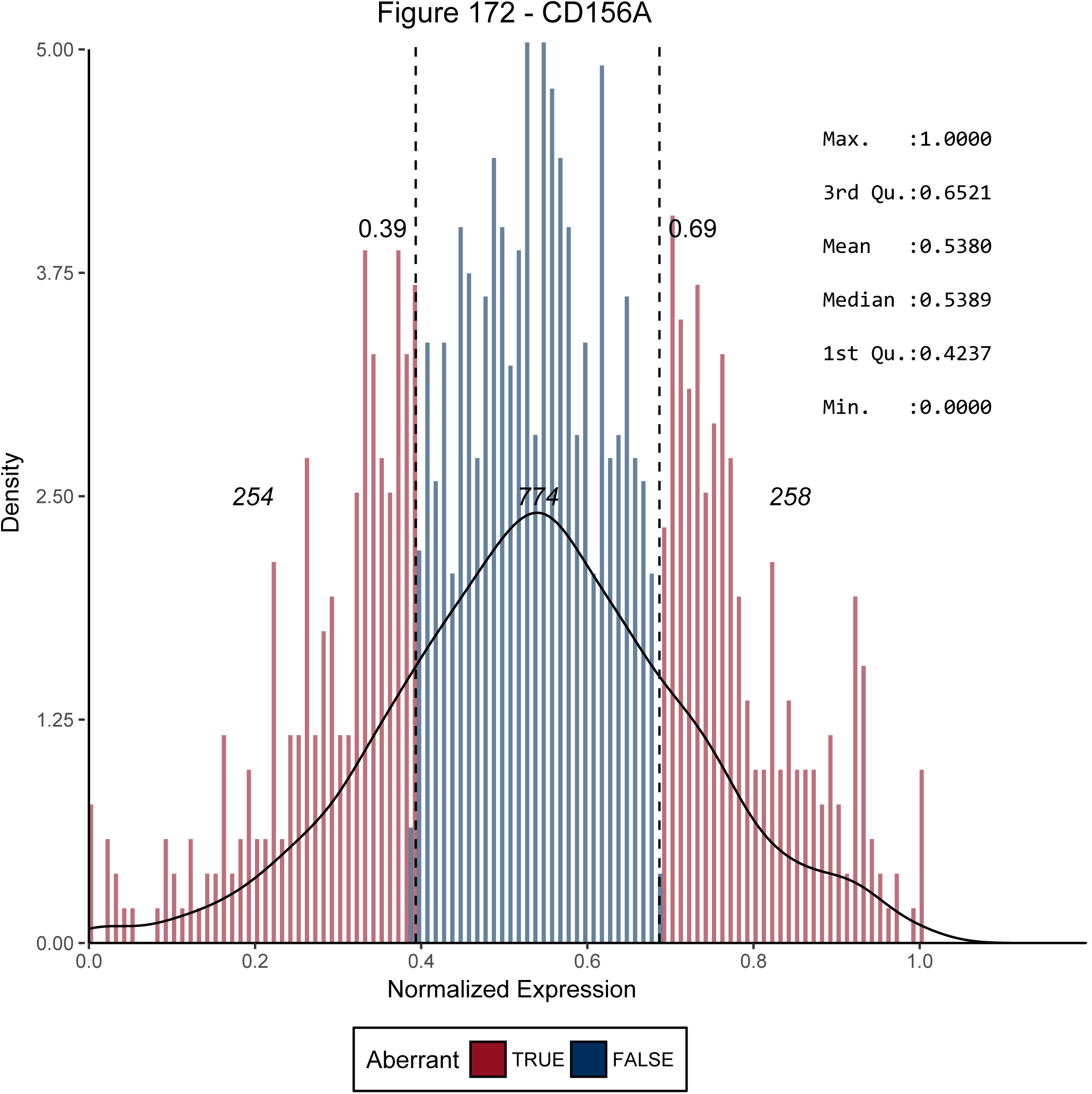
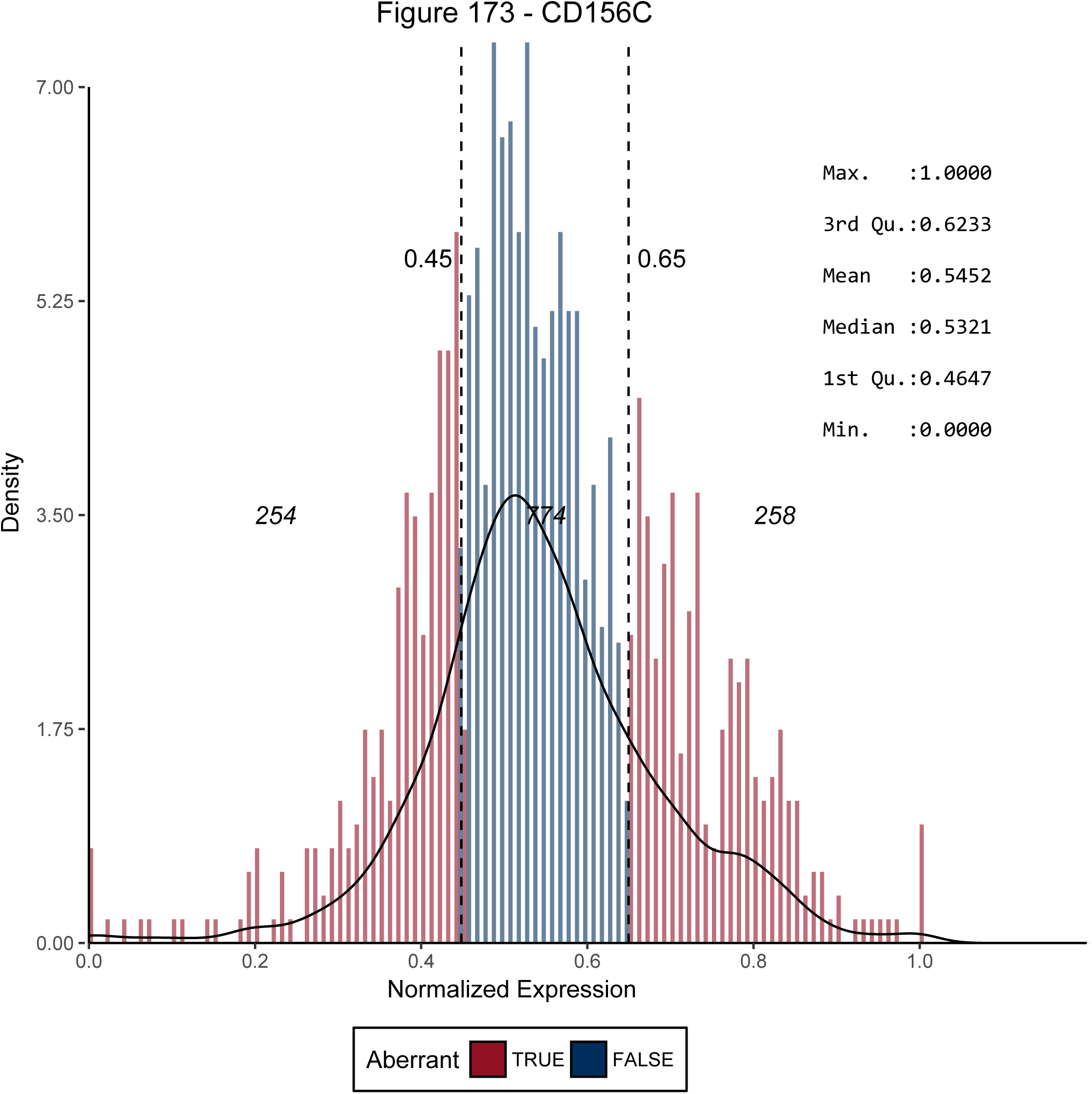
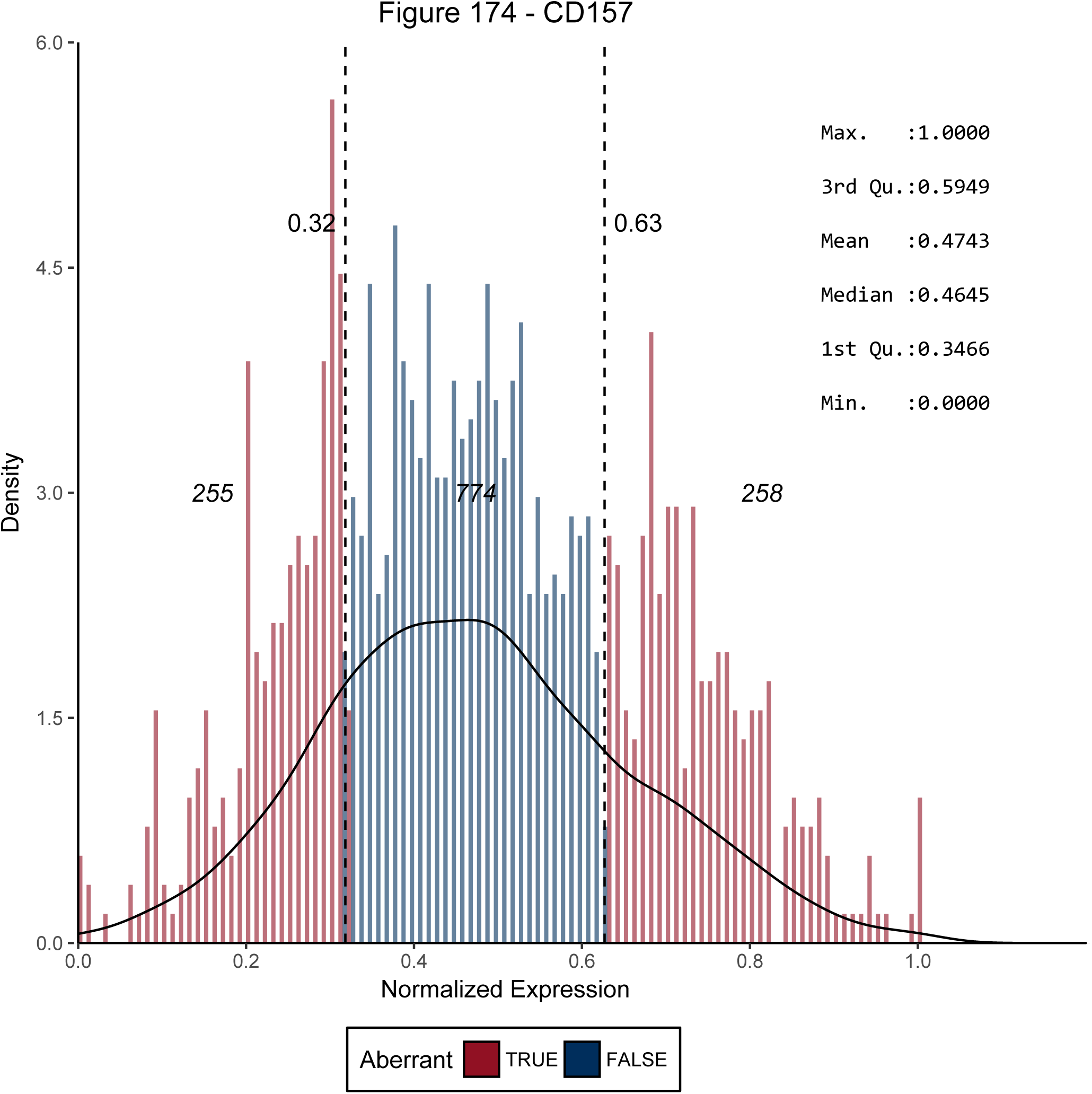
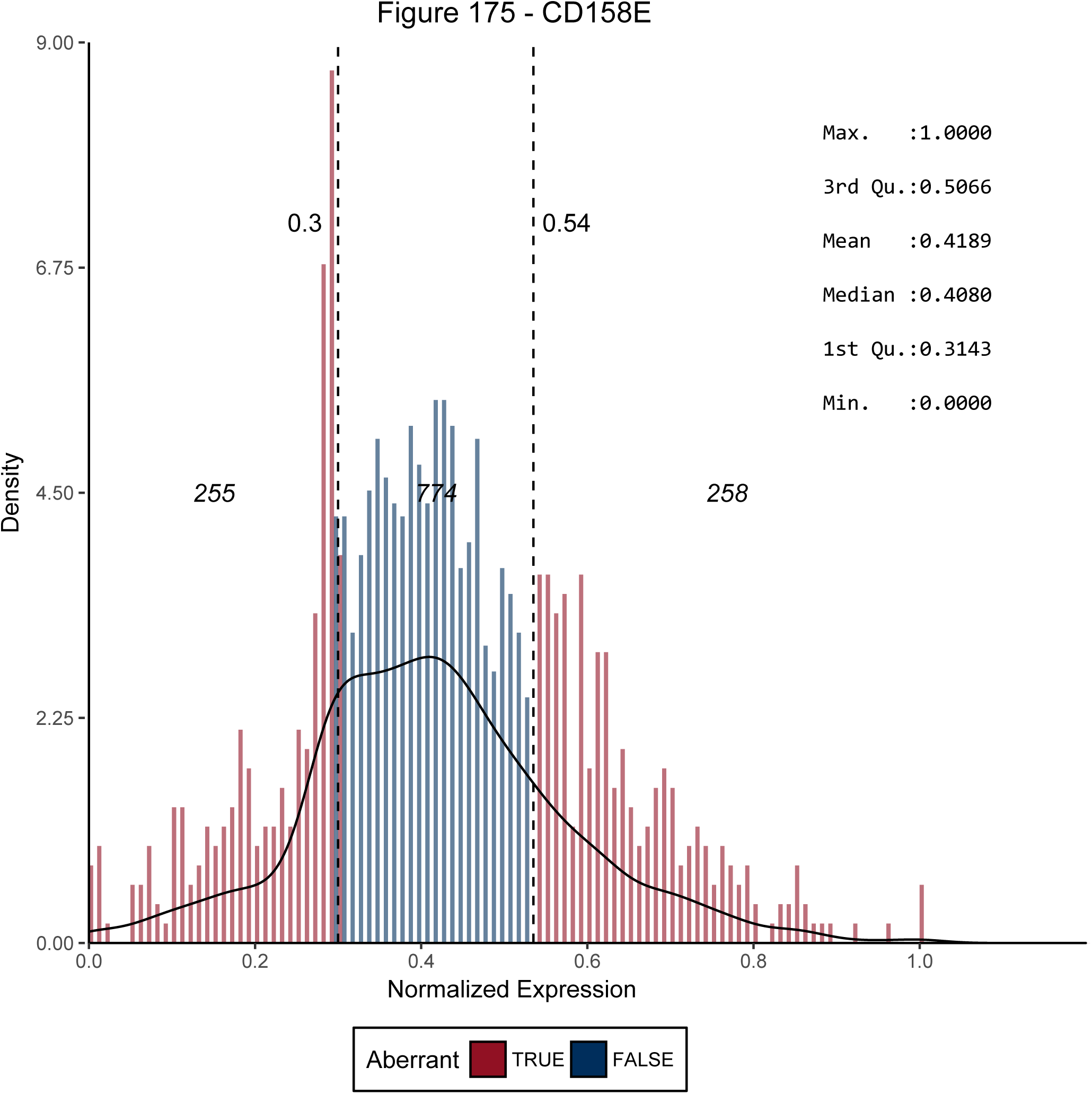
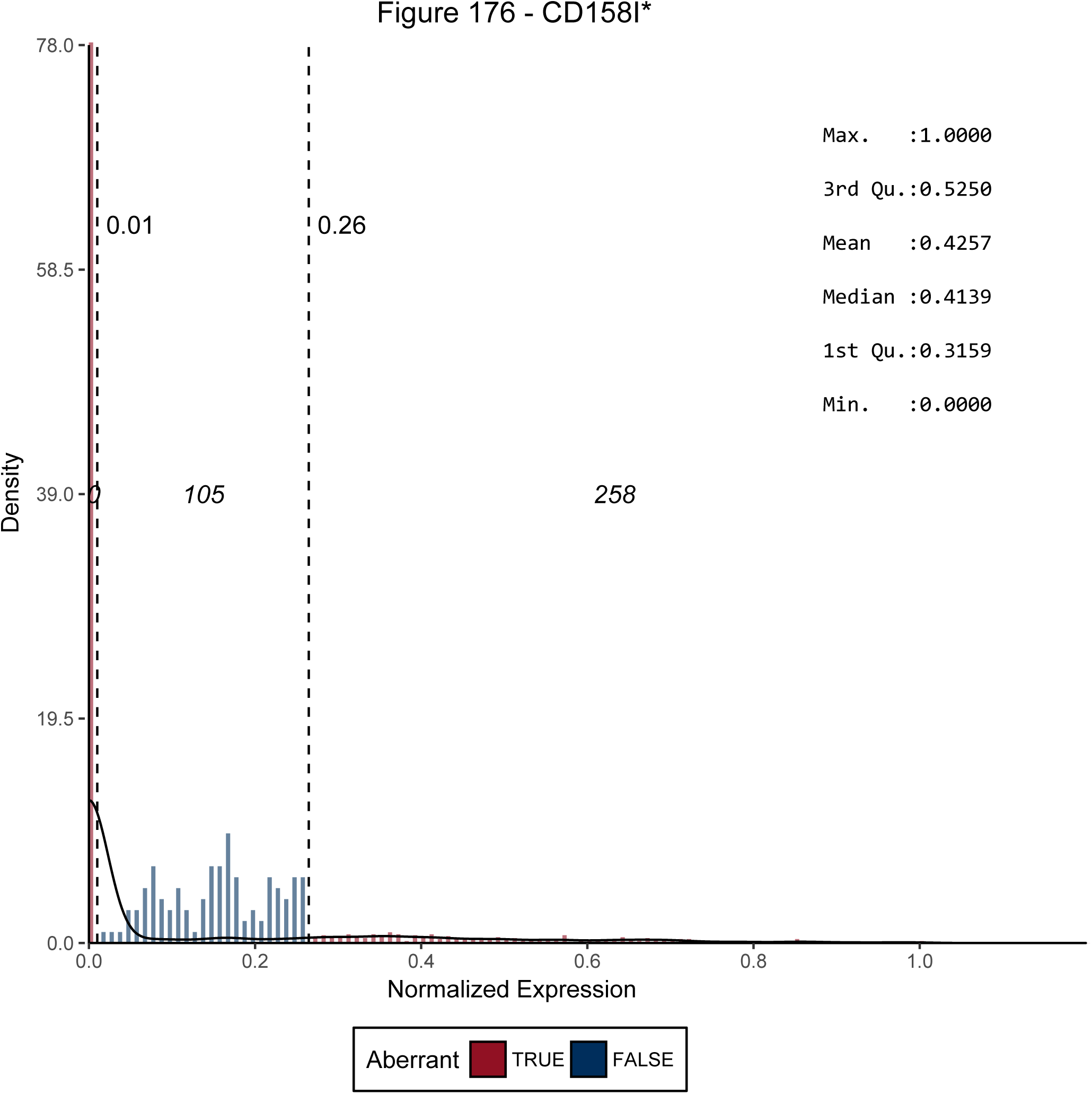
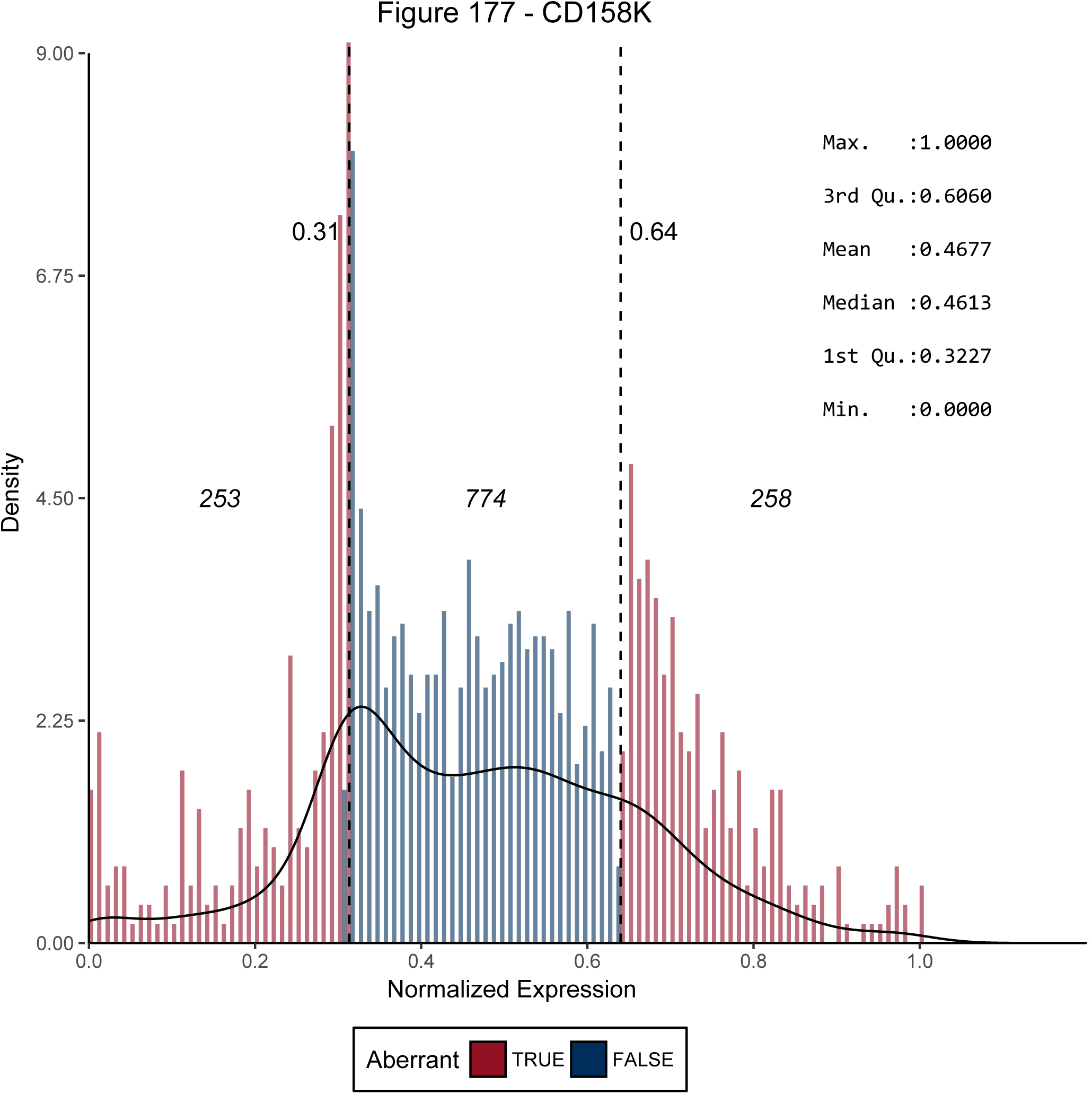
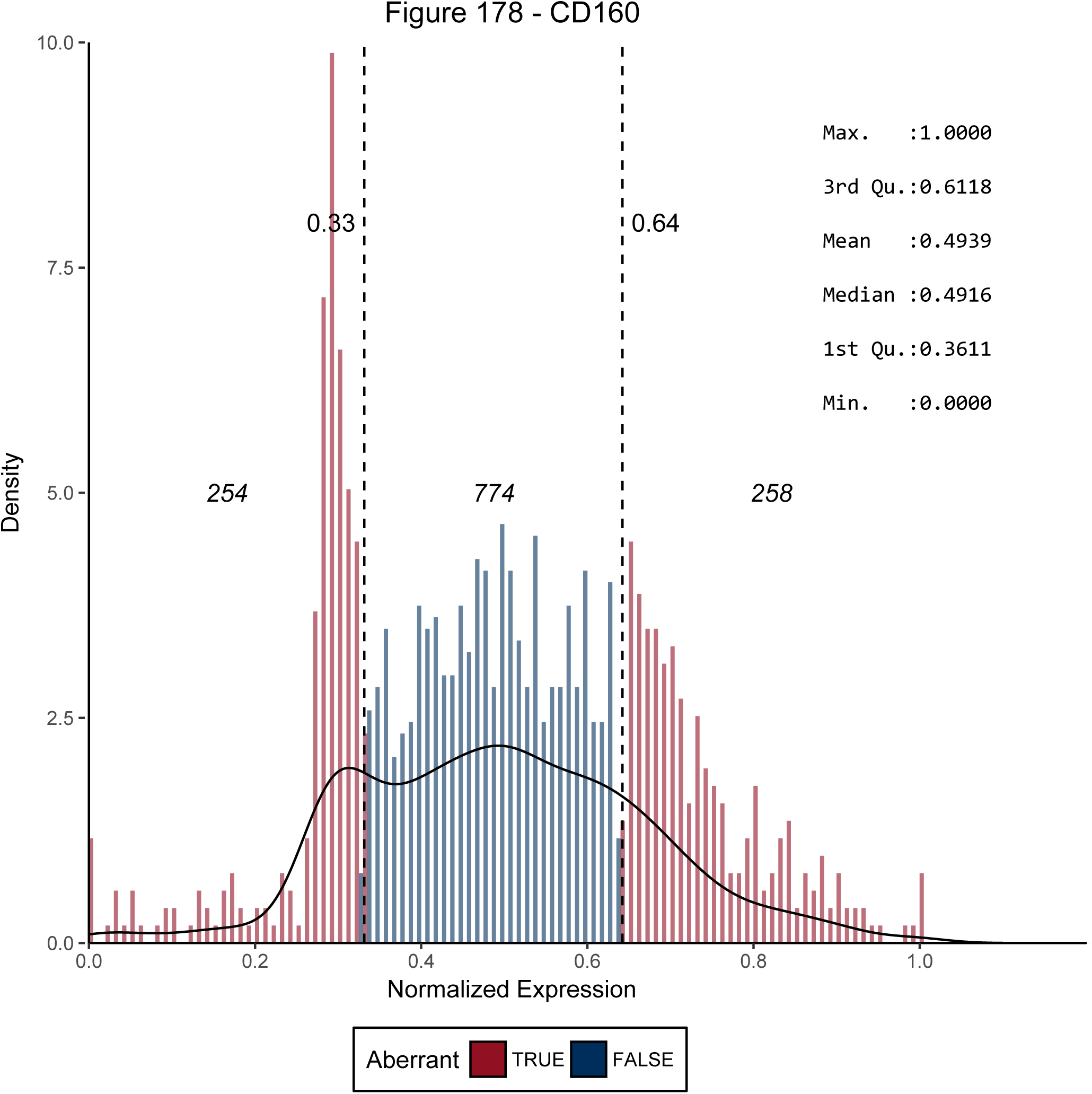
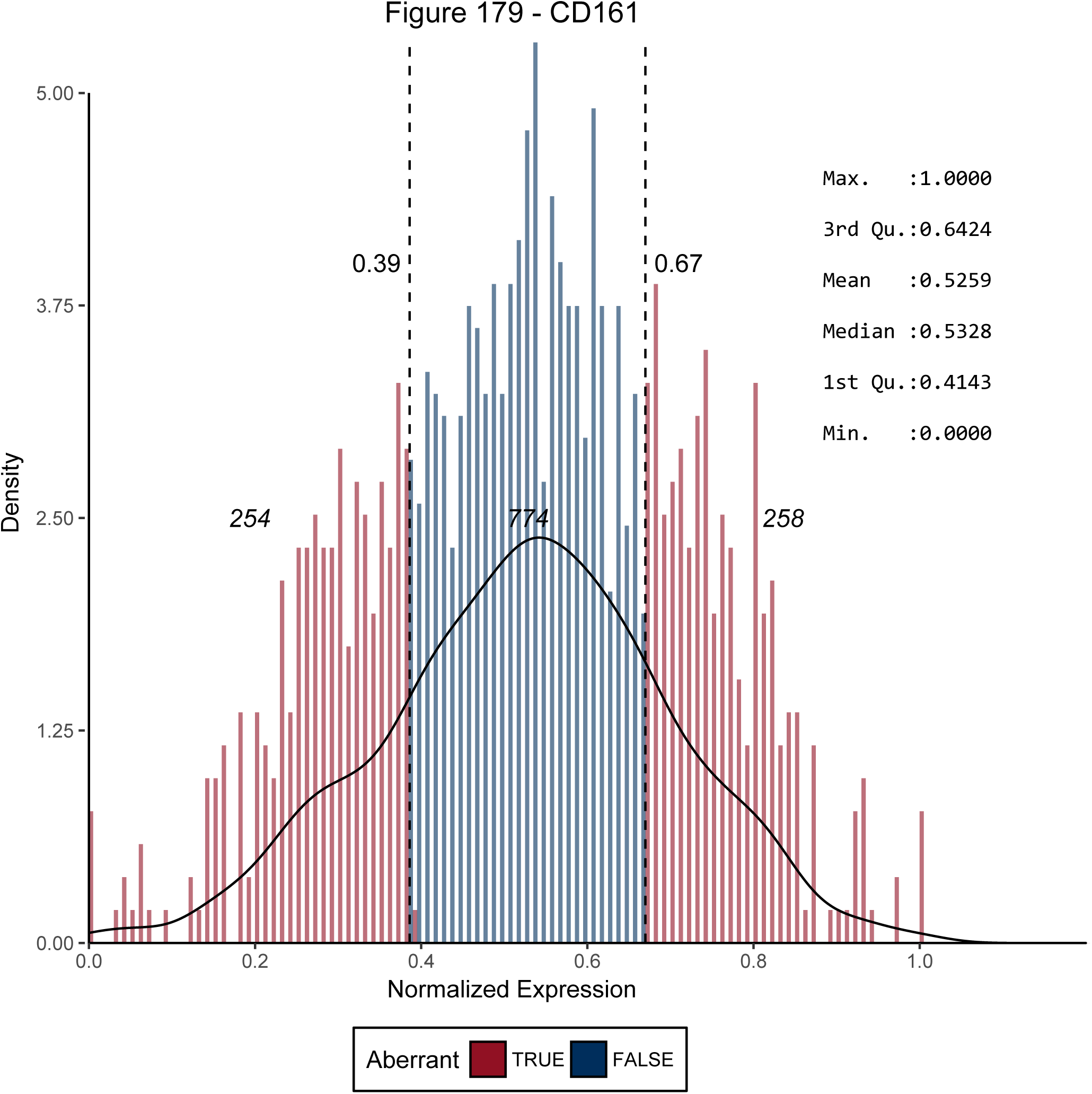
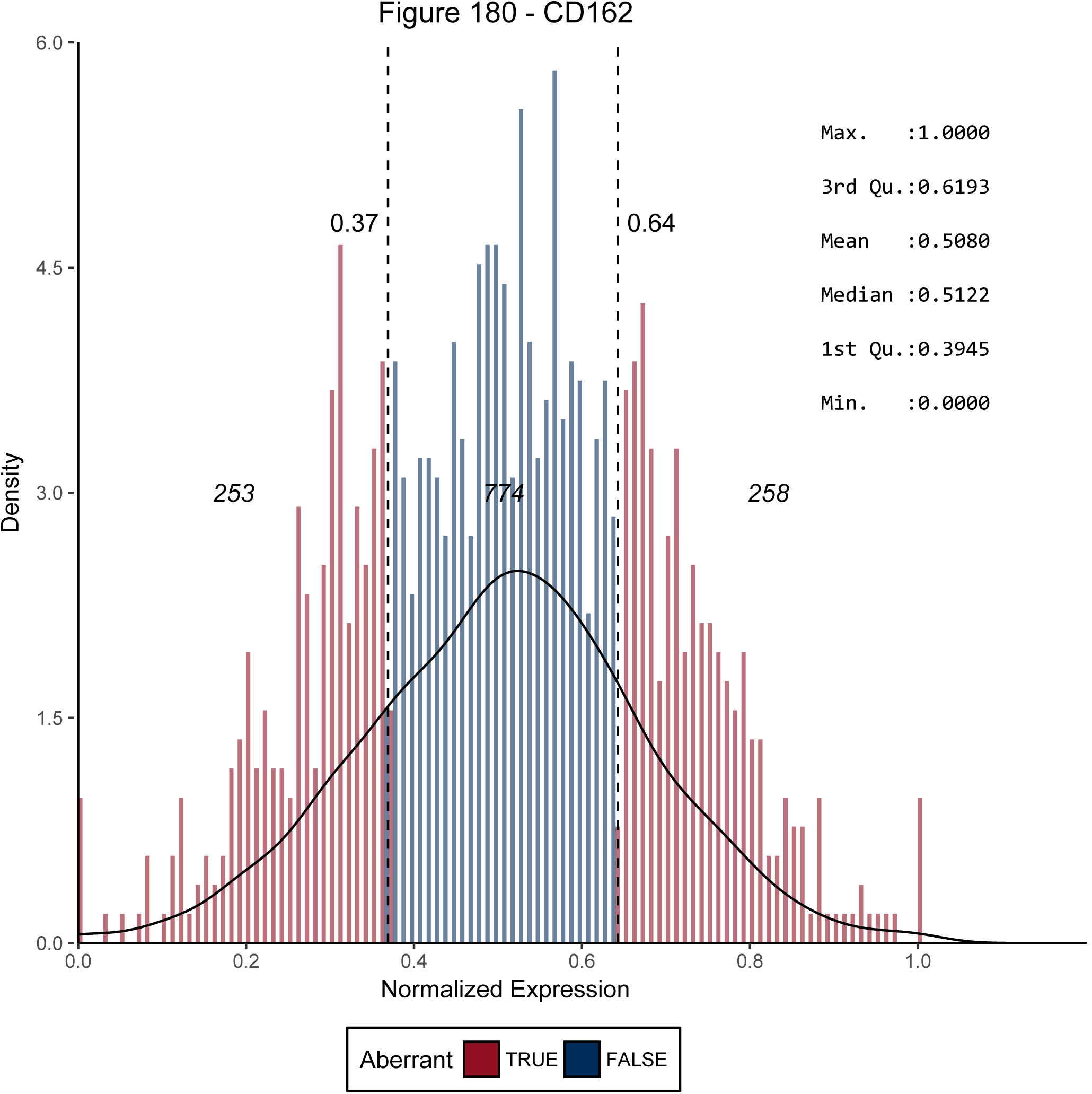
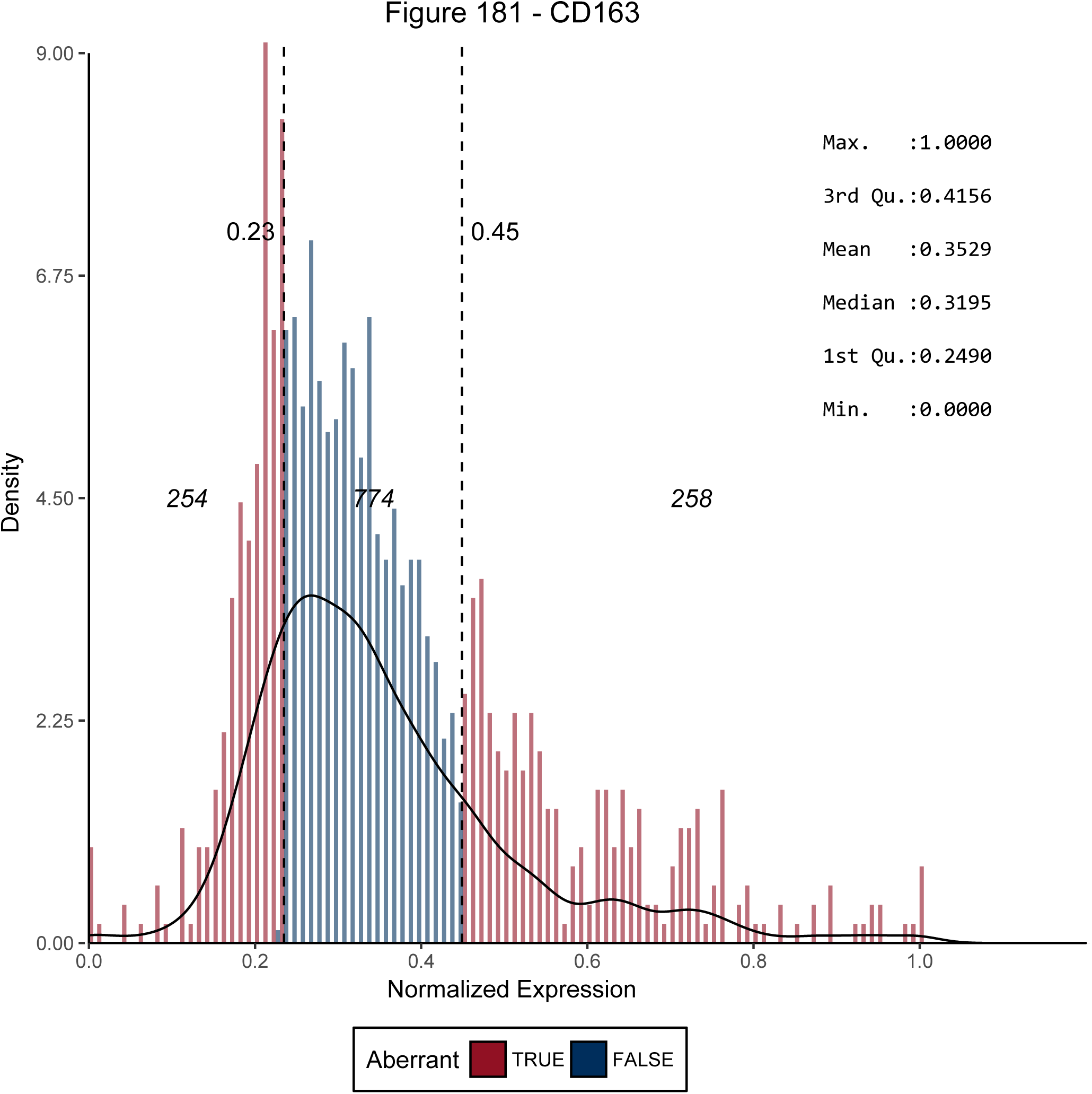
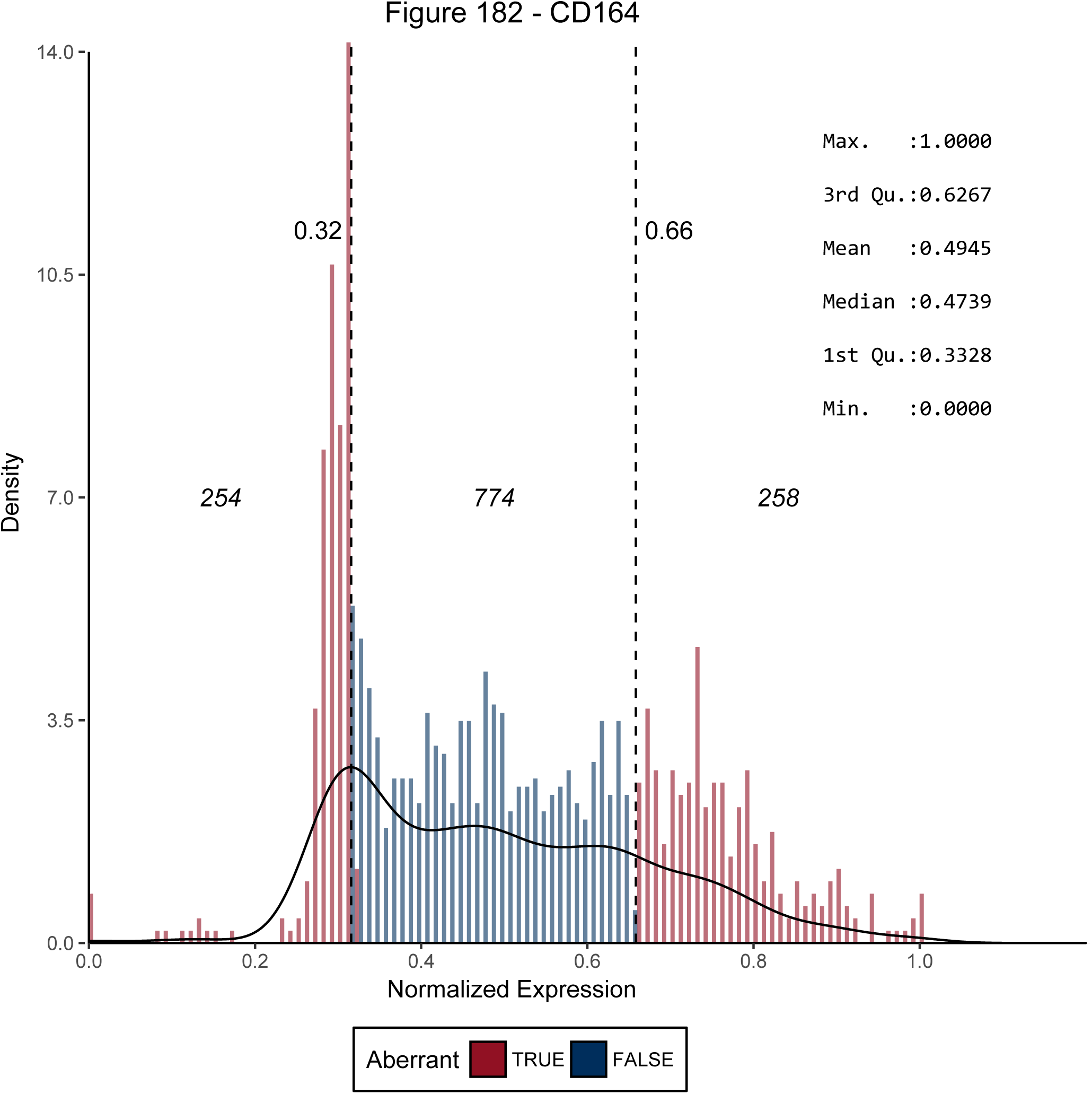
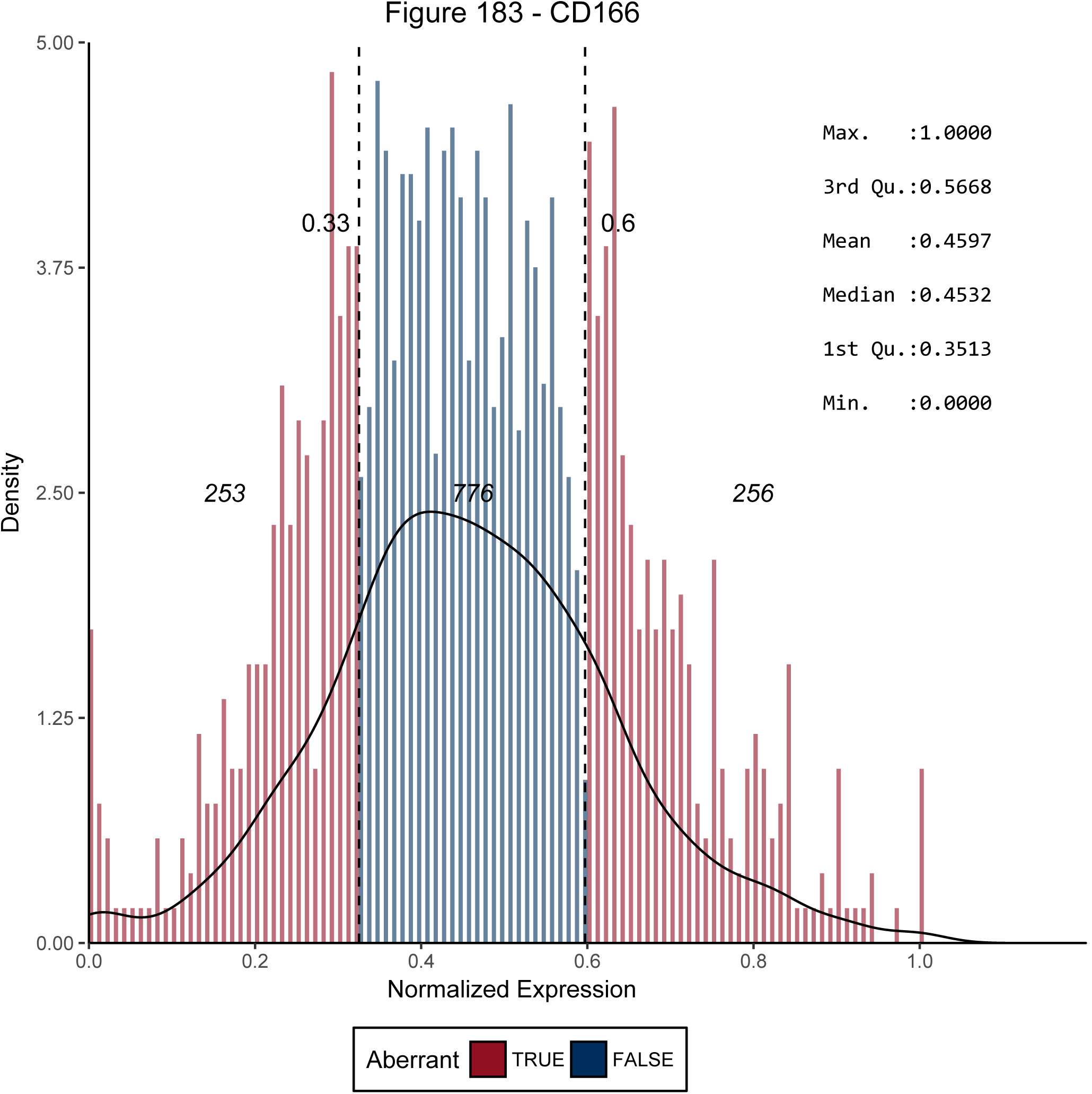
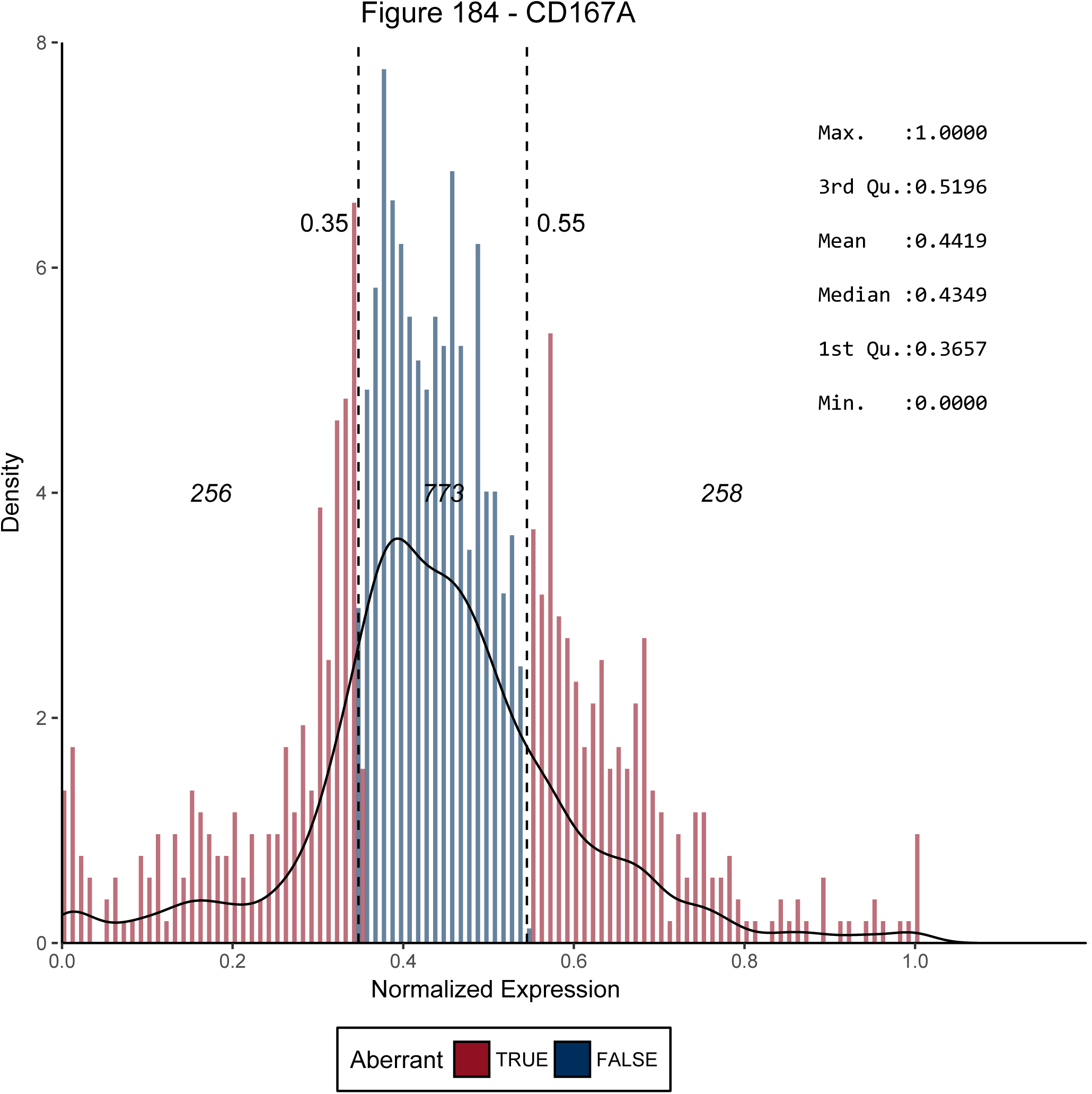
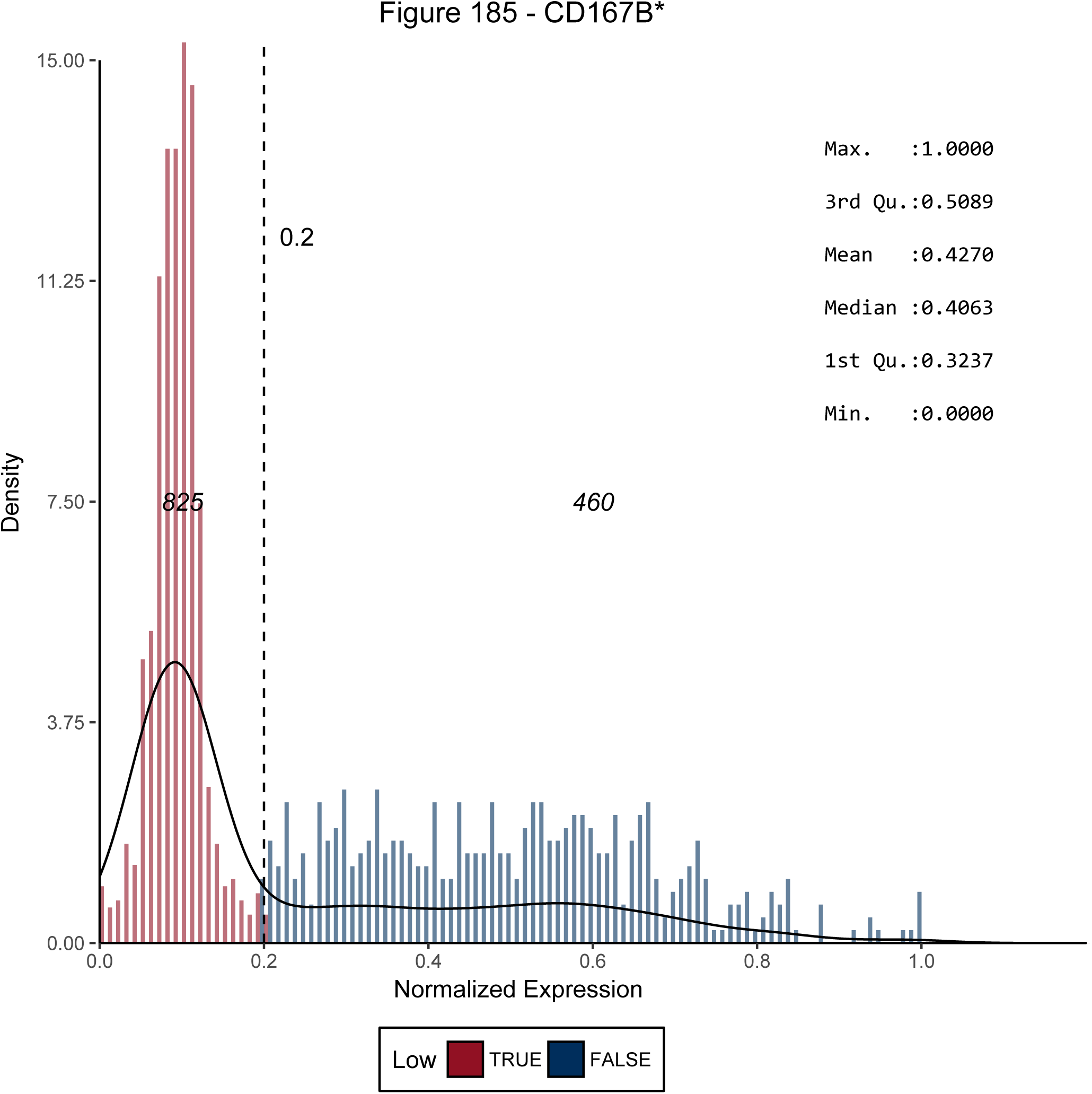
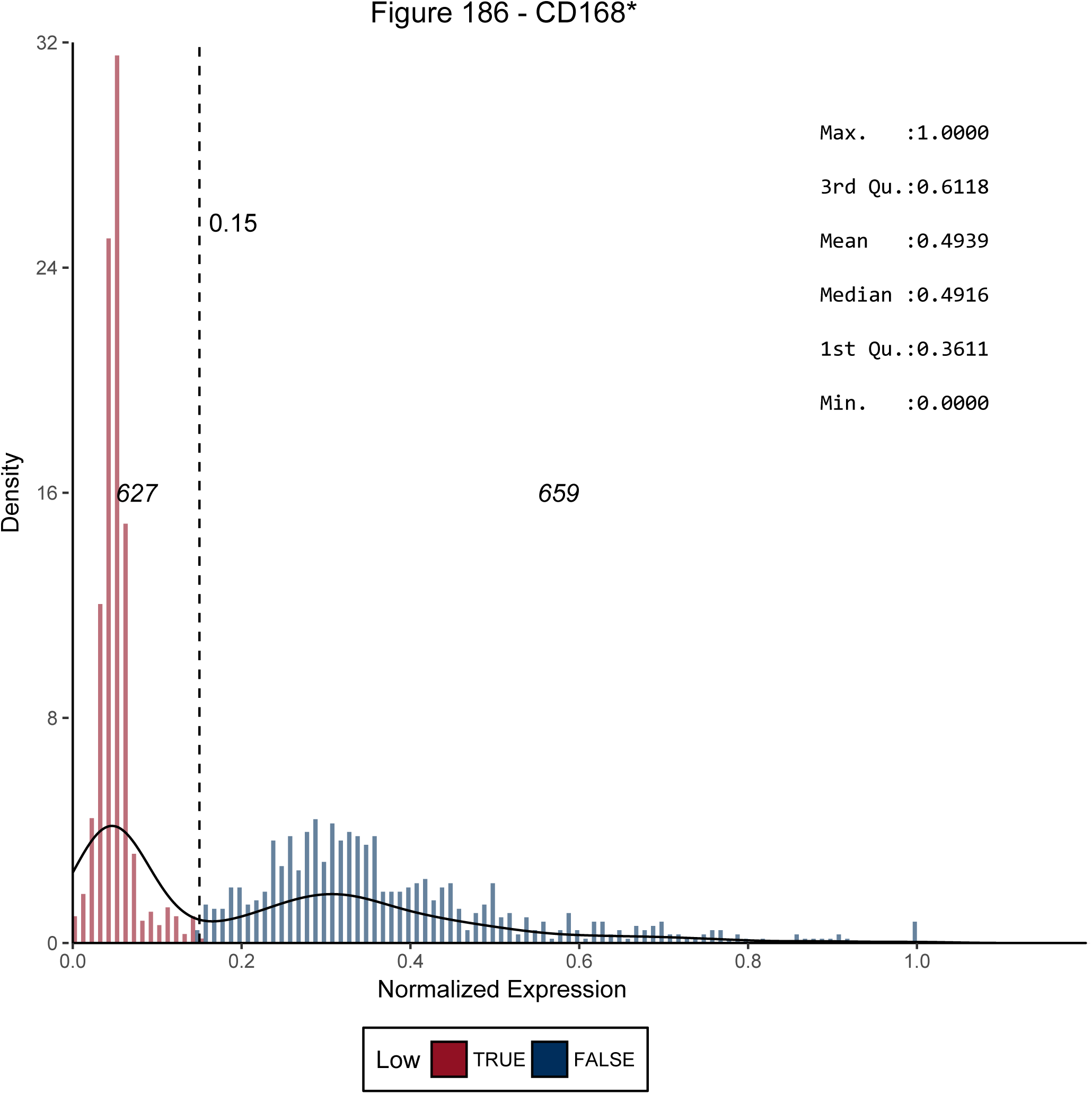
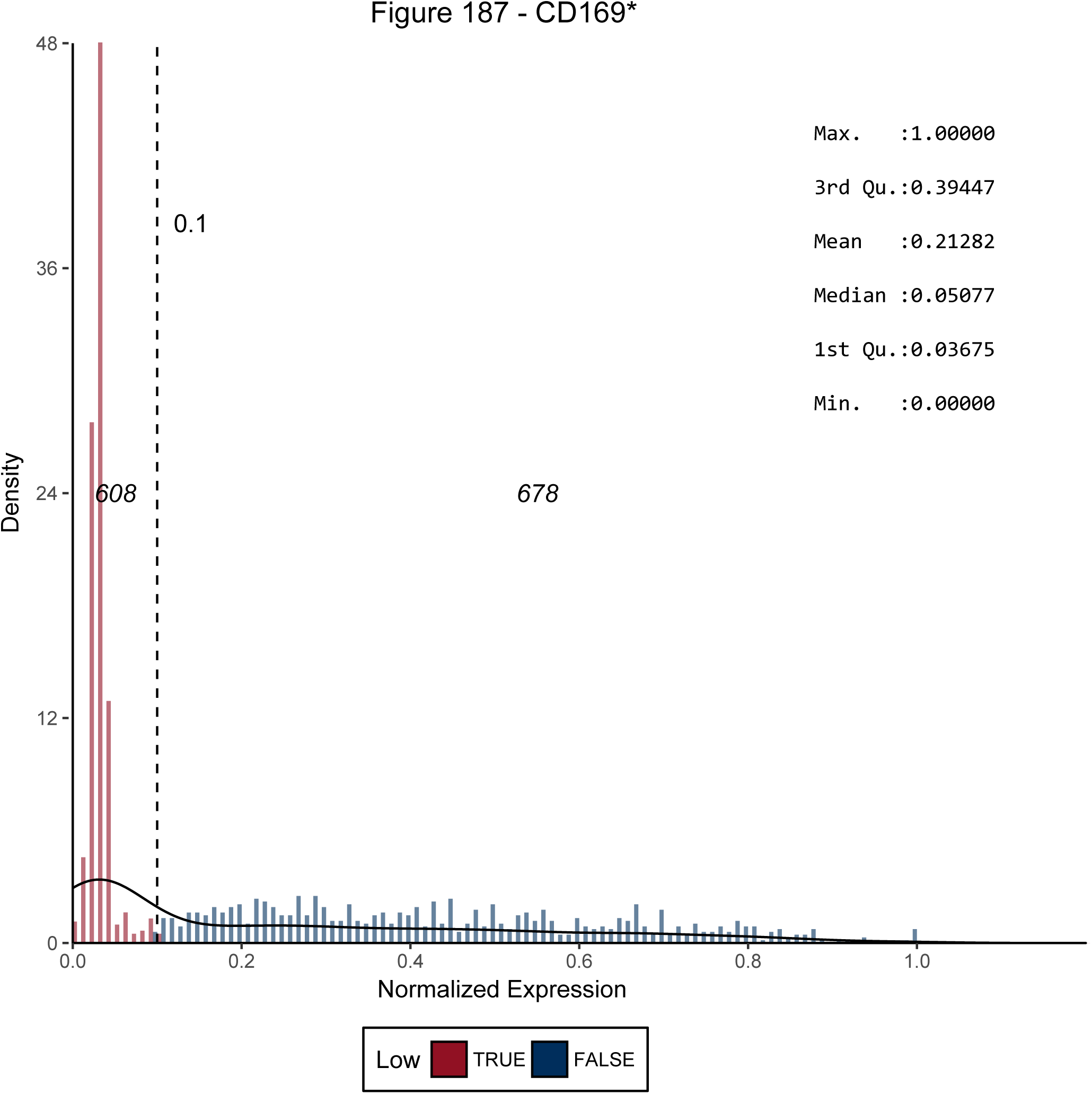
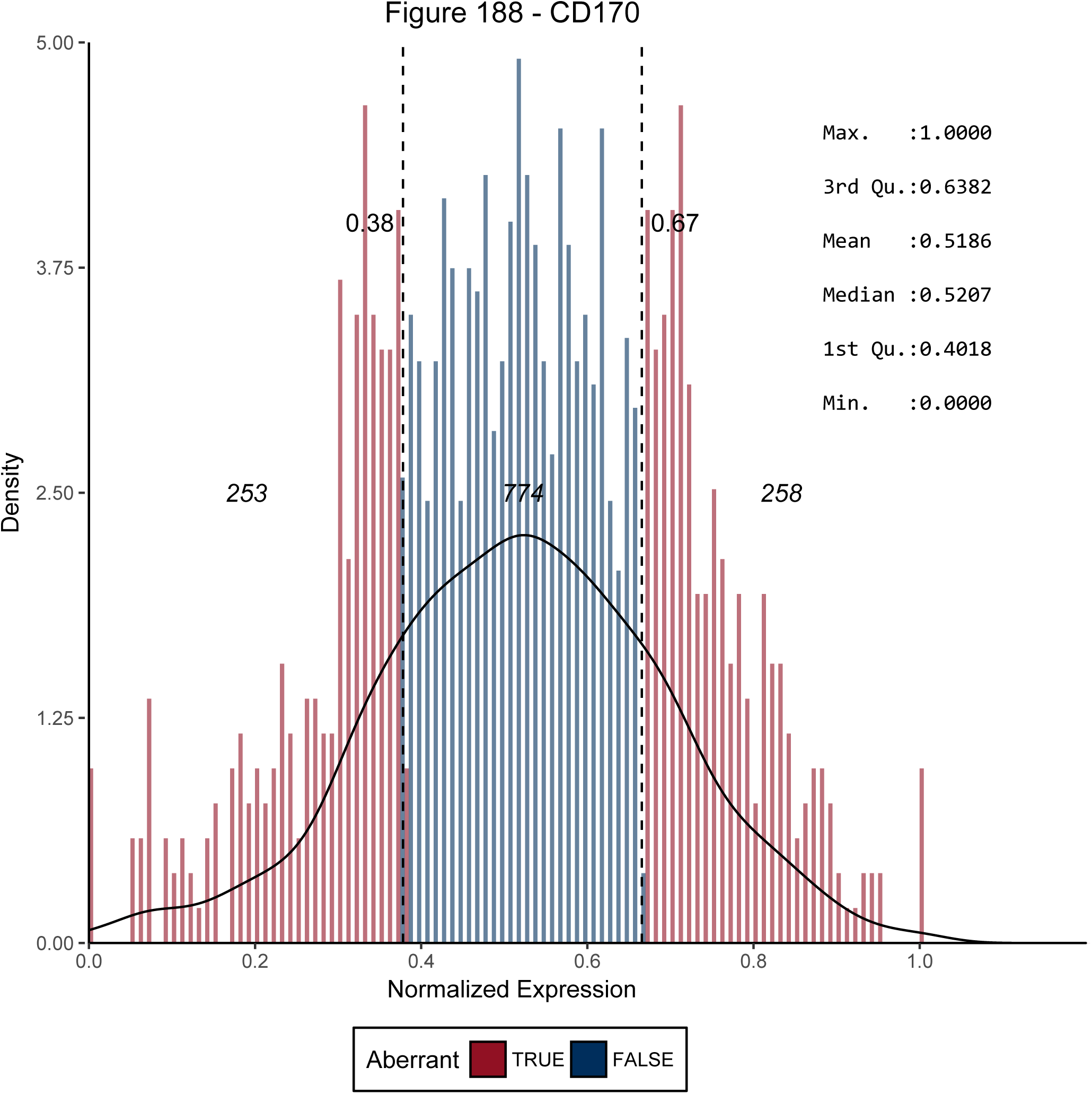
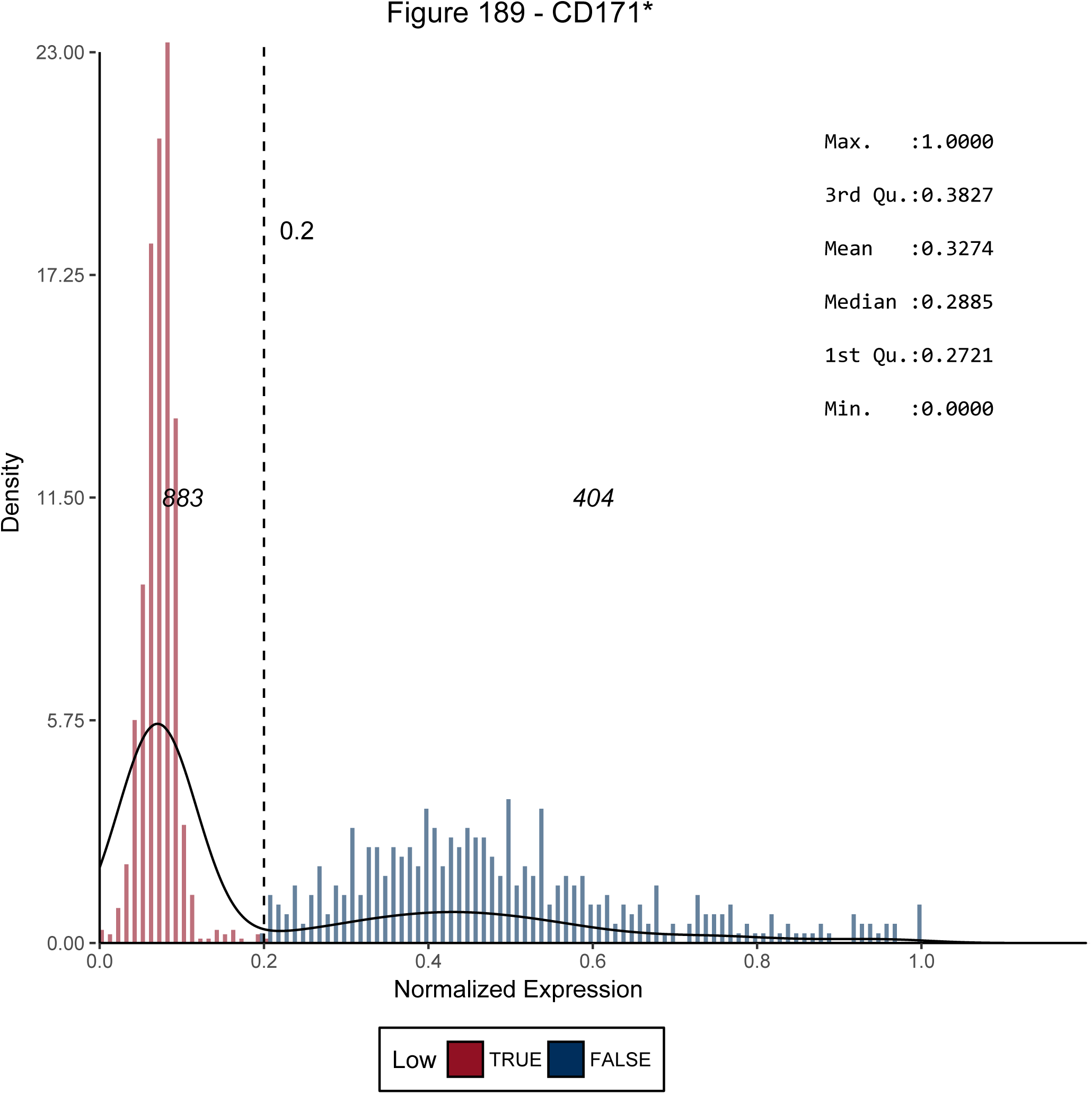
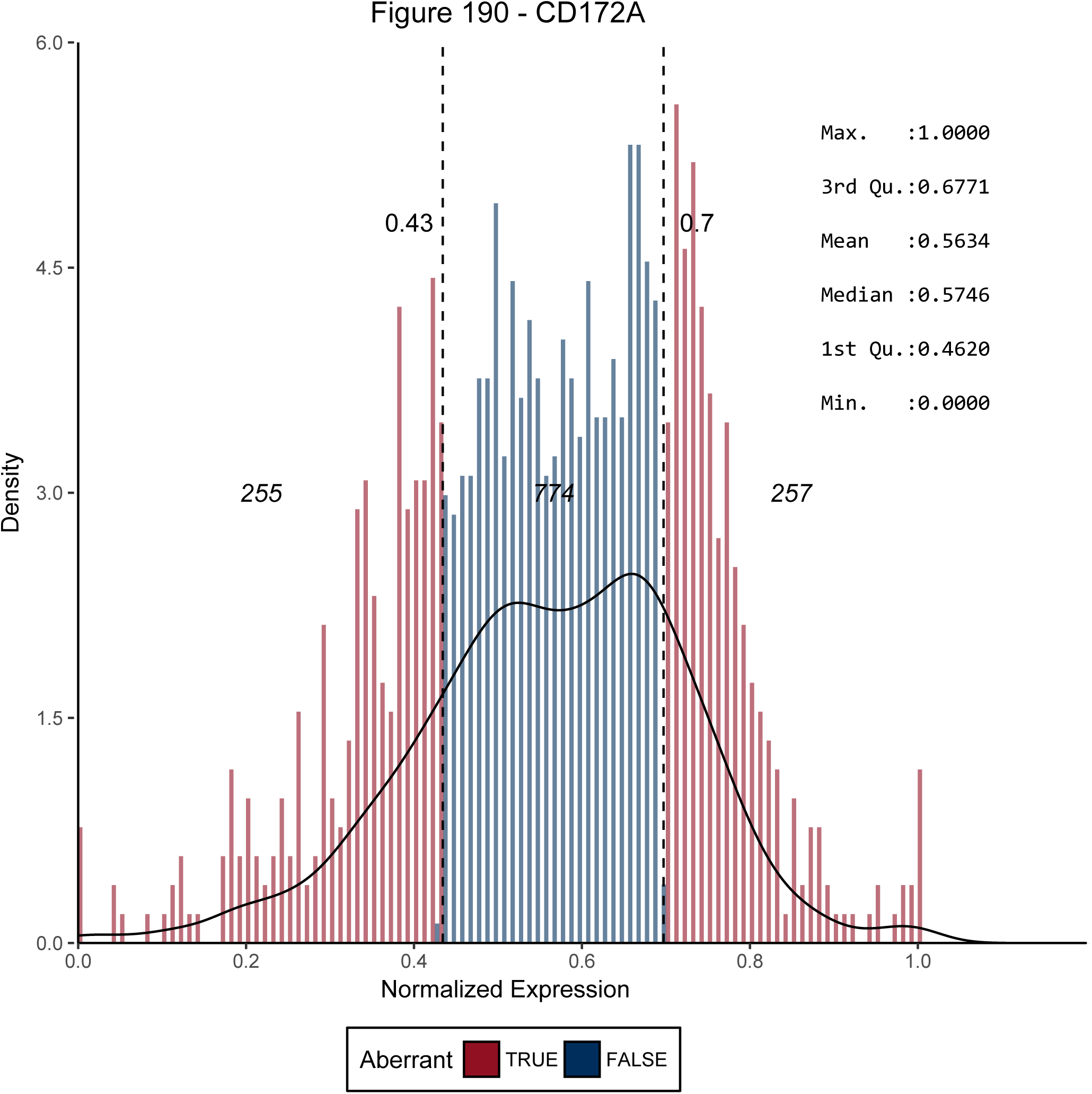
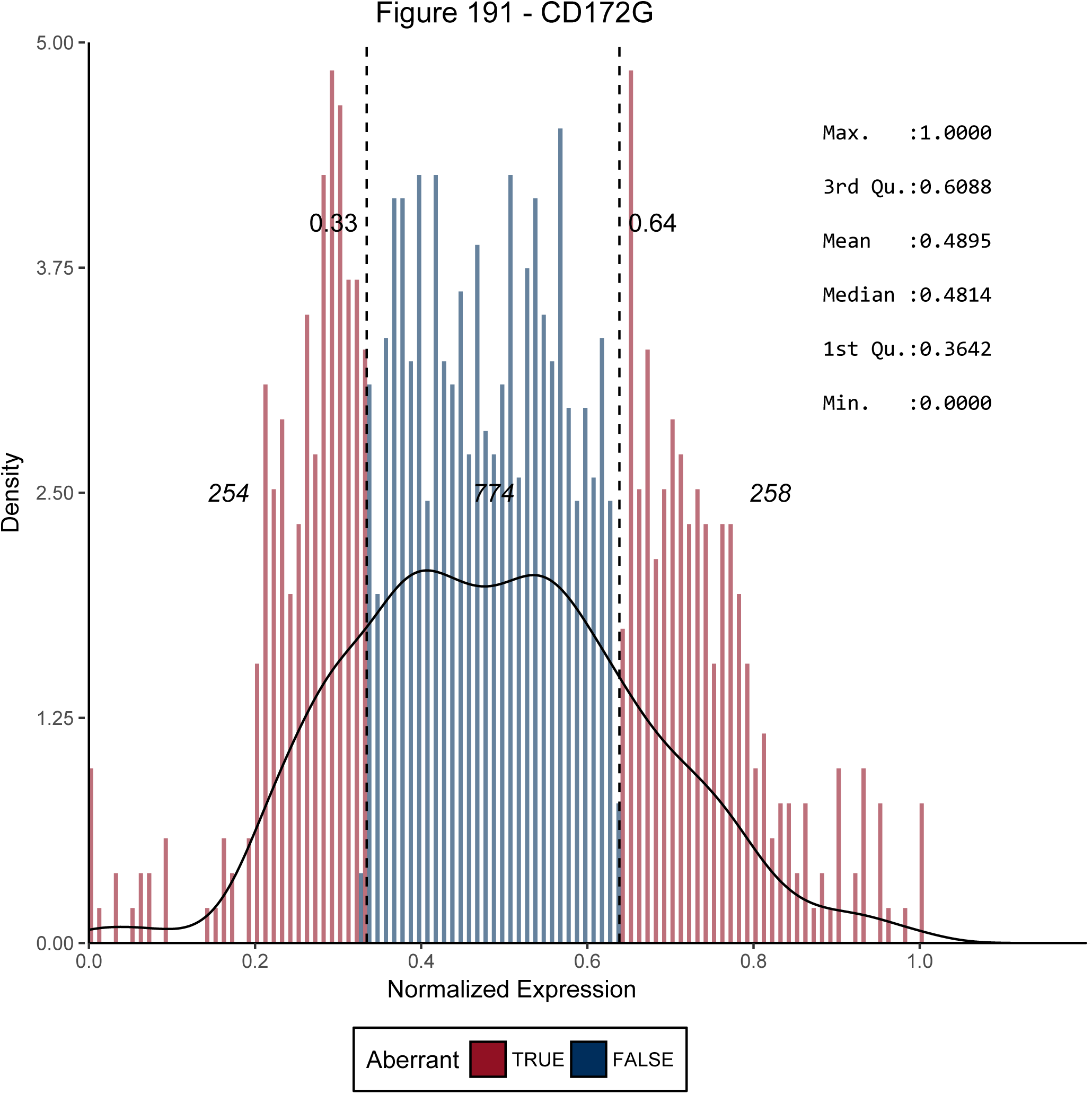
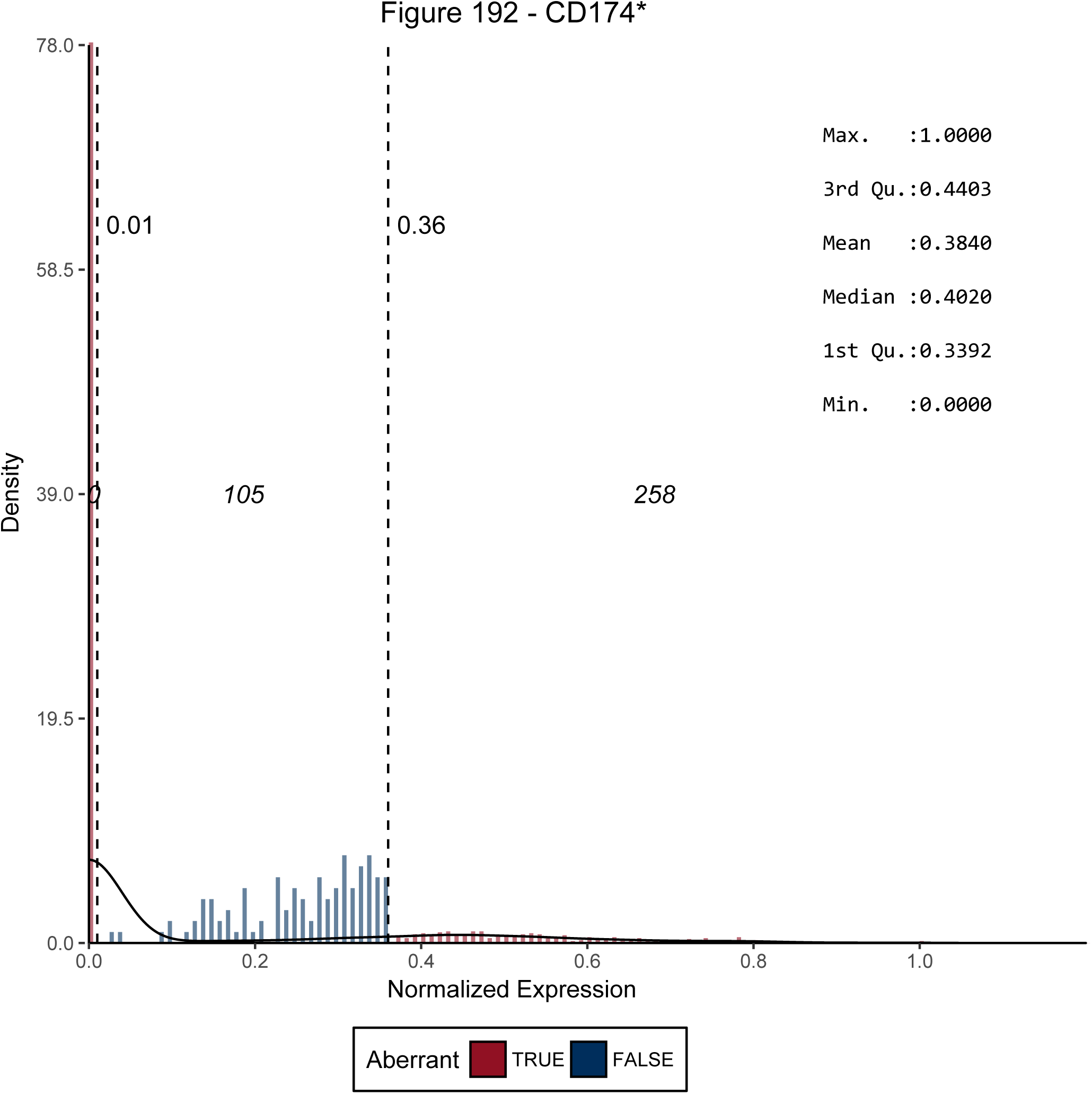
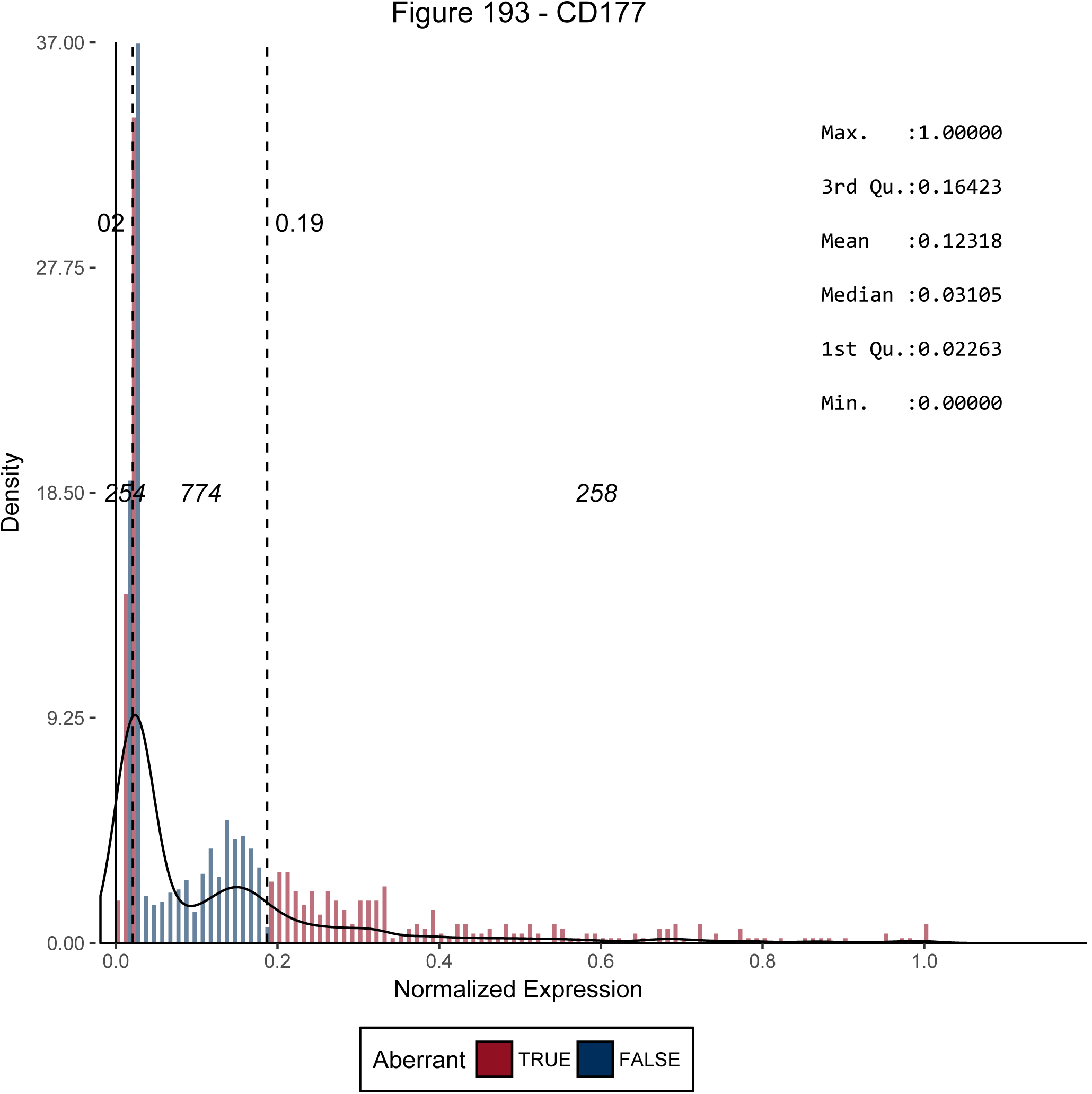
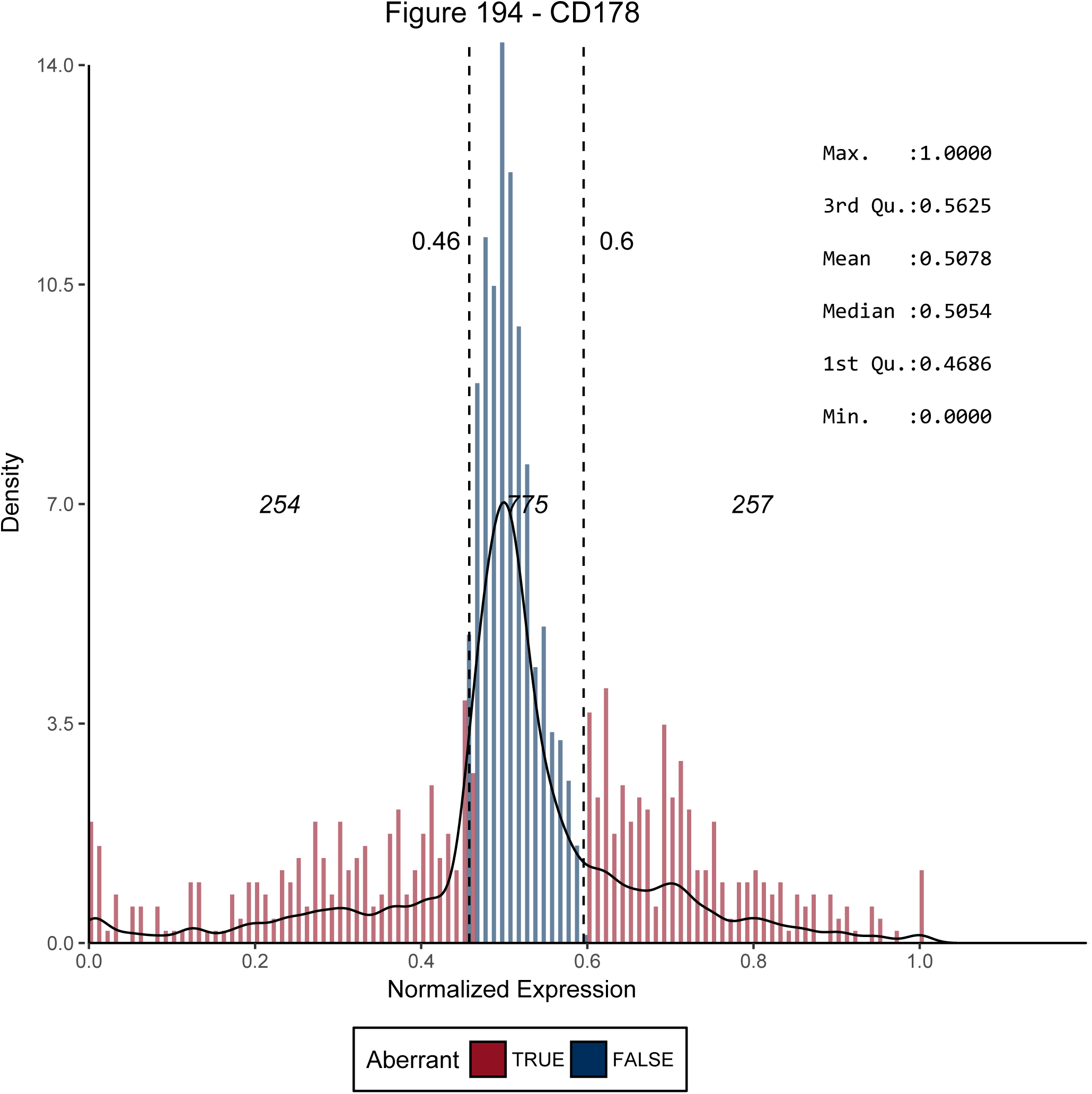
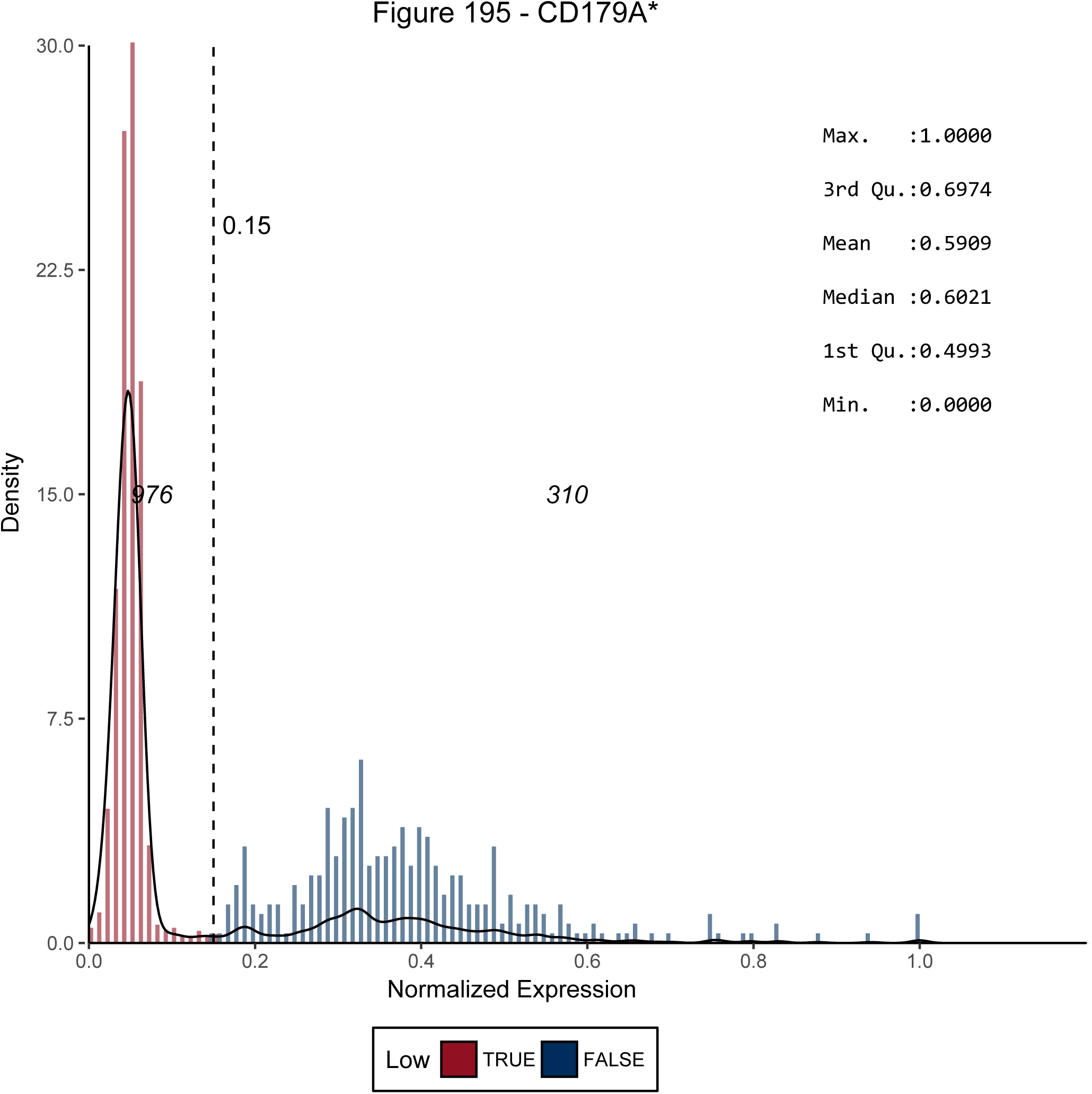
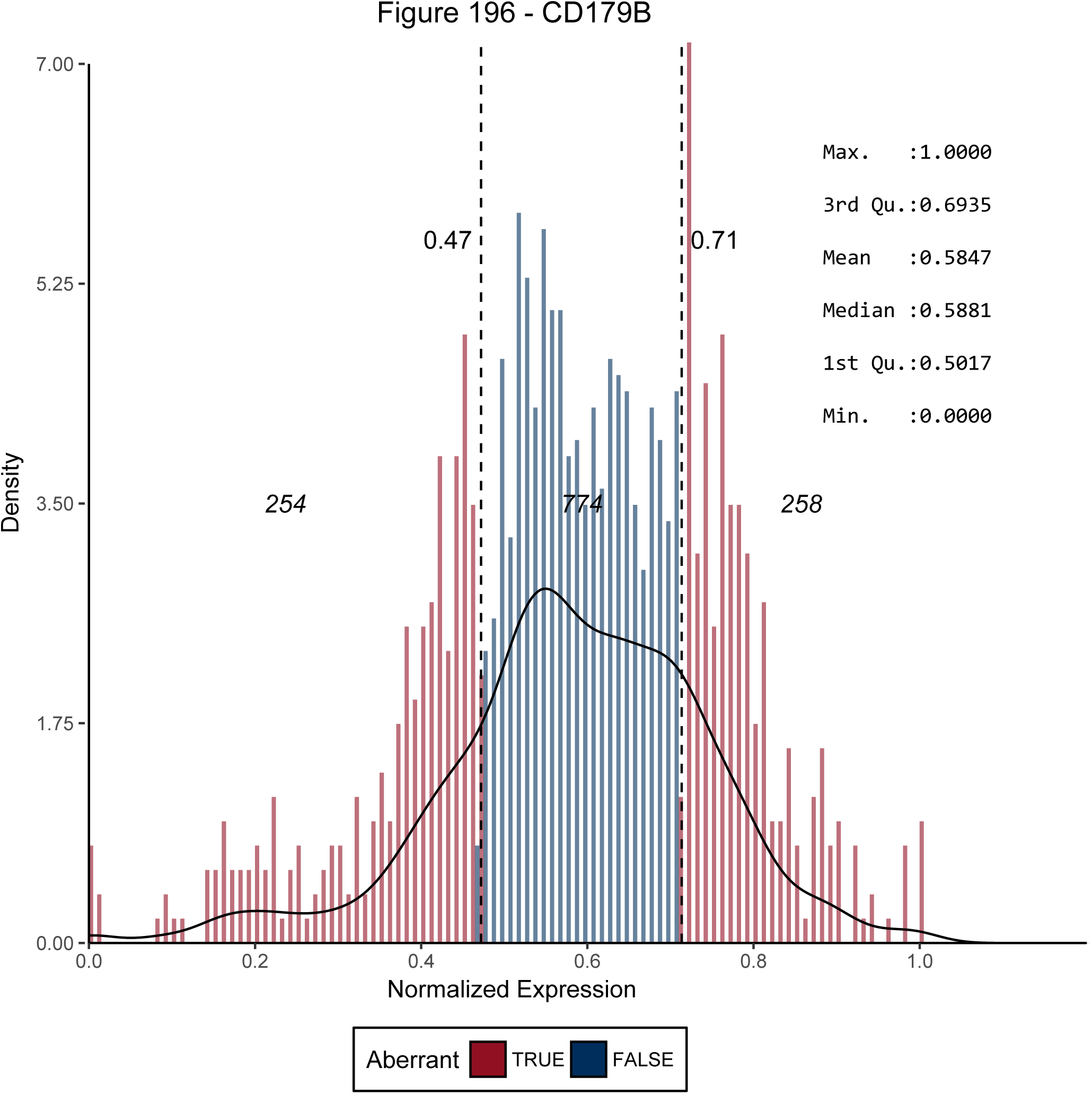
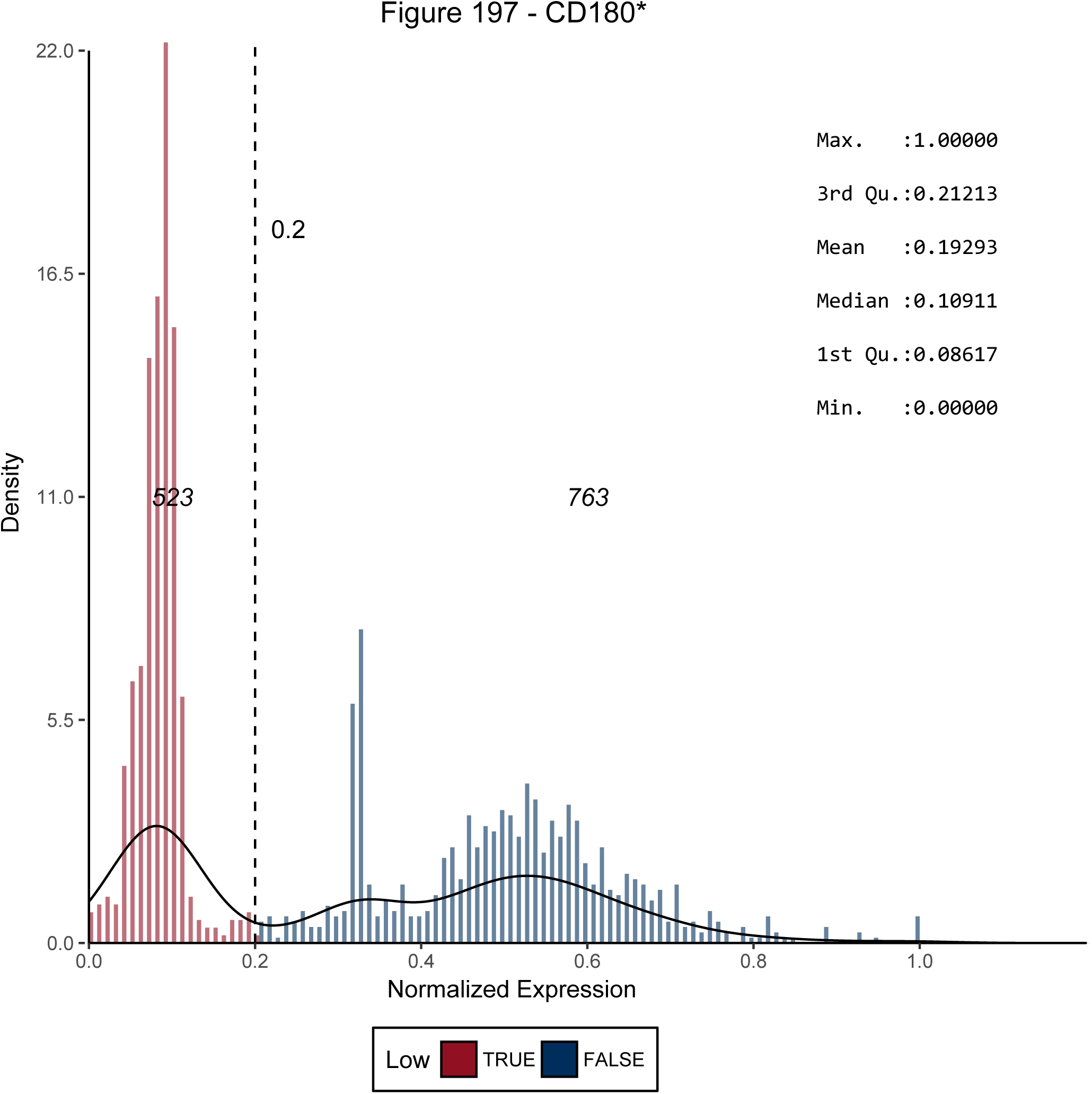
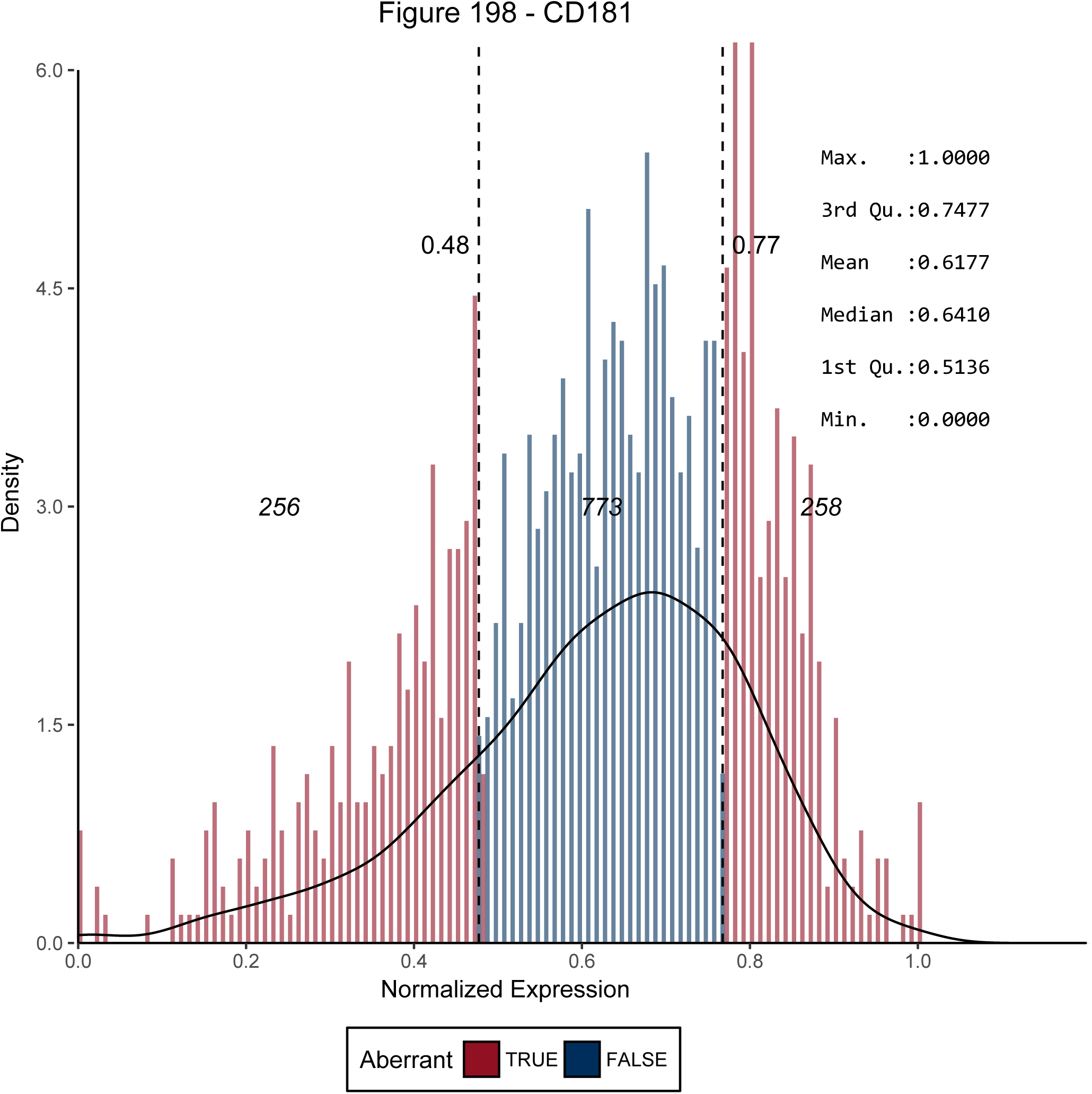
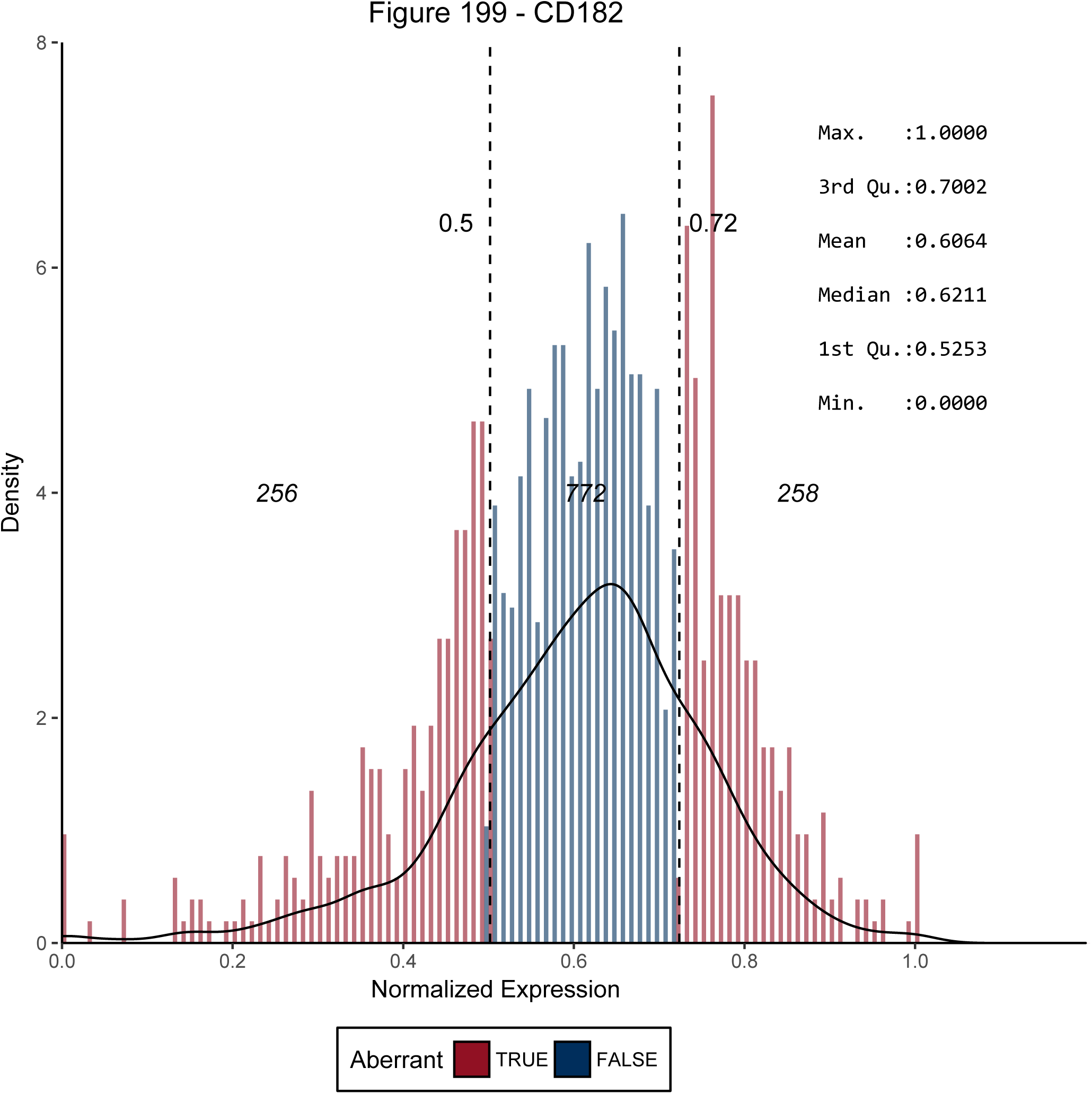
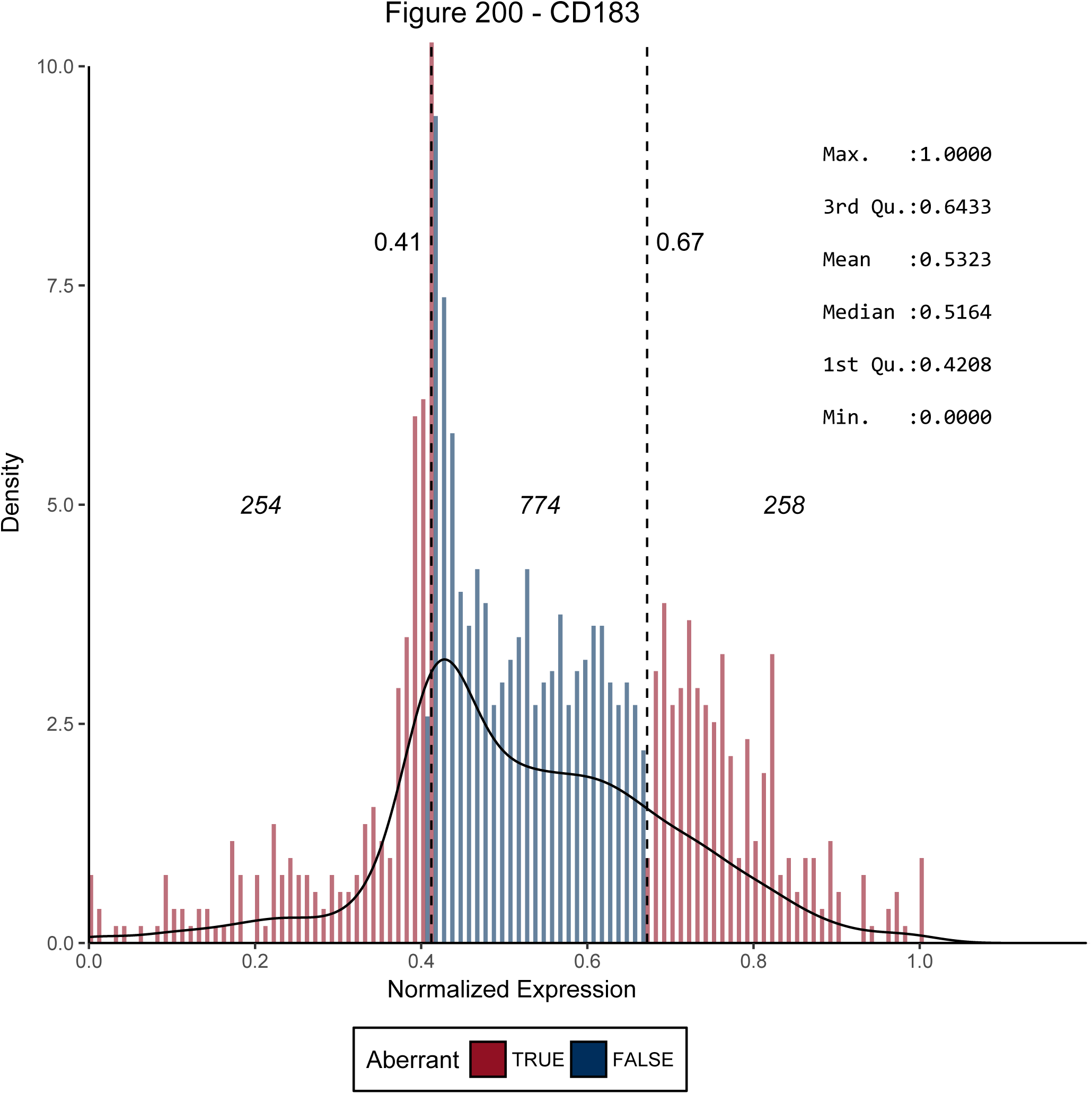
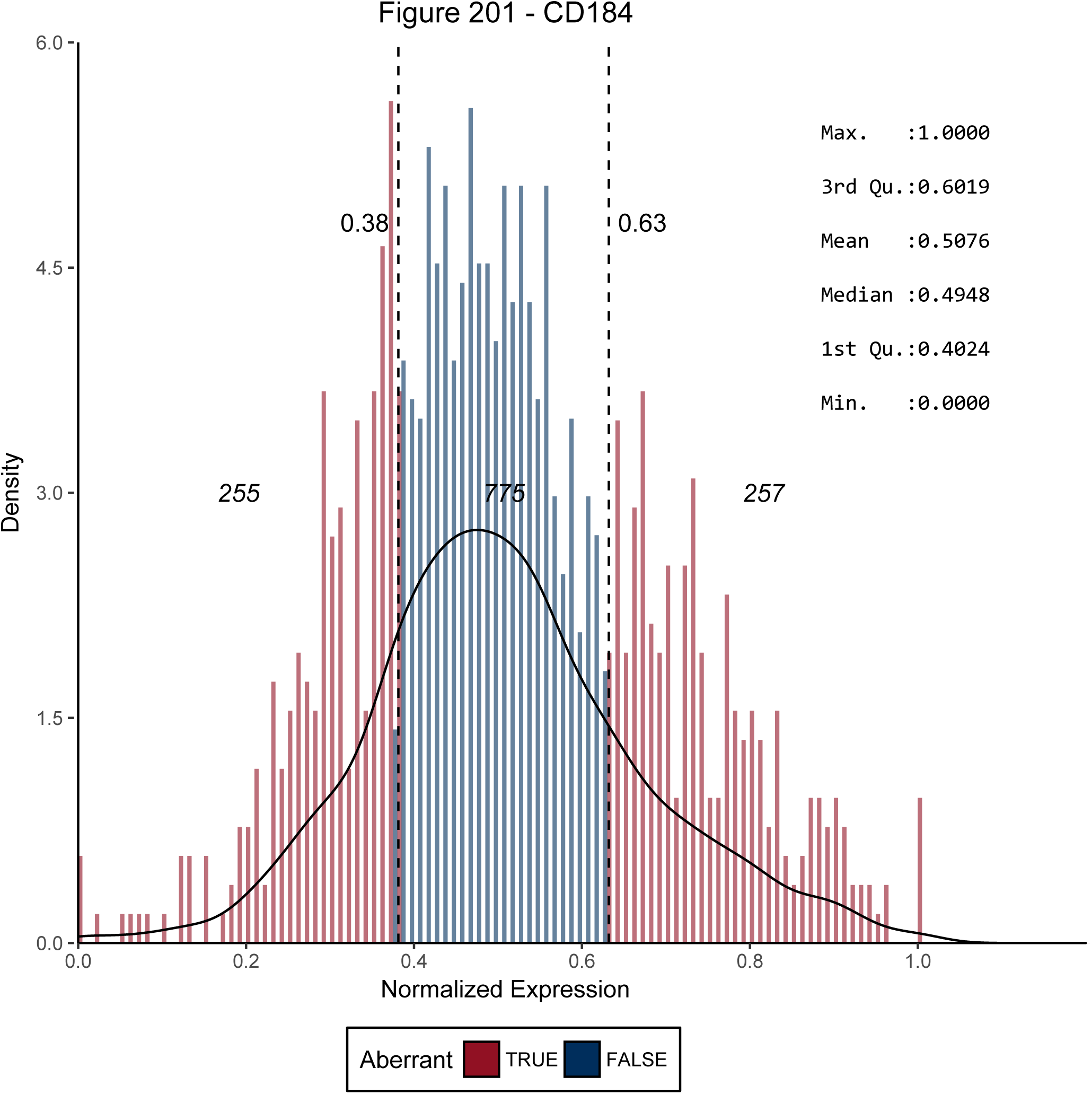
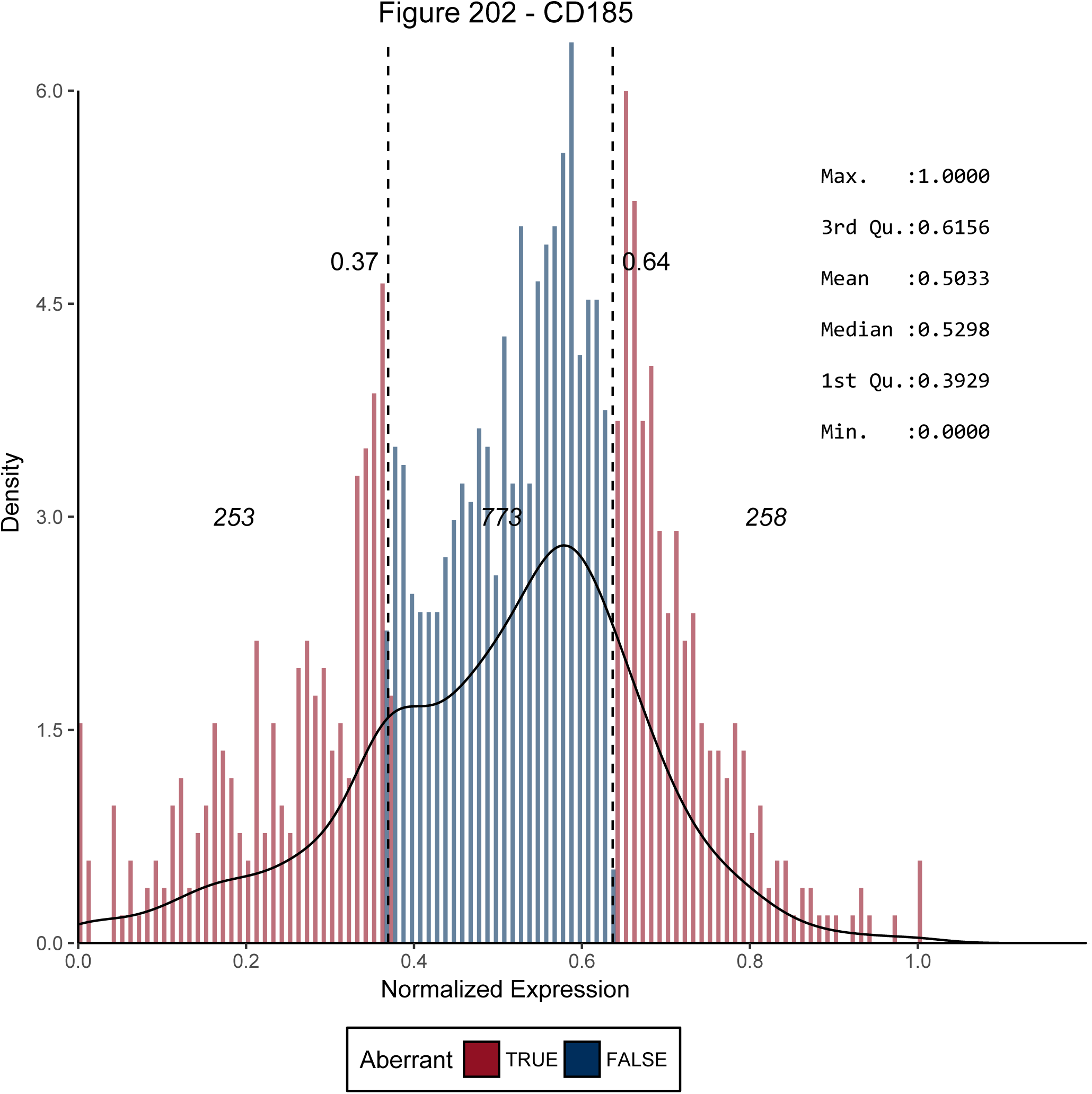
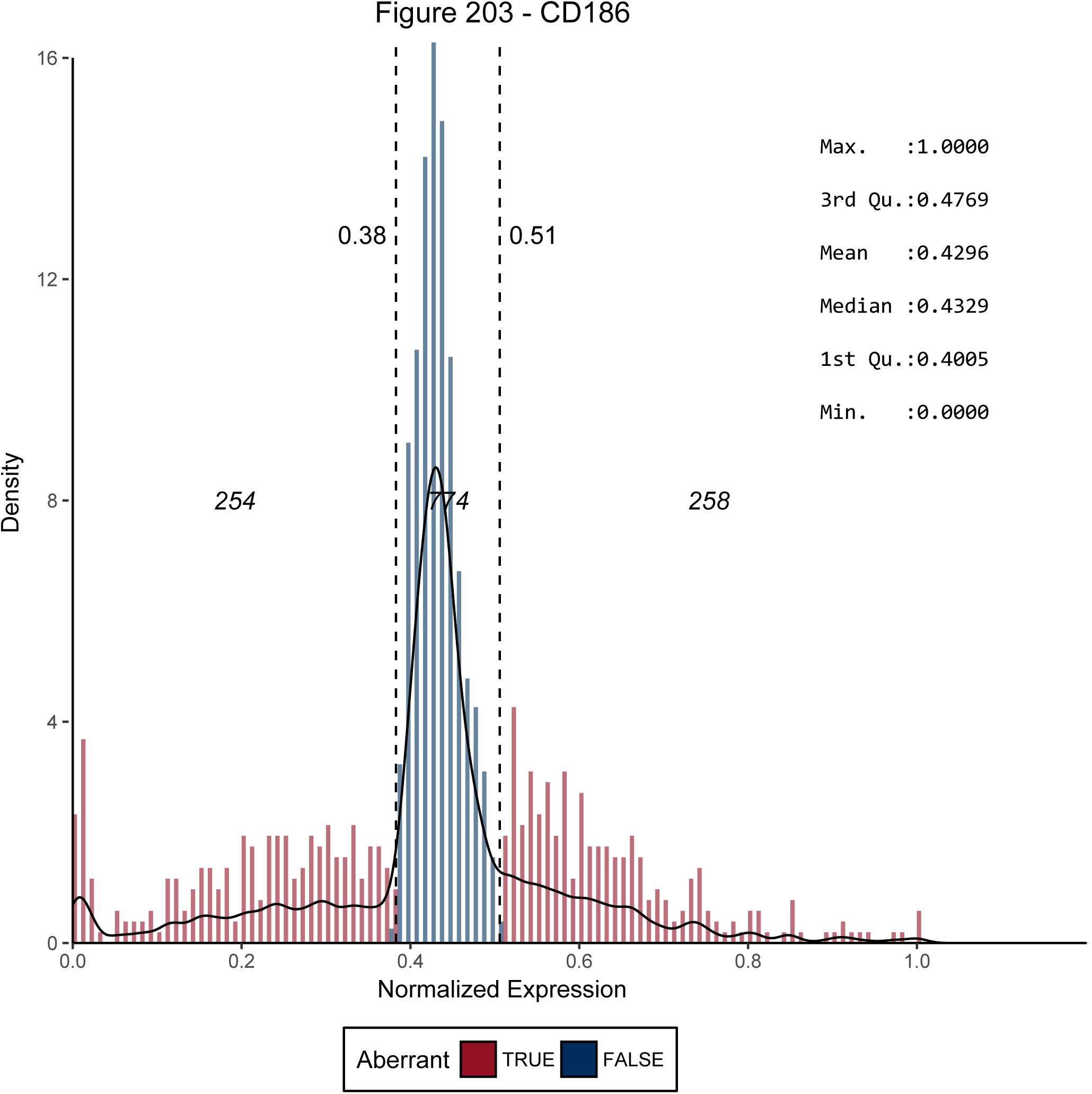
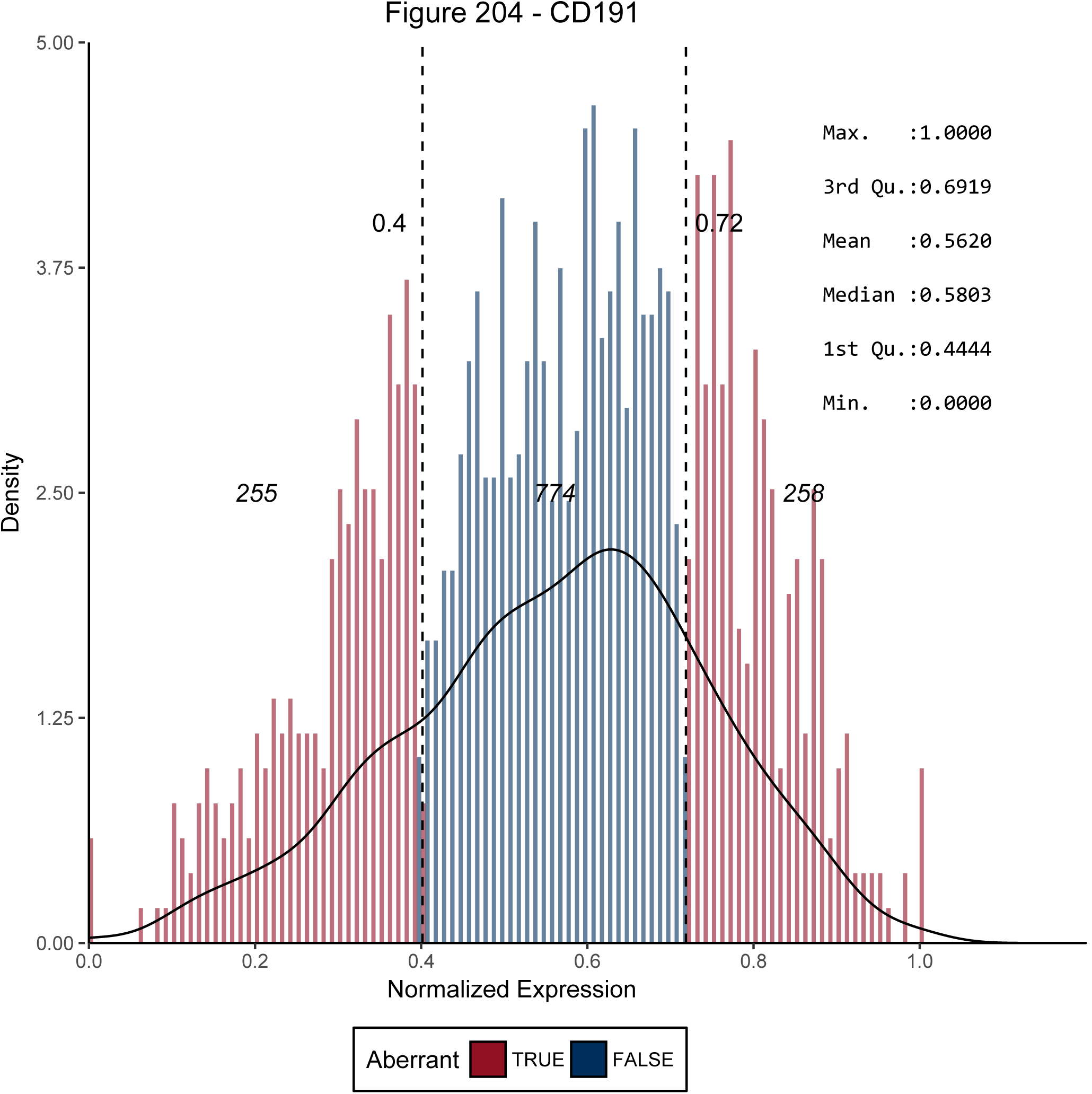
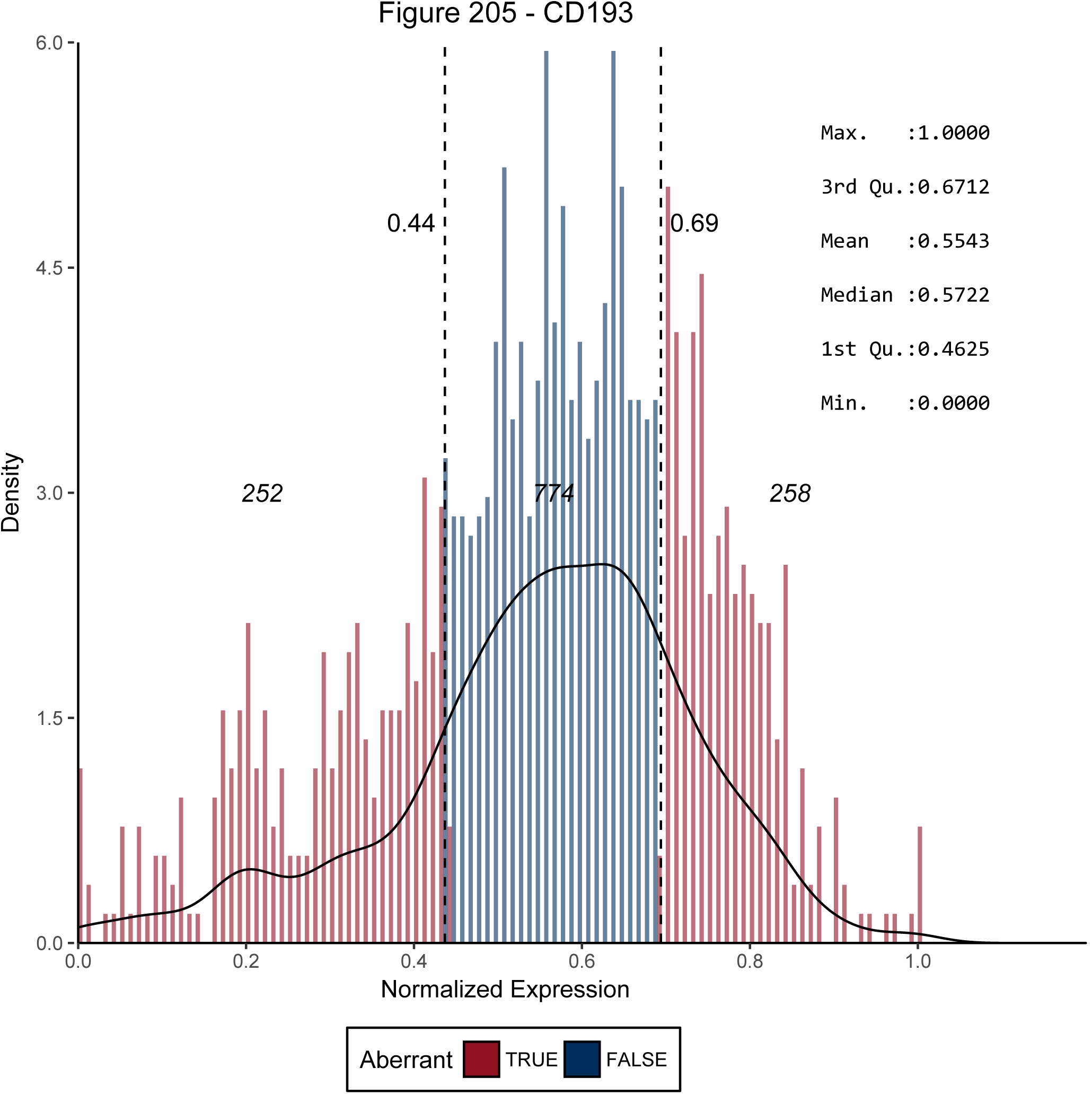
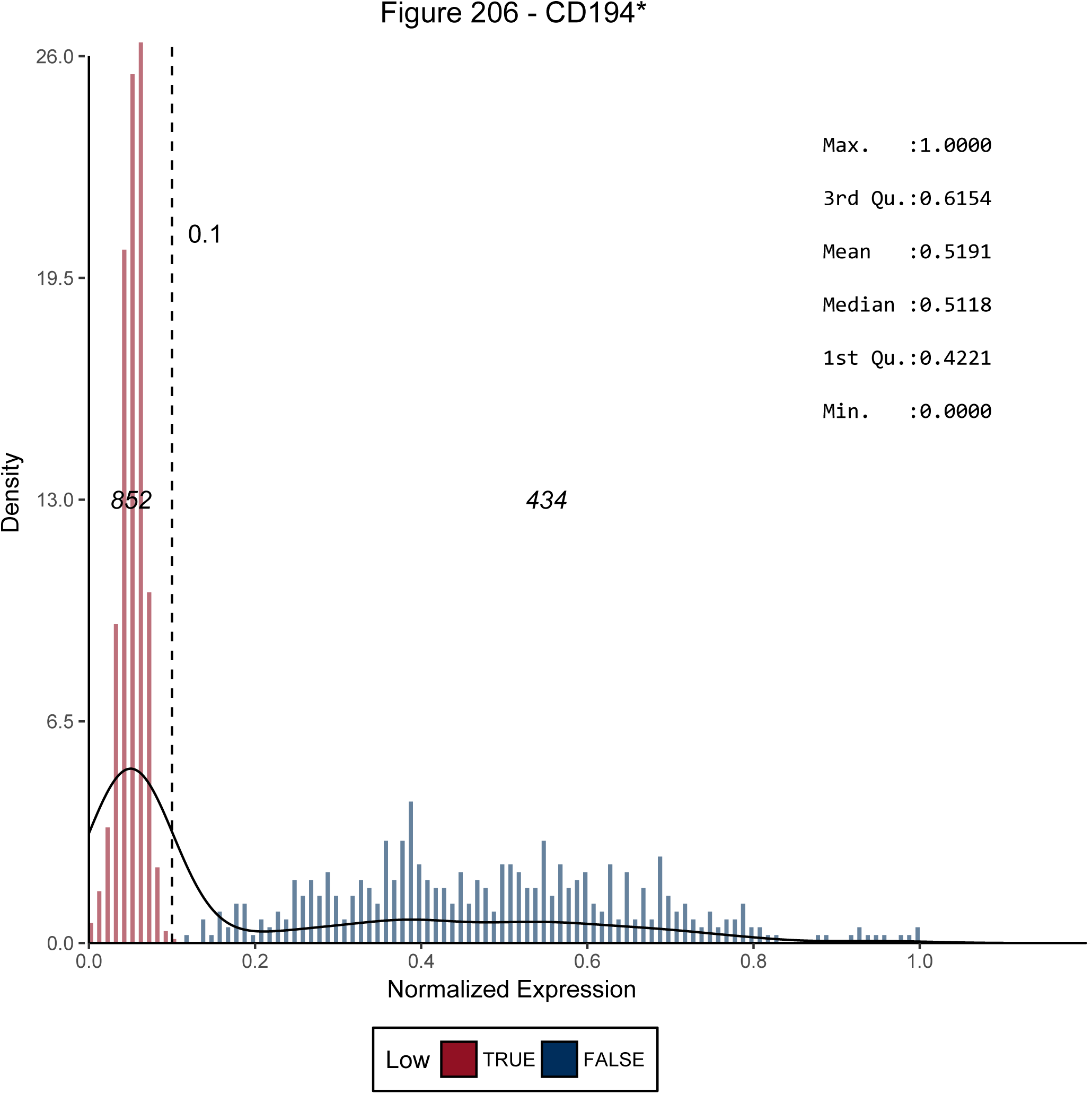
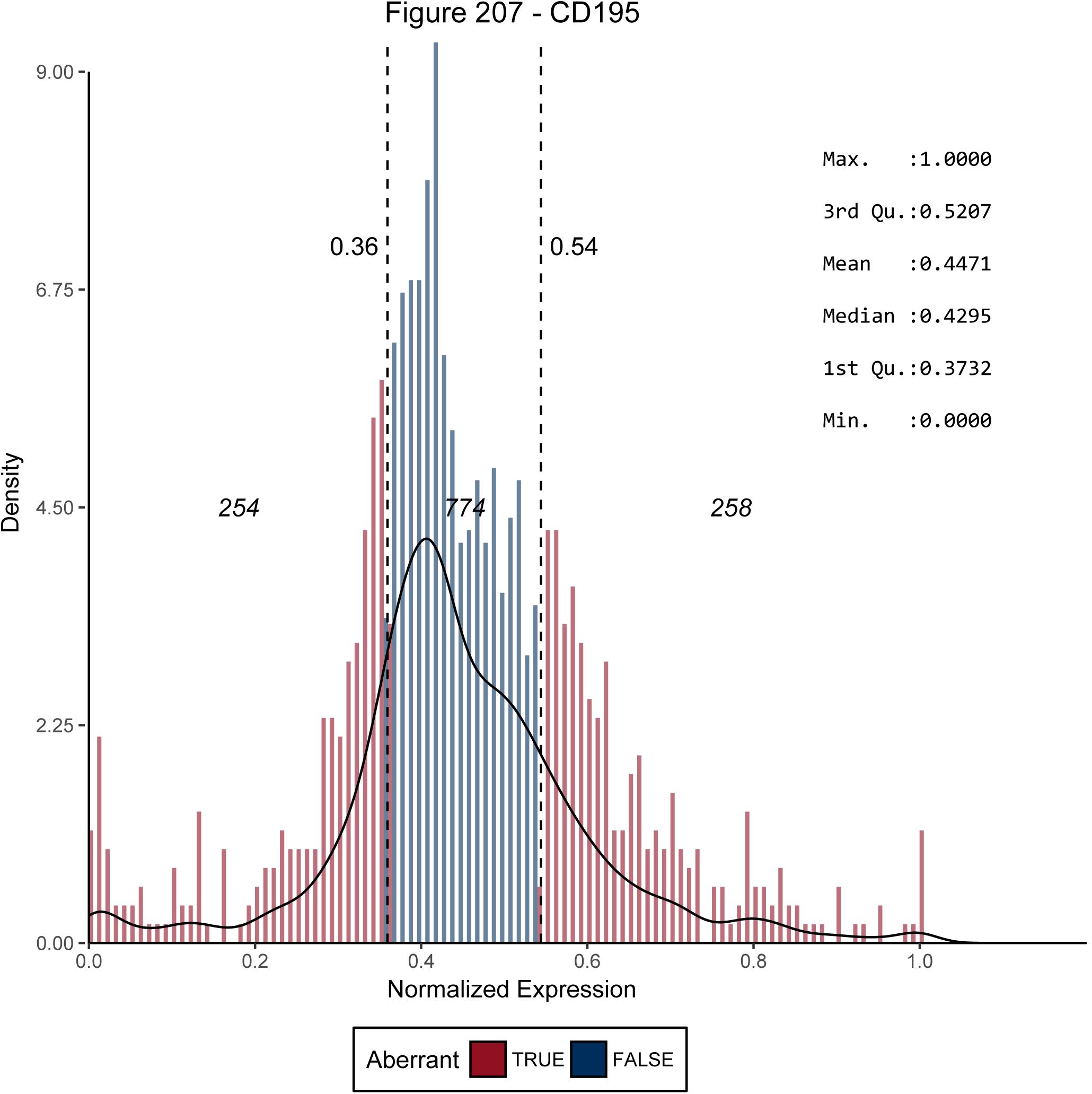
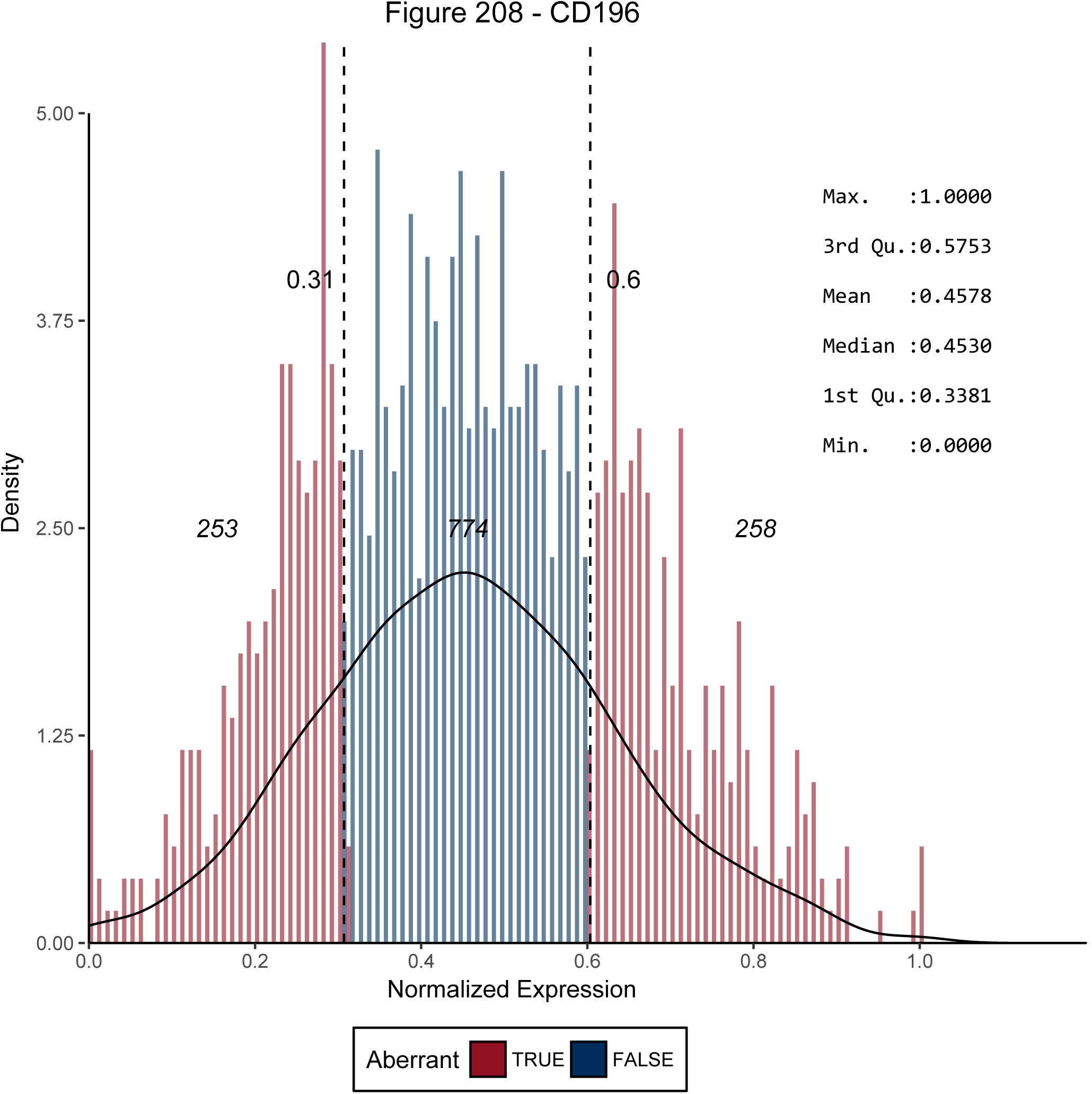
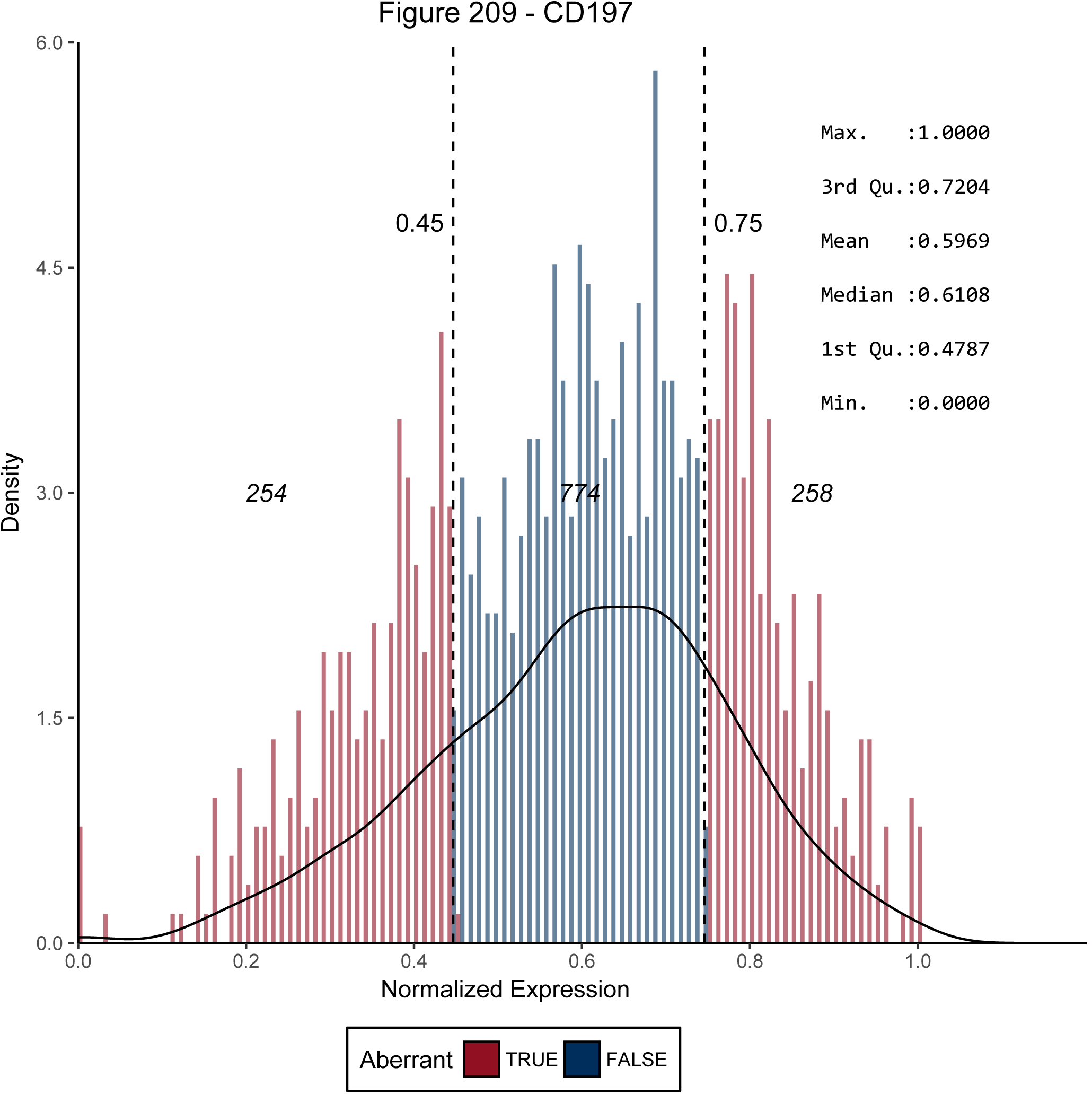
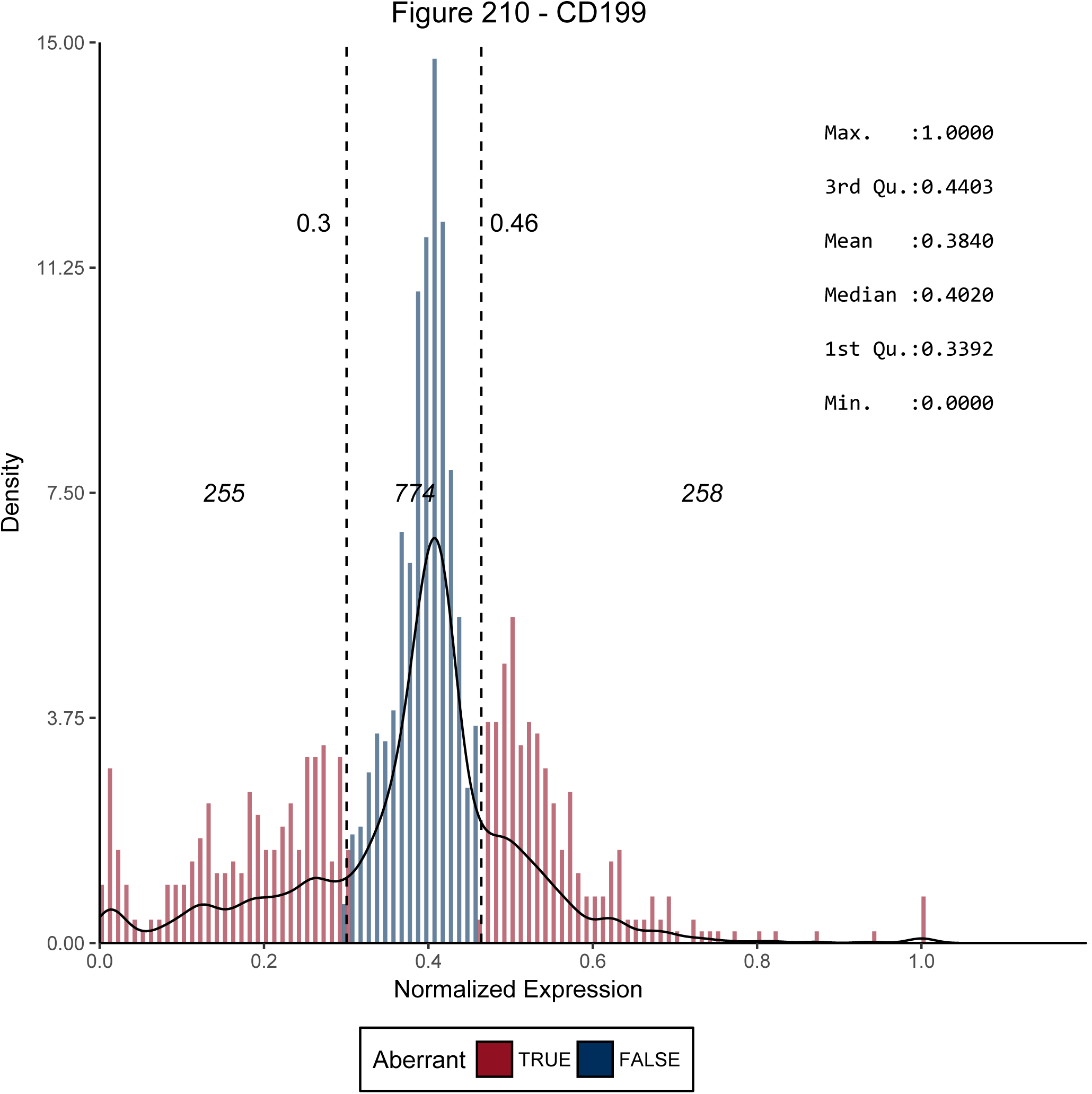
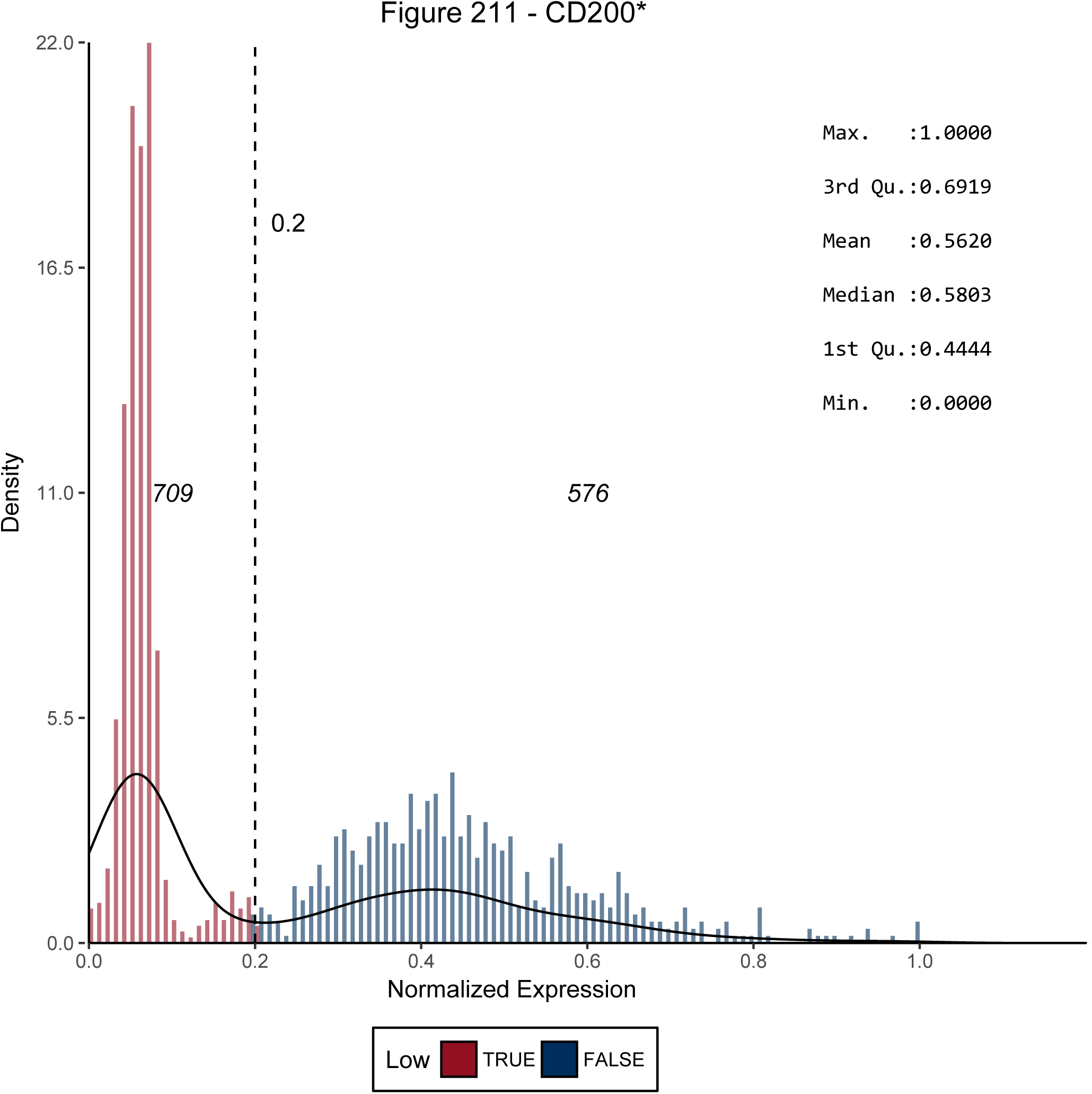
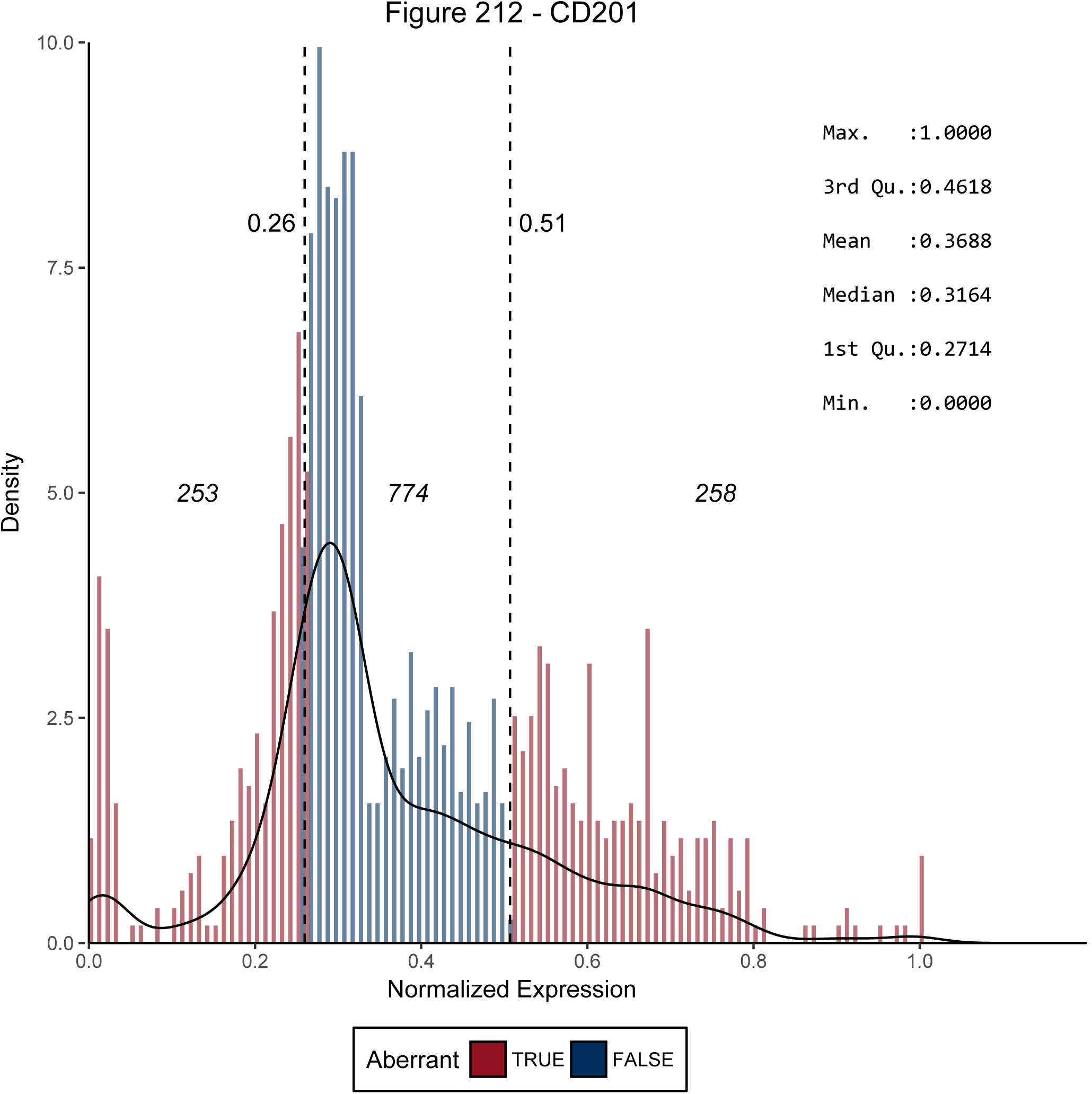
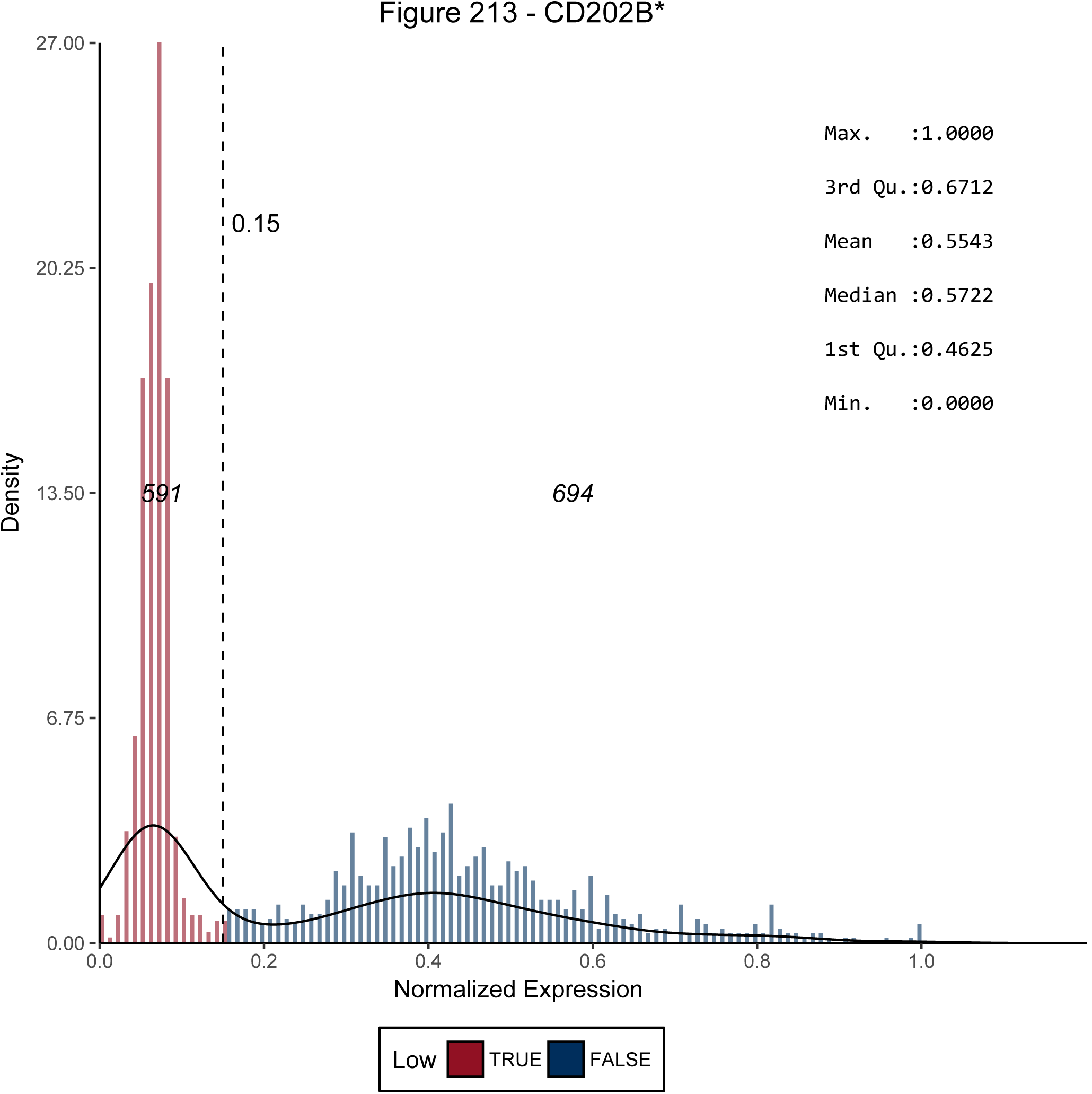
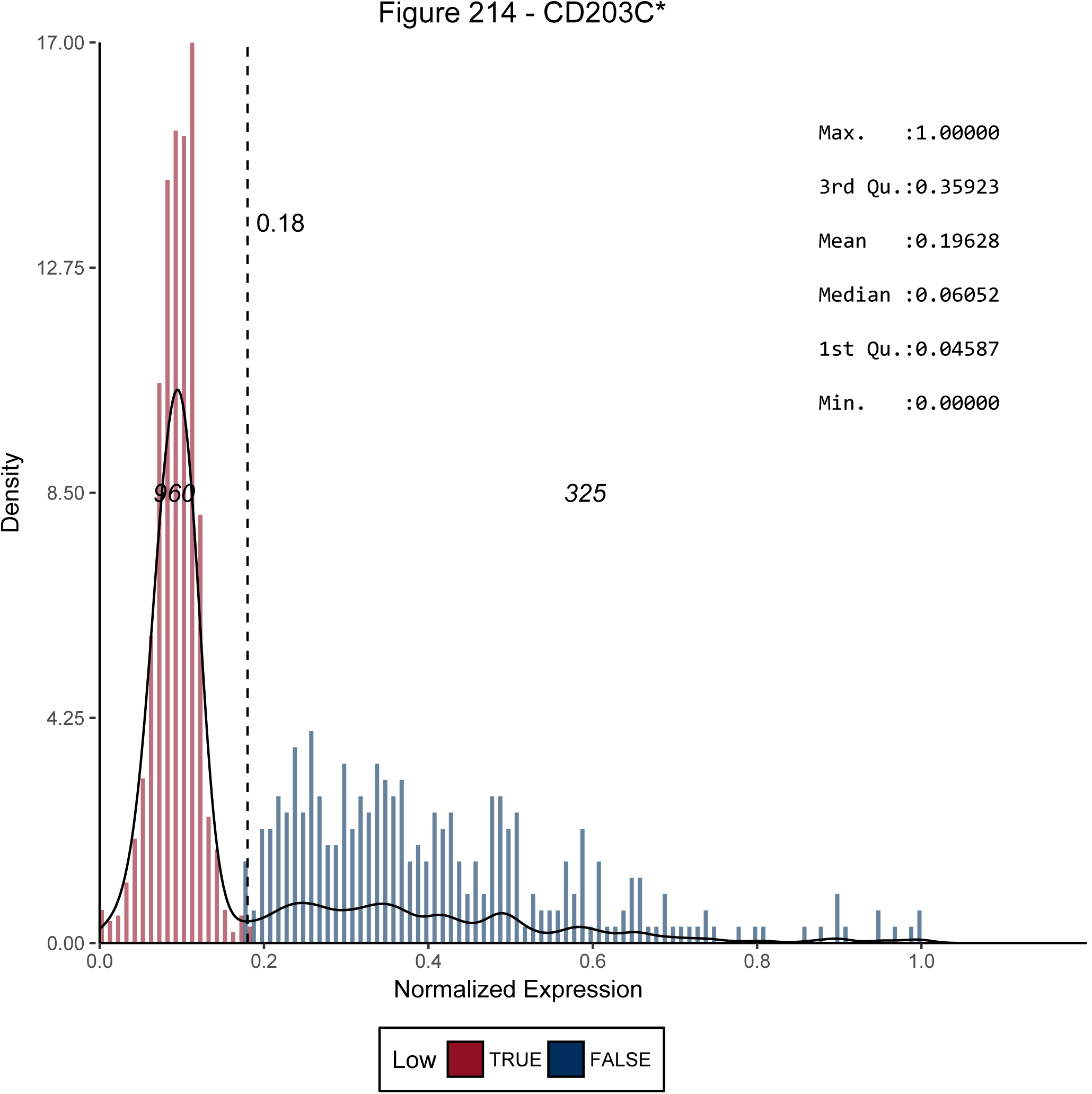
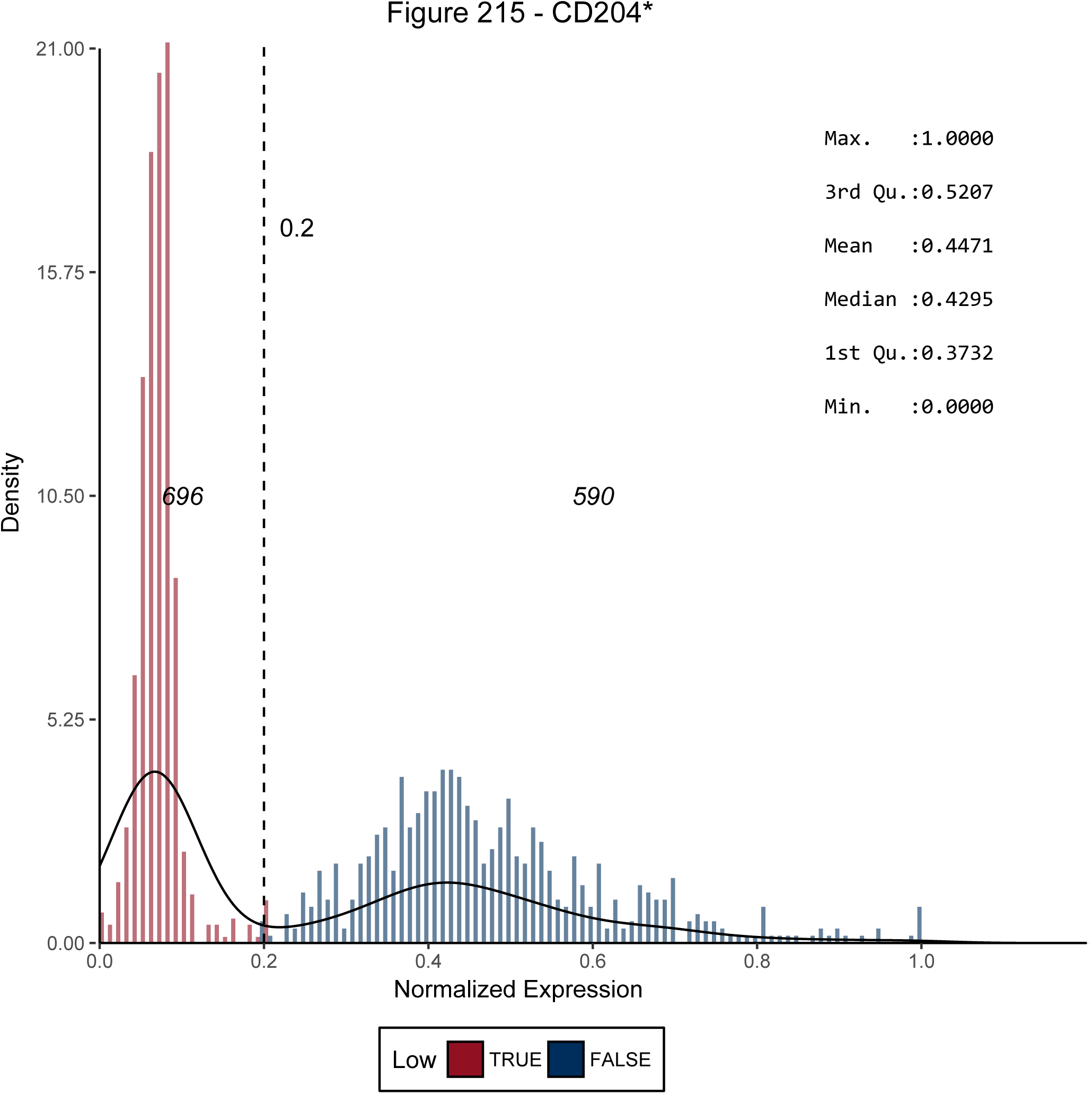
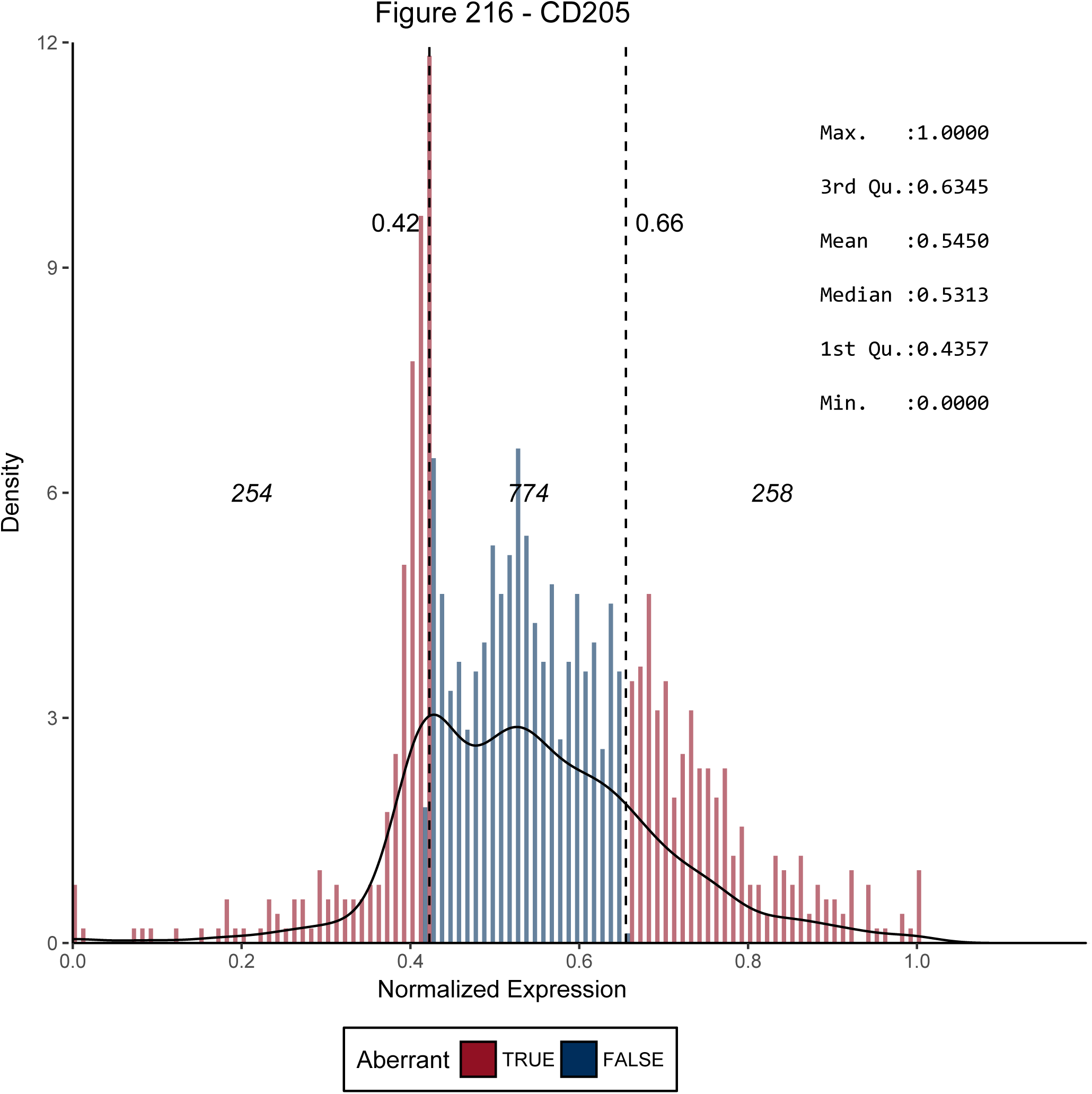
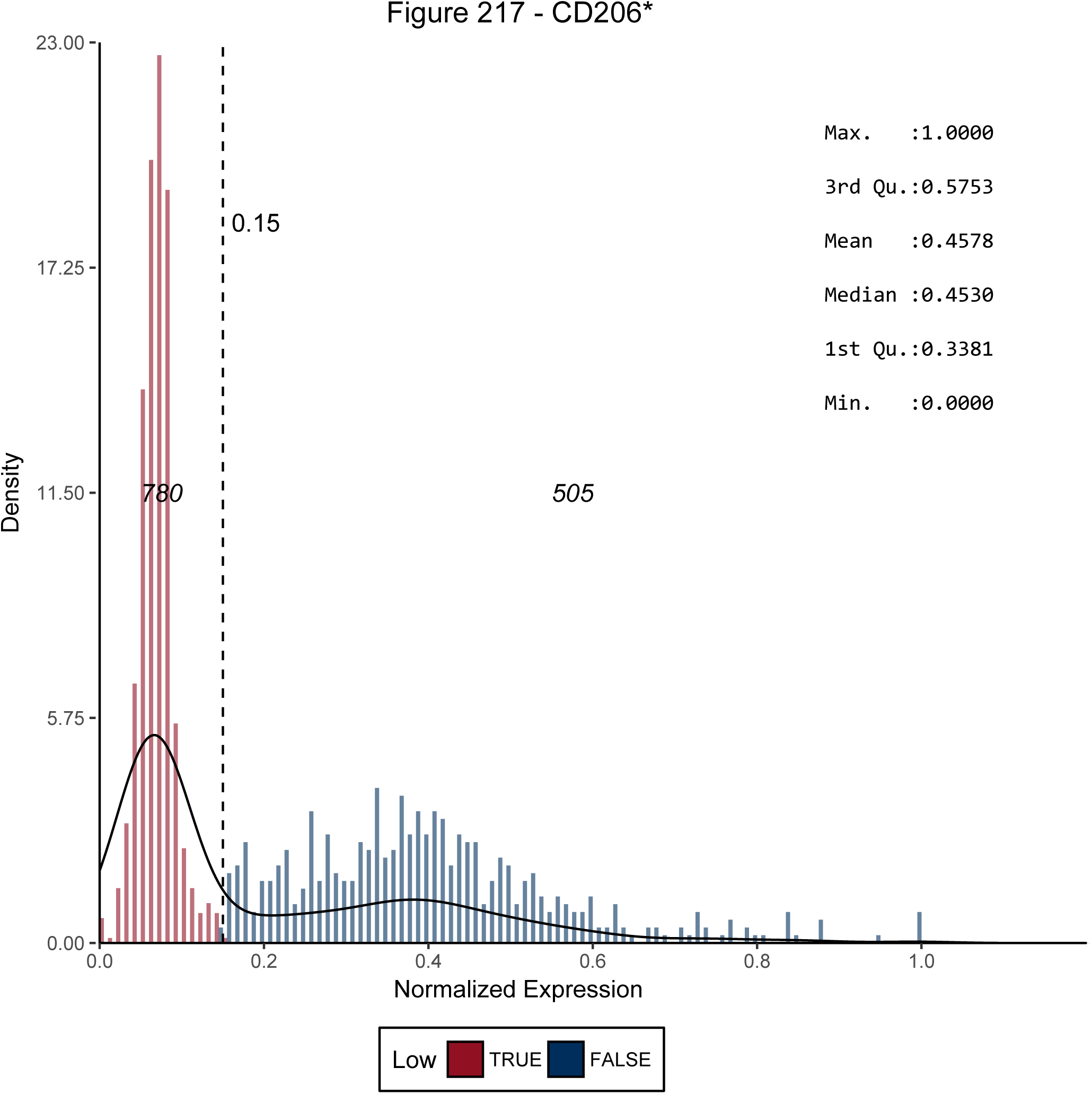
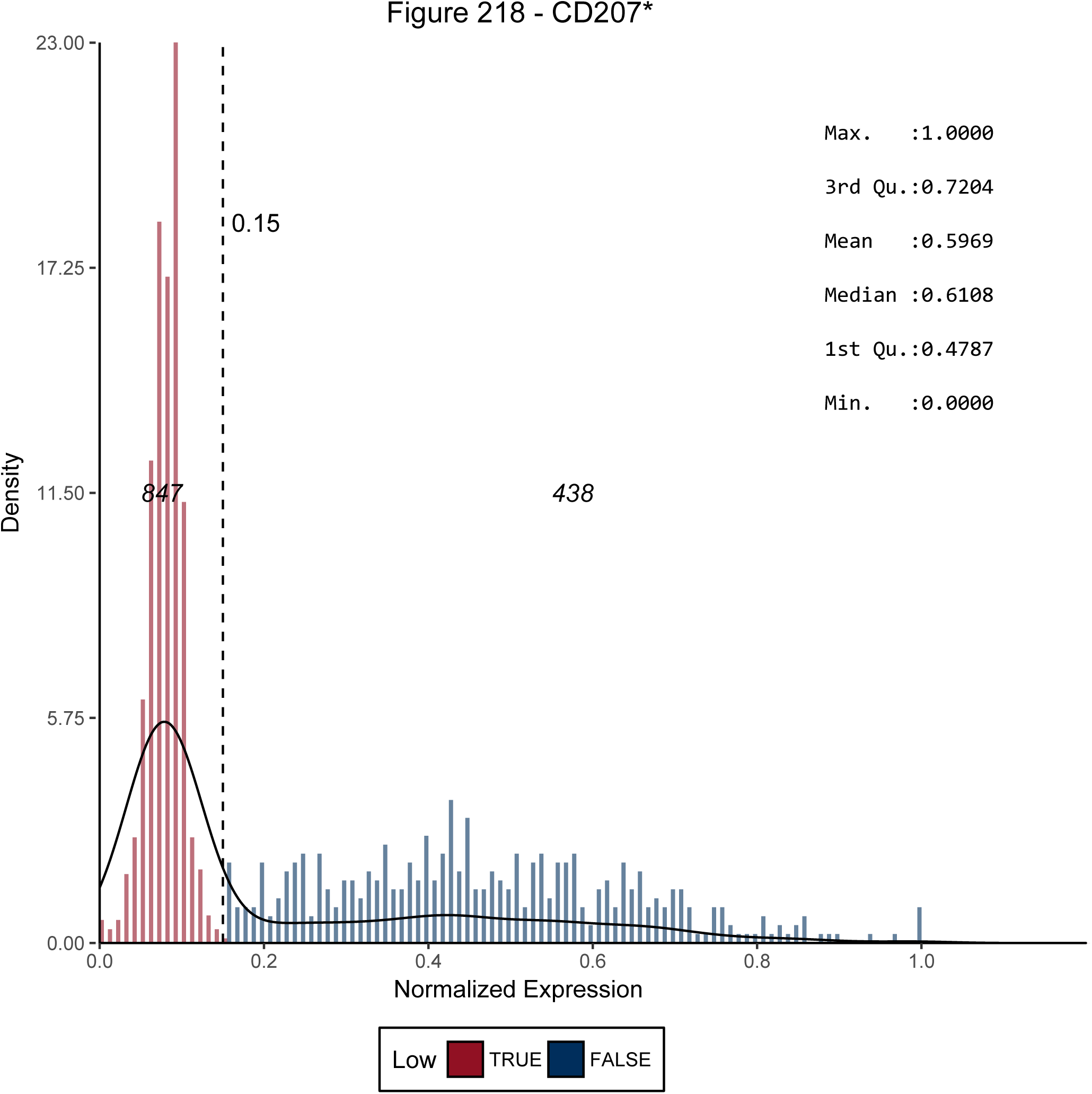
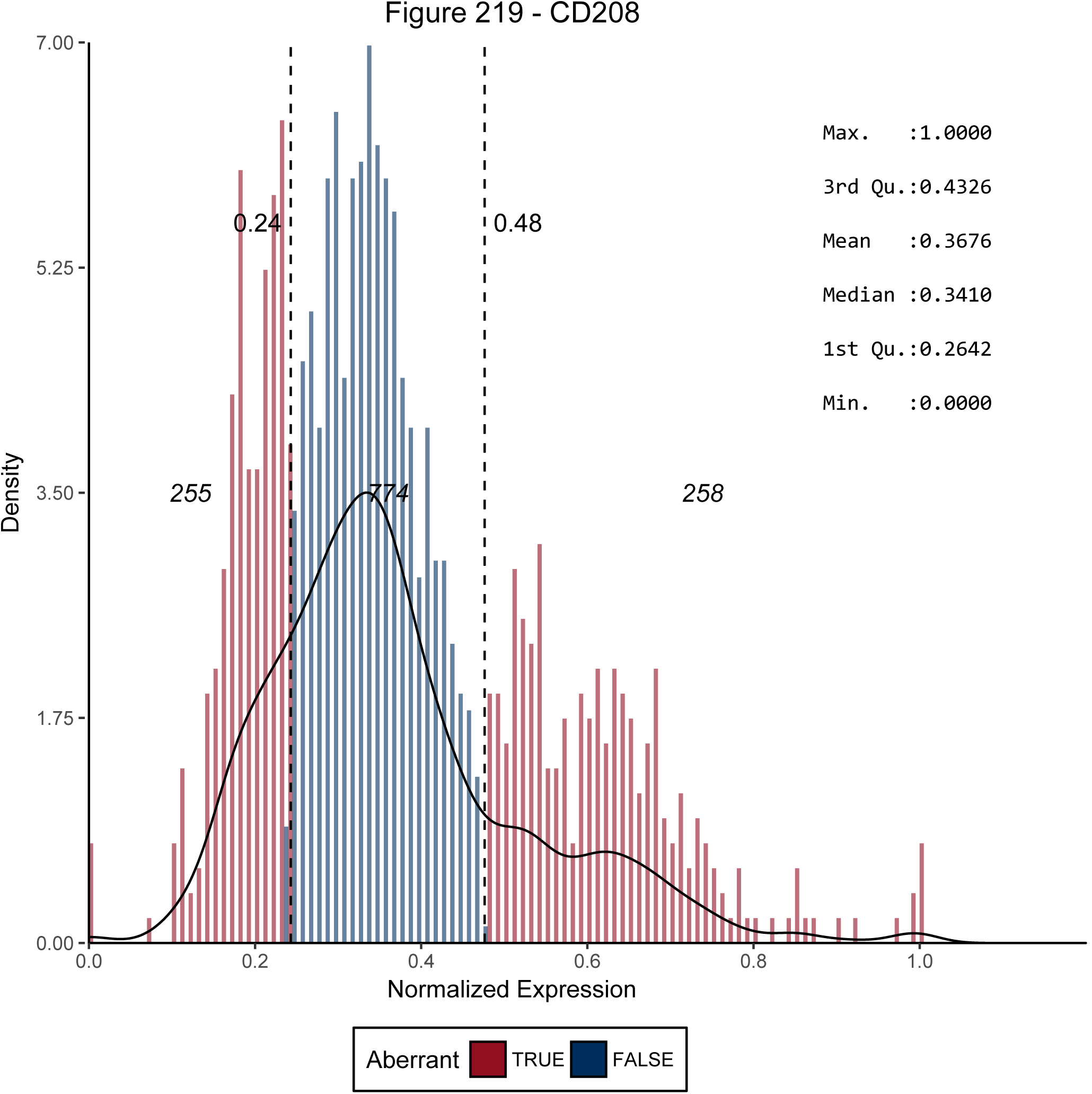
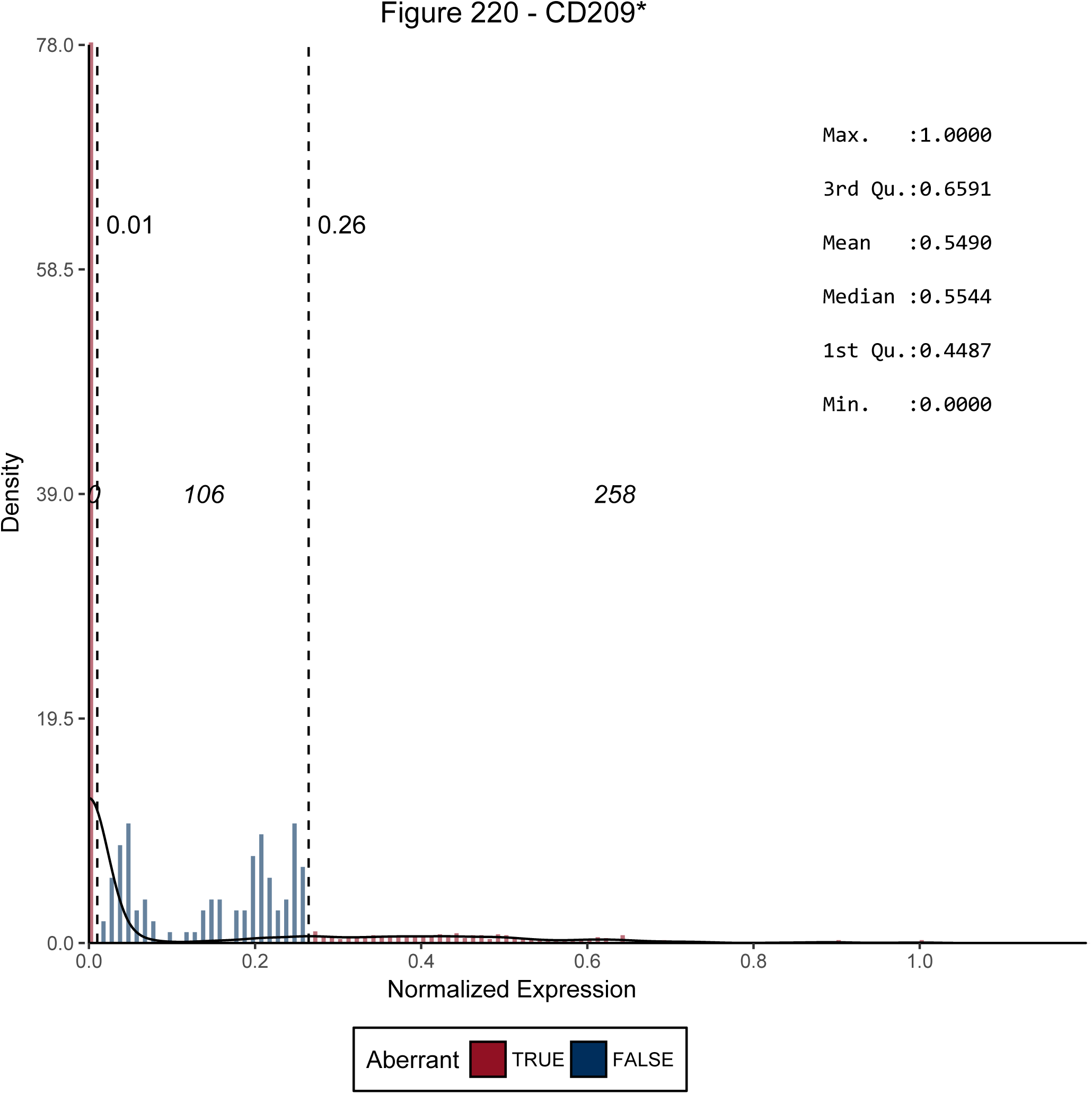
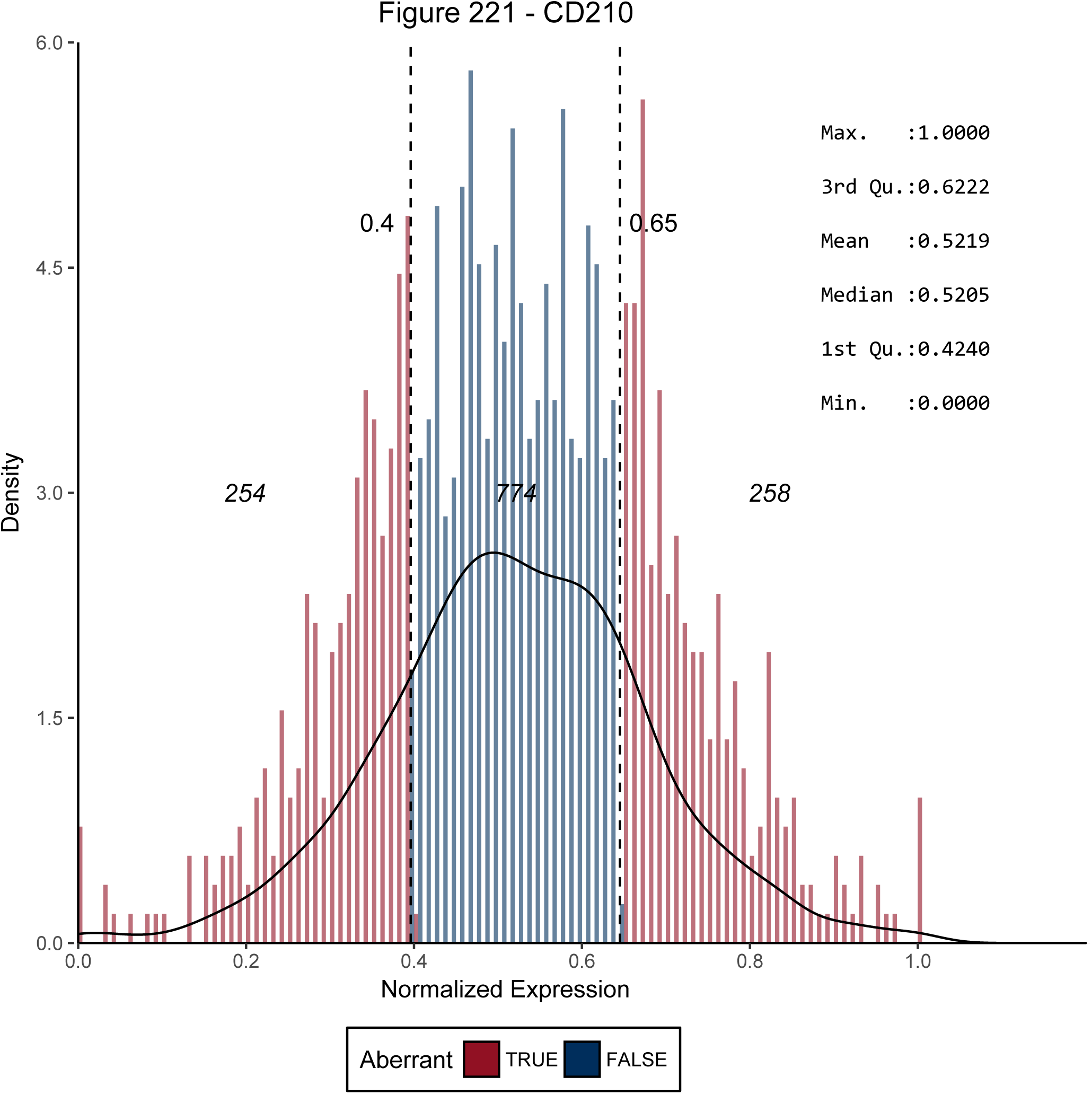
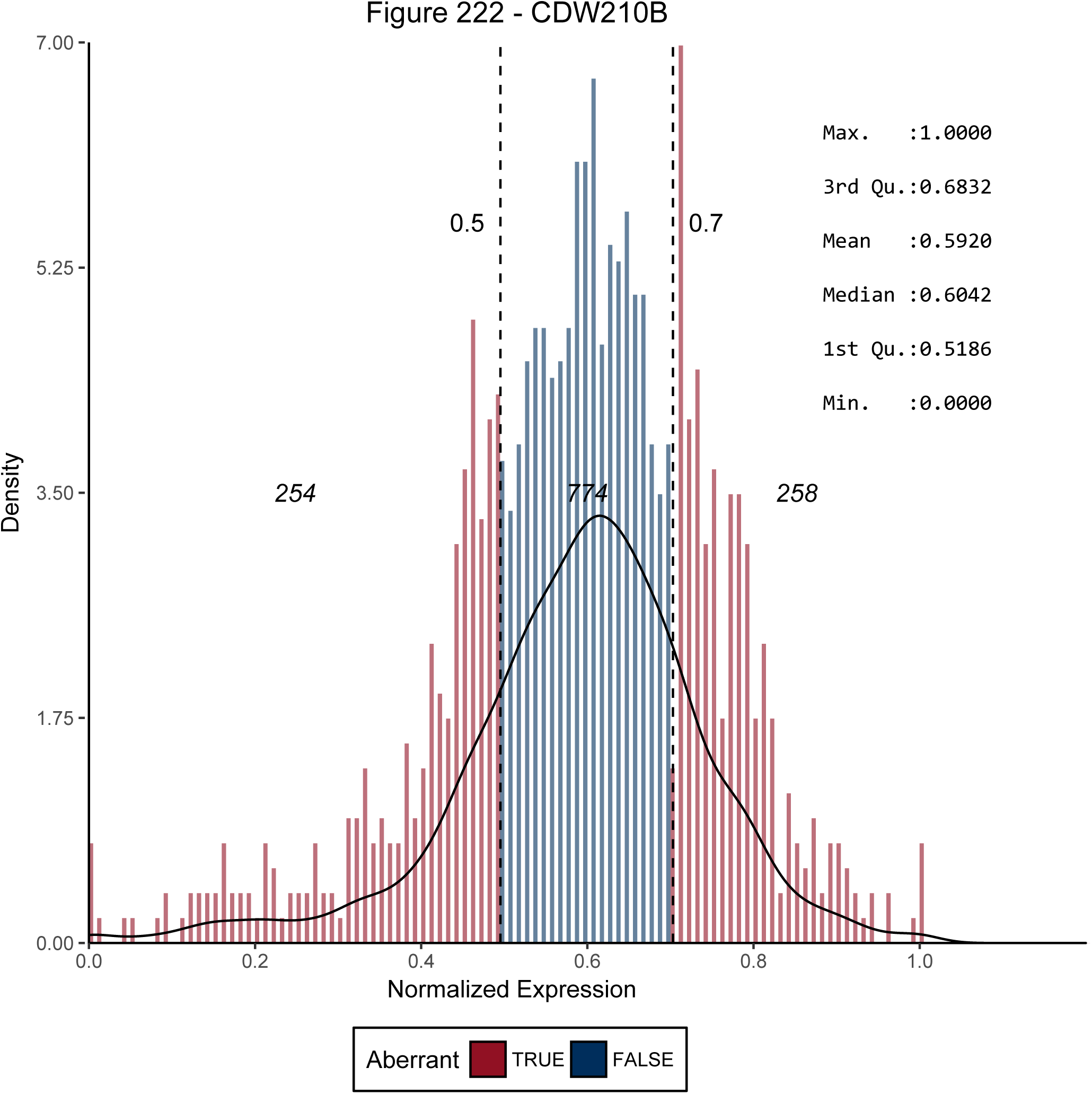
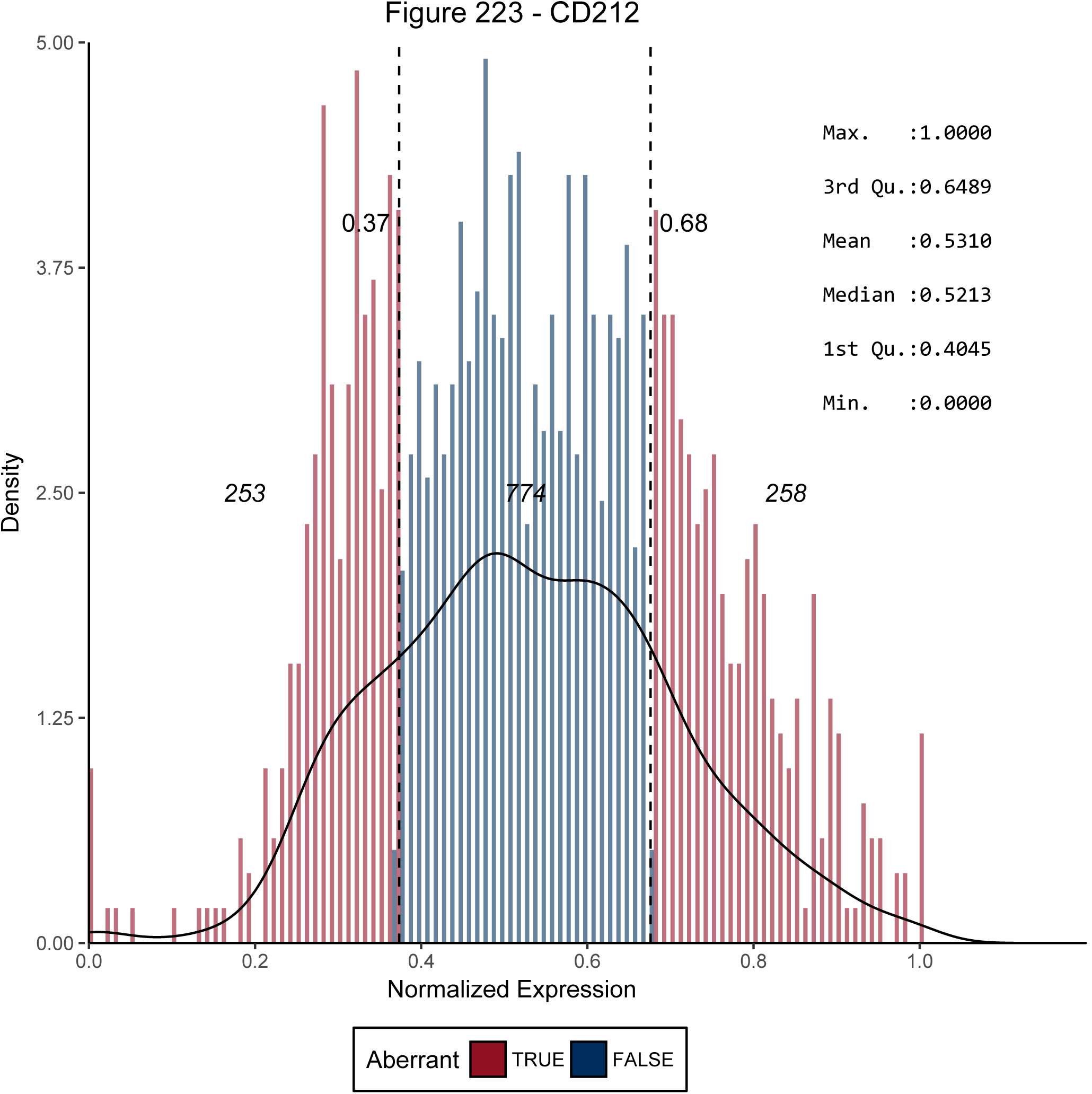
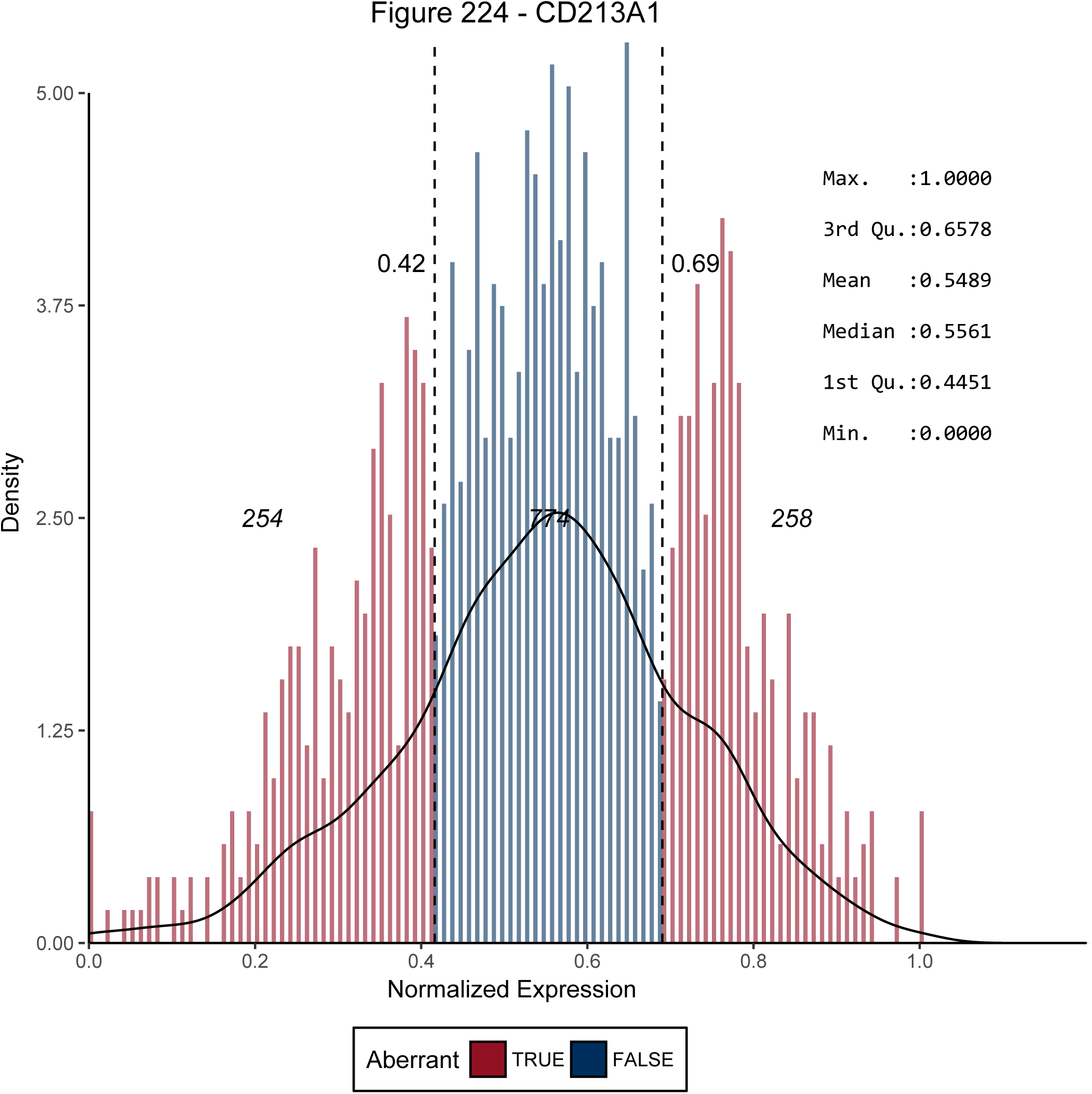
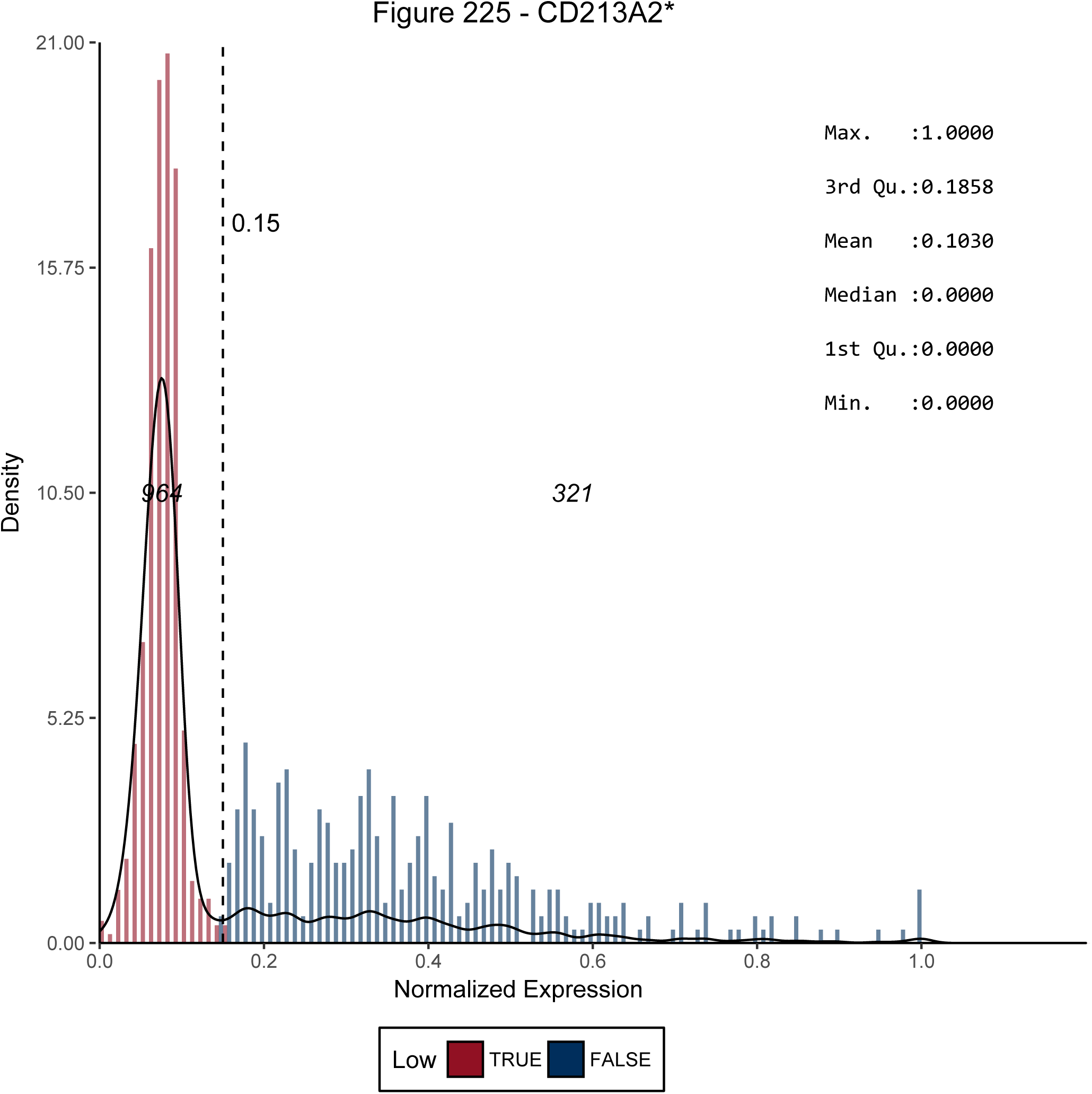
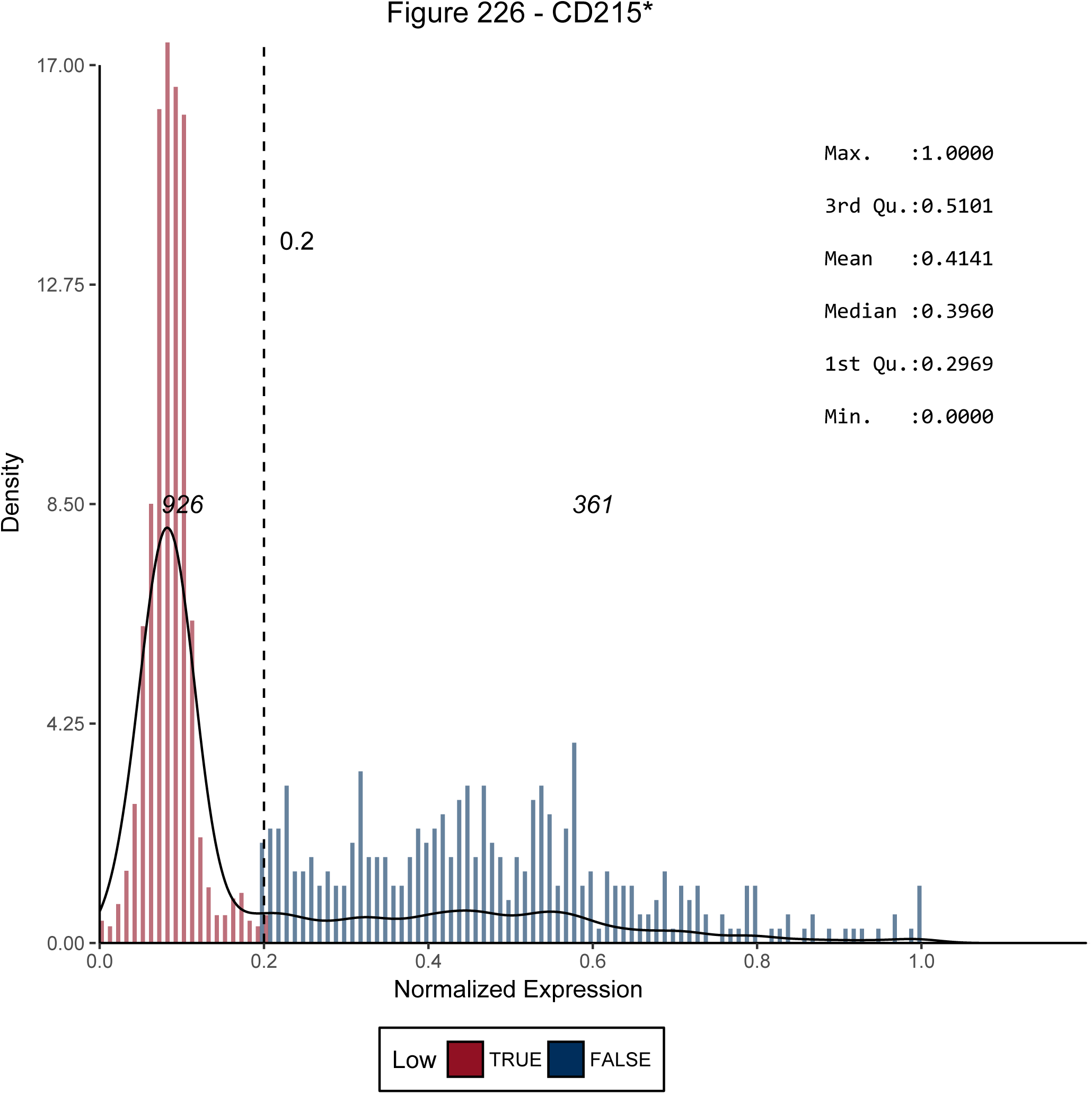
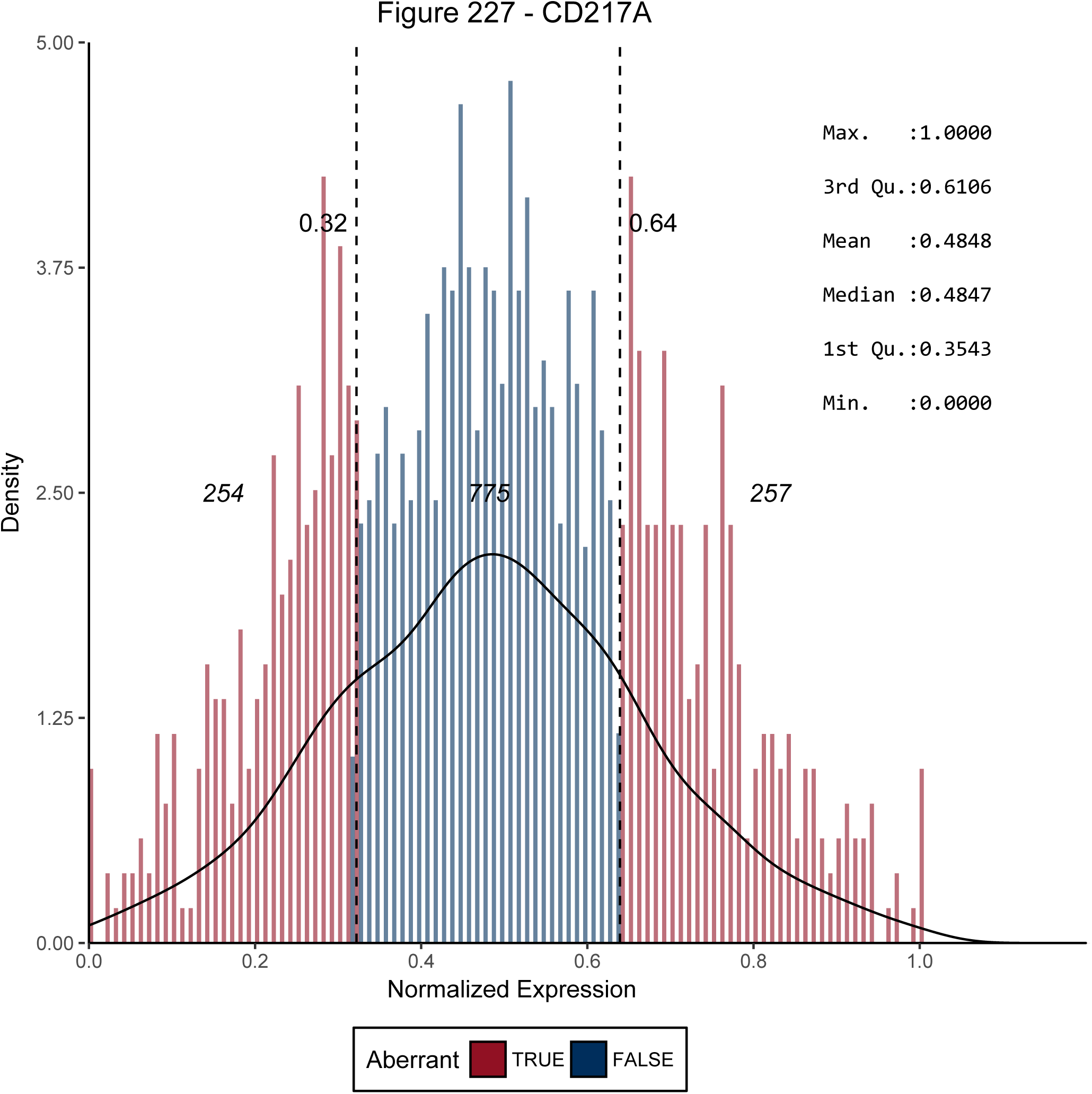
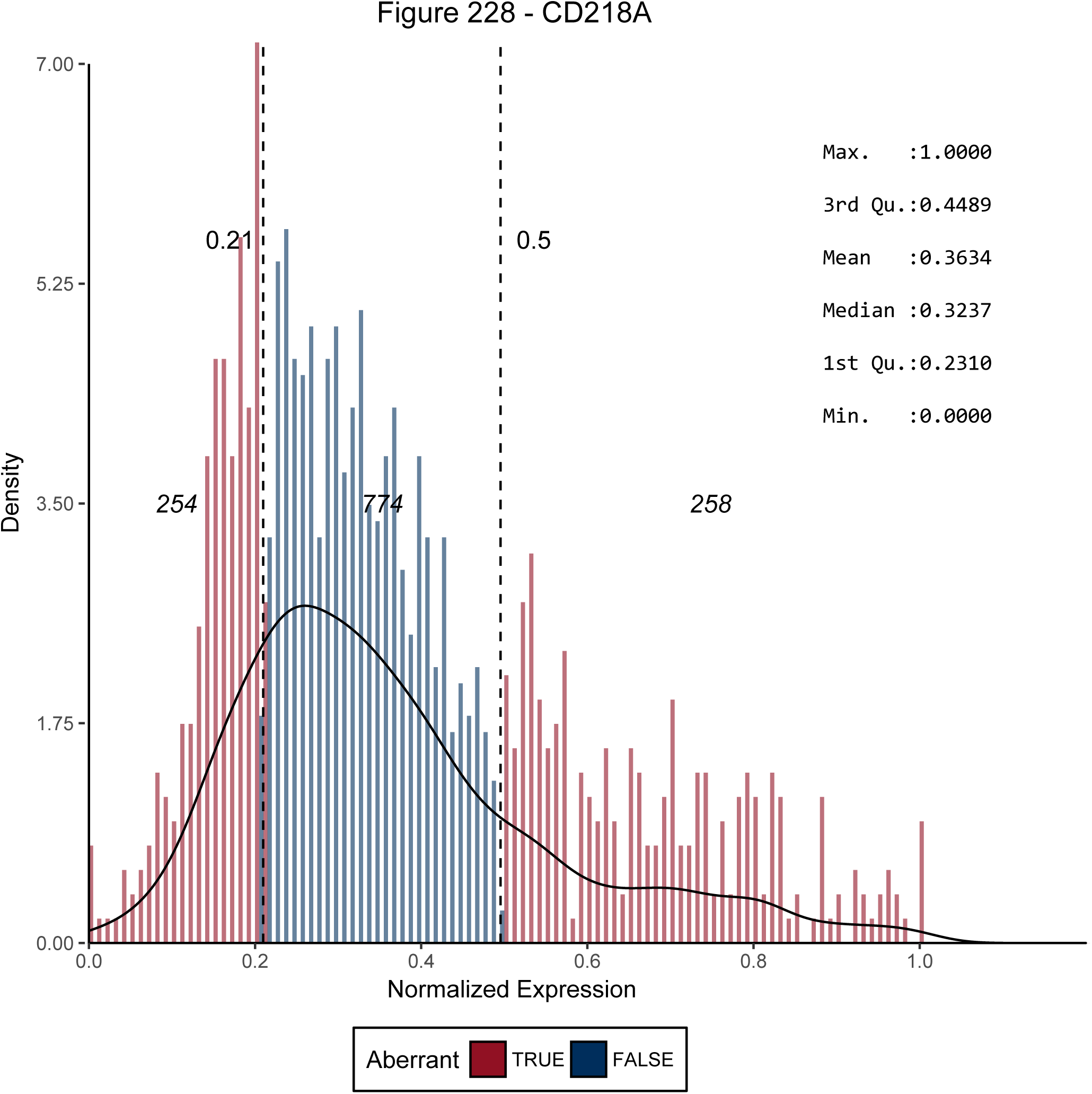
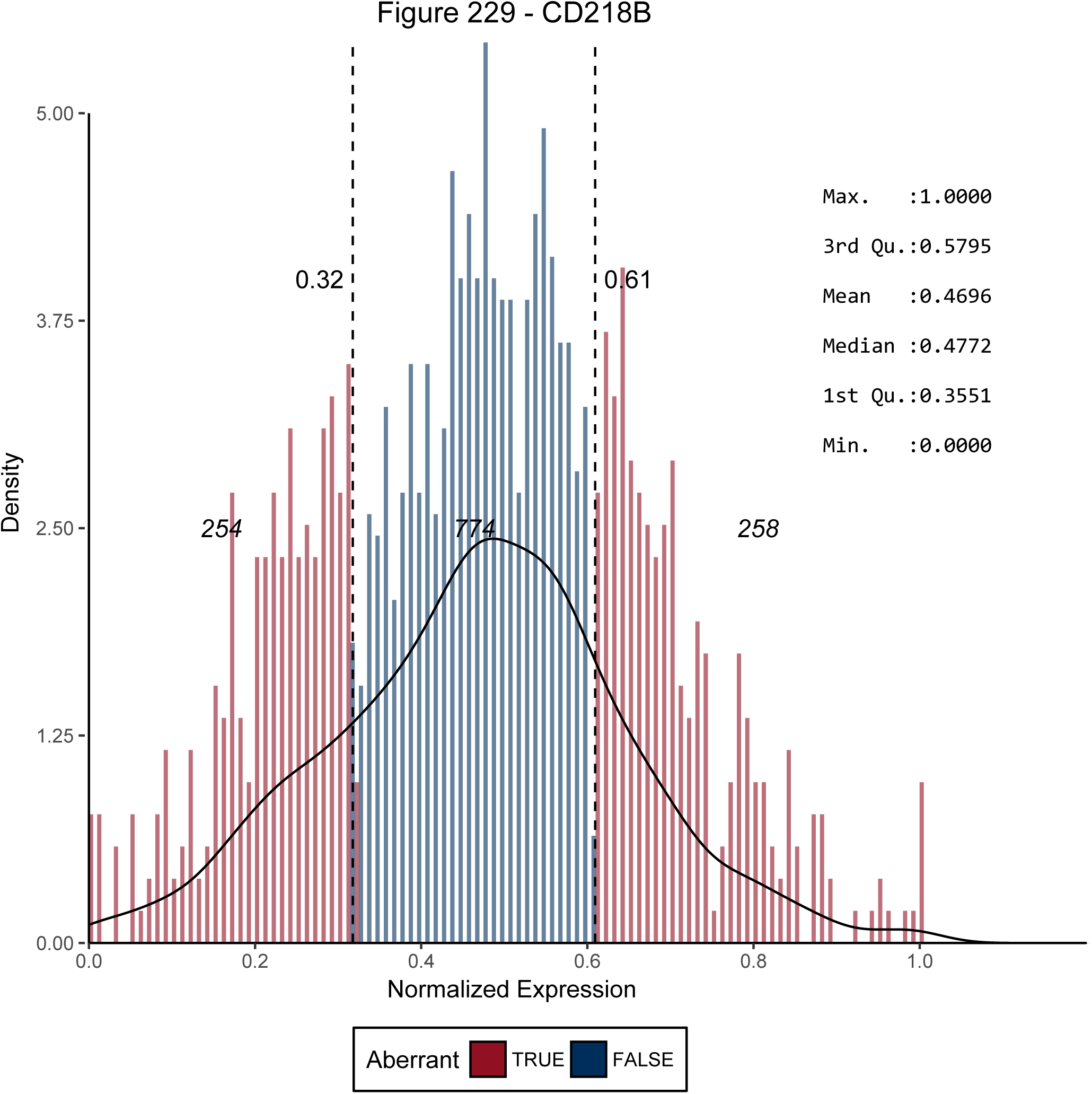
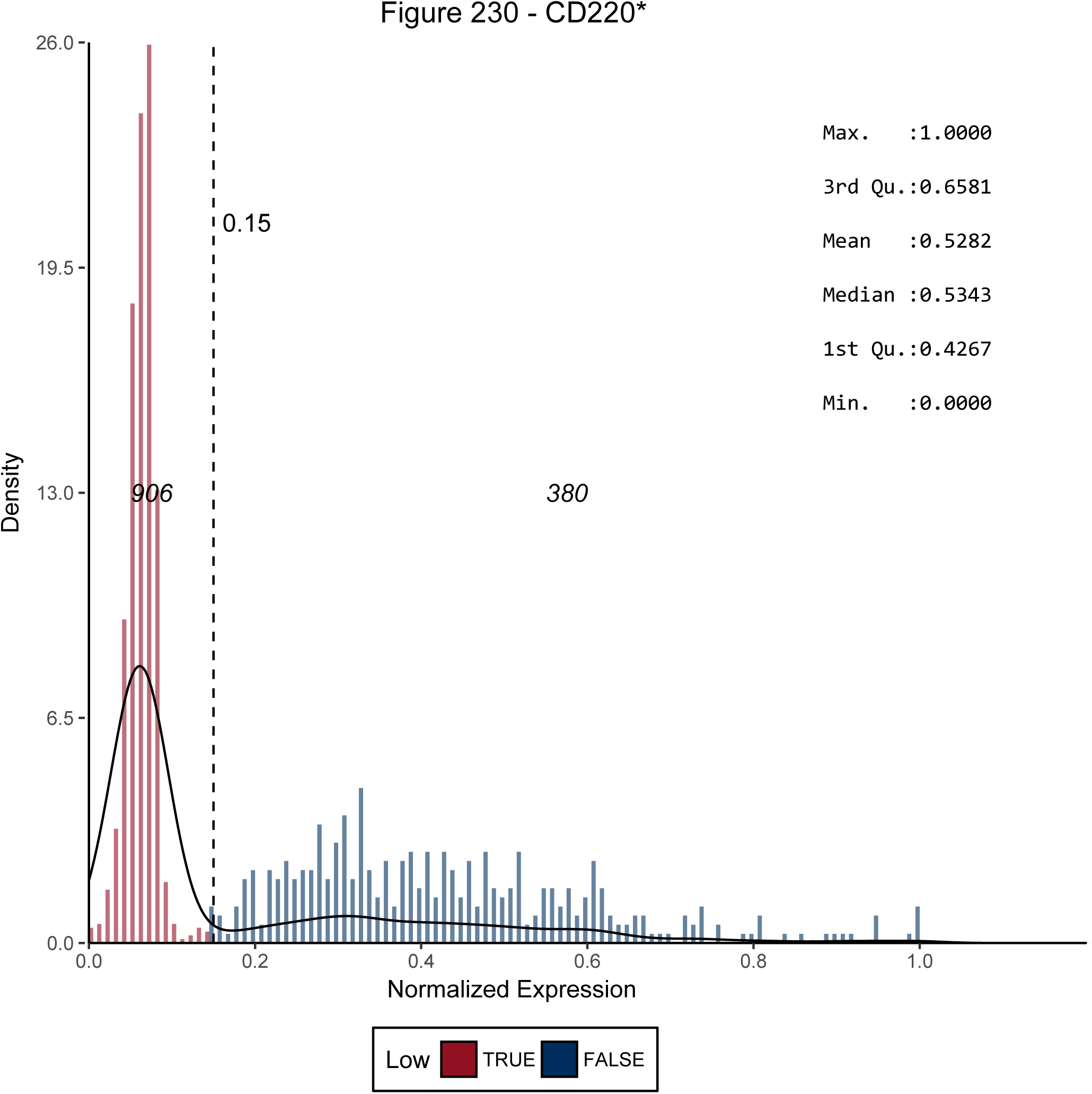
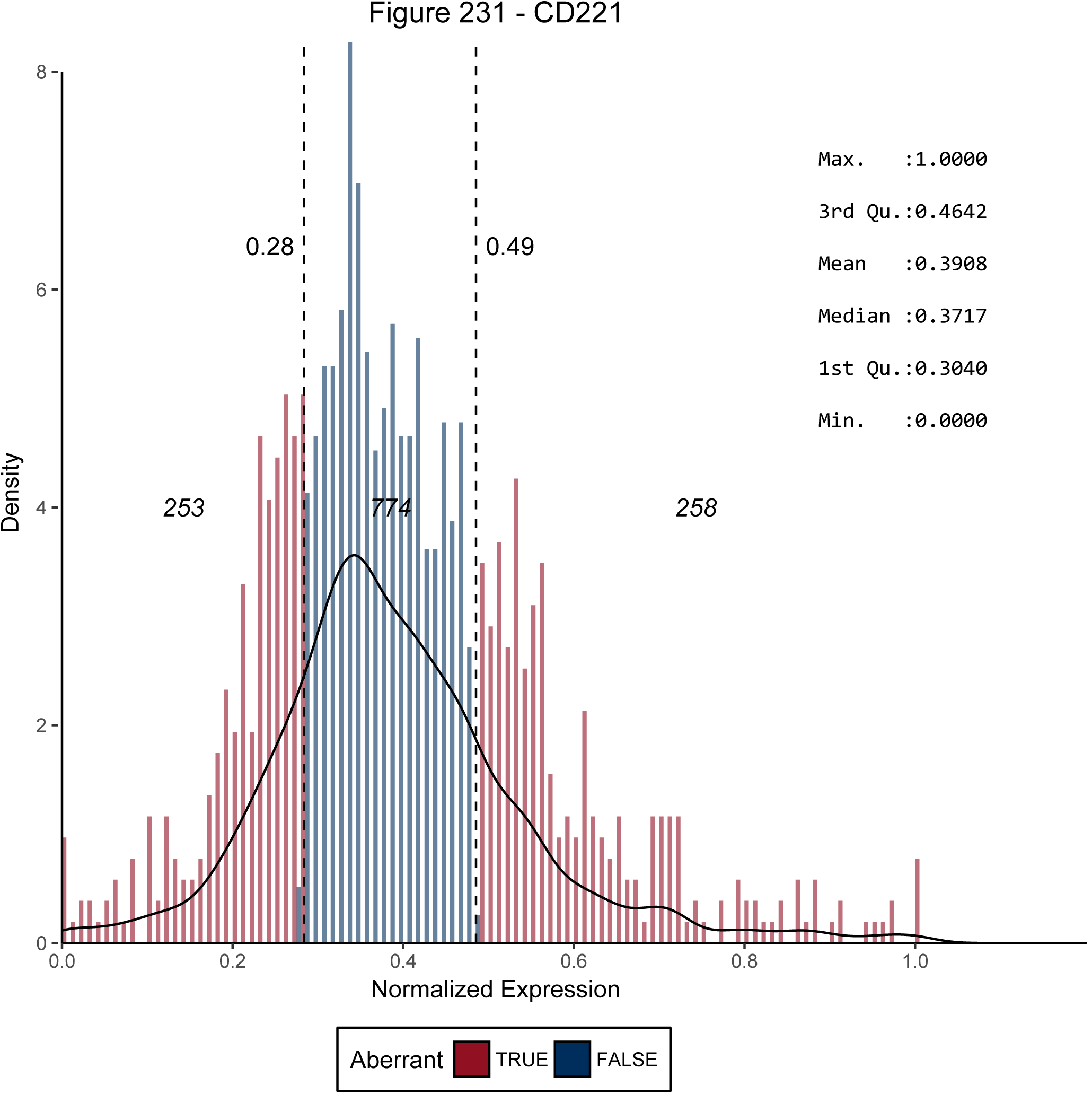
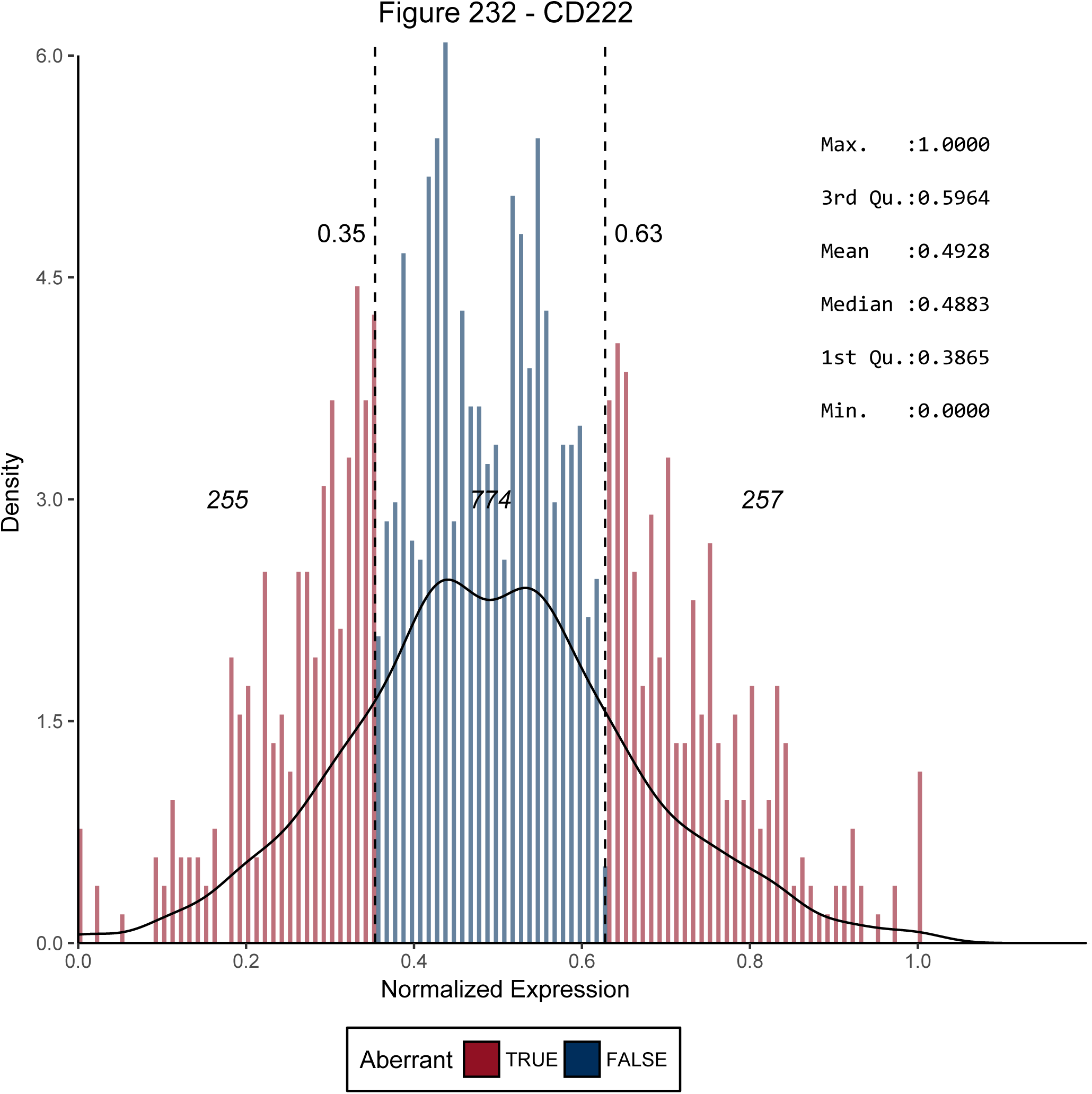
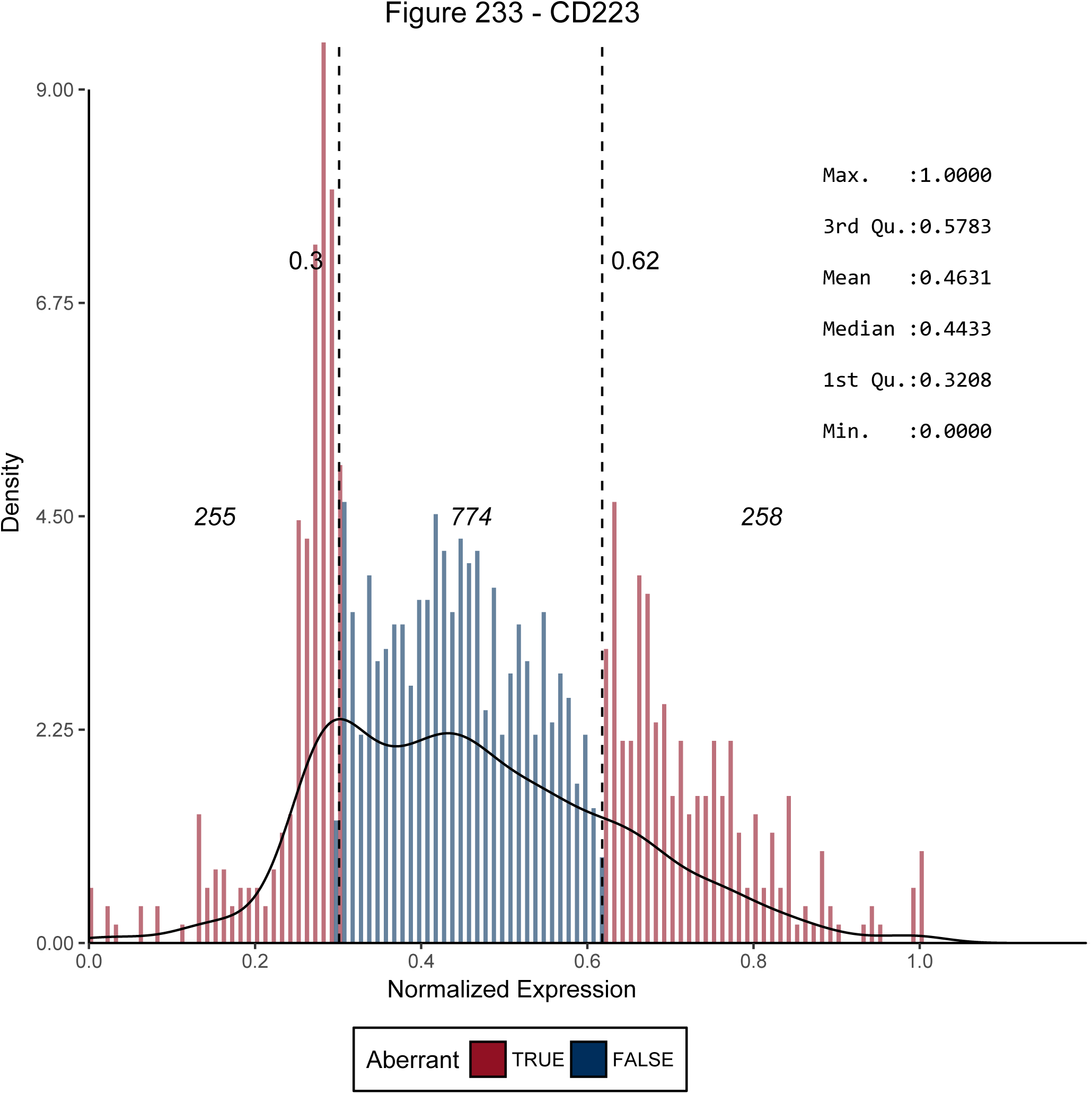
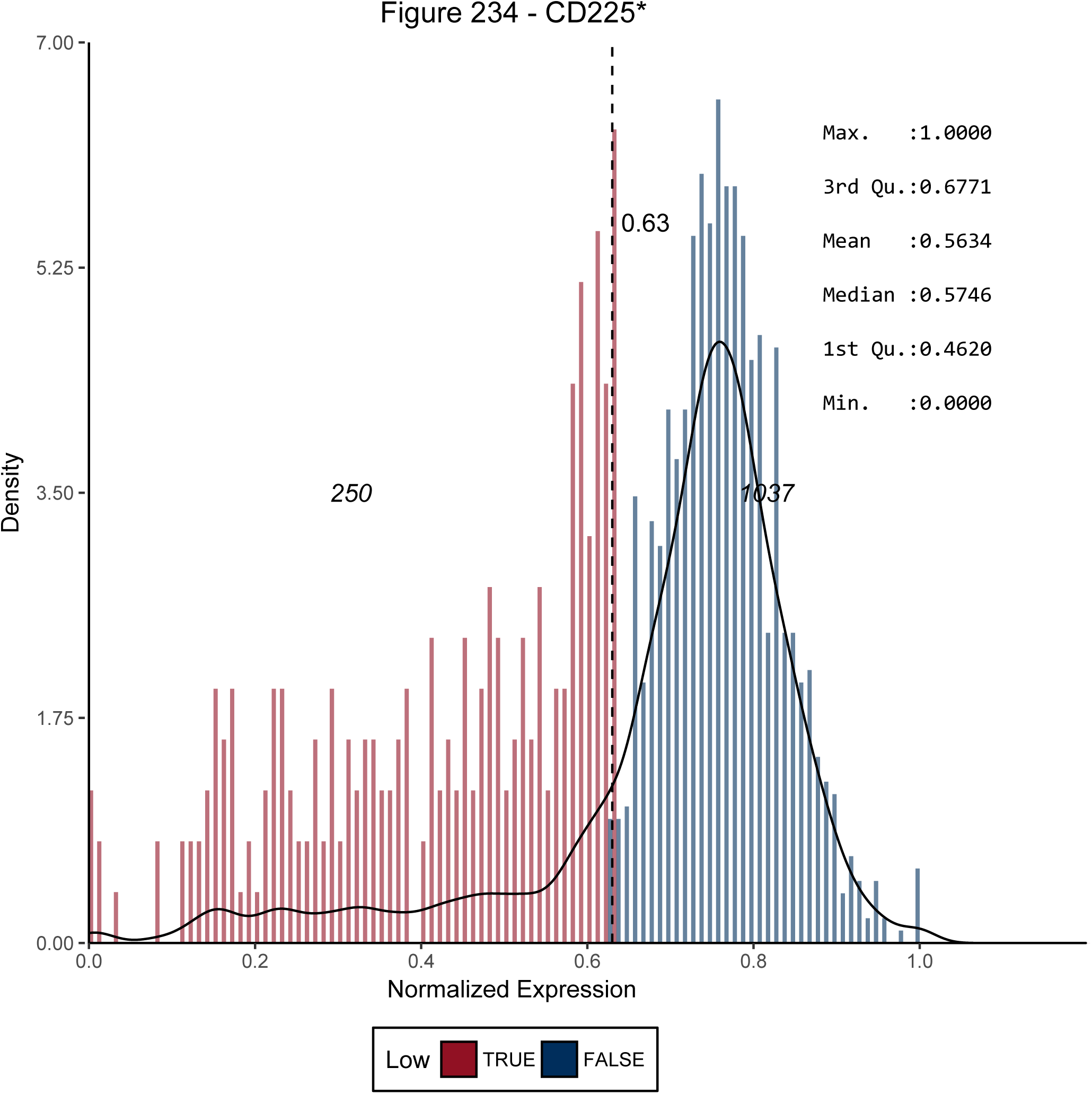
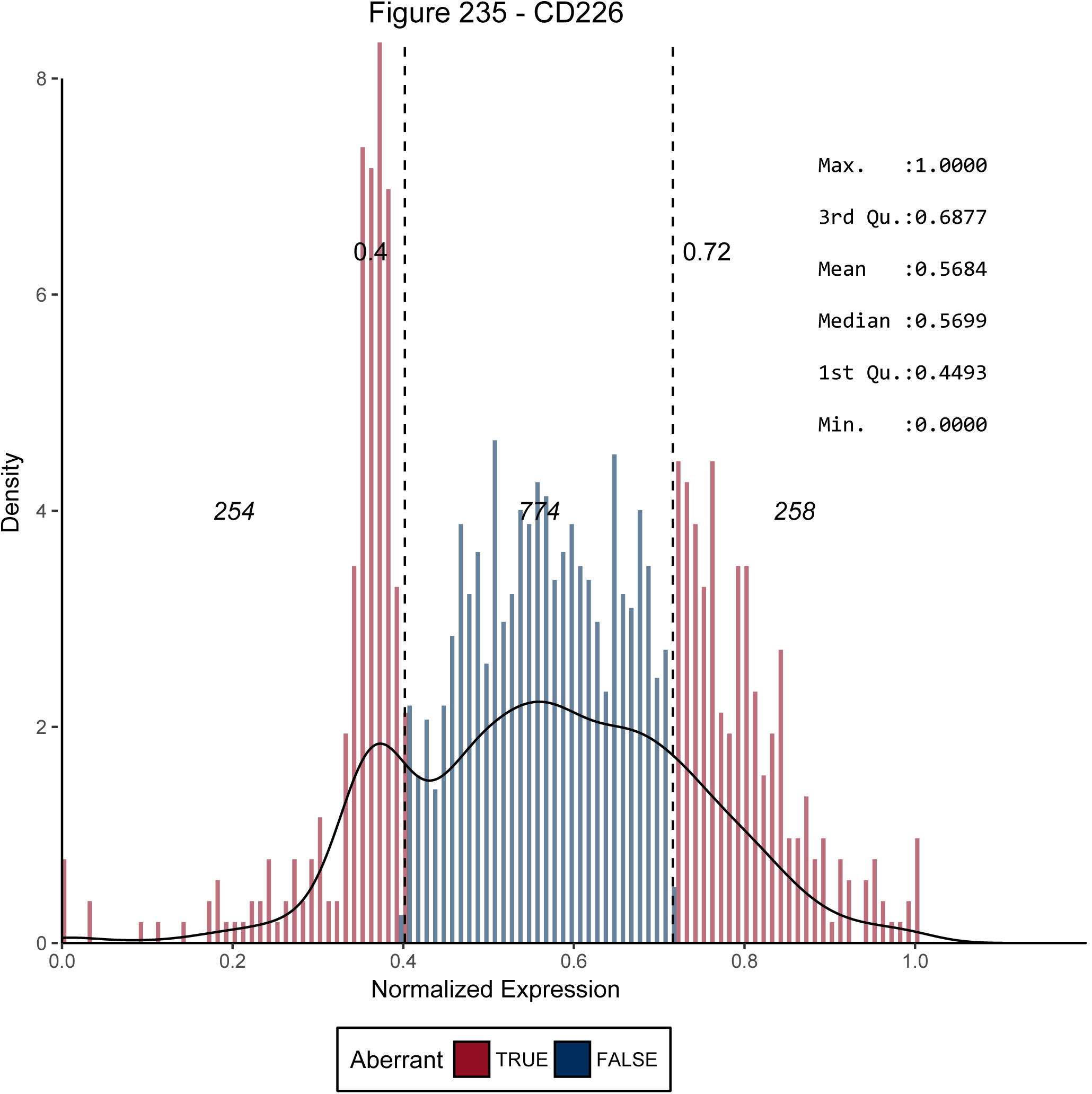
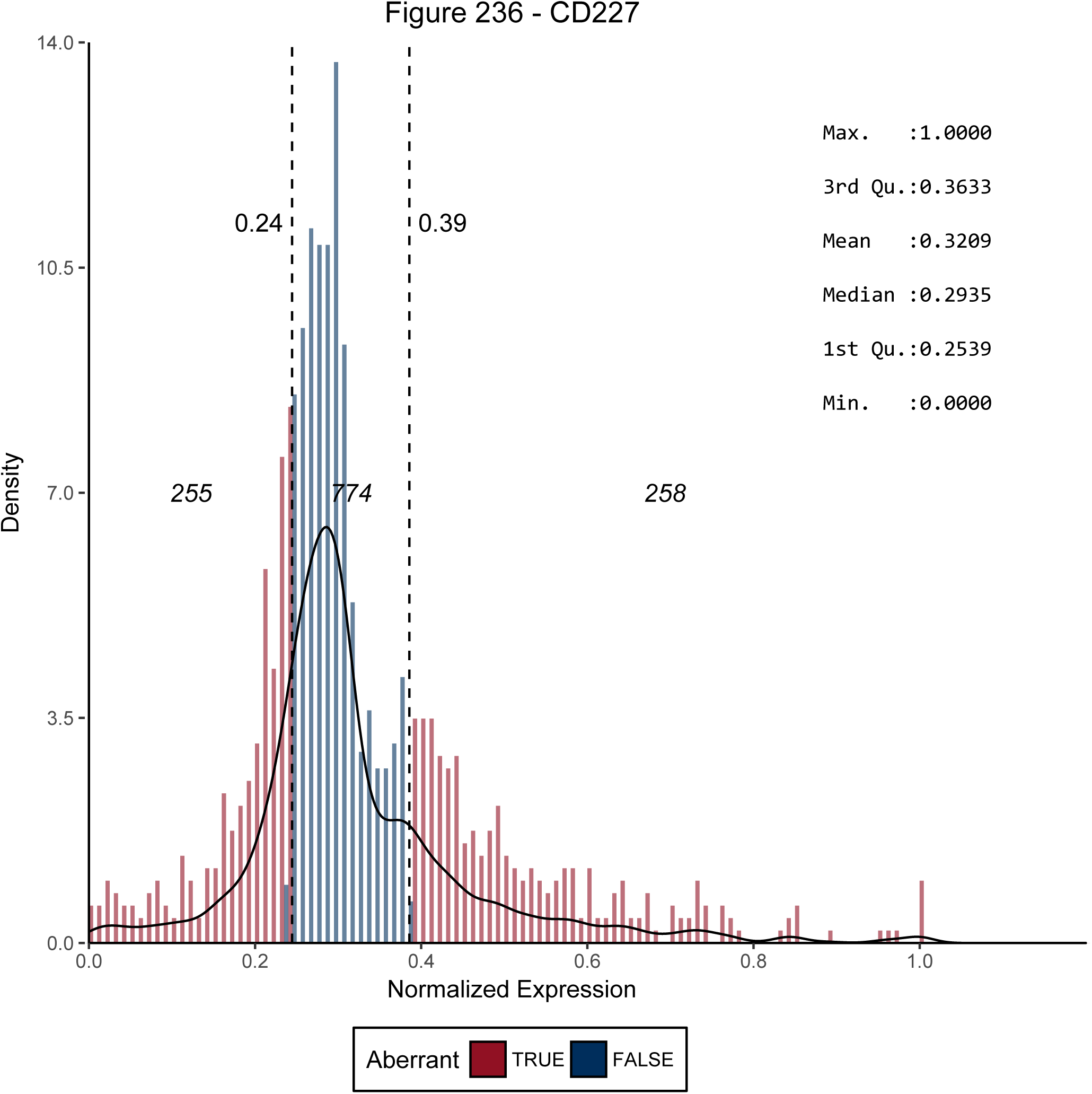
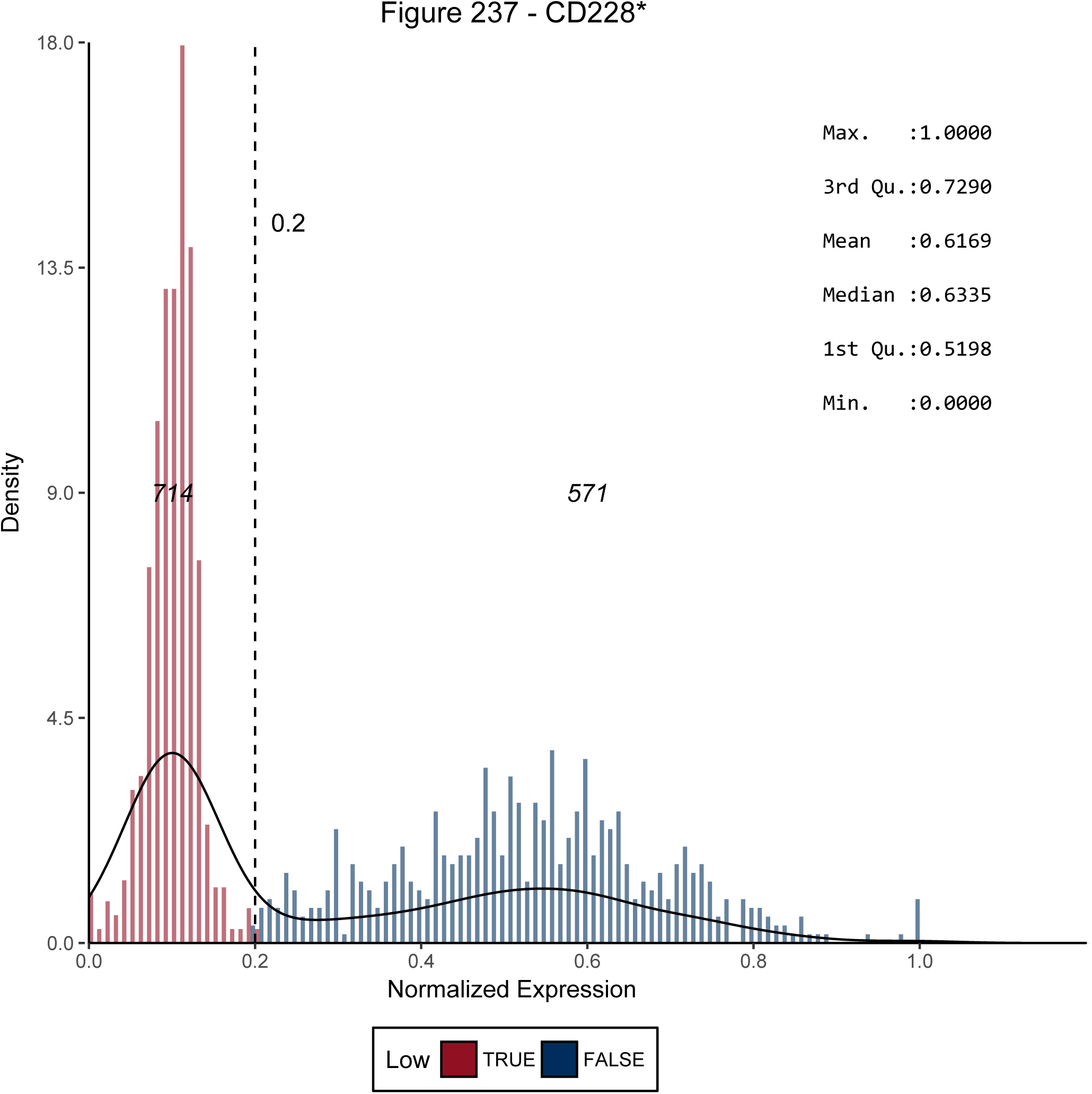
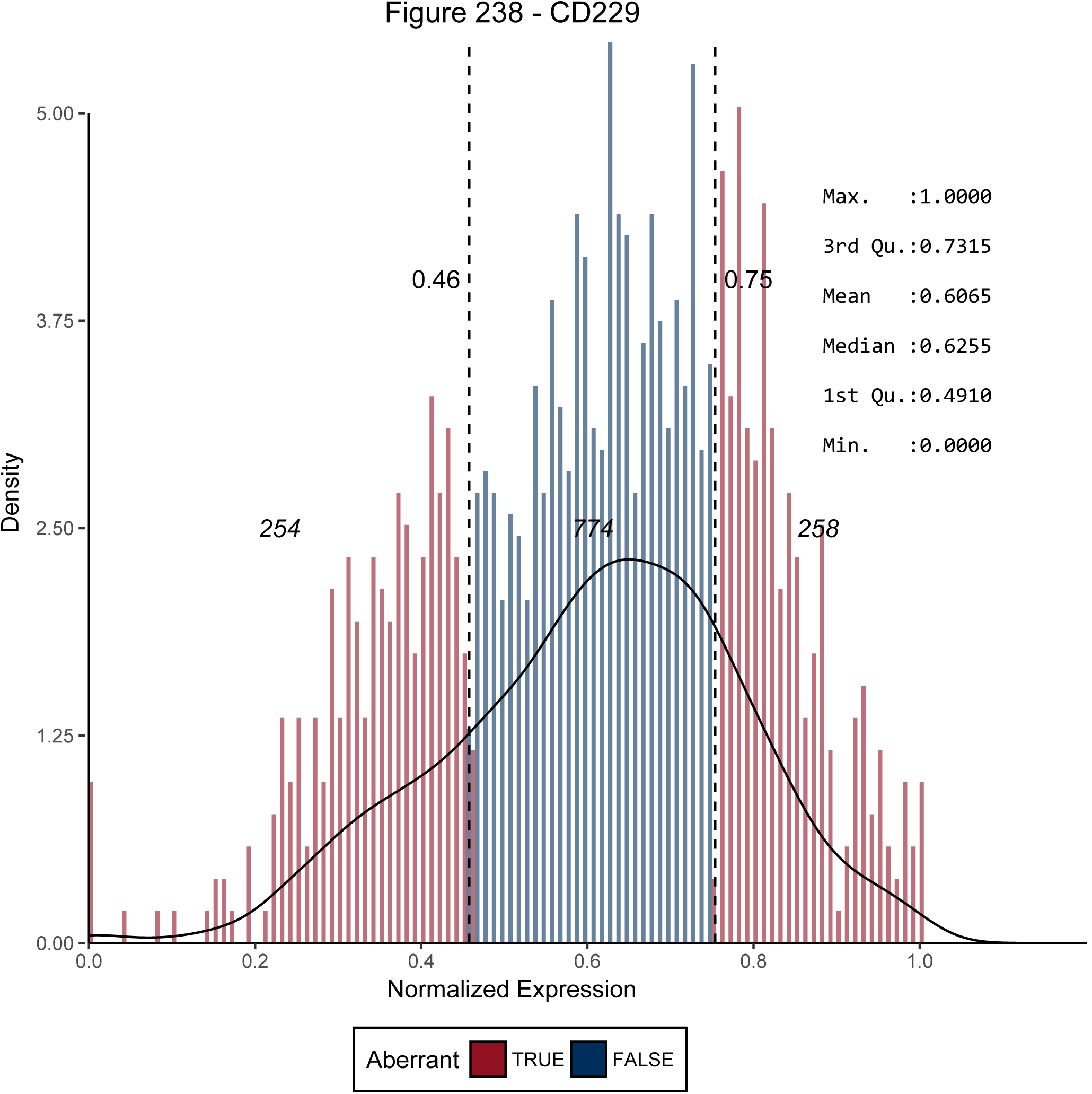
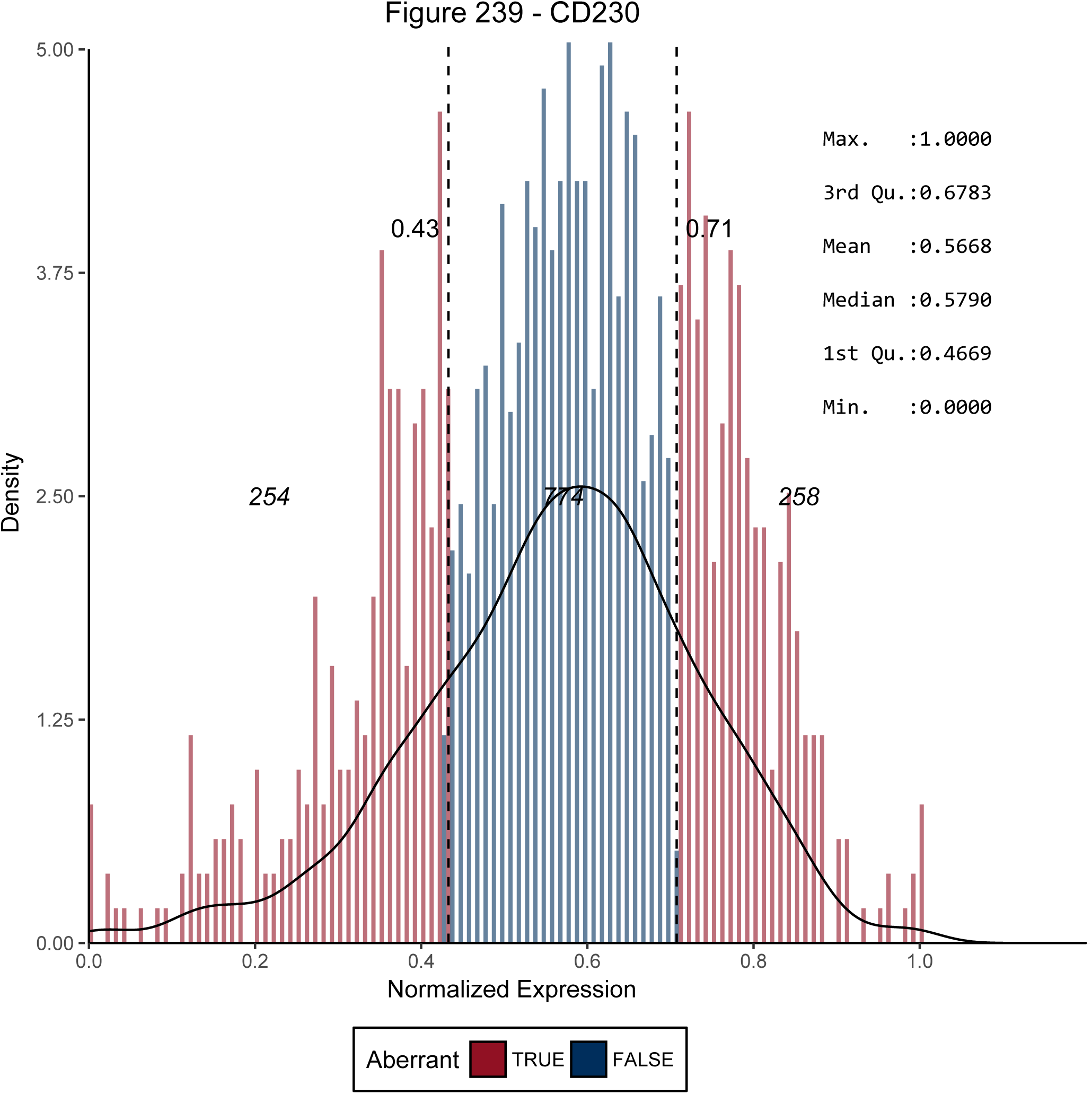
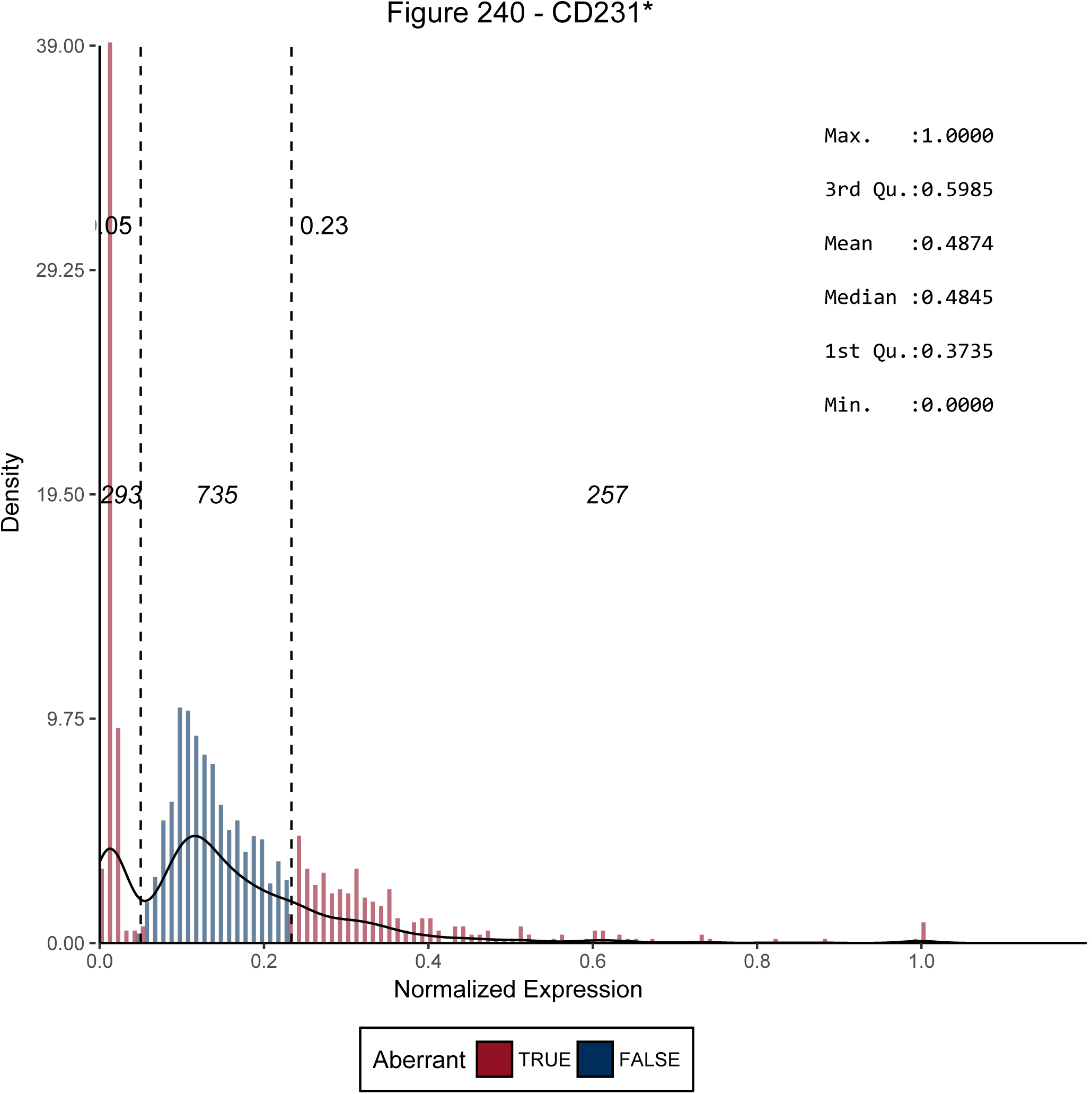
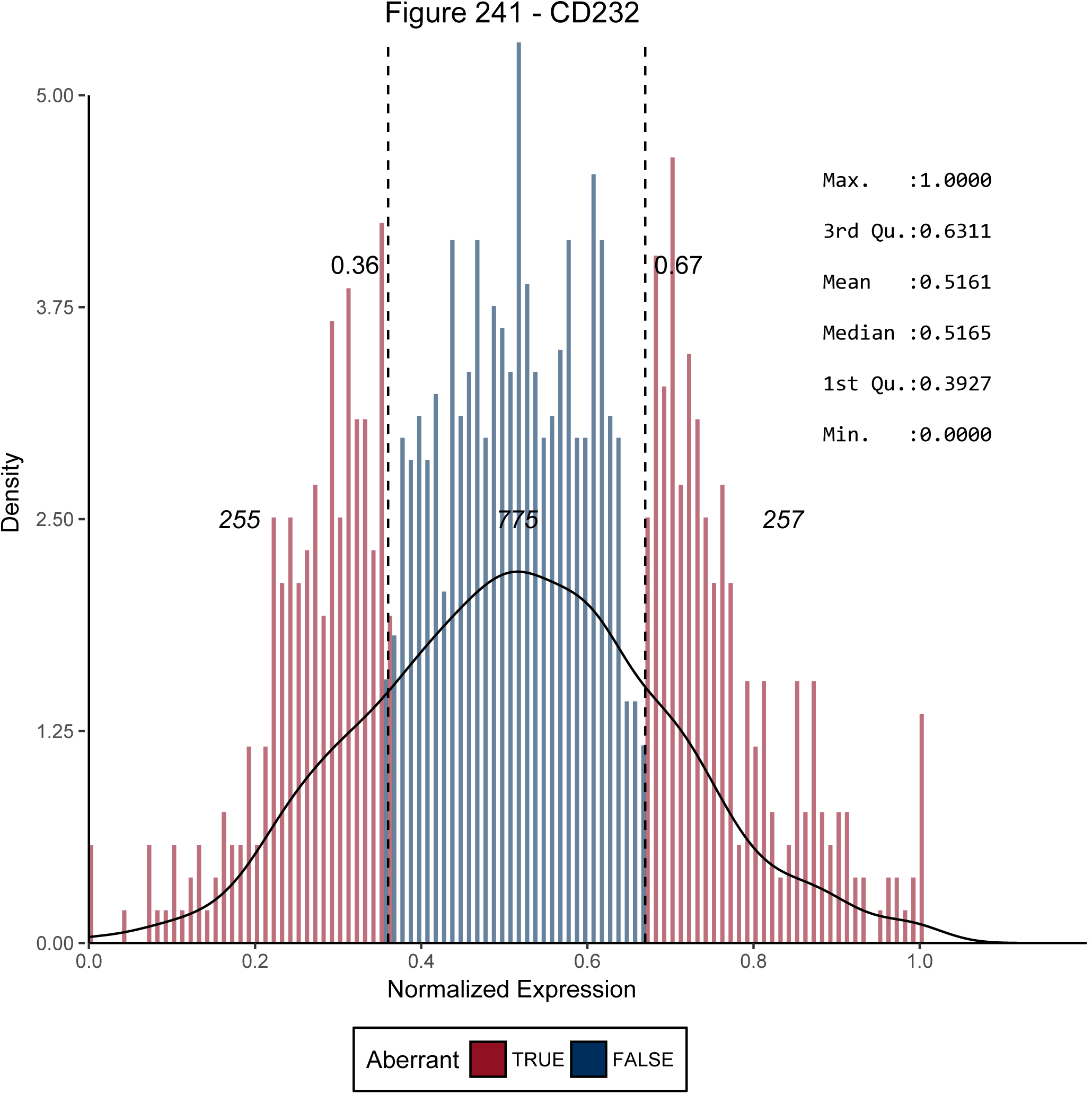
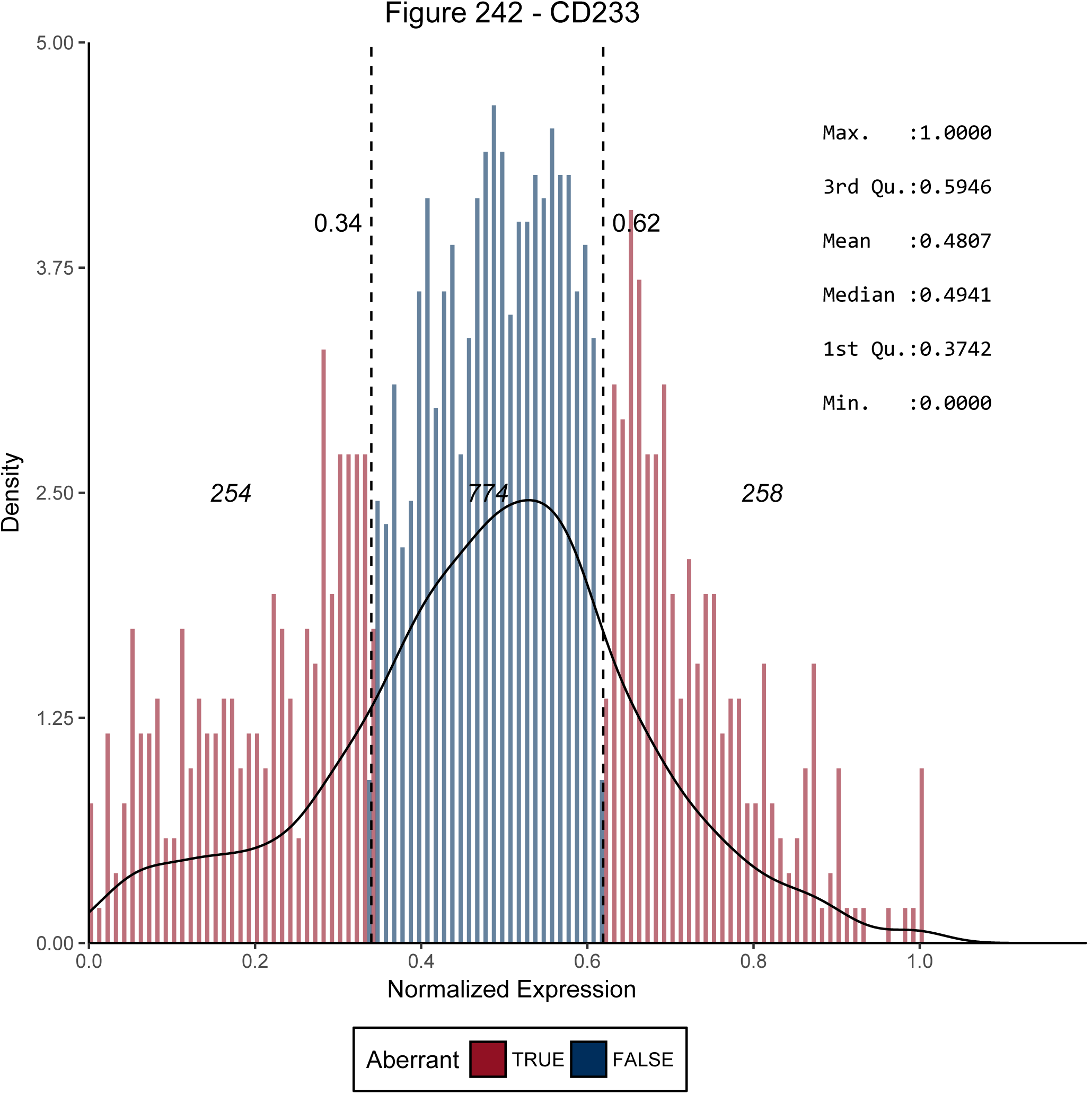
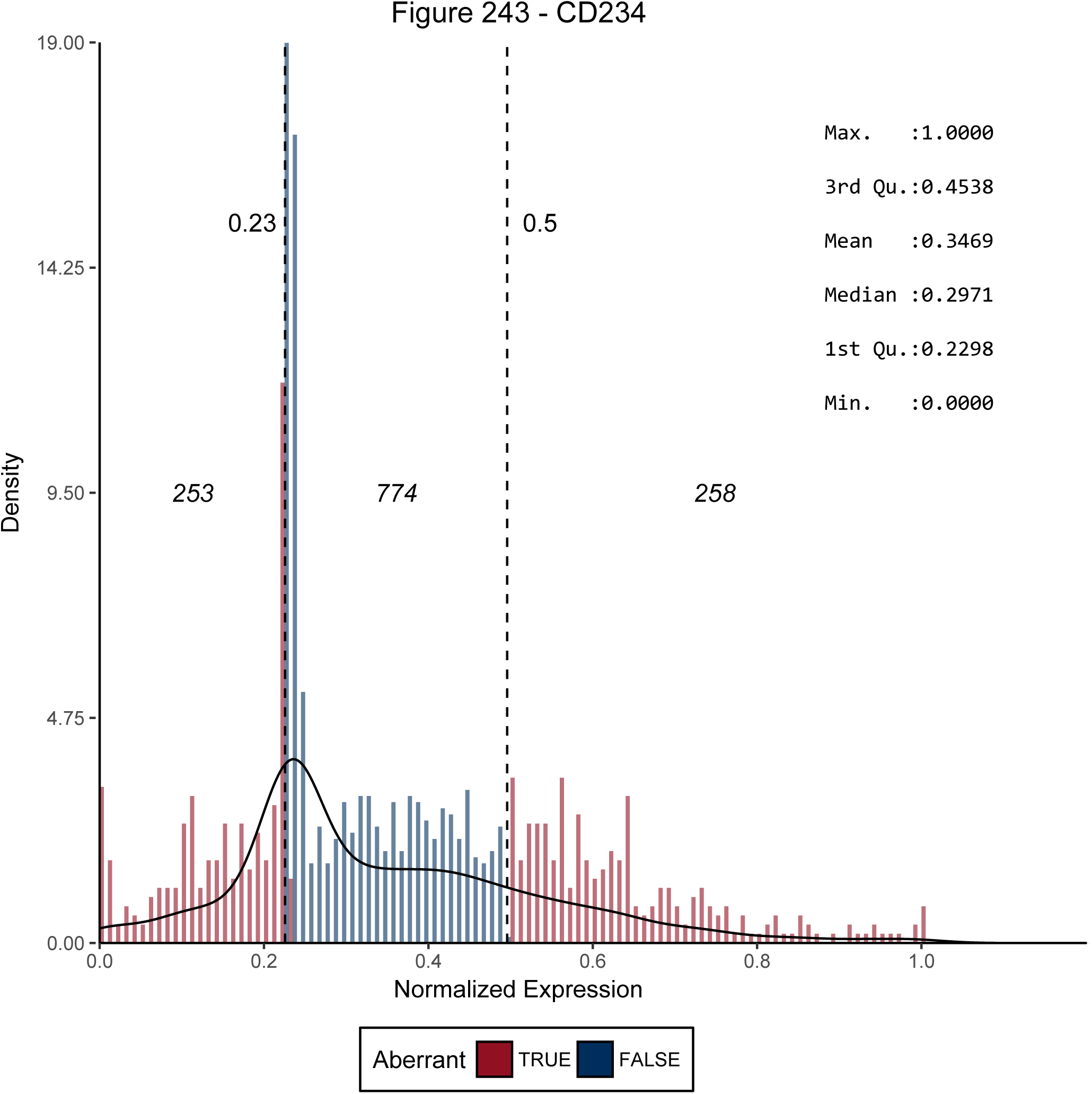
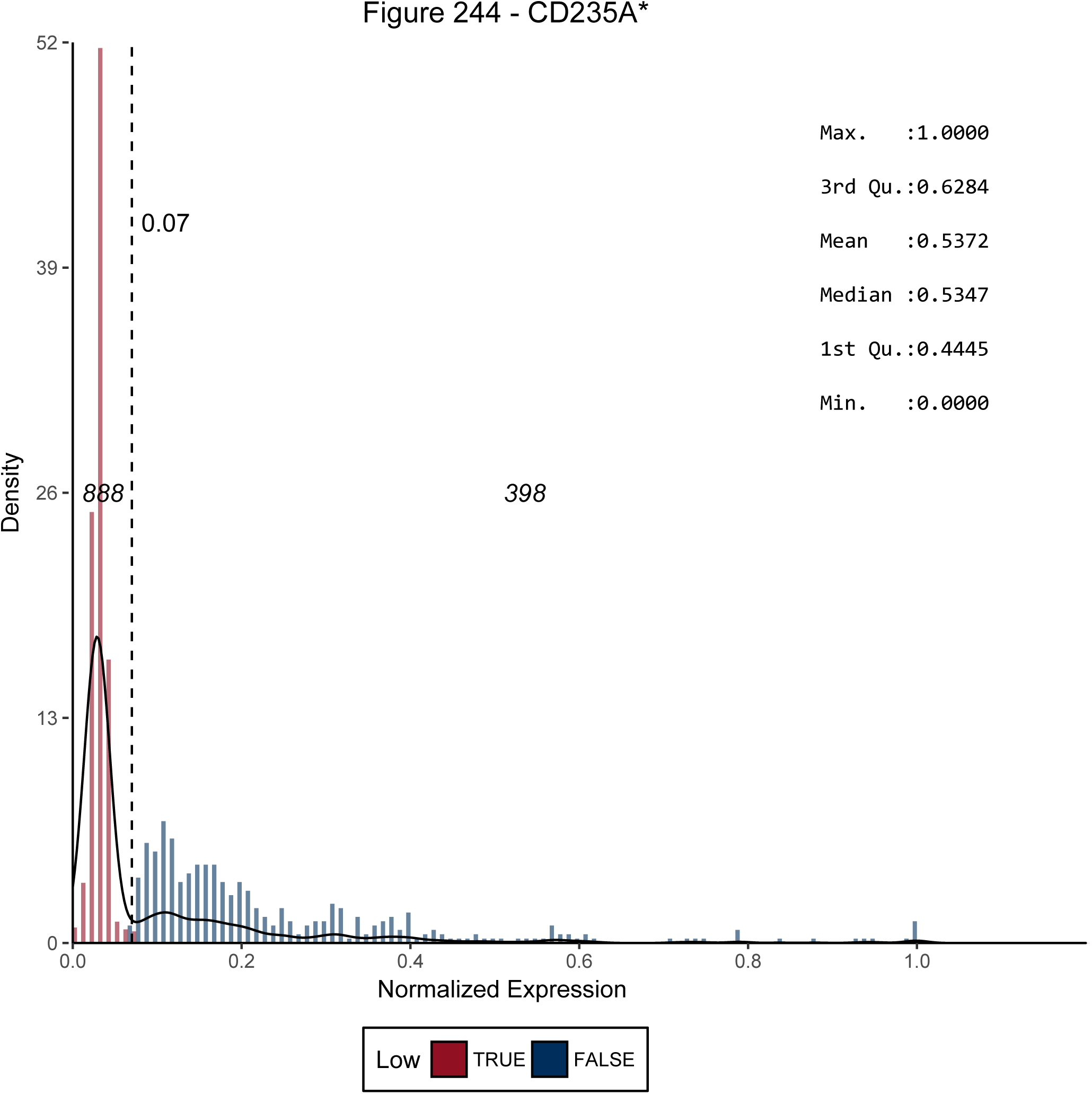
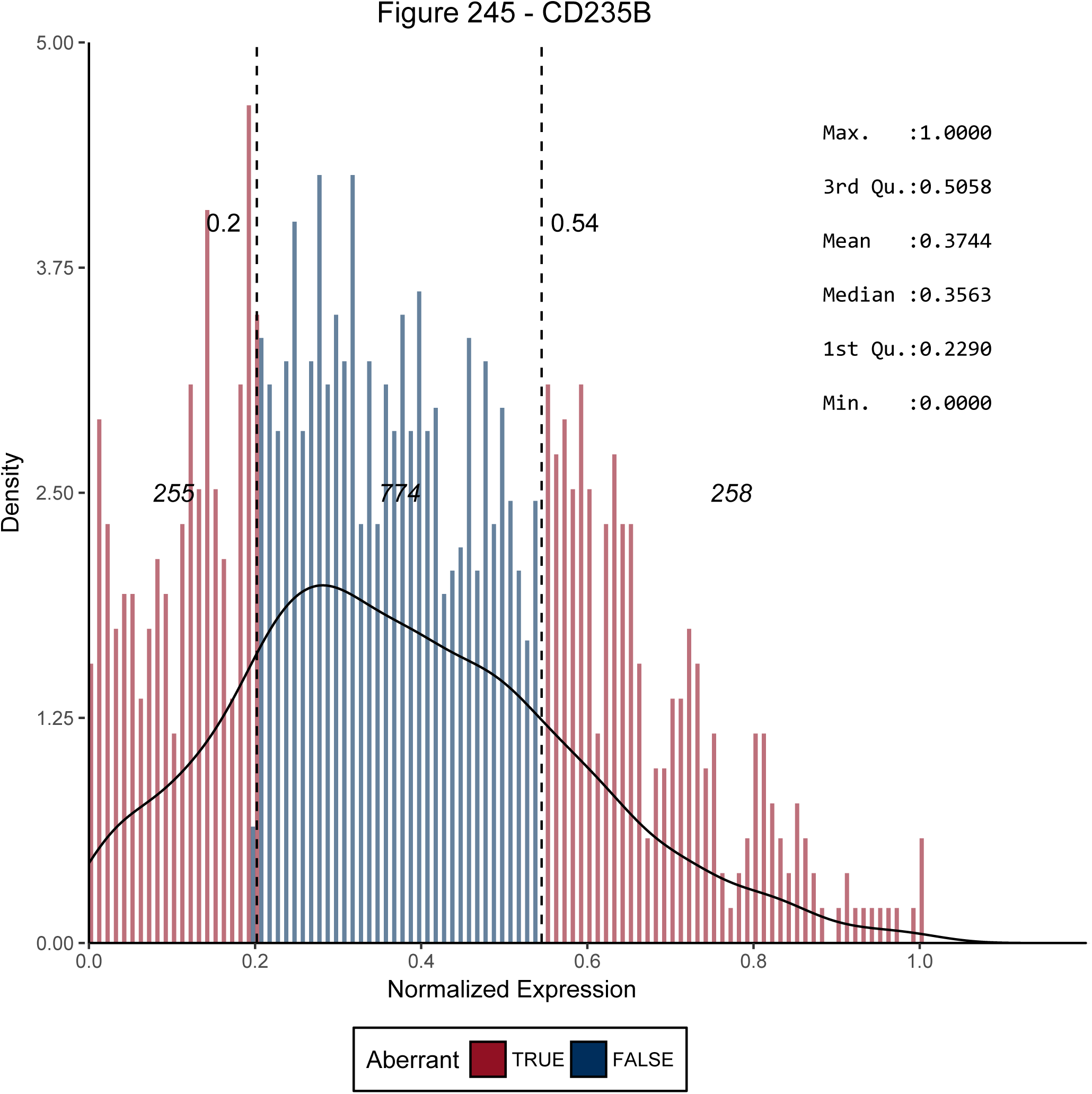
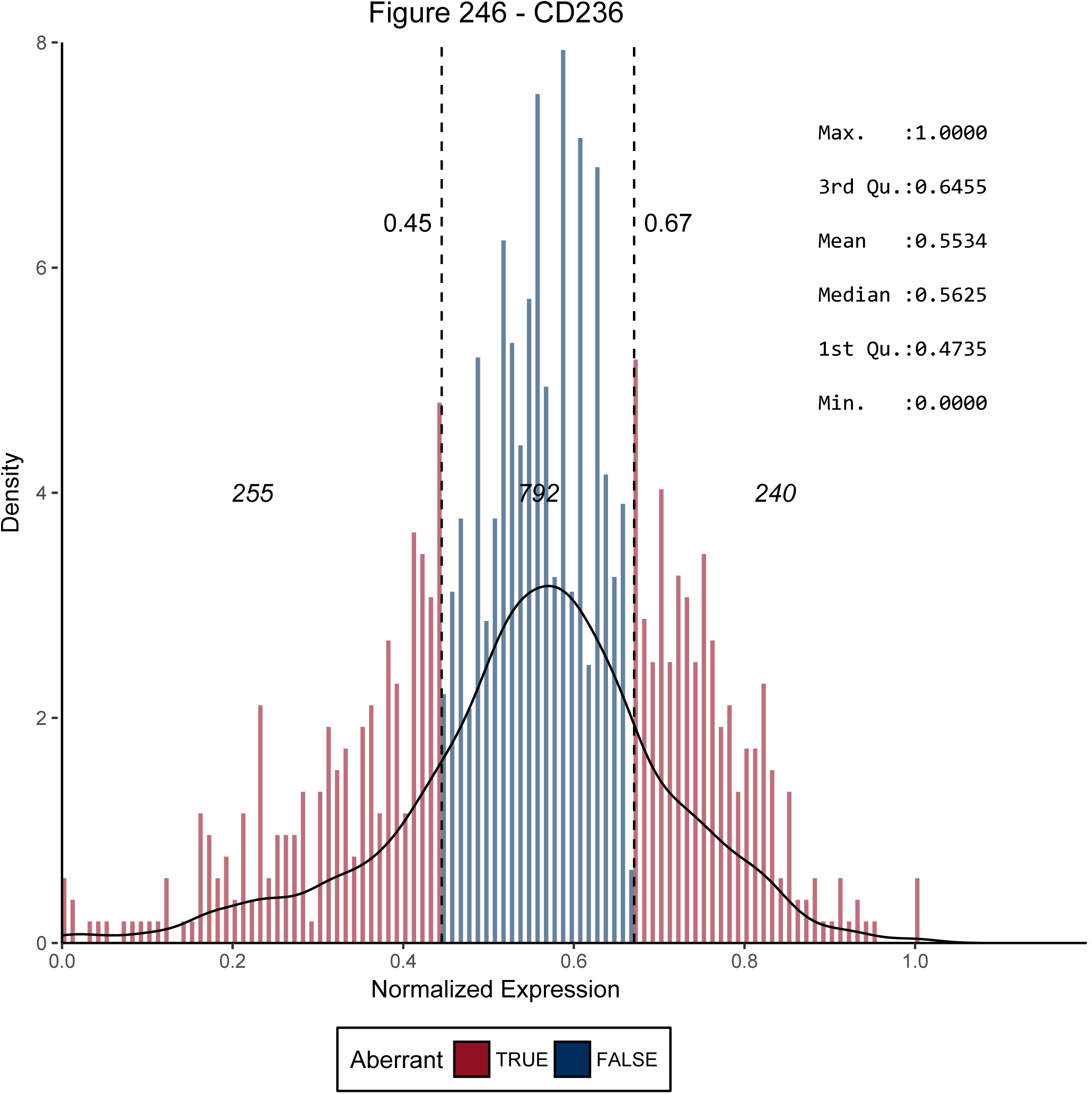
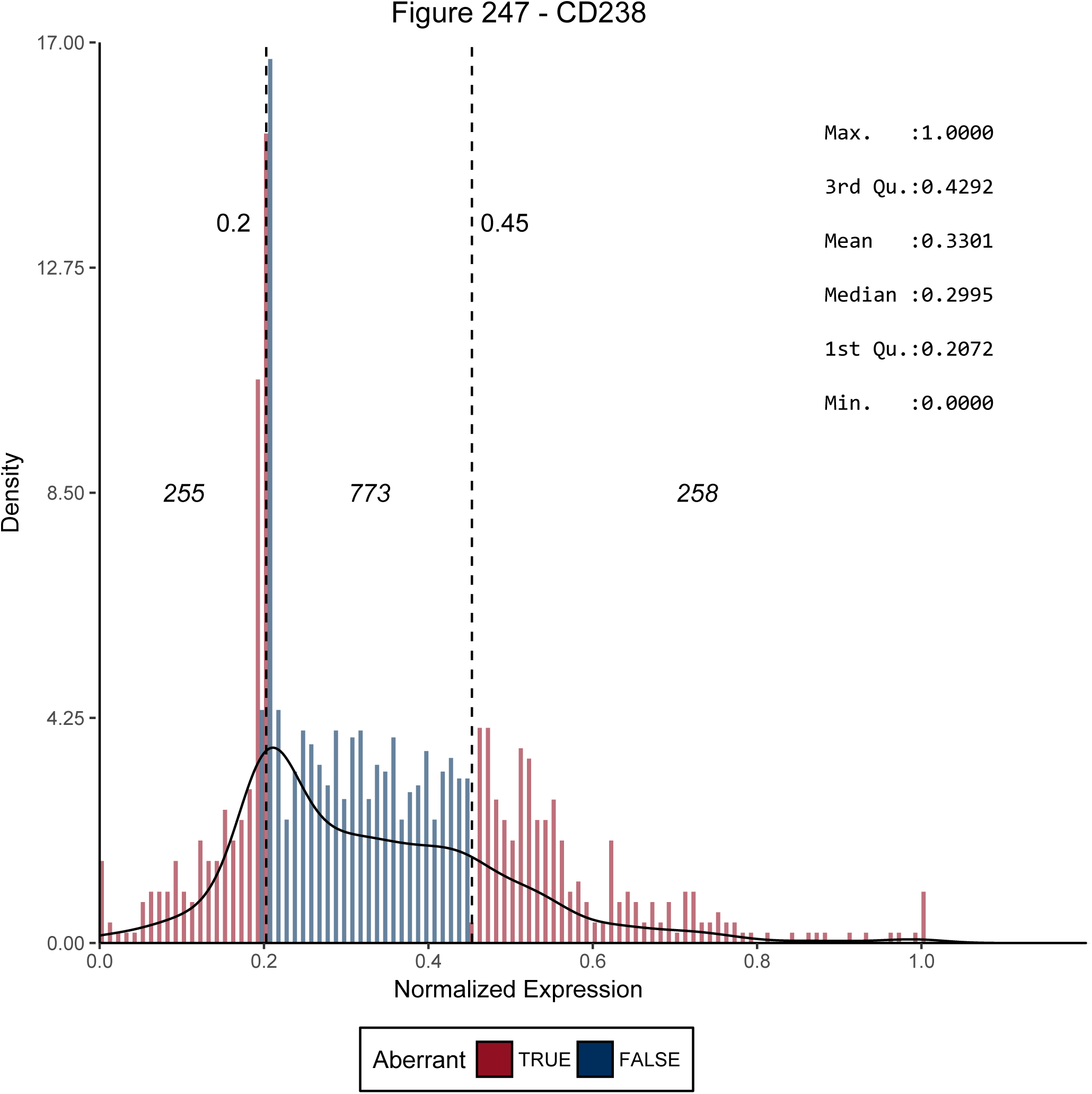
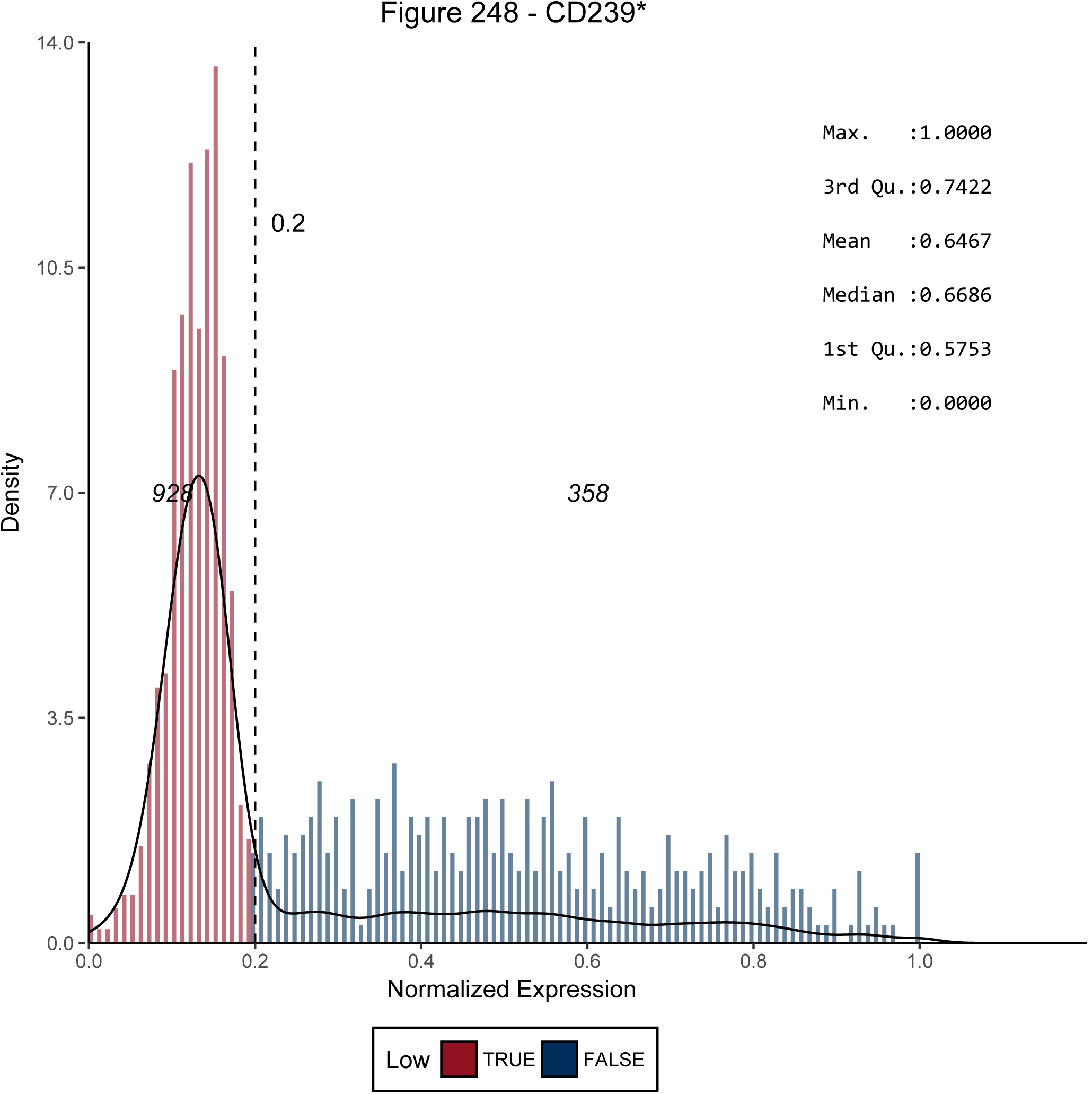
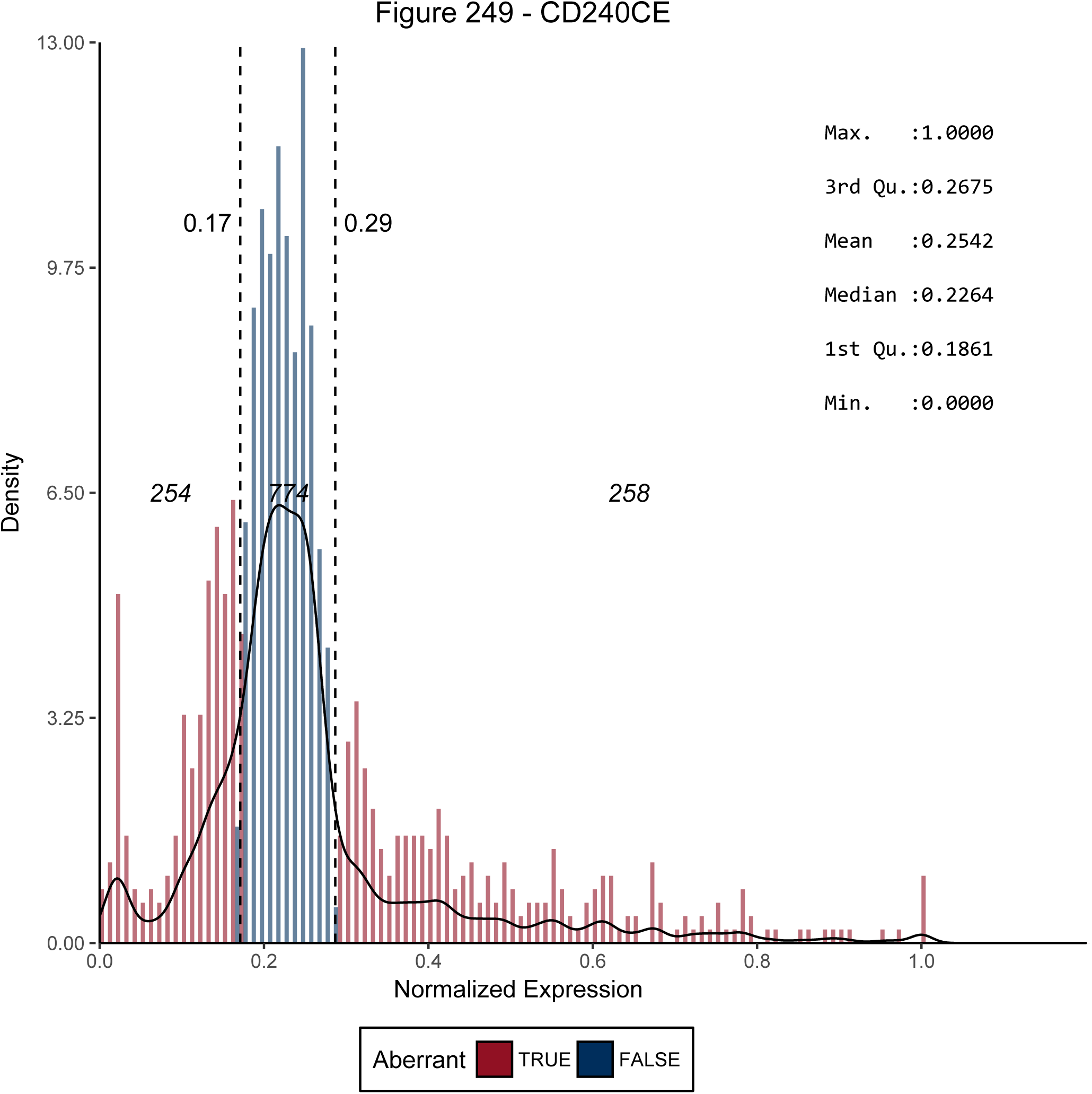
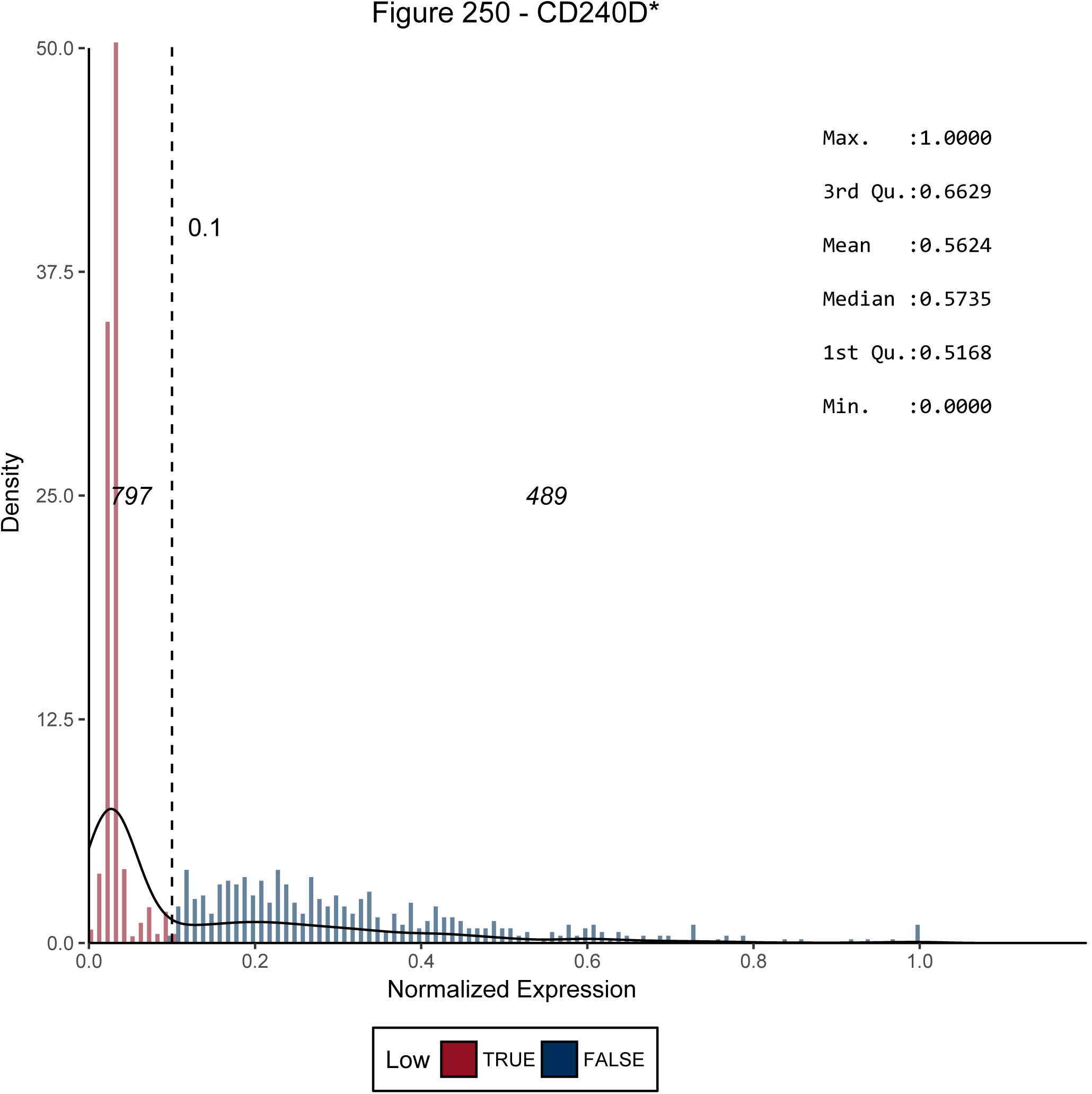
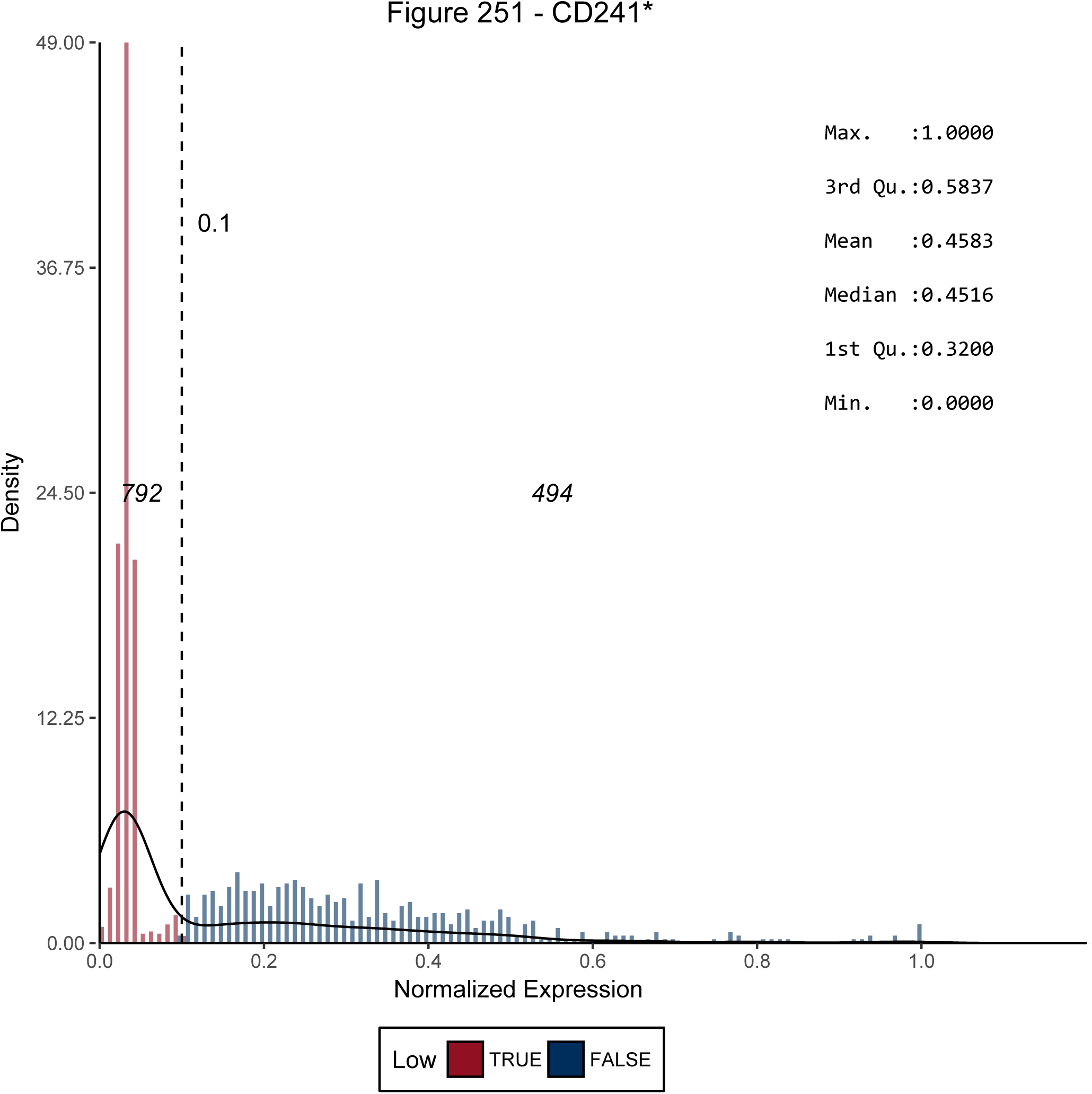
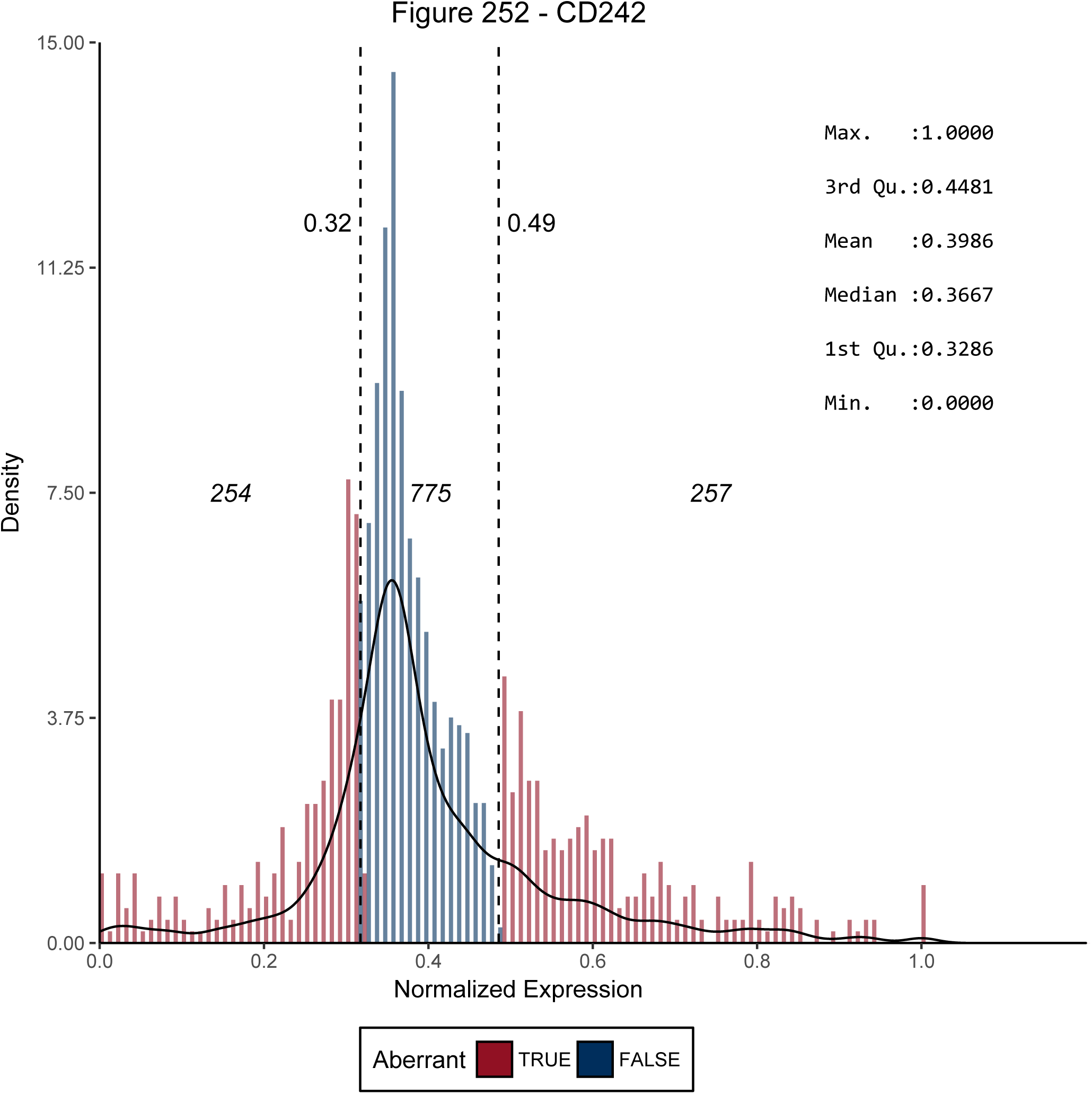
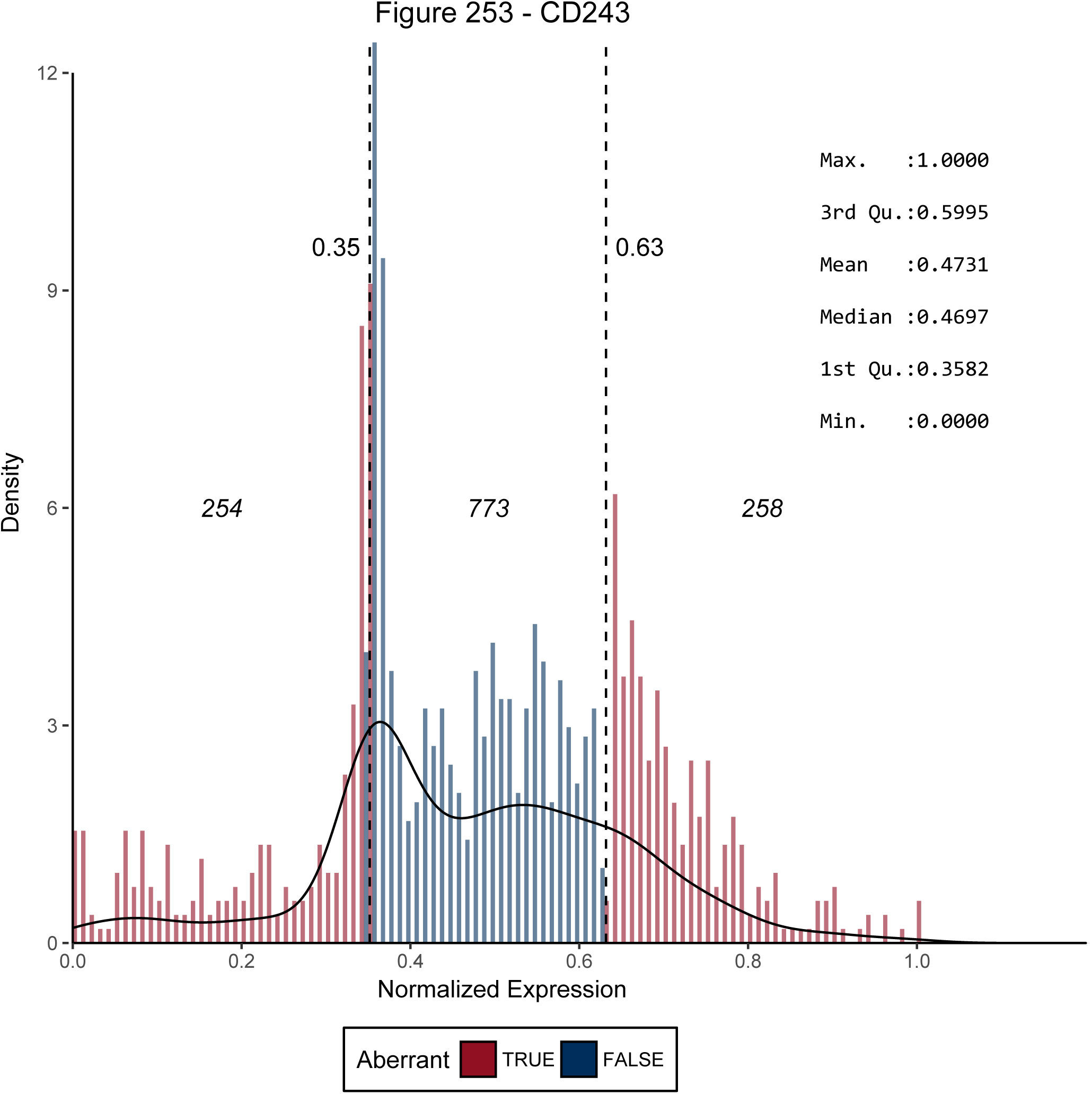
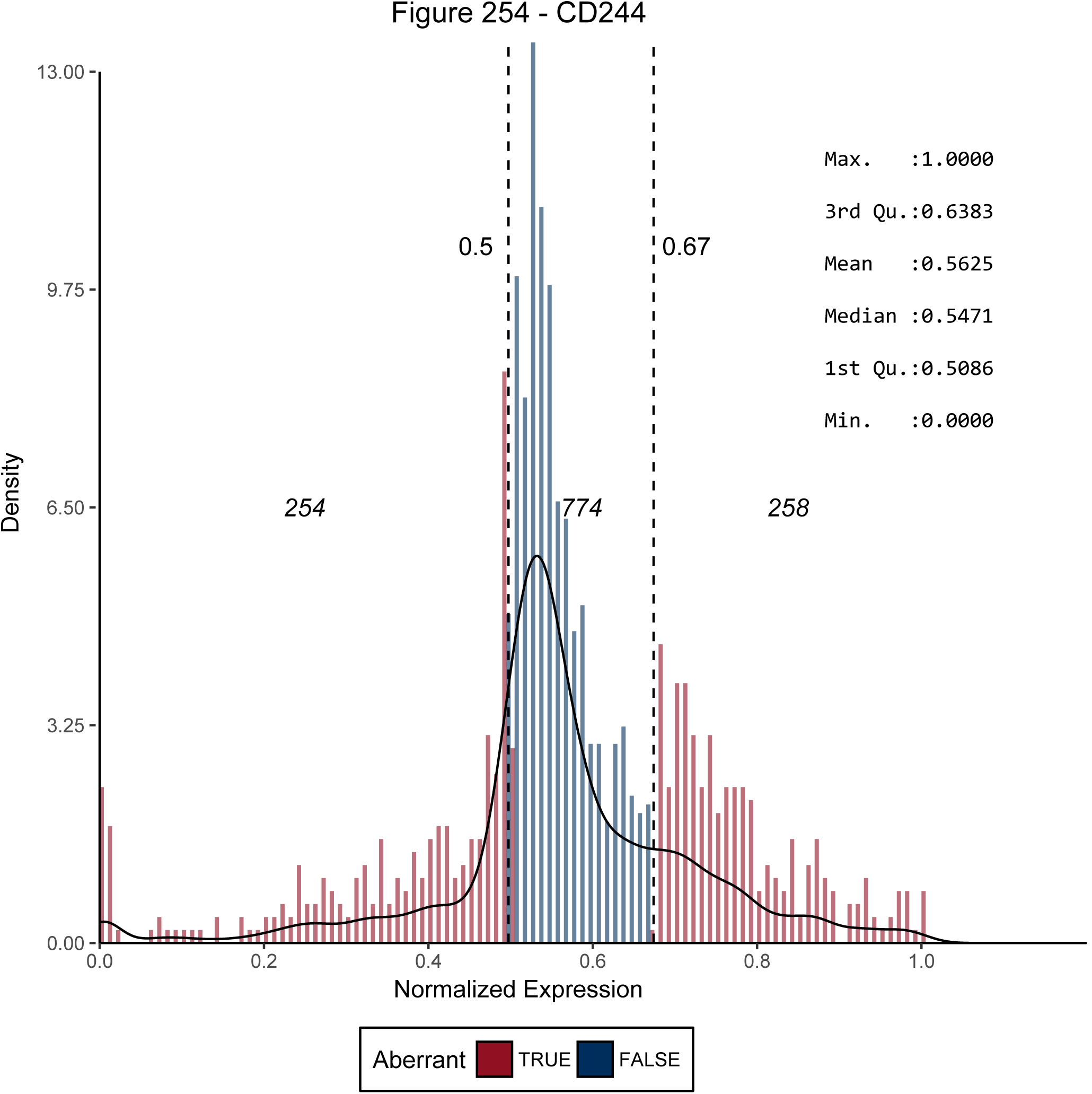
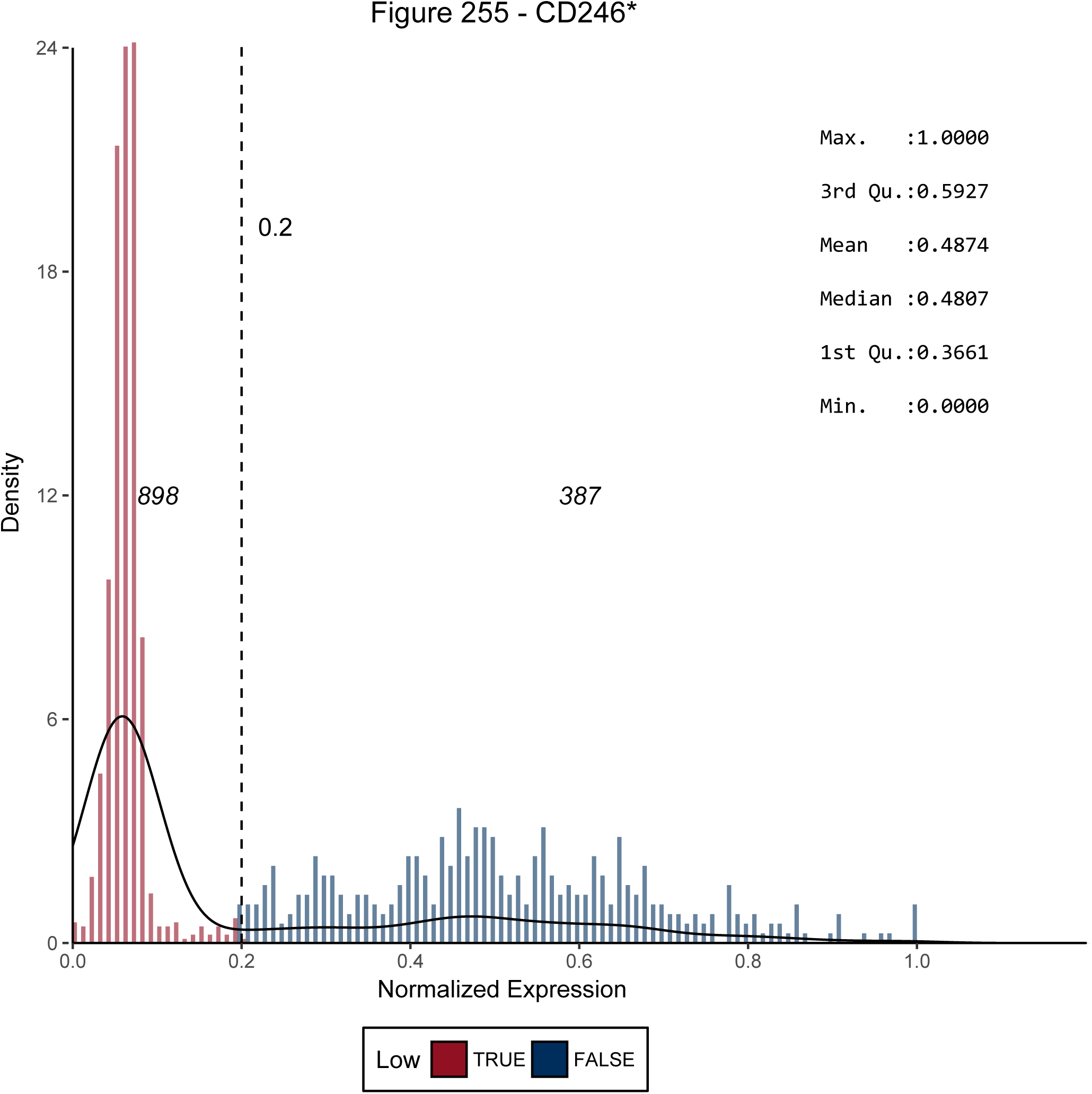
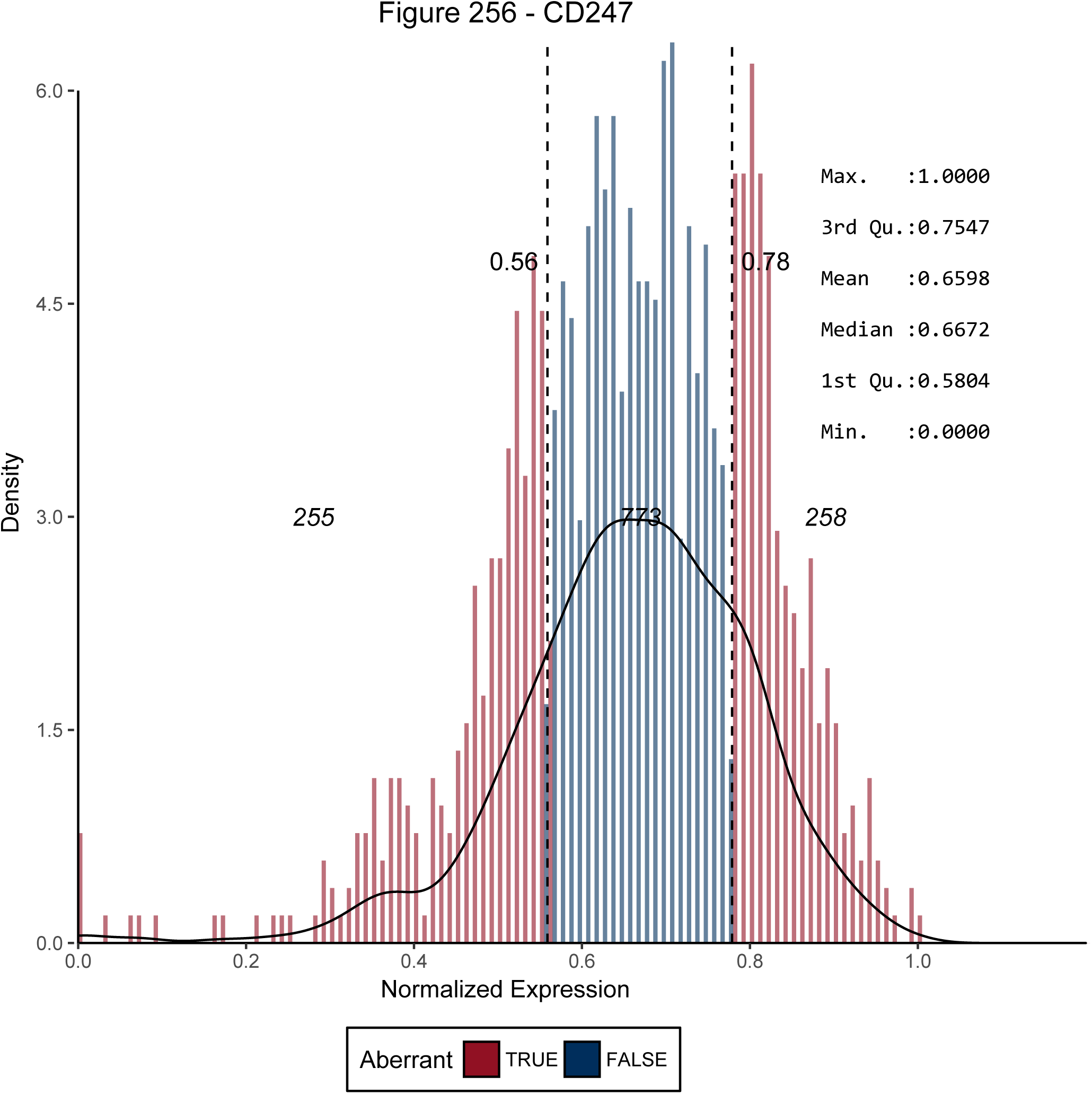
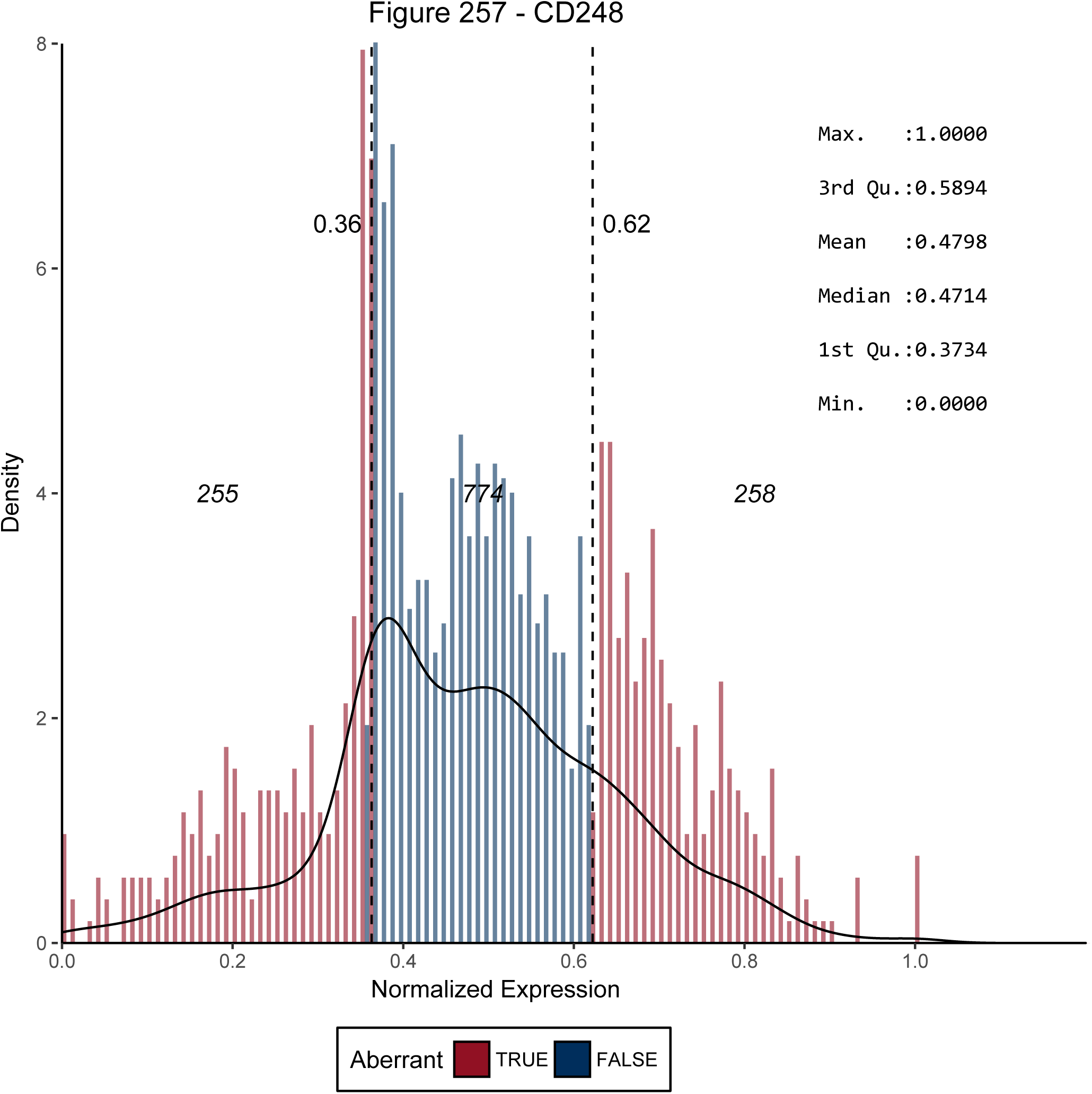
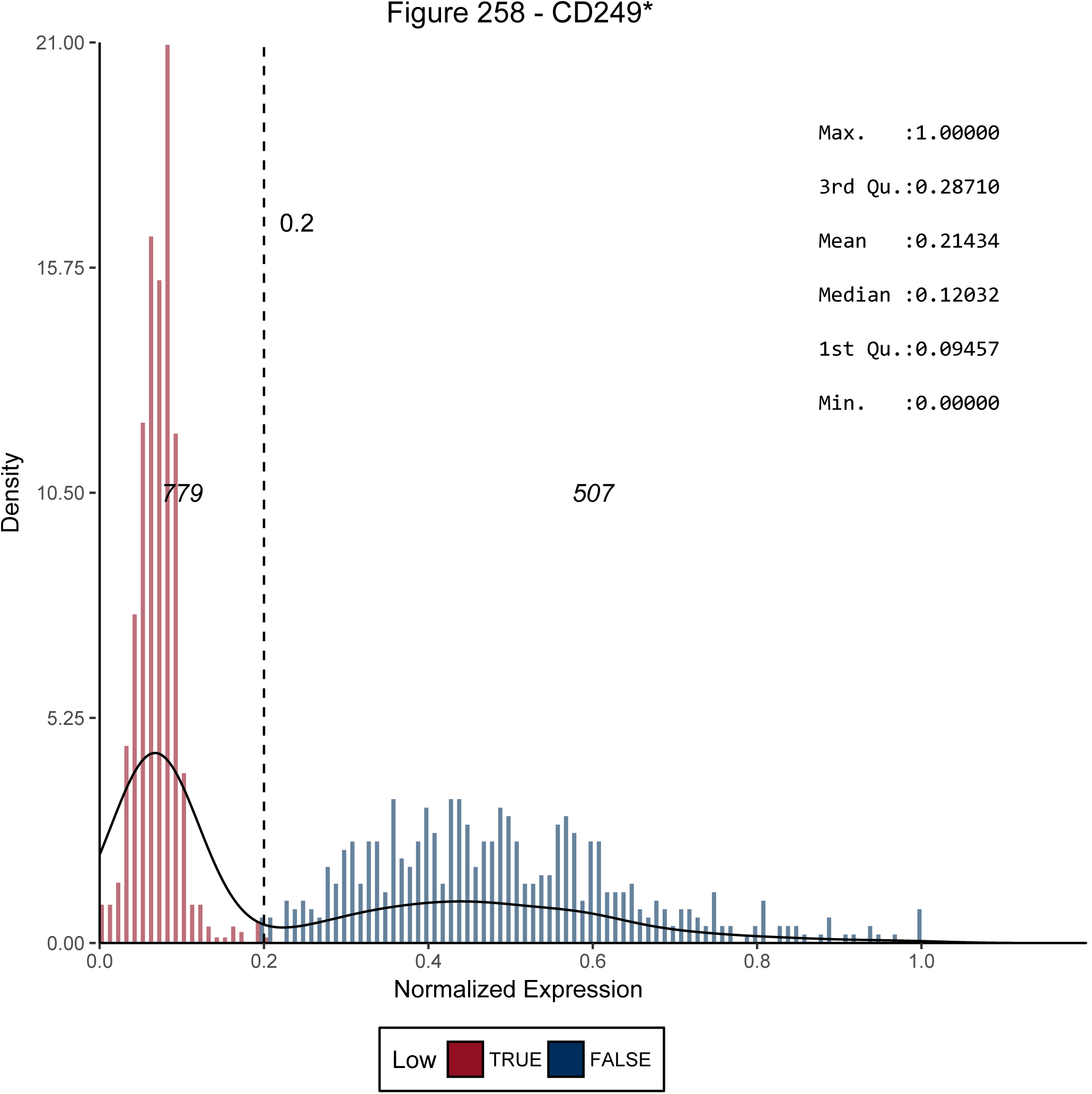
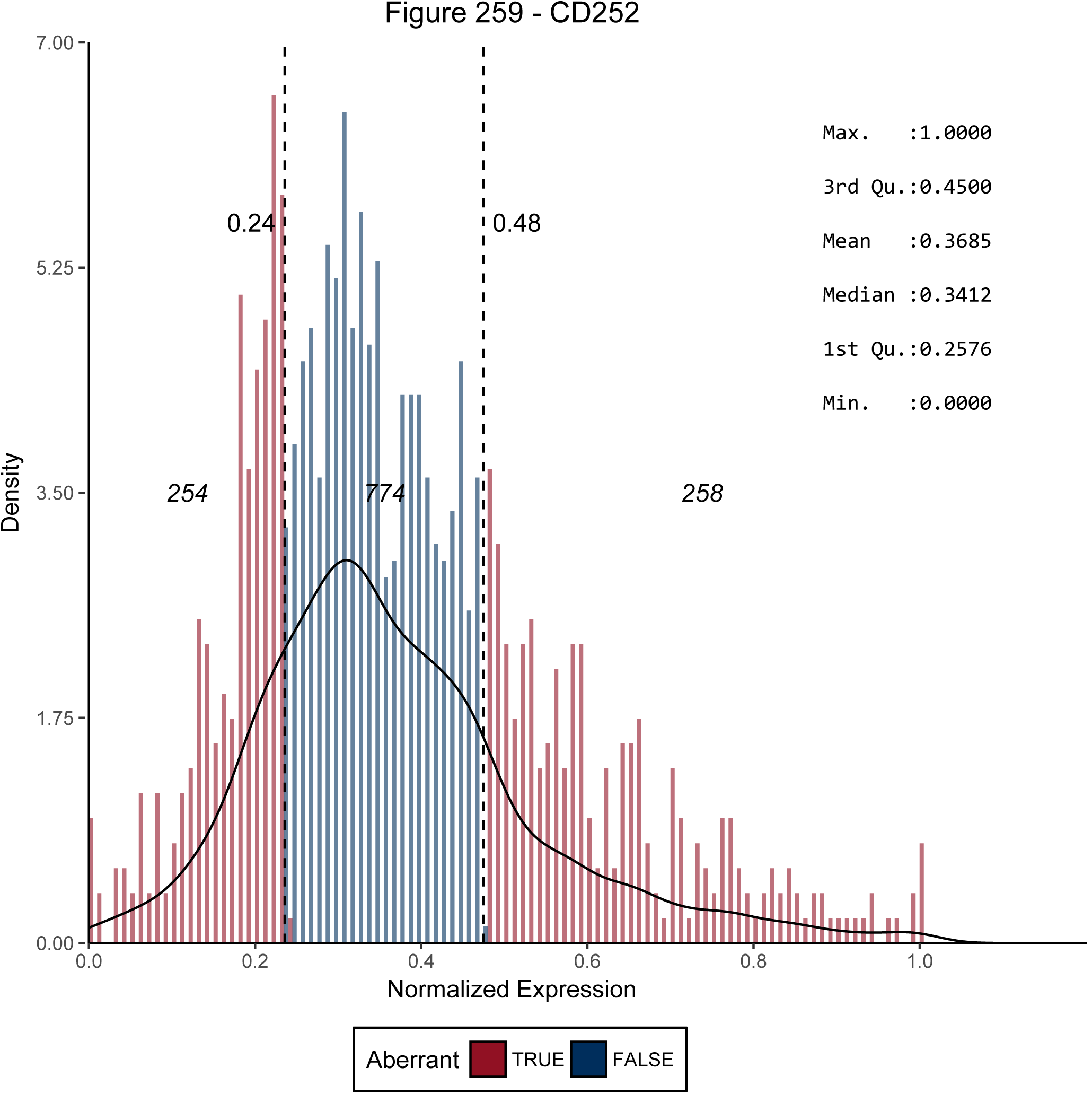
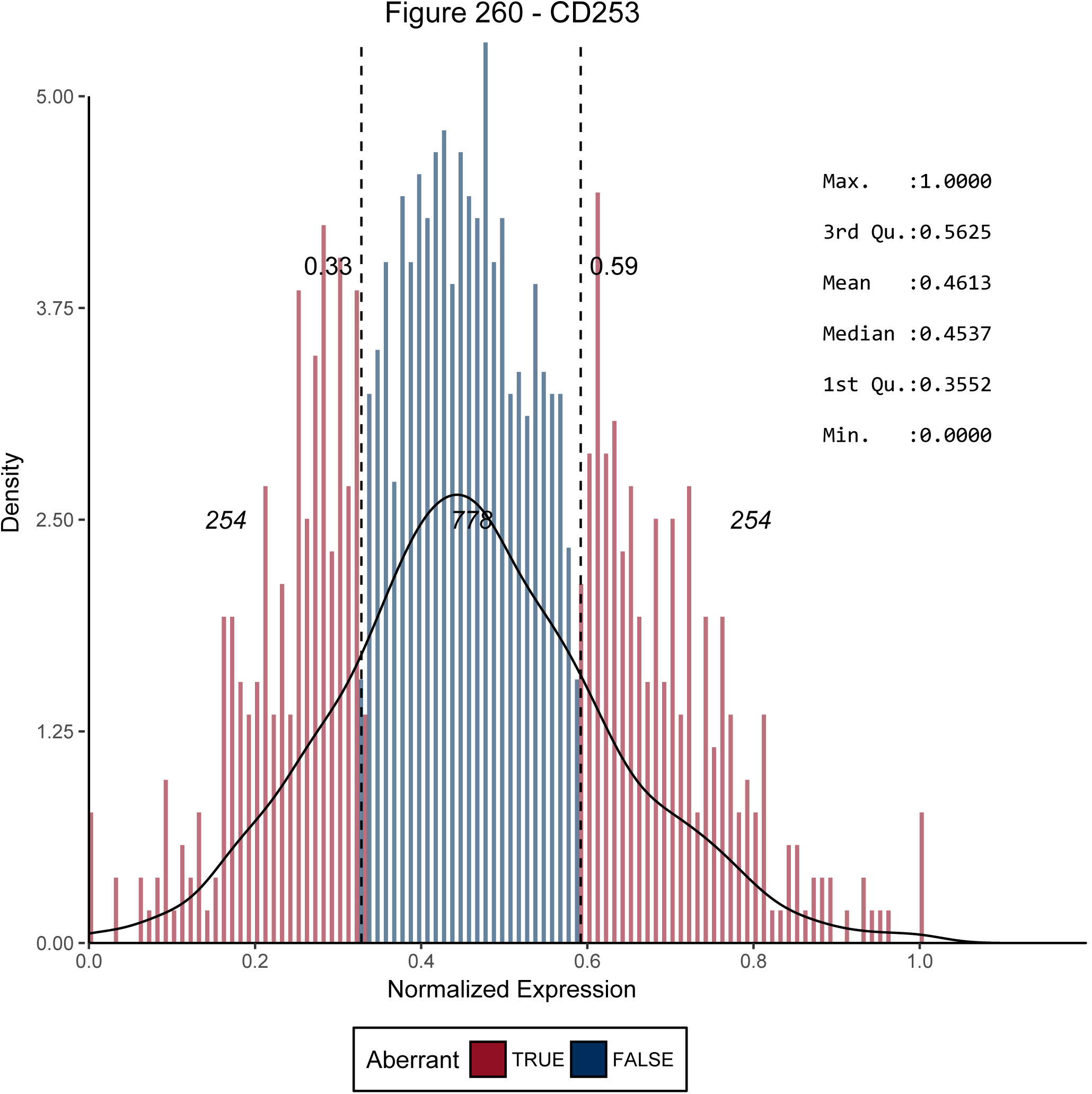
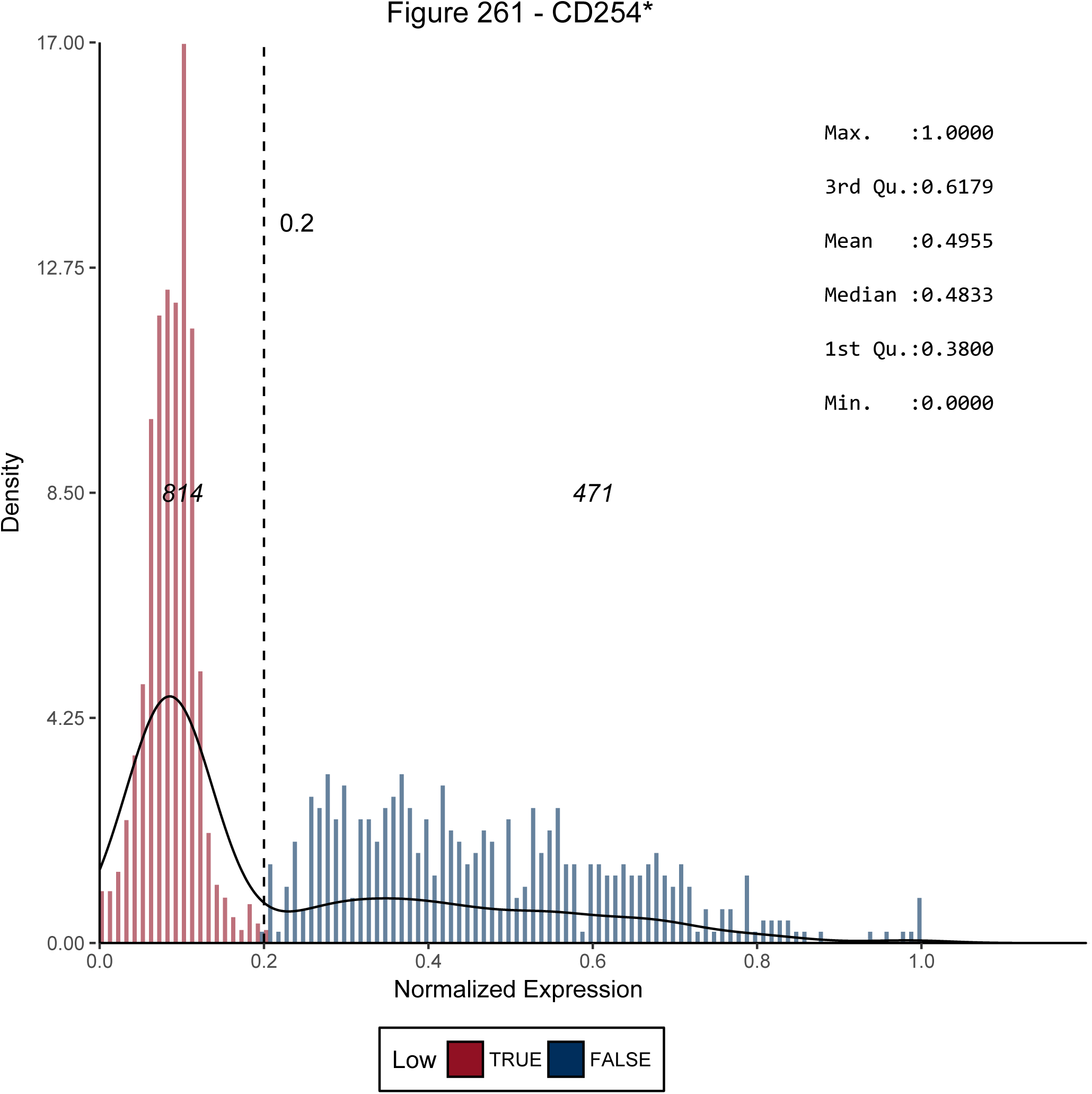
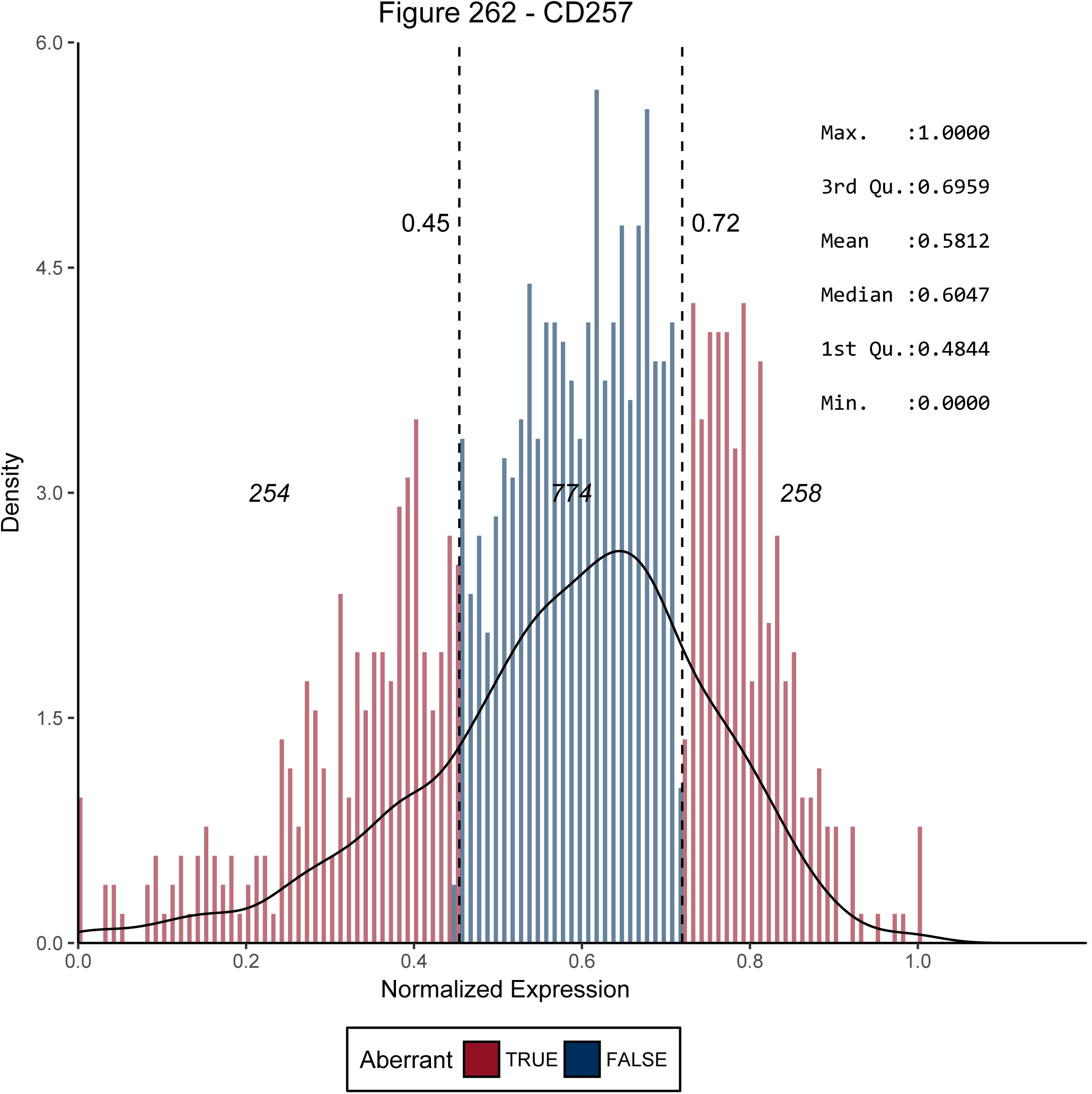
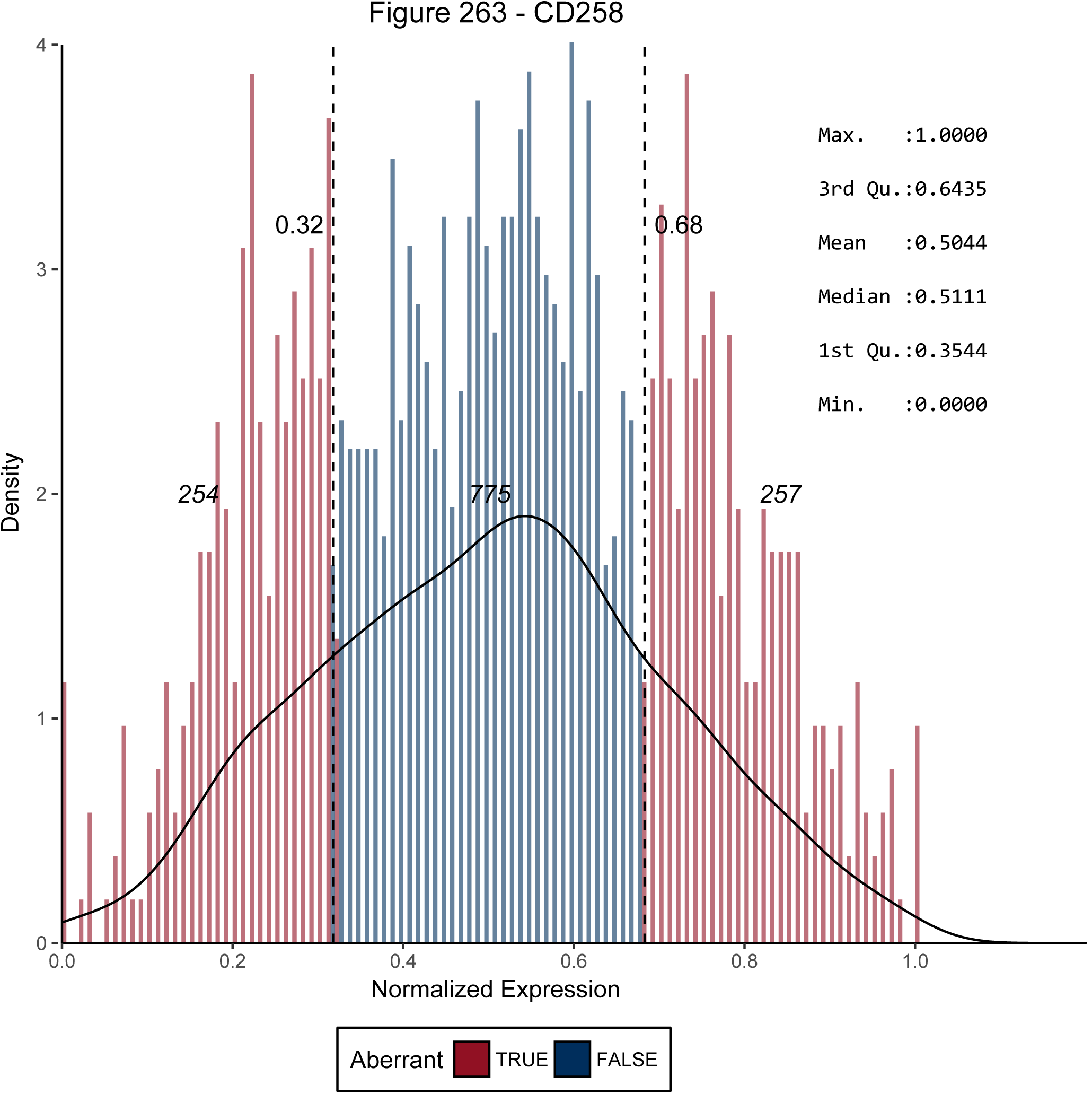
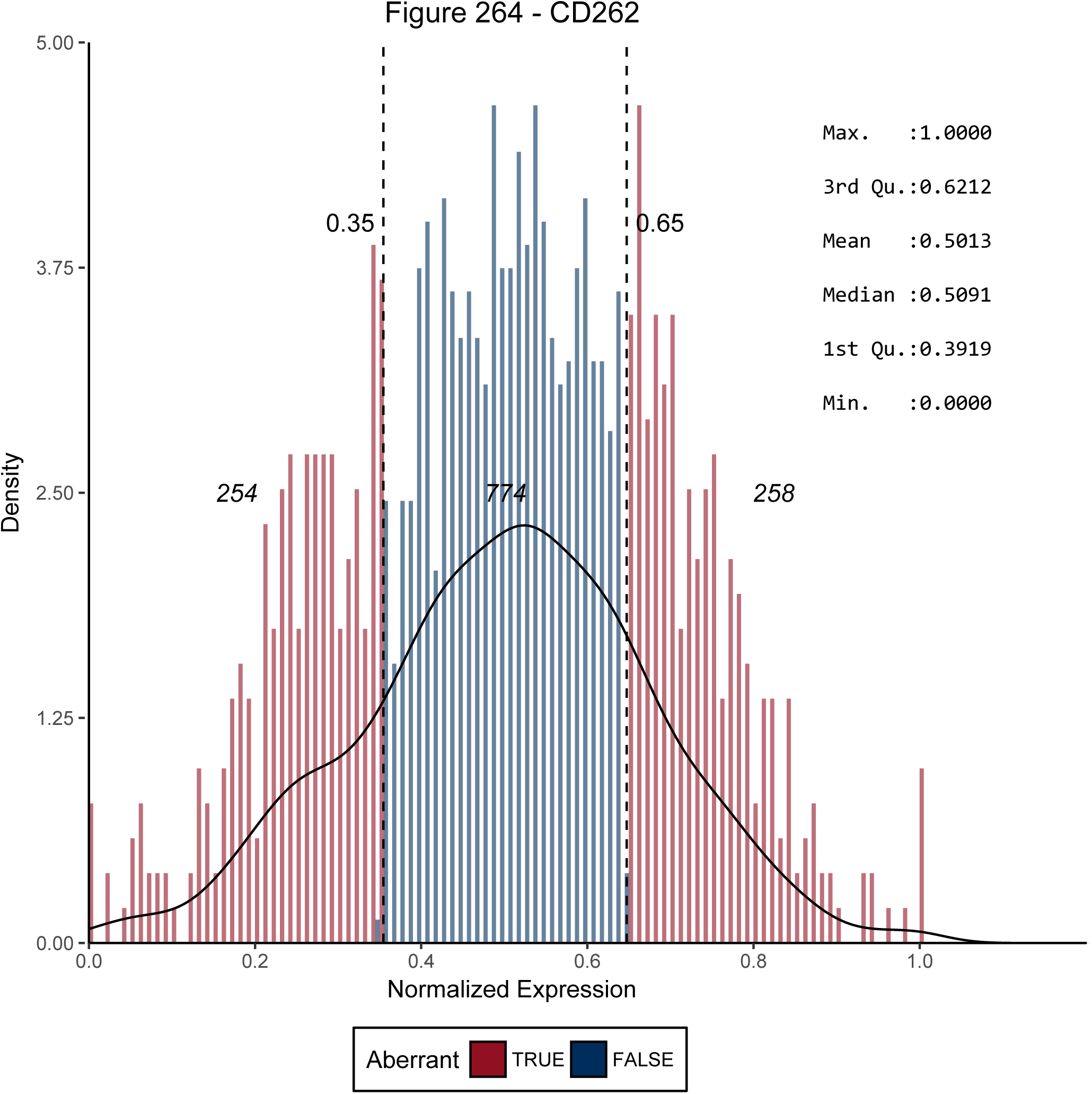
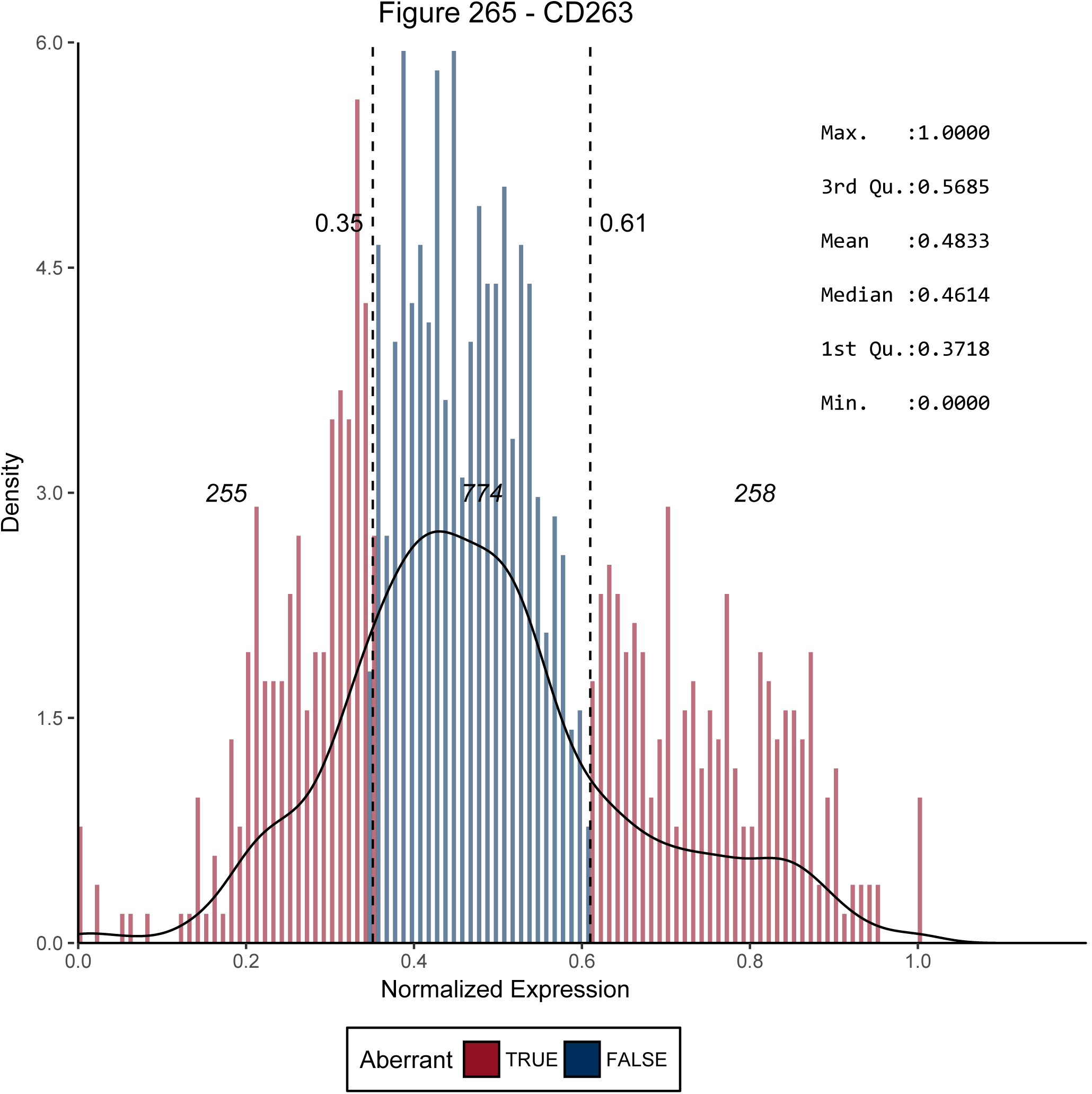
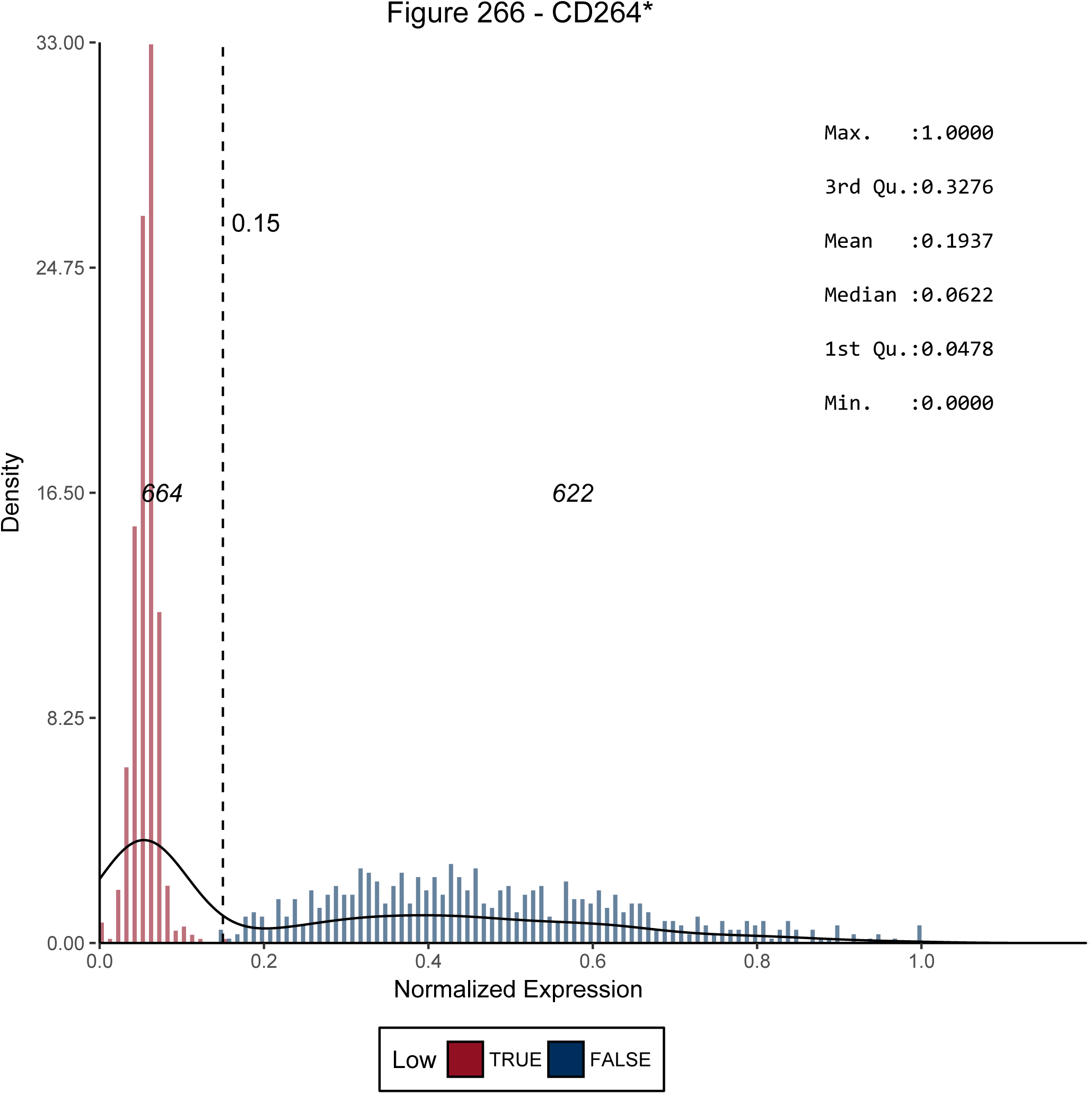
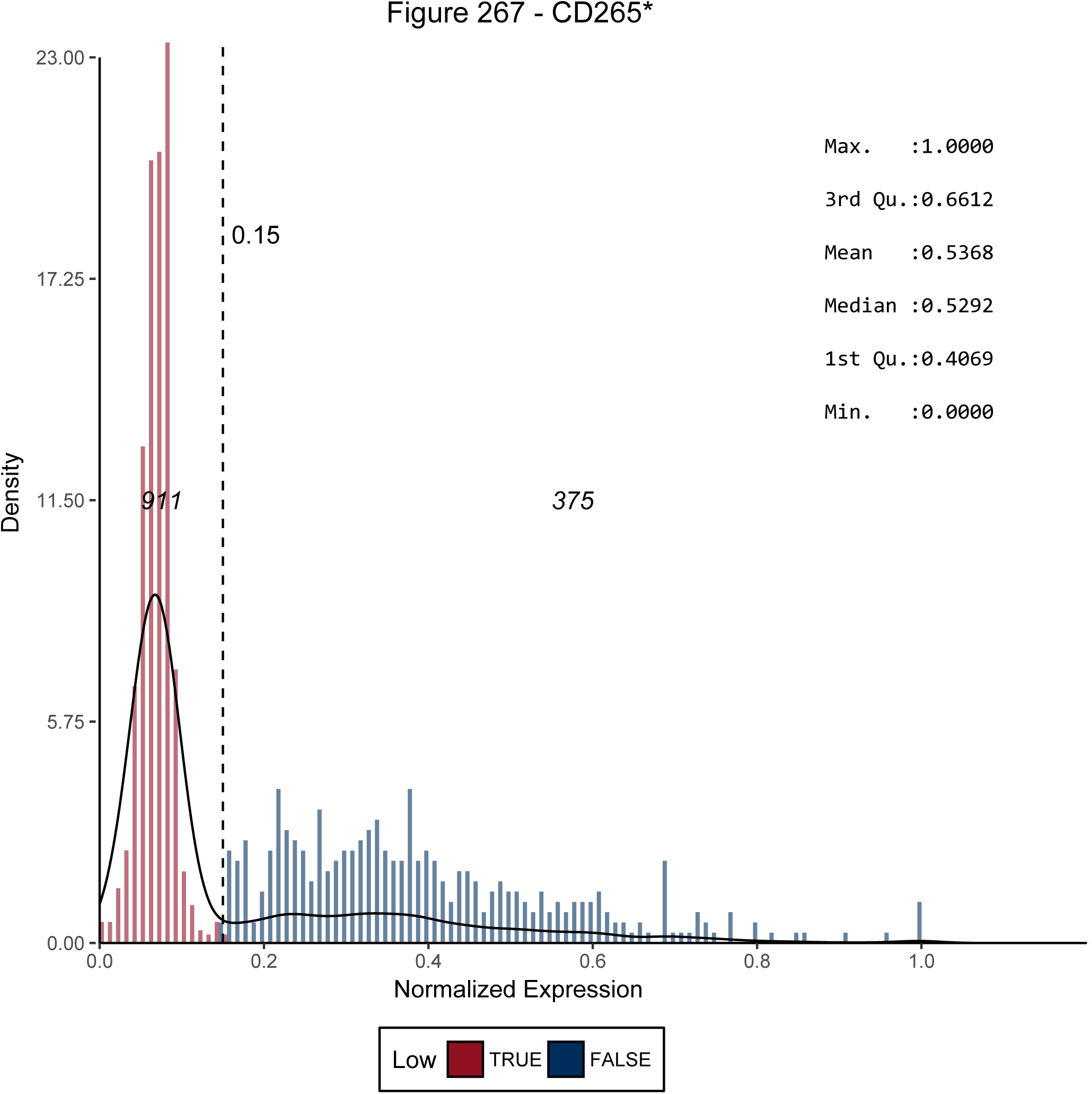
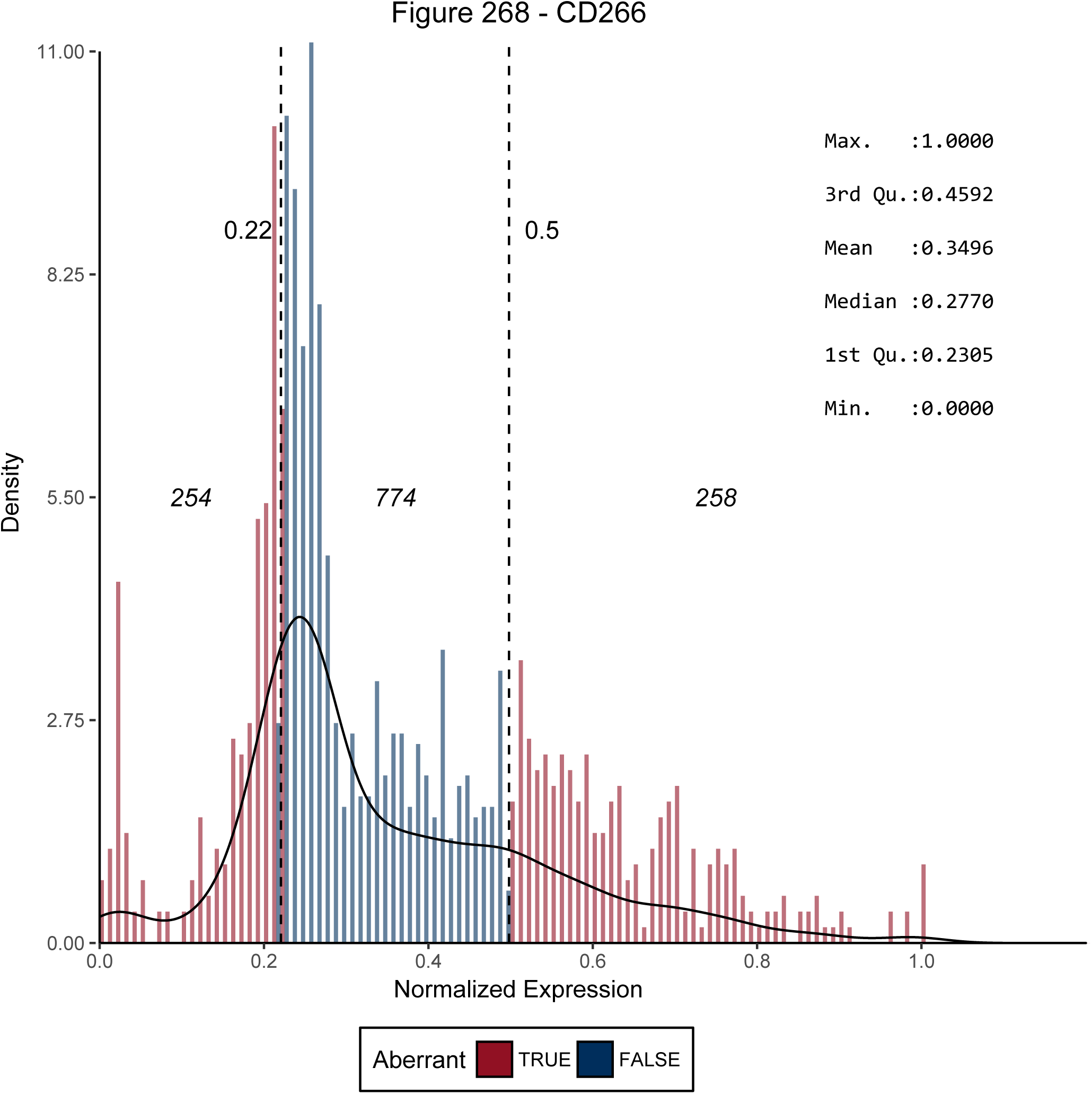
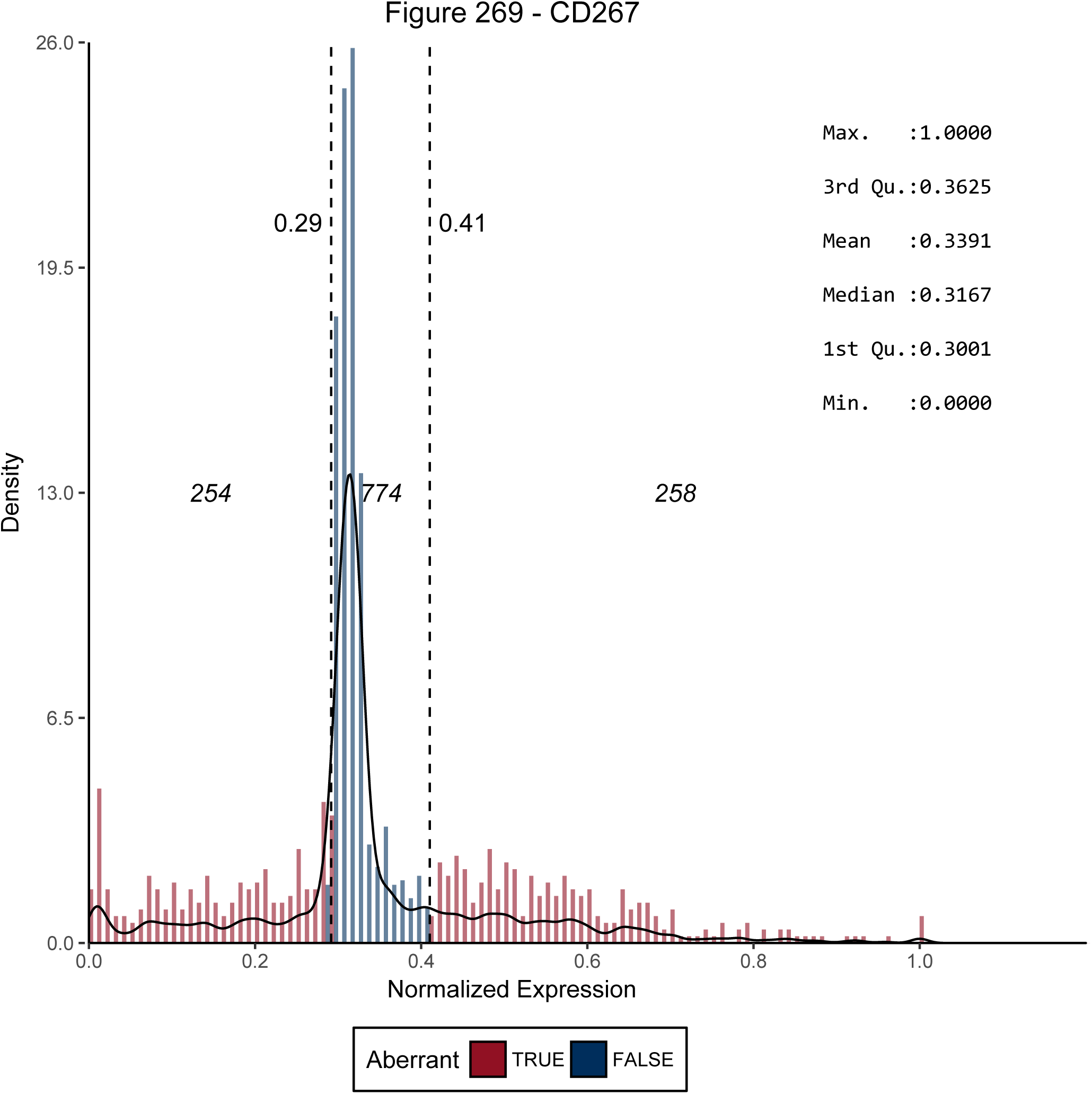
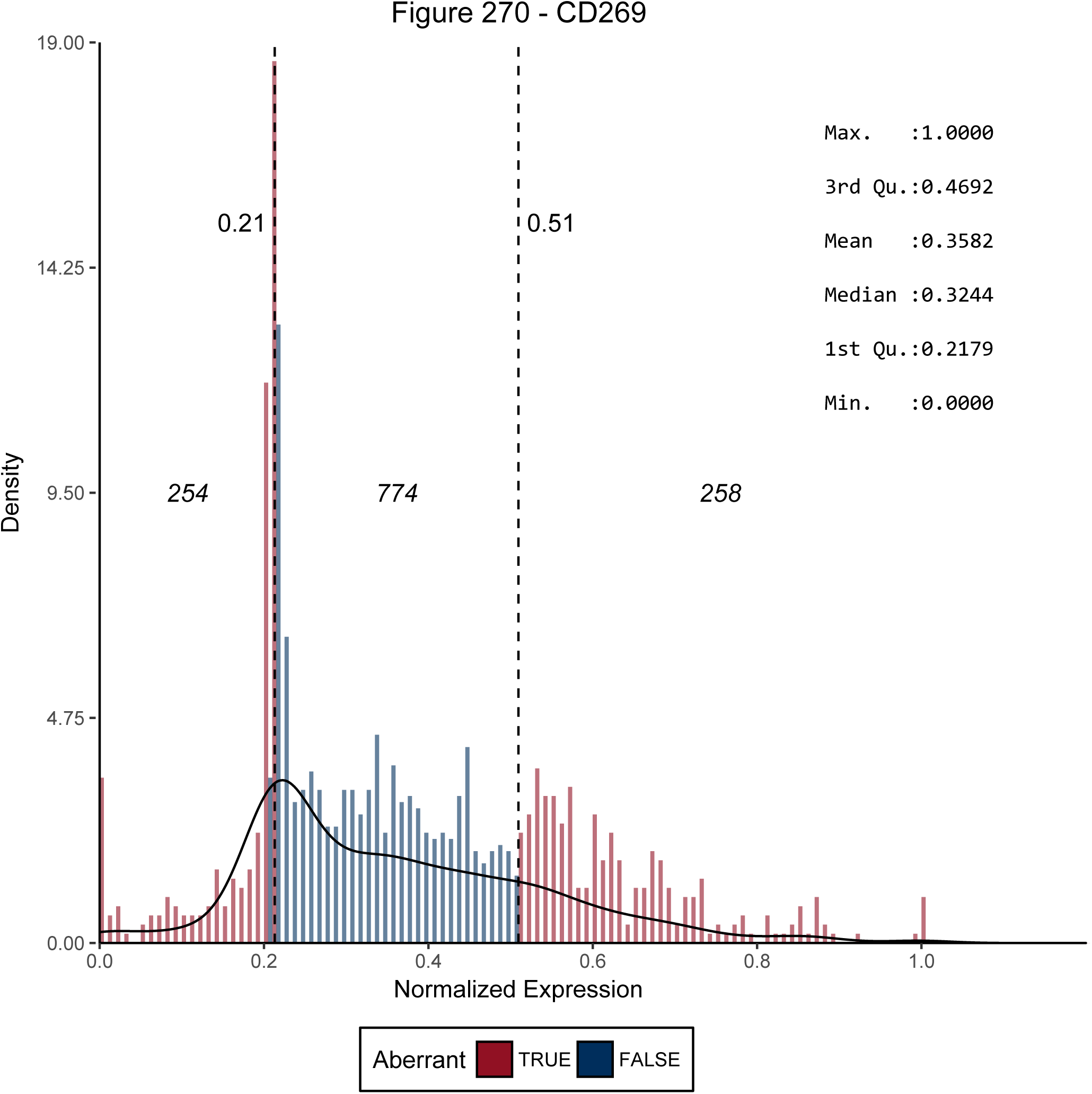
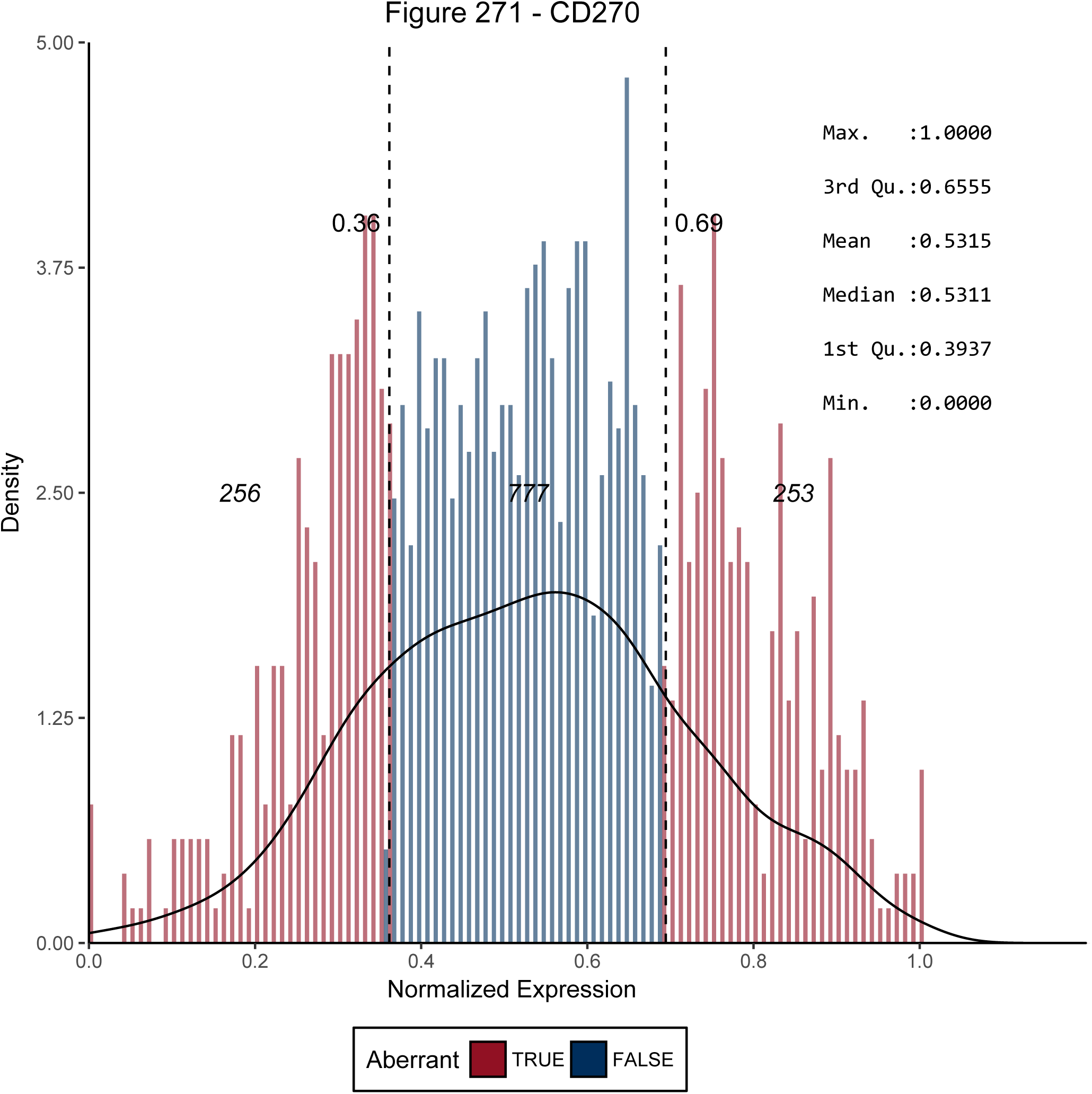
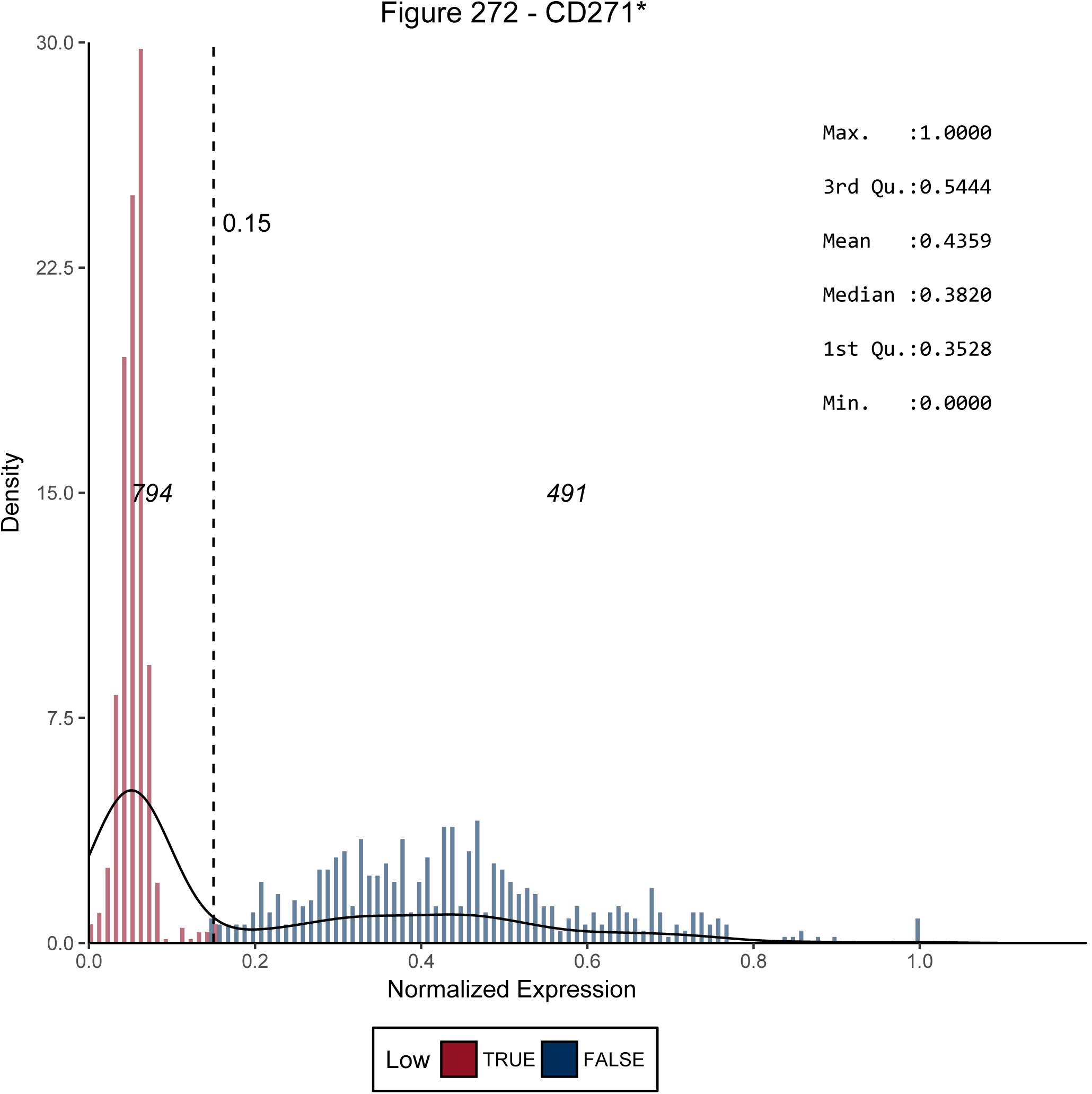
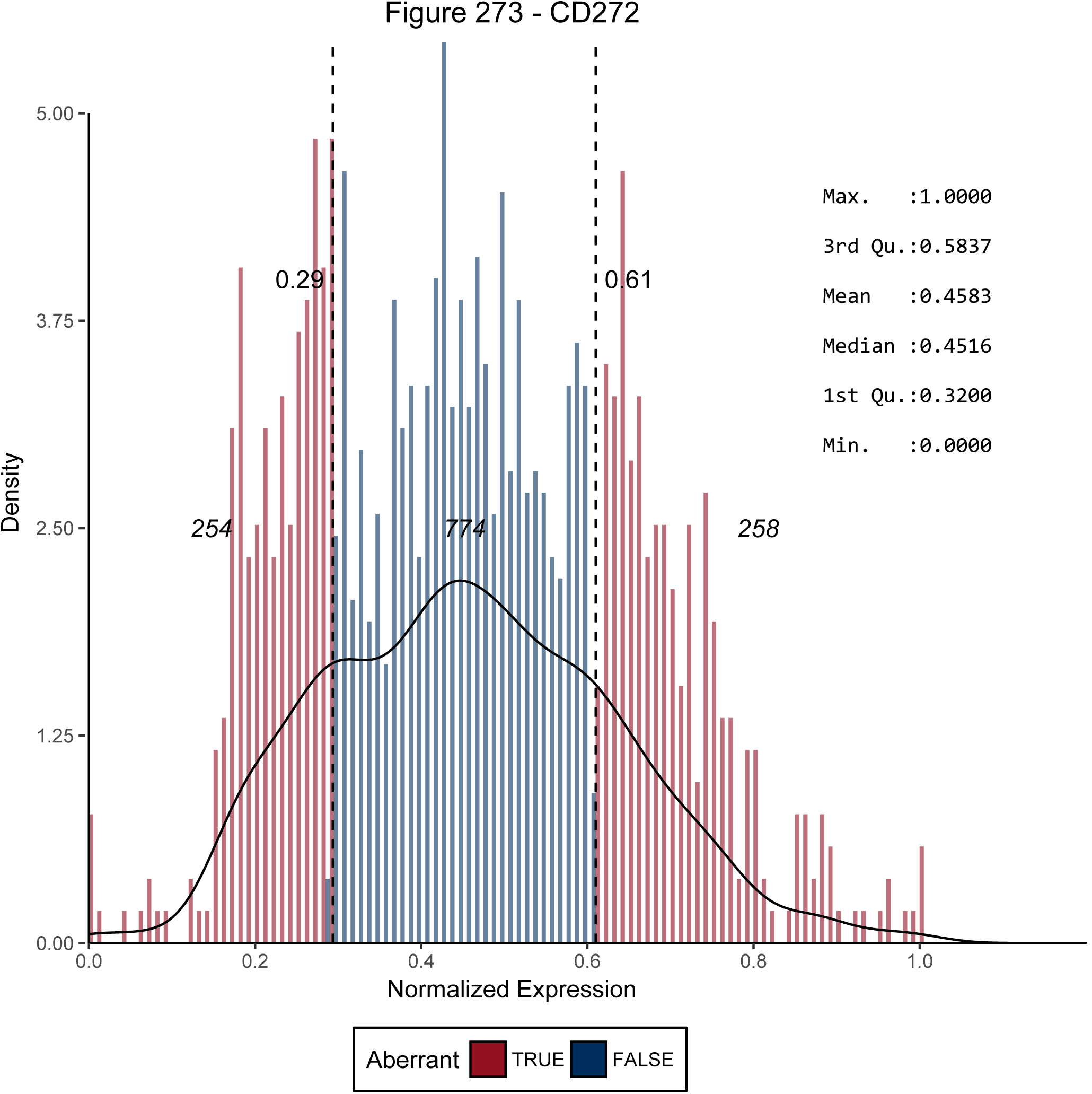
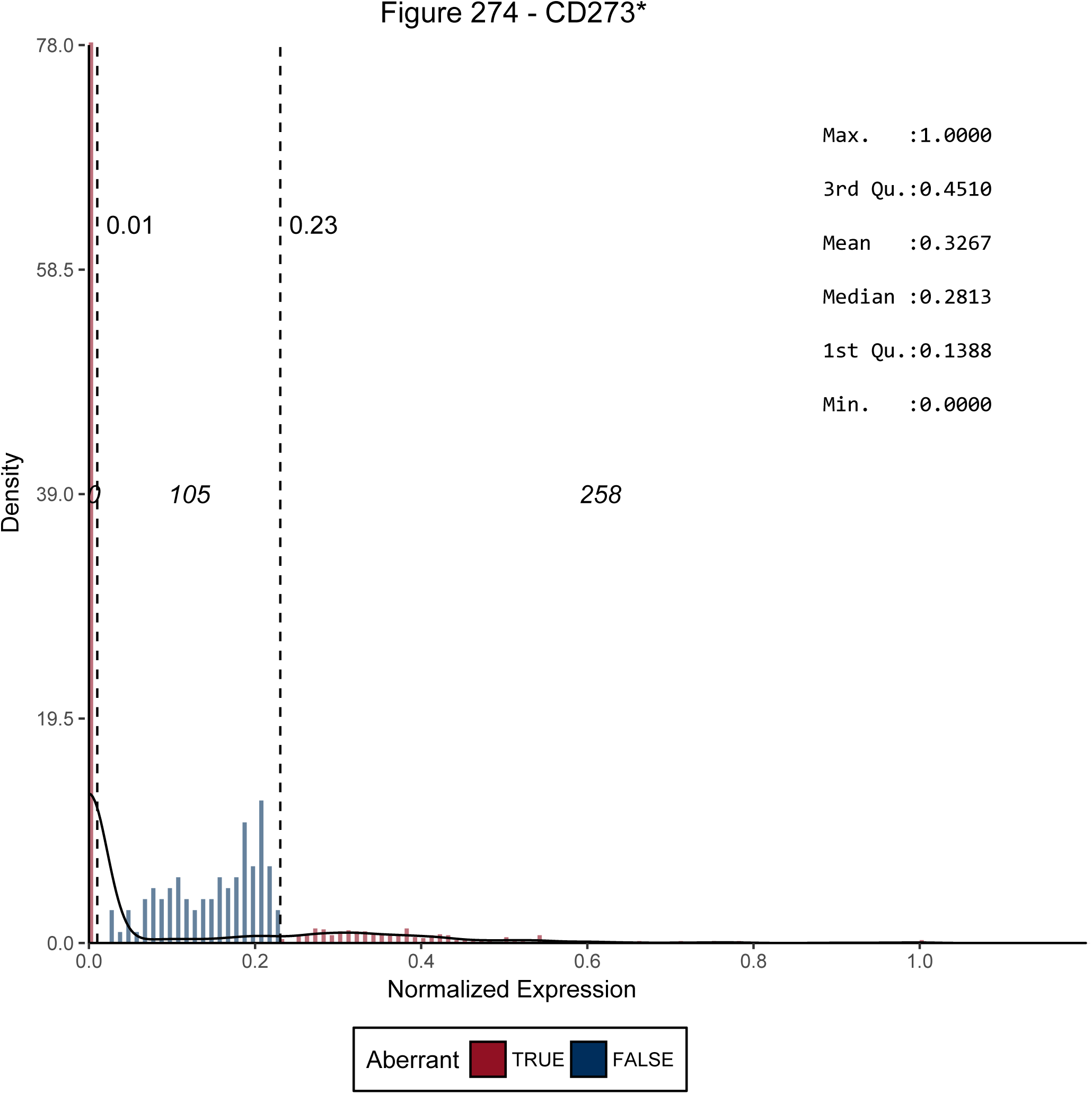
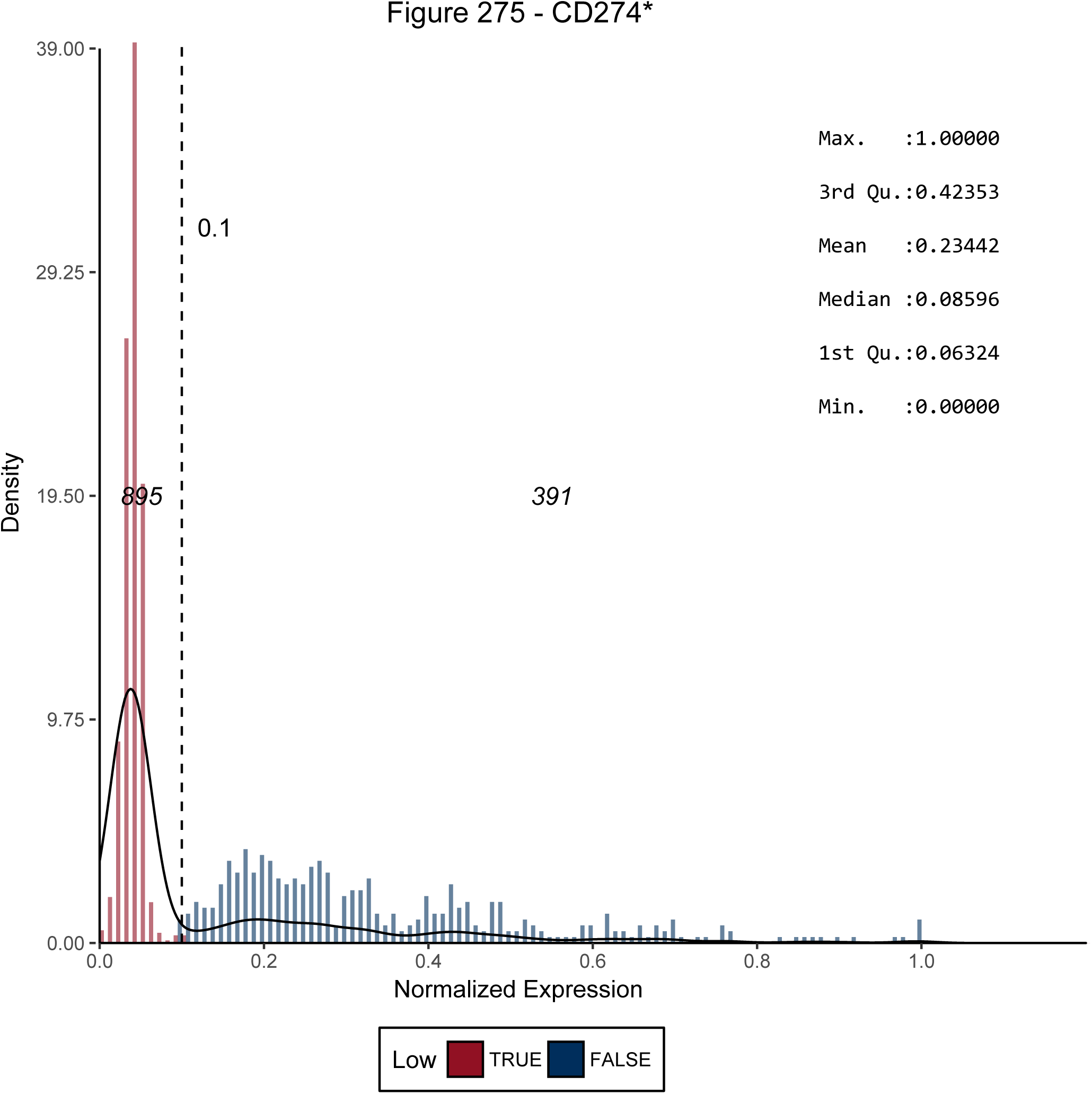
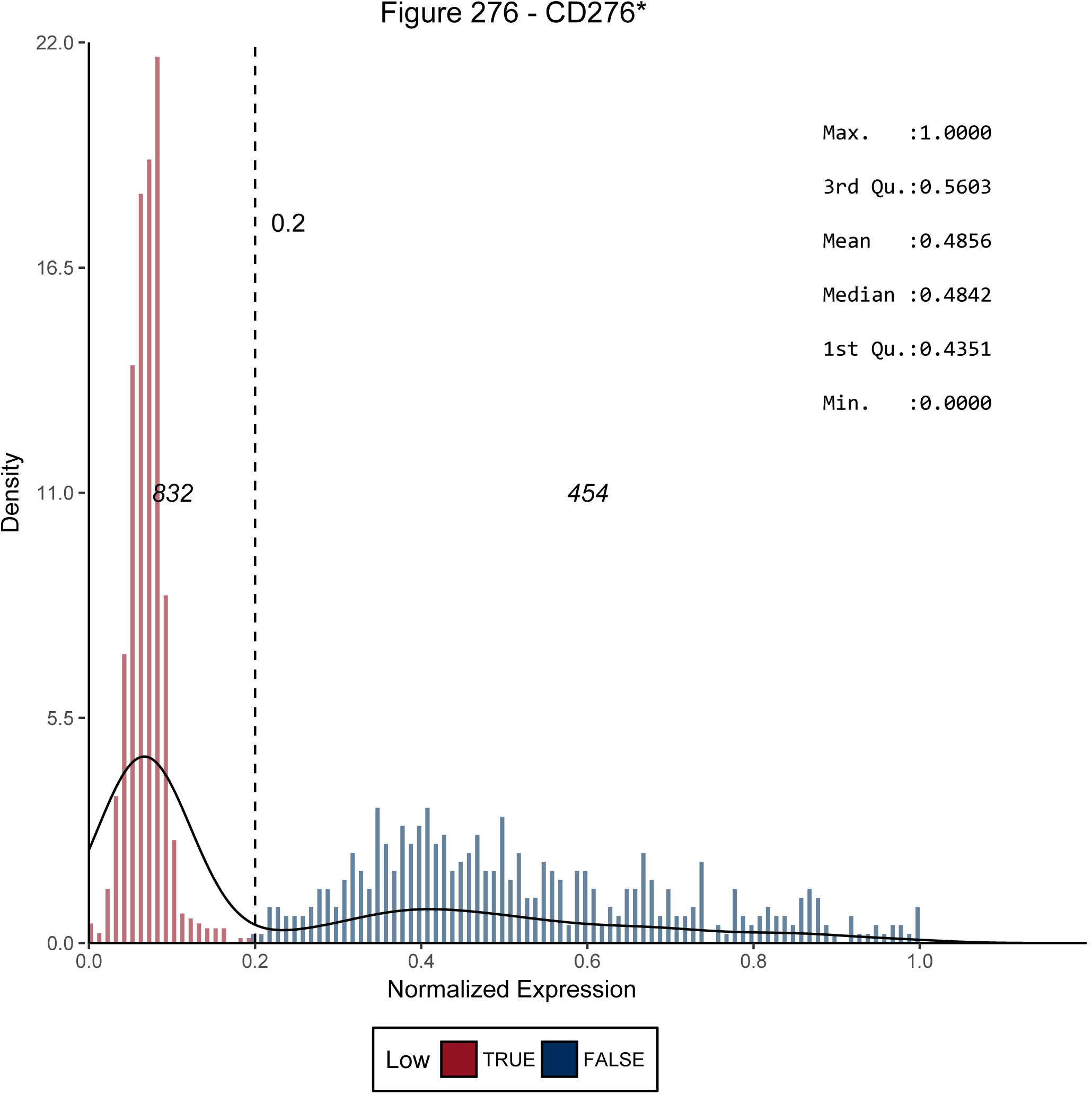
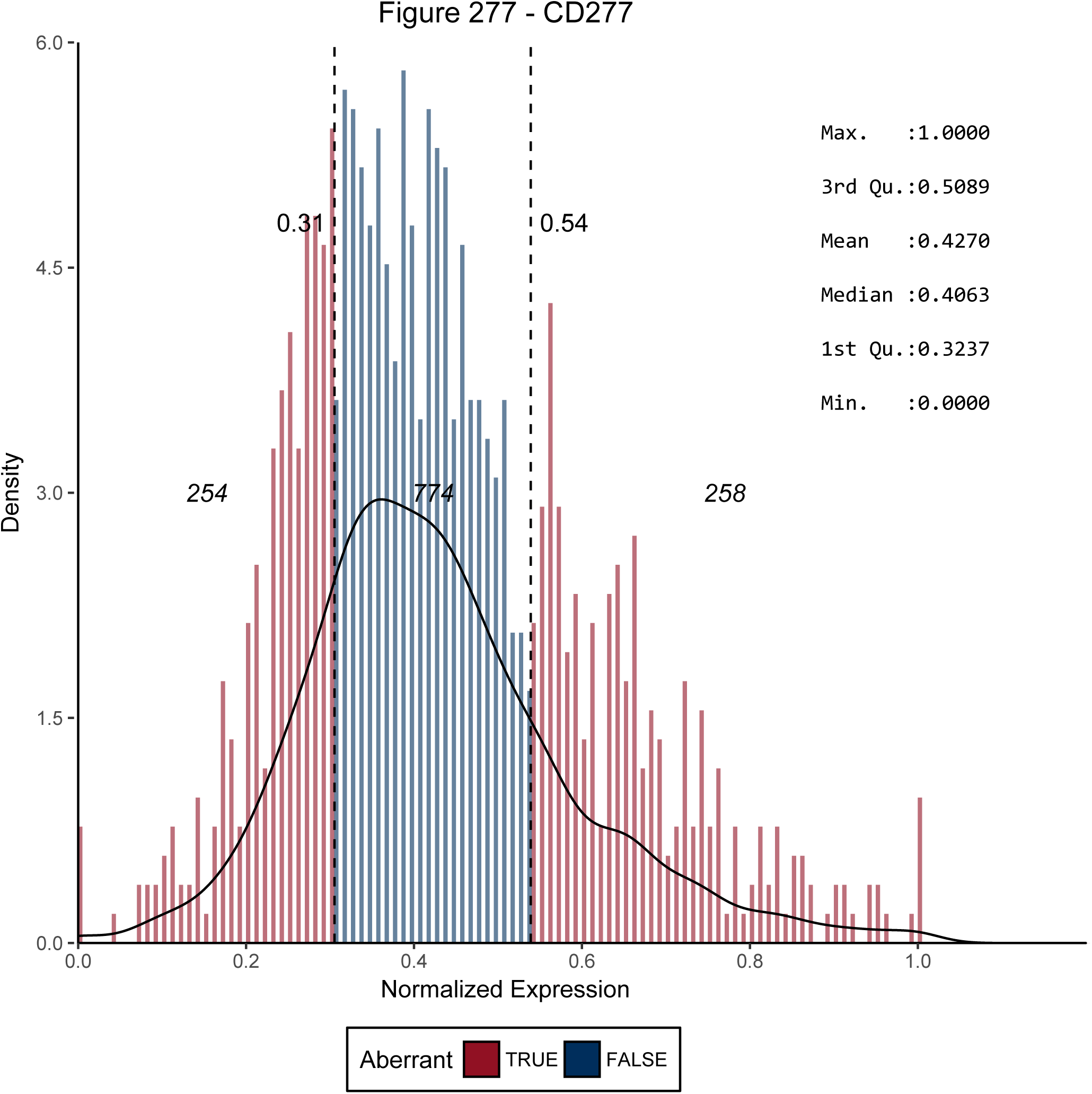
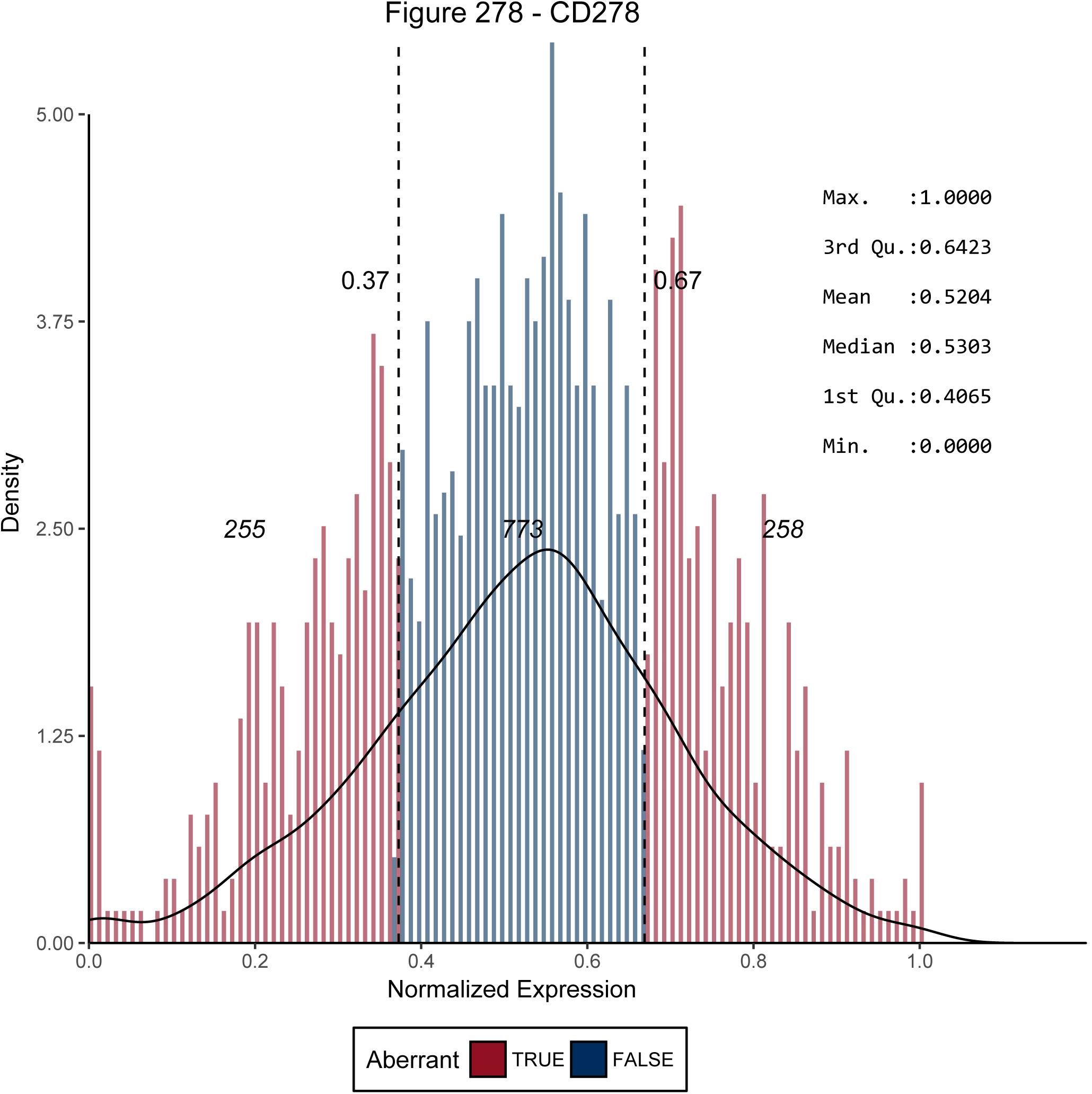
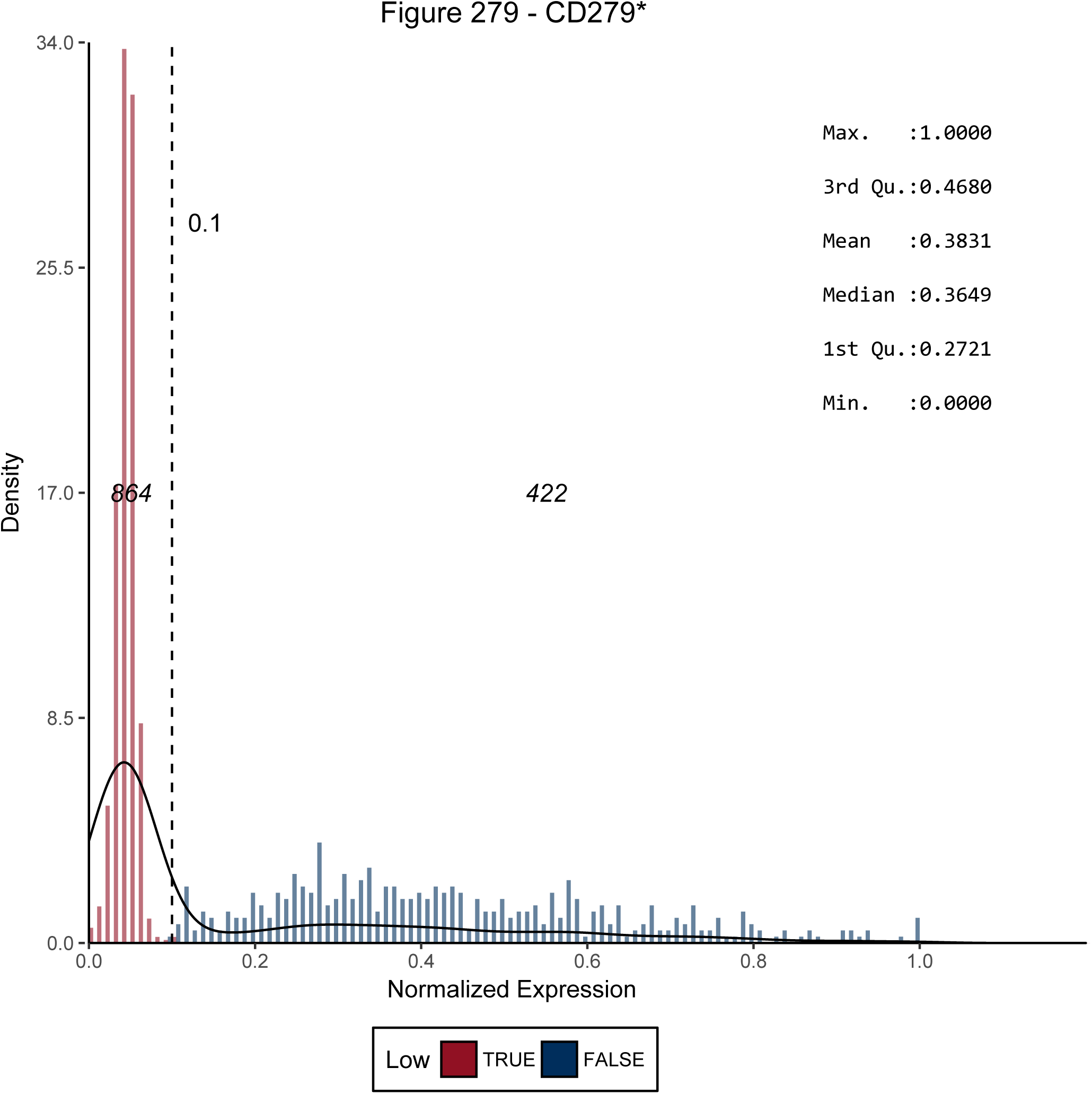
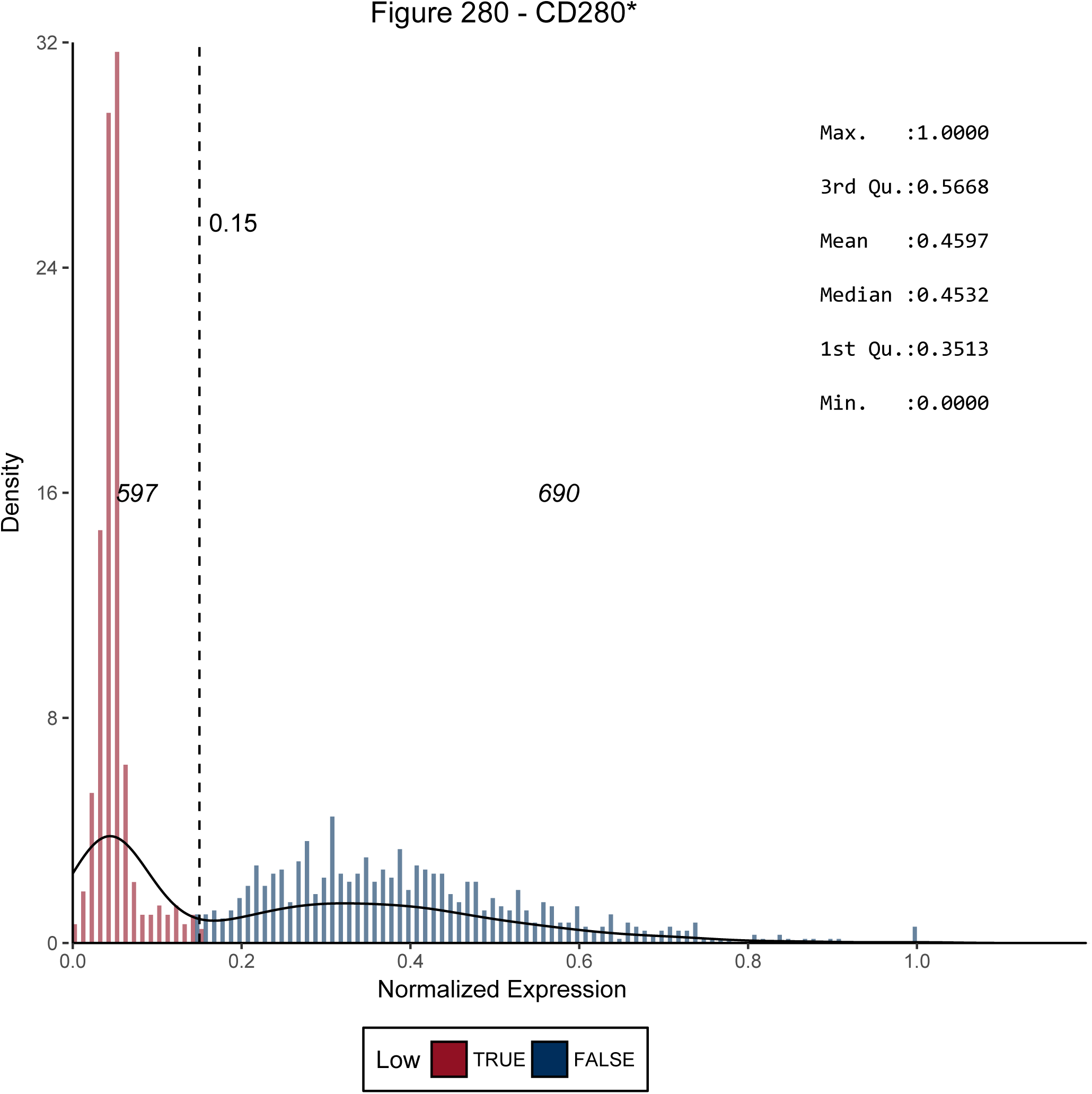
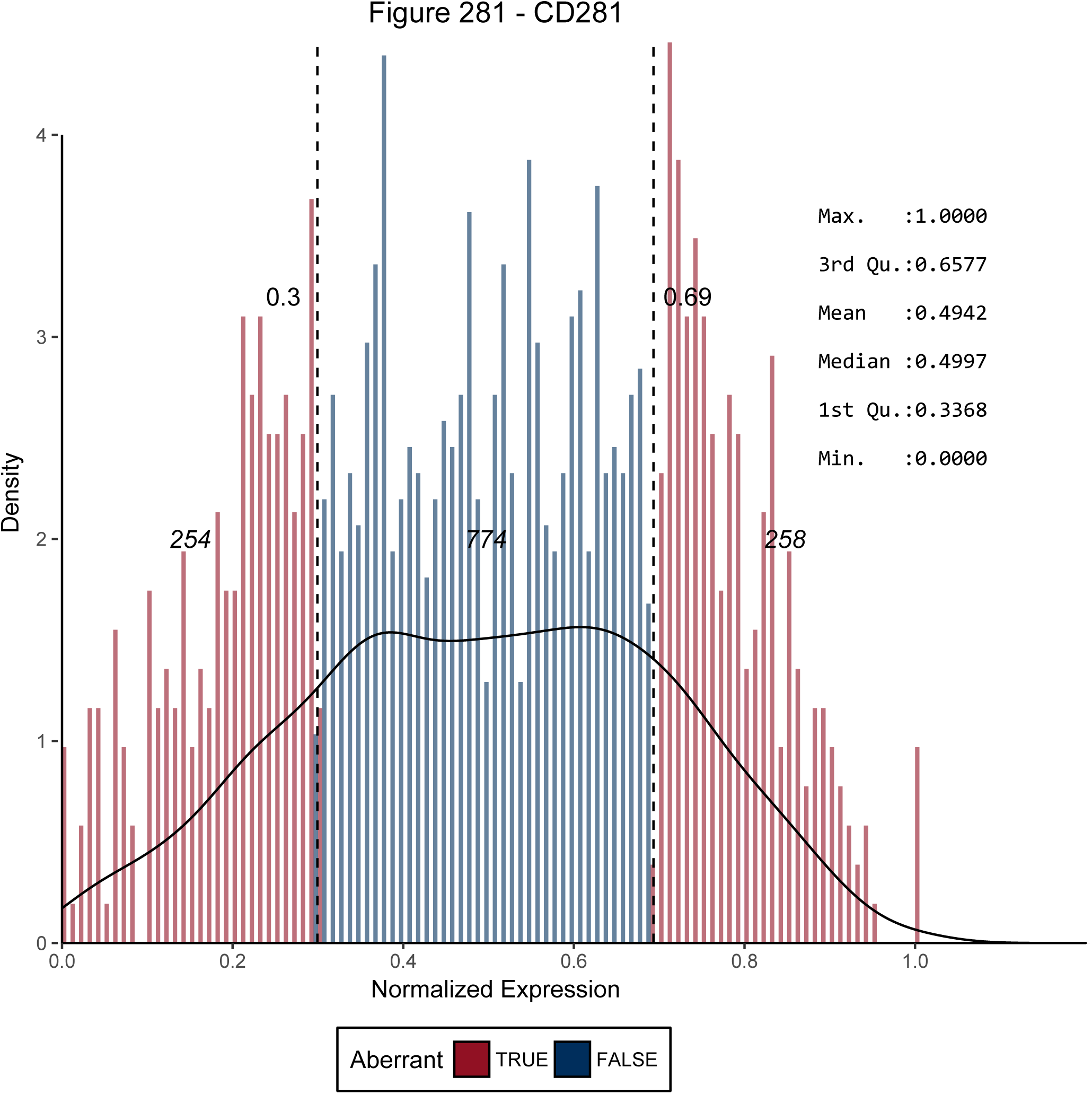
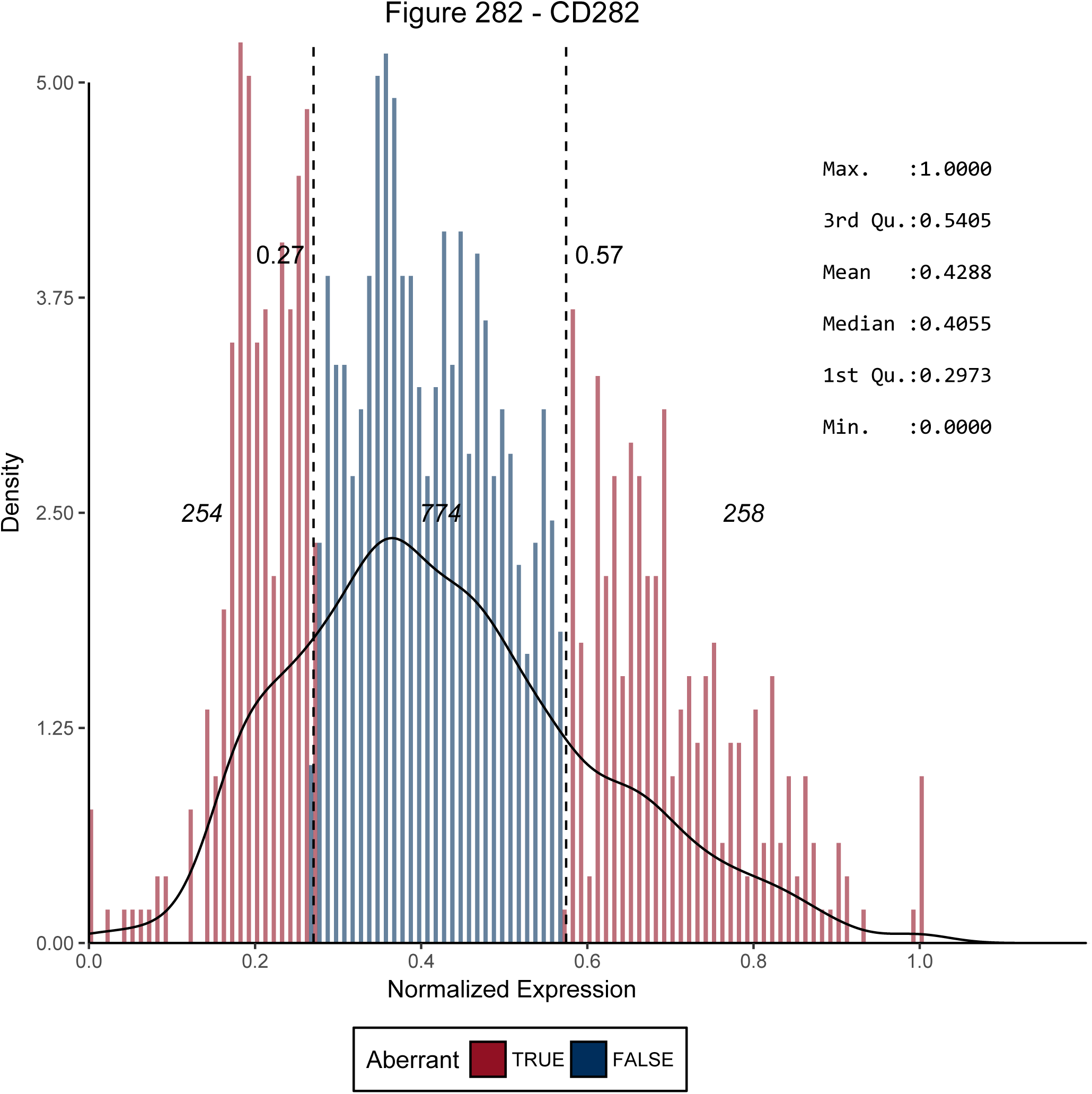
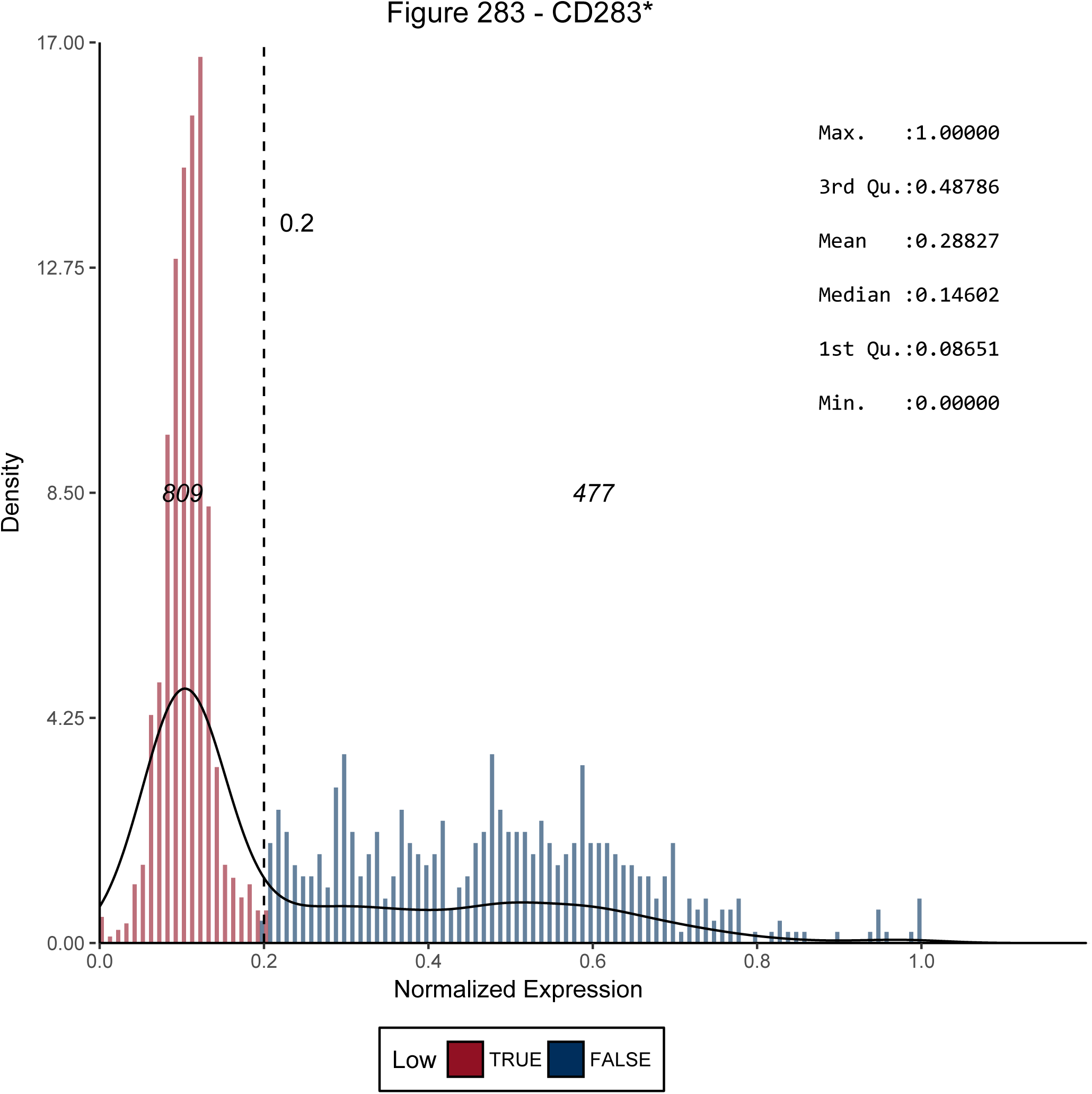
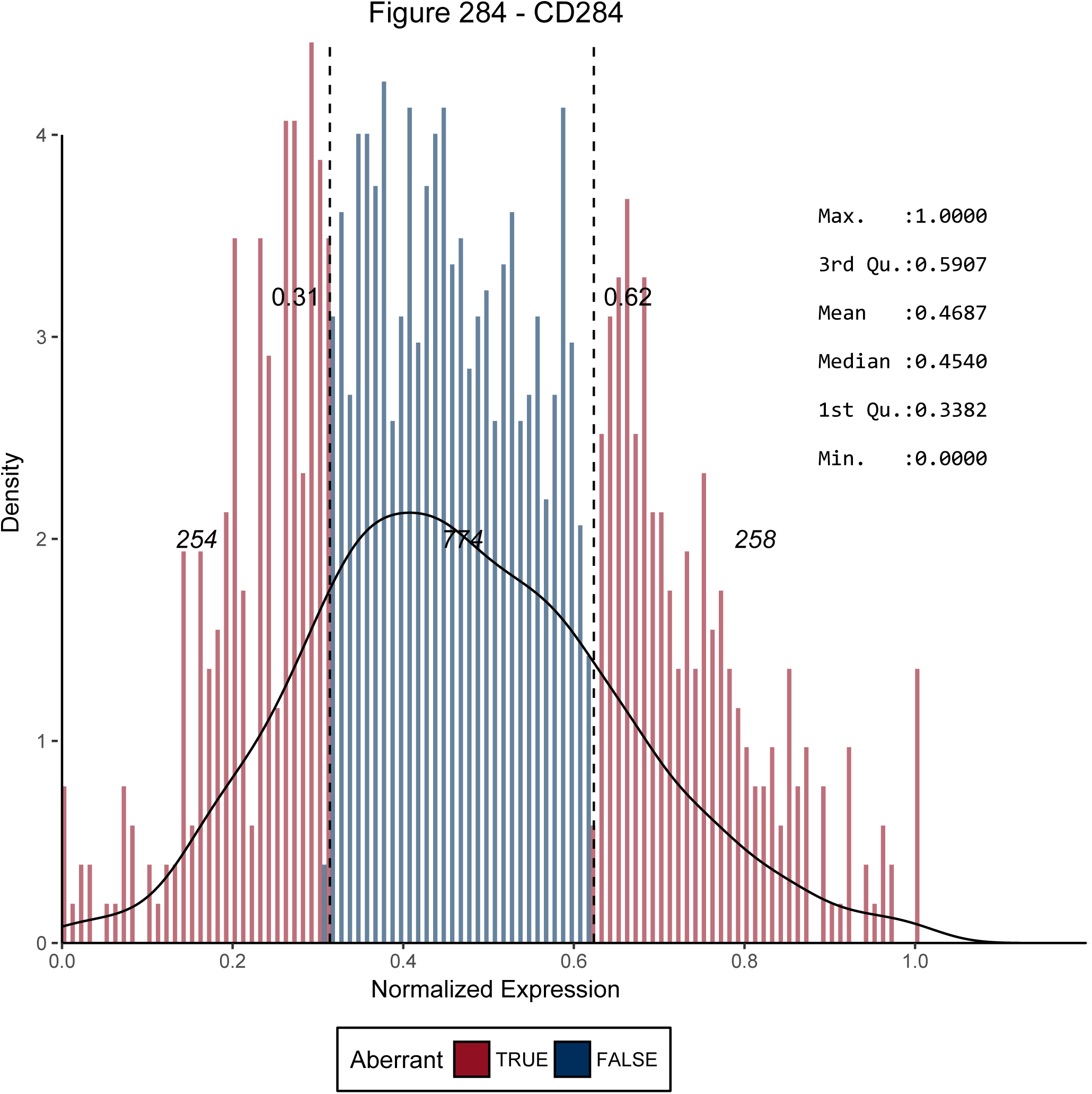
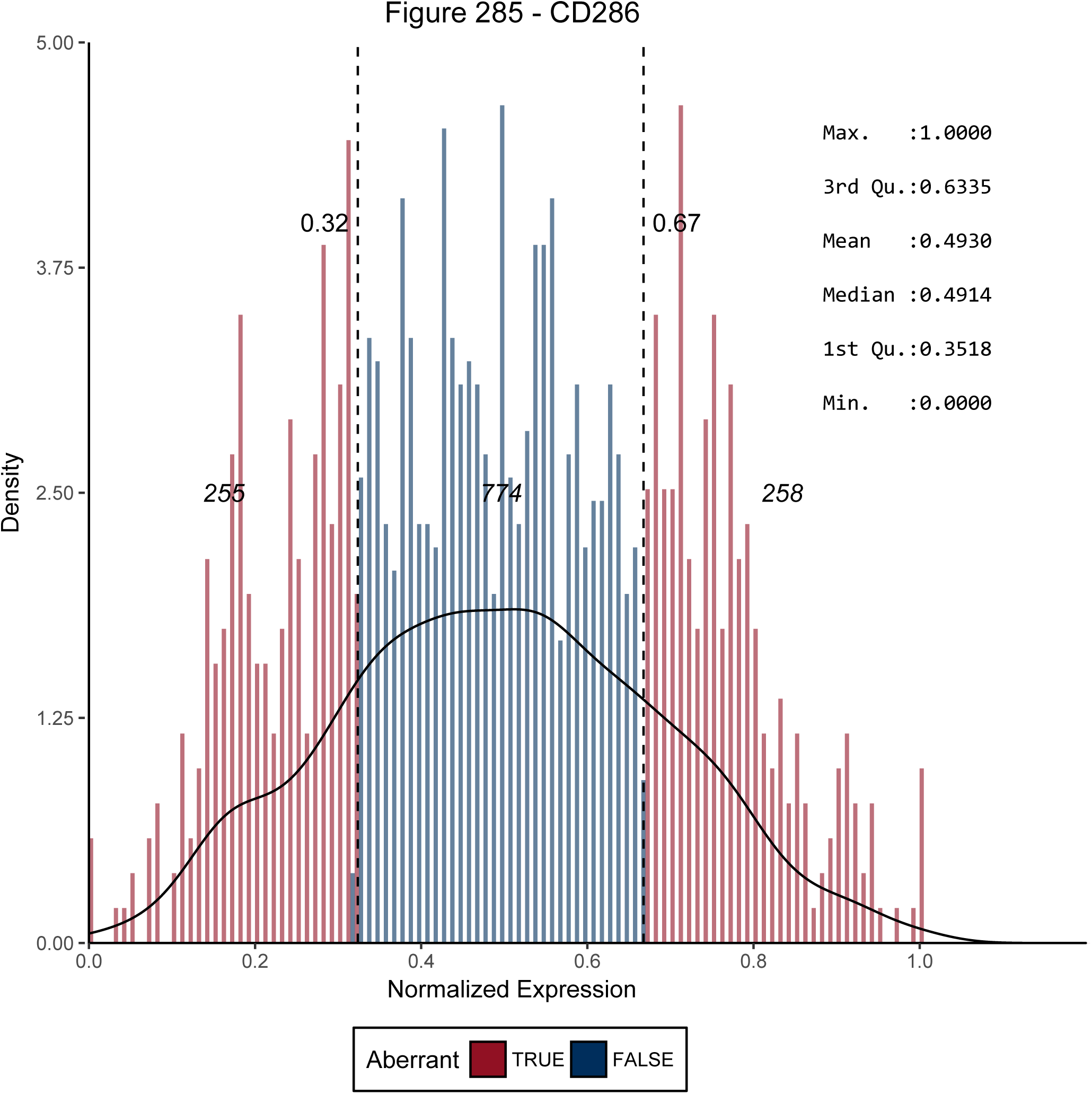
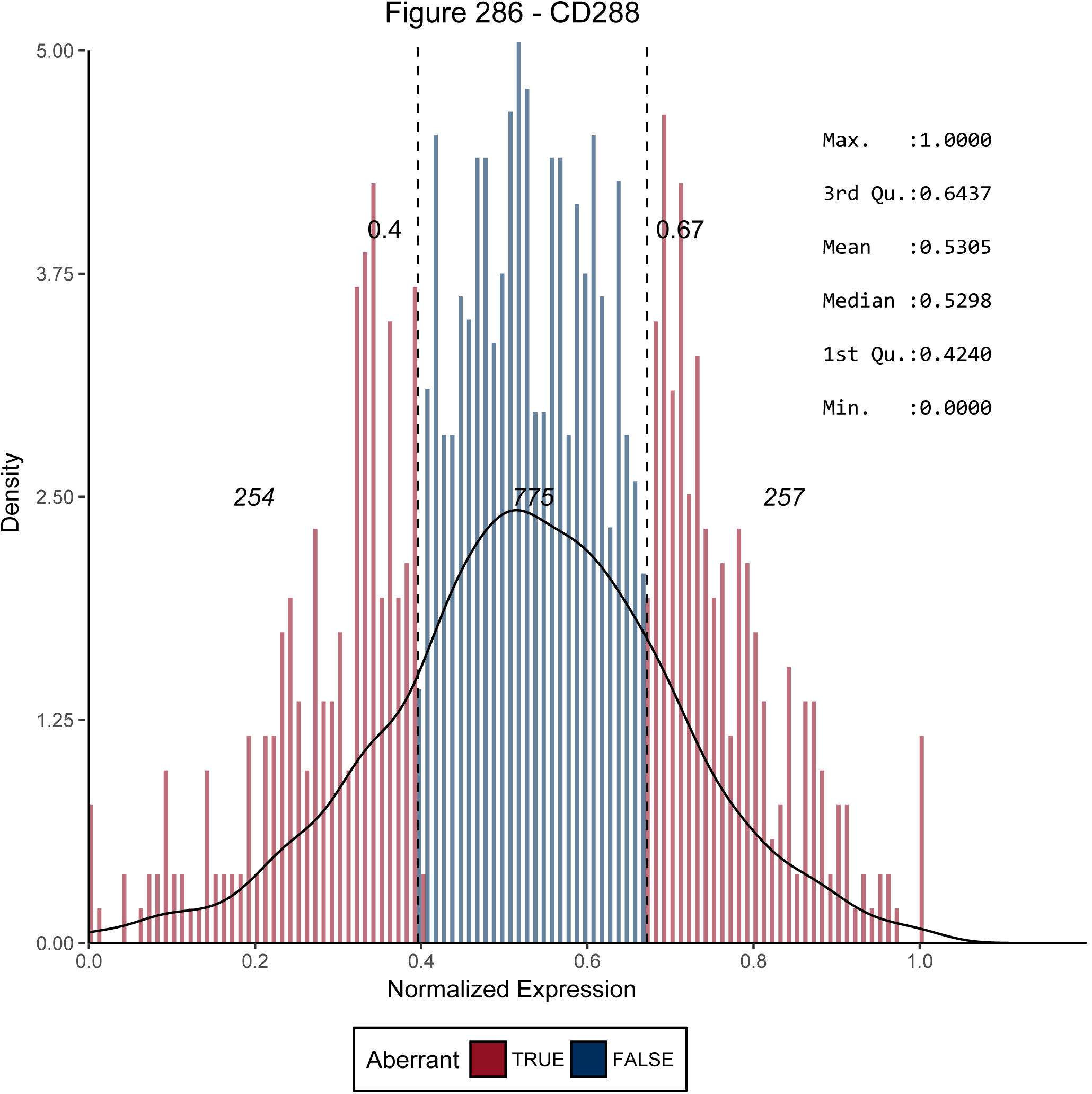
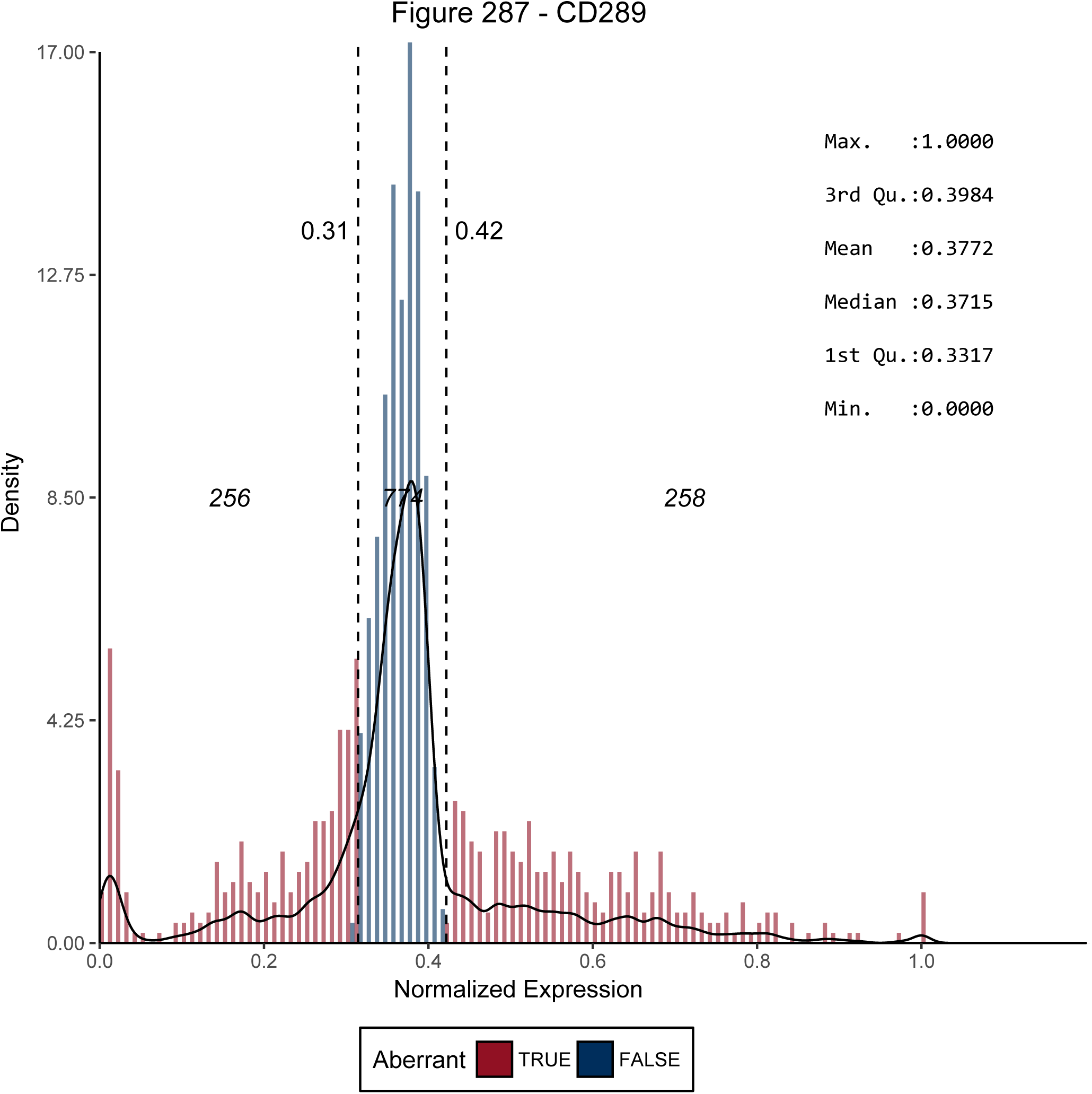
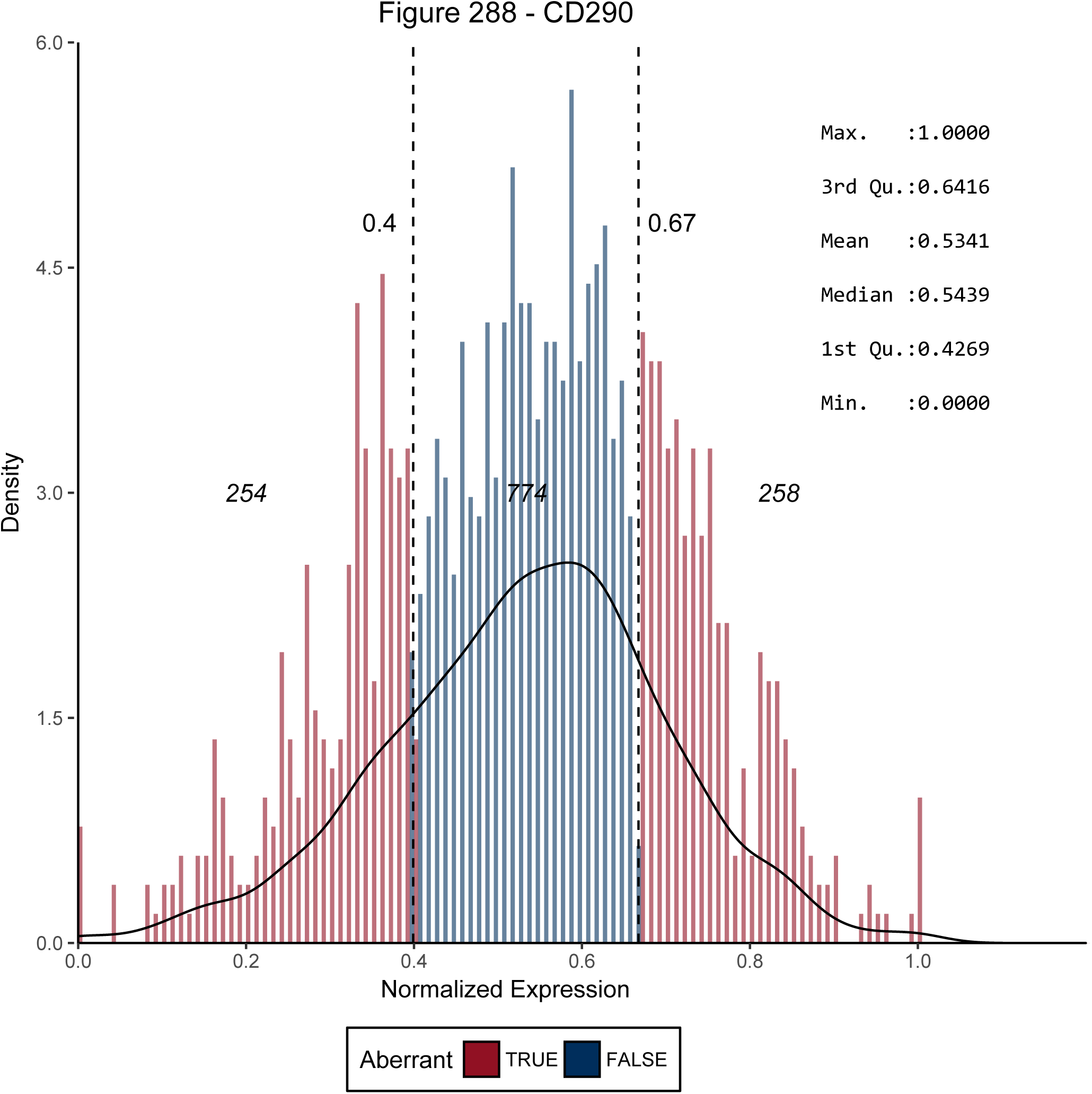
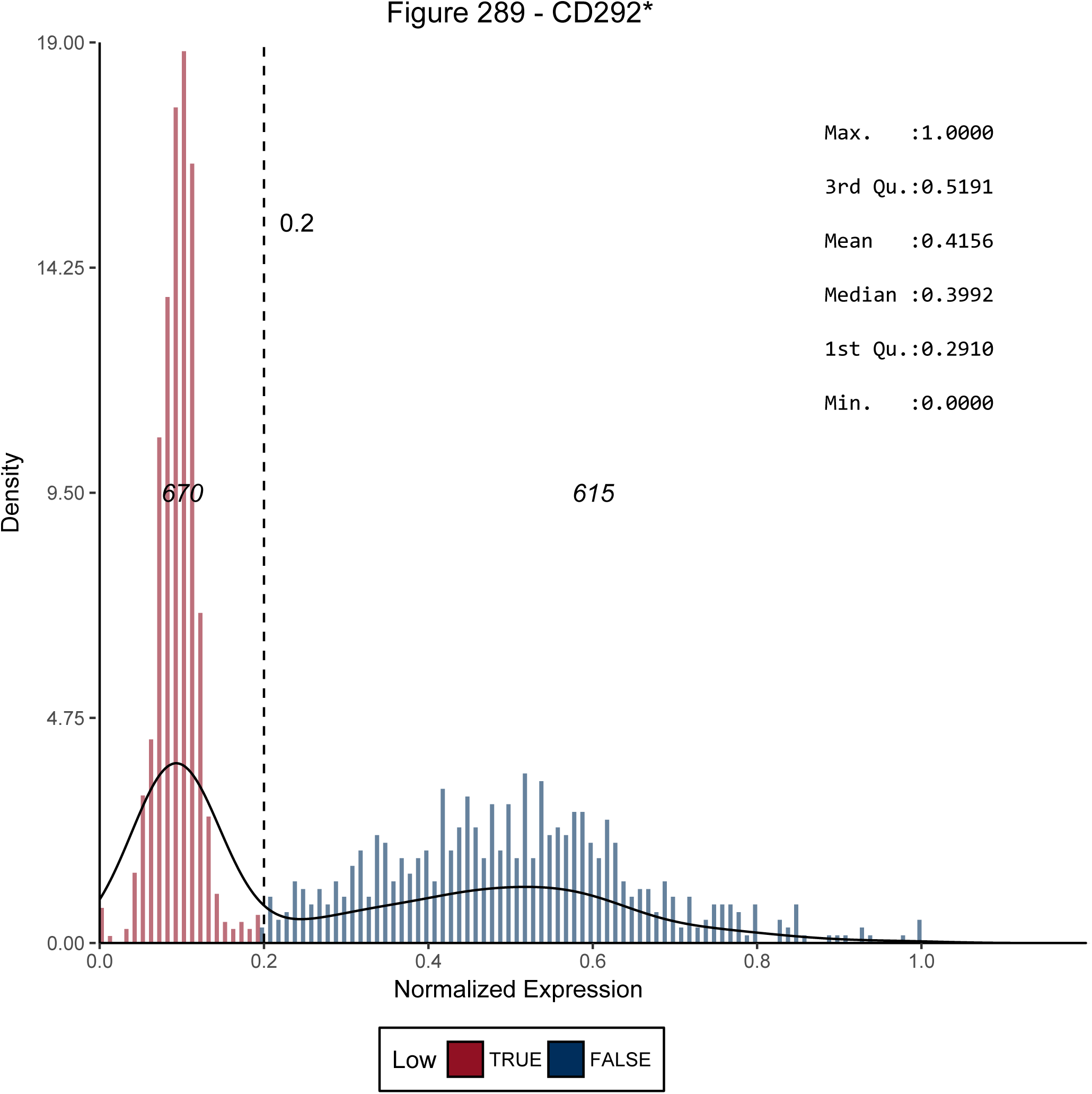
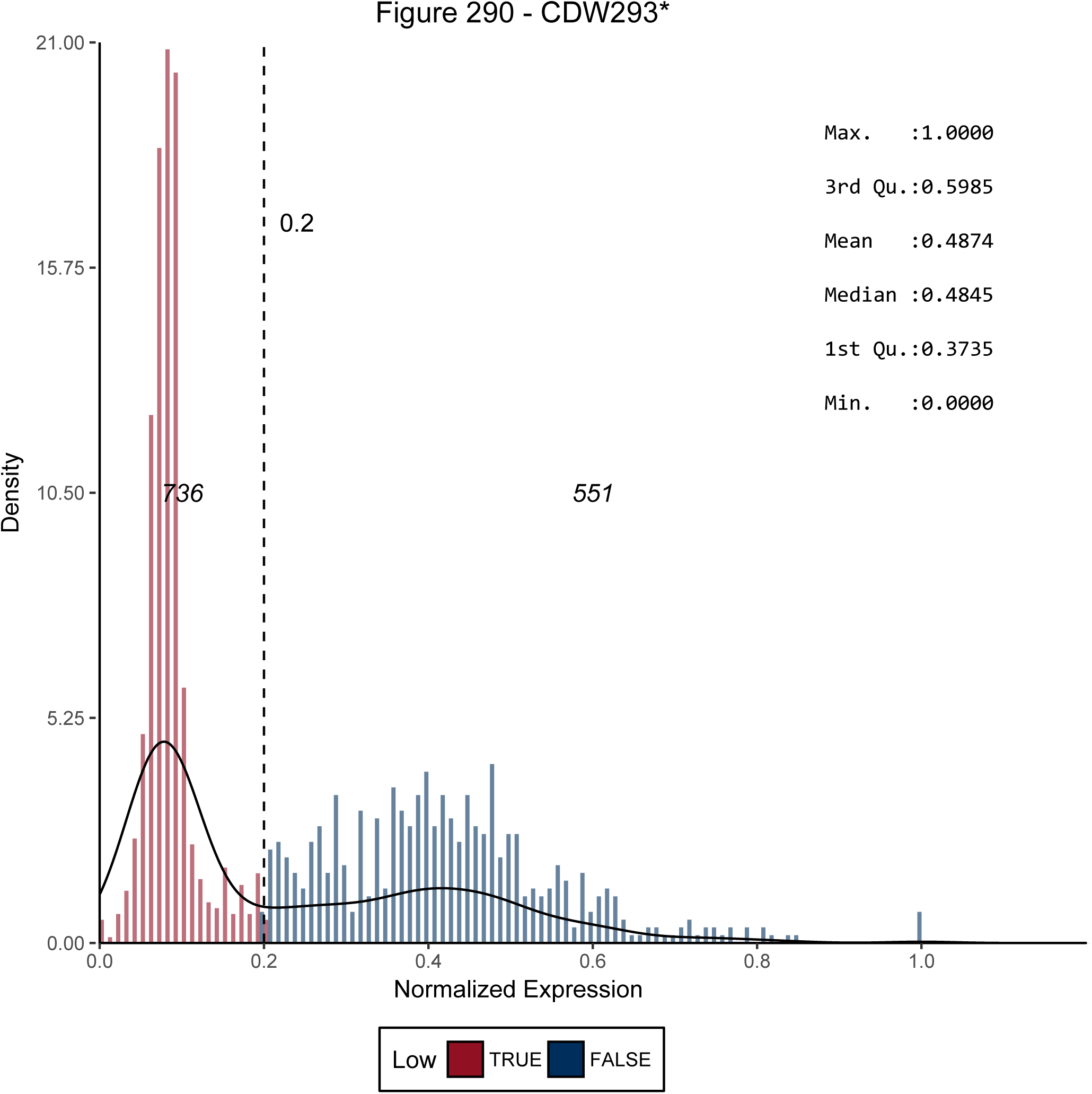
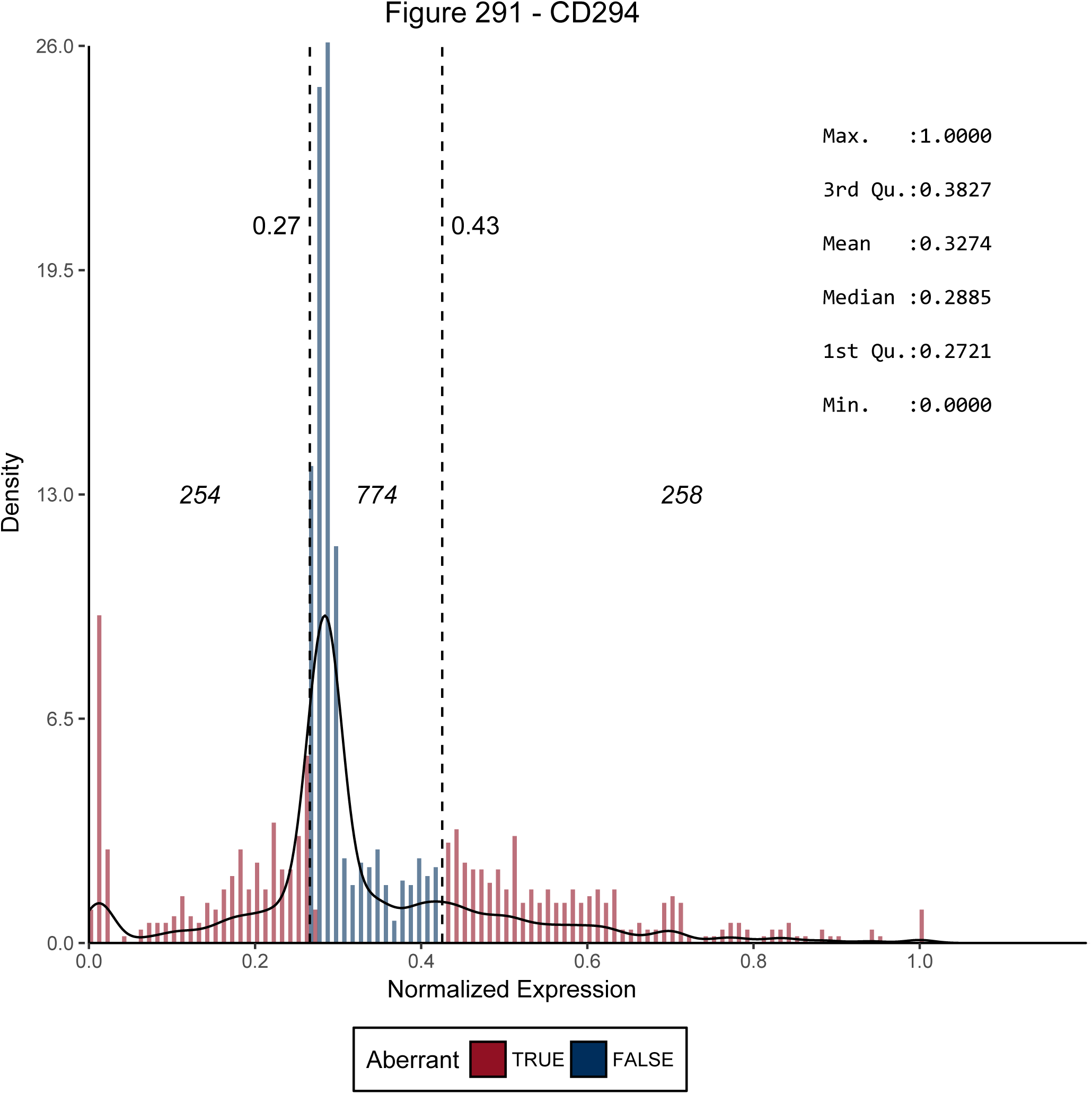
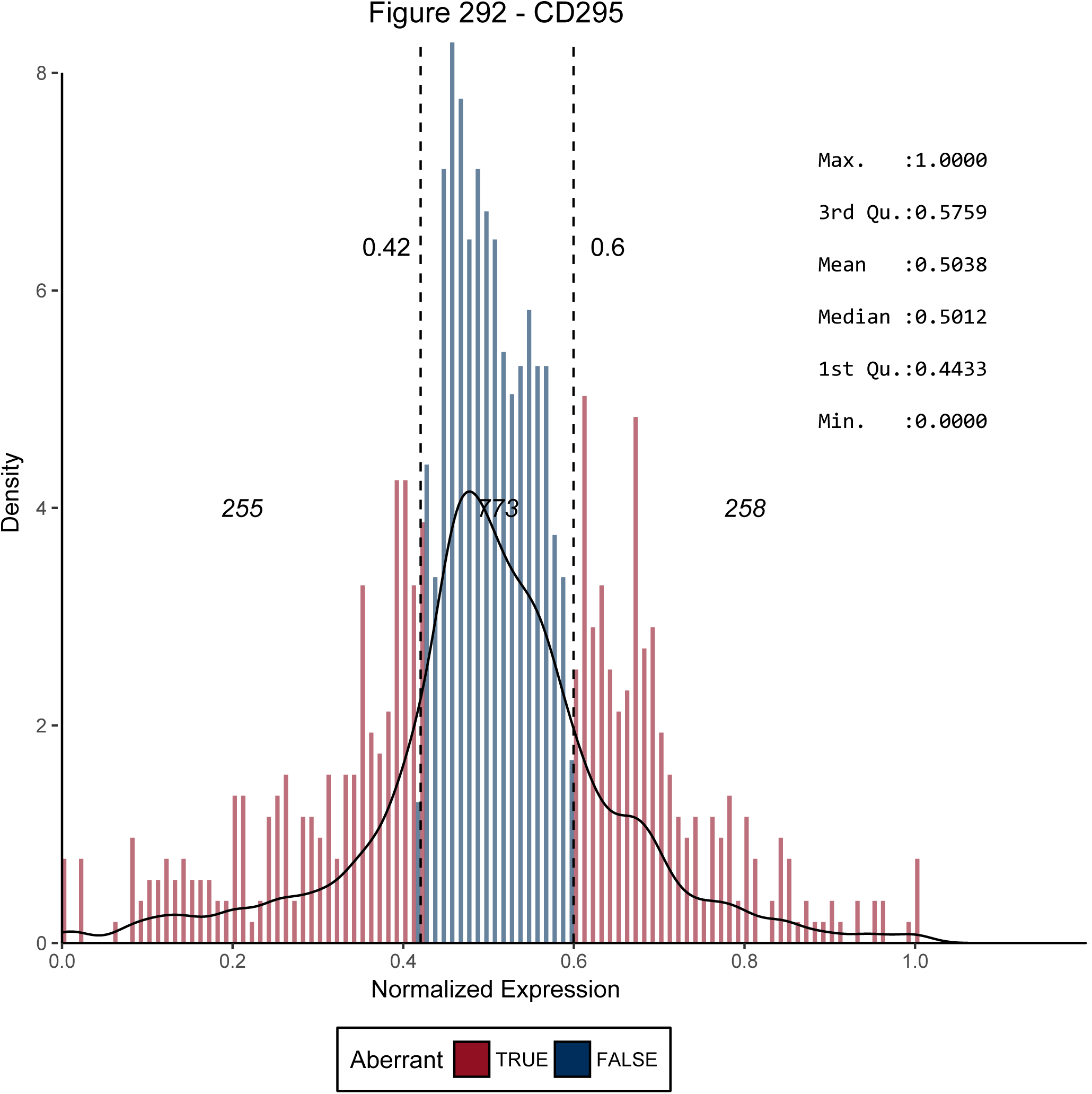
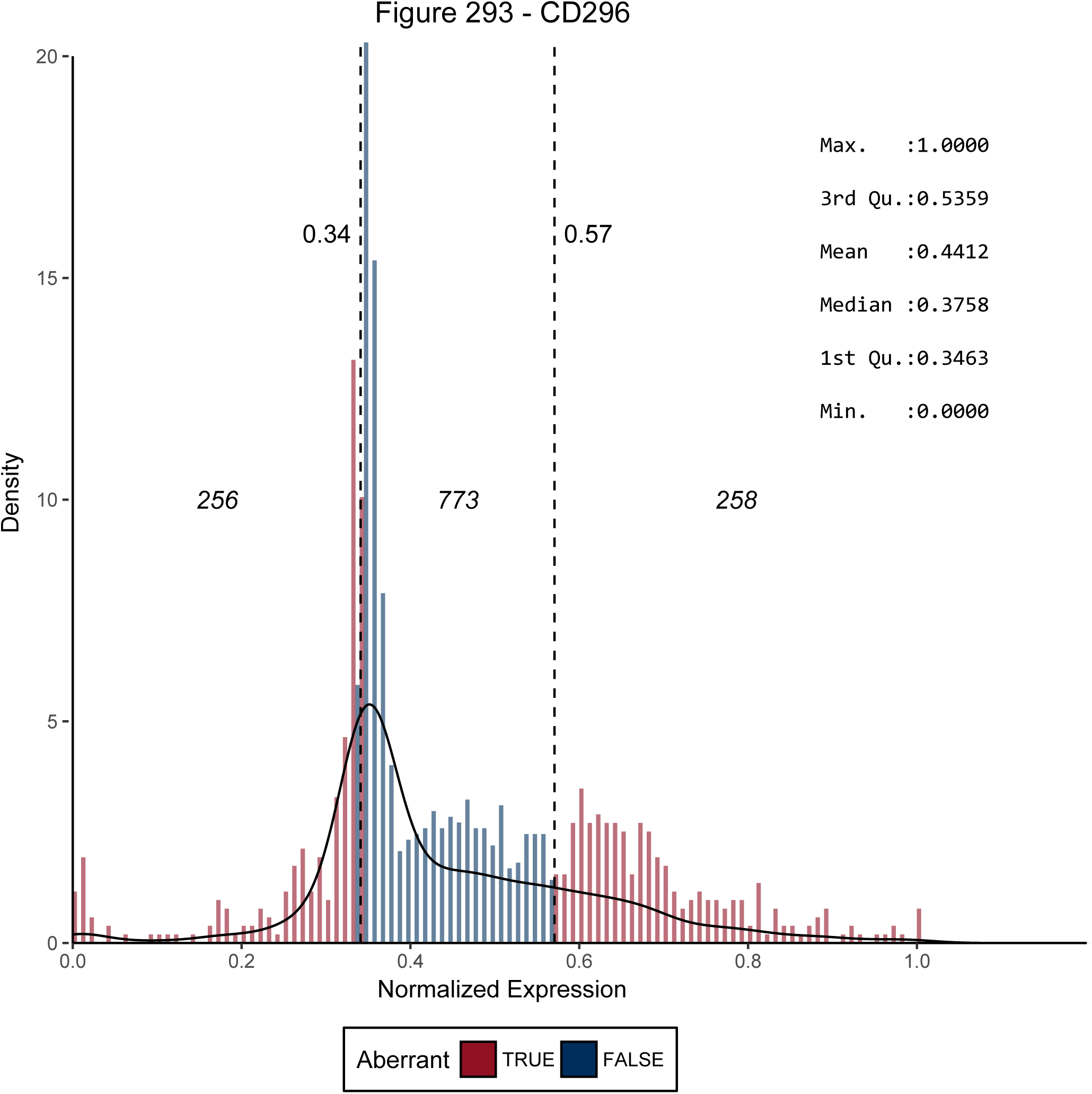
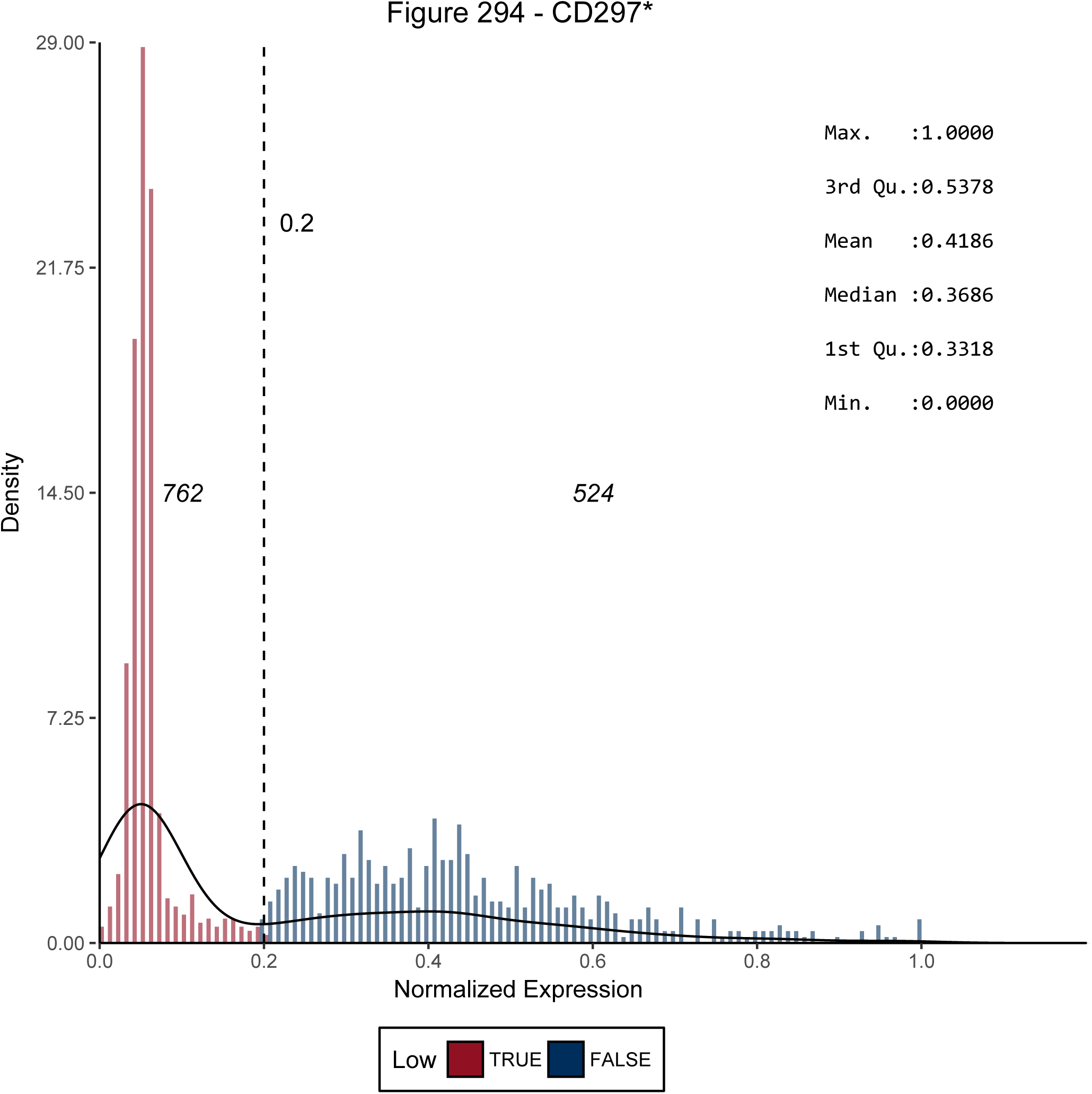
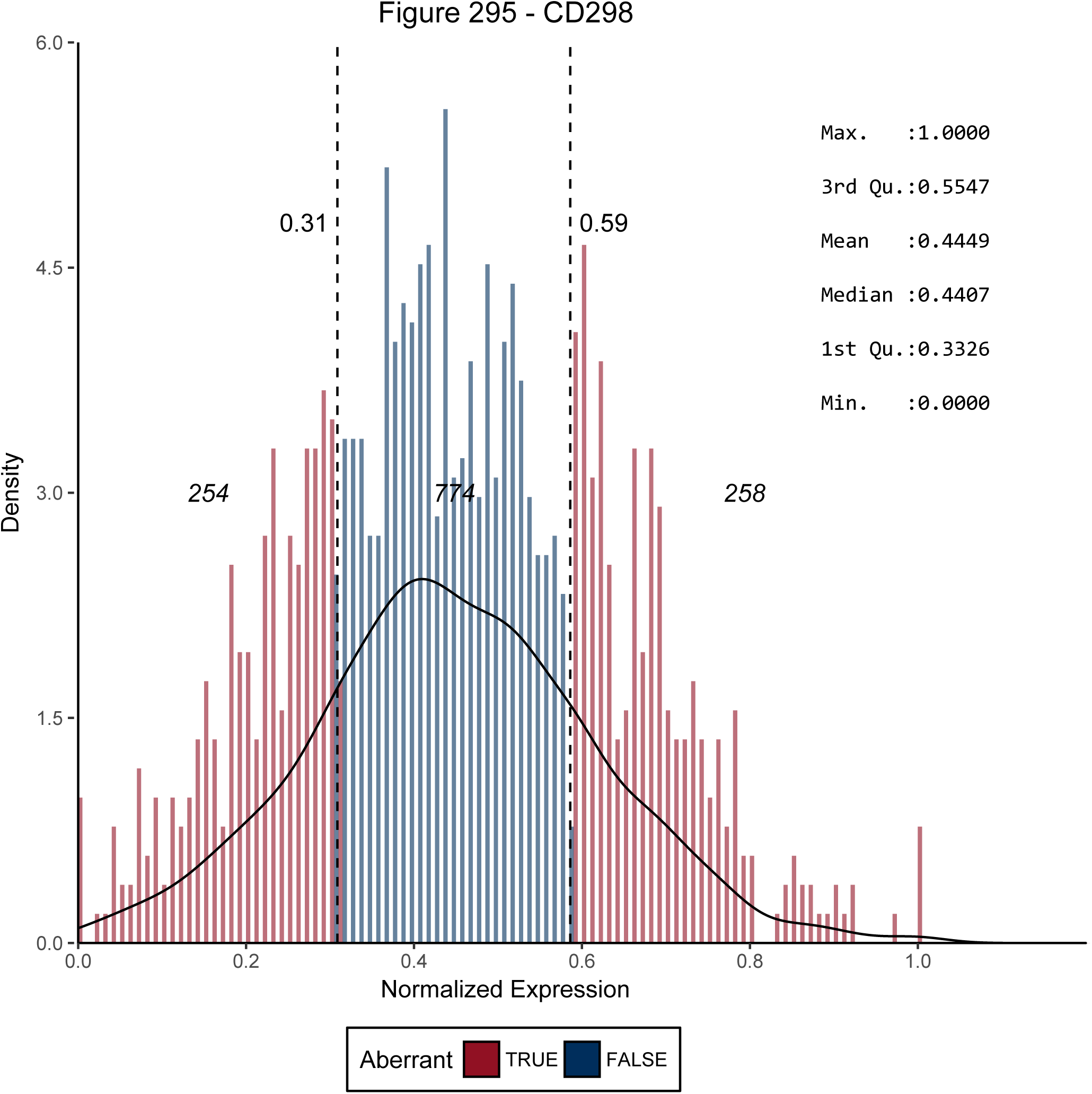
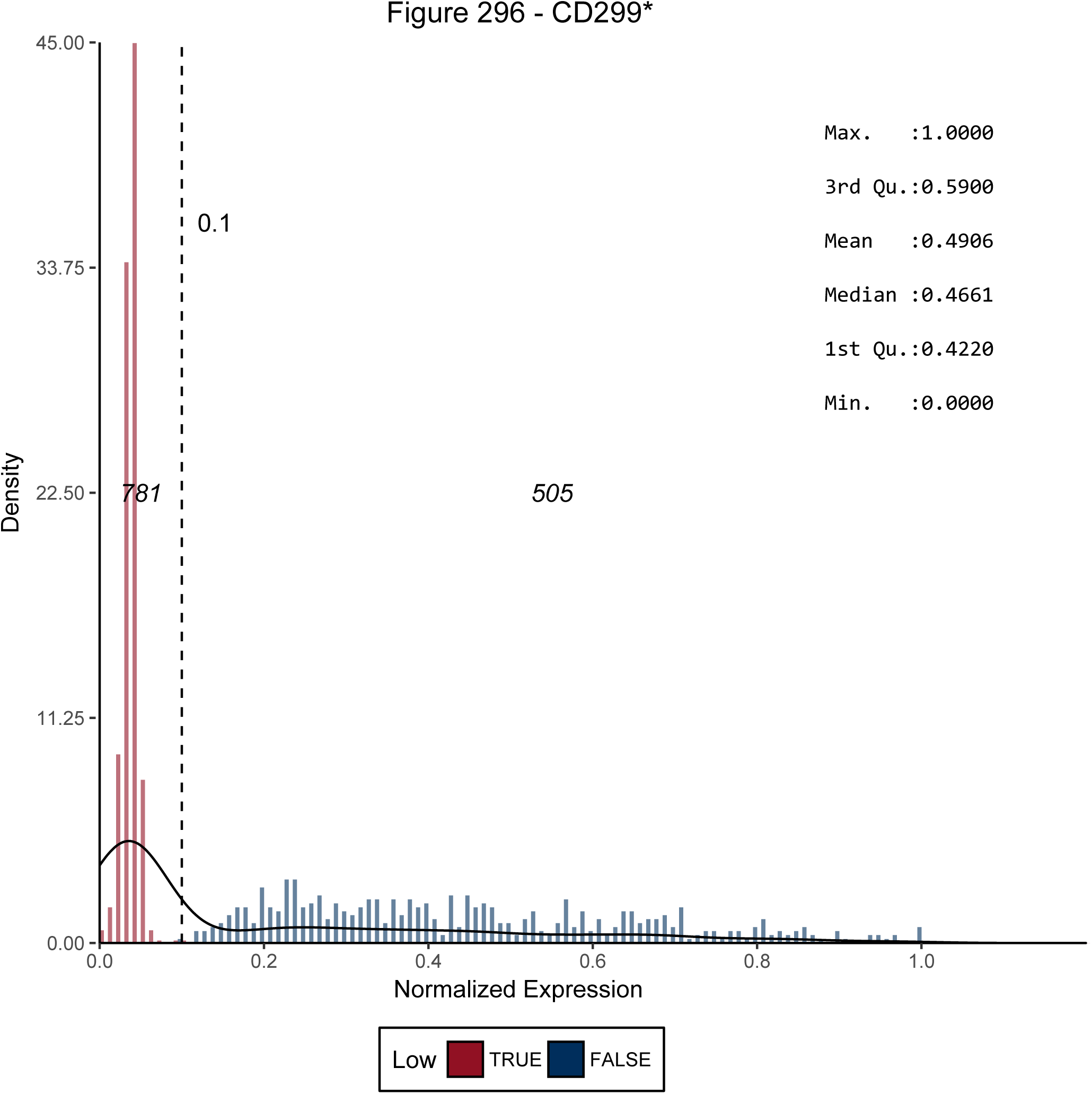
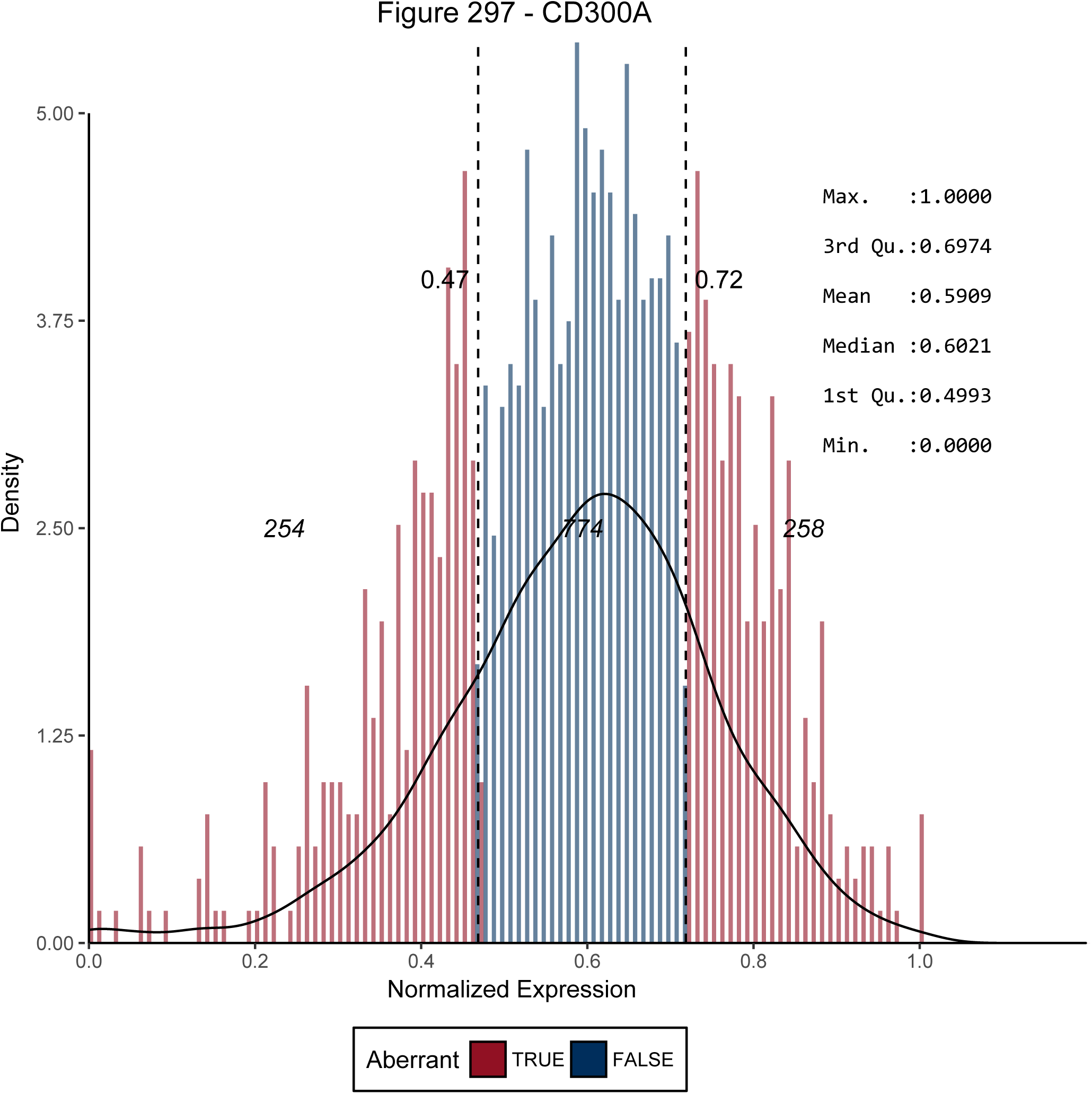
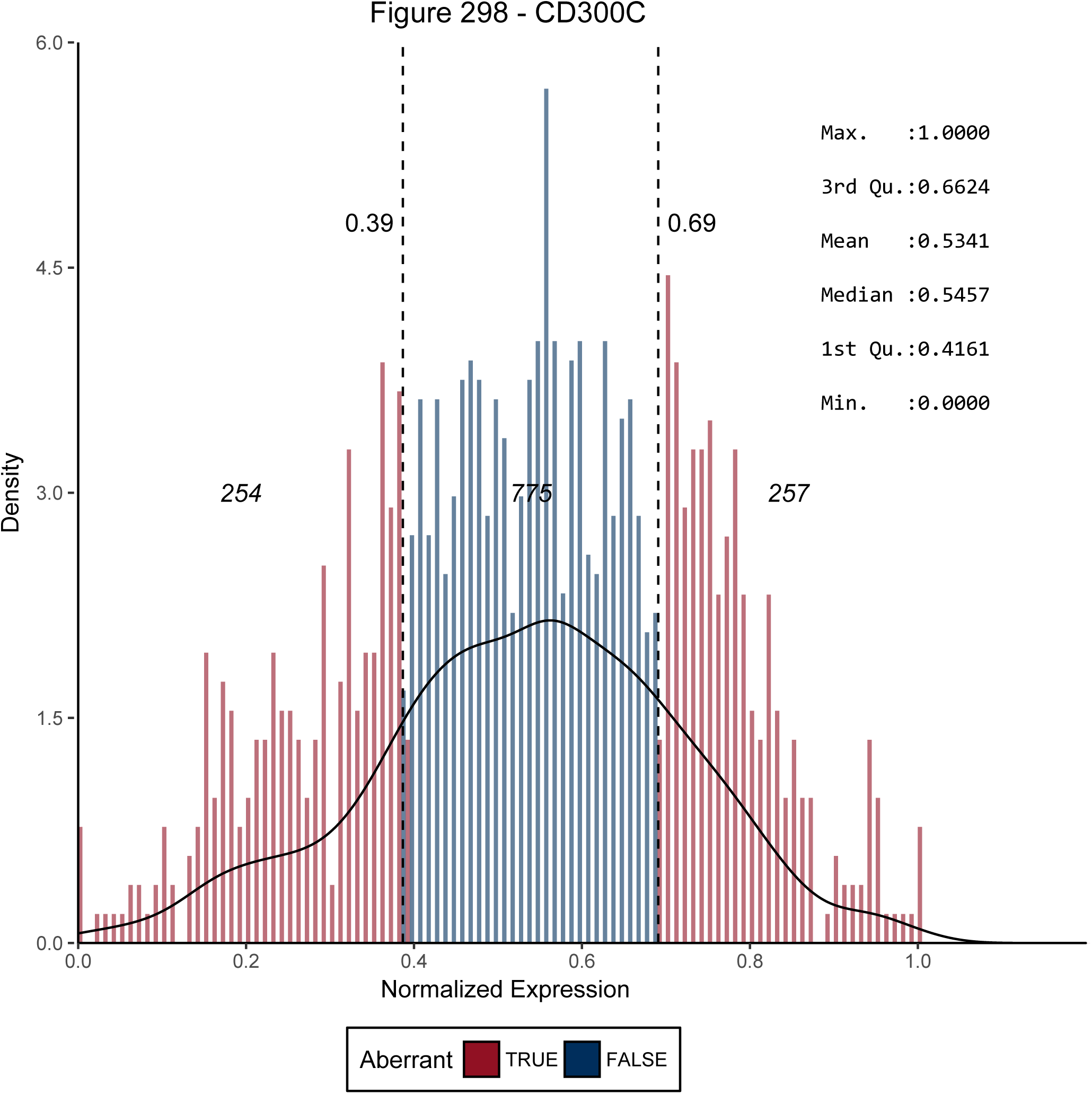
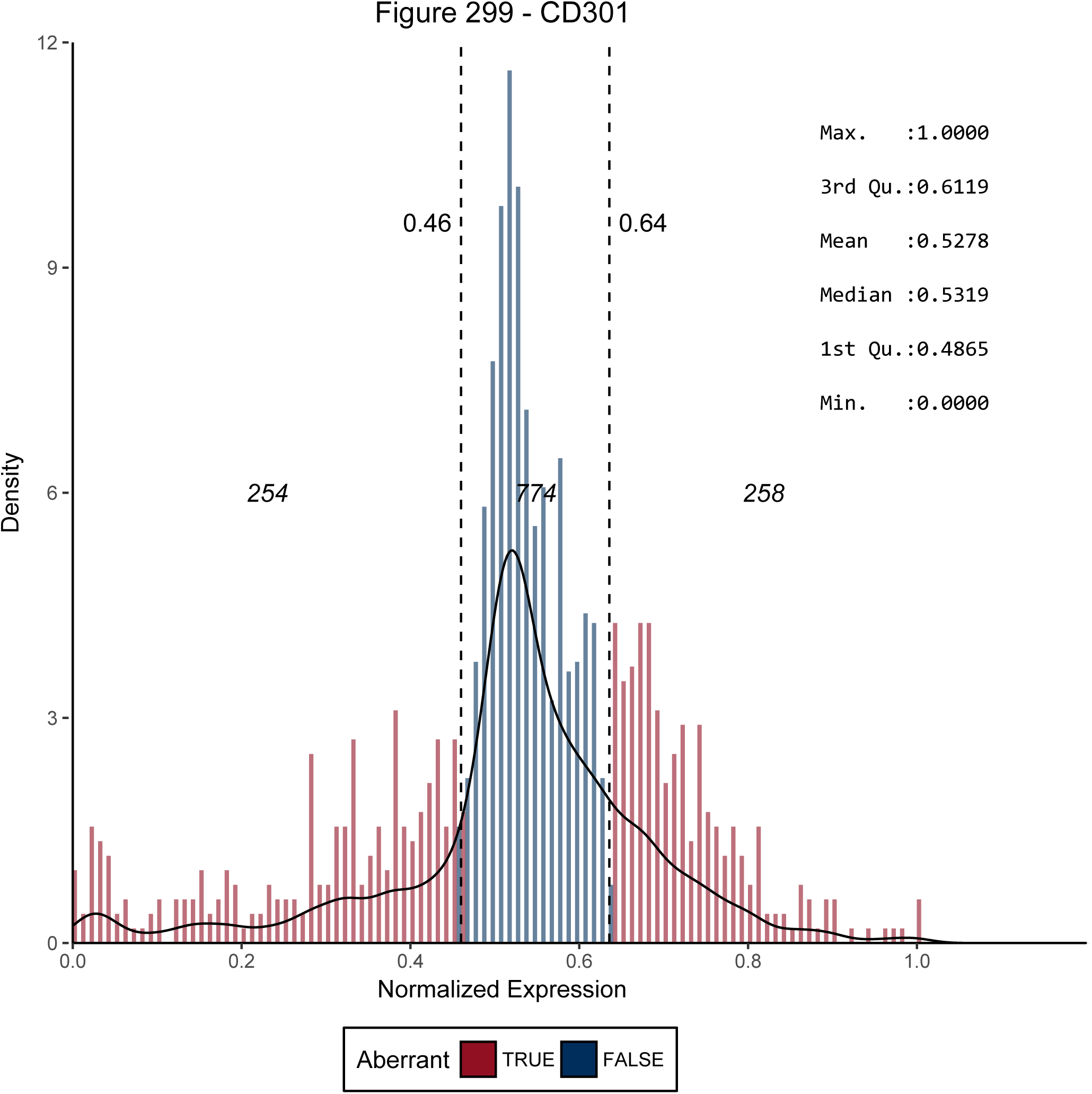
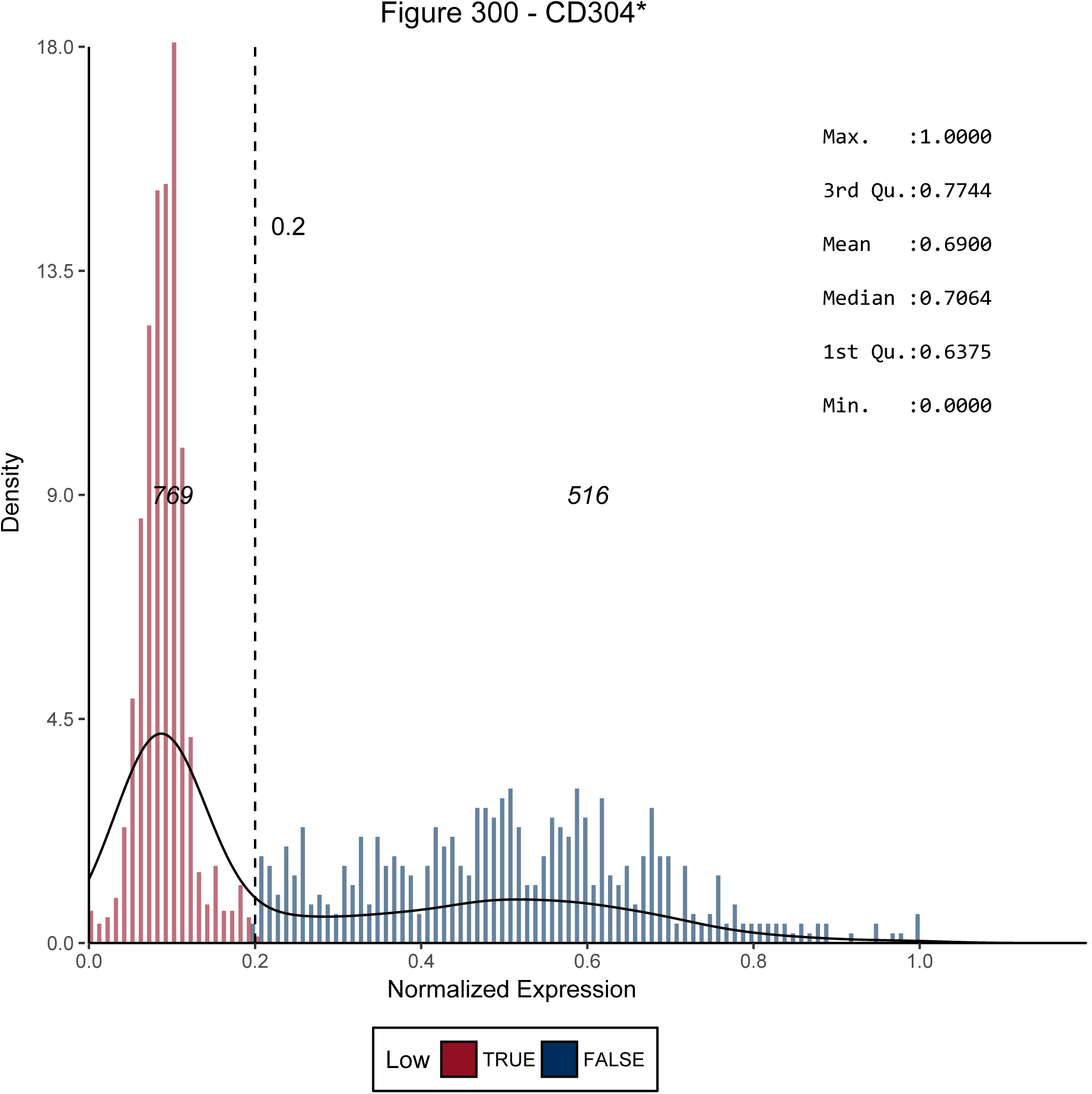
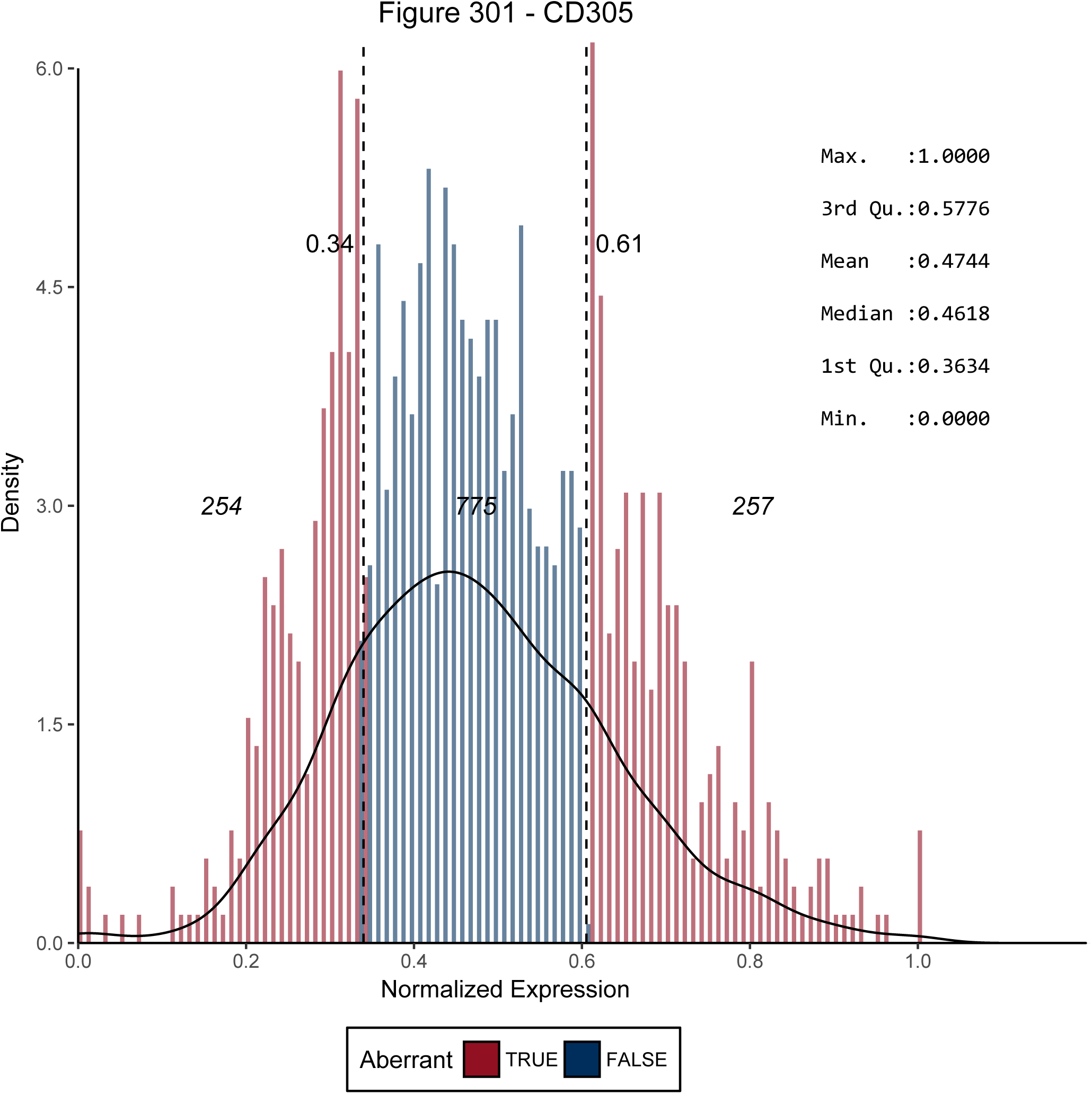
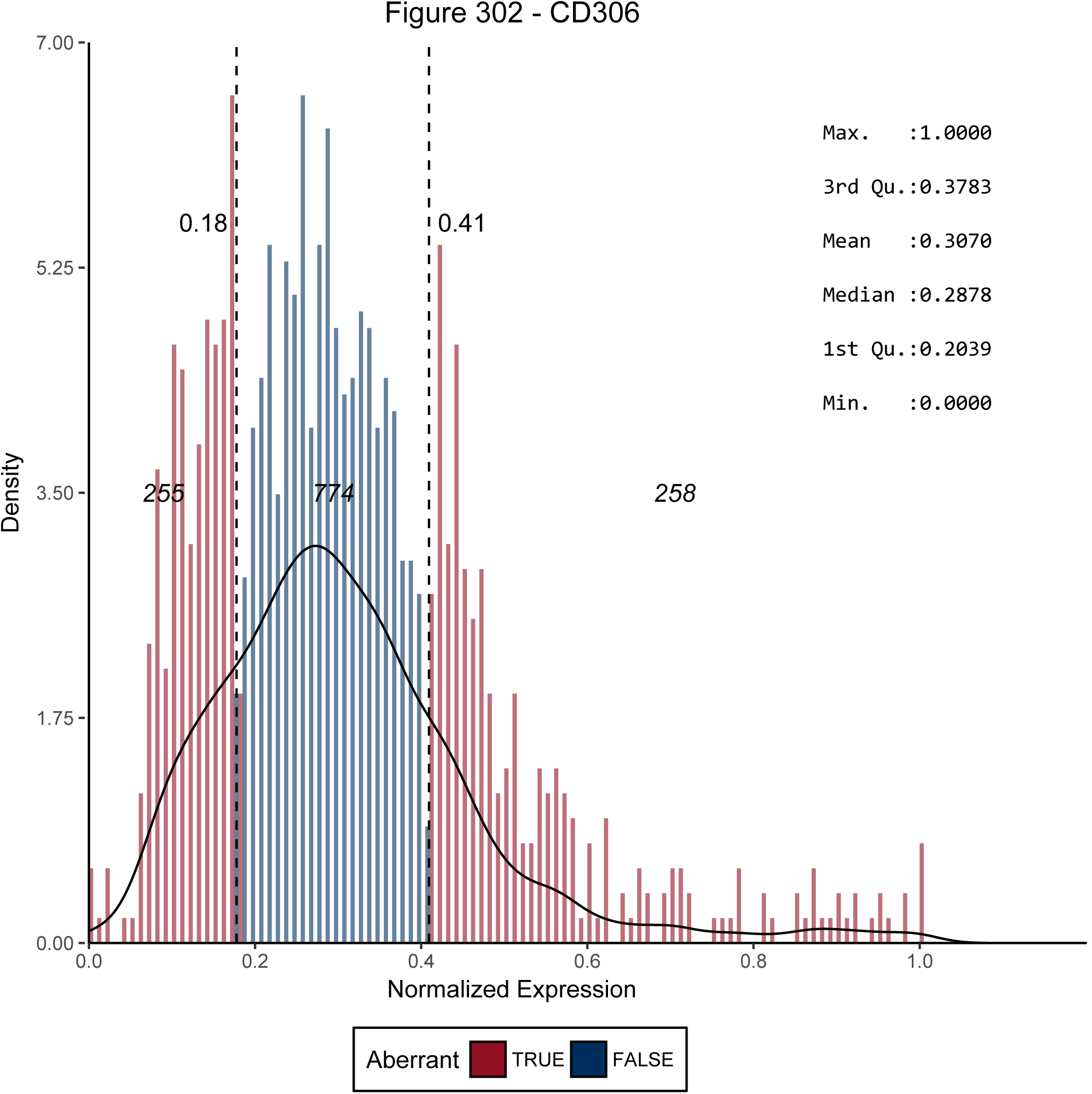
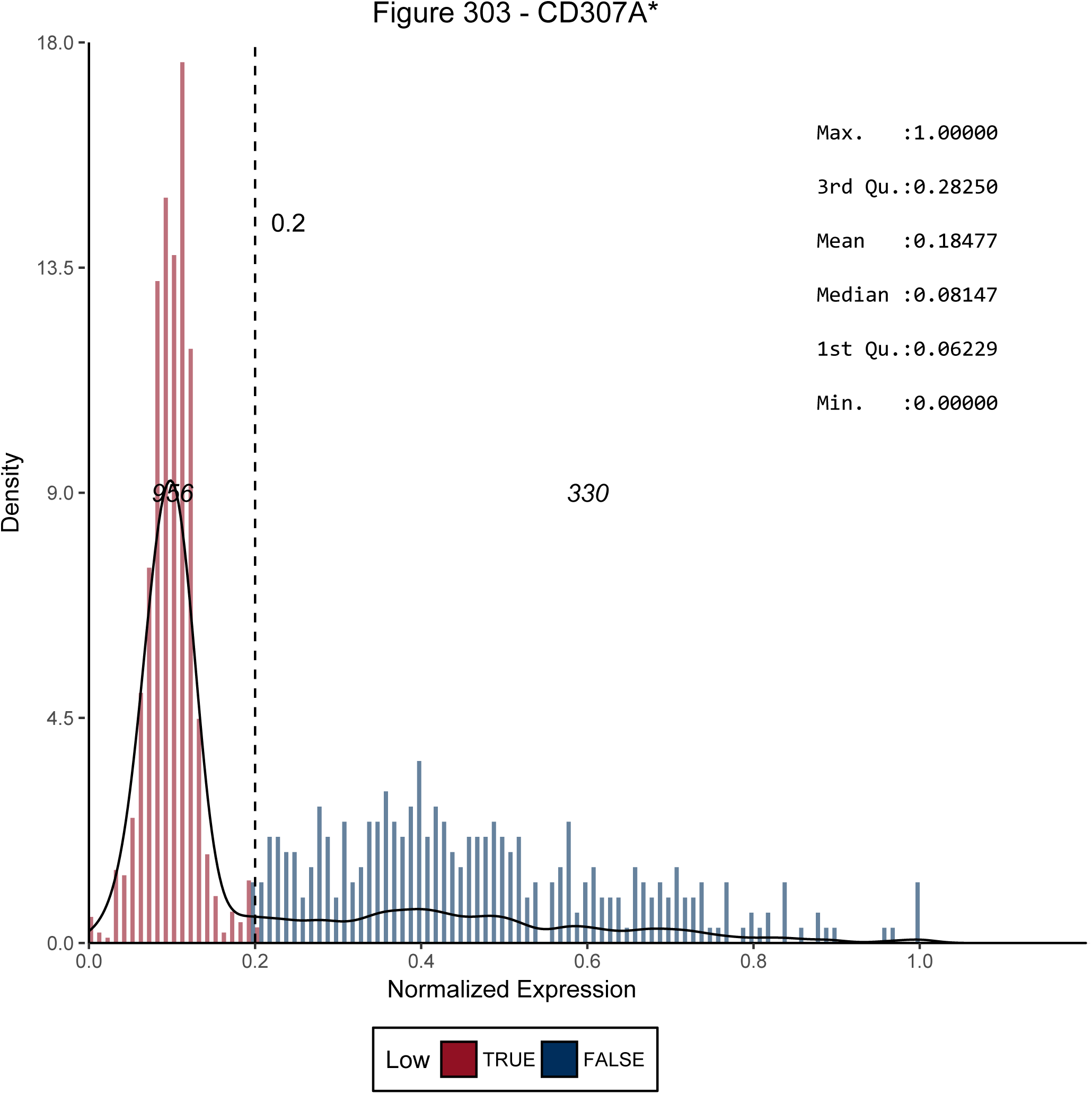
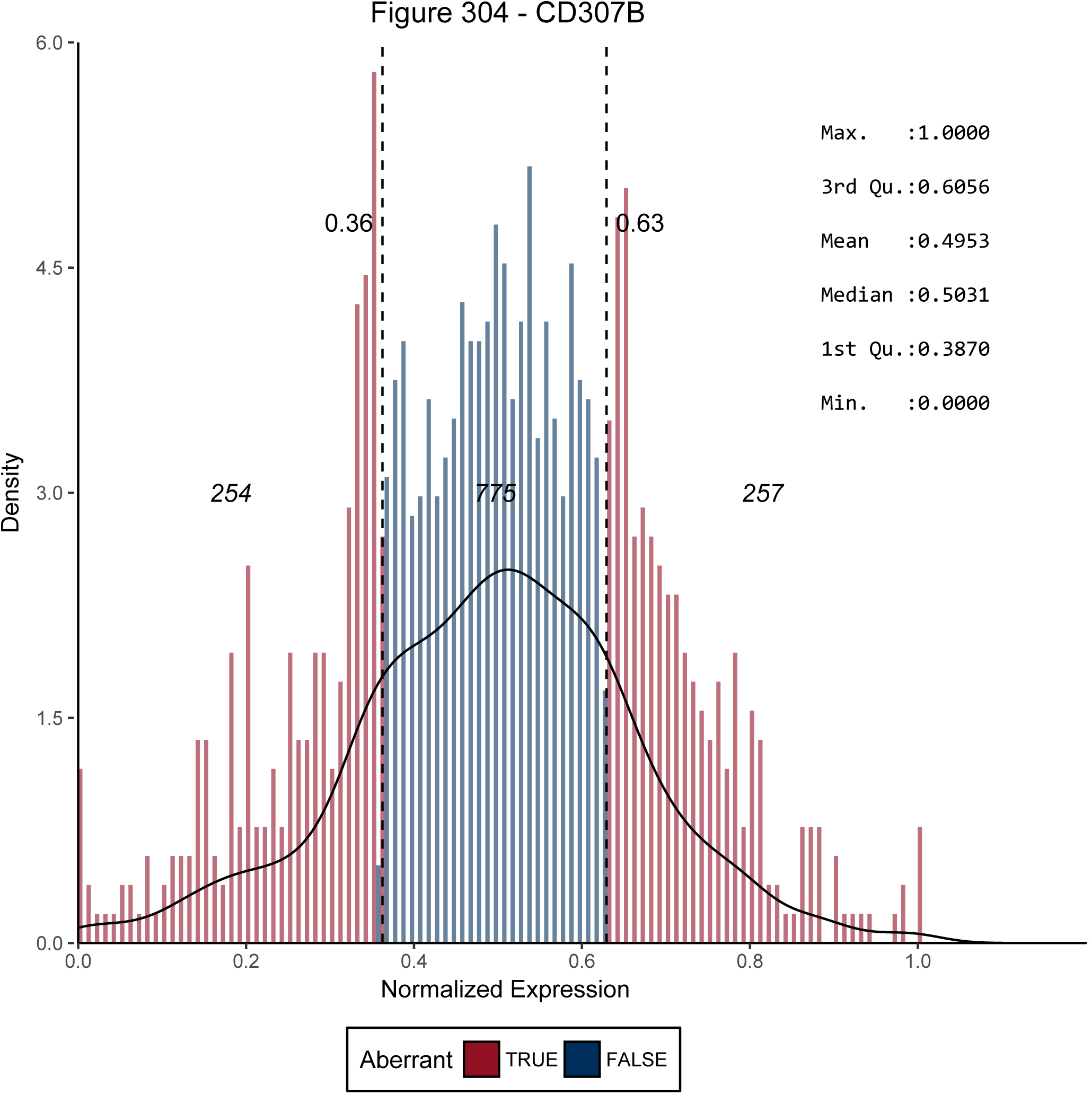
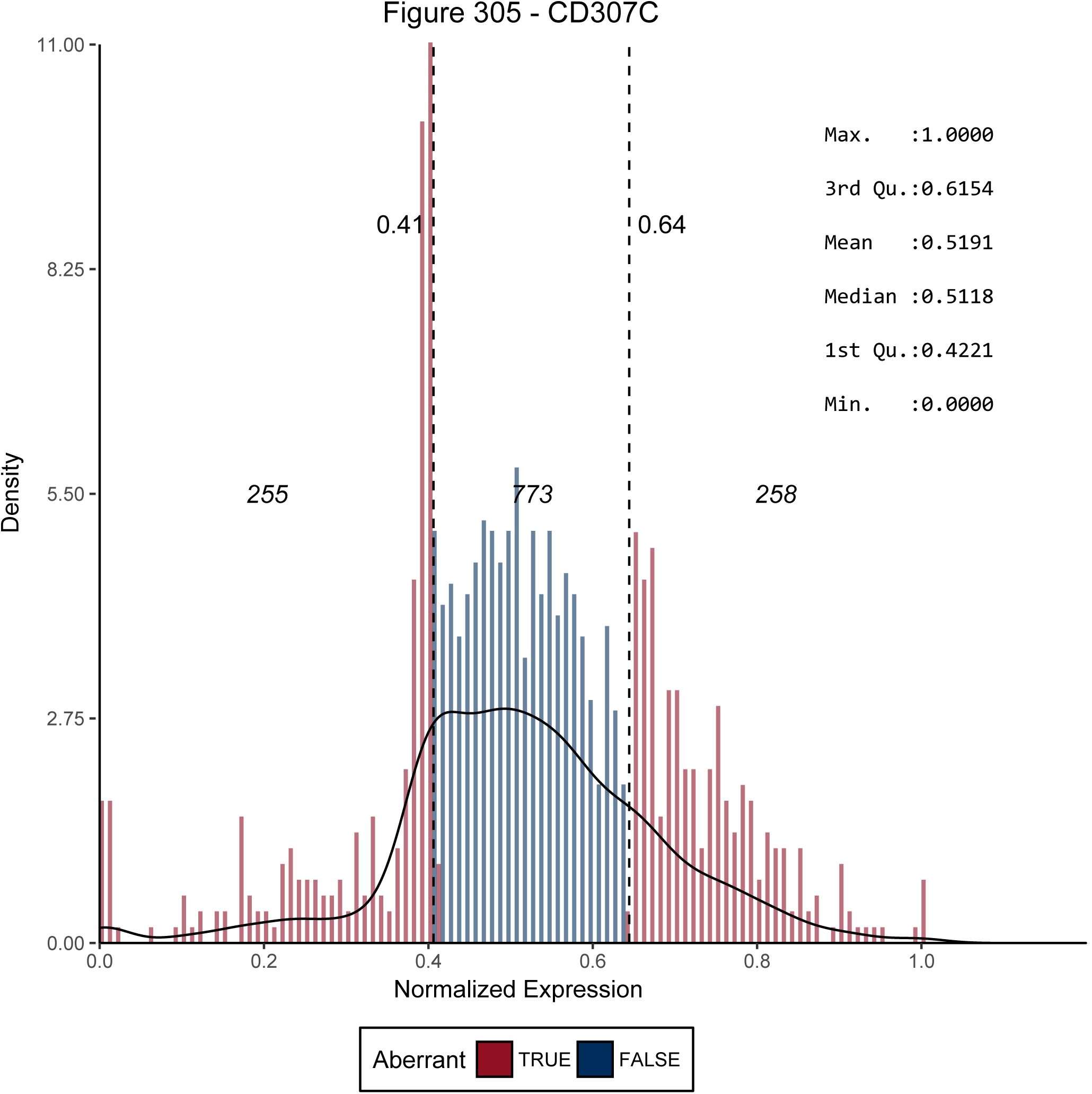
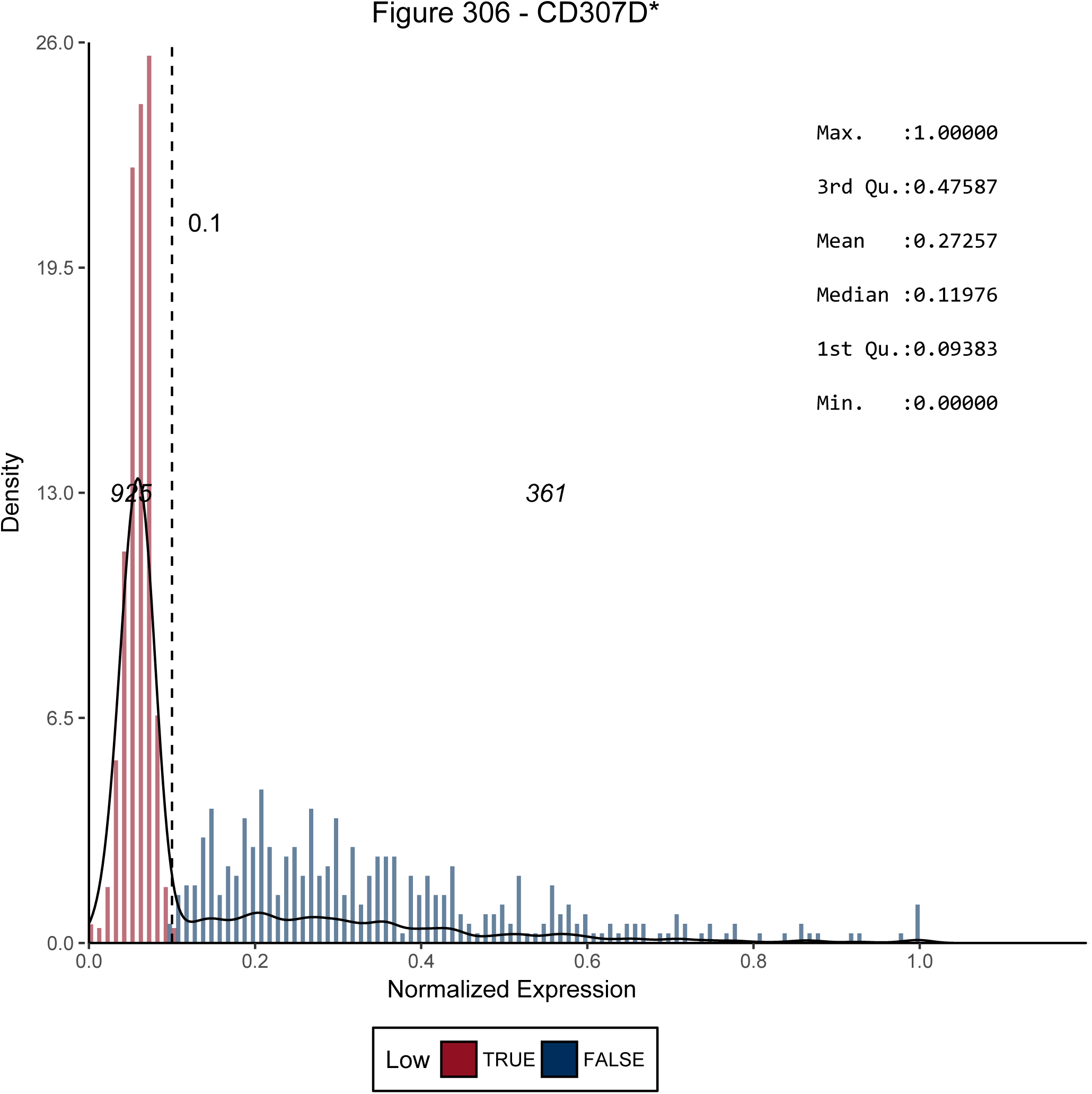
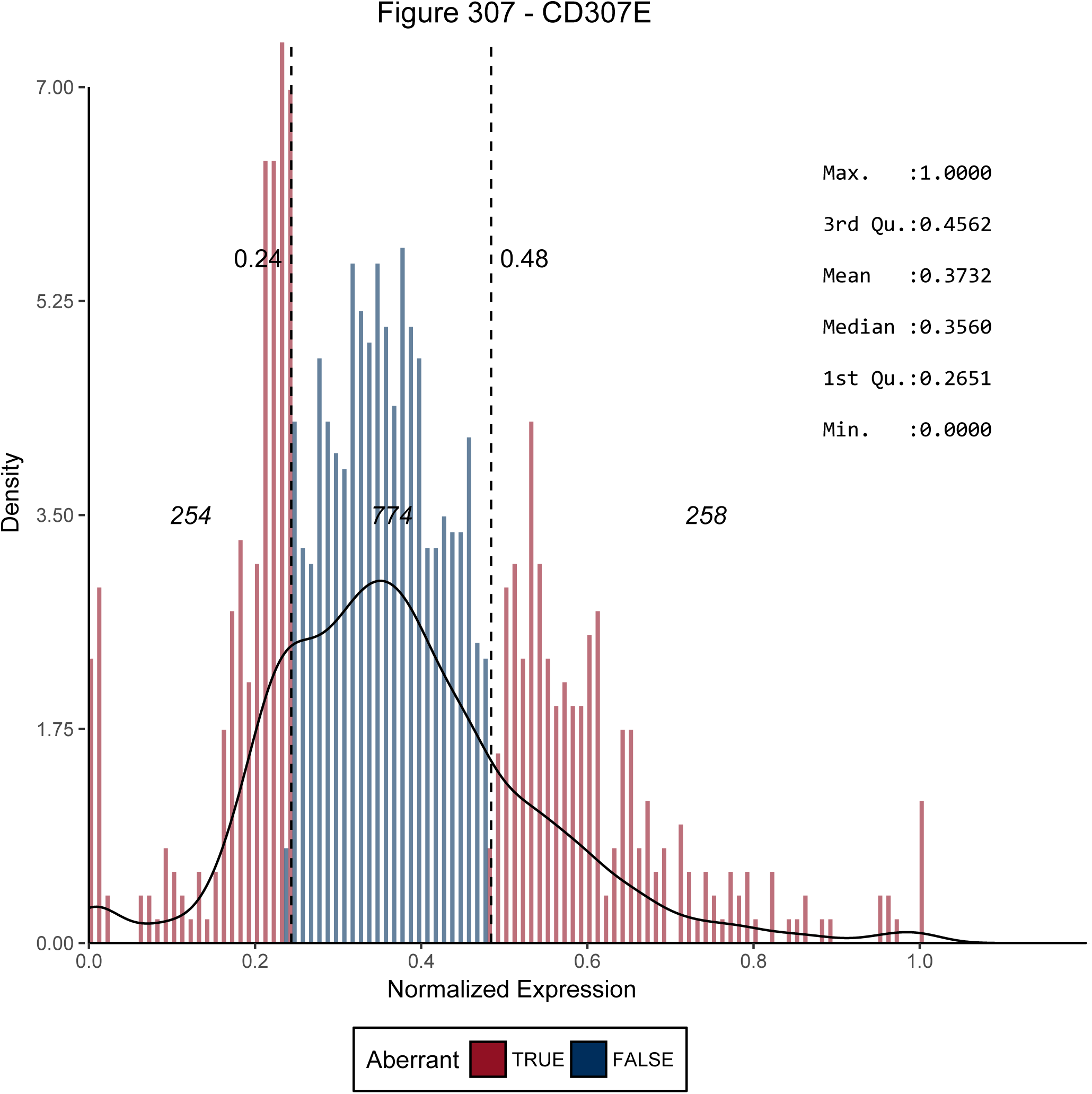
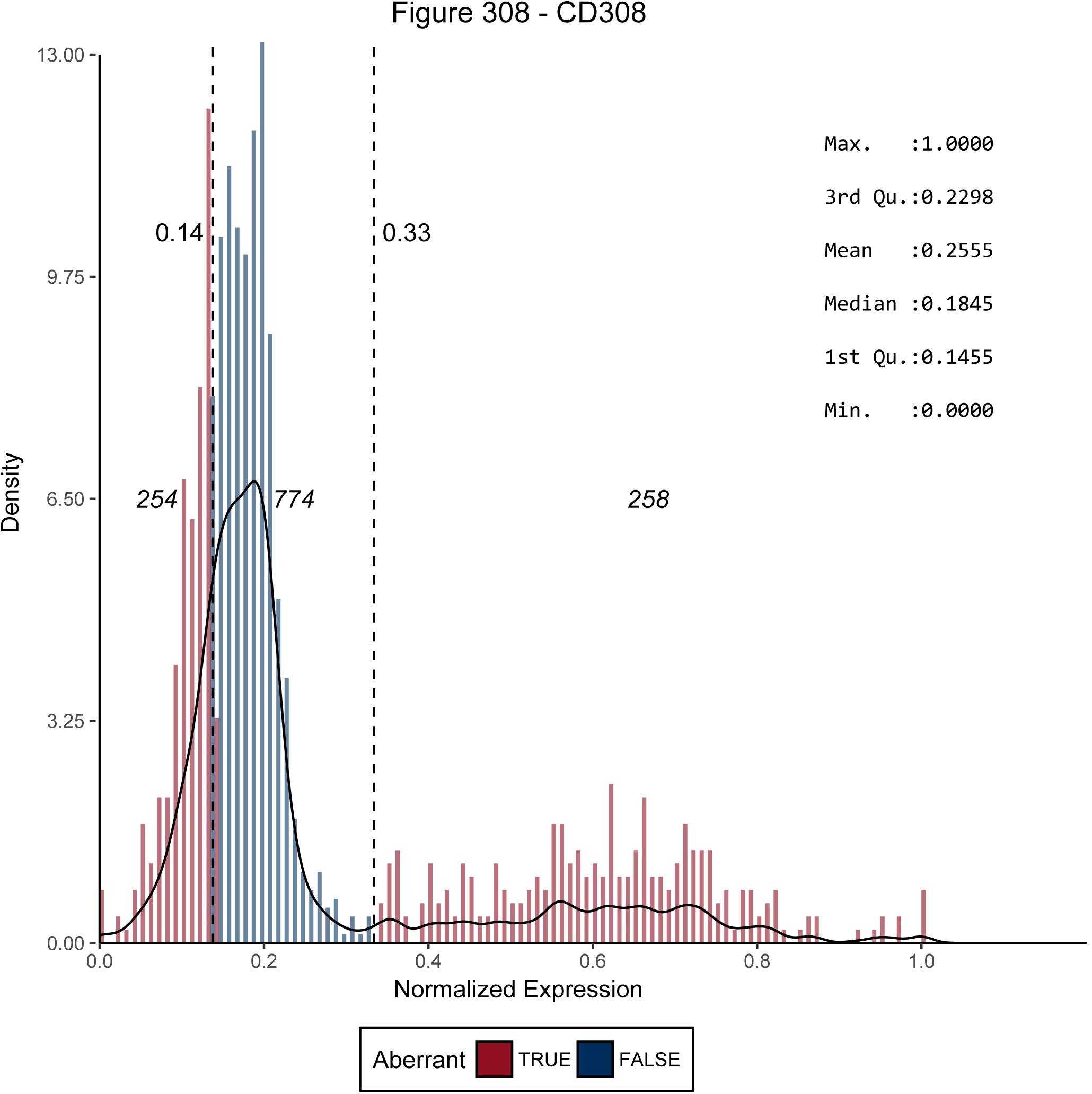
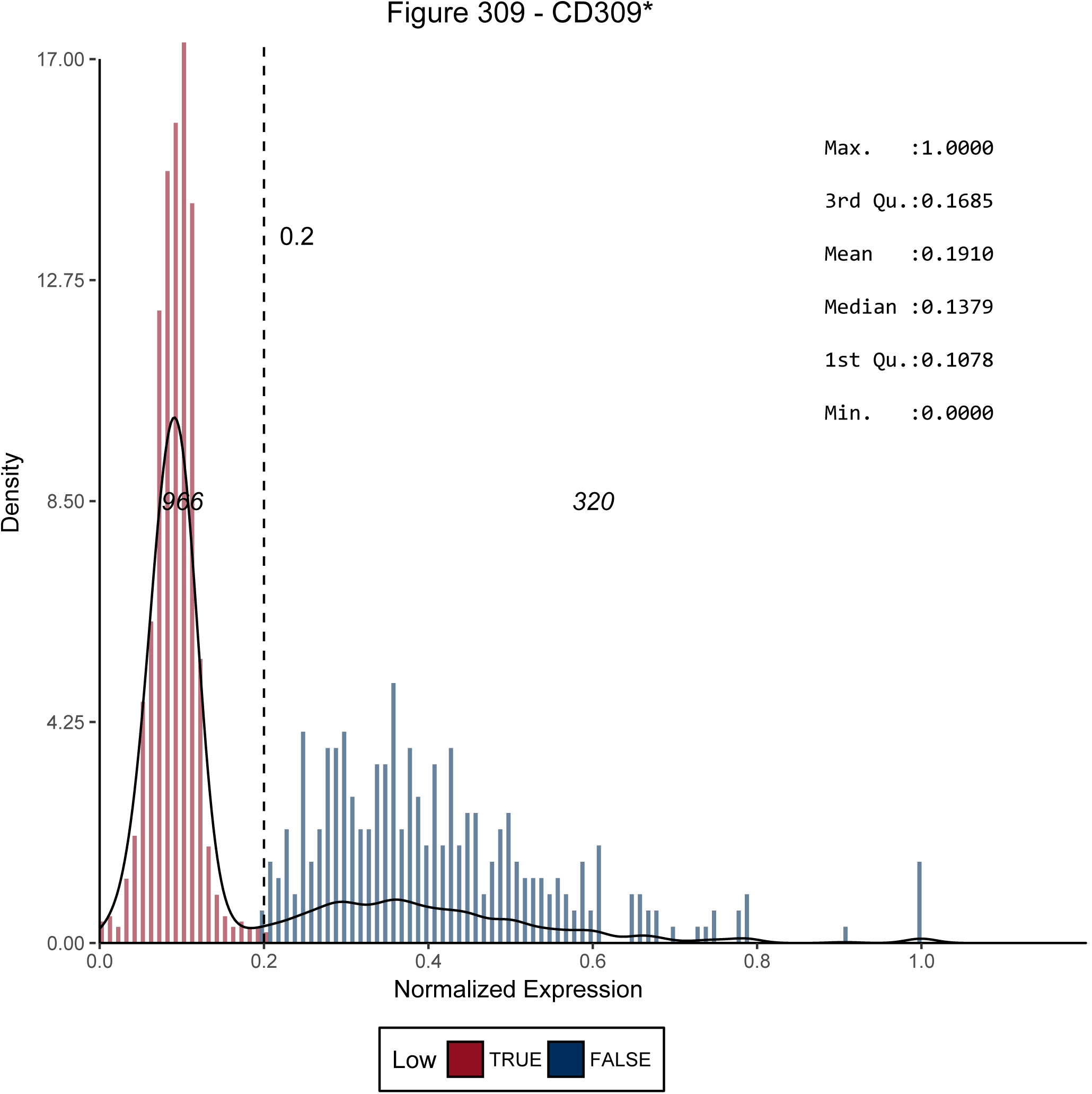
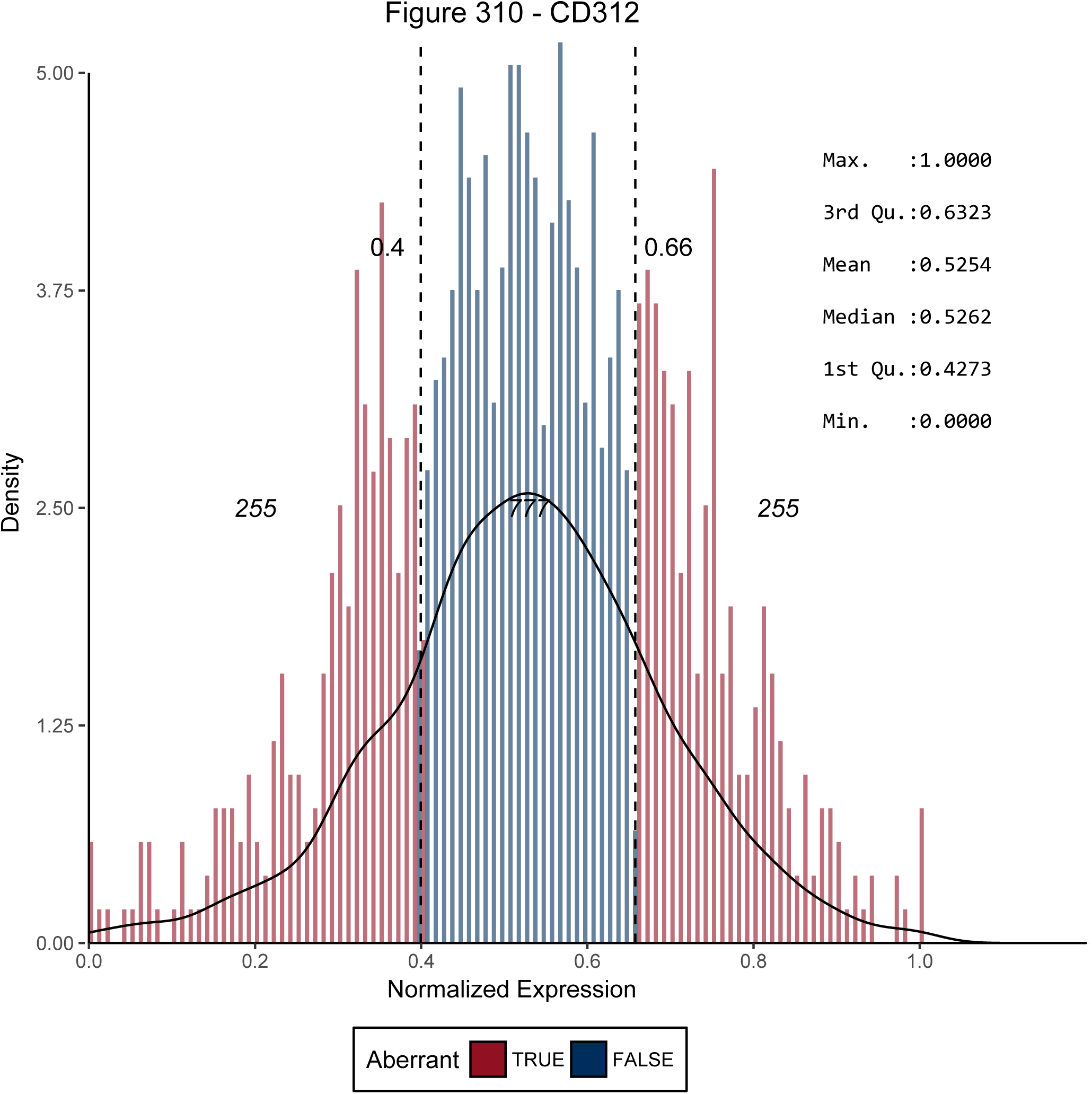
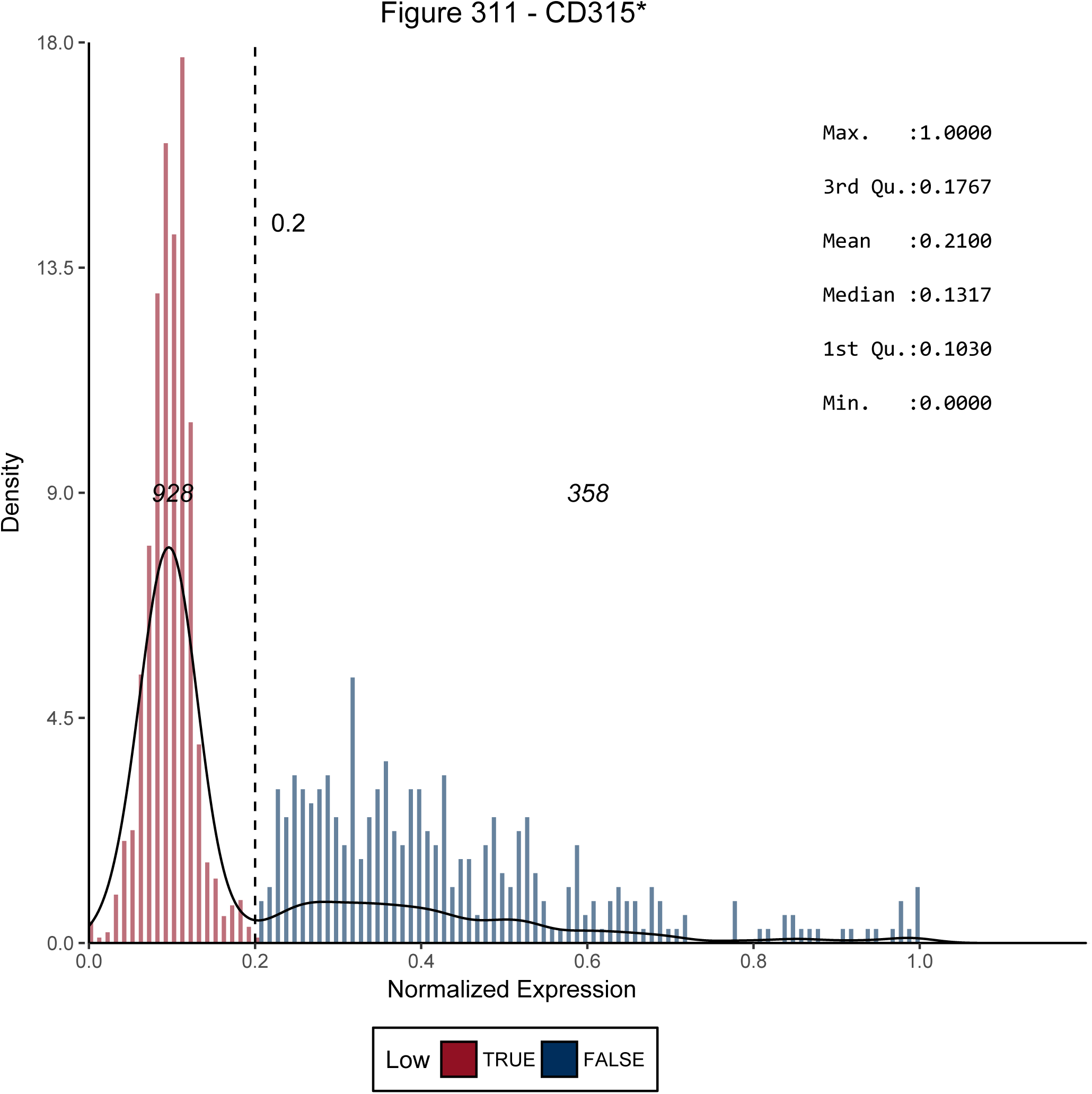
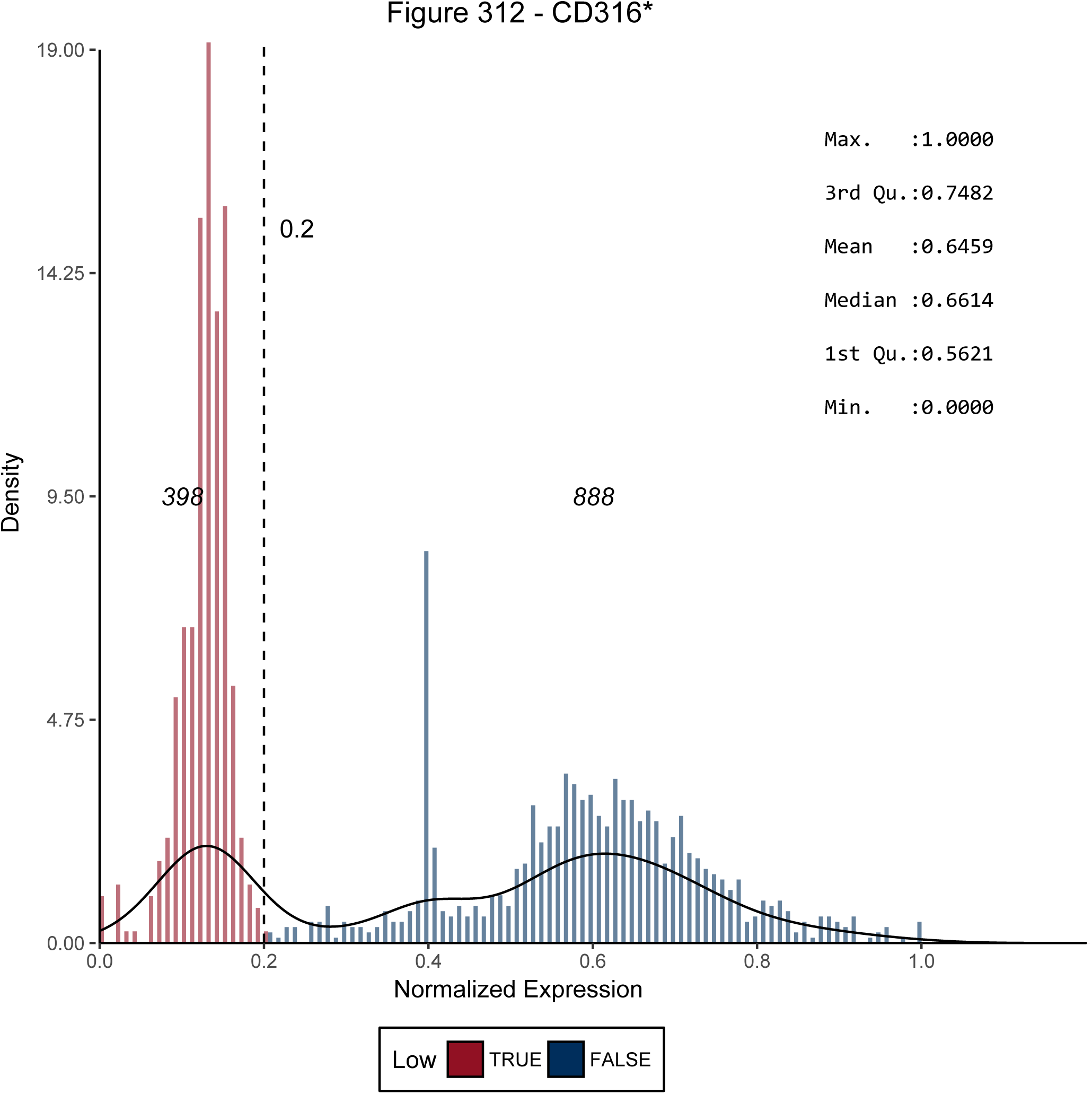
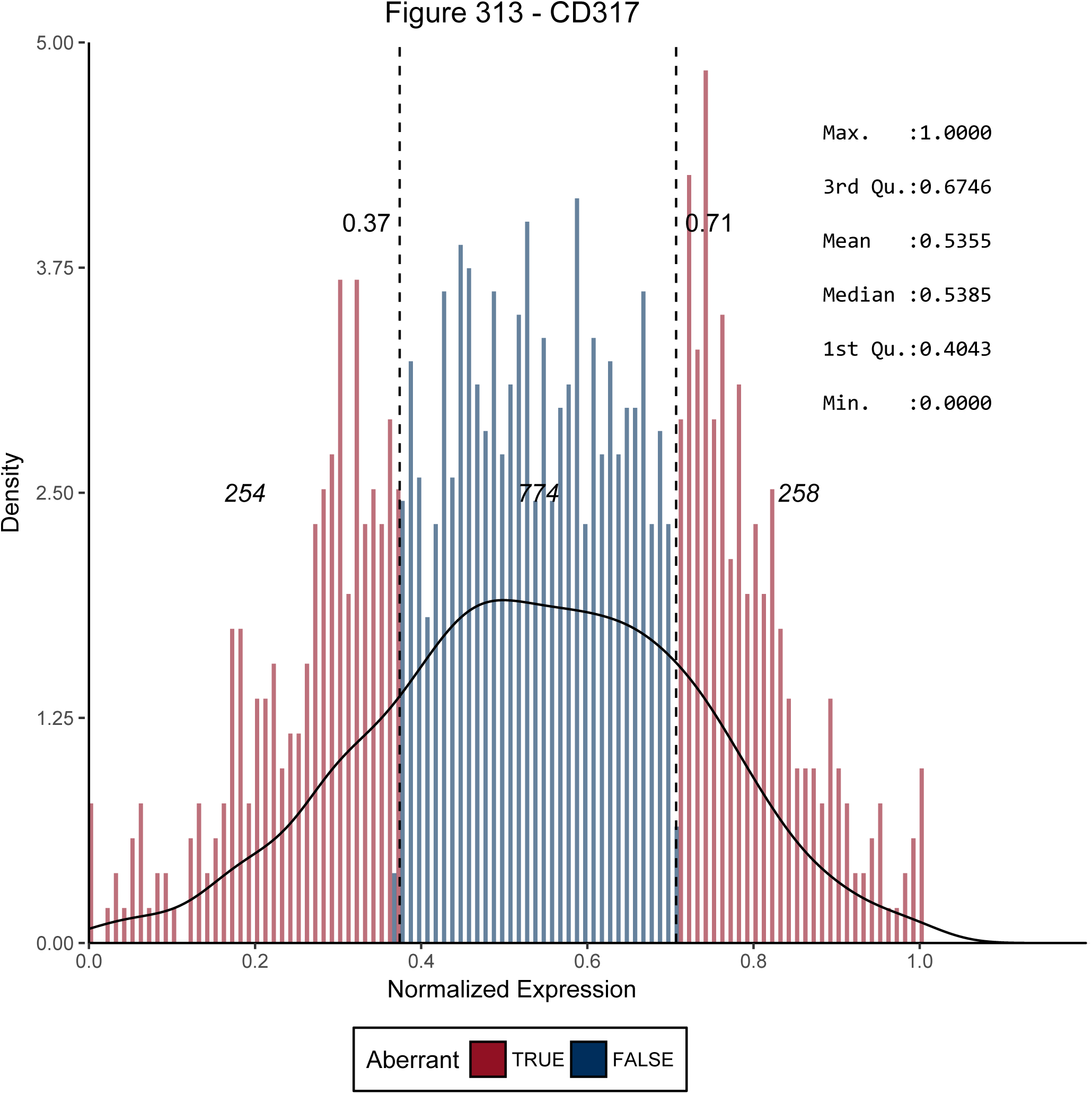
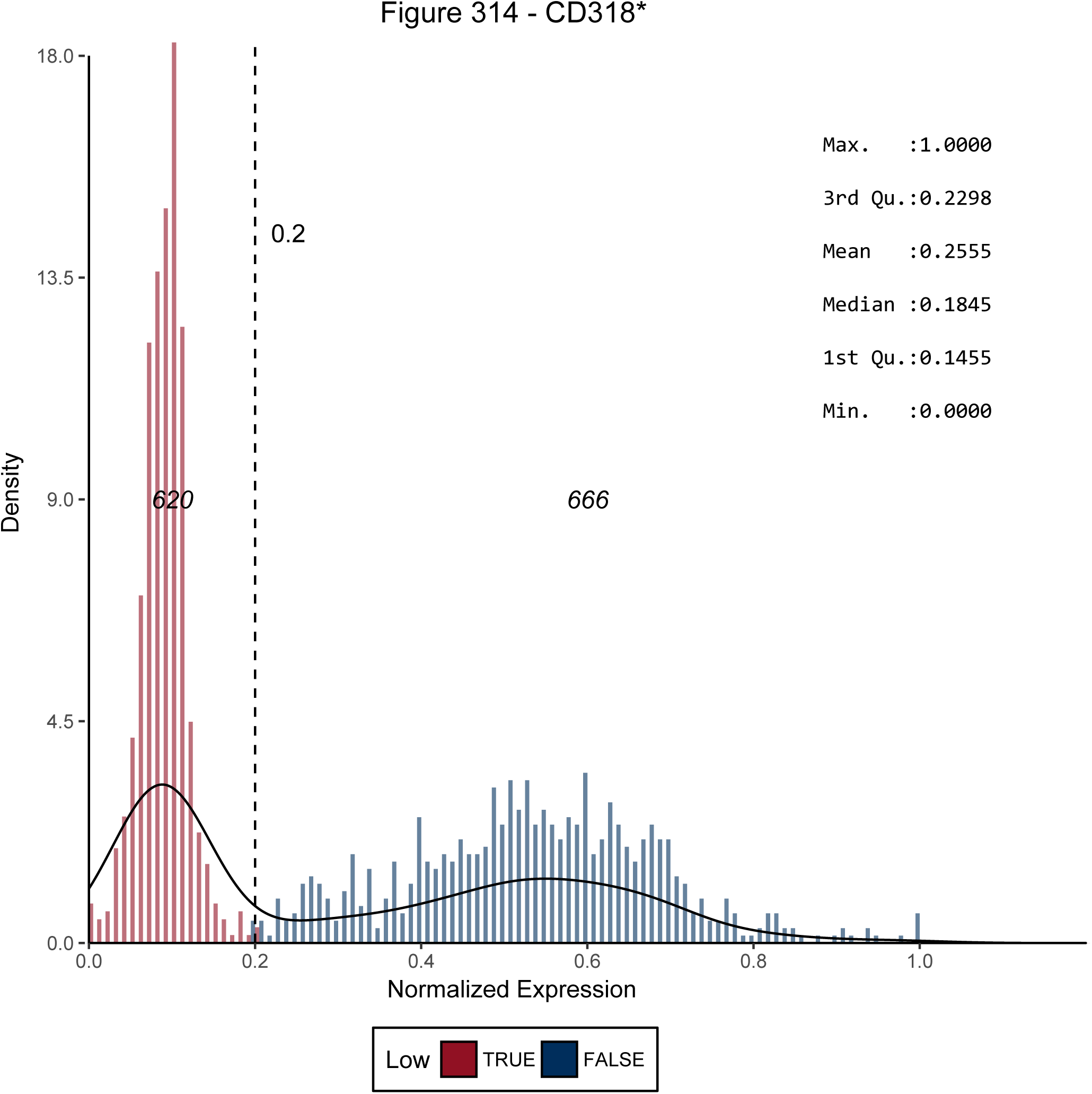
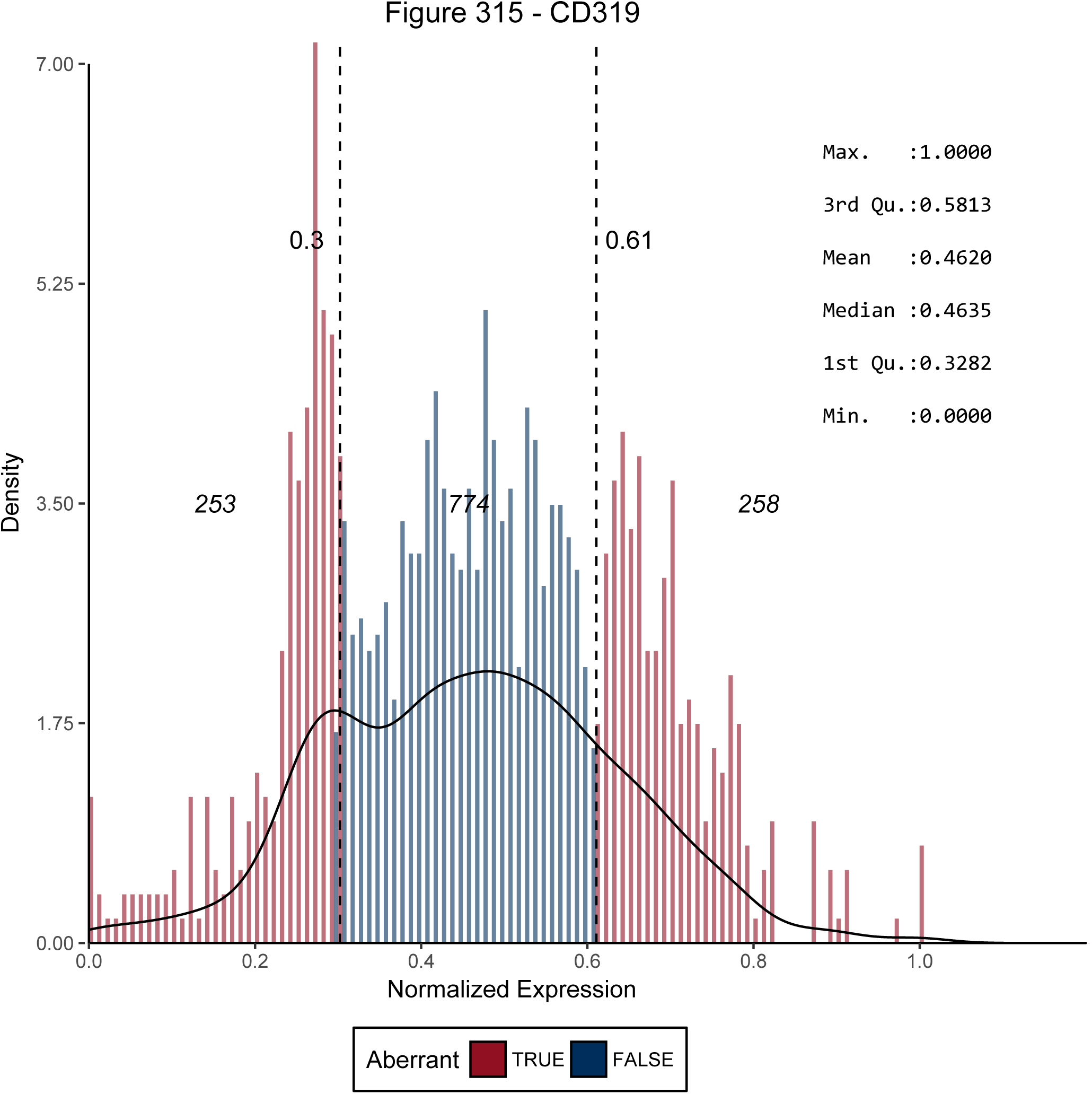
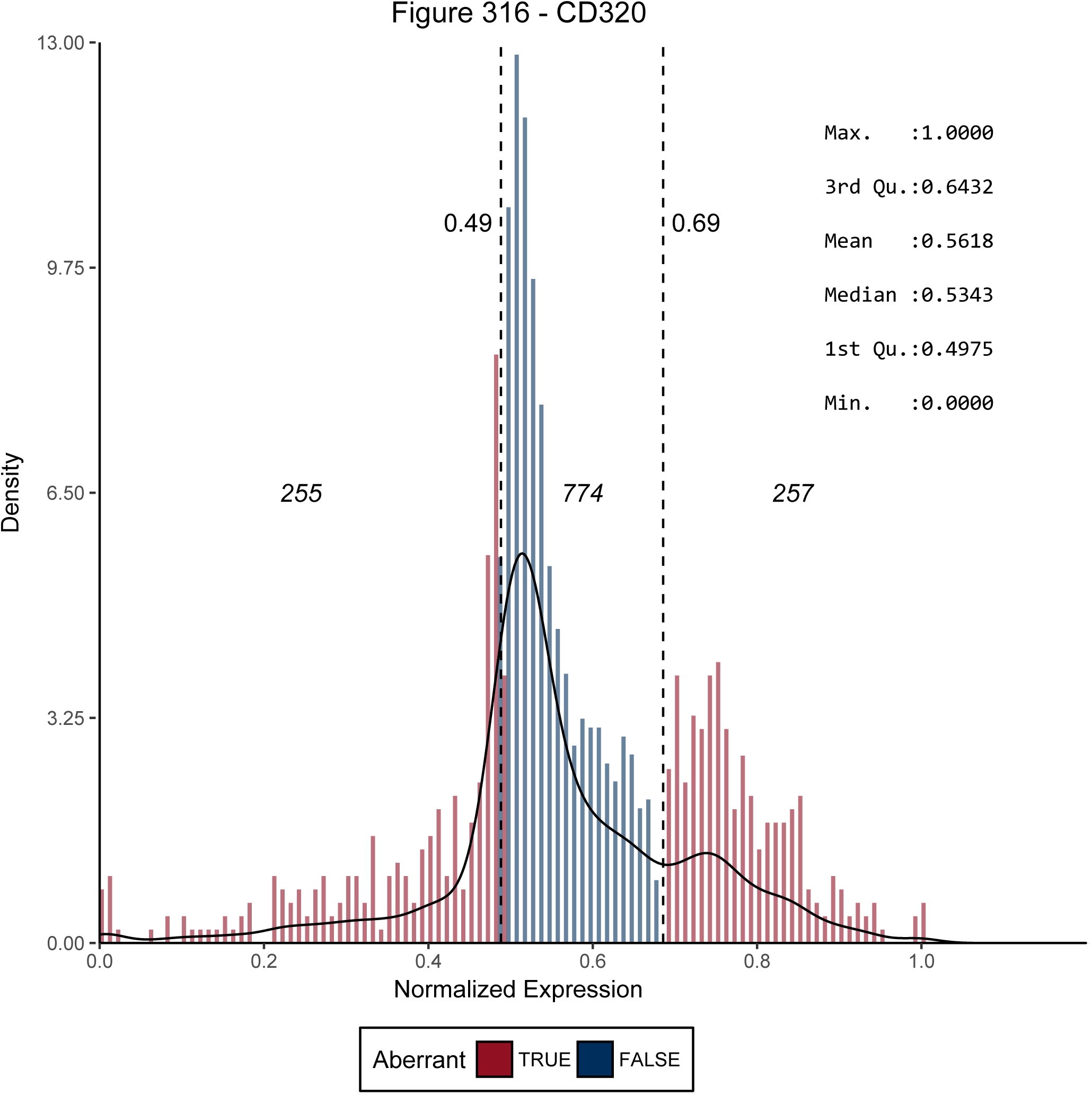
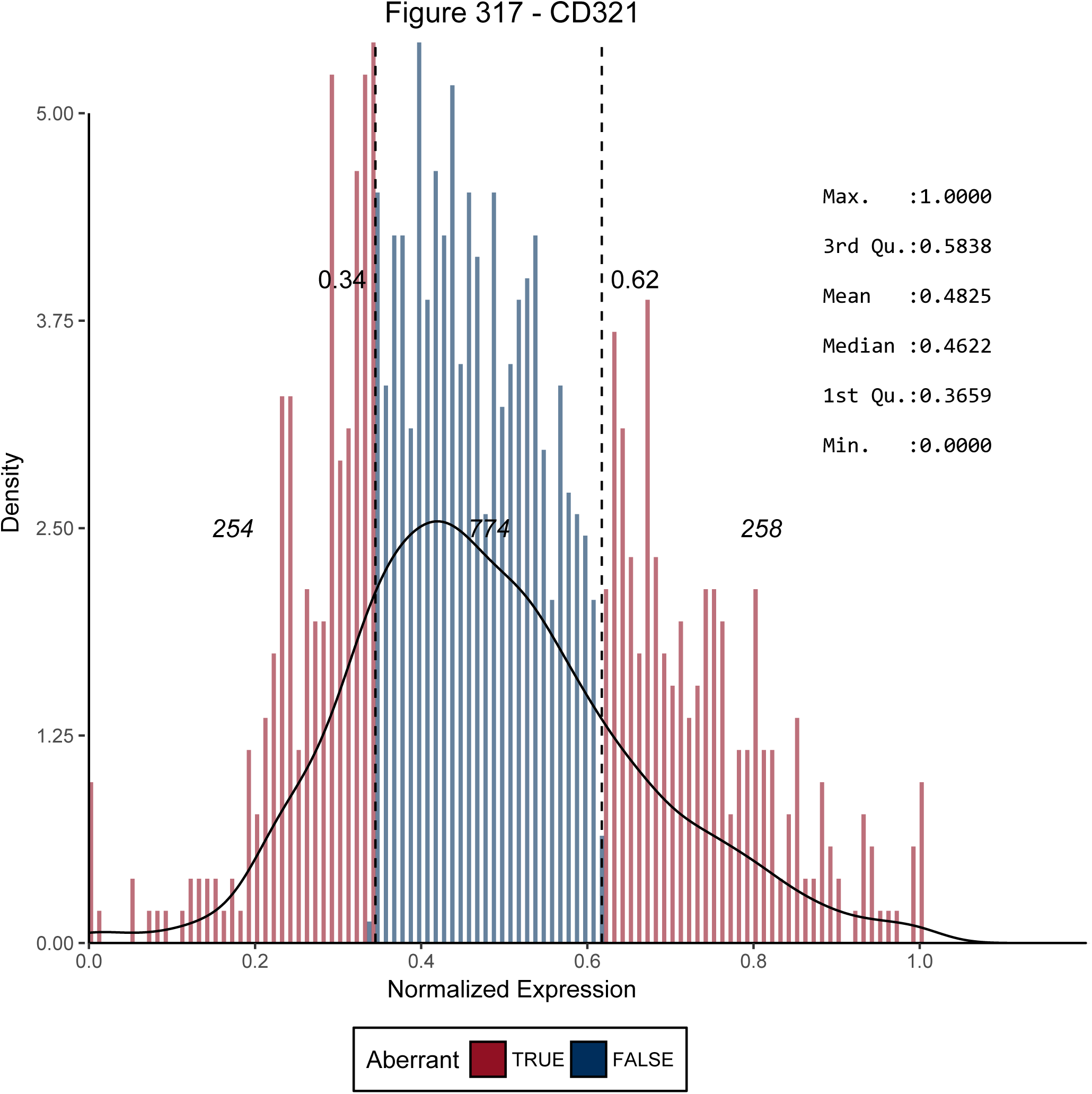
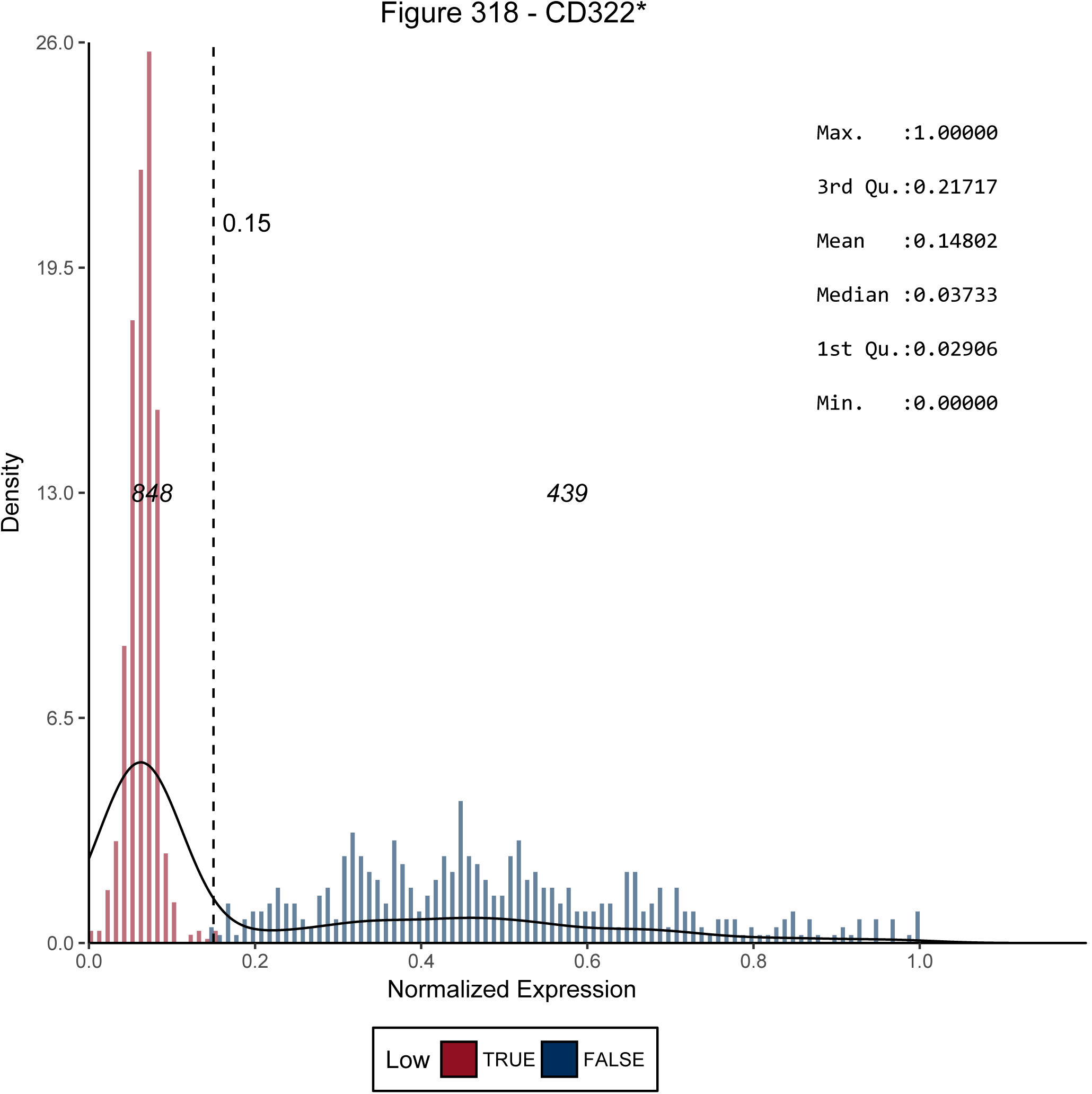
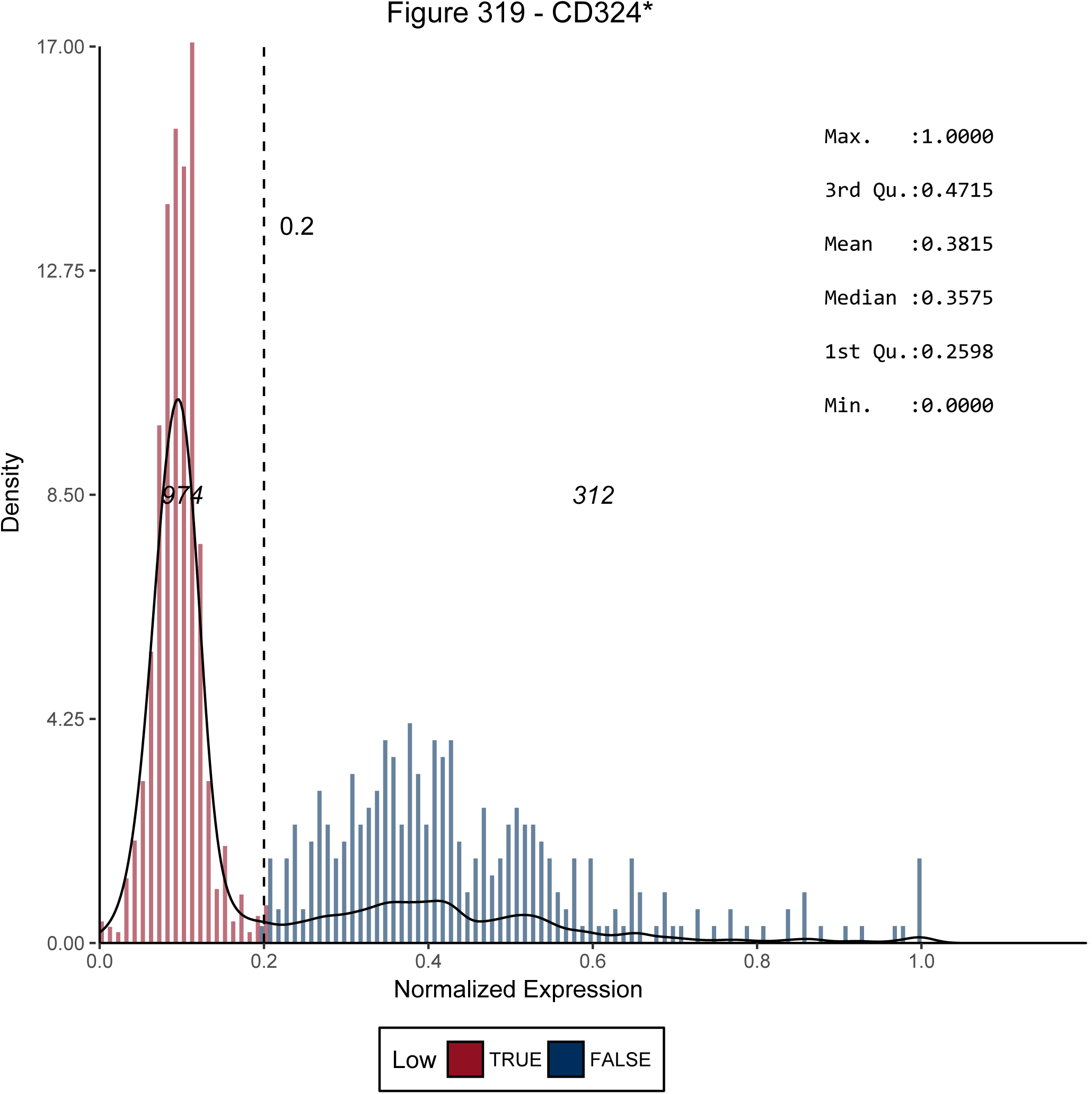
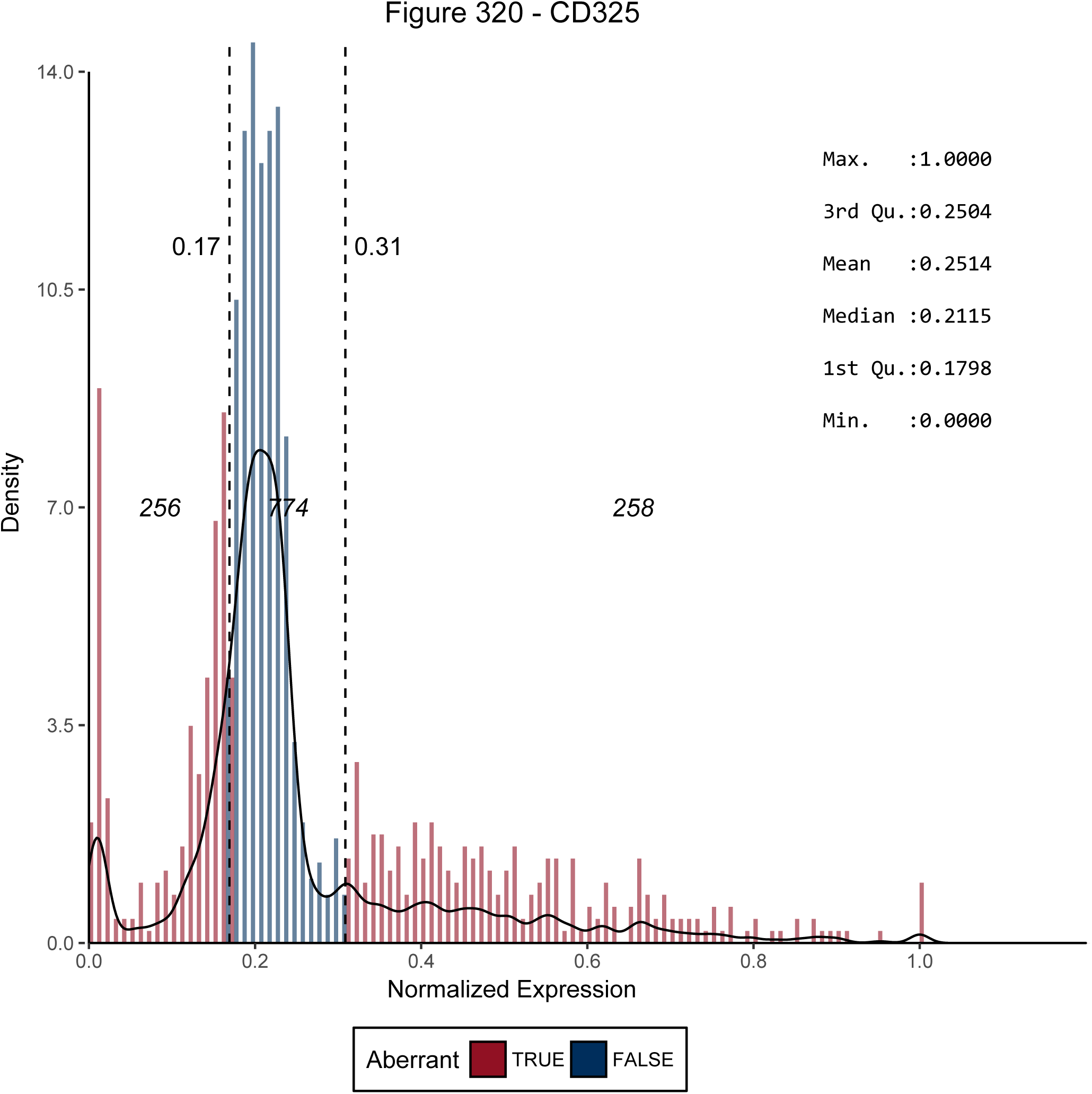
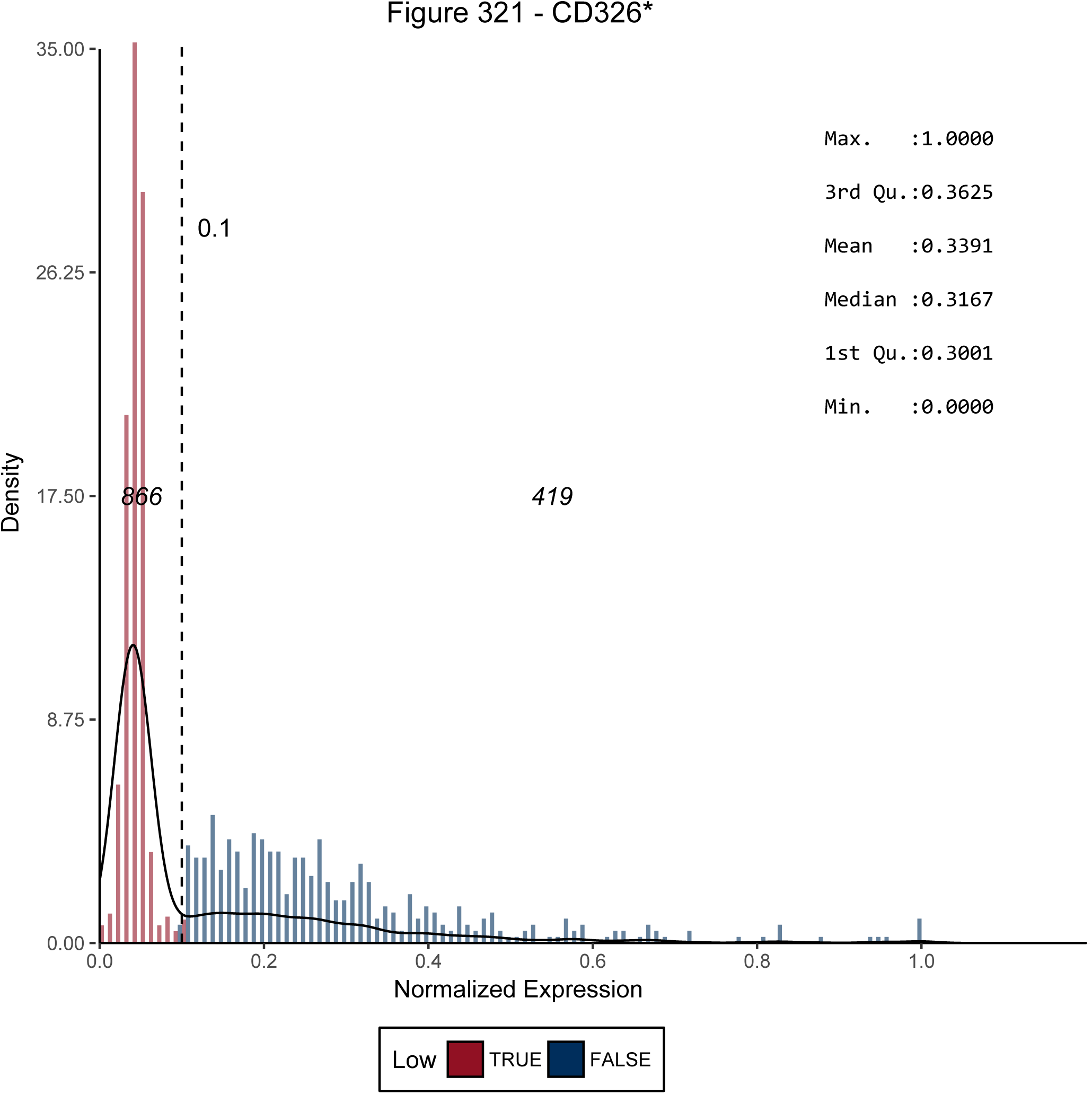
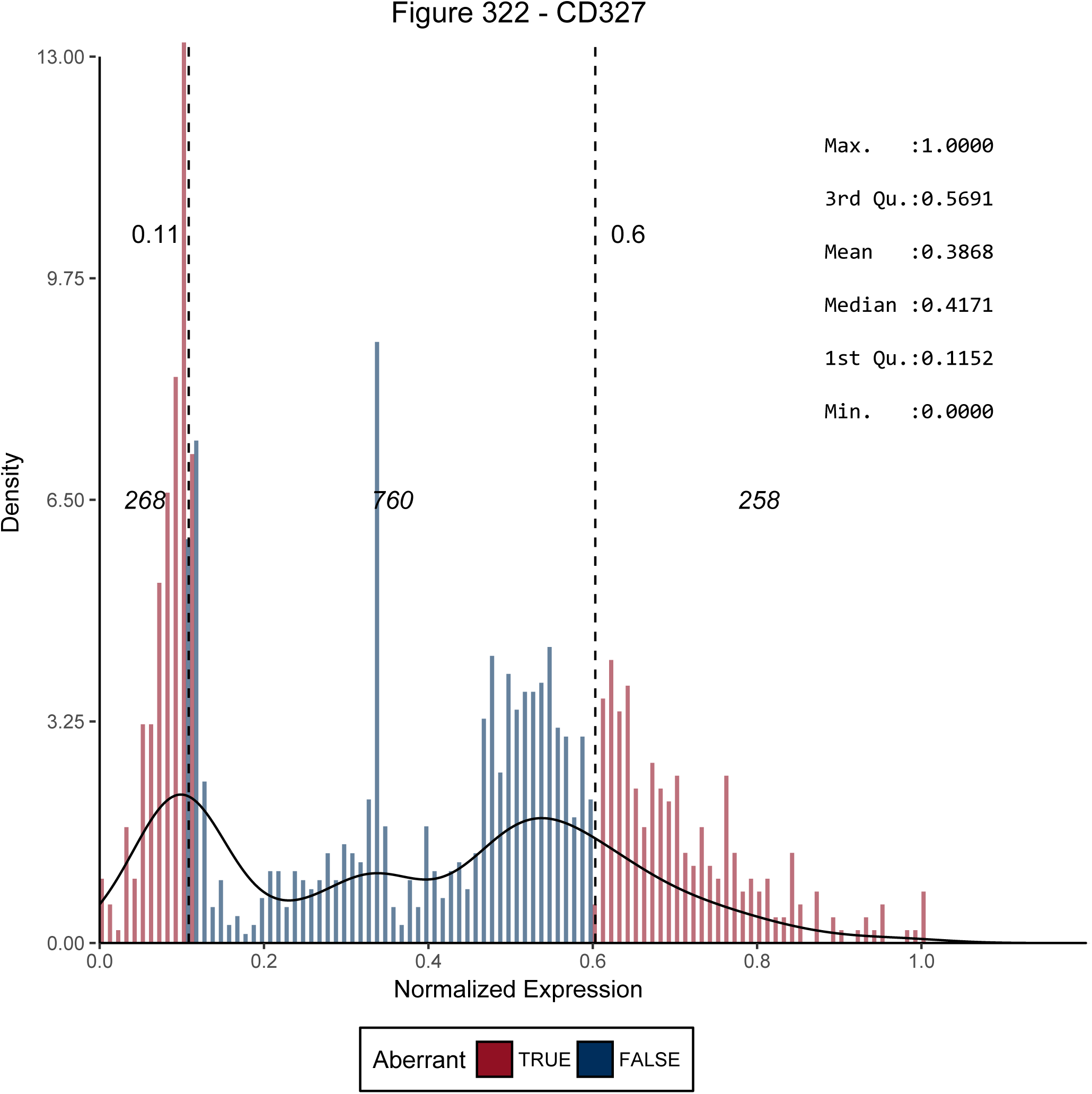
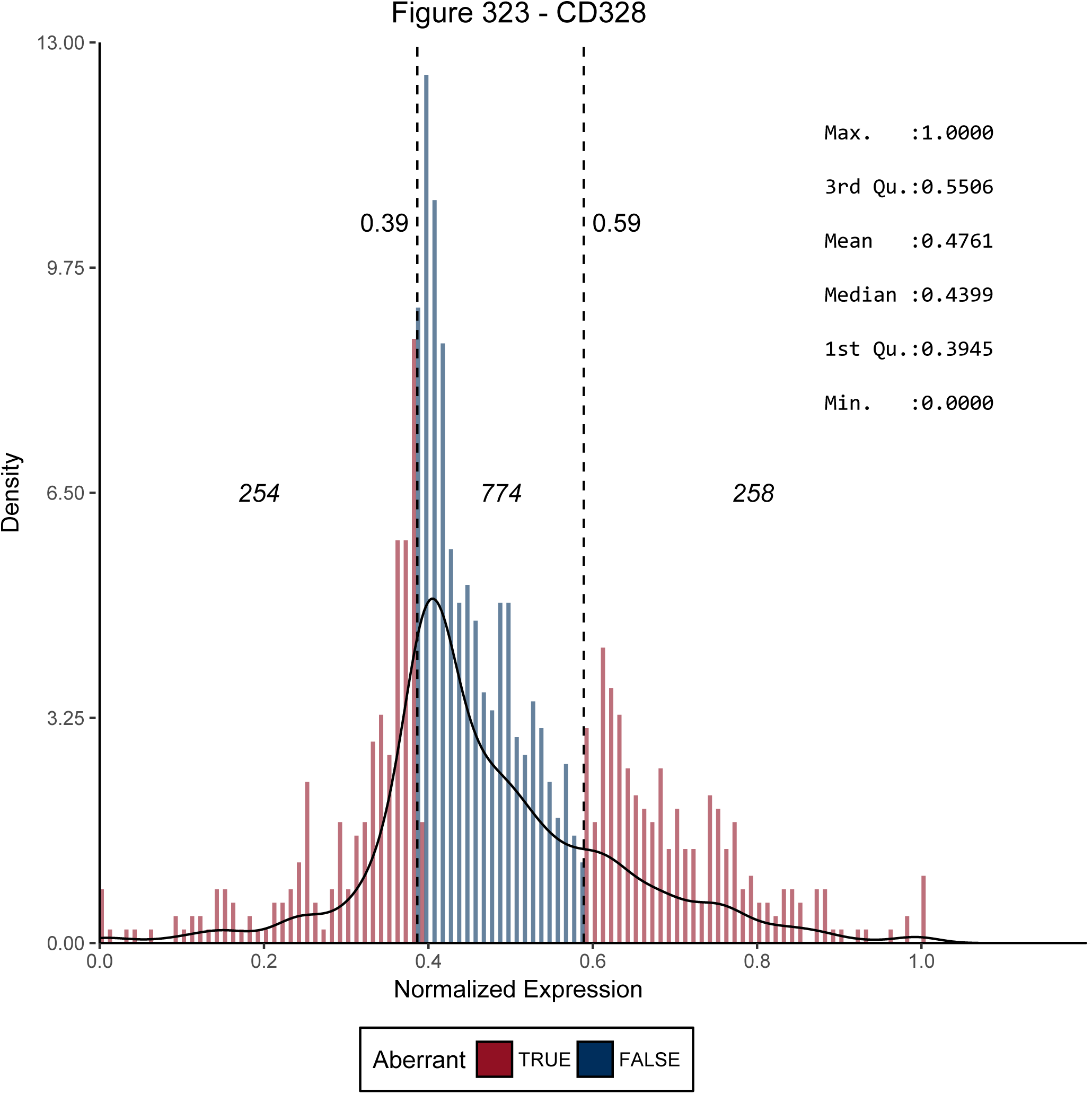
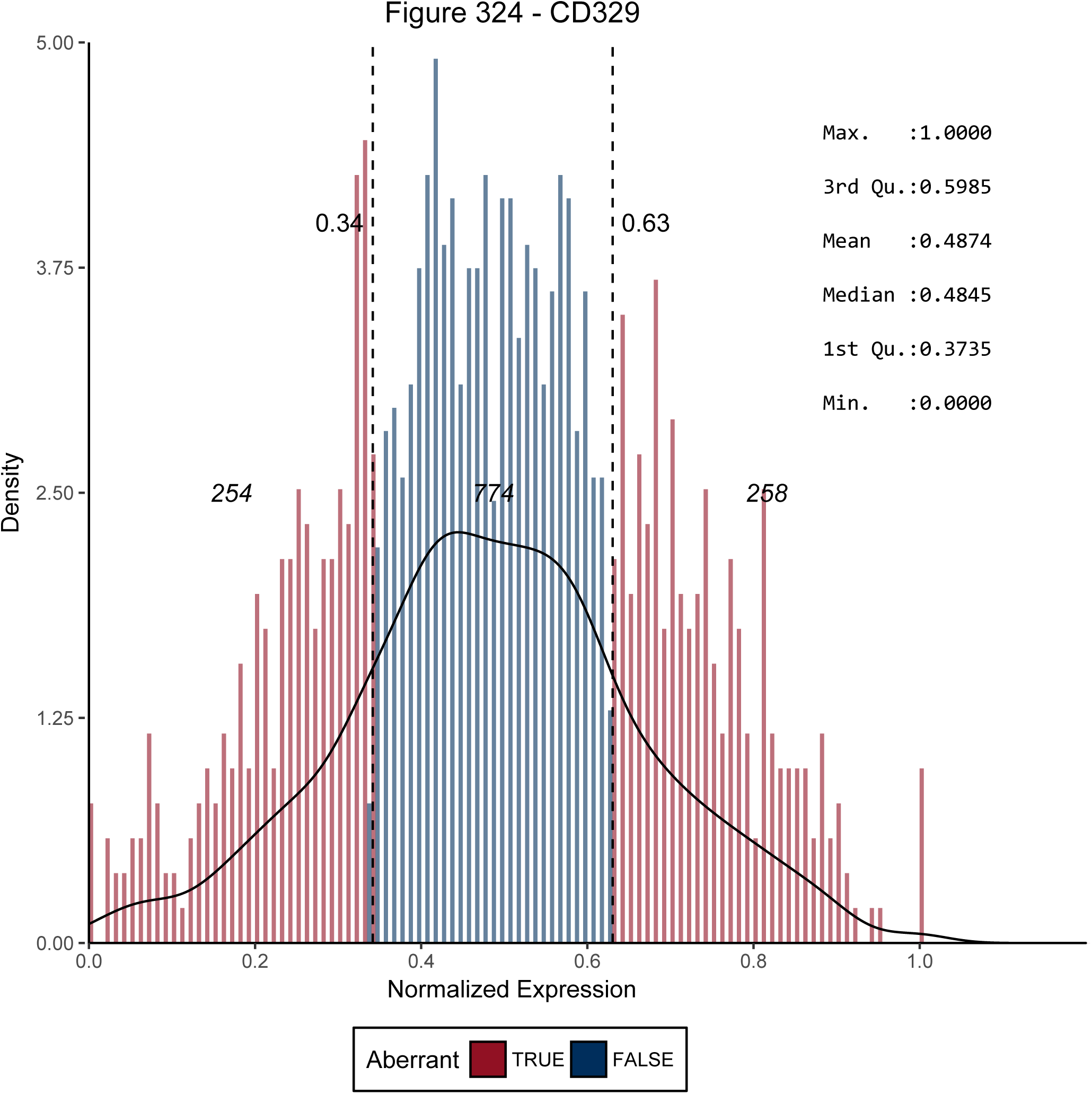
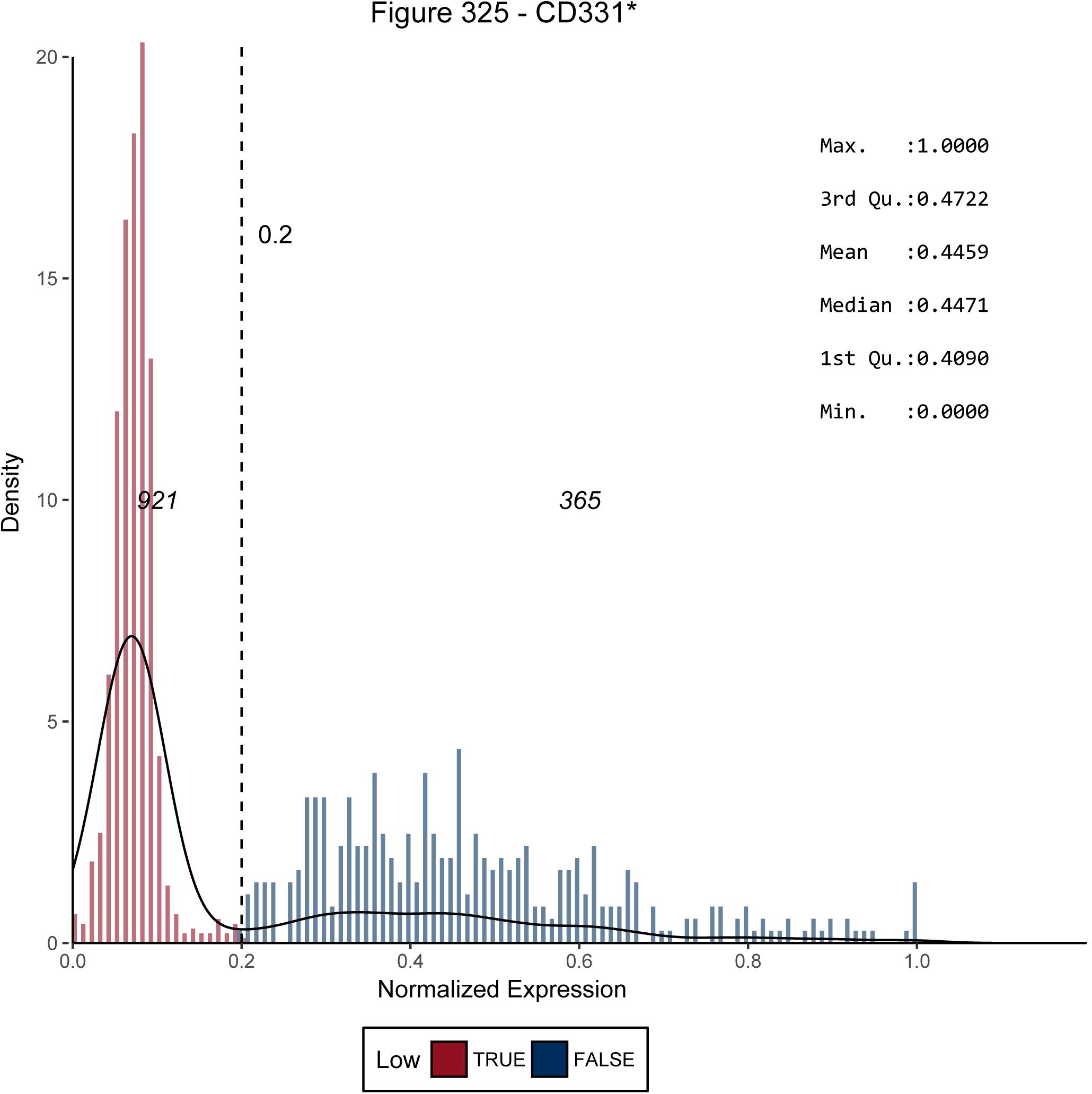
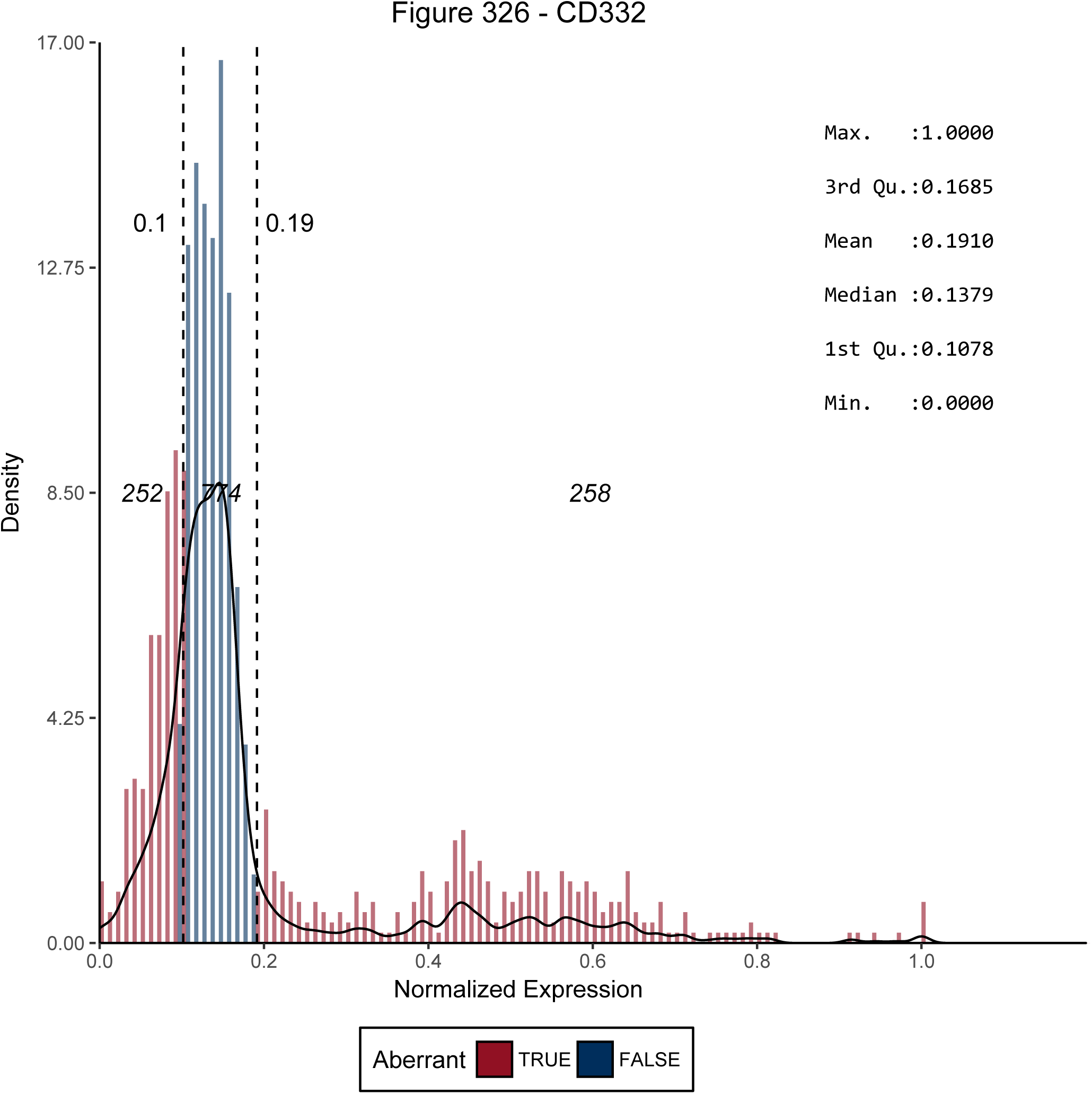
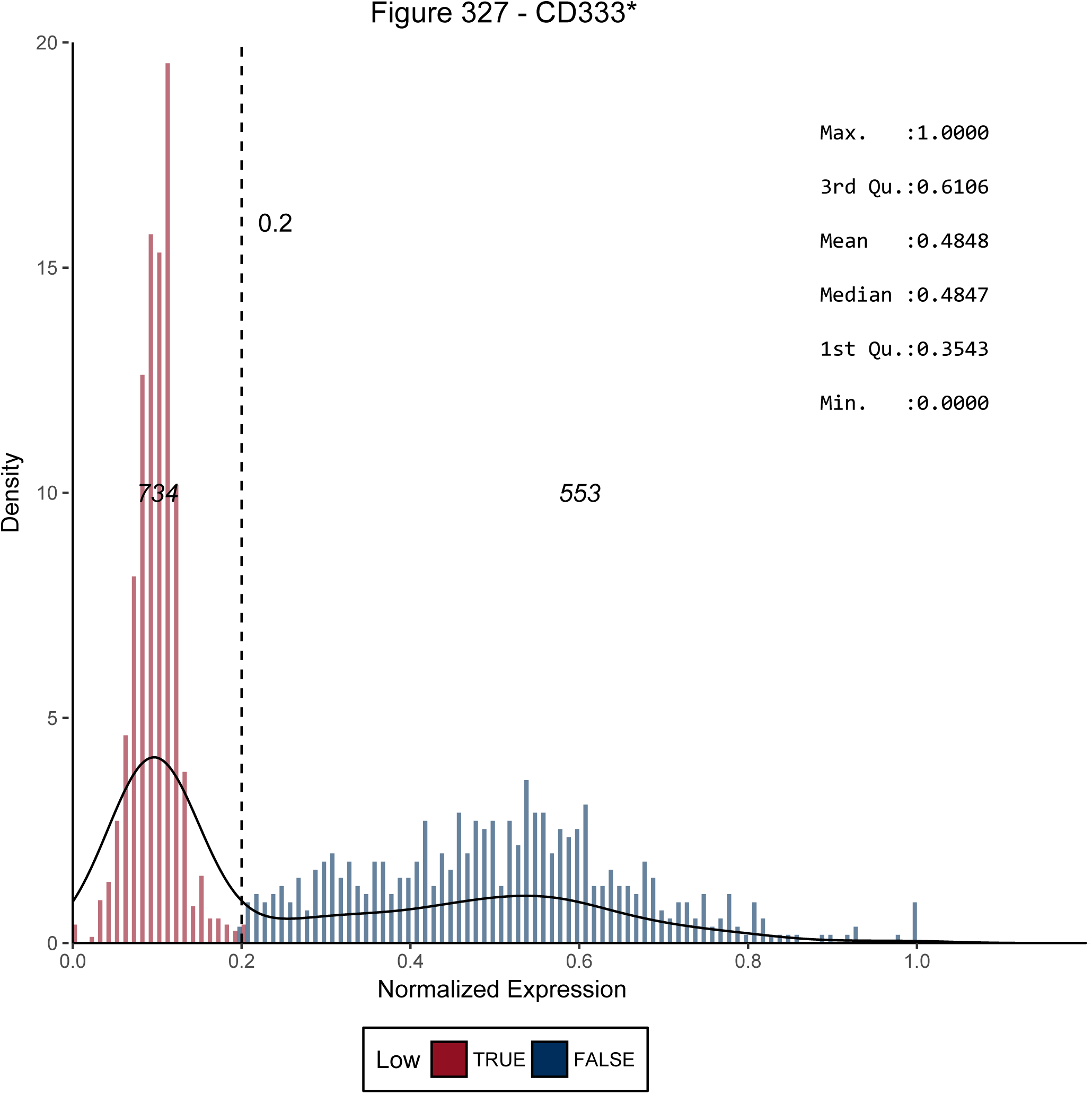
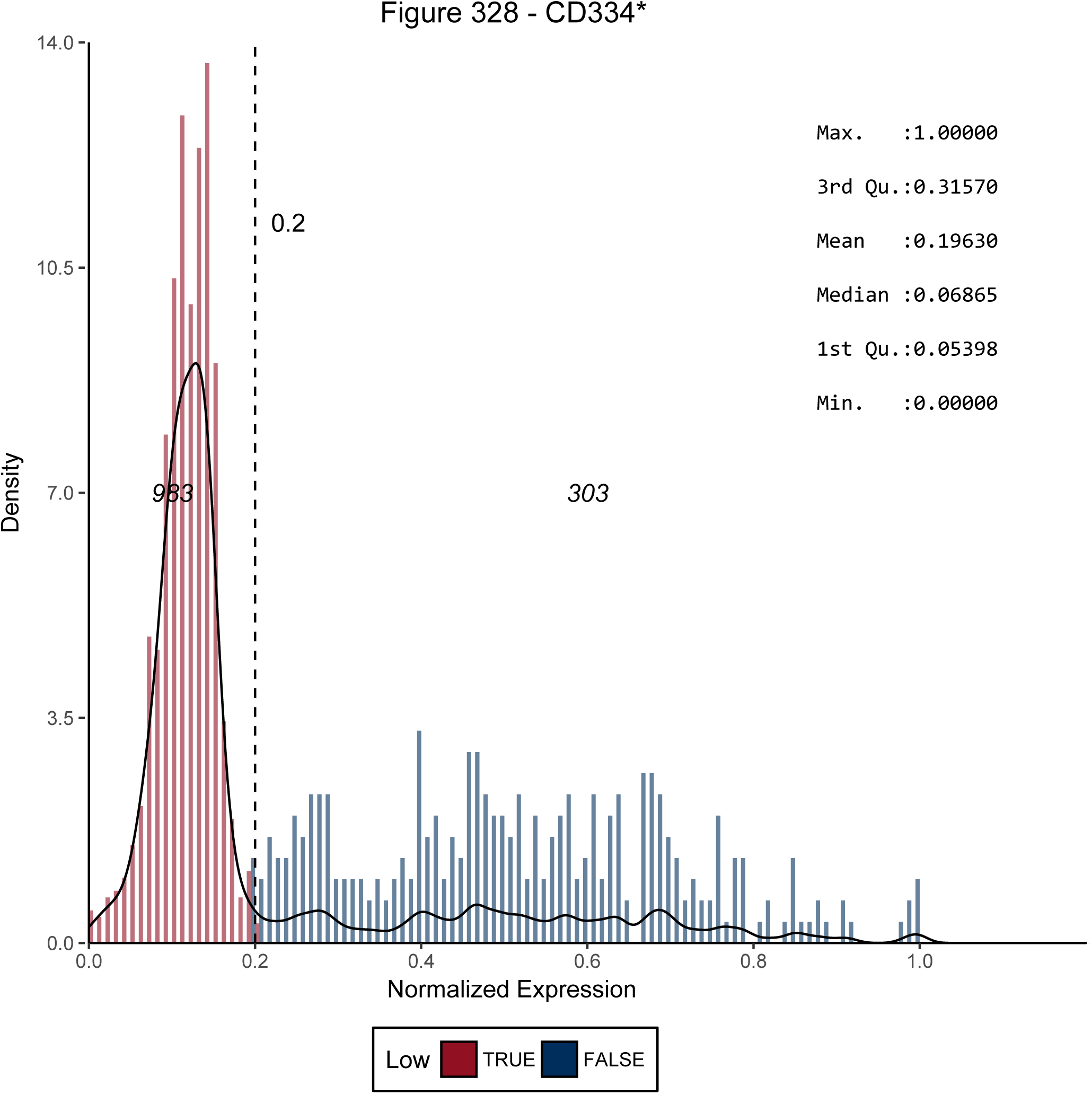
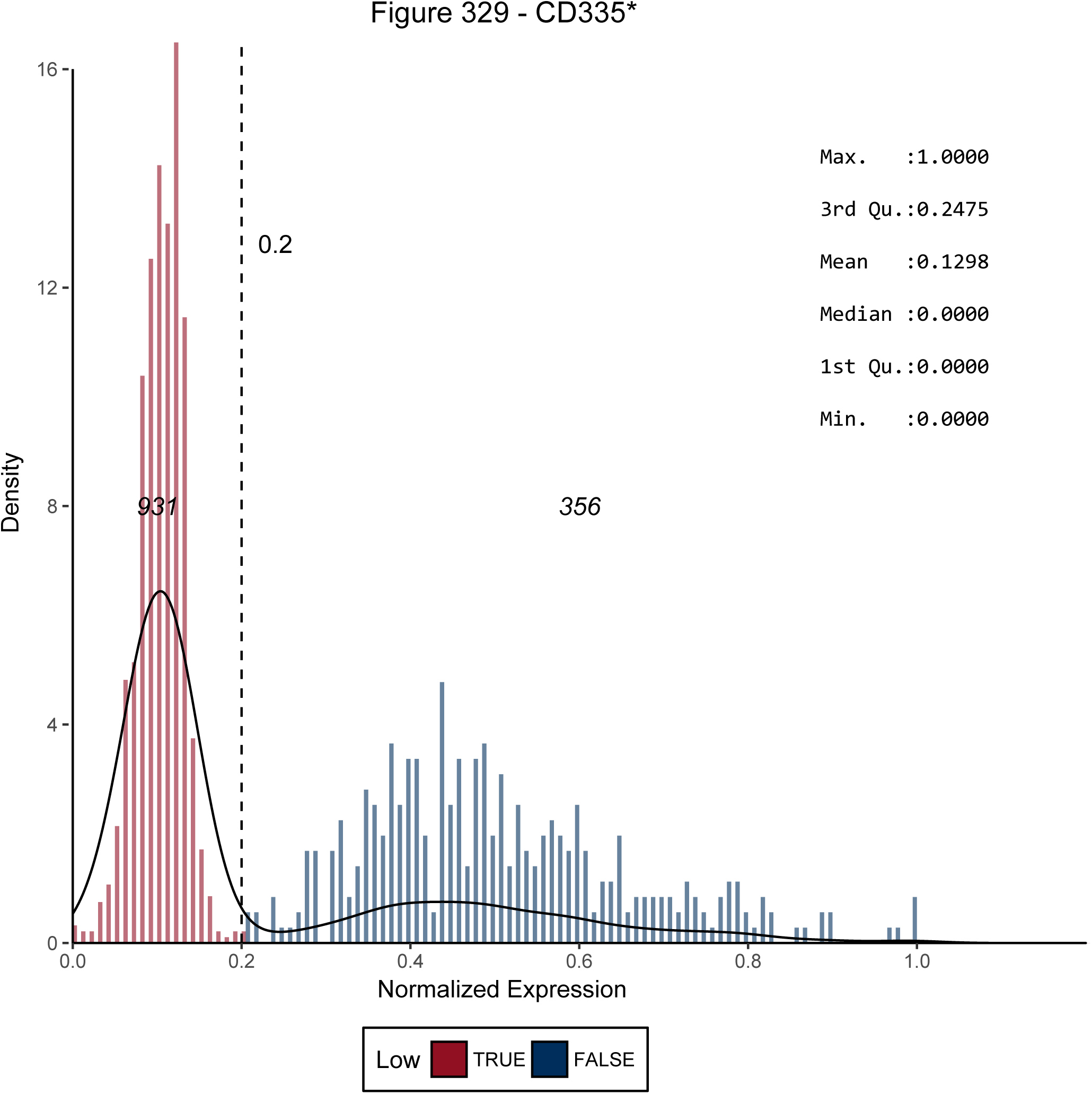
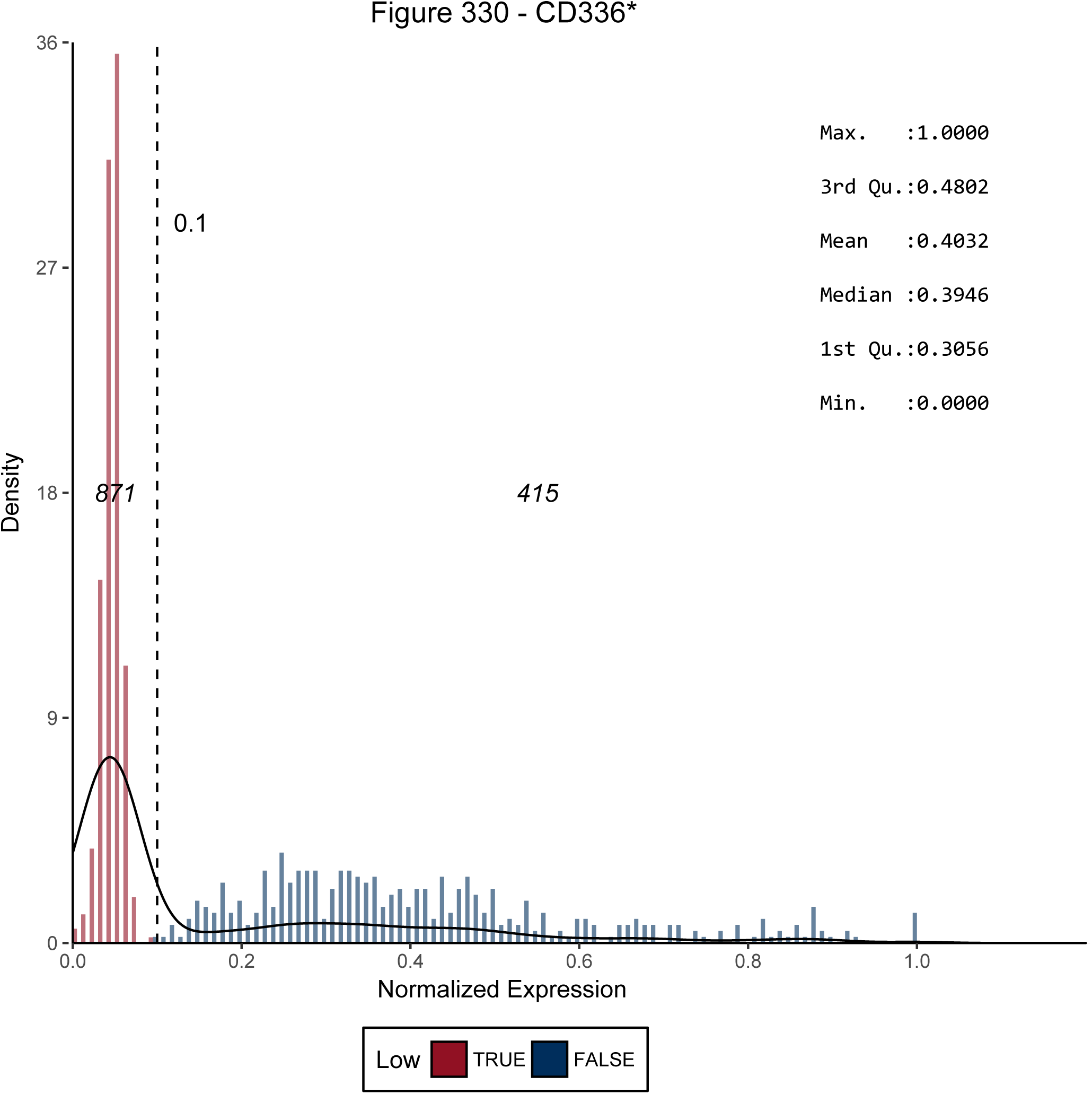
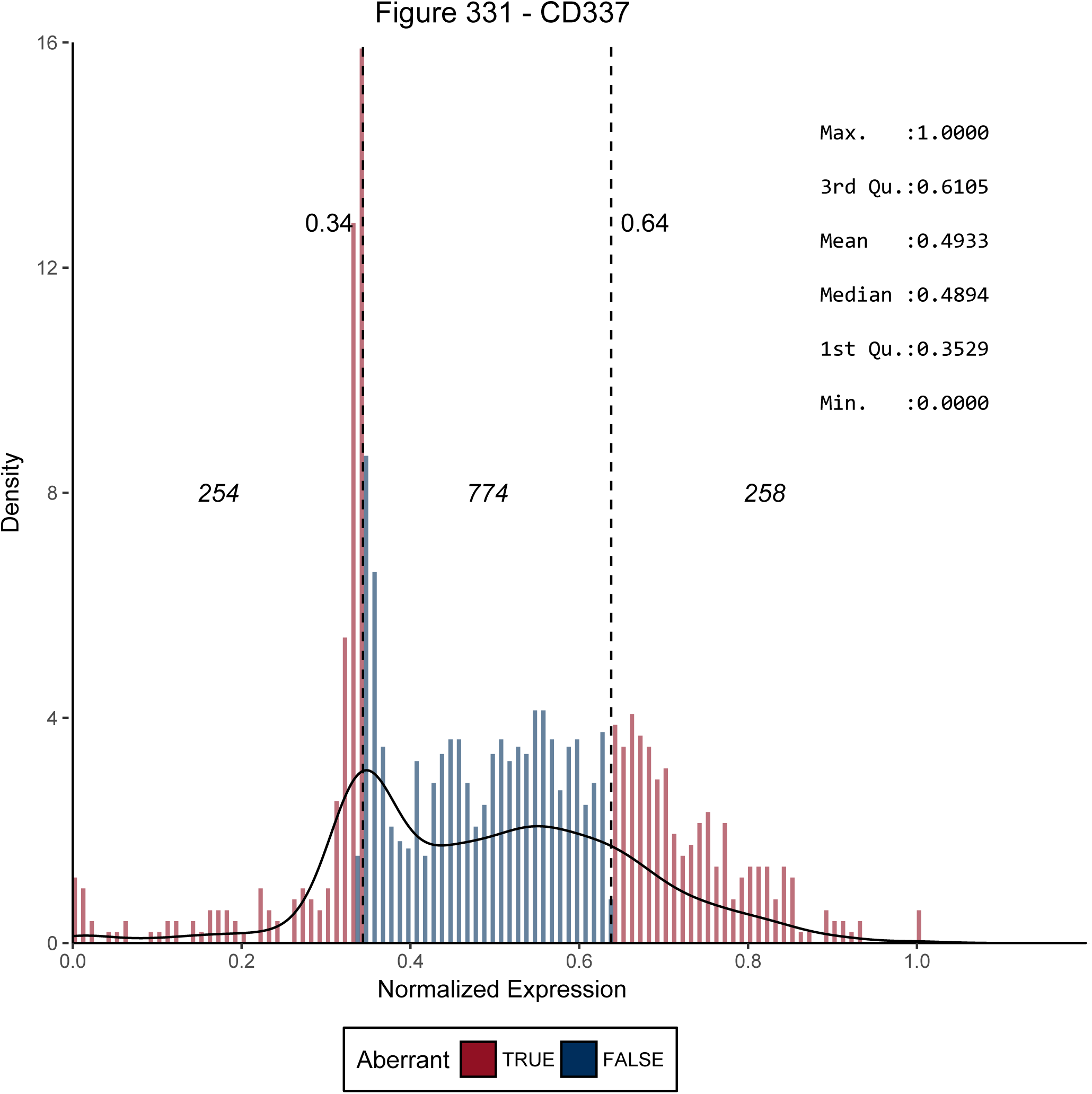
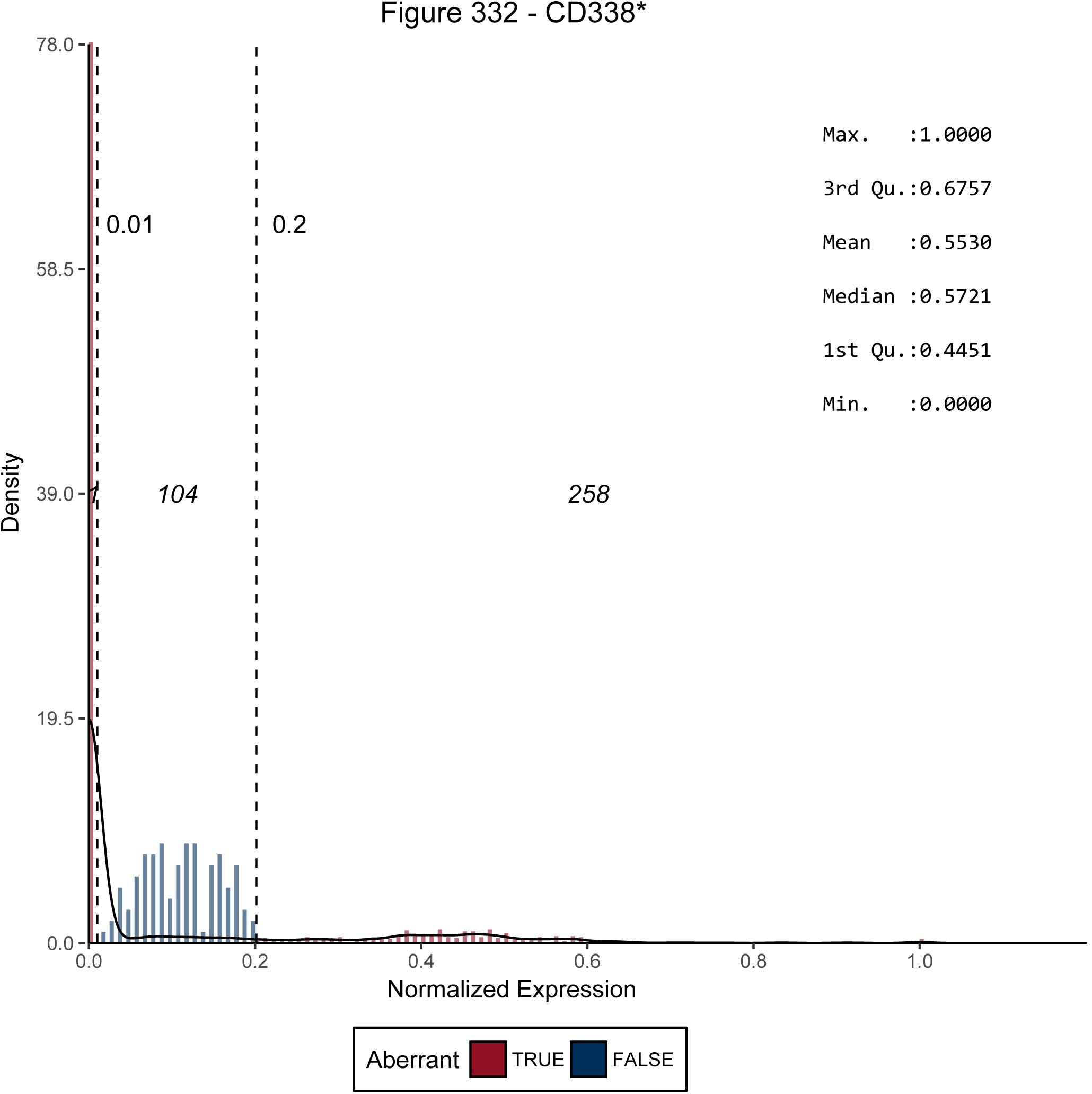
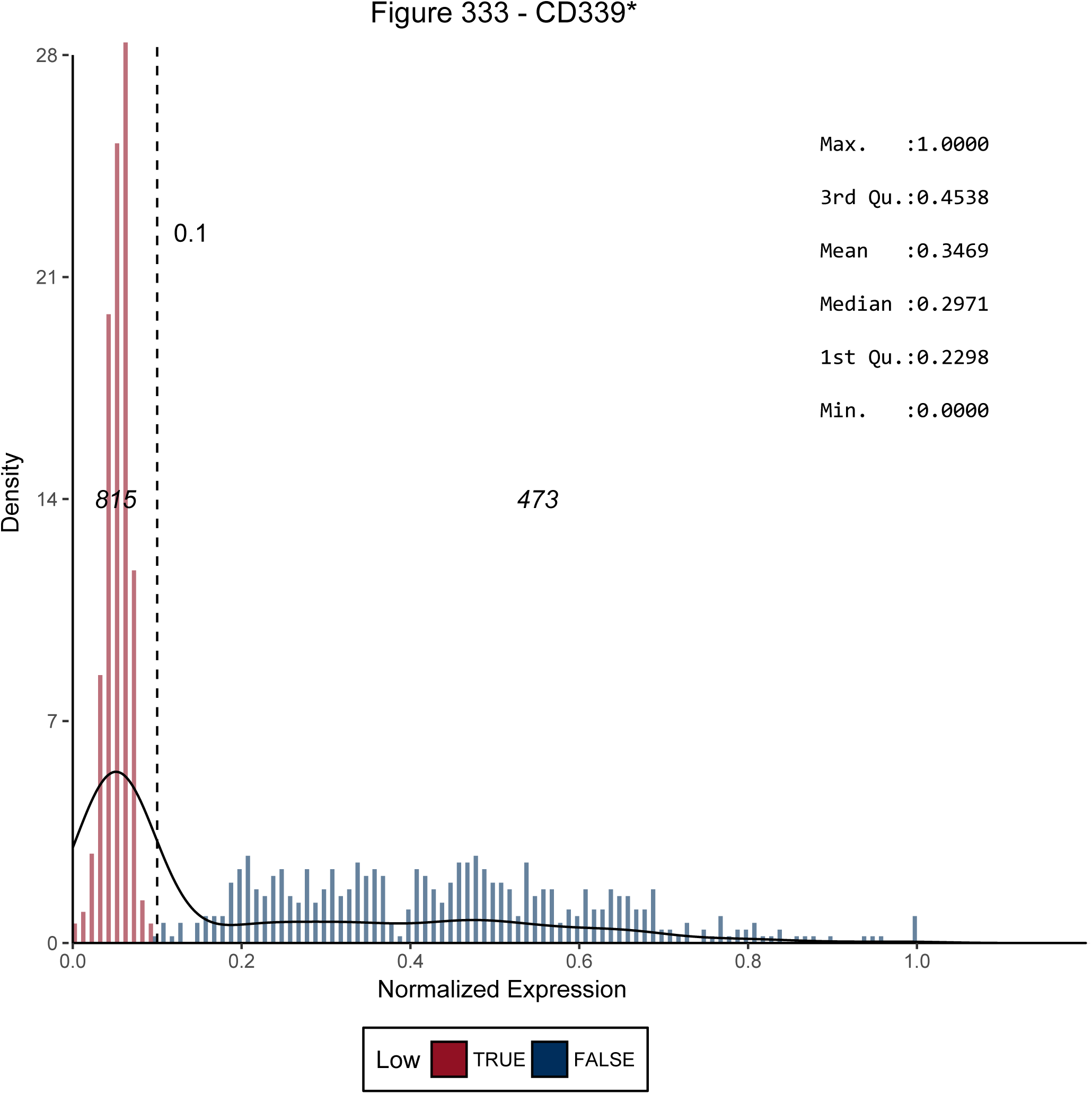
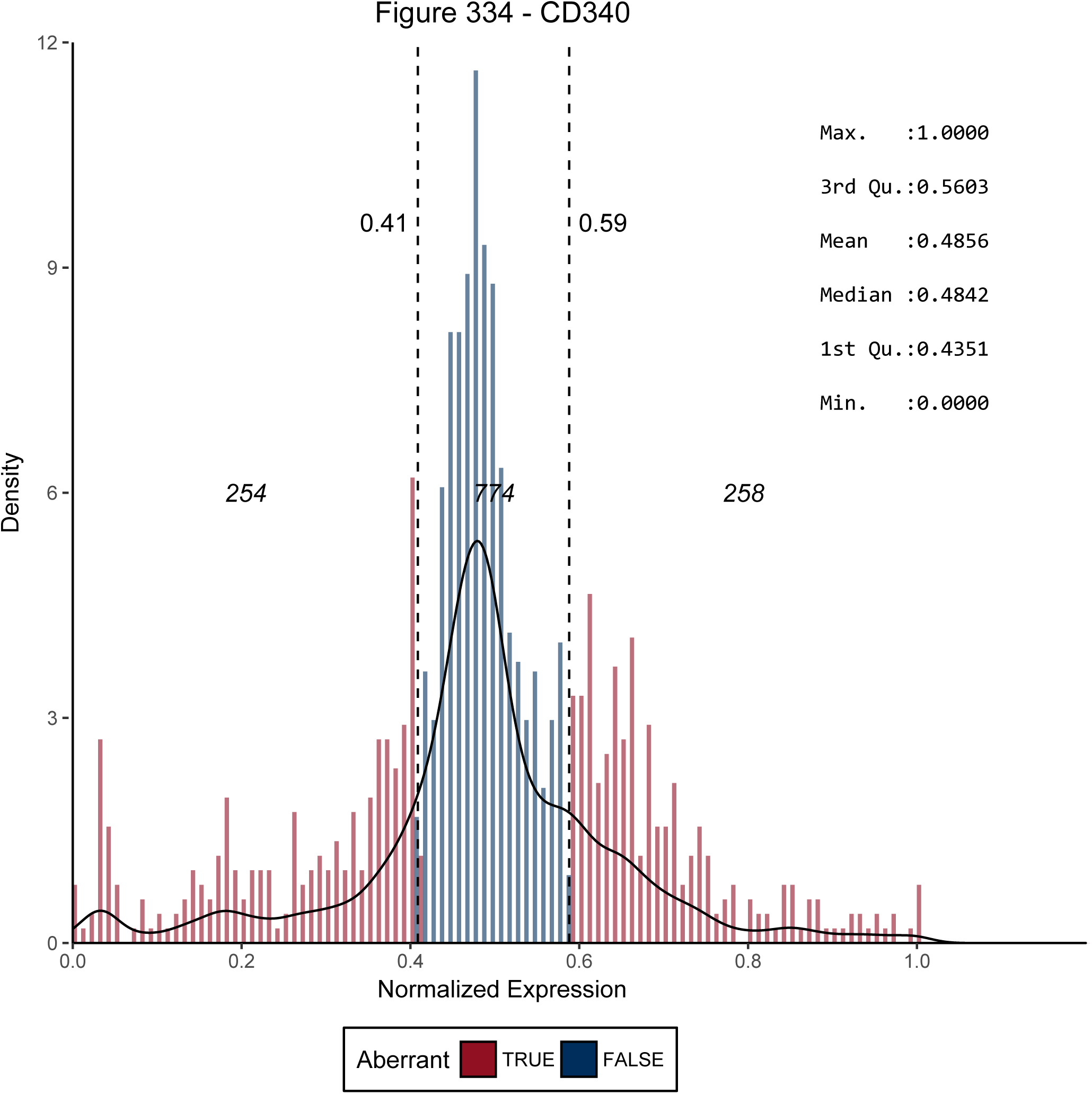
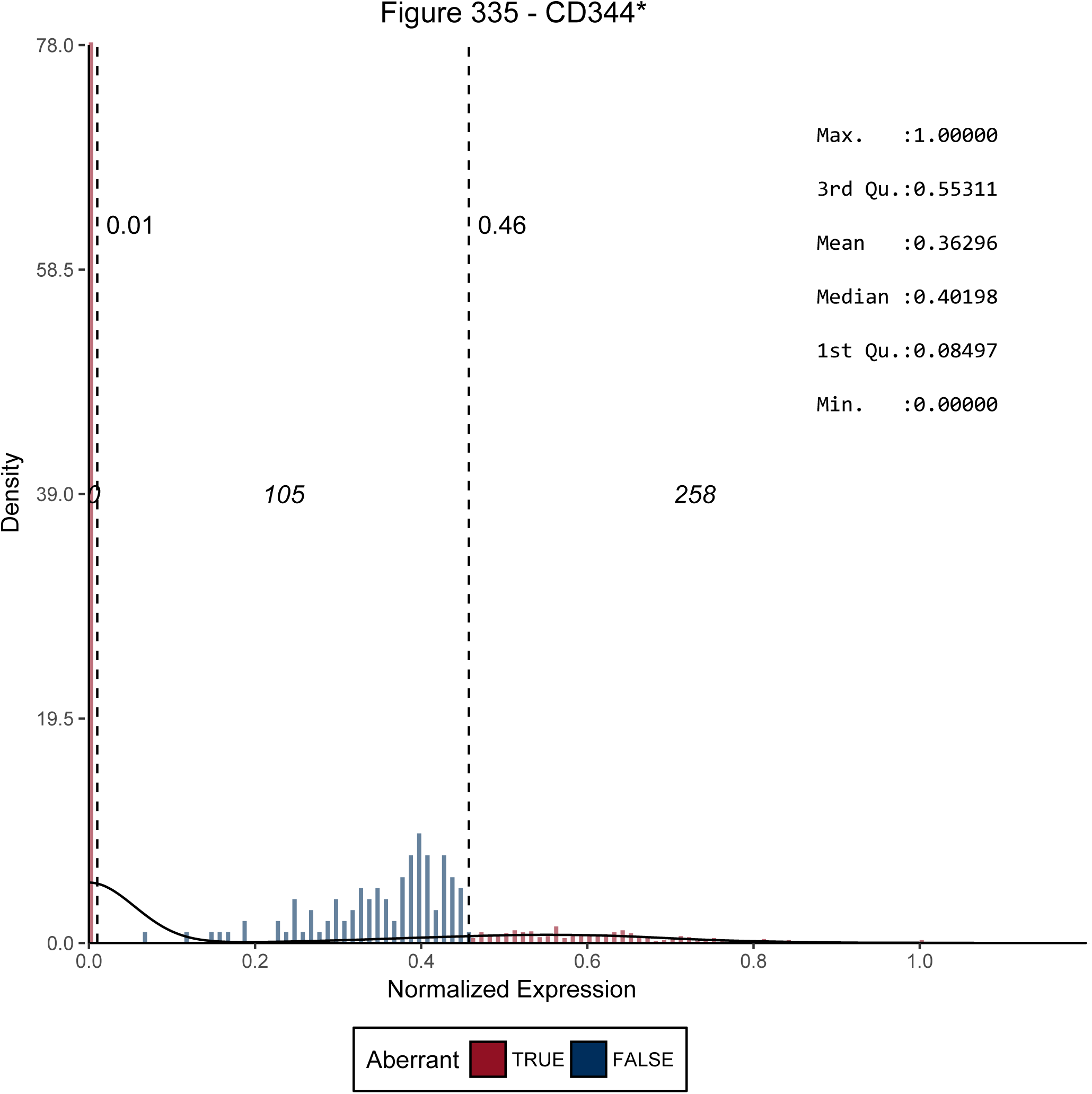
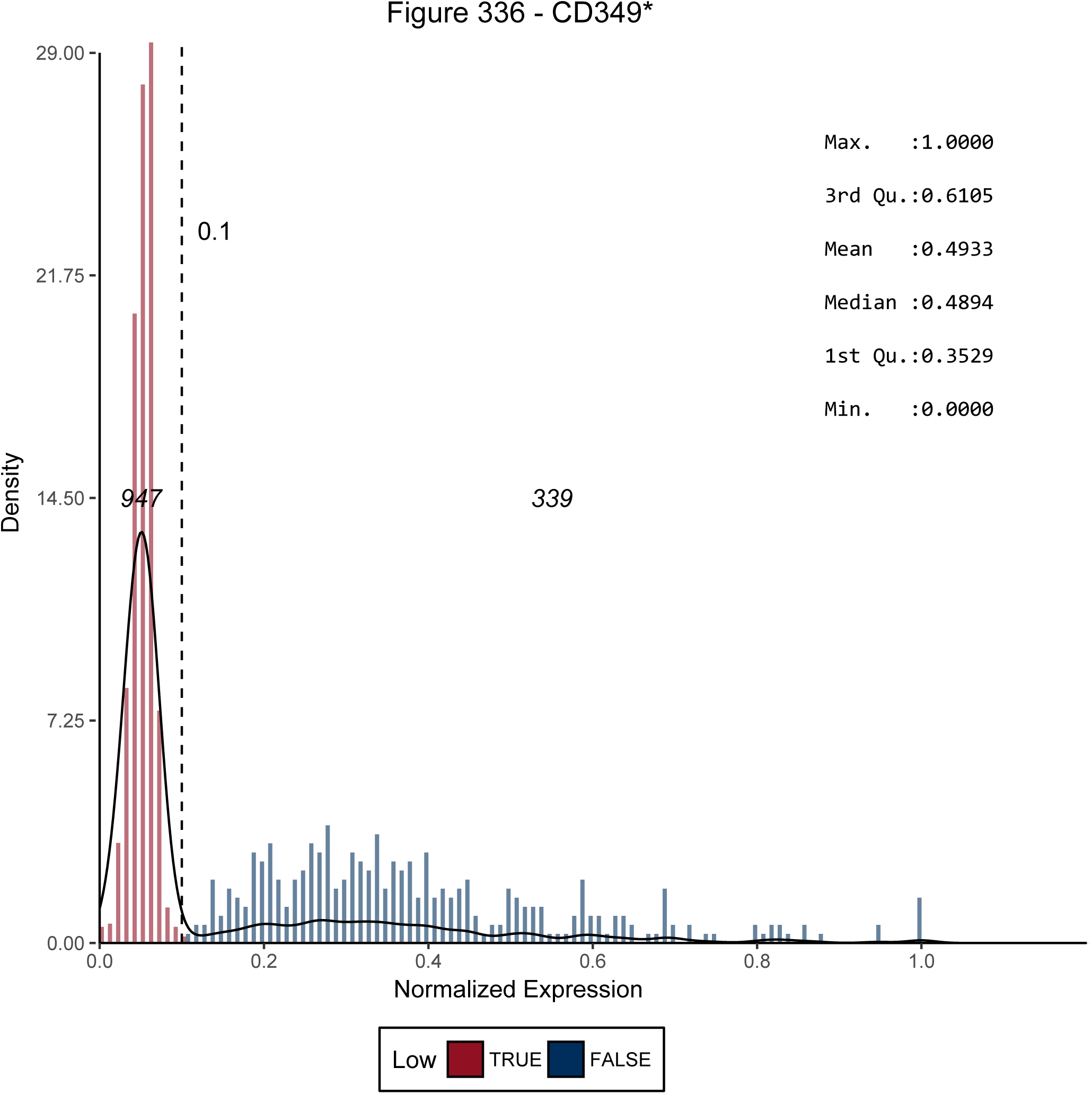
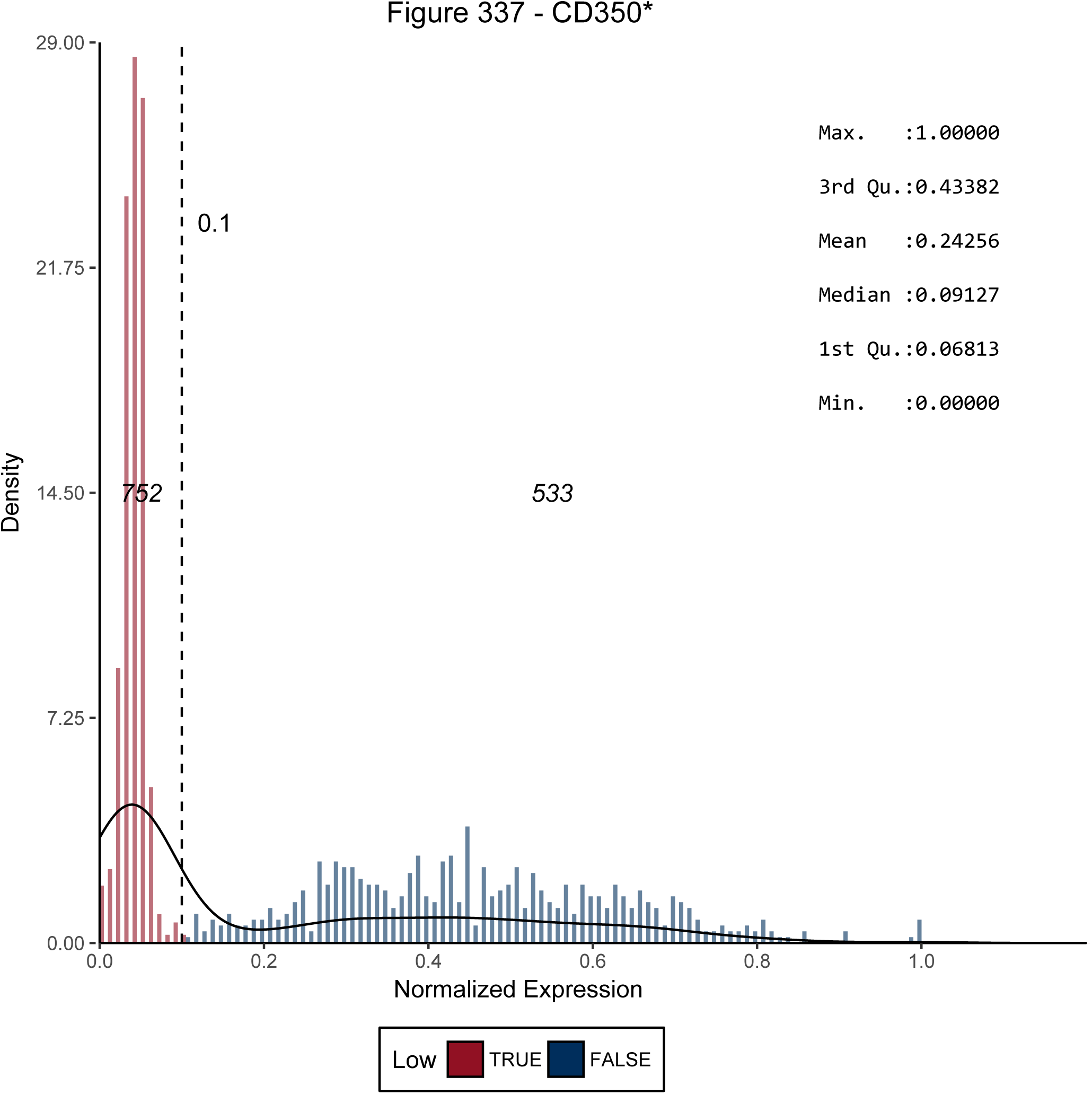
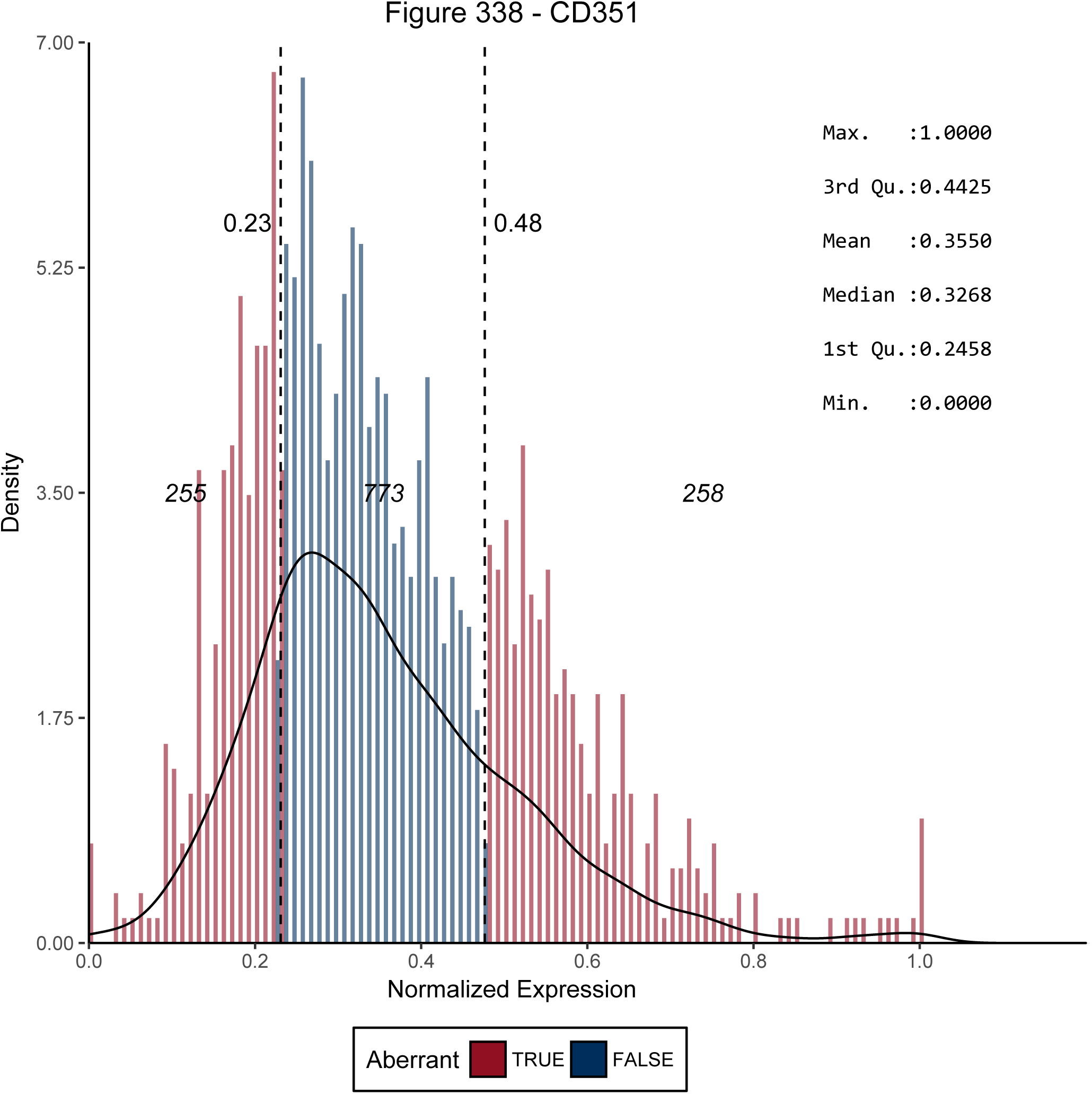
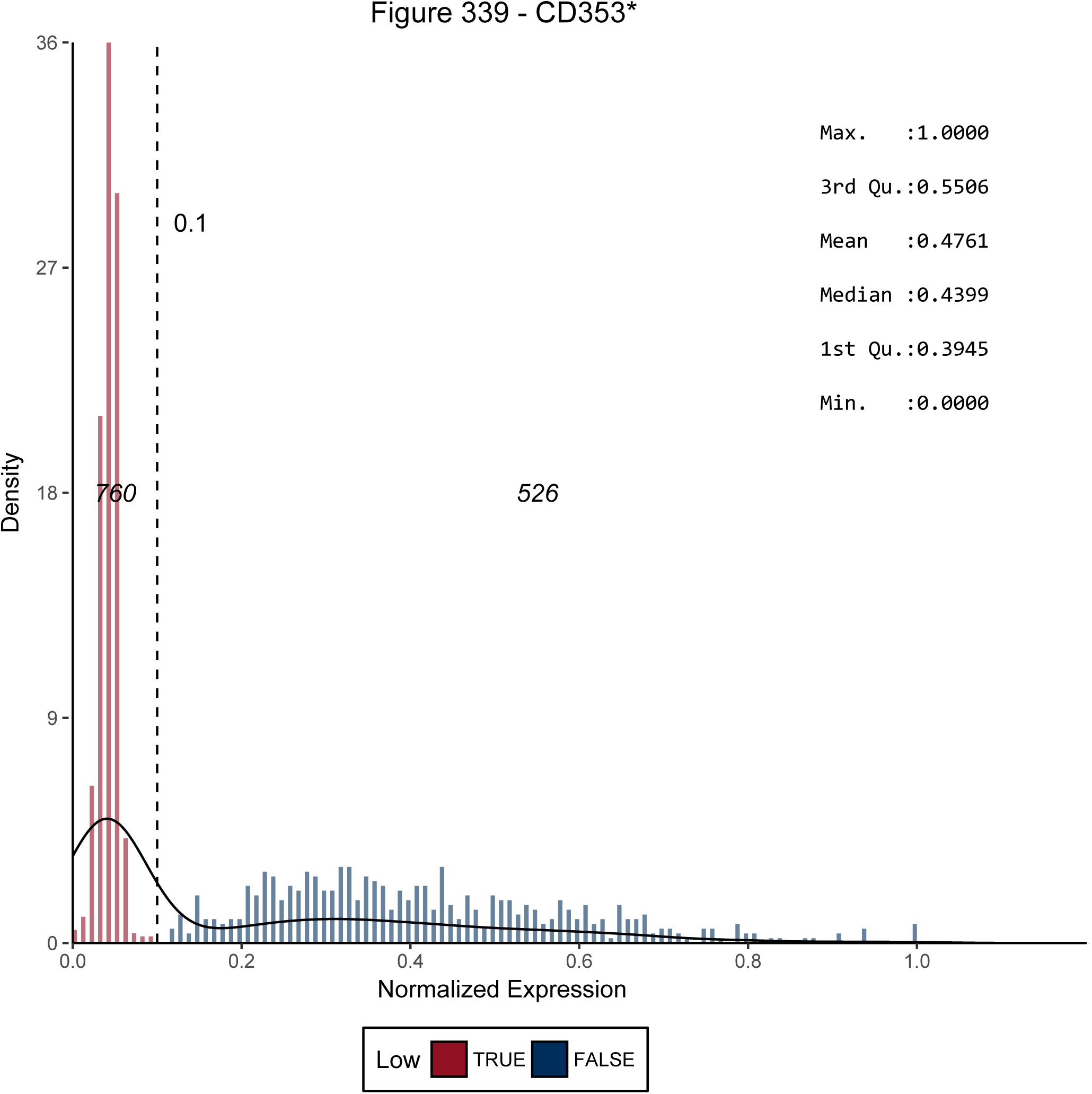
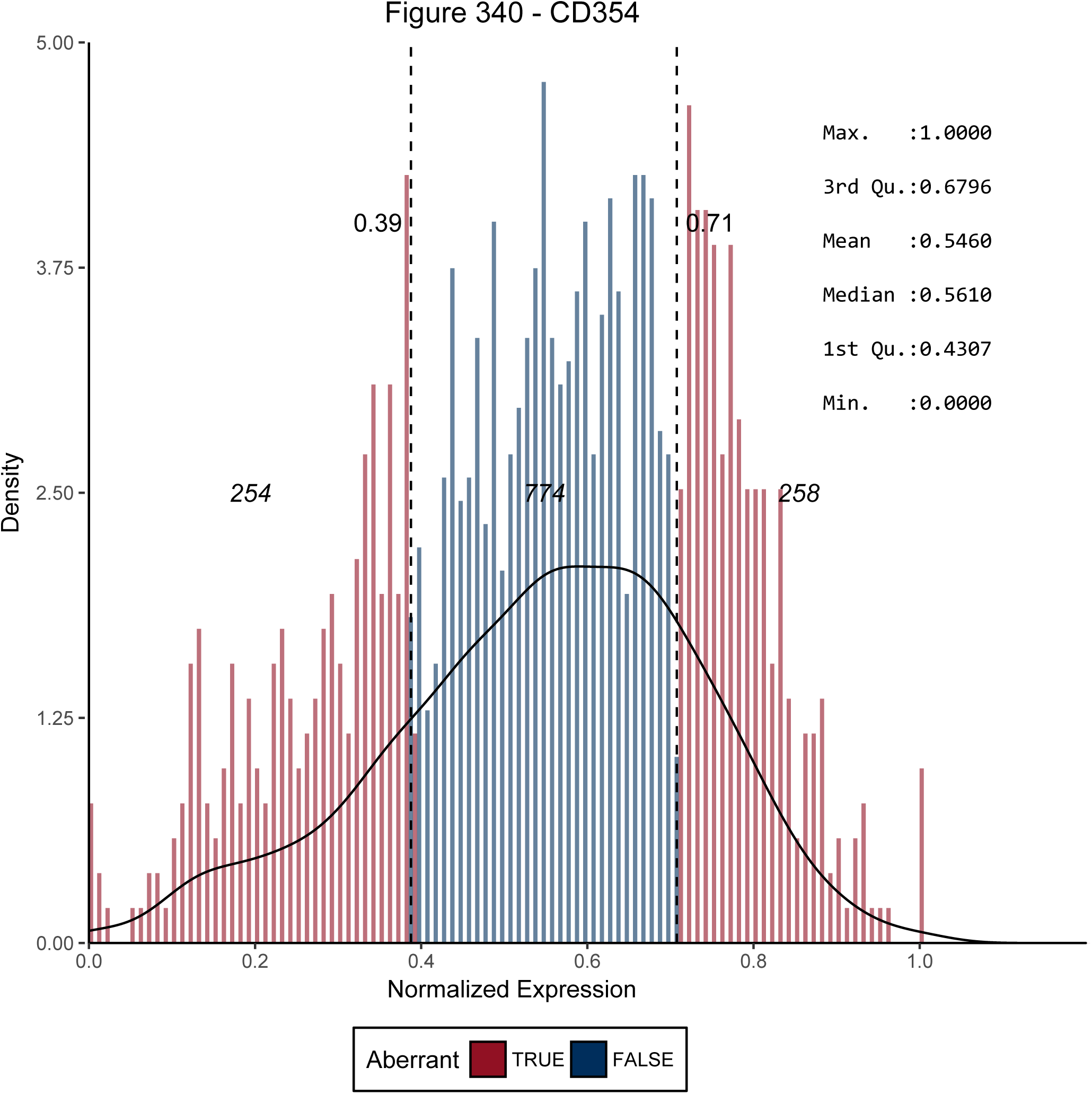
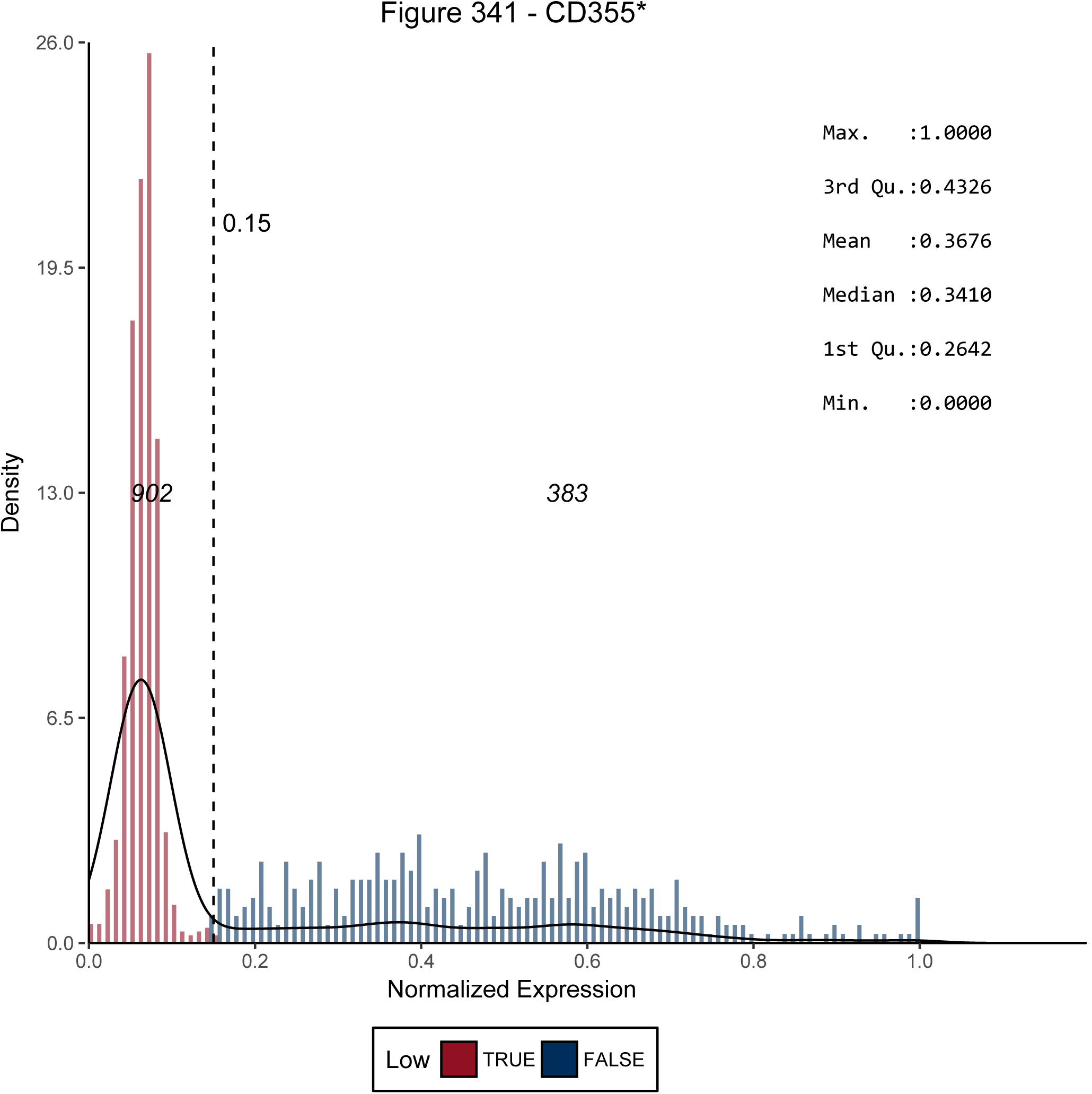
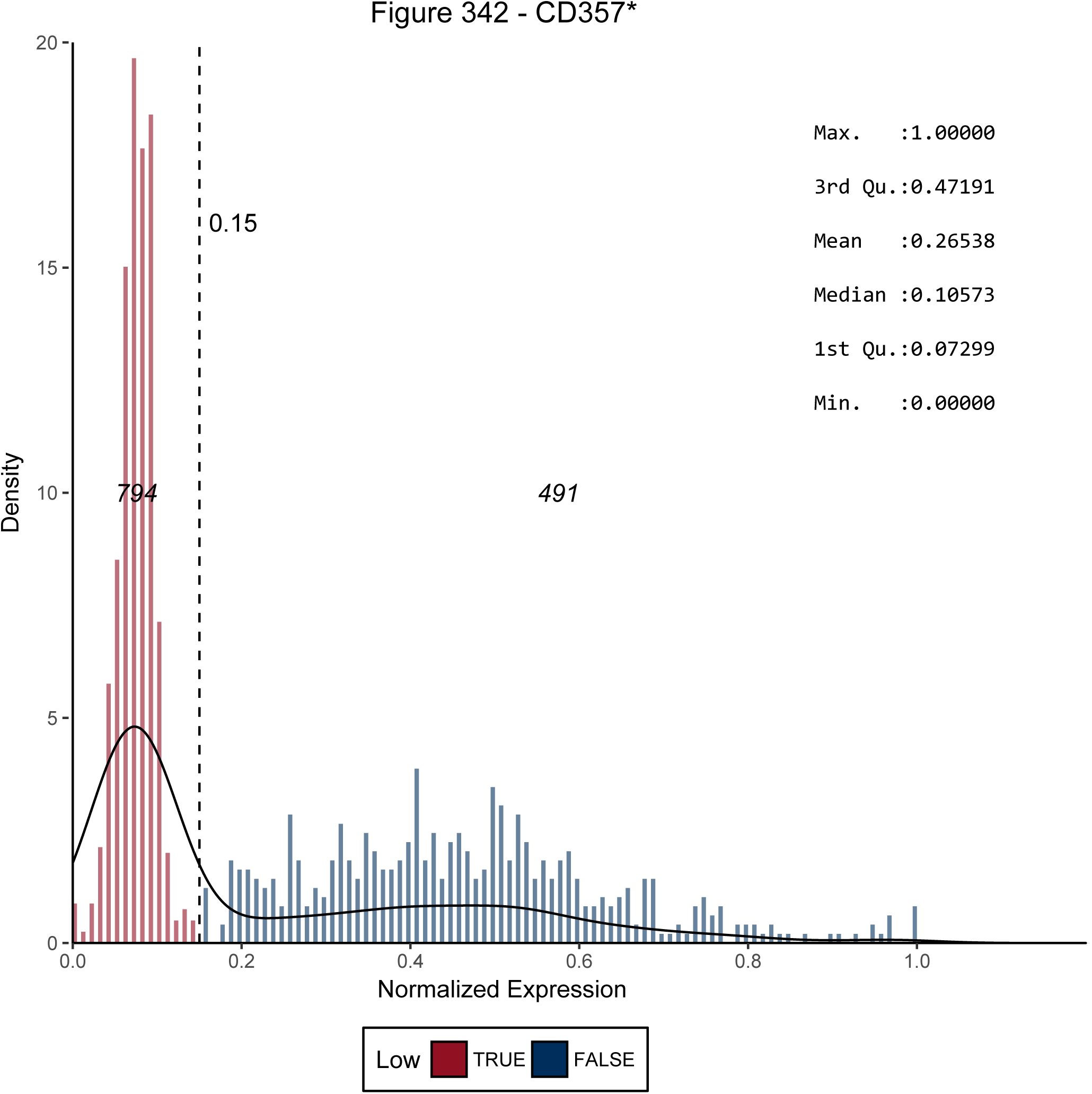
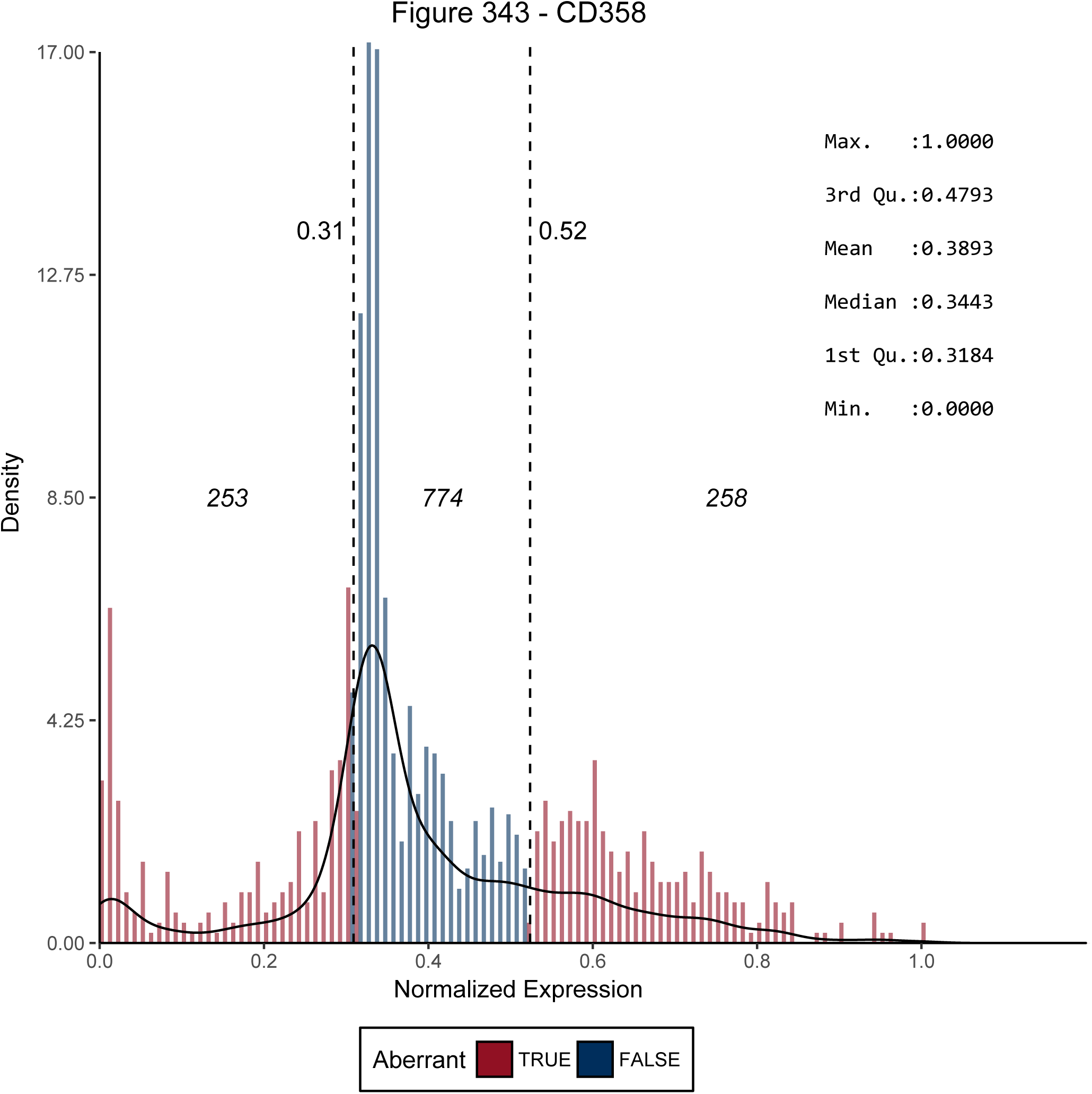
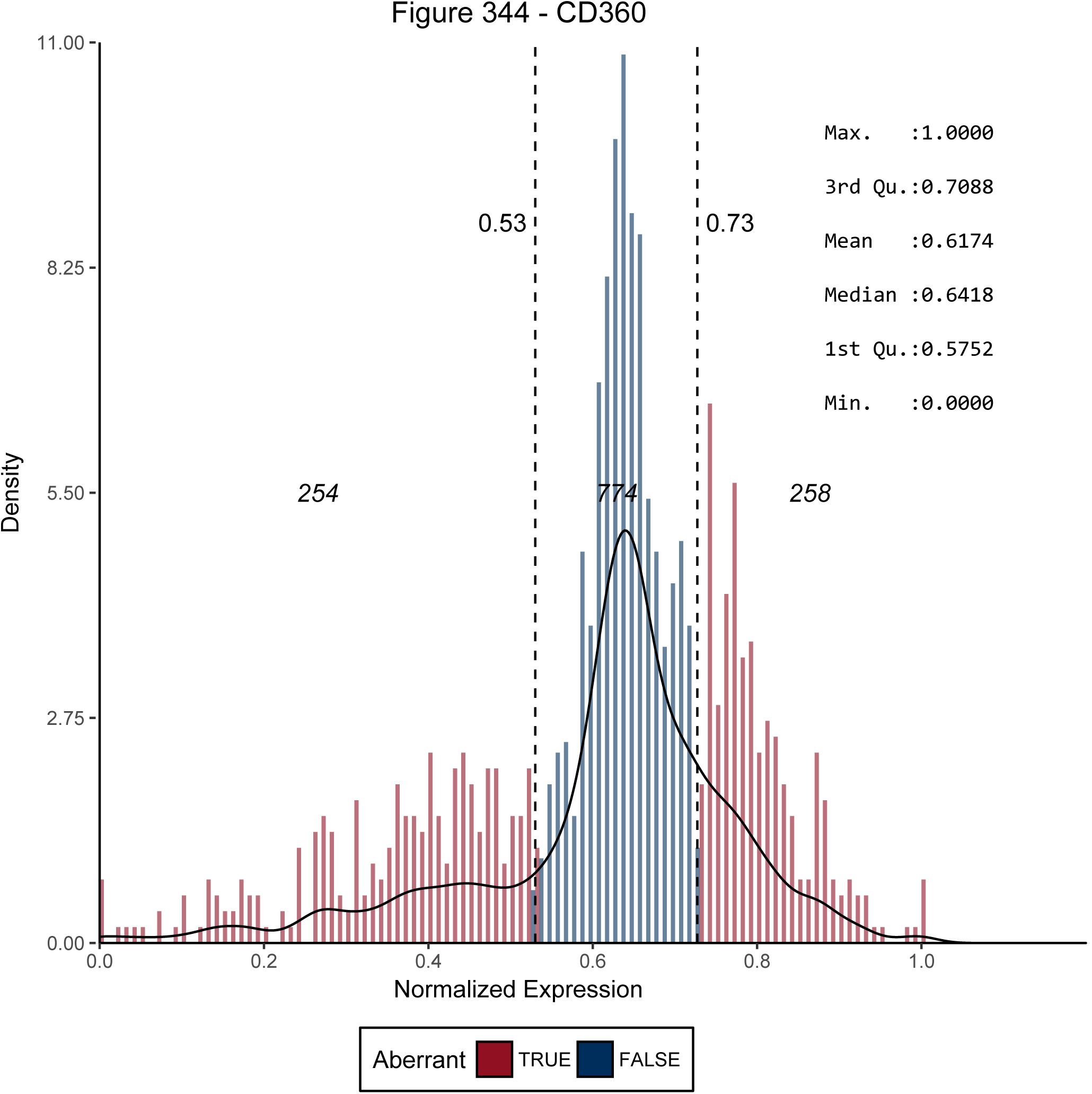
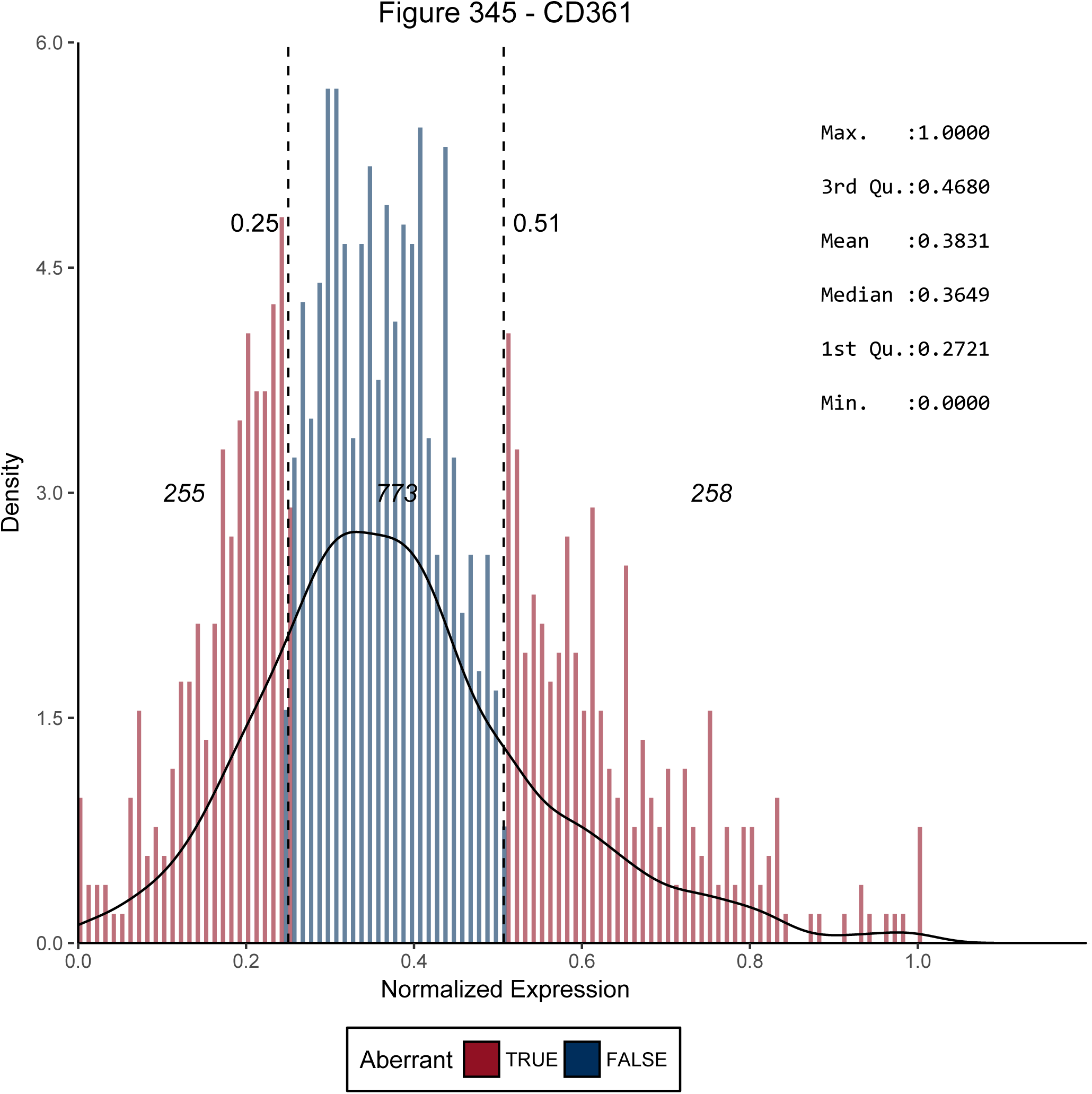
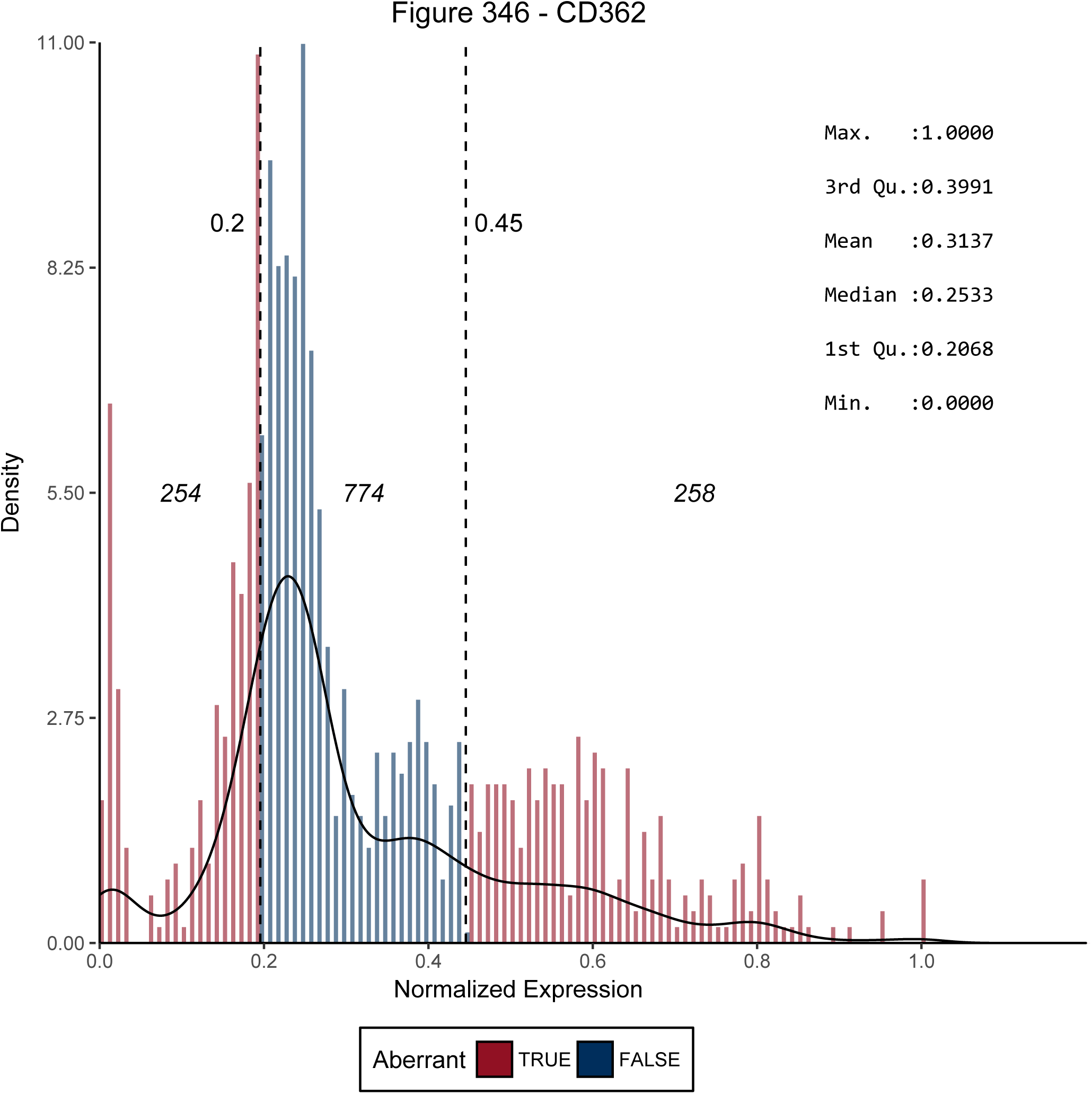
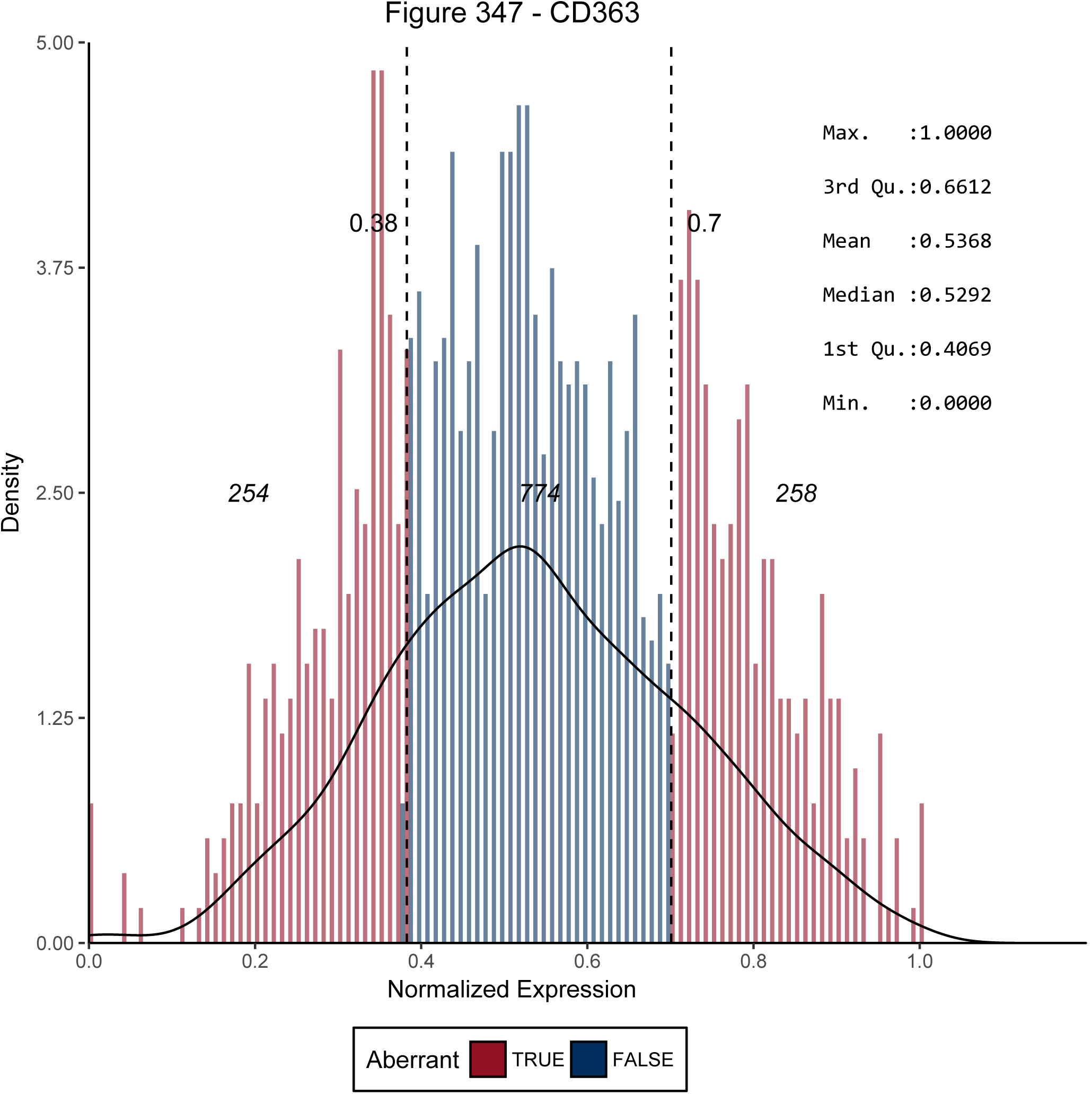
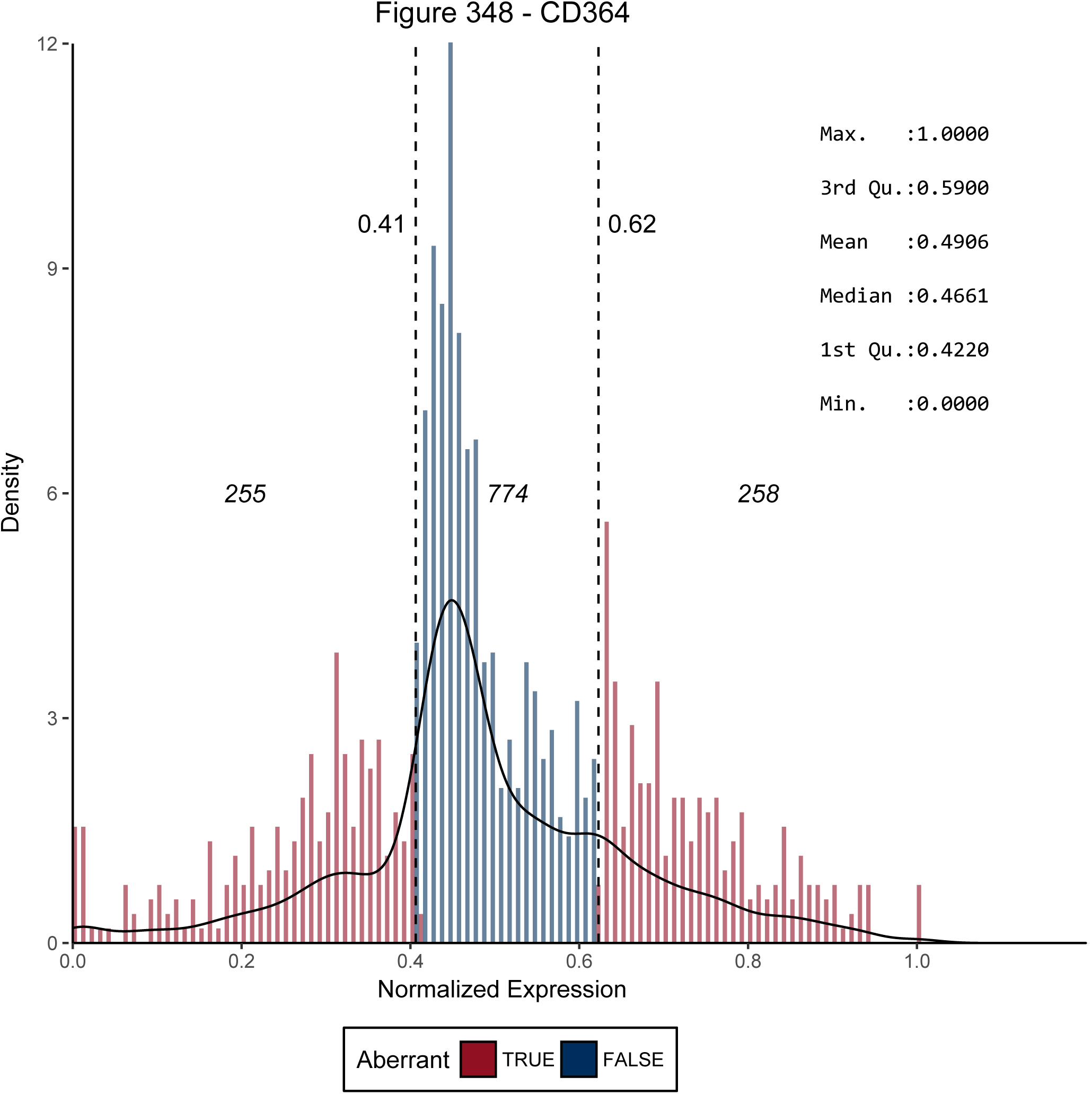
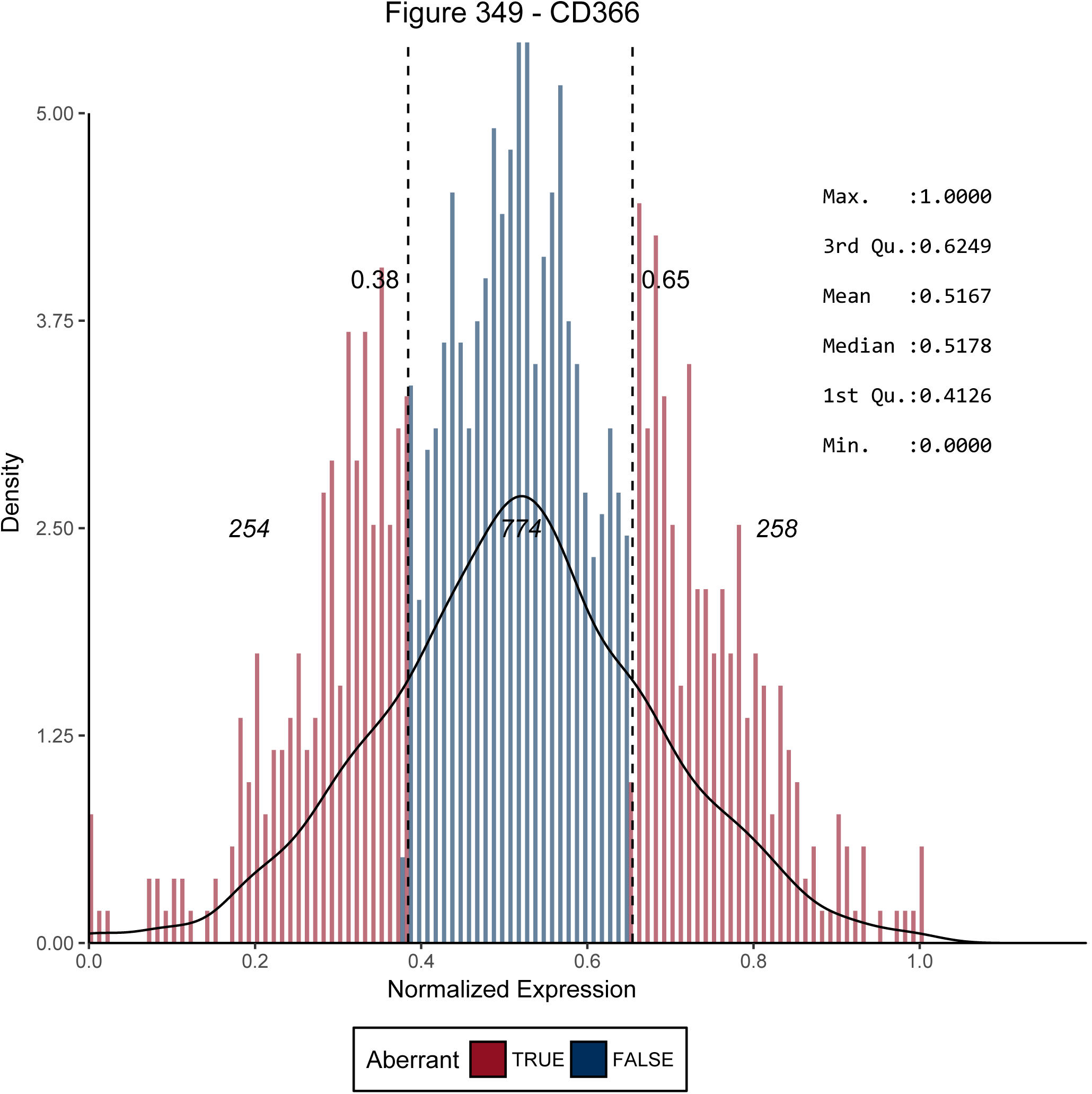
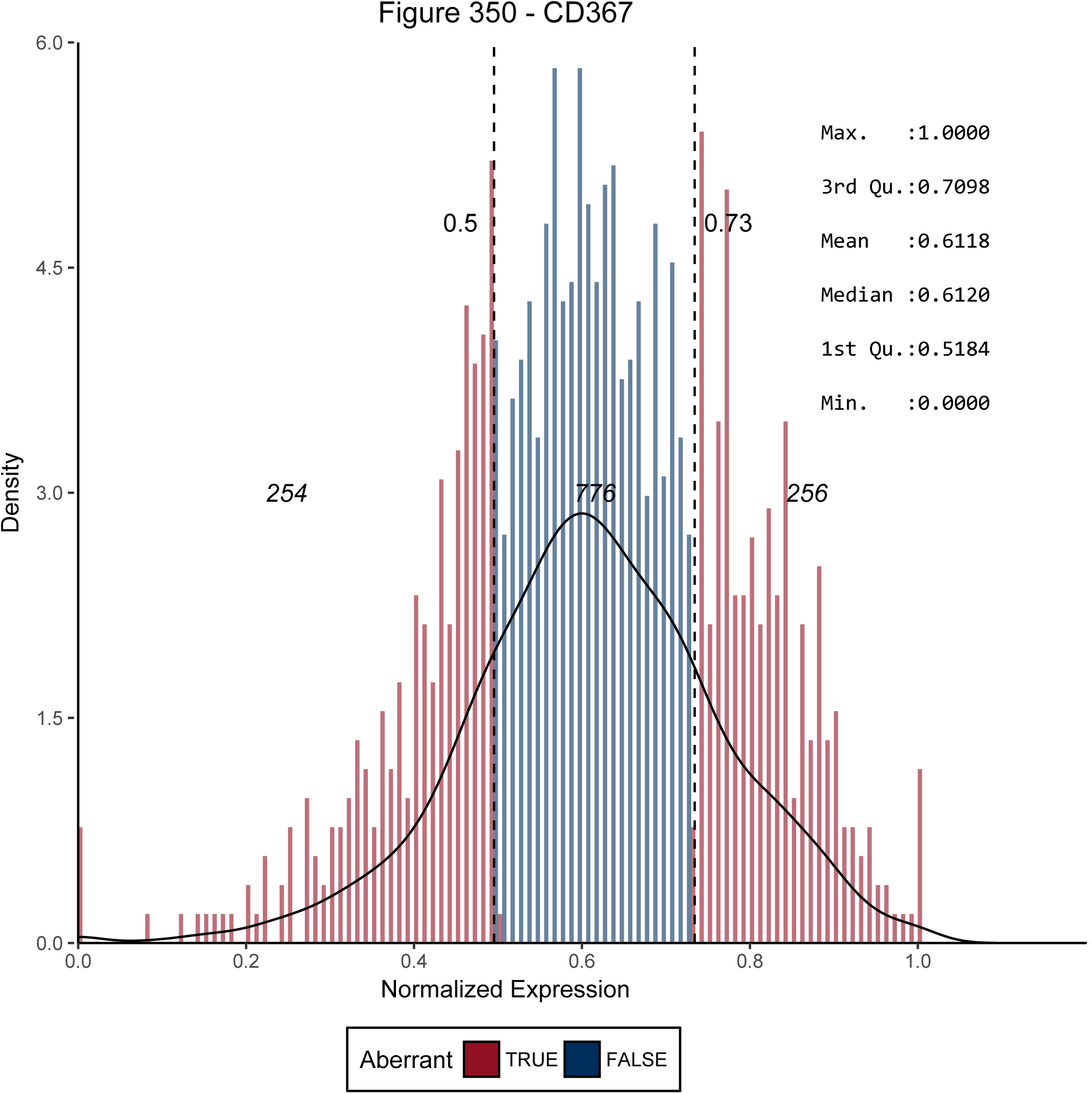
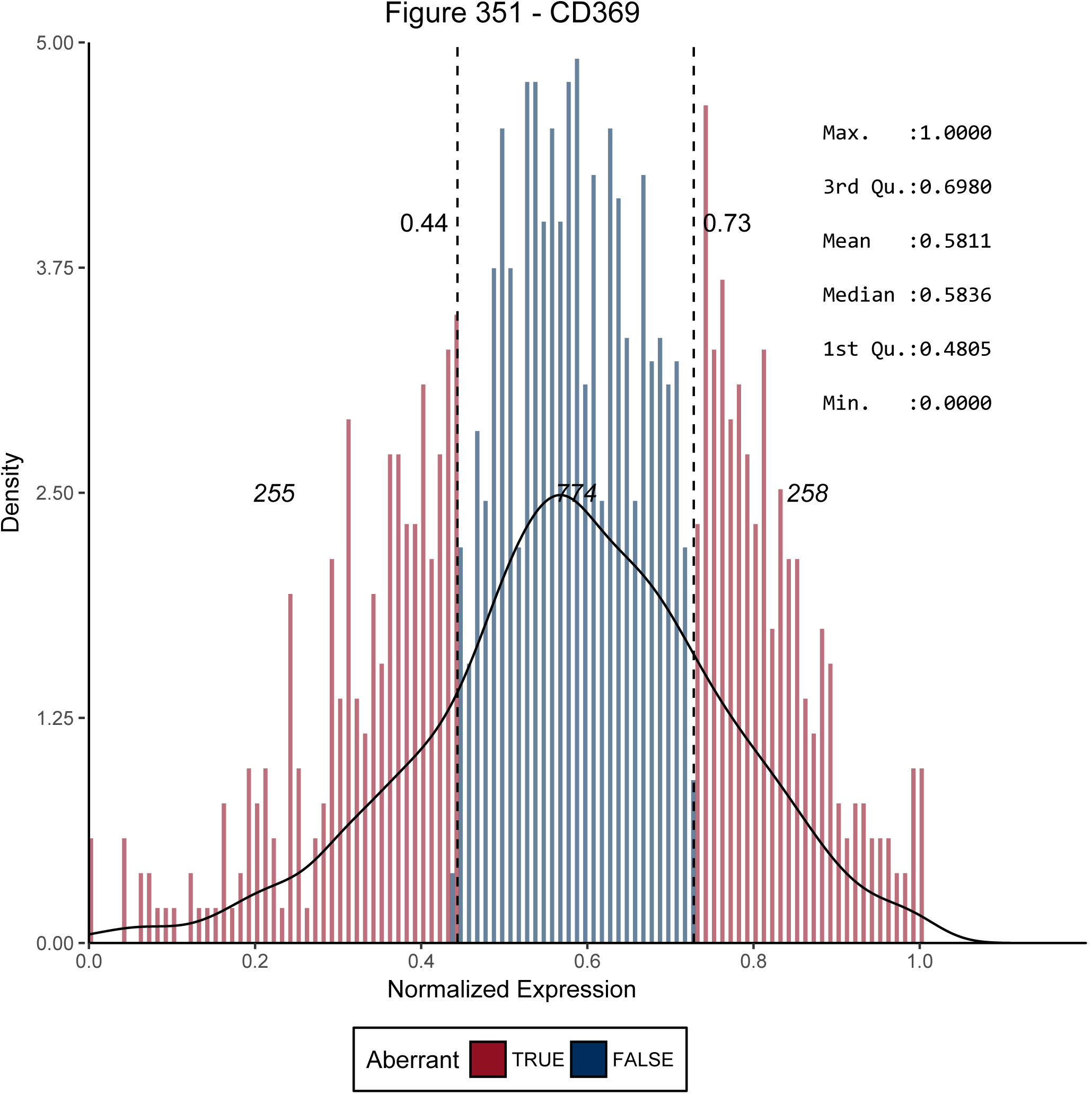
Distributions Clusters of Differentiation and labels of aberrant gene expression.

